# CAD-C: An engineered nuclease enables repair-free *in situ* proximity ligation and nucleosome-resolution chromosome walks in human cells

**DOI:** 10.64898/2025.12.22.695891

**Authors:** Jan Soroczynski, Lauren Anderson Westcott, Wu Zuo, Arnold Ou, Hera Canaj, James Hickling, Joanna L. Yeung, Hide A. Konishi, Ellie B. Campbell, Conor Whelan, Jennifer Balacco, Giulio Formenti, Viviana I. Risca

## Abstract

Chromosome conformation capture (3C)-derived methods have become an indispensable tool in the study of gene regulation. The three-dimensional contacts probed by 3C methods depend strongly on the properties of the enzyme used to fragment chromatin prior to proximity-driven ligation. Micrococcal nuclease (MNase), used in Micro-C, increases resolution at the expense of low ligation efficiency and the need for extensive enzyme titration. To overcome these limitations, we engineered a TEV protease-activatable caspase-activated DNase (CAD) to enable an efficient, low-sequence-bias, and high-resolution proximity ligation assay we call CAD-C. CAD-C was successful on the first attempt for each human cell line tested and the resulting datasets capture loops, TADs, compartments, and stripes similarly to Micro-C. However, compared to Micro-C and Hi-C, CAD-C shows enhanced sensitivity for promoter-enhancer loops. Leveraging the ligation-competent DNA ends produced by CAD cleavage, we show that CAD-C is compatible with a highly streamlined, repair-free protocol and produces multi-step CADwalks, consecutive ligations between nucleosomal or sub-nucleosomal fragments. With these walks, we probe local chromatin fiber folding contacts, nucleosomal and sub-nucleosomal footprints, and long-range nuclear organization regimes in human cell lines. CAD-C is an efficient, robust chromatin structure assay that can span sub-nucleosomal to chromosomal length scales in a single experiment.

## Introduction

Chromosome conformation capture (3C^1^) and its high-throughput derivatives (Hi-C^2,3^ and Micro-C^4–6)^ have become essential parts of the toolkit for mapping genome folding from megabase compartments to kilobase loops and nucleosome-resolved regulatory architectures that are providing mechanistic insights into the regulation of transcription and other DNA-based processes^7–10^. These assays share a common logic: crosslinking chromatin, fragmenting DNA within intact nuclei, and using proximity ligation to convert spatial nucleosome neighborhoods into sequencing-readable DNA junctions. 3C-based readouts measure population-averaged proximity/ligation frequencies rather than distance in three-dimensional space^11–13^. The fragmentation step strongly influences both the resolution of the data and its interpretability^3,11,12,14^. Restriction enzyme–based workflows impose site spacing and often rely on detergent conditions that can perturb native chromatin. In contrast, MNase-based Micro-C improves spatial resolution and in a few selected cell lines, has revealed intriguing mesoscale features of chromatin folding near genes that were previously not captured with Hi-C^3,5,6,15^. However, Micro-C introduces pronounced accessibility bias, a sequence composition bias and a narrow practical working range: modest changes in MNase activity or exposure can drive a transition from under-digestion to over-digestion, eroding informative fragment distributions and compromising reproducibility^16–19^. The ratio of MNase to chromatin used in an experiment can dramatically affect the sequence composition of the final library, sometimes producing considerable artifacts^20,21^. In addition, MNase generates heterogeneous DNA termini that require end processing to restore efficient ligation substrates^22^, adding steps that can further influence library composition and reduce efficiency. Together, these limitations have circumscribed the applications of Micro-C to samples in which input cell lines are homogeneous and abundant for titration and have reduced the interpretability of short-range features of contact maps in which sequence bias cannot be easily normalized^8^. Much of the potential of Micro-C remains unrealized due to these limitations.

In parallel with improvements in pairwise contact mapping, a growing set of approaches has emphasized that chromatin domains and regulatory assemblies are often better described as multiway contact neighborhoods rather than collections of independent pairwise edges. Long-read proximity-ligation methods, such as MC-3C and Pore-C, directly sequence concatemers to recover higher-order contacts within single molecules^23–25^, while combinatorial barcoding strategies, like SPRITE, capture multiplex hubs without relying on proximity ligation^26,27^. Proximity-tagging approaches such as Proximity Copy Paste (PCP) further demonstrate that nucleosome-scale chromosome structure can be recorded through barcoding^28^. Together, these efforts underscore both the biological relevance of higher-order interactions and the remaining technical challenge: recovering multi-way information at nucleosome resolution while retaining strong sensitivity to regulatory chromatin.

We address this technological gap and the limitations of Micro-C with a chromatin conformation capture strategy, CAD-C, that repurposes a caspase-activated DNase (CAD; also known as DFF40, the 40-kDa subunit of the DNA fragmentation factor). CAD is an eukaryotic endonuclease best known for producing internucleosomal DNA fragmentation during apoptosis^29,30^. In non-apoptotic cells, CAD is maintained in an inactive state through association with its chaperone–inhibitor ICAD (also known as DFF45); upon caspase-3 activation, ICAD is cleaved, releasing CAD to dimerize, activating its endonuclease activity to cleave chromatin into oligonucleosomes^31,32^. CAD-C leverages CAD’s ability to generate ligation-ready blunt ends^33–35^ and low sequence sensitivity to generate nucleosome-resolution contact maps that retain higher-order chromatin features while strengthening recovery of contacts associated with active regulatory elements, including promoters and transcription-linked regions. Second, we exploit CAD’s propensity to ligate multiple nucleosome-scale fragments into single concatemers, in conjunction with short and long-read sequencing, to generate CADwalks. This per-molecule framework preserves ordered, multi-fragment co-occurrence rather than collapsing observations into independent pairs. Together, CAD-C and CADwalks provide a unified platform for interrogating genome folding across both scale and interaction order, enabling multi-scale and multiway views of the regulatory landscape and directly linking fine-scale architecture to transcriptionally relevant chromatin organization.

## Results

### Engineered CAD robustly cleaves chromatin between nucleosomes with low bias

We set out to find an improved nuclease for sequence-independent chromatin fragmentation between nucleosomes prior to proximity ligation. CAD has several properties that make it an attractive candidate. Unlike MNase, which is thought to engage with DNA and cleave one strand at a time before engaging in exonucleolytic digestion of the DNA, leaving frayed ends^18,36^, CAD is activated as a scissor-like homodimer whose active sites sit in a deep catalytic cleft; it cuts dsDNA endonucleolytically (without processive end resection), producing blunt or near-blunt breaks with 5′-phosphate/3′-hydroxyl ends^33–35^ (Fig. 1a-c). CAD produces blunt 5’-phosphorylated ends at its cleavage sites which are substrates for T4 DNA ligase^22^ and is not thought to overdigest past the mononucleosome scale^37^. Moreover, CAD can be deployed under chromatin structure-preserving buffer conditions enabling nucleosome-scale mapping without disrupting higher-order chromatin organization^38–40^. CAD has already been successfully employed for nucleosome and polymerase footprinting and for chromatin immunoprecipitation, demonstrating its potential as a probe of chromatin structure^41,42^.

**Figure 1.**
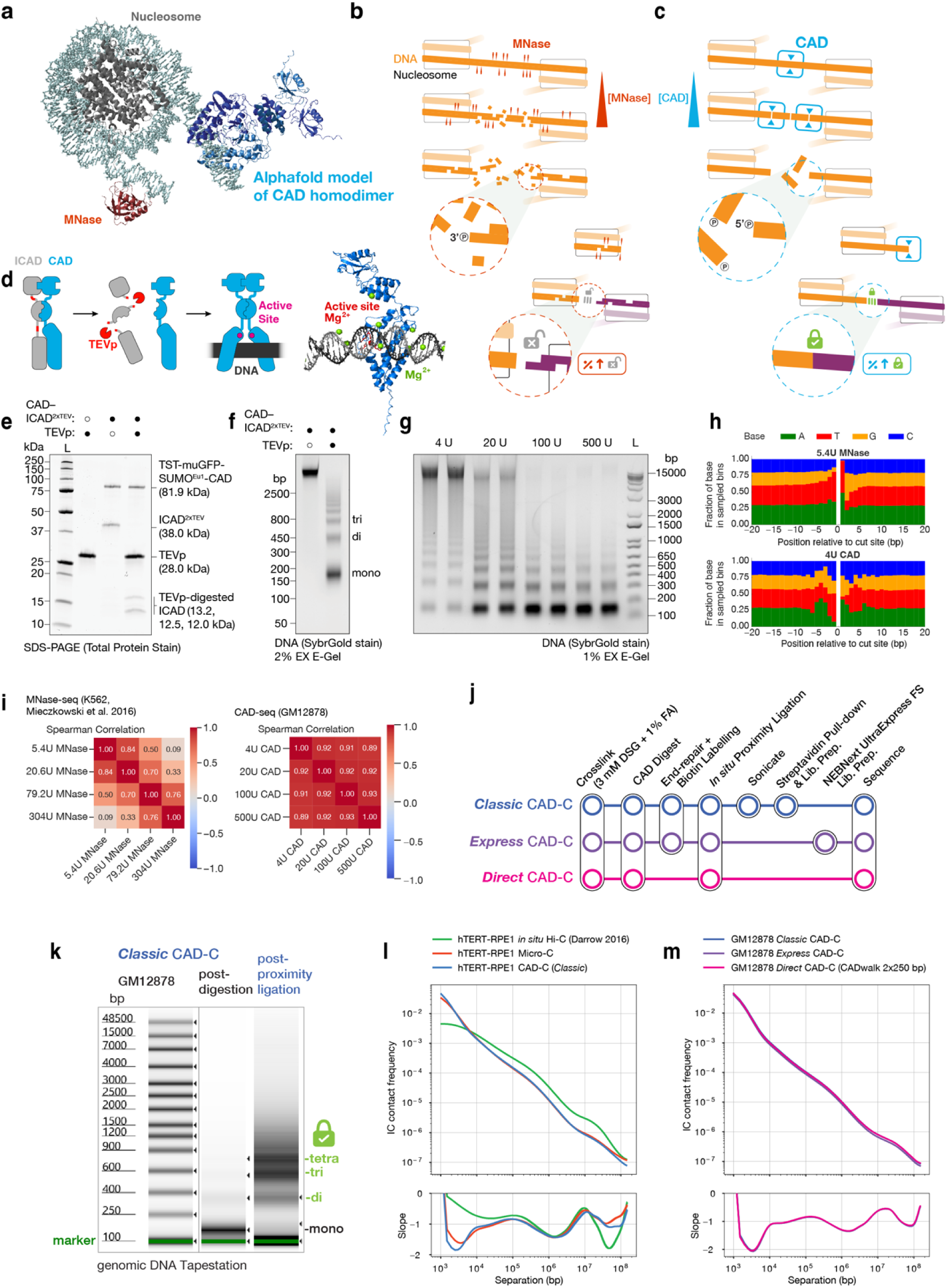
TEVp-gated activation of CAD enables controlled nucleosomal fragmentation with reduced end bias and streamlined repair-free CAD-C workflows. (a) Structural context for chromatin digestion assays: cryoEM mononucleosome (PDB 7PEU, grey), Alphafold model of the CAD homodimer (blue) and MNase (PDB 1STN, red). (b-c) Schematic contrasting MNase, which combines endonuclease, exonuclease and nickase activities that generate frayed 3’-phosphorylated ends at nucleosomal margins^22^, with CAD, which is a strict endonuclease yielding blunt 5’-phosphorylated ends^33–35.^ ^(d)^ Schematic depicting activation of ICAD2xTEV–CAD proenzyme by TEV protease. (e) Validation of TEVp-cleavage of ICAD^2xTEV^:TwinStrepTag-muGFP-SUMO^Eu1^. (f) TEVp-dependent activation of CAD DNase activity on a fixed chromatin substrate. SDS-PAGE gel stained for total protein. (g) 1% EX E-Gel of GM12878 nuclei (4 × 10^6^ per lane) fixed with 3 mM DSG and 1 % FA and digested with 4 U, 20 U, 100 U and 500 U CAD. Mono-, di- and tri-nucleosome bands are indicated. (h) Per-position A/C/G/T frequencies from −20 to +20 bp relative to each fragment start (inferred cut) for a K562 MACC 5.4 U MNase dataset and a GM12878 CAD digestion library prepared with 4 U CAD (CAD-seq). (i) Spearman correlation matrix of genome coverage of digested fragments with indicated concentrations of MNase^47^ and CAD. (j) Overview of Classic, Express and Direct CAD–C workflows. (k) Capillary electrophoretogram (genomic TapeStation) of the DNA fragment length distribution from GM12878 cells post-digestion and post-Classic CAD-C proximity ligation. (l) Intrachromosomal (IC) contact frequency as a function of genomic separation (top) and corresponding local slope (bottom) for hTERT-RPE1 datasets. Curves were computed from 500 bp binned, ICE^48^ balanced cis matrices (cooler^49^/cooltools^50^) and show in situ Hi-C^51^ (green), Micro-C (orange), and Classic CAD-C (blue). (m) P(S) and slope for GM12878 CAD-C datasets, comparing Classic (blue), Express (purple) and Direct CAD-C / CADwalk 2×250 bp libraries (magenta). Illustrations in (b-d) created by Oscar Alejandro Perez.

MNase has dominated as the enzyme of choice for chromatin cleavage in part because it is easily activated by the addition of calcium^18,19^. The physiological activation of CAD requires caspase cleavage of its inhibitor ICAD, with which it must be co-expressed because ICAD is required for proper CAD folding^32,43^ (Fig. 1d). Caspase-based activation is impractical for genomics applications, due to caspase’s many endogenous protein substrates present across cell types^44^. Previous studies have engineered human CAD to be cleavable by TEV protease (TEVp)^41,45,46^. Inspired by these studies, we used a similar strategy to engineer TEVp-cleavable sites into the full-length murine ICAD sequence and designed a dual-cassette, codon-optimized co-expression construct for full-length murine CAD with an N-terminal fusion of TwinStrepTag, muGFP and SUMO^Eu1^ and murine ICAD^2xTEV^, that produces an excess of ICAD^2xTEV^ over TwinStrepTag-muGFP-SUMO^Eu1^-CAD (Fig. 1d, Extended Data Fig. 1a-d). This optimization ensures that all expressed CAD is inhibited and properly folded by its interaction with ICAD. After affinity purification and polishing by size exclusion chromatography, we found that the majority of the TEVp-cleavable ICAD and CAD eluted in the size exclusion void volume, likely due to binding of bacterial DNA (Extended Data Fig. 1e-h). In preliminary results from a subsequently optimized protocol, we have shown that increasing salt concentration eliminates this DNA binding and allows a purer fraction of CAD/ICAD^2xTEV^ to be eluted in a discrete peak with ∼2.5-fold higher TEVp-specific DNA cleavage activity (Supplementary Fig. 1; see Methods).

The void volume CAD/ICAD fractions exhibited the highest TEVp-activation-specific nuclease activity (Extended Data Fig. 1h). We pooled these fractions into a working stock (Fig. 1e, Extended Data Fig. 2a-b) with robust, titratable. and reliable TEVp-dependent DNA nuclease activity (Extended Data Fig. 2c-d). Digestion of chromatin with this CAD stock produced the expected oligonucleosomal DNA ladder (Fig. 1f, Extended Data Fig. 2d-e).

To assess the robustness of chromatin digestion by our engineered CAD, we digested disuccinimidyl glutarate (DSG) and formaldehyde (FA) fixed GM12878 nuclei, titrating CAD/ICAD in the presence of excess TEVp (Fig. 1g). Across a 125-fold range of CAD concentrations, the digested DNA resolved into mono-, di- and tri-nucleosome bands, with no detectable accumulation of sublZnucleosomal fragments, consistent with controlled fragmentation of crosslinked chromatin rather than extensive over-digestion. No digestion was observed in the absence of TEVp (Fig. 1f). Together, these results demonstrate that the engineered CAD enzyme digests fixed chromatin efficiently, and remains strictly dependent on TEVp for activation.

We next asked whether CAD shares the local nucleotide bias characteristic of MNase^18,19^, or displays a distinct sequence preference at its cleavage sites. We applied an identical fragment-end analysis to a 5.4 U K562 MNase accessibility (MACC) dataset^47^ and to a 4 U GM12878 CAD digestion library (Fig. 1h). Compared to MNase (Fig. 1h, top) CAD digestion dataset displays a more even distribution of A, C, G and T around the cut sites (Fig. 1h, bottom), with only modest variation in A+T versus G+C composition, but exhibits a purine–pyrimidine asymmetry across the cleavage sites (Extended Data Fig. 3a), consistent with previous reports^33,52^. The lower base composition bias of CAD cleavage is evident in comparing the base composition quantile-normalized coverage plots for cleaved fragment libraries (Fig. 1i, Extended Data Fig. 3b-d). Although CADlZcleaved libraries show some celllZtype–specific differences, potentially reflecting biological differences, their profiles are largely insensitive to enzyme concentration, in contrast to MNase. Coverage correlations across genomic bins remain high across CAD titrations, whereas MNase titration shows substantially lower concordance^47^ (Fig. 1i, Extended Data Fig. 3e-f). Similar trends were observed when comparing fragmentlZend profiles in hTERT-RPE1 cells (Extended Data Fig. 4a-c).

Lastly, to assess CAD-mediated chromatin cleavage at higher resolution, we analyzed the coverage profiles and length distributions of CAD-generated DNA fragments. We first verified that the nucleosome-sized fragments reproduce expected patterns^53,54^ around CTCF-bound sites (Extended Data Fig. 3g,j), hyperaccessible sites (Extended Data Fig. 3h), and transcription start sites (TSSs) (Extended Data Fig. 3i). We observed that sub-nucleosomal-sized fragments, thought to derive from footprints of other DNA-bound proteins such as CTCF, polymerase complexes, transcription factors and linker histones, were retained even at high enzyme concentrations–in contrast to the loss of these sub-nucleosomal fragments in MNase digestion (Extended Data Fig. 3j). This difference in the robustness of the digestion profile to changes in enzyme concentration is also observable in the fragment size distributions from MNase digests (Extended Data Fig. 3k) as compared to CAD digests (Extended Data Fig. 3l-o). Sub-nucleosomal fragments from CAD digests recapitulate ATAC-seq peaks (Extended Data Fig. 3p).

### CAD-C: CAD digestion enables a streamlined proximity ligation method without sacrificing performance

The robustness of CAD digestion prompted us to incorporate CAD into a proximitylZligation workflow, which we term CAD-C. We began with a protocol adapted from *in situ* Micro-C as optimized for Region Capture Micro-C (RCMC)^15^, modifying the chromatin digestion step to permit RNase treatment (because CAD is inhibited by RNA^55^), followed by incubation with CAD/ICAD heterodimer, and activation of CAD with TEVp cleavage of ICAD (Classic CAD-C; Fig. 1j-k). To test whether ligation-junction enrichment is required once CAD-C has already generated junction-dense concatemer DNA, we also prepared an Express CAD-C variant protocol in which purified post-ligation CAD-C DNA is converted to Illumina libraries by enzymatic fragmentation and one-tube library preparation without streptavidin capture (Fig. 1j; Methods). Finally, we implemented Direct CAD-C, which, given CAD’s ligation-competent cleavage ends, skips end repair and biotin fill-in (and therefore streptavidin-based enrichment) allowing sequencing of intact concatemers that give rise to CADwalks (Fig. 1j), discussed in more detaile below.

We applied Classic CAD-C to three cell lines, hTERT-RPE-1, which were arrested in G0/G1 by contact inhibition and GM12878 and K562, which were asynchronously cycling (Supplementary Fig. 2). We also applied additional variations of the protocol to GM12878. In all CAD-C experiments, we used a saturating concentration of CAD, i.e. a concentration beyond which no further chromatin digestion occurs. Excitingly, the protocol produced high-quality libraries from all three cell lines on the first attempt.

CAD-C libraries reproduced the low nucleotide composition biases and high-fidelity capture of nucleosomal fragments evident in CAD-C digests, exhibiting nucleosome footprints in both genome-wide tracks and pileups around CTCF binding sites or transcription start sites (TSSs) (Extended Data Fig. 4d-h).

To compare the datasets from these protocols against each other and against the field-standard Hi-C and Micro-C^15,56^, we computed intrachromosomal contact–probability curves L(L) from 500 bp, ICE-balanced cis matrices to obtain distance-stratified cis expectations on chromosome arms. In hTERT-RPE1, the Classic CAD-C L(L) closely tracked the Micro-C curve over separations from ∼10^3^ to ∼10^7^ bp, with both non-restriction enzyme-based assays showing a steep decay that extended into the short-range regime (Fig. 1l). By contrast, the *in situ* Hi-C dataset displayed a flatter decay at the shortest distances, consistent with limits imposed by the spacing of MboI restriction sites. Interestingly, CAD-C showed a slightly stronger shoulder in the *P*(*s*) derivative (Fig. 1l bottom) at ∼10kb, which may indicate an increased sensitivity to short loops and interactions between chromatin elements on this length scale, which we explore in more detail below. In comparing the three versions of CAD-C (Fig. 1j,m), we found that they performed nearly identically, showing that there is no loss in data quality in the Direct CAD-C protocol. Therefore, for analyses requiring the deepest data, such as loop calling or short-range pileups, we merged data from different protocols.

Quality control metrics of contact distance distributions (Supplementary Fig. 2) indicated that CAD-C produces more contacts within ∼1 kb than Micro-C or Hi-C. To compare the performance of CAD-C against Hi-C and Micro-C in more detail for chromatin compartment and topologically associated domain (TAD) analysis, we normalized the datasets based on the coverage of contacts beyond the length of 1kb as these long-range features are based on contacts away from the contact matrix diagonal.

The self-organization of chromatin into active (A) and inactive (B) compartments is evident in a 100-kb resolution contact map (Fig. 2a), in which we observe comparable compartmentalization patterns upon comparison between CAD-C with Hi-C and Micro-C in the same cell line. To assess how well CAD-C captures this segregation compared to standard methods, we performed saddle-plot analysis on ICE-balanced contact matrices for both hTERT–RPE1 (Fig. 2b) and GM12878 (Extended Data Fig. 5a). In hTERT–RPE1, the Classic CAD-C dataset yields a compartment strength score of 5.33, compared to 1.89 for in situ Hi-C and 2.84 for Micro-C (Fig. 2b). The corresponding saddle heatmaps show that the CAD-C matrix displays deeper depletion in the off-diagonal (A–B) blocks and stronger enrichment in the homotypic (A–A and B–B) corners than either benchmark, indicating that CAD-C resolves the compartmentalization of the RPE1 genome with higher contrast. This may be due to some enrichment of CAD cleavage at active chromatin that is not fully compensated by matrix balancing, or more likely, higher efficiency of capture of contacts between active regions discussed below. Inspection of CAD-C coverage over regions containing accessible sites indicates that this bias toward accessible chromatin is not causing strong over-representation of these regions compared to Micro-C (Extended Data Figure 4d).

**Figure 2.**
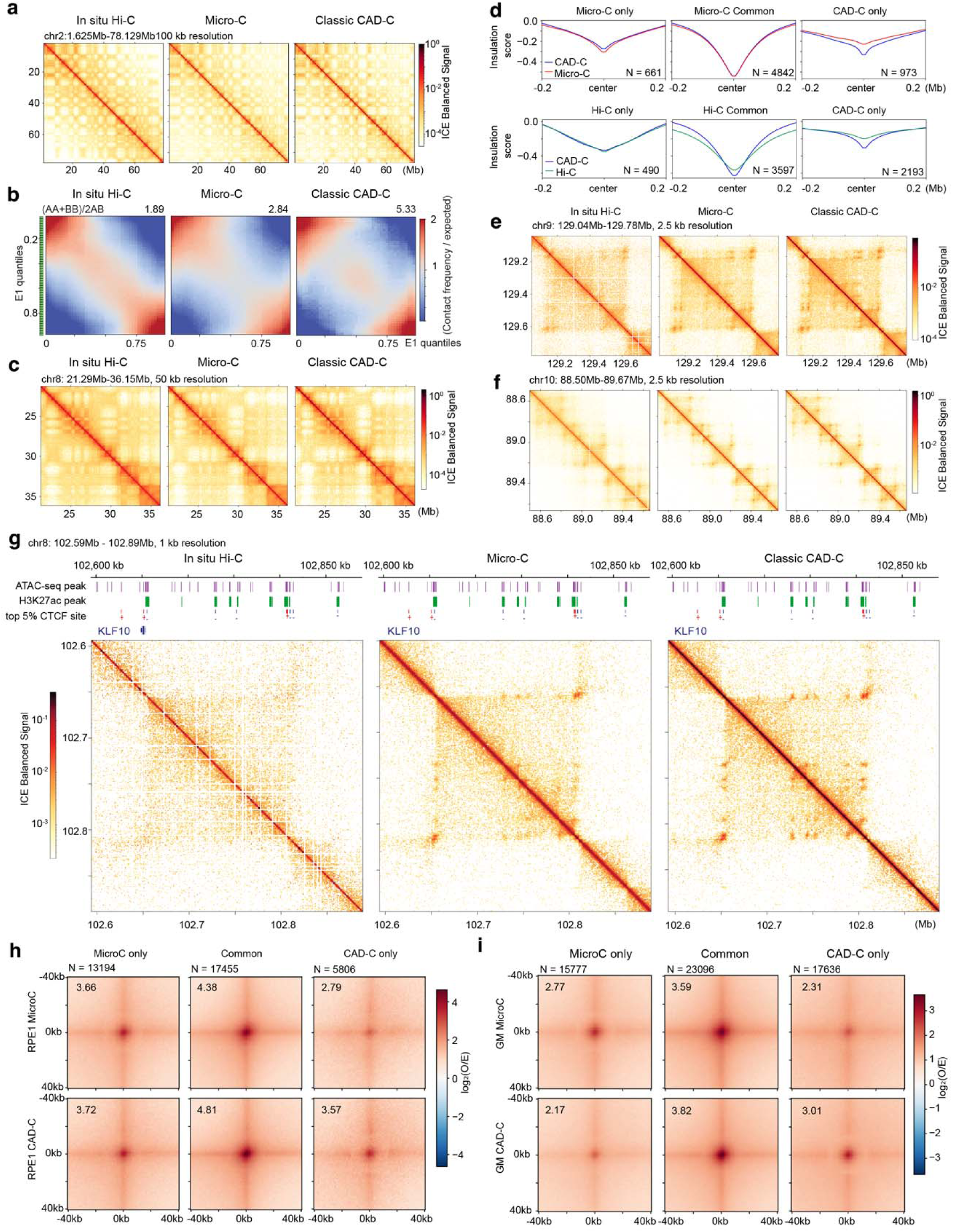
Classic CAD-C matches canonical chromatin features from Hi-C/Micro-C, shows stronger A/B compartmentalization, and identifies an overlapping but non-identical set of boundaries and loops. (a) Contact maps for in situ Hi-C^51^, Micro-C and *Classic* CAD-C in hTERT-RPE1 cells at the resolution of 100 kb for chr2:1,625,634–78,128,894. Datasets were subsampled to match their coverage depth for contacts > 1kb. (b) Eigenvector-based saddle plots for in situ Hi-C^51^, Micro-C and Classic CAD-C in hTERT–RPE1. The compartment-strength score (AA + BB)/2AB is reported above each panel (hTERT– RPE1 in situ Hi-C: 1.89; hTERT–RPE1 Micro-C: 2.84; hTERT–RPE1 Classic CAD-C: 5.33). The datasets were subsampled as in (a). (c) hTERT-RPE1 heatmap 50 kb resolution maps for chr8:21,293,200–36,154,099, subsampled as in (a). (d) CAD-C insulation pileups at hTERT-RPE1 TAD boundaries compared with Micro-C and Hi-C. Upper panel: Comparison of CAD-C (blue) and clinical Micro-C (red) insulation profiles at boundaries called only in Micro-C, common between CAD-C and Micro-C, or only in CAD-C. Lower panel: Comparison between CAD-C (blue) and in situ Hi-C (green) at Hi-C-only, common, and CAD-C only boundaries. datasets for the comparison were subsampled as in (c). (e-f) 2.5 kb resolution maps for hg38 chr9:129,042,198–129,780,074, and hg38 chr10: 88,496,048-89,665,189 bp. Left: public *in situ* Hi-C dataset. Middle: *in situ* Micro-C dataset. Right: Classic CAD-C. Data was subsampled as in (a). (g) 1 kb resolution maps for hg38 chr8:102,594,246–102,889,522 centred on the *KLF10* locus comparing Left: public *in situ* Hi-C dataset^51^, Middle *in situ* Micro-C, and right CAD-C plot. IGV tracks shown above the heatmaps indicating KLF locus, ATAC-seq narrow peaks (purple), H3K27ac ChIP-seq (GSM5899629) narrow peaks (green) and occupied CTCF motifs +(positive strand in red) – (negative strand in blue). Data was subsampled as in (a). (h) Loop pileup over +/-40kb for *in situ* Micro-C only, common, and CAD-C only loops, from hTERT-RPE1 cells at 1kb resolution. Data was subsampled to match total contact depth (regardless of contact distance). (i) Loop pileup over +/-40kb for *in situ* Micro-C only, common, and CAD-C only loops, from GM12878 cells at 1kb resolution. GM12878 Micro-C data was obtained from Hong *et al.* ^57^.

To assess CAD-C’s ability to identify topologically associating domains (TADs), evident in 50-kb resolution contact matrices (Fig. 2c) we used cooltools^50^ to call TAD boundaries in hTERT RPE1 cells at 10 kb resolution from matched in situ Hi-C, Micro-C and CAD-C datasets (Fig. 2d). At comparable contact matrix total count depth, >80% of CAD-C boundaries overlapped those identified by Micro-C, and CAD-C detected substantially more boundaries than in situ Hi-C (Fig. 2d). To further evaluate boundary concordance across methods, we compared insulation scores genome-wide and computed pairwise correlations. The correlation heatmap revealed the strongest agreement between Micro-C and CAD-C, indicating highly consistent insulation landscapes (Extended Data Fig. 5b). A similar boundary analysis in GM12878 cells also showed a high degree of consistency between Micro-C and CAD-C (Extended Data Fig. 5c). We additionally performed local pileup analyses centered on called boundaries. All three boundary sets exhibited canonical boundary signatures, with a reproducible dip in insulation at boundary centers and higher insulation values on either side (Fig. 2d).

At higher resolutions, the Hi-C map becomes noticeably more diffuse, whereas the Micro-C and Classic CAD-C maps preserve more punctate loop-like focal enrichments along the diagonal (Fig. 2e-f). This difference is even more evident in a 1 kb view centered on the KLF10 locus on chromosome 8 (hg38 chr8:102,594,246–102,889,522), where Classic CAD-C reveals sharp, discrete contact peaks that, as expected, are weak or not clearly resolved in the matched-depth Hi-C map (Fig. 2g). Several of these CAD-C focal contacts coincide with H3K27ac and ATAC-seq narrow peaks in the overlaid IGV tracks (Fig. 2g), consistent with contacts involving putative enhancer elements in this region.

Given the functional importance of looping contacts between regulatory elements and genes or between regulatory elements themselves, like enhancers and promoters^9^, we sought to further investigate the performance of CAD-C in detecting loops in comparison to Micro-C. We called loops using mustache^58^ on total contact depth-matched datasets–for all contact distances–for *in situ* Micro-C and CAD-C in hTERT-RPE1 cells at ∼1B pairs (Fig. 2h) and in GM12878 cells at ∼4B pairs (Fig. 2i)^57^. We observed an overlap in the majority of loop calls by mustache and cooltools at 2.5 kb resolution, but in both cases, there were several thousand loops called by only one method (Fig. 2h-i). To compare loop-detection sensitivity, we generated subsampled datasets spanning a range of unique valid-pair counts and called loops at 2.5-kb resolution. Loop yield increased monotonically with depth for all assays and CAD-C performed similarly to the state-of-the-art *in situ* Micro-C protocol in GM12878 cells (Extended Data Fig. 5d,e). Taken together, these multi-scale comparisons indicate that, at matched depth, CAD-C produces hTERT-RPE1 contact maps that recapitulate large-scale compartment and domain organization seen by Hi-C and Micro-C while providing high sensitivity for loop detection, comparable to *in situ* Micro-C.

### CAD-C preferentially captures loops anchored at active transcription regulatory regions

We chose to further benchmark loop analysis on GM12878 libraries for several reasons. First, GM12878 cells are well-annotated in terms of both their epigenome and some functional gene regulation data, which facilitates validation of loops in terms of their functional relevance. Second, two types of Micro-C public datasets are available for comparison, providing a useful contrast: one, produced using the Dovetail commercial kit (Cantata Bio), and another ultra-deep *in situ* Micro-C dataset associated with a recent preprint^60^.

In analyzing contact matrices from our deepest merged CAD-C dataset (∼4 billion contacts in GM12878 cells) at resolutions from 2.5 kb to 0.5 kb (Fig. 3a-c), we observed that there is strong signal on the diagonal, as would be expected based on a high contact probability of nearby loci captured by the high ligation efficiency of CAD-C. Despite this concentration of signal on the diagonal (contact distance < 5-10 kb), we also observe high-contrast loops off-diagonal, coinciding with a subset of accessible DNA sites. For example, the loop in Fig. 3c spans two active promoters.

**Figure 3.**
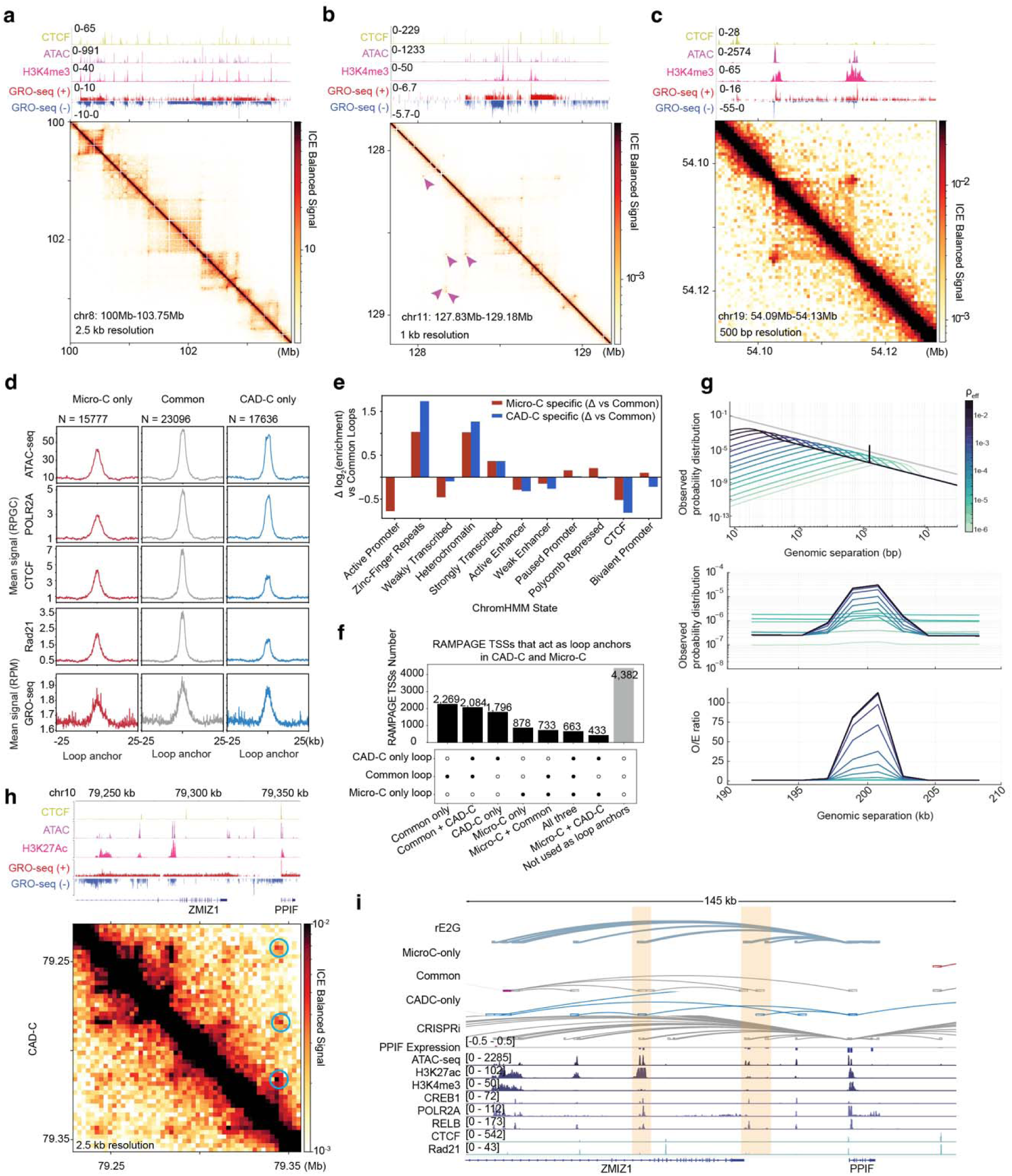
CAD-C sensitively detects loops between active regulatory regions. (a-c) ICE-balanced intrachromosomal contact maps from the GM12878 merged CAD-C dataset with a shared logarithmic color scale for ICE-balanced signal. (a) 2.5 kb resolution map for hg38 chr8:100,000,000–103,750,000, showing similar multi-megabase organization with many focal contacts aligning with CTCF/ATAC peaks. IGV tracks display CTCF ChIP–seq, ATAC–seq, H3K4me3 ChIP–seq and GRO–seq signal (plus and minus strands). (b) 1 kb resolution map for hg38 chr11:127,832,229– 129,177,988. Magenta arrowheads mark selected punctate contacts that are supported by Mustache loop calls and coincide with local CTCF and ATAC peaks. and IGV tracks display CTCF ChIP–seq, ATAC–seq, H3K4me3 ChIP–seq and GRO–seq signal (plus and minus strands). (c) 500 bp resolution map for hg38 chr19:54,093,528–54,128,248, highlighting sharp near-diagonal focal contacts at positions with clustered ATAC and H3K4me3 peaks. IGV tracks display CTCF ChIP–seq, ATAC–seq, H3K4me3 ChIP–seq and GRO–seq signal (plus and minus strands). (d) Chromatin accessibility, H3K27ac, POLR2A, CTCF, cohesin subunit Rad21, and nascent transcription (GRO-seq) pileup line plots at Micro-C only, common, CAD-C only loop anchors called at the resolution of 2.5 kb with equal depth of CAD-C and Hong *et al.* GM12878 *in situ* Micro-C data, as in Fig. 2i^57^. (e) ChromHMM state enrichments at loop anchors for the 4 billion-contact matched GM12878 CAD-C merged data and GM12878 *in situ* Micro-C^57^ loop sets (as in Fig. 2i), corresponding differences (Δ log_2_) in enrichment for CAD-C–specific and Micro-C–specific anchors relative to the Common set. (f) UpSet plot summarising these TSS-level intersections (union-only view). Each bar reports the number of RAMPAGE TSSs that are used as loop anchors in the indicated combination of loop sets (black dots in the lower panel). TSSs counts for uniquely anchored in CAD-C–specific loops (CAD-C only) or Micro-C–only loops, or Common loop sets that were called with Mustache on 4 billion-contact, ICE-balanced GM12878 CAD-C merged data and GM12878 *in situ* Micro-C^57^, as in Fig. 2i. (g) Simulated observed contact probability distributions Pobs(S) (top and middle) and corresponding observed/expected (O/E) ratios (bottom) for the same set of ρ_eff_ values. (See Supplementary Note for details.) (h) CAD-C loops at the Nasser PPIF IBD risk locus with IGV-style composite view at the PPIF locus followed by ATAC, H3K27ac, H3K4me3, transcription factor and cohesin tracks (CREB1, POLR2A, RELB, CTCF, Rad21), RNA-seq, GRO-seq and gene annotation. (i) The PPIF locus centred on the intronic IBD risk variant rs1250566, adapted from Nasser et al., 2021^62^. Top loop track: rE2G predicted loops^63^ from the activity-by-contact (ABC) model^62^. Loop tracks 2-4 from top: GM12878 Micro-C-only, Common and CAD-C-only loops called at 2.5 kb resolution. Bottom loop track: CRISPRi sgRNA positions and differential expression (CRISPRi DE) from Nasser et al., 2021^62^. Below: PPIF expression change upon CRISPRi treatment at each putative enhancer locus and epigenome feature tracks. Orange bars highlight CAD-C-only loops that coincide with functionally validated PPIF enhancers.

Chromatin loops can be formed by multiple mechanisms, including cohesin pausing at inwardly-oriented CTCF-bound sites and cohesin-independent interactions between active regulatory regions^61^. To compare the sensitivity of CAD-C to these two loop classes, we explored the propensity of CAD-C-specific loops to occur between regions of hyper-accessible chromatin (a proxy for regulatory regions) *versus* CTCF-bound sites (Extended Data Fig. 6a), and then analyzed the epigenetic state at loop anchors. CAD-C-specific loops exhibit a stronger association with chromatin accessibility (ATAC-seq, ∼39% stronger) and active transcription (RNA polymerase, ∼50% stronger and GRO-seq signal, ∼5% stronger), and a weaker association with loop extrusion boundaries (CTCF, ∼14% weaker and RAD21, ∼14% weaker) (Fig. 3d) as compared to Micro-C-specific loops. We used Hi-ChIP, an orthogonal method that uses antibody pull-down to examine contacts between targeted regions, to cross-validate that CAD-C-specific loops have contacts comparable to those found by Micro-C at both cohesin-associated loops and H3K27ac-associated loops (Extended Data Fig. 6b).

We wondered whether the enhanced sensitivity of CAD-C for loops between active regulatory regions may be a result of a failure of balancing in which all open chromatin regions within a certain distance may appear to interact. However, examination of neighborhoods of ATAC-seq peaks and their corresponding loops showed that it is not the case—not all ATAC-seq peaks interact (representative example shown in Extended Data Figure 6c).

We found that the promoter or enhancer preference of CAD-C *vs.* Micro-C depends on the type of Micro-C protocol used. It is less than two-fold stronger than *in situ* Micro-C when depth-matching is performed across all contacts (Fig. 3d). We also compared loop calls at 5 kb resolution between CAD-C and Micro-C when depth matching was performed based on contacts > 1 kb, because CAD-C data has a higher fraction of contacts < 1 kb than Micro-C (Supplementary Fig. 2). In this type of analysis, CAD-C-specific loops as compared to those called on the kit, commercial Micro-C data (Dovetail/Cantata Bio) are more focal and more enriched in active chromatin marks (Extended Data Fig. 7a-d). For example, the enrichment of ATAC-seq signal and RNA polymerase ChIP-seq signal was more than two-fold at CAD-C specific loop anchors as compared to commercial Micro-C specific loop anchors (Extended Data Fig. 7). A similar comparison against our own *in situ* Micro-C in hTERT-RPE1 cells shows a less dramatic but still present contrast (Supplementary Fig. 3a-c).

Consistent with these results, we observed that active promoters are similarly enriched at CAD-C-specific and common loop anchors relative to all anchors (log2-fold enrichment=1.73), but are less enriched at Micro-C-specific loop anchors (log2-fold enrichment=0.96, difference in log_2_-fold enrichment = −0.77) (Fig. 3e). Furthermore, most of the active transcription start sites (active promoters) that overlap with loop anchors do so in CAD-C-specific loops (1,796 TSSs) or common loops detected by both CAD-C and *in situ* Micro-C (2,269 TSSs), as compared to 878 TSSs found at Micro-C-specific loop anchors (Fig. 3f).

We wondered why CAD-C may be more sensitive to loops bridging two anchors with DNA accessibility and active transcription. Part of the answer may be a preference of CAD cleavage for accessible chromatin (Extended Data Fig. 3h, Extended Data Fig. 4d,g). However, we did not find a difference between the ligation junction coverage produced by CAD-C and Micro-C over each set of loop anchors, indicating that this is not likely to be the responsible mechanism (Extended Data Fig. 8). We therefore turned to the contribution of the high ligation-competence of CAD-cleaved fragments. Using a simple model of loop capture by proximity ligation that depends on the effective density of ligatable sites (ρ_eff_), we show that increasing the density of ligatable sites is sufficient to both create a stronger signal near the diagonal of the contact matrix, and to strengthen the contrast of observed loops (Fig. 3g and Supplementary Note). This result provides us with a basis for interpreting both the enrichment of sub-kilobase contacts and the higher contrast of loop calls at low-frequency long-range interactions such as promoter- and enhancer-associated loops as consequences of the higher density of ligatable DNA fragment ends produced by the CAD-C assay.

Finally, we asked to what extent the CAD-C-specific loops we detect may provide functional insights into promoter-enhancer communication. Although the correlations with active epigenetic states, DNA accessibility, and transcription machinery were promising, we sought to determine whether CAD-C can perform better in identifying functionally validated enhancer-promoter loops. In GM12878 cells, the PPIF locus has been profiled using CRISPRi at a large number of predicted enhancer sites^62^. We observed that at this locus, some loops are well-resolved on CAD-C contact matrices at 2.5kb resolution (Fig. 3h). When comparing against the state-of-the-art *in situ* Micro-C dataset^57^, the CAD-specific loops specifically connected the PPIF promoter to the enhancers shown to have a functional effect on gene expression when perturbed (Fig. 3i, orange bars).

Taken together, our results demonstrate that CAD-C not only faithfully recapitulates structural chromatin features such as TADs and compartments with a simplified and robust protocol, but also offers a contrast and sensitivity advantage in detecting transcriptionally active chromatin loops.

### CAD-C enables multi-step nucleosome-resolution walks through local chromatin contacts

We sought to explore if it was possible to bridge multiple length scales within the same nucleus by combining both short-range contacts among nucleosomal DNA fragments and long-range contacts that can report on features like loops, insulation domains, and compartments. Standard 3C assays reduce concatemers to pairs through size selection or computational pair-wise analysis, making this goal impossible. CAD’s ability to produce 5’ phosphorylated/3’ hydroxyl ends allows fragments to be ligated without end-repair, enabling us to study chromosome walks at the resolution of individual nucleosomes. We used Direct CAD-C to produce long concatemers of CAD digest fragments, called CADwalks (Fig. 1j, Fig. 4a).

**Figure 4.**
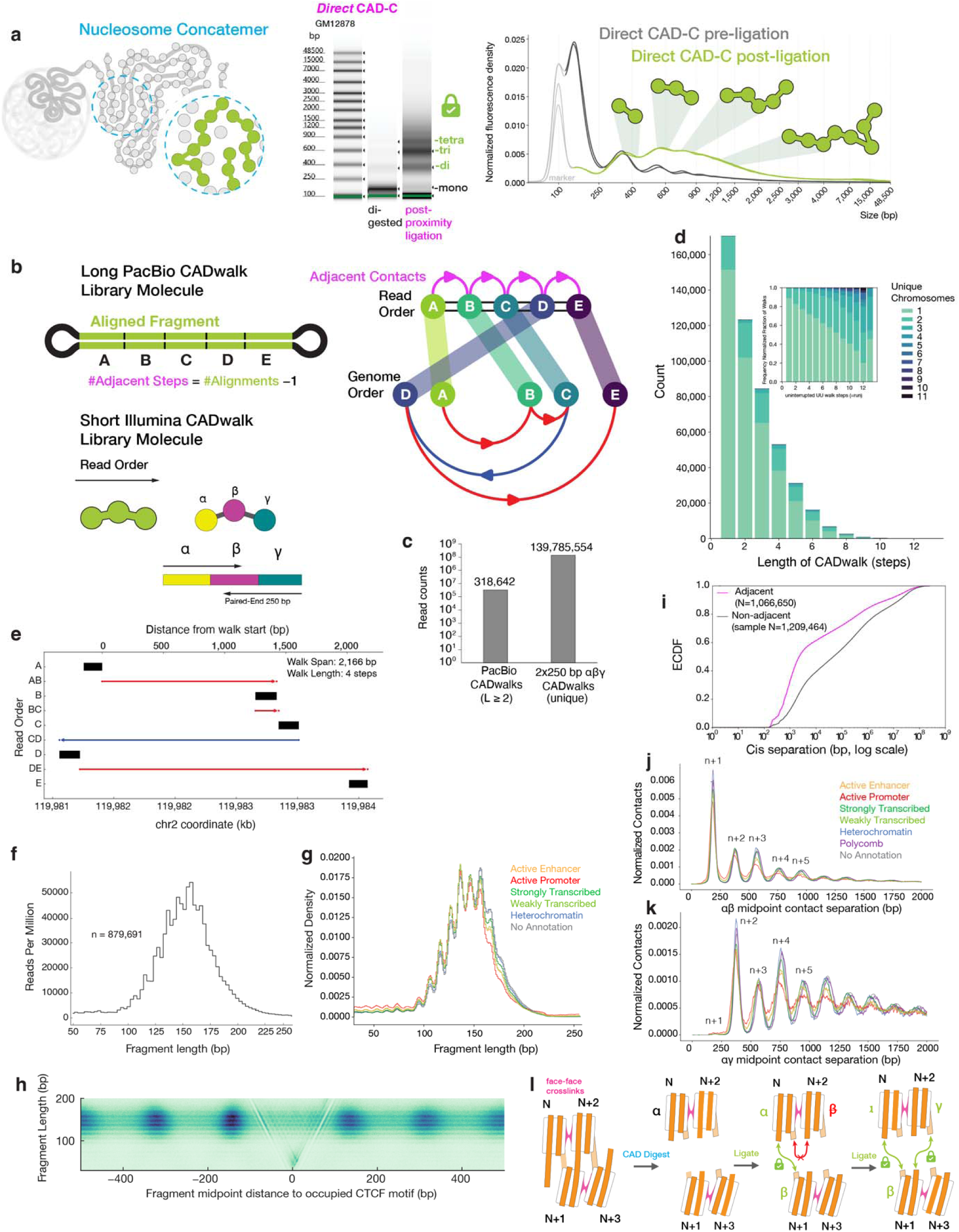
CADwalk concatemers integrate nucleosome footprints with chromatin structure. (a) Diagram of *in situ* nucleosome concatemer, TapeStation electrophoretogram and trace before & after ligation with Direct CAD-C. (b) Left: diagram of CADwalk concatemer reconstruction post-sequencing and alignment for long-read CADwalks (PacBio sequencing) and short CADwalks (250PE Illumina sequencing). Right: Diagram of a long-read CADwalk showing how read order of ABCDE nucleosomes relates to genome order. Genomic coordinates of the molecule illustrated are plotted in (e). (c) Number of observations of CADwalks with >2 steps across long-read and short-read sequencers. (d) Number of unique chromosomes explored per run with an inset of normalized fractions. Histogram is capped at 13 steps. (e) Representative cis-chromosomal long-read CADwalk with five fragments (a-e), four steps. Schematic of ordering for this representative walk is shown directly above in (b, right). Black rectangles mark fragment positions along the chromosome (bottom x-axis, hg38 genomic coordinate in bp). Horizontal segments denote red (positive) and blue negative step along the genomic coordinate. (f) Histogram of all cis-chromosomal long-read CADwalks, bin length = 3bp, normalized to reads per million. (g) Fragment length distribution frequency plot of cis-chromosomal two-step CADwalks fragment length distributions stratified by ChromHMM state, normalized to reads per million. ChromHMM states depicted: Active Enhancer (orange), Active Promoters (red), Strongly transcribed (green), weakly transcribed (light green), no annotation (gray). (h) V-plot of CADwalk fragment midpoint coverage around occupied CTCF motifs (oriented relative to the hg38 plus strand) in GM12878 cells. (i) The empirical cumulative distribution functions (ECDFs) for adjacent and non-adjacent ligations. Distances are computed from fragment midpoints. (j) Plotting from two-step α-β-γ walks, normalized contact frequency plots of direct αβ contacts calculated using midpoints of fragments. Walks were subset by ChromHMM state annotation of fragment α of the given walk. Only uni-directional walks of α-β-γ geometry are plotted, ensuring displacement αγ = αβ + βγ. ChromHMM states depicted: Active Enhancer (orange), Active Promoters (red), Strongly transcribed (green), weakly transcribed (light green), Polycomb regions (purple), no annotation (gray). (k) As in (j), but plotted for indirect αγ contacts. (l) Schematic of proximity ligation at stacked nucleosome fiber showing model of zig-zag local chromatin fiber structure directing the ligation trajectories of CADwalks, providing a model for the stronger observed N+2 periodicity in indirect αγ contacts as observed in (k-j). Illustrations in (a-b) created by Oscar Alejandro Perez.

We reconstructed the CADwalks by two sequencing approaches, first by sequencing long (length >1 kb) concatemers by PacBio long-read sequencing, termed “long-read CADwalks” (Fig. 4b, upper), and secondly by sequencing shorter (length 380 bp-580 bp) concatemers with paired-end 250 bp Illumina sequencing, termed ‘short-read’ CADwalks (Fig. 4b, lower) (see Methods). We define a CADwalk as the longest uninterrupted series of high-confidence ligation steps on a single concatemer read, retaining only steps where the two unique alignments are adjacent (“UU” as defined by the pairtools suite, with ≤30 bp intervening unaligned sequence). As such, each step reflects a true ligation junction adjacency within the same molecule. We refer to long-read cis CADwalk constituent fragments as A,B,C,D,etc. PacBio reads usually cover the entirety of the library molecule, obviating the need to pre-filter long-read CADwalks or enforce any expected walk length. For short CADwalks, we filtered for directly detected ligation junctions and restricted analysis to three-fragment, two-step walks, which we call α-β-γ walks (see Methods). From longlZread CADwalks, 1,215,079 unique steps were obtained (87.79% cis; 12.21% trans, where a trans step is an adjacent fragment pair mapping to different chromosomes). In total, 489,053 longlZread walks were reconstructed, including 318,642 walks with ≥2 steps (i.e., ≥3 fragments) and 139,785,554 unique short αβγ CADwalks (Fig. 4c). The maximum observed walk length was 15 aligned steps (Fig. 4d, Extended Data Fig. 9a).

CADwalks sequenced with both long-read and short-read approaches are predominantly composed of mono-nucleosome sized fragments, but also contain sub-nucleosome fragments and oligo-nucleosome sized CAD digest fragments, which map to the chromosome with steps going in both positive and negative coordinate directions (Fig. 4e-g, Supplementary Fig. 12b). Active promoter and enhancer states show a somewhat higher contribution of short fragments compared to repressed or heterochromatic states, consistent with CAD preserving transcription-factor and non-canonical nucleoprotein footprints in addition to nucleosomes (Fig. 4g). Strikingly, CADwalk concatemers faithfully report on nucleosome-scale and sub-nucleosomal CTCF footprinting, demonstrating that walk reconstruction by proximity ligation and sequencing does not noticeably distort fine-scale chromatin structure information (Fig. 4h). Thus, an advantage of the CADwalk approach is the ability to capture walk trajectories–which are necessarily within the same nucleus, as ligation happens *in situ*–and nucleosome structure with the same measurement.

To compare the relationships between directly ligated (adjacent) steps and indirectly connected (non-adjacent) steps, we calculated the empirical cumulative distribution functions (ECDFs) of successive cis-chromosomal separations for adjacent versus non-adjacent steps within each long-read walk (Fig. 4i). The adjacent-step curve is left-shifted relative to the non-adjacent curve across several orders of magnitude in separation as reported previously^25^, indicating that directly ligated segments are heavily enriched in fragments below 1 kb, consistent with relatively rare long-range ligation events connecting local nucleosome neighborhoods.

Given that CADwalk concatemers capture short fragments and short contact separations with high efficiency, we sought to understand how the *cis* contact probabilities of adjacent and non-adjacent steps shift across epigenetic states. We filtered the deeply sequenced (118M high-confidence walks) two-step α-β-γ walks for walks with unidirectional steps along the chromosome coordinate and calculated their short-range contact probability distributions as a function of their overlap with imputed 12-state chromHMM^68^ epigenetic states (Fig. 4j-k). We observe that direct contacts (α to β) have a decreasing contact probability among successive nucleosomes down the fiber that makes two-nucleosome steps, similar to Micro-C^6^ (Fig. 4j). On the other hand, the indirect contact separation density (α to γ) has a contact probability curve with enrichment of even nucleosome contacts within an envelope of decreasing contact frequency, which is consistent with alternating nucleosome stacking, or a partially ordered “zig-zag” chromatin fiber conformation (Fig. 4k). This increased sensitivity to the geometry of the chromatin fiber evidenced by indirect steps in CADwalks may be due to constraints on ligation placed by the geometry of crosslinked, stacked nucleosomes (Fig. 4l), which are relieved by allowing an intervening nucleosome to complete the contact. Across all of our data, indirect contacts similarly enable high-resolution footprinting readout of local nucleosome spacing (Extended Data Fig. 9b). We note that the n+1 step is missing in indirect contacts because we enforced that each step is unidirectional along the chromosome coordinate. Epigenetic states show some variation in the short range contact probability, with the strongest difference evident at the active promoter state, with the weakest ‘zig-zag’ contact probability, which may reflect a more open acetylated chromatin state or more disordered nucleosome positioning^69–71^. Collectively, these results indicate that the ability of CAD to cut nucleosomal and sub-nucleosomal fragments enables CADwalks to detect high-resolution chromatin fiber folding states^71^.

### CADwalks span multiple chromatin organization length scales, exhibiting both local confinement and focal long-range contacts

Following up on the observation that indirect contacts have a longer genomic separation distribution than directly ligated fragments (Fig. 4i), we asked whether and how CADwalks reflect chromatin structure constraints across different genomic length scales. Defining the “walk span”, as the maximum cis-chromosomal separation in linear genomic distance within a CADwalk molecule as a metric, we asked how it scales with the number of steps in a CADwalk (Fig. 5a). Individual walks, even with the same number of steps, can have a range of walk spans varying orders of magnitude (10^3^ - 10^8^ bp) (Fig. 5a). Additionally, walk spans of any step number partition into three prominent regimes that reflect orders of genome organization in the single-step *P*(*s*) curve: Regime 1 (0–10 kb, chromatin fiber, nucleosome clutch and gene domain scale), Regime 2 (10 kb–3.16 Mb, loop extrusion scale), and Regime 3 (3.16 Mb–1 Gb, compartmental scale). These different regimes are also evident in the net displacement distribution of two-step walks (αγ distance) (Fig. 5b).

**Figure 5.**
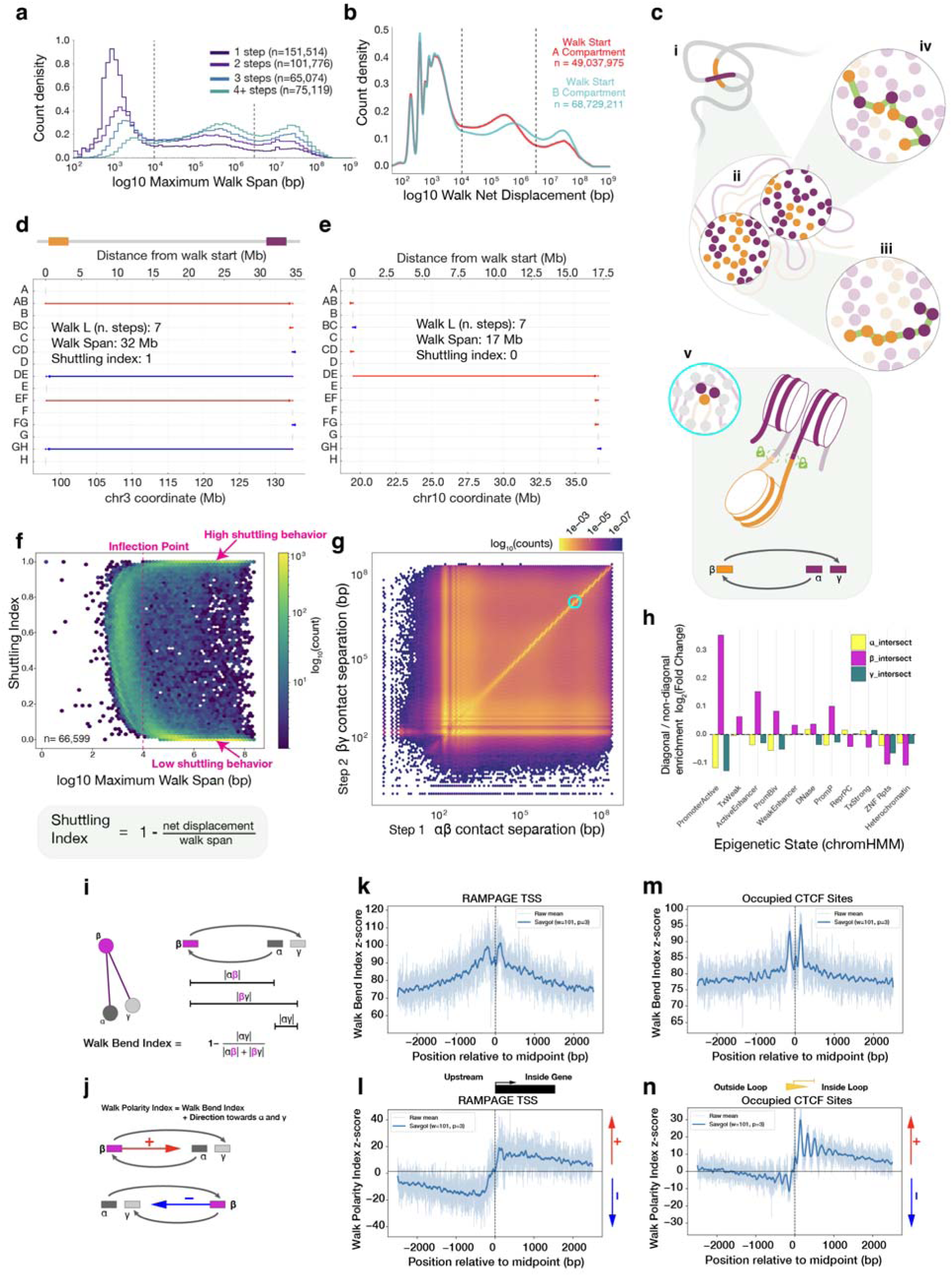
CADwalk shuttling. (a) Walk span, calculated as the maximum within-run genomic separation (Dmax) distributions for cis-confined, single chromosome CADwalk runs, stratified by number of steps in a walk (uninterrupted UU-run walk length, L). (b) Net displacement within cis-chromosomal two-step α-β-γ walks calculated from abs(γ_start-α), where α is always the fragment with the lowest coordinate in genome order. Walks are stratified by A and B compartment annotation by α fragment genomic coordinate from compartment calls (Fig. 2). Dotted lines represent the genomic distance regime boundaries. (c) Diagram of nucleosome ligation within the nucleus: i. Genome regions in proximity in 3D space, ii. Nucleosomes from different genome regions in situ, iii. Ligation of nucleosomes from different genome regions in situ with low shuttling, iv. Ligation of nucleosomes from different genome regions in situ with high shuttling, v. Tri-nucleosome contact in situ, zoomed in on proximity ligation of cartoon α-β-γ walk where β is an intervening, venturing nucleosome (from a distal genomic region to proximally located α and γ). (d) Representative CADwalk with high shuttling index. Walk segments are labeled A-H in read order (schematic shown in Fig. 4b). (e) As above, but showing a representative CADwalk with low shuttling index. (f) Distribution of run-level shuttling index (also called tortuosity) (Equation 4), and its relationship to walk span, calculated as the maximum within-run cis separation, D_max_. Shuttling index versus log_10_ (D_max_ [bp]) for runs with L=2 (2 steps). D_max_ is the maximum same-chromosomal span of fragment midpoints within a run, taken per chromosome and then maximized across chromosomes. Shuttling index is defined per walk run as 1-(net displacement/*D*_max_) where steps within a cis-chromosomal walk have lengths > 0. (g) Joint distribution of αβ distance vs βγ distance from cis-chromosomal two-step α-β-γ walks. Hexbin intensities are normalized to 1 and color values are shown with a log10 transform. (h) Log₂ enrichment of walk density in diagonal walks vs all other walks by position (α, β, or γ) stratified by fragment intersection with ChromHMM state. Diagonal walks were defined by step-length similarity, where (|log αβ distance|-|log βγ distance| < 0.2). Enrichment was calculated to all cis-chromosomal walks annotated to ChromHMM states as a baseline. (i) Definition of the Absolute Walk Bend Index (AWB). For a three-fragment walk with steps αβ and βγ and span αγ, F is near zero for linear α→β→γ paths and increases toward 1 as the walk doubles back so that β lies outside the α–γ interval and the two legs are of similar length (symmetric bend). (j) Definition of the Walk Polarity Index (WPI), which multiplies F by the sign of β’s offset from the α–γ midpoint, assigning positive values when β lies upstream of the midpoint and negative values when β lies downstream. Together, AWB and WPI provide per–walk, per–site bend and polarity scores. (k) Metaplot of zAWB (z-score of the AWB) around RAMPAGE–defined TSSs, oriented such that transcription proceeds to the right (“Inside gene”). Thin curves show the raw mean and the dark trace shows Savitzky–Golay^73^–smoothed values. (l) zAWB around occupied CTCF motifs, oriented by motif strand so that the positive side lies to the right (“Inside loop”). (m) Signed polarity zWPI (z-score of the WPI) around TSSs. (n) zWPI around occupied CTCF motifs, revealing a strongly asymmetric polarity: the interior (+1, +2, +3) nucleosomes show a focused positive WPI, whereas the upstream (–1) nucleosome exhibits a weaker negative polarity. Illustration in (c) created by Oscar Alejandro Perez.

We asked to what extent the net displacement distribution of two-step walks (our deeper dataset) depends on the chromatin compartment within which they begin, the A (active) and B (inactive) compartments. We observe that the net displacement distribution is nearly the same for the smallest regime of contacts, < 10 kb, but diverges, with shorter-range contacts favored in the A compartment (Fig. 5b). While this shift to shorter distances in the A compartment is consistent with previous observations using lower-resolution chromosome walk mapping^72^, our data reveals that there is a smaller-scale regime < 10 kb that does not shift, indicating a length-scale-specific behavior.

As expected, the distribution of maximum walk span shifts to longer distances as walks with more steps are considered (Fig. 5a). However, we wondered whether this scaling with step number was consistent with a random walk with step sizes sampled from the single-step distribution evident in our *P*(*s*) curve. We constructed null distributions by shuffling individual steps into synthetic walks (Supplementary Fig. 4a-b). By comparison, the maximum walk span distributions of the real CADwalks with 3 or 4 steps were shifted toward shorter distances than the corresponding null model (Supplementary Fig. 4a-b), indicating that walks are more confined than would be expected from the *P*(*s*) curve and a random stepping model.

### CADwalk shuttling at long-range contacts

Motivated by the observation that chromatin contacts occur in different step size regimes with different behaviors, we next asked whether CADwalks can reveal the nature of chromatin interactions at distal locus contacts. To what extent do long-range chromatin looping contacts consist of intermingling of the short-range chromatin domains at each anchor (Fig. 5c)? What types of chromatin are involved in short- or long-range contacts?

We first visualized long-multi-step CADwalks from long-read sequencing using what we term “runway plots” of consecutive steps mapped onto the genomic coordinate (Fig. 5d-e). We observed that many multi-step long-range walks consist of consecutive steps of similar magnitude but opposite direction, which we term “shuttling” (Fig. 5c-iv,e). To quantify shuttling, we defined the shuttling index as *1 − (net displacement / walk span)* (Fig. 5c,f), which is invariant to the direction of the walk. Intuitively, shuttling index approaches 0 for straight paths (Fig. 5c-iii,d), and approaches 1 for trajectories in which successive steps nearly cancel the total displacement (Fig. 5c-iv,e). Visualizing how the shuttling index changes with total walk span in two-step CADwalks, we observed that there is a distinct change in behavior from a broad range of values (as would be characteristic of walks like that shown in Fig. 4e) to a strong dichotomy between very high and very low shuttling index (exemplified by Fig. 5d-e) at walk spans beyond ∼10 kb. Analyzing walks with 3 or 4 steps, we observe similar behavior, with a novel population of shuttling index values appearing for walks with odd numbers of long-range steps (Supplementary Fig. 4c-d).

To gain further insights into this type of shuttling behavior and the relationships between consecutive step sizes, we turned to our two-step, three fragment (α–β–γ) short-read CADwalk dataset (Fig. 4b-c). We calculated the joint probability distribution as a function of step size for the two consecutive steps within each short-read CADwalk (Fig. 5g). We observed a strong density on the diagonal, consisting of walks with the same step size. Further analysis showed that this observation is consistent with the high-shuttling index walks we observed in the long-read CADwalks (Fig. 5f), as can be seen by segmenting the two-step long-read CADwalks by shuttling index profile and plotting their joint step size probability (Supplementary Fig. 4e-h). Interestingly, the diagonal emerges from a broader regime of different step size combinations beyond the ∼ 10 kb length scale, recapitulating our results showing a shift of behavior at ∼10 kb from the long-read walk analysis (Fig. 5a,f). This is also observable more clearly in a calculation of the correlation between step sizes for walks on and off the diagonal (Extended Data Fig. 10a), where the correlation begins to rise at ∼10 kb.

We asked to what extent this kind of strong correlation between step sizes might be expected from the distribution of walk steps and walk spans in our CADwalk libraries. We built a synthetic three-fragment, two-step (α–β–γ) walk model by sampling step sizes and walk start locations from our ultra-deep short-read data to test whether successive cis separation steps are independent (Supplementary Note, Supplementary Fig. 5). In the simplest version of the model (Model 1), we took all steps to be independent, while in a α–γ distance-limited version of the model (Model 2), we required that the α–γ distance distribution follow the empirical data. In both cases, we did not observe as strong a correlation between step sizes or a strong diagonal in the joint probability matrix as in the experimental data (Supplementary Fig. 5c-d).

The high prevalence of shuttling walks may be due to the enhanced likelihood of subsequent crosslinks to a captured distal locus once one ligation has been established. However, the inherent asymmetry of a three-fragment walk between the middle β nucleosome “venturing” to make a distal contact and its flanking nucleosomes α and γ, which are relatively close together (Fig. 5c-v), presented us with an opportunity to ask about the biological identity of the “venturing” nucleosome as opposed to its flankers. Interested in what biological factors were correlated with these shuttling long-range two-step walks (Fig. 5g, diagonal), we segmented them based on the symmetry between walk steps (|log αβ step length|-|log βγ step length| < 0.2; see Methods for details). For each of the three positions in these on-diagonal CADwalks, we calculated the enrichment of each epigenetic state (imputed 12-state chromHMM^68^) relative to all other off-diagonal CADwalks. We find that on-diagonal shuttling walks have middle (β) fragments that are more likely to be from active regulatory regions such as enhancers or promoters (Active promoter ∼0.3 log_2_FC, or ∼23% enrichment, and active enhancer ∼0.1 log_2_FC ∼7.18% enrichment) (Fig. 5h). Our results indicate that these venturing nucleosomes that make distal contacts are more likely to come from open chromatin regions than their flanking nucleosomes.

To formalize venturing behavior per molecule, we defined two per-walk indices. The Absolute Walk Bend (AWB) compresses the full two-step geometry of each α–β–γ walk into a bounded bend (venturing) magnitude (Fig. 5i), and the Walk Polarity Index (WPI) attaches a sign that records whether the β fragment lies upstream or downstream of the α–γ midpoint (Fig. 5j). Unlike classical Hi-C directionality or insulation measures, which are computed directly from aggregated upstream versus downstream pairwise contacts, AWB and WPI are assigned once per short CADwalk and only then averaged across molecules.

Genome-wide robust Z-scored AWB and WPI tracks revealed sharp, nucleosome-scale signatures at both TSSs (Fig. 5k-l) and occupied CTCF motifs (Fig. 5m-n). AWB peaks adjacent to both landmarks, with slightly sharper profiles at CTCF. Polarity was more diagnostic: TSSs showed a near-antisymmetric WPI pattern with opposite signs upstream and downstream and a steep sign change at the TSS, consistent with venturing returns arriving from both sides. In contrast, oriented CTCF sites showed a pronounced asymmetry between the −1 and +1 nucleosomes, with stronger positive polarity on the loop-interior side. When rescaled around convergent CTCF loop domains, polarity reversed sign across the two anchors, consistent with an oriented, CTCF-aligned bias that is compatible with uni-directional cohesin loop extrusion (Extended Data Fig. 10b). Together, these analyses connect run-level shuttling in long-read CADwalks to locus-resolved venturing in short CADwalks, enabling the generation of nucleosome-scale bend and polarity genome tracks (Extended Data Fig. 10c).

## Discussion

Despite the broad utility and ongoing development of proximity ligation-based techniques demonstrated over the last decade^74^, the limited enzymatic repertoire available for the chromatin fragmentation step remains a common constraint across all 3C-based methods. Here, we demonstrated that CAD, when engineered to be specifically cleavable by TEVp, can be seamlessly integrated into a Hi-C/Micro-C-like workflow and can further simplify that workflow by obviating the need for end repair, end labeling or ligation junction enrichment. CAD digestion of chromatin preserves sub-nucleosome sized fragments, has reduced GC sequence content bias relative to MNase and is robust to changes in enzyme concentration. This robustness is underscored by the fact that once we determined saturating enzyme concentrations, generating a successful library in a new cell type required minimal titration. The simple and efficient proximity ligation after CAD digestion allowed us to generate CAD-C libraries that yielded consistent results despite variations in the protocol.

We anticipate that CAD-C will extend the applicability of nucleosome-resolution proximity ligation to investigate, for example, enhancer-promoter looping or chromatin domain boundaries in rare or limited samples which have heretofore been incompatible with the high-input requirements of Micro-C, due to its need for extensive MNase titration and less efficient protocol that requires end-repair of DNA fragments.

The high ligation efficiency of CAD-C drives several beneficial properties of the assay. Rare interactions such as loops involving promoters are detected at higher frequency and there is an enrichment for sub-kilobase distance contacts, which we interpret as reflecting the physiological reality that most chromatin interactions are local. CAD-C preferentially captures enhancer- and promoter-associated loops, which are critical for gene regulation, thereby providing a valuable tool for studying both 3D genome architecture and gene regulatory networks, as shown at the PPIF locus.

The same ligation efficiency also enables a novel analysis mode: nucleosome-scale chromosome walks, or CADwalks, read out by sequencing the multi-contact concatemers that form upon proximity ligation in CAD-C experiments. We observed that on the local scale, CADwalk segments can be directly used for footprinting of both nucleosomes and sub-nucleosomal particles, and that even short two-step CADwalks can reveal differences in chromatin fiber folding by probing indirect contacts between nucleosomes that may be sterically incompatible with efficient direct ligation. The zig-zag contact signature we observed in GM12878 cell CADwalks was stronger in indirect CADwalk contacts than in direct CADwalk contacts and is consistent with other observations of alternate nucleosome stacking contacts^6,71,75^. Although we and others have begun to use short-range Micro-C contacts as an indicator of local chromatin compaction^76–78^, small changes in the over-digestion of nucleosomes by MNase between samples or between genomic loci have the potential to cause artifactual changes in the contact probability curve^78^. CAD-C’s improved robustness to enzyme concentration therefore holds promise for proximity ligation becoming a more reliable tool for short-range chromatin compaction measurements.

On longer length scales, CADwalks are multi-scale objects that span both local exploration of a nanoscale chromatin domain and long-range interactions. Our analysis of step-size correlations revealed that symmetric walks often occur, with two equal steps venturing to a distal locus and coming back to the same chromatin neighborhood. We found that venturing nucleosomes are enriched in open chromatin and active regulatory regions. The *in situ* nature of CAD-C allows us to infer that a single walk must have come from a single nucleus (if not necessarily from a single sister chromatid or homologous chromosome). Nucleosome-scale features of walks can therefore report on the extent of intermingling between chromatin fibers from distal interacting loci. Such intermingling has been observed for chromatin fibers with 10n+5 bp nucleosome spacing^79^, but its relevance at specific genomic loci is not yet understood. The shuttling behavior observed in long-read CADwalks is suggests that such close interdigitation does indeed occur, although the relatively low throughput of the long-read data set makes it difficult to obtain statistical enrichments at this point. CADwalks are well-poised to enable future studies to investigate this point in greater detail with deeper data sets, and to reveal the physical nature of chromatin intermingling at both silenced loci and at interacting active regulatory regions.

Local features of walks can reveal novel phenomena. For example, we observed that the venturing middle nucleosomes in two-step walks are enriched at open chromatin sites. Taking a per-walk molecular view to analyze walk polarities, we found that there is strong, hyper-local, nucleosome-scale insulation and directional facilitation of contacts at CTCF-bound insulation boundaries, consistent with the loop extrusion model^80,81^. This manifests as a nearly symmetric divergence of walk polarity on either side of a TSS, as expected for an insulating element, and as walks that are on the loop side of a CTCF site strongly being polarized to occur within the loop. The ability to observe this insulation at the walk and the nucleosomal scale reinforces the highly local effects of CTCF stalling of the cohesin complex and its insulation properties, down to the nucleosome scale. Finally, we found that at around 10kb, there is a shift from random exploration of a local chromatin neighborhood with a low correlation between step sizes to a shuttling regime in which long-ranging steps are often paired with similarly sized steps back to the original locus. This length scale coincides with the stronger shoulder observed in the *P*(*s*) curve. It may mark a shift from a partially disordered chromatin globule in which contacts between nucleosomes are common, toward a regime in which chromatin interactions become less random and perhaps need to be facilitated by long-range mediators of looping such as loop extrusion. Interestingly, this length scale is similar to the ∼20 kb distance over which enhancer-promoter interactions become dependent upon loop extrusion by cohesin^82^.

The examples of observations described above show that there is still much to be understood and measured at the mesoscale organization of chromatin. In addition to the multi-scale measurements we performed as proof-of-principle in this paper, there are many additional potential applications for the unique benefits of CAD-C. The simplified Direct CAD-C protocol does not require capture of biotinylated fragment ligation junctions if digestion of chromatin into mostly mono-nucleosomal fragments is efficient. It can therefore serve as a platform for derivative assays that involve biotin-labeling and capture to target sequencing to regions of interest. Furthermore, CAD has the potential to serve as a platform for RNA-DNA interaction assays. Activated DFF40/CAD is competitively inhibited by intact nuclear RNA (and other polyanions)^55^. CAD-C therefore includes an RNase step after fixation and before digestion to ensure robust, reproducible fragmentation. Because CAD does not degrade RNA, this step decouples DNA fragmentation from RNA loss (in contrast to MNase, which cleaves both DNA and RNA^83^) and could be treated as a tunable control variable. In an experiment without RNase treatment, CAD activity dropped by ∼10-fold yet still yielded predominantly mononucleosome-scale fragments (Supplementary Fig. 4m), motivating potential “RNA-retaining” CAD-C variants that could naturally with RNA–DNA proximity and multiway approaches (ChAR-seq^84^/GRID-seq^85^; RD-SPRITE^86^) that highlight RNA’s role in nuclear organization.

Although the integration of CAD with a Micro-C protocol is straightforward, and our protocol for expressing and purifying the recombinant engineered CAD/ICAD heterodimer has been optimized for high yield, producing the enzyme with this protocol requires access to fast protein liquid chromatography. We have not yet explored the full range of optimizations of the CAD-C protocol that may increase its efficiency and broad accessibility to a variety of research teams. In particular, it is likely that the optimal sample crosslinking conditions for CAD-C, the importance of which to 3C data quality has been explored previously^3,87–90^, may differ from those for Micro-C, which we used here.

Overall, we find through the proof-of-concept experiments described here, that CAD-C is a flexible and robust tool for probing the multi-scale organization of chromatin, from sub-nucleosomal footprints to long-range looping, compartmental, or trans-chromosomal interactions. Its relative ease of use and simplified protocol is likely to enable a new generation of proximity ligation assays targeting the still poorly understood mesoscale domain of chromatin organization.

## Supporting information

Supplementary Note on Ligation Gating Model

Custom code

## Acknowledgments

We are grateful to Dr. Anders Hansen, Dr. Viraat Goel, Dr. Deena A. Oren and Dr. Elphege Nora for helpful discussions. We thank the Rockefeller University Reference Genome Resource Center, Genomics Resource Center, and High Performance Computing Resource Center for technical support. We thank Oscar Alejandro Perez for assistance with figure illustrations and formatting. This work was supported by the NIH Office of the Director (New Innovator Award 1DP2GM150021 to V.I.R.); Irma T. Hirschl/Monique Weill-Caulier Trust (Career Scientist Award to V.I.R.); The Rita Allen Foundation (Scholar Grant to V.I.R.); Stavros Niarchos Foundation Institute for Global Infectious Disease Research at Rockefeller University (grant to V.I.R.); Robertson Technology Development Fund (Grant to J.S. and V.I.R.). Boehringer Ingelheim Fonds (PhD Fellowship to J.S.). NSF GRFP (PhD Fellowship to L.A.W.); Human Frontier Science Program (Postdoctoral Cross-Disciplinary Fellowship to A.O); NSERC (Postgraduate Scholarships to H.C. and J.L.Y.); and Japan Society for the Promotion of Science (Overseas Research Fellowship to H.A.K.). We thank Dr. Iestyn Whitehouse and Dr. Axel Delamarre for sharing pre-publication results describing their use of a CAD-C variant in budding yeast.

## Disclosures

V.I.R. and J.S. are inventors on a related patent application covering CAD-C (PCT application filed 2024).

## Author contributions

These authors contributed equally: L.A.W. and W.Z. J.S.: conceptualization, methodology, software, investigation, formal analysis, visualization, writing-original draft, data curation, supervision, funding acquisition. L.A.W.: conceptualization, methodology, software, investigation, formal analysis, visualization, writing--original draft, data curation, writing--review and editing; W.Z.: conceptualization, software, methodology, investigation, visualization, formal analysis, writing--original draft, data curation, validation, writing-review and editing. A.O.: investigation, formal analysis, data curation, writing--review and editing. H.C.: conceptualization, methodology, writing—original draft, writing-review and editing. J.H.: methodology, investigation, visualization. J.L.Y.: investigation. H.A.K.: methodology, resources. E.B.C.: investigation. C.W.: investigation. J.B.: methodology, supervision, resources. G.F.: methodology, supervision, resources, writing-review and editing. V.I.R.: conceptualization, methodology, data curation, writing--review and editing, supervision, project administration, funding acquisition.

## Code availability

Analysis code is available as Supplementary Information and more details on how custom code was used to analyze CADwalks is available as Supplementary Methods.

## Data and reagent availability

Plasmids for CAD/ICAD expression are available upon request. A CAD/ICAD expression construct sequence and map are available as Supplementary Information. Raw data generated in this study has been deposited to GEO under accession numbers: CAD-C: GSE318108; CADwalk: GSE318239; CAD-seq: GSE318105; Micro-C: GSE318103; CUT&Tag: GSE318107; ATAC-seq: GSE318018.

## Methods

### CAD-ICAD^2xTEV^ construct design and expression

Full-length Mus musculus CAD (Dffb/DFF-40; UniProt O54788) and ICAD (Dffa/DFF-45; UniProt O54786) coding sequences were retrieved from UniProt^91^ and reverse-translated with Escherichia coli–optimized codons using GenSmart™, followed by manual refinement in SnapGene. To enable protease-controlled activation, ICAD was engineered with two TEV protease recognition sites replacing segments of the endogenous caspase cleavage sites (aa 114–116 and 221–223 deleted; ENLYFQS inserted), yielding a 339-aa ICAD^2×TEV^ variant. The modified sequences were synthesized as gene blocks (IDT) and cloned into a custom dual-expression pRSFDuet-1–derived vector, incorporating optimized T7 promoters^92^ and translational leader sequences (TIR-2) for high-level E. coli expression. CAD was N-terminally fused to a Twin-Strep-tag, muGFP^93^, and SUMO^Eu1^^94^ separated by flexible linkers (including GGSGGSGGS and GDGAGLIN sequences), facilitating affinity purification of the CAD:ICAD heterodimer.

### Protein Expression and Purification

pRSFDuet-1_ICAD2xTEV_TwinStrepTag-muGFP-SUMOEu1-CAD plasmids was first propagated in NEB Stable E. coli to minimize leaky expression and then transformed into NEB T7 Express lysY/Iq cells (NEB C3013) with 50 µg/mL kanamycin selection for protein production. Cultures started from single colonies were grown in Terrific Broth supplemented with 0.4% glycerol, 50 µg/mL kanamycin, 0.005% antifoam 204 agent at 37 °C to OD600 1.0–1.4, induced with 1 mM IPTG, and shifted to 18 °C for 18–24 h with shaking at 280-320 rpm in double-baffled flasks. Cells harvested by pelleting at 5,000-10,000 xg at 4 °C and stored frozen at −80°C in pellets corresponding to 500 mL culture. Thawed pellets were washed, and lysed by sonication in 4 mL lysis buffer (CLysB: 20 mM HEPES-NaOH pH 7.8, 100 mM NaCl, 1 mM EDTA, 10% glycerol, 0.01% Triton X-100, 10 mM βME, 1x cOmplete EDTA-free (Roche), 200 U/mL Dr. Nuc (Syd Labs)) per gram of wet pellet. Clarified lysates (centrifuged at 21,000 xg at 4 °C for 30 minutes and 0.22 µm filtered) were applied to a Strep-Tactin XT 4Flow affinity column (IBA Lifesciences), washed with CSWB buffer (20 mM HEPES-NaOH pH 7.8, 150 mM NaCl, 1 mM EDTA, 10% glycerol, 0.01% Triton X-100, 10 mM βME), and eluted with biotin-containing IBA-BXT buffer (100 mM Tris-HCl pH 8.0, 150 mM NaCl, 1 mM EDTA, 50 mM biotin, 20 mM βME). Fractions containing the highest muGFP fluorescence were pooled and dialyzed overnight (10 MWCO) into CSEC buffer (20 mM HEPES-HCl pH 7.5, 100 mM NaCl, 1 mM EDTA, 10% glycerol, 0.01% CHAPS, 10 mM DTT).

The CAD:ICAD^2×TEV^ complex was further purified by size-exclusion chromatography (HiLoad Superdex 200 pg 16/600 column on an ÄKTA Pure instrument) in CSEC buffer. Void volume peak fractions corresponding to intact heterodimers were pooled, concentrated 14-fold (Amicon Ultra-15 30 MWCO), supplemented with 25% v/v glycerol (UltraPure, Invitrogen), aliquoted into protein LoBind tubes (Eppendorf) and stored at −20 °C, and used for downstream activity assays. This procedure was used to prepare CAD/ICAD for the assays described in this paper, unless otherwise stated.

We have since optimized high-salt preparations (Lot 3) with the following changes. We used larger culture volumes, freezing pellets from 1L of culture each. Lysis was performed at 500 mM NaCl in 4 mL/g CLysB–500 (20 mM HEPES–NaOH pH 7.8, 500 mM NaCl, 1 mM EDTA, 10% v/v glycerol, 0.01% Triton X-100, 10 mM β-mercaptoethanol, 1× cOmplete EDTA-free protease inhibitor). After sonication, MgCl_2_ was added to 3 mM and Benzonase Salt Tolerant Endonuclease (Millipore Sigma) was added to 100 U/mL, then incubated 60 min at 4°C with gentle rocking prior to clarification and filtration (0.2 µm). Affinity purification was performed with a 5 mL StreplllTactinXT 4Flow high-capacity column washed with CSWB–500 (20 mM HEPES–NaOH pH 7.8, 500 mM NaCl, 1 mM EDTA, 10% v/v glycerol, 0.01% Triton X-100, 10 mM β-mercaptoethanol) and eluted with six 0.5 column volume fractions of freshly prepared Buffer CEB: 10× IBA Buffer BXT (1 M Tris-Cl, 1.5 M NaCl, 10 mM EDTA, 500 mM biotin, pH 8) diluted to 1× with CAD Elution Diluent (CED; 20 mM HEPES-HCl pH 7.5, 100 mM NaCl, 1 mM EDTA, 10% glycerol, 20 mM β-mercaptoethanol). Fractions with the highest muGFP signal were held at 4°C, pooled and concentrated ∼two-fold (Amicon Ultra-15 30 kDa MWCO) (Supplementary Fig. 1a). Size exclusion chromatography was performed as above. 2.5 mL fractions comprising the second peak (Supplementary Fig. 1b-d) were supplemented to 25% v/v glycerol, snap-frozen and stored at −80 °C, yielding ∼1 mg/mL tagged CAD at ≥62.5 U/µL as measured with the activity assay described below. To minimize aberrant heat-induced cleavage during routine SDS–PAGE, samples were typically denatured at a lower temperature (70–75°C).

### CAD Activity Assay

The enzymatic activity of soluble CAD:ICAD^2×TEV^ was measured using purified λ genomic DNA in 1× WDB-70 buffer (70 mM KCl, 3 mM MgCl₂, 1 mM DTT, 10 mM HEPES–KOH, pH 7.5). Standard reactions contained 2.5 µg λ DNA (NEB) in 25 µL, initiated by adding CAD (typically 12.2 µM tagged; 81.9 kDa) together with TEV protease and incubated 15 min at 30 °C followed by 45 min at 37 °C, then briefly heated to 65 °C to denature CAD. Reactions were quenched with 0.77% (w/v) SDS and analyzed on 1% agarose gels. One CAD unit (1 U) was defined as the amount of tagged CAD (estimated as 0.041 µg, ∼0.5 pmol for Lot 1) required to digest 1 µg λ DNA to fragments <100 bp under these conditions. Stock preparations, including Lot 3, were titrated using this assay, with Lot 3 showing ≥2.5-fold higher activity than Lot 1 (Supplementary Figure 2). Assay variations (e.g., reduced DTT, addition of BSA, or shortened incubation) were applied for specific QC measurements.

### Disuccinimidyl glutarate (DSG) synthesis

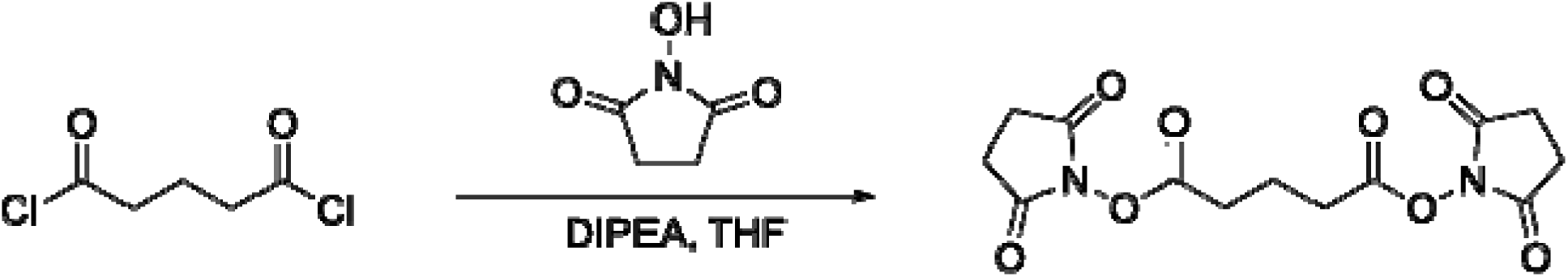

DSG was synthesized based on a previously reported protocol^95^, with slight modifications. In a 2-neck flask under argon, N-hydroxysuccinimide (1.5 g, 13 mmol), dry tetrahydrofuran (50 mL) and a stir bar were added. The flask was cooled to 0 °C (ice bath) and N,N-diisopropylethylamine (5 mL, 28.6 mmol) was added slowly with stirring. Glutaryl chloride (755 uL, 5.92 mmol) was then added dropwise. The reaction (now an off-white suspension) was removed from the ice bath, then warmed to room temperature. After 4 hours, the reaction was stopped and the THF was removed *in vacuo*. The residue was resuspended in dry dichloromethane (50 mL), washed with brine, water, dried over anhydrous sodium sulfate and filtered. The product was purified by recrystallization in dichloromethane/isopropanol and then dried overnight *in vacuo*. The product was obtained as a white crystalline solid (1.49 g, 77%). Analytical data was in accordance to the previously reported compound^95^.

^1^**H NMR** (600 MHz, CDCl_3_): 2.84 (br, 8H), 2.80 (t, *J* = 7.2 Hz, 4H), 2.19 (quint, *J* = 7.2 Hz, 2H).

**^13^C NMR** (150 MHz, CDCl_3_): 169.1, 167.8, 29.7, 25.7, 19.7.

**HRMS** (ESI+): m/z calculated for C_13_H_15_N_2_O_8_^+^ [M+H] 327.0823 and found 327.0823

### CAD Digestion Assay (CAD-seq)

Fixed chromatin (1% methanol-free formaldehyde, 10 min at room temperature) CAD digestions were performed as described below to validate TEVp-dependent activation on nuclei. K562 (ATCC CCL243), hTERT RPE-1 (ATCC CRL-4000), and GM12878 (Coriell) cells were cultured under standard conditions and harvested at defined densities, with hTERT RPE-1 cells induced into a quiescent (G0) state by contact inhibition. Cells were washed with pre-warmed DPBS and fixed either with 1% methanol-free formaldehyde (Thermo Fisher Scientific, 28908) alone or using dual crosslinking with 3 mM DSG (freshly prepared 300 mM stock in DMSO) for 35 min followed by 1% formaldehyde for 10 min. Fixation was quenched with Tris–HCl pH 7.5 to a final concentration of 750 mM, and cells were pelleted, washed twice with ice-cold DPBS, resuspended to 4 × 10^6 cells/mL, and snap-frozen in liquid nitrogen for storage at −80 °C.

Frozen pellets were lysed on ice in 1× WDB-10 buffer (10 mM HEPES–KOH pH 7.5, 10 mM KCl, 3 mM MgCl₂, 1 mM DTT) supplemented with 0.2% NP-40 to isolate nuclei, which were then treated with RNase (RNase A, Thermo Scientific EN0531, 0.36 µg/µL for hTERT RPE-1; RNase T1, Thermo Scientific EN0542, 10 U/µL for K562 and GM12878) at 30–37 °C for 15 min. CAD digestion was initiated by adding 20 µL Peak 1 TST–muGFP–SUMO^Eu1^–CAD:ICAD^2×TEV^ proenzyme and 5 µL TEV protease (NEB, P8112S, Lot 3) per reaction, incubated for 20 min at 30 °C followed by 1 h at 37 °C with shaking at 1000 rpm. Reactions were quenched with 1 mM ZnCl₂ and washed with 1× WDB-10 + 0.2% NP-40. For titration experiments, GM12878 nuclei (4 × 10^6 per replicate) were digested with CAD at 4–500 U in 1× WDB-10. DNA was deproteinated, crosslinks reversed with Proteinase K (NEB, P8107S, 20 µg/µL) in the presence of SDS and NaCl overnight at 65 °C, and purified using the Genomic DNA Clean & Concentrator-10 kit (Zymo, D4011) with a 1:5 sample-to-binding-buffer ratio to retain low-molecular-weight fragments. Purified DNA (50–700 bp) was used to prepare libraries with the NEBNext Ultra II DNA Library Prep Kit for Illumina (NEB, E7645L) using recommended modifications to retain short fragments (excess adapters were removed using ∼2.3× the reaction volume of Ampure XP beads). Adapter ligation, USER enzyme treatment, and size selection with Ampure XP beads were performed according to the manufacturer’s instructions, and libraries were indexed (NEB, E6441A) with 4 PCR cycles, cleaned with 1.2× Ampure XP beads, and quantified by Qubit dsDNA HS and Agilent HS D1000 TapeStation analysis. CAD-seq libraries were sequenced on Illumina NextSeq 1000 (2×60 bp) or NovaSeq X/X Plus (2×150 bp) platforms.

### Classic CAD-C

Crosslinked cell pellets (5 × 10^6 cells) were thawed on ice and resuspended in 1 mL ice-cold 1× WDB-10 buffer (10 mM HEPES–KOH pH 7.5, 10 mM KCl, 3 mM MgCl□, 1 mM DTT) supplemented with 0.2% NP-40. After a 20 min incubation on ice with intermittent pipetting, nuclei were pelleted (1,750 × g, 5 min, 4 °C), washed twice with 1× WDB-10 + 0.2% NP-40, and resuspended in 200 µL WDB-10. RNase treatment was performed using RNase A (Thermo EN0531, 7.5 µL of 10 mg/mL stock; hTERT-RPE1) or RNase T1 (Thermo EN0542, 10 U/µL; K562 and GM12878) for 15 min at 30–37 °C with shaking. CAD digestion was initiated by adding 20 µL recombinant TST–muGFP–SUMO^Eu1^–CAD:ICAD^2×TEV^proenzyme (25 U/µL; Lot 1) and pre-incubating at 30 °C for 5 min, followed by TEV protease activation (NEB P8112; 5 µL, 10 U/µL). Reactions proceeded for 20 min at 30 °C and 60 min at 37 °C with shaking, then quenched with 1 mM ZnCl□. Nuclei were washed sequentially in WDB-10 with and without 0.2% NP-40 and resuspended in WDB-70. A fraction was deproteinized using SDS and Proteinase K (Thermo Scientific, 20 mg/mL) overnight at 65 °C as a digestion control.

Nuclei were pelleted (5 min, 2,000×9, 4 ◦C) and the supernatant was removed. End repair was performed by adding 55 µL H2O, 10 µL 10× NEBuffer 2.1, 20 µL 10 mM ATP, and 5 µL 100 mM DTT to each pellet (90 µL total), followed by 5 µL T4 polynucleotide kinase (10 U µL −1; NEB M0201L) and 10 µL DNA Polymerase I, Large (Klenow) Fragment (5 U µL −1; NEB M0210), giving a 105 µL reaction. Reactions were incubated for 30 min at 37 ◦C with shaking (1,000 rpm). Biotinylated nucleotides were incorporated by supplementing each 105 µL end-repair reaction with 10 µL 1 mM biotin-dATP (Jena Bioscience NU-835-BIO14-S), 10 µL 1 mM biotin-dCTP (NU809-BIOX-S), 1 µL 10 mM dTTP, 1 µL 10 mM dGTP, 5 µL 10× T4 DNA ligase buffer, 0.25 µL 20 mg mL−1 BSA, and 22.75 µL H2O (50 µL supplement; 155 µL total). Reactions were shifted to 25 ◦C and incubated for 1 h in a thermomixer (Eppendorf). Nuclei were pelleted (2,000×9, 5 min, 4 ◦C), washed once with 1 mL WDB-10, and resuspended for proximity ligation.

Nuclei were pelleted, washed, and subjected to in situ proximity ligation in a 500 µL reaction with 10,000 U T4 DNA ligase (NEB, 400 U/µL) in T4 DNA ligase buffer containing 0.1 mg/mL BSA, incubated overnight at 25 °C. Nuclei were pelleted (3,000×9, 5 min, 4 ◦C) and residual 3′ biotin on unligated ends was removed by Exonuclease III treatment (NEB, 100 U/µL; 1000 U in a 200 µL reaction incubated 15 min at 37 ◦C with interval mixing). Crosslinks were reversed by adding 26 µL 20 mg mL−1 proteinase K, 26 µL 10% SDS, 10.4 µL 5 M NaCl, and 2.6 µL 10 mg mL−1 RNase A to each 200 µL Exonuclease III reaction (265 µL total) and incubating at 65 °C for 48 h. DNA was purified either by phenol–chloroform–isoamyl alcohol extraction with ethanol precipitation or using the Genomic DNA Clean & Concentrator-10 kit (Zymo, D4011). Purified DNA was sheared to ∼220 bp (Covaris S220, Covaris microTUBE AFA Fiber Pre-Slit Snap-Cap, 180 seconds), biotinylated fragments captured on MyOne Streptavidin C1 Dynabeads (Thermo Fisher), and on-bead libraries prepared using the NEBNext Ultra II DNA Library Prep Kit (NEB E7645L/E6441A). Libraries were indexed, cleaned with Ampure XP beads, and quality-controlled by Agilent TapeStation and Qubit dsDNA HS prior to Illumina NovaSeq X sequencing (2 × 150 bp).

### Express CAD-C

Express CAD-C libraries (GM12878) used purified, crosslink-reversed post-Exonuclease III DNA from the Classic workflow. Downstream sonication and streptavidin-capture steps were replaced by NEBNext UltraExpress FS library preparation (NEB). Each technical replicate was assembled with 200 ng input DNA, fragmented at 37 °C for 5 min followed by 65 °C for 15 min, and ligated to NEBNext hairpin adapters. Dual-unique indexing primers (NEB E6440) were added to assign indices, and libraries were cleaned and amplified with limited-cycle PCR (6 cycles for 200 ng input), targeting ∼275 bp insert size. Libraries were eluted in 35 µL 0.1× TE, quantified by Qubit, and analyzed with Agilent TapeStation before sequencing on a single Illumina NovaSeq X Plus 25B flow cell (2 × 150 bp).

### Direct CAD-C / CADwalk

Direct CAD-C (“CADwalk”) libraries were generated from GM12878 nuclei. CAD digestion and proximity ligation were performed as in the Classic workflow, omitting end repair and biotin labeling. For PacBio long-read libraries (Large concatemers, >1 kb), ligated concatemer DNA was gel-purified (500 ng concatemer DNA per well, Invitrogen 2% E-Gel EX agarose Gel, Zymo DNA Gel Recovery Kit), adapter-ligated with NEBNext Ultra II hairpin adapters, indexed using dual-unique primers, PCR-amplified with Watchmaker polymerase (10 cycles, 5 min extension), and purified with Ampure XP beads. Libraries were used for PacBio Sequel IIe HiFi SMRTbell preparation and sequencing at the Vertebrate Genome Laboratory (VGL), Rockefeller University using a Sequel Iie platform. Illumina Direct-CADwalk libraries (Small ∼300–500 bp; Medium ∼500–1,000 bp) were size-selected from the same concatemer pool, prepared with the NEBNext Ultra II hairpin-adapter protocol, and PCR-amplified with Watchmaker Master Mix. Extension times were adjusted according to the expected insert size: ∼35–45 s for Small, ∼60 s for Medium, and 5 min (300 s) for Large. Libraries were pooled equimolarly and sequenced on a single Illumina NovaSeq 6000 SP flow cell (2 × 250 bp).

### Micro-C

Micro-C was performed following a protocol from Slobodyanyuk et al.^56^ In brief, confluent RPE-1 hTERT (CRL-4000) cells were dissociated with TrypLE, washed and harvested. Cells were crosslinked with 1% formaldehyde for 15 minutes, quenched with Tris-HCl (final conc. 750 mM), and then crosslinked with 3 mM DSG (homemade, see SI), and quenched with Tris-HCl pH 7.5 (final conc. 750 mM). Doubly crosslinked cells were washed with PBS and then aliquoted into tubes and flash frozen (1 M cells per tube). MNase (NEB, lot no. 10155112) was then added to digest the chromatin to approximately 80:20 to 90:10 ratio of mono:di-nucleosome. The NEB Ultra DNA library prep kit was used to prepare micro-C libraries. Micro-C libraries were sequenced on the Novaseq X (Novogene) to approximately 1.4 B total fragments (150×150 PE) among 12 libraries.

### ATAC-seq

ATAC-seq was performed following the Omni-ATAC^96^ protocol using pG-Tn5 prepared as described in Soroczynski et al.^97^ ATAC-seq libraries were sequenced on the NextSeq1000 (50×50 PE). Nextera adapters were trimmed using cutadapt and then aligned to GRCh38 using bowtie2 (--end-to-end --very-sensitive --no-mixed --no-discordant --phred33 -I 10 -X 700). Peaks were called using MACS2 using default settings.

### CUT&TAG

CUT&TAG was performed following the protocol of Kaya-Okur et al.^98^ with some modifications (wash buffers were supplemented with digitonin, 0.05% w/v). pG-Tn5 was purified from E. coli following Soroczynski et al.^97^ In brief, cells were harvested, and nuclei were isolated. 100,000 nuclei were bound to Concanavalin-A beads (EpiCypher) and incubated overnight (4 °C) with primary antibodies (H3K4me1 (Abcam; 1:100), Pol2ser5 (CST; 1:50)). The next day, the beads were washed, incubated with Guinea pig anti-rabbit IgG (antibodies Online, ABIN101961), pAG-Tn5 (1:100) and activated with magnesium. The tagmentation reaction was incubated at 37 °C for 1 hour, washed with TAPS buffer, and released with SDS buffer. SDS was quenched with Triton-X100 (final 0.67% v/v), and libraries were directly PCR-amplified with NEBNext High Fidelity Master Mix (14 cycles). Libraries were purified with AmpureXP (0.9x) and quantified by Tapestation and Qubit. CUT&TAG libraries were sequenced on the NextSeq1000 (50×50 PE) to approximately 5-10M reads per biological replicate. Adapters were trimmed using cutadapt and then aligned to GRCh38 using bowtie2 (--end-to-end --very-sensitive --no-mixed --no-discordant --phred33 -I 10 -X 700). Peaks were called using MACS2^99^.

### CAD-seq Analysis

Paired-end BAMs were converted to RPGC-normalized bigWig coverage using deepTools bamCoverage (hg38, autosomes only). Fragment coverage was calculated by extending read pairs, and insert size-stratified tracks were generated for nucleosomal (100–200 bp) and sub-nucleosomal (<100 bp) fragments. Midpoint coverage was obtained using the -MNase option. Metaprofiles were computed as the mean signal around feature centers, including occupied CTCF motifs (JASPAR^100^ MA0139.1; top 5% by ChIP-seq or Cut&Run signal), ENCODE^64^ DNase footprints, and TSS/TES regions. Fragment-length distributions were derived from properly paired autosomal reads (MAPQ ≥30), downsampled to equal depth across replicates, and quantified as alignment file |TLEN| histograms with counts per million normalization. GC-balanced fragment-end composition and genome-wide GC-coverage bias were assessed by stratifying 1 kb windows into GC quantiles, sampling equal numbers of fragments per stratum, and normalizing coverage to the mean of filtered bins. Pairwise concordance among coverage tracks was evaluated by tiling the genome into 5 kbp bins, extracting mean signal, excluding zero or extreme bins, and calculating Pearson, Spearman, and Kendall correlations after mean-normalization. All scripts recorded parameters, contig sets, and software versions to ensure reproducibility.

### CAD-C Data Processing and Analyses

Computational processing of CAD-C datasets involved alignment, quality control, and multi-scale contact mapping. Paired-end Illumina and PacBio HiFi reads were aligned to the human reference genome (hg38) using bwa-mem or bwa-mem2^101^, and converted to BAM with SAMtools^102^. BAM files were processed with a SLURM-wrapped pairtools workflow (Python 3.12.11, pairtools^103^ 1.1.3, SAMtools 1.22) to parse contacts (pairtools parse2 -max-inter-align-gap 30, -min-mapq 30, -drop-sam, -report-position junction, -report-orientation junction, -add-pair-index, -no-expand, -no-flip), correct orientations, remove duplicates, and generate ligation-junction–aware .pairs files. Junction nucleotide composition, fragment coverage, and GC-stratified metrics were computed with custom Python scripts.

Downstream analyses included generation of single-base BigWig coverage tracks, orientation-aware local contact pileups anchored at CTCF motifs, and visualization of ligation-gated profiles. Contact matrices were binned at 500 bp, normalized using iterative correction (ICE), and assembled into multi-resolution .mcool files for genome-wide and locus-specific analysis. “Direct” walk types were selected using the predicate (walk_pair_type == “R1” or “R2” or “R1&2”) and unique–unique pairs were sub-selected and again de-duplicated to generate the “LJO” pairs subset containing walks with ligation junctions observed. All scripts and workflows recorded software versions, parameters, and provenance to ensure reproducibility and traceability. Code writing, refactoring, debugging, and documentation were assisted by OpenAI ChatGPT Pro (models o1, o3, GPT-5, GPT-5.1, and GPT-5.2 Pro). All model outputs were reviewed by the authors; final code and outputs/figures were validated against expected controls.

### *P(s)* plots

Intrachromosomal contact frequency–distance curves were derived from 500 bp .mcool matrices using a custom Python script. The script calculated cis contact frequencies, converted them to base-pair separations to generate the contact-probability function, and estimated the local slope of log *P(s)* versus log separation.

### Compartment analysis

Compartments were computed from 10-kb resolution cooler matrices using the “eigs-cis” program in the cooltools package (v0.7.1)^50^, with the parameter “--ignore-diags 1”. The first principal component (eigenvector 1, E1) for selected chromosomes or genomic regions was generated using a custom R script. Genomic regions with E1 > 0 were assigned to the A compartment, while those with E1 < 0 were assigned to the B compartment. To quantify compartment strength across different methods, saddle plots were generated using the cooltools “saddle” script with “--qrange 0.02 0.98”. The compartment strength, defined as (AA + BB) / (2AB), was calculated by extracting the top and bottom 10% (5 bins each) from the saddle matrix and computing their mean values.

### Insulation analysis

The insulation profiles of GM12878 OmniC (cantatabio.com), Dovetail Micro-C (cantatabio.com), *in situ* Micro-C (https://zenodo.org/records/15303879)^57^, Hi-C (4DN: 4DNFIXP4QG5B)^59^, and CAD-C, as well as hTERT-RPE1 in situ Micro-C, CAD-C, and Hi-C (GEO: GSE71831)^51^, were processed using cooler matrices, and insulation scores were computed with the cooltools “insulation” function at 10-kb resolution. Insulation indices were obtained by averaging signals within a 500-kb insulation window. The Pearson correlation coefficient of the insulation indices was used to quantify similarity between datasets.

TAD boundaries were identified using default settings, and a boundary strength threshold of 0.1 was applied to retain the most prominent boundaries. These boundaries were further classified into Micro-C-only / Hi-C-only, common, and CAD-C-only categories, with a flank region size of 20 kb, using bedtools^104^. Insulation score profiles across different subgroups were analyzed using computeMatrix from deepTools (v3.5.3) within a ±200-kb window centered on the TAD boundaries, with a 10-kb bin size. Pileups of 500 kb genomic regions flanking the TAD boundaries were performed on Micro-C and CAD-C datasets, binned at a 10 kb resolution, using coolpup.py (version 1.1.0) in local mode^105^. Normalization was conducted based on expected interaction probabilities at each genomic distance.

### Loop calling

Loop calling for CAD-C and Micro-C data was performed using mustache^58^. Mustache identified loops at bin sizes of 1kb, 2.5 kb, 5 kb, and 10 kb, with a sparsity threshold of 0.7 to filter out loops in sparse regions. The P-value threshold for interactions was set at 0.1. Loop anchors were classified using the pairtopair function of bedtools v2.31.1^104^, with flank regions defined as two bin sizes.

### ChromHMM state enrichment analysis of loop anchors

ChromHMM state enrichment at loop anchors was quantified using a custom Python workflow. Loop anchors from each category (Micro-C–specific, Common, and CAD-C–specific) were converted to fixed-width BED intervals and intersected with a GM12878 12-state ChromHMM segmentation to compute base-pair overlaps per state (hg38, https://egg2.wustl.edu/roadmap/data/byFileType/chromhmmSegmentations/ChmmModels/core <u>Marks/jointModel/final/</u>). For each state, observed anchor coverage was normalized by total anchor base pairs and compared to the genome-wide fractional coverage of that state to derive log2 enrichment values. Enrichment was computed independently for each loop category using the same ChromHMM map, enabling direct comparison across assays. All intermediate overlaps and summary statistics were recorded for reproducibility.

### Loop–feature intersection and cross**lZI**set comparison

We quantified overlap between genomic features and chromatin loops with a custom Python script. For each of the three loop sets (Micro-C–specific, Common, and CAD-C–specific), loop anchors were derived from BEDPE files and intersected with feature annotations (e.g., ENCODE RAMPAGE TSS) using base-pair overlap. At the loop level, anchors were scored binarily for feature overlap, and loops were classified as having zero, one, both, or any anchors overlapping a feature. Anchor- and loop-level summary metrics were computed for each set. For cross-set comparisons, feature identities detected in each loop category were combined to define disjoint overlap classes, which were visualized using UpSet plots. Method-specific differences in feature detection were evaluated using paired statistical tests on the union of detected features, with effect sizes reported as odds ratios and confidence intervals. All analyses were performed with a reproducible, logged workflow.

### RAMPAGE TSS–Loop Anchor Intersections and Cross-Set Analysis

Chromatin interaction loops were analyzed for their association with transcription start sites (TSSs) using a custom computational workflow. TSSs were represented as genomic intervals with unique feature identifiers. Loop anchors were defined directly from the original interaction coordinates and intersected with TSS intervals using a base-pair–level overlap criterion. Loops were classified according to whether zero, one, or both anchors overlapped at least one TSS.

Analyses were performed separately for Micro-C–specific, shared, and CAD-C–specific loop sets. For each set, we summarized the TSS features associated with loop anchors, stratified loops by TSS overlap class, and quantified anchor-level hit rates and feature densities. To compare TSS association patterns across datasets, features were assigned to shared or dataset-specific categories; when a reference TSS set was used, features not associated with any loop set were classified as neither. Statistical differences between Micro-C and CAD-C datasets were evaluated using paired or binomial tests, as appropriate. Overlap patterns were visualized using set-based intersection plots, and all analyses were conducted with parameter, input data and version logging for reproducibility.

### Contact matrix pileups

Pileup analysis of Hi-C / Micro-C / CAD-C /HiChIP contact matrices was performed using a custom Python wrapper built around the coolpuppy package^105^. Contact matrices were analyzed from multiresolution .mcool files at a common bin size (specified per analysis). Loop-centered pileups were generated from BEDPE loop coordinates, and feature-centered pileups from BED intervals, using fixed genomic windows or rescaled coordinates as indicated. When available, genome-wide cis expected values were used for normalization; otherwise, pileups were normalized to randomly shifted control regions. Pileup heatmaps were rendered using matplotlib.

The HiChIP data for Cohesin and H3K27Ac originate from GSE80820 (GSM2138324, GSM2138325, GSM2138326, GSM2138327) and GSE101498 (GSM2705041, GSM2705042)^66,67^. HiCLift was used to convert these datasets from the hg19 to the hg38 genome assembly^106^, followed by cooler to merge sample replicates.

### Loop anchors pileup analysis

To investigate differences in chromatin loops detected by Micro-C and CAD-C in GM12878 and hTERT-RPE1 cells, we analyzed loop anchors classified as Micro-C-only, common, and CAD-C-only. For GM12878, we utilized datasets from ENCODE^64^, including ATAC-seq (ENCFF667MDI) and ChIP-seq for various factors: CTCF (ENCFF680XUD), Rad21 (ENCFF407HJR), POLR2A (ENCFF322DRU). For hTERT-RPE1 cells, we referenced GSE196727^107^, which includes ChIP-seq data for CTCF (GSM5899605), Rad21 (GSM5899607) and our in-house ATAC-seq and Pol2ser5 CUT&Tag. To examine signal distribution across different loop anchors, we applied the computeMatrix function from deepTools (v3.5.3), centering on loop anchors and aggregating signals within a ±25 kb flanking region, with a bin size of 100 bp^108^. Furthermore, GRO-seq data for GM12878 was obtained from GEO accession GSM1480326^65^. After lifting over from hg19 to hg38 with CrossMap^109^ and merging plus and minus strands, we analyzed nascent transcription signals across different loop anchor groups using computeMatrix from deepTools (v3.5.3) within a ±25 kb flanking region centered on the anchors, with a bin size of 1 kb.

### Short, two-step α-β-γ CADwalk analyses

To analyze short-range three-fragment chromatin walks, we focused on two consecutive ligation junctions derived from individual proximity-ligation concatemers, corresponding to A–B–C fragment configurations (represented as α-β-γ in figures to avoid confusion with A/B nuclear compartments). Reads were restricted to those containing exactly two consecutive junctions, ensuring that the two interaction steps shared a single intermediate fragment. Candidate walks were required to traverse a single digestion fragment at the intermediate position, enforced by genomic contiguity within 500 bp and strand-orientation constraints (opposite strands if coming from separate reads), thereby isolating bona fide two-step walks that share a common CAD-cleaved B fragment of less than 500 bp, rather than independent ligation events. Validated A–B–C walks were reconstructed into three genomic fragments with standardized coordinates and orientations. Records were deduplicated, flipped as necessary to a consistent genomic order (to enforce an A start coordinate lower than the C start coordinate), and separated into cis and trans configurations based on chromosomal identity. Cis walks were further stratified by the relative genomic ordering of the three fragments (ABC, ACB, or BAC), enabling positional analysis of multi-fragment contact structures. Analyses were performed using pairtools parse2 and custom python scripts.

Fragments from ABC walks were annotated using BedTools and custom python scripts by intersecting fragment start positions with genomic state annotations, including chromatin compartments and 12-state GM12878 hg38-liftover ChromHMM states (sourced from: https://egg2.wustl.edu/roadmap/data/byFileType/chromhmmSegmentations/ChmmModels/coreMarks/jointModel/final/). A custom R script used to assign each in the three-fragment walk (A, B, C) an A/B compartment label based on its intersection with nuclear compartments called using GM12878 Micro-C. Fragments that did not intersect with A or B compartment were treated as missing/invalid. For walk-type–stratified analyses, records were retained only if AB_distance, BC_distance, and AC_distance were present and the walk-type label contained no missing compartment calls.

These annotations were used to stratify subsequent analyses by chromatin context. Fragment length distributions were computed separately for A, B, and C fragments within each genomic state, providing state-specific fragment size profiles. To characterize the spatial organization of two-step walks, we analyzed the distances between fragments and local contact patterns. Separation histograms were truncated to short genomic ranges of less than 5000 bp between fragment midpoints to emphasize local contact structure and were visualized as state-stratified density profiles to generate indirect and direct contact probability curves. For cis walks, fragment midpoint–midpoint contacts were aggregated within fixed windows centered on genomic features of interest, yielding local contact matrices for direct (A–B, B–C) and indirect (A–C) interactions. Strand-specific profiles were combined by symmetry to generate strand-agnostic representations.

### Two-step α-β-γ CADwalk analyses around CTCF-bound sites

We further examined motif-centered organization by aligning ABC walks to fixed windows around CTCF motifs. Fragment coverage, midpoint distributions, and fragment length–distance relationships were quantified relative to motif centers, producing aggregated coverage profiles and V-plots without further normalization. Aligned CAD fragments were potentially counted more than once if they contributed to more than one window around a motif center. All analyses pooled fragments across walk positions unless otherwise specified and were performed with a reproducible custom python script. The script does not take CTCF motif or TSS strand into account. We therefore separately calculated coverage for minus- and plus-strand CTCF motifs and flipped minus strand results before combining them. For two-dimensional plots of density at fragment-pair midpoint coordinates, pairs were plotted only when both midpoints fall within the same coordinate window being represented. As for profiles, matrices were calculated separately for plus and minus strand window centers, and minus strand matrices flipped before combining.

For pseudo-4C analysis of walks intersecting viewpoints at three loop anchors of nested loops, three-fragment walks were selected in which any of the three fragments ovelaps the viewpoint region. Pileups were then generated for each viewpoint and walks were shown as a horizontal gray span from minimum start to maximum end of its three fragment coordinates, and the fragments intersecting any viewpoint were shown in magenta.

### Directionality and B-enrichment analysis

Genome-wide directionality of three-fragment CAD walks was quantified using a B-Enrichment Index (BEI), which measures the relative enrichment of intermediate (B) fragments compared with anchor (A and C) fragments at a particular locus. Walks were oriented such that fragment A lies upstream of fragment C, and fragment midpoints were aggregated into fixed-width genomic bins with default sizes of 1 kb for whole-genome BigWig files and 1 bp for per-site walk metric summaries (e.g. at CTCF binding sites or TSSs). For each bin, counts of A, B, and C fragments were converted to compositional proportions using Laplace smoothing (Equation 1), and BEI (Equation 2) was defined as a linear transformation of the B-fragment proportion, with positive values indicating enrichment of the interior nucleosome fragment (B), negative values indicating enrichment of the flanking nucleosome fragments (A+C).

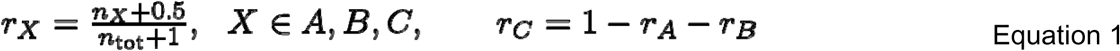

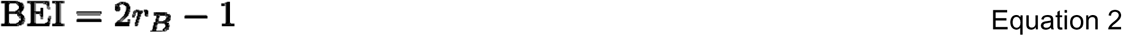

To remove large-scale genomic trends, log-ratio metrics comparing anchor and interior fragments were detrended separately for each chromosome using a linear model, and residuals were converted to robust Z-scores based on the median absolute deviation. Bins with fewer than 50 total fragment counts were excluded from log-ratio calculations. BEI values were similarly standardized on a per-chromosome basis. Genome-wide tracks of detrended and standardized metrics were exported as BigWig files at the analysis bin resolution and used for downstream visualization and pileup analyses.

### Walk polarity, genome-wide aggregation, and walk-class analyses

Non-linear returning three-fragment CAD walks were quantified using a Walk Polarity Index (WPI), defined as a scale-free fold fraction of the fragment separations in absolute genomic distance F (Equation 3):

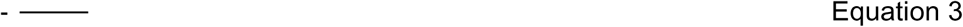

signed by the displacement of fragment B relative to the midpoint of A and C (WPI∈[−1,1]). F=0 for walks in which B lies between A and C and increases with the distance of B away from the A-C interval and it is positive when B is to the left in genomic coordinate relative to the A-C midpoint. Walks were pre-oriented with A to the left of C, and WPI values were assigned to the B-fragment midpoint and aggregated genome-wide; site-level means were converted to per-chromosome robust Z-scores using median absolute deviation, producing single-base, zero-centered BigWig tracks. Strand-aware meta-profiles were computed by piling up BigWig tracks over genomic annotations, including CTCF motifs and RAMPAGE TSSs, with midpoint-centered windows aligned to strand and directional signals flipped for minus-strand features (i.e. the sign of the signal was inverted for the minus-strand profile for WPI, but not for BEI). Profiles were smoothed with a Savitzky–Golay filter^73^.

For each three-fragment walk (A, B, C), pairwise cis separations were calculated as absolute differences between fragment start coordinates A_start, B_start, C_start: AB_distance = abs(B_start-A_start), BC_distance = abs(C_start-B_start), and AC_distance = abs (C_start-A_start). Cis-chromosomal walk geometry was classified using pairwise distances (AB, BC, AC), with diagonal walks defined by similar step lengths (|log_10_(AB) – log_10_(BC)| < 0.2) and remaining walks labeled as other, and for some purposes sub-classified into linear (with the B fragment lying between A and C along the chromosome coordinate) or non-linear. Step-length relationships were visualized with log_10_-scaled hexbin plots with a log_10_-scaled color scale, and correlations assessed via Spearman statistics in AB distance quantile bins. Fragments were annotated by A/B compartment and ChromHMM state, enabling stratification by compartment transitions (e.g. A→B→A) and per-position log-enrichment of ChromHMM states relative to genome-wide baseline. Diagonal log2 enrichment (over full data) was divided by “other” log2 enrichment (over full data) to visualize differences in per-position ChromHMM state. Walks with fragments for which the compartment assignment was “NA” were excluded.

### Synthetic two-step walk models

Synthetic two-step α–β–γ null models were generated using custom R code by inverse transform sampling AB and BC distances from spline-fitted empirical distributions. The log-distance histogram with 500 equal bins was fitted with a 20-knot cubic regression spline. The first anchor (“A”) was randomly sampled from the pool of empirical α-β-γ fragment anchors annotated to a nuclear compartment, and subsequent anchor locations were based on the distances sampled from the log-distance distribution spline model. An independent-assortment model sampling AB and BC step distances independently to yield implied AC distances, and a constrained model additionally enforcing AC distances consistent with the observed distribution. Walks with coordinates falling outside the starting chromosome were not accepted (i.e. *trans* walks were not modeled). A minimum step size of 100 bp was enforced, and 100M synthetic walks were generated for each model. These synthetic walks served as references for geometric and enrichment comparisons.

### Long-read CADwalk analyses

PacBio CADwalks were parsed into uninterrupted unique-unique (UU) runs using pairtools parse2 and represented in a canonical orientation based on terminal fragment geometry, enabling consistent representation of both fragment contiguity with a walk pair index and genome coordinate order across reads. Each run was decomposed into fragment-, step-, and run-level representations, from which we quantified fragment midpoints, *cis/trans* composition, longest cis blocks, alternation rates, longitudinal genomic span (D_max_), and a run-level shuttling (tortuosity) index reflecting a combination of directional step reversals and step sizes.

For each uninterrupted *cis* block *b* within a run, and for a given geometry *x* ∈ {*J*, *M*}, we write each step *j* as a signed length (*d^x^*, *s^x^*) with *d^x^* > 0 and *s^x^* ∈ {±1} indicating genomic direction. In the junction geometry (*x*=*J*), lengths are absolute junction separations 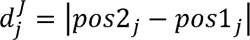; in the midpoint geometry (*X=M*), we first assign each fragment a genomic midpoint *m* and then take 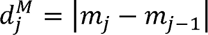. For each block *b* we define the net signed path length and accumulated path length as

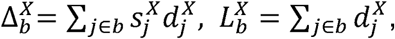

and the run-level shuttling index (tortuosity) in geometry *X* as

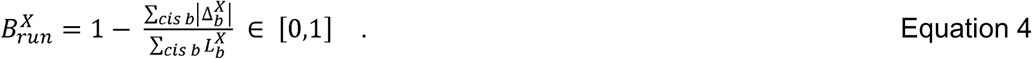

This construction is orientation-invariant and weights each *cis* block in proportion to its length. We used the midpoint geometry unless otherwise stated.

Fragment-length and cis-separation distributions were aggregated across runs and technical replicates, and trans-chromosomal behavior was summarized as run-length–dependent trans-step rates and chromosome novelty. Cis-only runs were further analyzed by projecting stepwise genomic separations into log–log space and stratifying by robustly Z-scored shuttling index, visualized using hexbin density maps. To assess whether observed longitudinal spans exceeded geometric expectations, empirical D_max_ distributions within fixed run-length strata were compared to resampled null models generated by step-length resampling under fragment-size and cis-geometry constraints (see above). Finally, individual CADwalks were visualized on genome coordinates using standardized A–B–C–D–E fragment labels based on their order within a read, ensuring internal consistency between fragment order, step polarity, and genomic coordinates.

---

**Extended Data Figure 1.**
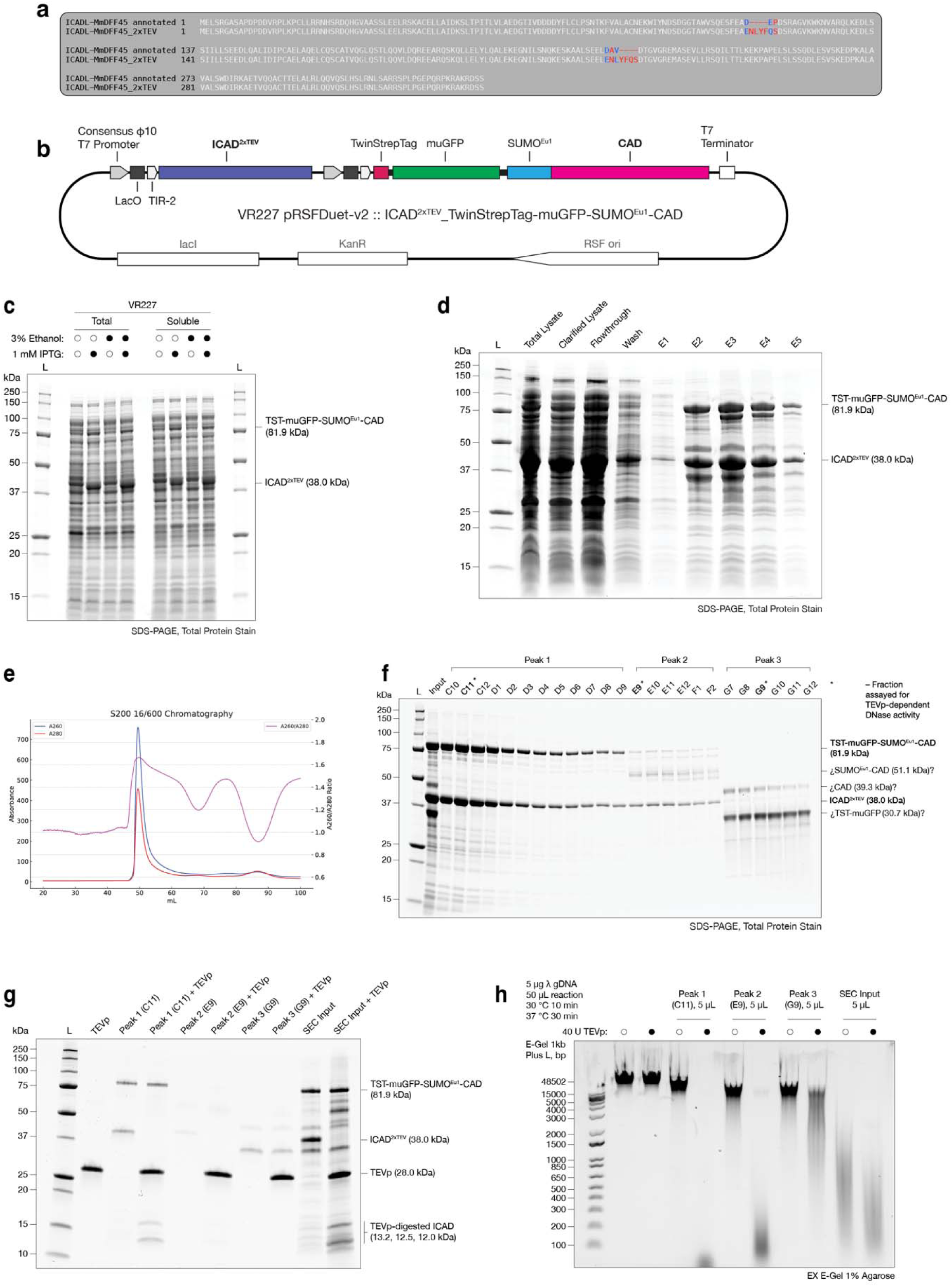
Engineering design, express and purification of pRSFDuet-1_ICAD^2xTEV^_TwinStrepTag-muGFP-SUMO^Eu1^-CAD construct. (a) Sequence of wild-type *Mus musculus* ICAD (DFF45) (UniProt: O54786) and engineered, TEVp-cleavable ICAD^2xTEV^ protein sequence in pRSFDuet-1_ICAD^2xTEV^_TwinStrepTag-muGFP-SUMO^Eu1^-CAD construct. (b) Map of pRSFDuet-1_ICAD^2xTEV^_TwinStrepTag-muGFP-SUMO^Eu1^-CAD construct. (c) Validation of soluble over-expression of ICAD^2xTEV^:TwinStrepTag-muGFP-SUMO^Eu1^-CAD heterodimer in *E. coli*. Supplementation of culture with 3% final ethanol does not further increase solubility. (d) TwinStrepTag affinity chromatography of ICAD^2xTEV^:TwinStrepTag-muGFP-SUMO^Eu1^-CAD heterodimer. Clarified lysate from 4 g wet pellet was filtered (0.22 µm) and loaded onto one 1 mL Strep-TactinXT 4Flow column. (e) Size exclusion chromatography A260, A280 profile, using S200 16/600 column on an ÄKTA Pure instrument and separated with 1 mL/min flow of 1x CSEC into 0.8 mL fractions in 96 deep-well collection plates. (f) SDS-PAGE of select SEC fractions. Peak 1 apex = 49.3 mL, Peak 2 apex = 66.9 mL, Peak 3 apex = 86.1 mL. (g) SDS-PAGE of select SEC fractions corresponding to peaks 1, 2 and 3 incubated with TEVp. (h) Lambda genomic DNA assay for TEVp-specific DNase activity of SEC fractions from major peak apexes and SEC input material. DNA resolved on a 1% agarose gel stained with SybrGold stain (EX E-Gel).

**Extended Data Figure 2.**
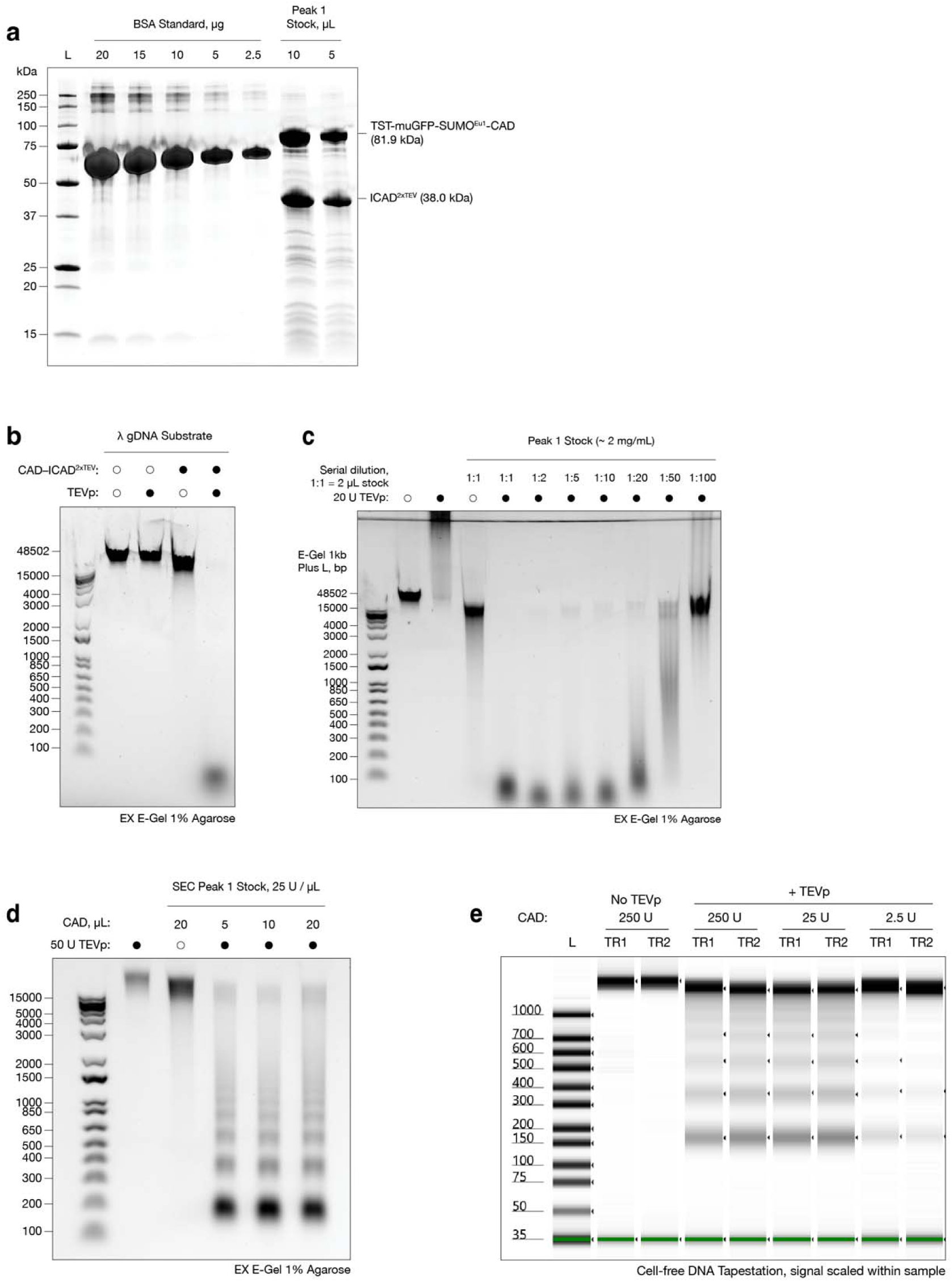
Characterization of ICAD^2xTEV^:TwinStrepTag-muGFP-SUMO^Eu1^-CAD stock. (a) Confirmation of TwinStrepTag-muGFP-SUMO^Eu1^-CAD:ICAD^2xTEV^ proenzyme heterodimer stock against BSA standard. (b) TEVp-activatable, specific activity of CAD enzyme assayed on lambda phage genomic DNA. (c) TEVp activation increases CAD enzyme activity ≈ 100-fold, showing low non-specific nuclease contamination of the ICAD^2xTEV^-CAD proenzyme. DNase activity assay using lambda genomic DNA. (d) CAD digestion of 3 mM DSG + 1% formaldehyde-fixed human hTERT-RPE1 cell line nuclei. Titration of CAD shows no discernable over-digestion across a 4-fold concentration range. (e) Digestion carried out on cells as in (d), but across a 100-fold CAD concentration range. Digestion material DNA analyzed using Agilent cell-free DNA Tape, signal scaled within sample.

**Extended Data Figure 3.**
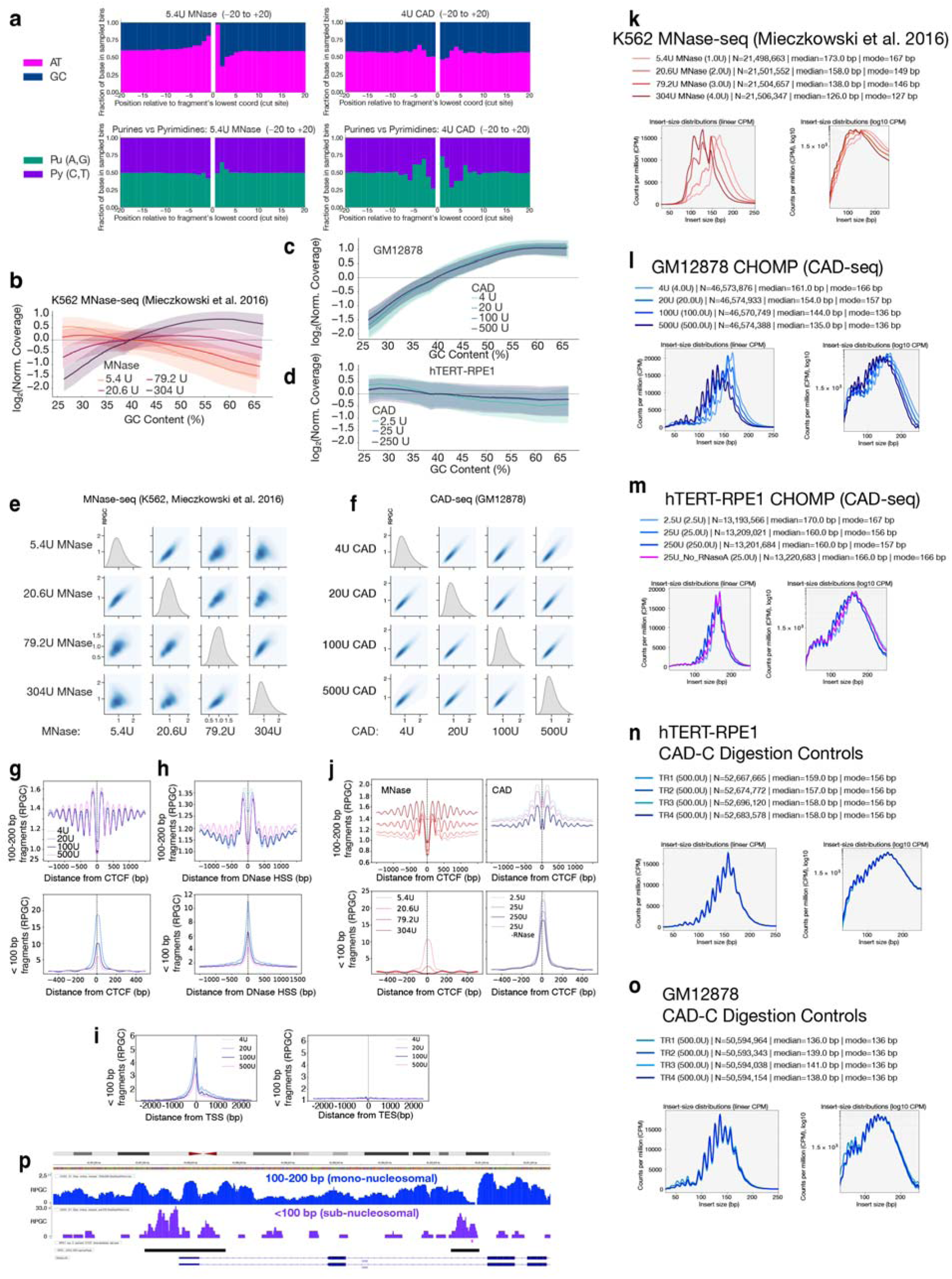
Characterization of chromatin digestion properties of ICAD^2xTEV^:TwinStrepTag-muGFP-SUMO^Eu1^-CAD and benchmarking against MNase-seq. (a) Per-position nucleotide frequencies from −20 to +20 bp relative to each fragment start (inferred cut) for a K562 MACC 5.4 U MNase dataset^47^ and a GM12878 CAD digestion library prepared with 4 U CAD as in Fig. 1(h), top row = A,T vs. G,C content, bottom row = purine (A,G) vs. pyrimidine (C,T) (b) K562 MACC MNase titration (5.4, 20.6, 79.2 and 304 U). (c) Cycling GM12878 cells digested with 4, 20, 100 and 500 U CAD. (d) Quiescent hTERT–RPE1 cells digested with 2.5, 25 and 250 U CAD (CAD-seq libraries). (e) Pairwise kernel-density plots of RPGC coverage in 5 kb autosomal bins for the K562 MACC MNase titration (5.4–304 U). (f) Pairwise kernel-density plots for the GM12878 CAD digestion titration (4–500 U CAD). (g) GM12878 CAD titration (4–500 U) fragment coverage, stratified by mapped fragment length: 100–200 bp mononucleosomal fragments flank occupied CTCF motifs (top) and <100 bp sub-nucleosomal fragments (bottom). (h) GM12878 CAD titration (4–500 U) as in (g) plotted around ENCODE DNase hypersensitive sites (HSS). (i) GM12878 <100 bp CAD fragments around annotated transcription start sites (TSS) and transcription end sites (TES). (j) Fragment size-stratified coverage at occupied CTCF sites. K562 MACC MNase titration (5.4–304 U) 100-200 bp fragment coverage (top) <100 bp fragment coverage at the motif center (bottom). Right: hTERT-RPE1 CAD titration (2.5–250 U; the 25 U –RNase A trace is overlaid) 100-200 bp fragment coverage (top) <100 bp fragment coverage (bottom). (k) Fragment-length (insert-size) distributions for a K562 MACC MNase-seq titration (5.4, 20.6, 79.2 and 304 U), from autosomal, properly paired primary alignments (MAPQ ≥ 30) after downsampling each series to the shallowest eligible-pair count; CPM normalization uses 0–750 bp fragments as the denominator, and curves are shown for the 30–250 bp range on linear and log10 CPM axes (Methods). (l) GM12878 CAD-seq CAD titration (4, 20, 100 and 500 U), processed identically to (k); two technical replicates per condition were merged prior to plotting. (m) hTERT–RPE1 CAD-seq CAD titration (2.5, 25 and 250 U), processed identically to (k), with a matched 25 U digest prepared without RNase A overlaid. (n) hTERT–RPE1 Classic CAD-C digestion-control technical replicates (TR1–TR4; 500 U CAD concentration). (o) GM12878 Classic CAD-C digestion-control technical replicates (TR1–TR4; 500 U CAD concentration). (p) In hTERT-RPE1 cells, 100–200 bp (blue) and <100 bp (purple) CAD digest fragment coverage at the CDC6 locus. hTERT-RPE1 ATAC-seq narrow peaks are shown in black.

**Extended Data Figure 4.**
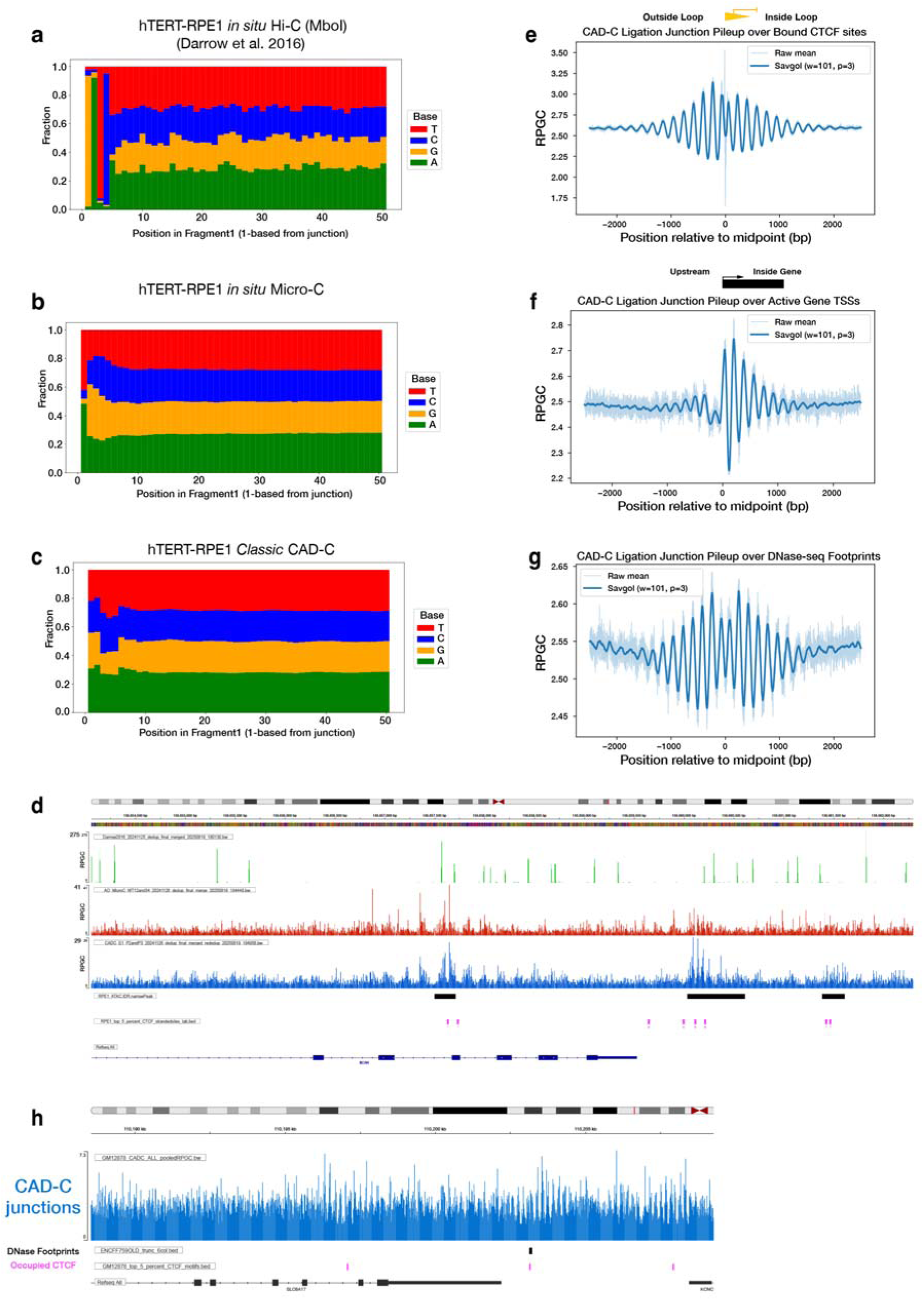
Analysis of biases and genome coverage in CAD-C library ligation junctions. (a–c) Junction-proximal base composition from ligation junctions in hTERT–RPE1 for (a) in situ Hi-C^51^ (MboI), (b) in situ Micro-C, (c) and Classic CAD-C. Stacked A/C/G/T fractions are shown for positions +1 to +50 bp into Fragment 1 (1-based from the ligation junction), using the same sampling logic used to enrich for local end-sequence bias in Fig. 1(h), here adapted from inferred cut sites to ligation junctions. (d) Ligation-junction coverage at an ATAC-accessible, occupied CTCF site in hTERT–RPE1 cells. Tracks show in situ Hi-C^51^ (green; MboI), in situ Micro-C (red), and Classic CAD-C (blue) ligation-junction coverage (scaled as reads per genomic content, RPGC), displayed on separate y-axis scales with different maxima (Hi-C: 275; Micro-C: 41; CAD-C: 29). Below, hTERT–RPE1 ATAC-seq IDR narrow peaks are shown, followed by top 5% occupied CTCF motifs. (e) CAD-C ligation-junction pileup over bound CTCF sites (GM12878 merged data). Mean CAD-C ligation-junction coverage (scaled as RPGC) is aggregated in a ±2.5 kb window centered on bound CTCF motif midpoints, oriented such that the loop interior (“Inside Loop”) is to the right. The raw mean (light) is Savitzky–Golay–smoothed (w=101, p=3; dark). (f) CAD-C ligation-junction pileup over active gene TSSs (GM12878 merged data). Mean CAD-C ligation-junction coverage (RPGC) is aggregated in a ±2.5 kb window centered on transcription start sites, oriented such that the gene body (“Inside Gene”) is to the right. The raw mean (light) is Savitzky–Golay–smoothed (w=101, p=3; dark). (g) CAD-C ligation-junction pileup over DNase-seq footprints (GM12878 merged data). Mean CAD-C ligation-junction coverage (RPGC) is aggregated in a ±2.5 kb window centered on ENCODE DNase-seq footprint midpoints. The raw mean (light) and Savitzky–Golay–smoothed trace (w=101, p=3; dark) are shown versus position relative to the midpoint (bp). All pileups in (e–g) were computed from the GM12878 merged LJO dataset (∼1.63 × 10^9^ pairs). (h) Genome-browser view of GM12878 merged data CAD-C ligation-junction coverage. The CAD-C ligation-junction density / coverage track (blue; RPGC, reads per genomic content) is shown alongside ENCODE DNase footprints (black) and top 5% occupied CTCF motifs (magenta), with RefSeq gene annotations below.

**Extended Data Figure 5.**
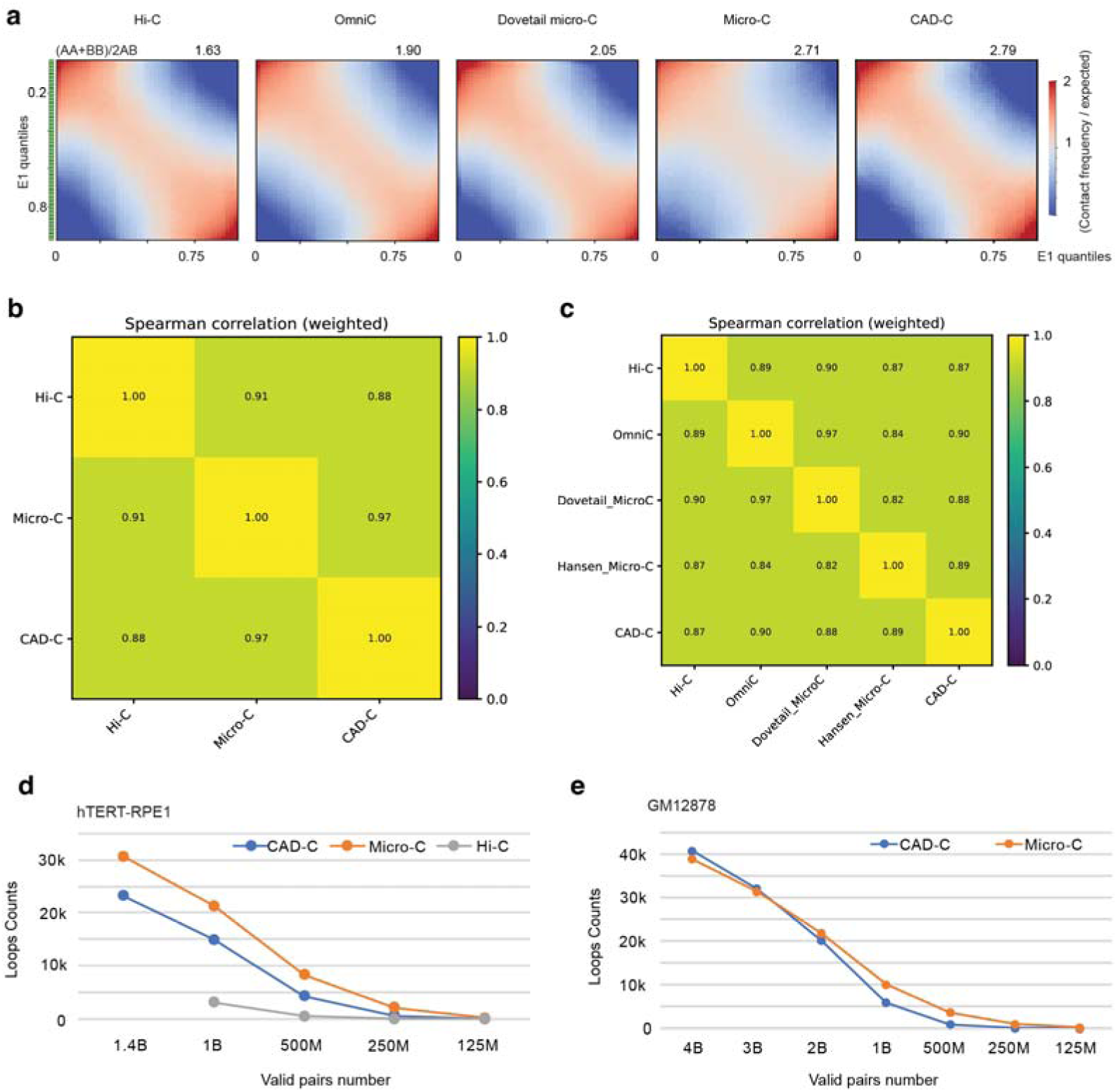
Comparative analysis of chromatin architecture across HiLC, MicroLC, OmniLC, and CADLC assays in GM12878 and hTERTLRPE1 cells. (a) Saddle plots and scores for GM12878 datasets: in-situ Hi-C^59^, OmniC (Dovetail), Micro-C (Dovetail), in situ Micro-C^57^ and CAD-C merged data. Compartment strength as follows, CAD-C (2.79), high-depth in situ Micro-C (2.71) and commercial Micro-C (2.05), Omni-C (1.90) and standard Hi-C (1.63). (b) Spearman correlation heatmap of insulation scores derived from hTERT-RPE1 in situ Hi□C, in situ Micro□C, and classic CAD□C datasets. The heatmap displays pairwise Spearman correlation coefficients calculated from genome□wide insulation scores across the three chromatin□conformation assays. (c) Spearman correlation heatmap of insulation scores derived from datasets in panel (a). The correlation matrix illustrates pairwise Spearman correlation coefficients computed from genome□wide insulation scores for all five chromatin□conformation datasets. (d) hTERT□RPE1 in situ Hi□C, Micro□C, and CAD□C subsampling analysis. Line plots depict the number of loops identified from subsampled subsets containing varying numbers of unique valid pairs. (e) GM12878 Micro□C and CAD□C subsampling analysis. Line plots show the number of loops identified from subsampled subsets with different numbers of unique valid pairs.

**Extended Data Figure 6.**
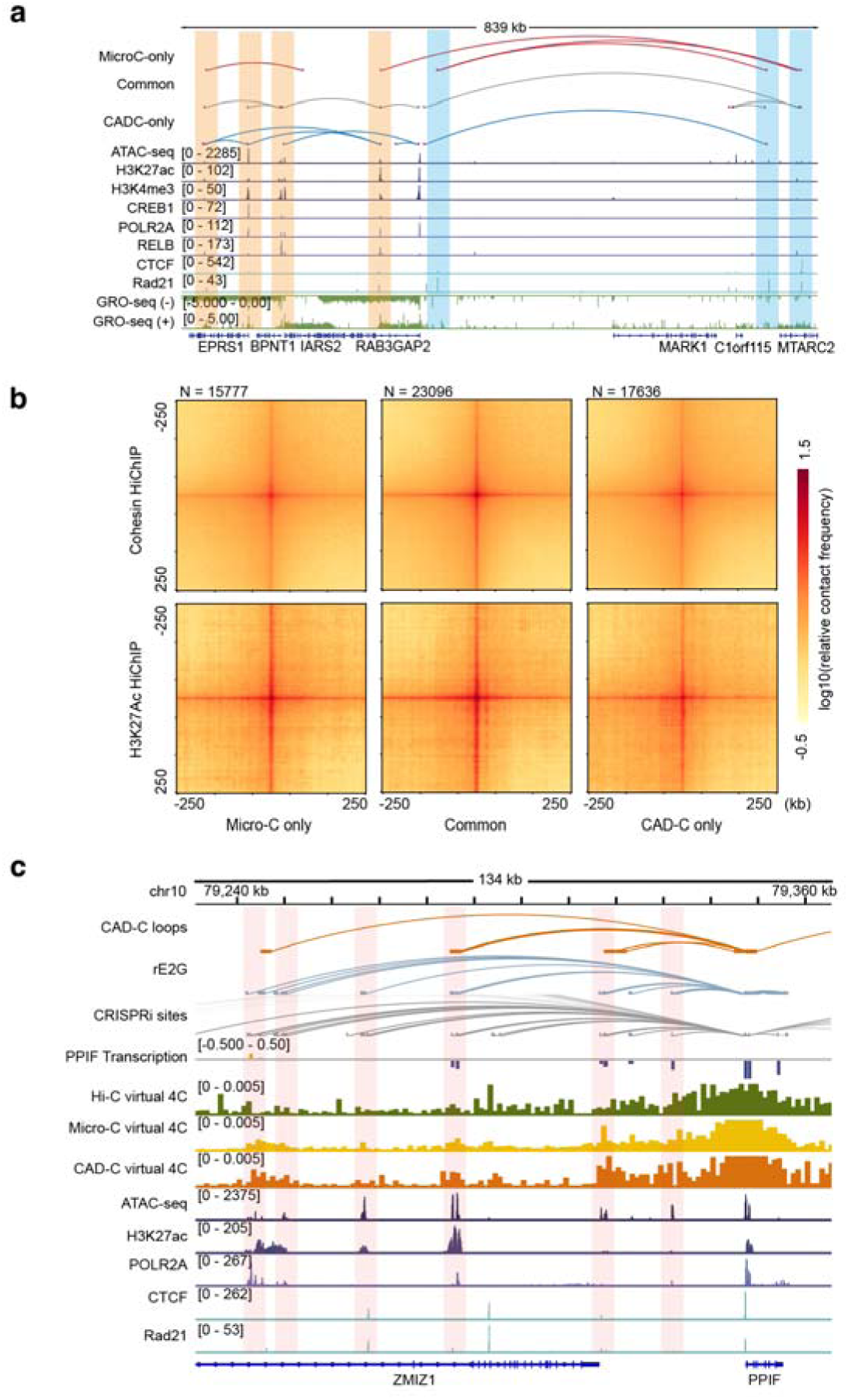
CADLC–specific chromatin loops reveal transcriptional activity beyond open chromatin. (a) Genomic tracks across the chr1:219.96–220.80 Mb region showing Micro□C and CAD□C loop calls together with ATAC□seq, H3K27ac, H3K4me3, CREB1, POLR2A, RELB, CTCF, Rad21^64^, and GRO□seq^65^ profiles. CAD□C–only loop anchors are highlighted with an orange background and display strong enrichment of active transcriptional signals, whereas Micro□C–only loop anchors exhibit prominent CTCF binding and robust cohesin (Rad21) occupancy. (b) Pileup heatmaps of the three loop groups (Micro-C only, common, CAD-C only) on CTCF and H3K27Ac HiChIP data from GM12878 cell line. All groups show typical aggregation patterns. CAD-C-only loops exhibit richer surrounding signals on H3K27Ac HiChIP data^66,67^. (c) CAD□C–detected chromatin loops surrounding the PPIF locus, shown together with virtual 4C tracks anchored at the PPIF promoter (viewpoint: chr10:79,347,550–79,347,750), ATAC□seq, H3K27ac, POLR2a, CTCF, Rad21, and CRISPRi tracks targeting candidate PPIF enhancers^62^. The pink shading marks all ATAC□seq peaks within the region, illustrating that the CAD□C-identified loops are not merely reflective of open chromatin.

**Extended Data Figure 7.**
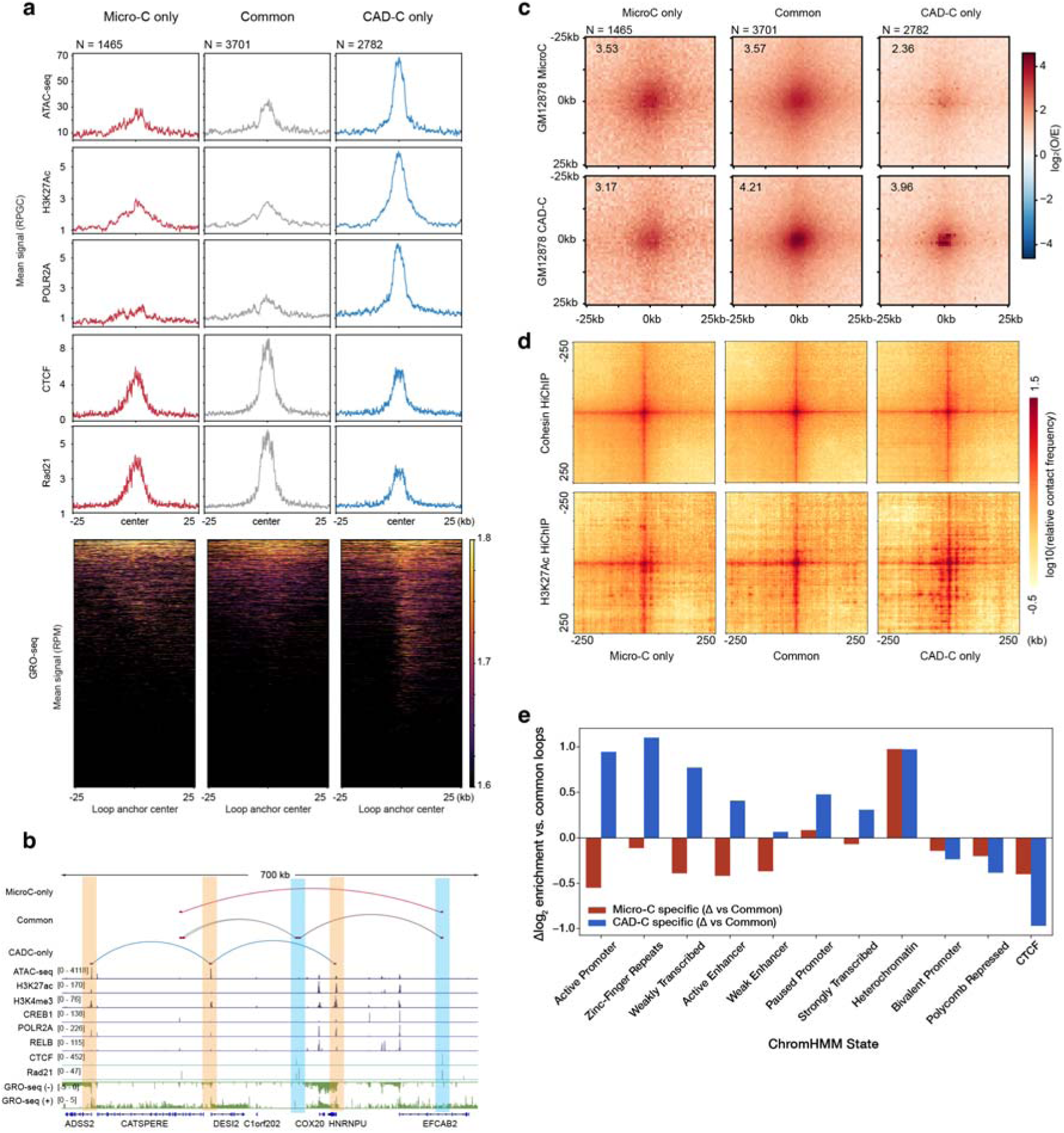
CAD-C preferentially captures active transcription loops compared to bead-based Micro-C. (a) Chromatin accessibility, H3K27ac, POLR2A, CTCF, cohesin subunit Rad21^64^, and nascent transcription (GRO-seq)^65^ pileup line plots at Micro-C only, common, CAD-C only loop anchors called at the resolution of 5kb with equal depth of CAD-C and bead-based Micro-C (Dovetail) in GM12878 cells. (b) Genomic tracks across the chr1:244.40–245.10 Mb region showing Micro□C and CAD□C loop calls together with ATAC□seq, H3K27ac, H3K4me3, CREB1, POLR2A, RELB, CTCF, Rad21, and GRO□seq profiles. CAD□C–only loop anchors are highlighted with an orange background. (c) Pileup analysis of the three loop groups in panel a, centered on loops (±25 kb). (d) Pileup heatmaps of the three loop groups from panel (a) on CTCF and H3K27Ac HiChIP data. All groups show typical aggregation patterns. CAD-C-only loops exhibit richer surrounding signals on H3K27Ac HiChIP data. (e) ChromHMM state enrichments at loop anchors for the 4 billion-contact matched GM12878 CAD-C merged data and GM12878 bead-based Micro-C (Dovetail) loop sets, corresponding differences (Δ log_2_) in enrichment for CAD-C–specific and Micro-C–specific anchors relative to the Common set.

**Extended Data Figure 8.**
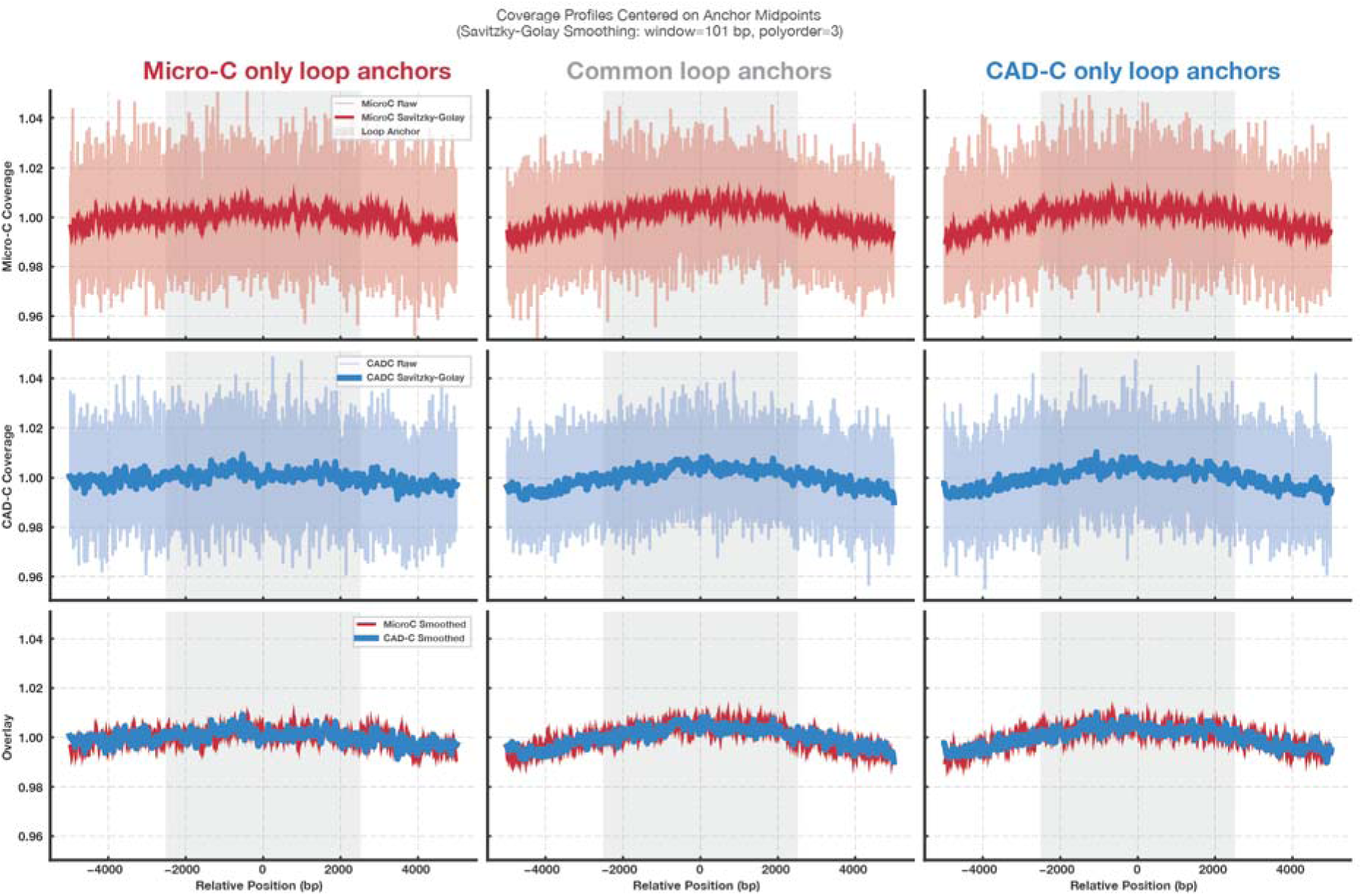
CAD-C and Micro-C ligation junction coverage over loop anchors is similar. Average ligation-junction coverage profiles centred on loop-anchor midpoints in GM12878, using the same depth-matched (with regards to cis >1 kb ligation contacts at .pairs.gz file level) loop sets at 2.5 kb defined for the Dovetail 800 M Micro-C versus CAD-C merged data comparison (Micro-C-only, Common and CAD-C-only loops; columns). Top row: Micro-C junction coverage (raw in light red, Savitzky–Golay–smoothed in dark red). Middle row: CAD-C junction coverage (raw in light blue, smoothed in dark blue). Bottom row: overlay of the smoothed Micro-C (red) and CAD-C (blue) profiles.

**Extended Data Figure 9.**
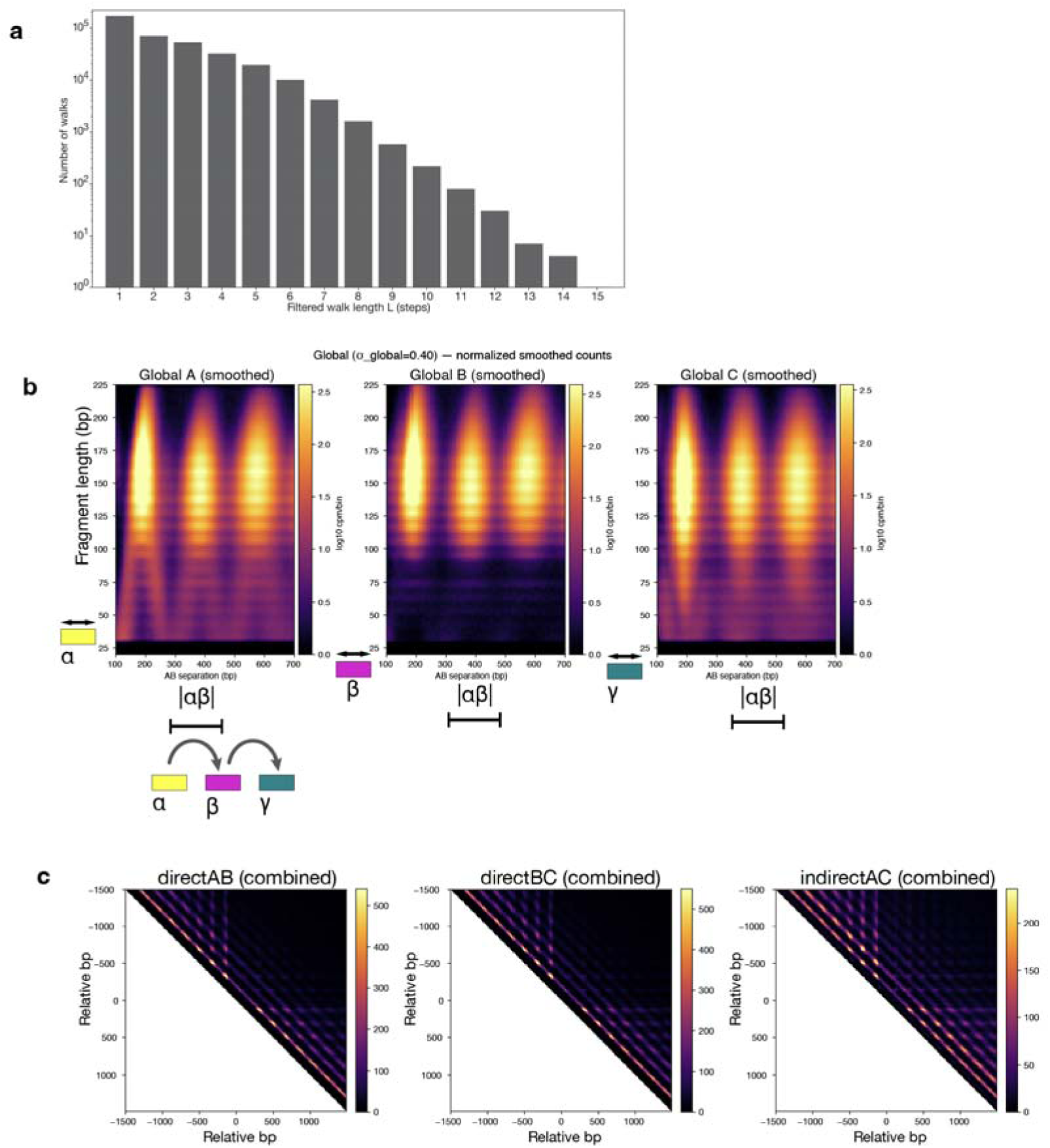
CADwalks consist of concatenated nucleosomal and sub-nucleosomal DNA fragments that report on local chromatin structure. (a) Number of walk steps (L) versus number of walks, plotted from long-read CADwalks filtered for walks with no fragment alignment. (b) Smoothed AB-distance × fragment-length maps for α, β, and γ fragments from cis-chromosomal αβγ CADwalks in GM12878, plotted as log10 counts per million per bin (cpm/bin) over AB midpoint separations of 100–700 bp and fragment lengths of 25–225 bp (global smoothing (J global = 0.40). (c) Midpoint-based CTCF contact matrices for three-fragment (αβγ) CADwalks. For each occupied CTCF motif in GM12878 (top 5% JASPAR motifs ranked by overlap with GM12878 CTCF ChIP–seq; seeMethods), bp-resolution local contact matrices (±1.5 kb around the motif centre) were constructed using fragment midpoints, strand-flipped and summed to obtain strand-agnostic signal. Panels show α–β (direct AB), β–γ (direct BC), and α–γ (indirect AC) contacts; upper triangles are plotted with axes in relative bp and colour indicating aggregated contact counts.

**Extended Data Figure 10.**
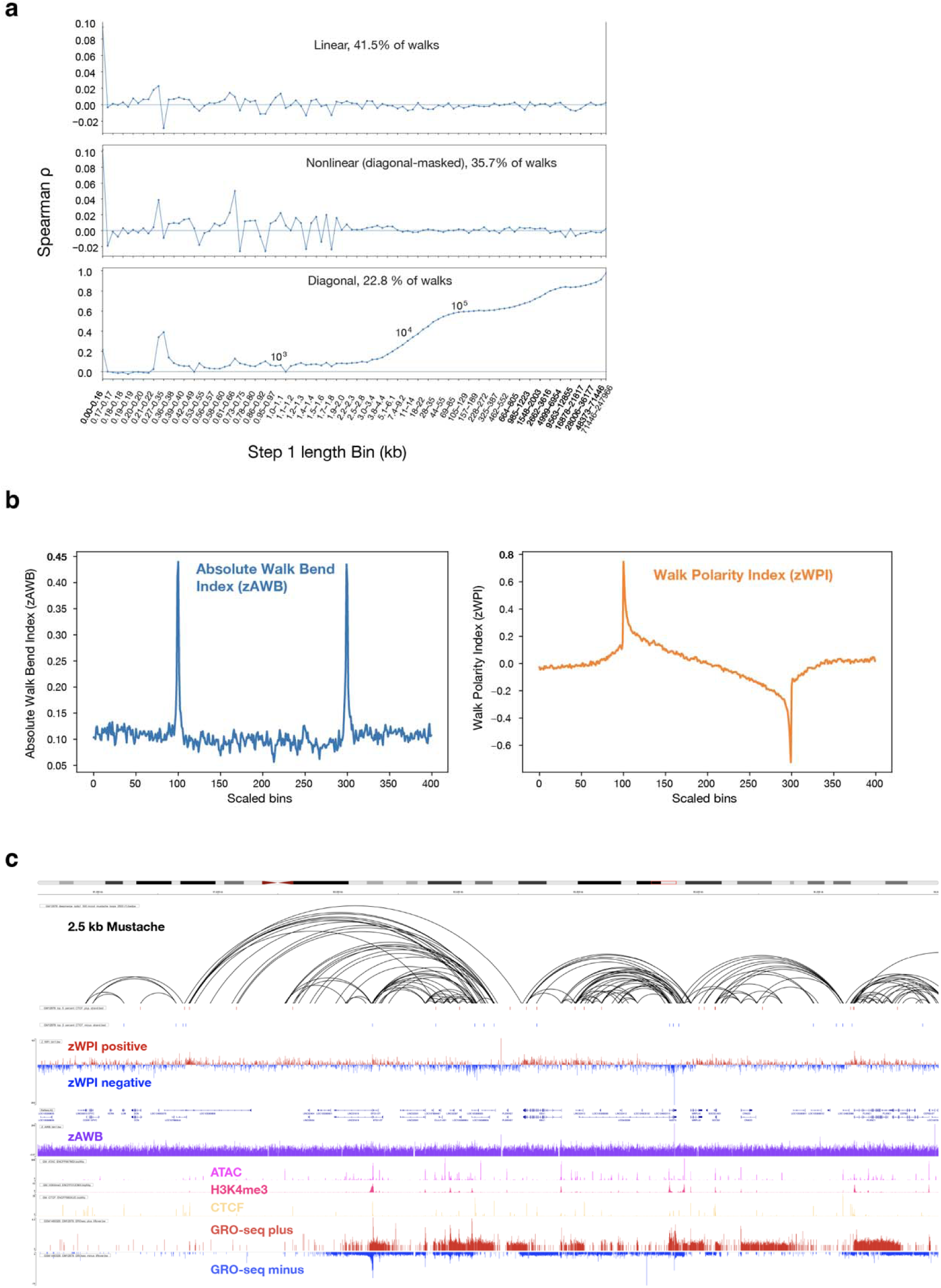
Step-size correlations and CADwalk-derived metrics across convergent CTCF loops. (a) Spearman correlation between consecutive step sizes |αβ| and |βγ| as a function of step-1(αβ) length bin, shown separately for linear, nonlinear, and diagonal walk classes (defined as in Methods). (b) Metaplots of zAWB (z-score of the Absolute Walk Bend index) (left) and zWPI (z-score of the Walk Polarity index) (right) across convergent CTCF loop domains in GM12878. Loop coordinates are rescaled so that scaled bins 100 to 300 span the loop interior (0-100%); flanking bins (0-100, 300-400) extend to −50% and 150% of loop length, respectively. (c) Example locus showing Mustache loop calls (2.5 kb), occupied CTCF motifs, zWPI track (positive/red; negative/blue), zAWB track, gene annotations, and chromatin marks (ATAC, H3K4me3, CTCF ChIP, GRO-seq).

## Notes

### Summary of Updates

The CAD/ICAD purification methods section has been updated to clarify and correct buffer recipes. Custom analysis code and additional detailed methods sections describing analyses have been added as supplementary material.

