## Supplementary Note on Ligation Gating Model for "CAD-C: An engineered nuclease enables repair-free *in situ* proximity ligation and nucleosome-resolution chromosome walks in human cells"

### 1 Ligation-junction coverage at loop anchors is similar in CAD-C and Micro-C

Loop-anchor ligation-junction coverage profiles are provided in the accompanying manuscript as Supplementary Fig. 11.

The genome-wide loop comparisons in this study revealed two separable differences between CAD-C and Micro-C at matched depth. First, CAD-C–specific loops were on average longer than Micro-C–specific loops, with the Common set intermediate; this right-shift arose from the non-TSS component rather than from promoter-engaged loops. Second, ChromHMM analysis showed that, relative to the Common set, CAD-C–specific anchors were biased towards Active Promoter/Enhancer states, whereas Micro-C–specific anchors were relatively enriched for Polycomb-repressed and heterochromatic states.

A natural explanation for these patterns is that CAD digestion, combined with native ligation competence, generated more ligation junctions in accessible regions. In Hi-C and related assays, locus "visibility" is known to depend on fragment length, GC content, sequence uniqueness and local chromatin accessibility, and several modeling frameworks explicitly correct for these systematic biases (Yaffe and Tanay 2011; Lajoie et al. 2015). In hTERT–RPE1, we previously showed that CAD-C ligation junctions were enriched at accessible chromatin and regulatory features, raising the possibility that elevated junction density at regulatory elements could increase the sensitivity of loop calling around those sites.

However, loop calling in this study was performed on ICE-balanced contact matrices, in which precisely this kind of one-dimensional visibility heterogeneity should be strongly attenuated. Iterative correction rescales rows and columns of the contact matrix until all bins have, by construction, equal total visibility (Imakaev et al. 2012), and bins that fail to converge are removed from downstream analyses. Independent work has shown that ICE and related methods largely flatten many visibility biases, even when they originate from differential restriction-enzyme accessibility, although some residual visibility signal can still correlate with chromatin state (Chandradoss et al. 2020). Under this scheme, a simple increase in ligation-junction density at accessible loci is unlikely, by itself, to generate CAD-C–specific loops once balancing has converged.

To test directly whether residual junction-density differences could nevertheless contribute to CAD-C–specific loops, we compared ligation-junction coverage around loop anchors in GM12878 CAD-C and Micro-C (accompanying manuscript, Supplementary Fig. 11). Using the same depth-matched GM12878 CAD-C DeepMerge and GM12878 Dovetail/Cantata Bio 800 M Micro-C datasets as in the APA analysis, we computed strand-agnostic ligation-junction coverage tracks from the `.pairs.gz` files for each assay and averaged coverage in ±4 kb windows around loop-anchor midpoints for three loop classes: Micro-C–only anchors, anchors of loops called in both assays (Common) and CAD-C–only anchors (Methods). Junction coverage was computed from ligation-junction–observed pairs and converted into RPGC-normalized single-base BigWig tracks before aggregation. Raw and Savitzky–Golay–smoothed mean profiles (window = 101 bp, polynomial order 3) were then overlaid for comparison.

Across all three anchor classes, both assays showed only modest and broadly similar enrichment of junction coverage in the immediate vicinity of the anchors (accompanying manuscript, supplementary Fig. 11). In the Micro-C tracks (top row), the smoothed profiles rose slightly above the flanking baseline within the central 1–2 kb and then returned towards baseline at larger distances. The CAD-C tracks (middle row) had essentially the same shape over the same y-axis range (0.96–1.04), with coverage varying within a narrow band around unity. When the smoothed Micro-C and CAD-C profiles were overlaid (bottom row), the traces almost completely coincided for Micro-C–only and Common anchors, and differed only subtly at CAD-C–only anchors, where CAD-C showed a slightly higher central bump. In no loop class was there a large assay-specific peak or trough at the anchor midpoint.

This similarity in loop-anchor coverage is notable in light of the biochemical differences between MNase and CAD. MNase is an endo-exonuclease that preferentially cuts A/T-rich linker DNA and can over-digest or resect nucleosomal DNA (Rushizky et al. 1960; Hörz and Altenburger 1981; Mieczkowski et al. 2016; Chereji et al. 2019), whereas CAD behaves as a double-strand-specific "one-cut" endonuclease with minimal exonucleolytic resection and generates ligation-ready 5′-phosphate/3′-hydroxyl ends (Widlak et al. 2000; Scholz et al. 2003; Zhelkovsky and McReynolds 2014). Given these digestion differences and the RPE1 analyses showing that CAD-C junctions were enriched at accessible chromatin, one might have expected a pronounced excess of junctions at CAD-C loop anchors. Instead, under the depth-matched and ICE-balanced conditions used here, the Micro-C and CAD-C junction tracks around anchors were more similar than the exonuclease/endonuclease contrast alone would suggest, and this similarity persisted even at CAD-C–only anchors that were nevertheless enriched for active ChromHMM states.

Two observations therefore need to be reconciled. First, the underlying nuclease biochemistry and footprinting behavior differ strongly between MNase and CAD, yet the one-dimensional ligation-junction coverage profiles at loop anchors in GM12878 were nearly indistinguishable between Micro-C and CAD-C (accompanying manuscript, Supplementary Fig. 11). Second, despite this similarity in anchor visibility and the use of the same ICE normalization and loop-calling framework, this study still reported more CAD-C loops whose anchors lay in TSS and active-chromatin states and captured regulatory examples such as the rs1250566–*PPIF* loop. Taken together, these observations suggest that the extra CAD-C loops were not well explained solely by "more junctions at anchors".

These results motivate a shift in perspective. The ligation-junction profiles analysed above are bulk, one-dimensional averages across molecules and cells: they report how often each genomic position participates in a junction, but they do not indicate which junctions occurred together in the *same* crosslinked complex or cell. By contrast, every count in a loop pixel ultimately arises from individual ligation events in which *both* participating loci were (i) present in the same crosslinked complex and (ii) simultaneously cut and ligation-competent in that cell at the moment of ligation. From this viewpoint, loop strength reflects a *joint* probability over both anchors rather than being determined solely by the marginal visibility of each anchor considered in isolation.

In this joint-probability view, CAD's chromatin digestion behavior—a double-strand-specific, non-processive "one-cut" nuclease that preserves ligation-ready ends at fixed offsets from crosslinked proteins and exhibits CHOMP-limited fragmentation—would be expected to increase the chance that both sides of a regulatory contact yielded intact, ligatable ends in the same cell before diffusion or overdigestion, especially at fragile, nuclease-sensitive regulatory nucleosomes. In contrast, MNase's endo-exonuclease activity and requirement for enzymatic end repair mean that, even if an anchor was accessible on average, a substantial fraction of underlying cells may have contributed damaged or resected ends that were less likely to participate in productive ligation at both sides of a given regulatory contact.

In the remainder of this note, we therefore adopt a *two-anchor ligation* view of CAD-C–specific loops: rather than being driven primarily by higher one-dimensional junction density at individual anchors, the excess of TSS- and active-chromatin-anchored loops in CAD-C is more consistent with an increased probability that *both* anchors of a regulatory contact were simultaneously ligatable in the same cell under CAD digestion. The minimal ligation-gating model introduced below formalizes this idea and provides a framework for relating end chemistry, CHOMP-limited fragmentation behavior and ligation efficiency to the observed contact distributions in CAD-C and related assays.

Taken together, these observations suggest that the key difference between CAD-C and Micro-C is not simply how often each anchor participates in ligation on average, but how often *both* anchors of a potential contact are ligatable in the *same* crosslinked complex. The one-dimensional ligation-junction tracks are bulk, marginal averages: they report how frequently each position ends up at a junction across many molecules and cells. In contrast, every count in a loop pixel reflects a stricter event, in which two loci are not only spatially proximate but also simultaneously carry ligation-competent ends at the moment of ligation. Two anchors can therefore look similarly "visible" in one dimension while still differing in how often they jointly pass this two-sided ligation gate. In the language introduced at the beginning of this study, this joint success probability is controlled by the *native ligation competence* (NLC) of each assay: the combination of raw cleavage density, chemical compatibility of the ends (5′-phosphate/3′-hydroxyl termini), and steric positioning of cuts relative to chromatin-bound proteins under CHOMP conditions. CAD's CHOMP-limited, native-end cleavage should raise the probability that *both* partners in a potential contact carry ligatable ends in the same crosslinked "cage", particularly at nuclease-sensitive, accessible chromatin, whereas MNase's exonucleolytic "chewing" is expected to lower it.

The ligation-gating model below is a minimal way to formalize this joint-probability view. It treats contact detection as the outcome of a polymer collision kernel filtered by a stochastic gate that depends on NLC and on the geometry of the crosslinked chromatin cage. Within this framework, changes in end chemistry and nuclease behavior enter through an effective anchor density $\rho_{\text{eff}}$, and we ask how realistic shifts in $\rho_{\text{eff}}$ might translate into the contact-scaling and loop differences observed between CAD-C and Micro-C.

### 2 A minimal ligation-gating model explains contact scaling and loop recovery

The analyses in the preceding sections highlight an apparent discrepancy. On one hand, CAD-C and Micro-C exhibit remarkably similar one-dimensional ligation-junction coverage at loop anchors (accompanying manuscript, Supplementary Fig. 11), suggesting that locus accessibility—or "visibility"—is broadly comparable between the two assays. On the other hand, the resulting two-dimensional contact maps differ in three important ways: CAD-C exhibits a massive enrichment of sub-kilobase contacts, a leftward shift in the $P(s)$ decay curve, and the specific recovery of active regulatory loops that appear faint or absent in Micro-C. Standard normalization procedures such as ICE balancing implicitly assume that once one-dimensional visibility biases are corrected, matrix entries are proportional to the underlying contact probabilities, and that the *efficiency* of capturing a contact does not itself depend on genomic distance beyond the polymeric decay.

Experimental and simulation work have begun to challenge this assumption. Proximity-ligation assays are probabilistic at multiple levels, reflecting both cell-to-cell heterogeneity in 3D conformations and stochasticity in crosslinking, digestion and ligation; consequently, detected contacts represent a population-average filtered by an assay- and context-dependent effective capture radius rather than a direct readout of a single underlying structure (O'Sullivan et al. 2013; Yang and Hansen 2024). Consistent with this view, successive 3C protocol refinements that increase capture efficiency (e.g. Hi-C 3.0 and Micro-C) shift recovered distance-decay toward shorter separations by improving recovery near the diagonal (Akgol Oksuz et al. 2021). Absolute quantification of 3C ligation products at the $\beta$-globin locus by Gavrilov and colleagues found that specific cross-ligation events typically occur at frequencies on the order of $\sim 1\%$, with self-ligation of the anchor fragment substantially more common (Gavrilov et al. 2013). More recently, Jusuf *et al.* calibrated Micro-C contact frequencies against live-cell imaging and reported genome-wide looping probabilities in the low single-digit percent range (Jusuf et al. 2025). Virtual Hi-C simulations that explicitly model crosslinking, digestion and ligation steps likewise show that digestion and ligation parameters can reshape $P(s)$ and contact statistics even for a fixed underlying polymer ensemble (Herrera et al. 2025). Together, these studies indicate that proximity-ligation assays operate in a regime where only a small fraction of potential ligation events succeed, and where that success probability depends on fragment length, end chemistry and the surrounding chromatin environment.

This framing resolves an apparent tension in benchmarking: increasing $\rho_{\text{eff}}$ is expected to elevate the near-diagonal background because short-range encounters dominate the collision pool, while simultaneously improving sensitivity to rare long-range regulatory interactions by increasing the probability that such encounters are captured as ligation products. Polymer models of looped chromosomes and finite detection kernels predict that loops can reshape distance-dependent contact scaling and introduce diagnostic features in $P(s)$ (K. E. Polovnikov et al. 2023; K. Polovnikov and Starkov 2025). Complementary live-cell imaging further indicates that distal enhancer–promoter communication is constrained by strongly subdiffusive chromatin motion and that cohesin-dependent dynamics can generate rare, longer-lived encounters that plausibly dominate productive regulatory interactions (Lee et al. 2025; Mazzocca et al. 2025; Ubertini et al. 2025). In this context, CAD-C's enrichment for sub-kilobase contacts and its increased recovery of distal regulatory loops can be understood as coupled consequences of operating in a higher-$\rho_{\text{eff}}$ ligation regime rather than evidence that the underlying polymer ensemble must differ between assays.

To formalize this idea in a compact way, we developed a minimal probabilistic framework, the *ligation-gating model*, which treats contact detection as a Poisson gating process superimposed on a simple polymer collision kernel. The model has a single tunable parameter, the *effective anchor density* $\rho_{\text{eff}}$ (anchors per base pair), which captures how efficiently a nuclease generates and preserves ligation-competent ends without eroding the underlying DNA substrate and how often those ends can find each other within a crosslinked cage. Within this framework, background contacts and structural loops differ only in their statistical requirements: generic background interactions require at least two ligatable anchors in the genomic interval, whereas loops, conditional on a fixed viewpoint, require only one successful ligation at the distal anchor.

#### 2.1 Model definitions and derivation

We model the observed contact probability $P_{\text{obs}}(s)$ as arising from two components: a polymer collision term that depends only on genomic separation $s$, and a *gating* term that encodes the probability of generating the necessary ligation-competent ends at the relevant loci. For the polymer part, we assume that potential contacts follow a power-law distance distribution

$$\gamma(s) \propto s^{-1} \tag{1}$$

between $s_{\min}$ and $s_{\max}$, with cumulative distribution $\Gamma(s)$ that is uniform in $\log s$ over this range. This log-uniform kernel is not intended as a precise fit to $P(s)$, but as a convenient stand-in for the fact that there are many more distinct locus pairs separated by large genomic distances than by short ones.

The effective anchor density $\rho_{\text{eff}}$ subsumes the microscopic factors introduced at the beginning of this study. For clarity, we write it schematically as

$$\rho_{\text{eff}} \;=\; \rho_0 \, \underbrace{\epsilon_{\text{chem}}\,\theta_{\text{steric}}}_{\text{NLC}} \, \phi_{\text{geom}}, \tag{2}$$

where $\rho_0$ is the microscopic cleavage density (cuts per base pair), $\epsilon_{\text{chem}}$ is the fraction of cuts that yield chemically compatible $5'$-phosphate/$3'$-hydroxyl termini, $\theta_{\text{steric}}$ is the fraction of cuts that occur at sterically favorable positions relative to chromatin-bound proteins (for example, at linker-proximal sites without overdigestion), and $\phi_{\text{geom}}$ is a dimensionless factor capturing the flexibility and orientation constraints of the crosslinked, fragmented chromatin cage. We refer to the product $\epsilon_{\text{chem}}\theta_{\text{steric}}$ as the *native ligation competence* (NLC) of a digestion protocol: it reflects, per cut, how often the resulting ends are both chemically and sterically ligatable. The geometry factor $\phi_{\text{geom}}$ then accounts for the additional probability that two such anchors can approach with favorable orientation within a crosslinked complex.

Under this formulation, CAD is naturally interpreted as increasing both $\epsilon_{\text{chem}}$ and $\theta_{\text{steric}}$ relative to MNase, by generating ligation-ready blunt ends and cutting at fixed offsets from chromatin-bound proteins under CHOMP conditions. MNase-based Micro-C, in contrast, starts from termini that are not directly ligatable and exhibits exonucleolytic resection. Overdigestion and resection can delete, shorten, or displace ends from the nucleosome–linker interface. These effects reduce both $\epsilon_{\text{chem}}$ (because MNase-produced ends require repair to be rendered ligatable by T4 DNA ligase) and $\theta_{\text{steric}}$ (because anchors are no longer positioned at fixed offsets from chromatin-bound proteins), so that the realized $\rho_{\text{eff}}$ for Micro-C is likely substantially below its microscopic maximum, even in accessible chromatin.

**Background gating ($\geq 2$ anchors).** For a random background interval of genomic length $s$ to be detected as a contact, both ends of the interval must be defined by ligatable anchors. Approximating anchors as a Poisson process with rate $\rho_{\text{eff}}$, the probability that an interval of length $s$ contains at least two anchors is

$$G_{\text{base}}(s;\rho_{\text{eff}}) = 1 - e^{-\rho_{\text{eff}} s}(1 + \rho_{\text{eff}} s). \tag{3}$$

This double-anchor gating function starts near zero at small $s$ or low $\rho_{\text{eff}}$ and rises sigmoidally with distance, approaching unity only once the interval is large enough that two anchors are likely. In the model, the gated background cumulative distribution is

$$D_{\text{base}}(s) = \Gamma(s)\, G_{\text{base}}(s;\rho_{\text{eff}}), \tag{4}$$

and the corresponding background contribution to the observed distance distribution is obtained by differentiation:

$$P_{\text{base}}(s) = \frac{\mathrm{d}D_{\text{base}}}{\mathrm{d}s}. \tag{5}$$

**Loop gating ($\geq 1$ anchor).** A structural loop, by contrast, is defined by a pre-existing bridge between two loci. When analyzing loops from the perspective of a fixed anchor (for example, a TSS-centred aggregate), the locus at one end of the loop is already specified. Under these conditions, detecting the loop requires only that at least one ligatable partner end be generated within a small genomic window around the distal anchor. If this window has span $s_{\text{window}}$, the corresponding single-anchor gate is

$$G_{\text{loop}}(\rho_{\text{eff}}) = 1 - e^{-\rho_{\text{eff}} s_{\text{window}}}. \tag{6}$$

Because Eq. 6 depends on $\rho_{\text{eff}}$ but not on the full separation $s$, and because it requires only a single event in a small window, it saturates more rapidly with increasing $\rho_{\text{eff}}$ than the background gate.

In the simulations, we represent a family of loops as a narrow band of genomic separations rather than as a single fixed distance. Concretely, we assume that the genomic distance between the two loop anchors is normally distributed with mean $s_{\text{loop}} \approx 200$ kb and standard deviation $\sigma \approx 1$ kb, and we denote this normalized Gaussian by $\ell(s)$. This construction does *not* set the contact or collision probability between the anchors; it simply acknowledges that experimentally called "200 kb" loops correspond to a small range of separations (due to binning, anchor uncertainty and local sliding of the contact point) rather than to a single base-pair distance. The actual probability that a loop contact is captured is controlled by the loop fraction $f_{\text{loop}}$ and the ligation gate $G_{\text{loop}}(\rho_{\text{eff}})$; the Gaussian $\ell(s)$ just specifies where, in separation space, the loop population is concentrated.

The loop contribution to the observed distance distribution is then

$$P_{\text{loop}}(s; \rho_{\text{eff}}) = f_{\text{loop}}\, \ell(s)\, G_{\text{loop}}(\rho_{\text{eff}}), \tag{7}$$

where $\ell(s)$ is the normalized Gaussian. By construction, we treat the loop state as already satisfying the spatial proximity requirement; conditional on being in the loop state, the polymer part contributes a fixed factor that is absorbed into $f_{\text{loop}}$, and the remaining stochastic requirement is the presence of a ligatable end in the distal window captured by $G_{\text{loop}}$.

The full observed distribution is a mixture of background and loop components,

$$P_{\text{obs}}(s; \rho_{\text{eff}}) = (1 - f_{\text{loop}}) P_{\text{base}}(s) + P_{\text{loop}}(s; \rho_{\text{eff}}), \tag{8}$$

and the observed/expected ratio at separation $s$ is defined as

$$\text{O/E}(s; \rho_{\text{eff}}) = \frac{P_{\text{obs}}(s; \rho_{\text{eff}})}{P_{\text{base}}(s; \rho_{\text{eff}})}. \tag{9}$$

### 2.2 Order-of-magnitude estimates for $\rho_{\text{eff}}$

Although $\rho_{\text{eff}}$ is treated as a free parameter in the simulations below, it is helpful to anchor its scale with simple back-of-the-envelope estimates. Here we focus on the *maximal* anchor densities implied by idealized fragmentations, before applying the NLC and geometry factors.

For an MboI-based Hi-C experiment in the human genome, the 4-bp GATC motif appears roughly once every $\sim 256$ bp in a random sequence, with a similar average spacing in *hg38*. Each cut generates two potential anchors, giving a theoretical microscopic cleavage density

$$\rho_0^{\text{Hi-C}} \approx \frac{2}{256} \approx 8 \times 10^{-3} \text{ anchors/bp.}$$

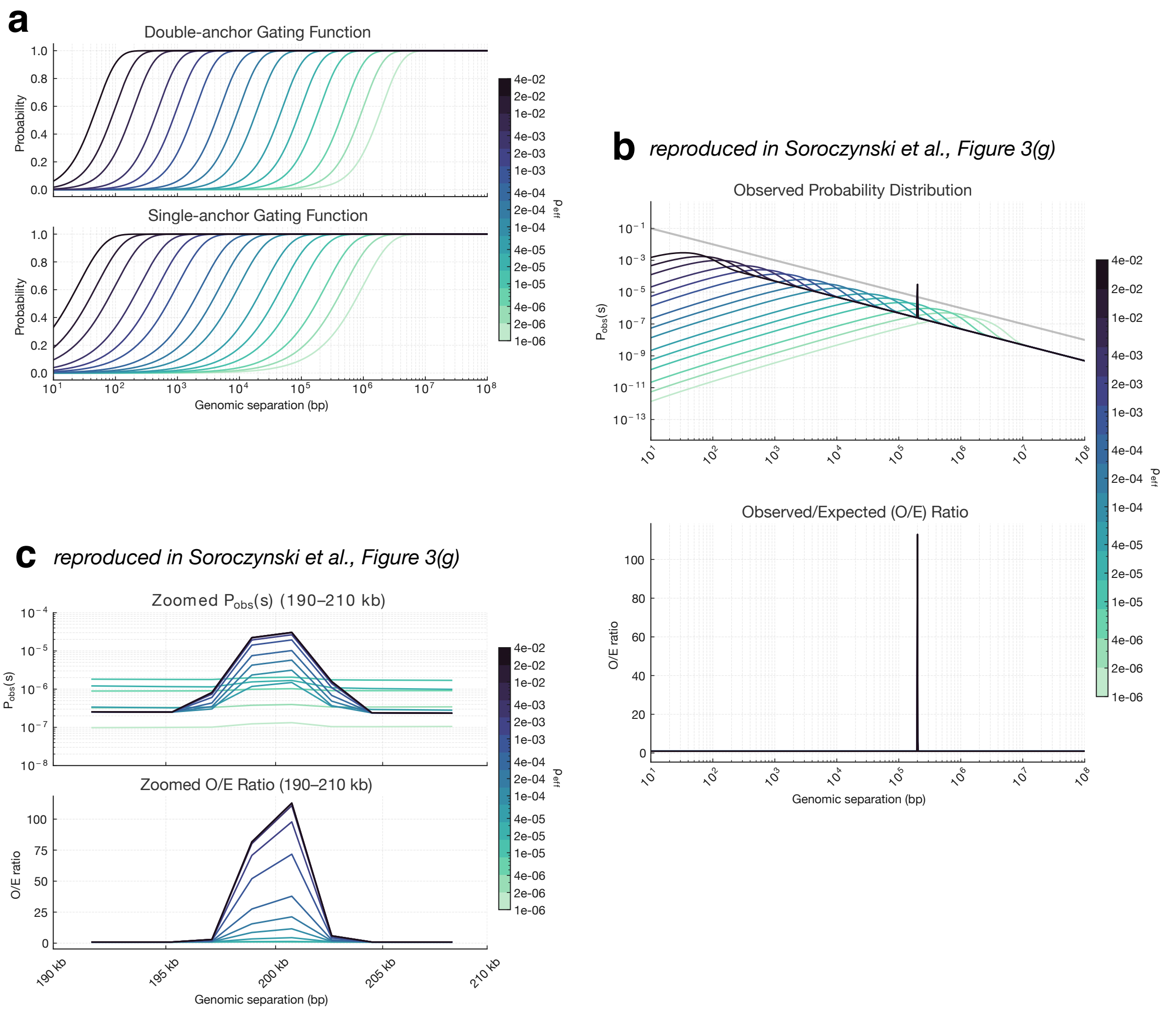


**Supplementary Note Fig. 1. A ligation-gating model links ligation efficiency to contact map features. (A)** Simulated gating functions for background contacts ($\geq 2$ anchors, top) and loop contacts ($\geq 1$ anchor, bottom) as a function of genomic distance $s$. Curves are colored by effective anchor density $\rho_{\text{eff}}$, increasing from low (light) to high (dark). Increasing $\rho_{\text{eff}}$ shifts the detection threshold for background contacts to shorter distances and causes the single-anchor loop gate to saturate more rapidly than the double-anchor background gate. **(B)** Simulated observed contact probability distributions $P_{\text{obs}}(s)$ (top) and corresponding observed/expected (O/E) ratios (bottom) for the same set of $\rho_{\text{eff}}$ values. As $\rho_{\text{eff}}$ increases, recovery of short-range contacts reweights probability mass towards small $s$ and steepens the decay of $P_{\text{obs}}(s)$, while the O/E ratio at the loop separation ($\sim 200$ kb) develops a pronounced peak. **(C)** Zoomed view of the loop region (190–210 kb) showing $P_{\text{obs}}(s)$ (top) and O/E (bottom). The differential saturation of the loop versus background gating functions produces a non-linear boost in contrast at the loop separation, illustrating how higher ligation efficiency can simultaneously increase short-range background and enhance loop prominence.

Gavrilov *et al.* measured absolute 3C ligation yields at a single locus and found that specific long-range cross-ligation products typically form at $\sim 1\%$ frequency, with self-ligation of the anchor fragment substantially more common (Gavrilov et al. 2013). This suggests that, once partial digestion, end repair, local structure and orientation constraints are taken into account, only a small fraction of potential ends contribute to successful proximity ligations. In the language of Eq. 2, this translates into an NLC$\times\phi_{\text{geom}}$ factor well below unity, such that the *effective* anchor density $\rho_{\text{eff}}$ relevant for Hi-C contact formation is much smaller than $\rho_0^{\text{Hi-C}}$.

For MNase-based Micro-C, we assume a uniform nucleosome repeat length of $\sim 190$ bp, decomposed into $\sim 150$ bp of nucleosome core DNA and $\sim 40$ bp of linker. In the idealized limit where digestion leaves only mononucleosome fragments and both of their ends are ligatable, each repeat contributes one 150 bp fragment with two ends. This corresponds to

$$\rho_0^{\text{Micro-C}} \approx \frac{2}{190} \approx 1.1 \times 10^{-2} \text{ anchors/bp},$$

slightly higher than the MboI case. In practice, MNase digestion involves both endonuclease and exonuclease activities; overdigestion and resection will often delete, shorten, or displace ends from the nucleosome–linker interface. These effects reduce both $\epsilon_{\text{chem}}$ (because MNase-produced ends require repair to be rendered ligatable by T4 DNA ligase) and $\theta_{\text{steric}}$ (because anchors are no longer positioned at fixed offsets from chromatin-bound proteins), so that the realized $\rho_{\text{eff}}$ for Micro-C is likely substantially below this theoretical maximum.

For CHOMP-limited CAD-C, we again assume an NRL of $\sim 190$ bp, but now take both the 150 bp nucleosome core and the 40 bp linker to survive as separate fragments with two ligation-competent ends each. In this idealized limit, each 190 bp repeat yields one "N" fragment and one "L" fragment, for a total of four ends:

$$\rho_0^{\text{CAD-C}} \approx \frac{4}{190} \approx 2.1 \times 10^{-2} \text{ anchors/bp}.$$

Because CAD directly generates blunt 5′-phosphate/3′-hydroxyl termini and CHOMP-limited digestion preserves fixed-offset cuts at nucleosome–linker interfaces, both $\epsilon_{\text{chem}}$ and $\theta_{\text{steric}}$ are expected to be closer to unity than in Micro-C. In Direct CAD-C experiments, we observed a $\sim 90\%$ depletion of the mononucleosome band after ligation, consistent with the majority of free mononucleosome fragments being consumed into ligation products rather than remaining as isolated monomers. While this does not by itself specify $\rho_{\text{eff}}$, it supports the view that CAD operates in a relatively high-NLC, high-anchoring regime.

Taken together, these simple estimates suggest the hierarchy

$$\rho_{\text{eff}}^{\text{Hi-C}} \lesssim \rho_{\text{eff}}^{\text{Micro-C}} \lesssim \rho_{\text{eff}}^{\text{CAD-C}},$$

with CAD-C enjoying both a higher microscopic anchor density $\rho_0$ and a higher NLC than Micro-C. In the simulations below, we therefore treat CAD-C-like conditions as corresponding to larger values of $\rho_{\text{eff}}$ than Micro-C-like conditions, without attempting to directly invert experimental data to obtain absolute $\rho_{\text{eff}}$ values.

### 2.3 Analytical results and interpretation

Sweeping the single parameter $\rho_{\text{eff}}$ across several orders of magnitude reproduces, in a qualitative sense, the three main features observed in CAD-C. First, increasing $\rho_{\text{eff}}$ shifts the onset of the double-anchor gate $G_{\text{base}}(s)$ towards shorter distances (Supplementary Note Fig. 1A, top). At low $\rho_{\text{eff}}$, very few background intervals at sub-kilobase separations contain two anchors, and $P_{\text{base}}(s)$ is relatively depleted near the diagonal. As $\rho_{\mathrm{eff}}$ rises, the gating function becomes non-zero for progressively shorter $s$, allowing the large pool of nearest-neighbor interactions ($< 1$ kb) to be converted into observable contacts. Under a fixed total number of cis contacts, this reweights probability mass towards short-range interactions and steepens the decay of $P_{\mathrm{obs}}(s)$ (Eq. 8), qualitatively matching the leftward shift seen in CAD-C relative to Micro-C. This behavior is consistent with more detailed polymer simulations in which digestion and ligation parameters are explicitly varied while the underlying polymer ensemble is held fixed (Herrera et al. 2025).

Second, the model explains the high-contrast recovery of loops. Because the single-anchor loop gate $G_{\mathrm{loop}}$ (Eq. 6) saturates more quickly than the double-anchor background gate at the loop separation, there is a regime of $\rho_{\mathrm{eff}}$ in which the loop detection requirement is essentially satisfied, while the surrounding background remains only partially gated. In this regime, the ratio $G_{\mathrm{loop}}/G_{\mathrm{base}}$ is maximized, and the loop peak in $P_{\mathrm{obs}}(s)$ becomes sharply defined against the rising background. The corresponding O/E curves (Eq. 9) display a pronounced spike at the loop position even though the local background under the peak has increased (Supplementary Note Fig. 1B,C). In other words, enhanced ligation efficiency can simultaneously raise short-range background and *increase* loop contrast. The choice of $f_{\mathrm{loop}}$ and $s_{\mathrm{loop}}$ in the simulations is deliberately simple: $s_{\mathrm{loop}} \approx 200$ kb sits within the typical range of CTCF/cohesin-mediated loops, and the loop fraction $f_{\mathrm{loop}}$ is chosen for visual clarity rather than to match the absolute loop probabilities of a few percent inferred from absolute quantification (Jusuf et al. 2025). We therefore interpret loop amplitudes in the model qualitatively, as a function of $\rho_{\mathrm{eff}}$, rather than quantitatively.

Third, this framework rationalizes why CAD-C preferentially recovers active loops. Active regulatory elements reside in nuclease-sensitive, accessible chromatin, where MNase-mediated resection can destroy or displace potential anchors, effectively lowering the local NLC and thus $\rho_{\mathrm{eff}}$ despite overall accessibility. By preserving fragment ends at fixed offsets from chromatin-bound proteins and generating ligation-ready termini without extensive exonucleolytic “chewing”, CAD maintains a high $\rho_{\mathrm{eff}}$ in exactly these regions. In the ligation-gating model, loops are the features most sensitive to such changes: because their detection criteria are governed by the single-anchor gate, modest increases in $\rho_{\mathrm{eff}}$ yield superlinear gains in loop signal relative to the surrounding background.

This minimal model is not intended as a full mechanistic description of chromatin folding or enzymology, and we do not attempt to fit its parameters directly to the CAD-C and Micro-C datasets. Instead, it provides a compact way to articulate how end chemistry, nuclease processivity and cage geometry, summarized here by $\rho_{\mathrm{eff}}$, can affect two-anchor ligation events. More elaborate polymer-based approaches that infer loop-extrusion parameters or loop densities from contact maps reach compatible conclusions: distance-dependent contact decay reflects a combination of polymer statistics, looped topology and a finite detection kernel that depends on fragment length and ligation geometry (K. E. Polovnikov et al. 2023; K. Polovnikov and Starkov 2025; Subic et al. 2025). Complementary live-cell imaging supports the same central constraint: chromatin motion is strongly subdiffusive and search times grow steeply with spatial separation, while cohesin-dependent dynamics can accelerate local search and can generate rare, longer-lived encounters (Lee et al. 2025; Mazzocca et al. 2025; Ubertini et al. 2025). Within this broader context, CAD-C’s enrichment for short-range contacts and its selective recovery of active regulatory loops can emerge naturally from operating in a higher-NLC, higher-$\rho_{\mathrm{eff}}$ ligation regime than MNase-based Micro-C.

Polovnikov, Kirill and Dmitry Starkov (Sept. 6, 2025). “Universal contact statistics of looped polymers resolve cohesin density and stoichiometry in vivo”. In: *Biophysics* biorxiv;2025.09.04.674214v1. (Visited on 09/07/2025).

Polovnikov, Kirill E et al. (Oct. 13, 2023). “Crumpled polymer with loops recapitulates key features of chromosome organization”. en. In: *Phys. Rev. X.* 13 (4), p. 041029.

Rushizky, G W et al. (Mar. 1, 1960). “A map of the products resulting from the action of micrococcal nuclease on thymus deoxyribonucleic acid and its use as a guide to specificity”. en. In: *Biochem. Biophys. Res. Commun.* 2 (3), pp. 153–158. (Visited on 11/20/2025).

Scholz, Sebastian R et al. (Aug. 12, 2003). “Experimental evidence for a beta beta alpha-Me-finger nuclease motif to represent the active site of the caspase-activated DNase”. en. In: *Biochemistry* 42 (31), pp. 9288–9294. (Visited on 11/20/2025).

Subic, Tina et al. (Oct. 17, 2025). “Mechanistic Genome Folding at Scale through the Differentiable Loop Extrusion Model”. In: *Genomics* biorxiv;2025.10.17.682904v1.

Ubertini, Mattia et al. (Sept. 24, 2025). “Loop extrusion creates rare, long-lived encounters underlying enhancer-promoter communication”. In: *bioRxiv.* (Visited on 10/05/2025).

Widlak, Piotr et al. (Mar. 17, 2000). “Cleavage Preferences of the Apoptotic Endonuclease DFF40 (Caspase-activated DNase or Nuclease) on Naked DNA and Chromatin Substrates*”. In: *J. Biol. Chem.* 275 (11), pp. 8226–8232.

Yaffe, Eitan and Amos Tanay (Oct. 16, 2011). “Probabilistic modeling of Hi-C contact maps eliminates systematic biases to characterize global chromosomal architecture”. en. In: *Nat. Genet.* 43 (11), pp. 1059–1065. (Visited on 04/25/2025).

Yang, Jin H and Anders S Hansen (July 2024). “Enhancer selectivity in space and time: from enhancer-promoter interactions to promoter activation”. en. In: *Nat. Rev. Mol. Cell Biol.* 25 (7), pp. 574–591.

Zhelkovsky, Alexander M and Larry A McReynolds (Nov. 28, 2014). “Polynucleotide 3’-terminal phosphate modifications by RNA and DNA ligases”. en. In: *J. Biol. Chem.* 289 (48), pp. 33608–33616. (Visited on 11/20/2025).
