## Supplementary material for "CAD-C: An engineered nuclease enables repair-free *in situ* proximity ligation and nucleosome-resolution chromosome walks in human cells": Custom code: bam_insert_size_fld_v2_20250927.py

```
#!/usr/bin/env python3
# -*- coding: utf-8 -*-
"""
================================================================================
bam_insert_size_fld.py — Insert-size (FLD) analysis & plots for one or more BAMs
================================================================================

Author: Jan Soroczynski (Risca Lab, The Rockefeller University)
Date: 2025_09_28
Version: 1.4.0

Overview
--------
This command-line tool computes **insert-size / fragment-length distributions (FLDs)**
from one or more **sorted, indexed, paired-end BAM** files. It is designed to be
**production-grade** and **replot-friendly**, with efficient multicore iteration.
It creates a **timestamped output subdirectory** and ensures that **all files share
the same timestamp**. PDFs are always saved with **fonttype=42** (TrueType).

Defaults & key capabilities
---------------------------
• **Autosomes-only by default**: FLDs use **main autosomes** only, `^(chr)?[0-9]+$`,
  excluding X/Y and scrappy/decoy contigs. Override with:
  - `--include-sex-chroms` (adds X/Y)
  - `--include-all-contigs` (disable filtering)
  - `--contig-regex` (custom allowlist)

• **Optional mitochondrial export**: `--export-chrM-bam` writes a separate BAM with
  **all alignments on mitochondrial contigs** (default names `chrM,MT`; override with
  `--mt-contigs`). Index is created if nonempty.

• **Count eligible pairs** per BAM (primary, proper pairs; MAPQ; duplicates optional;
  secondary/supplementary optional).

• **Downsampling**: either **to shallowest** (`--downsample-to-min`) or to a **goal**
  (`--goal-reads N`). Sampling is applied on-the-fly (no temporary BAMs).

• **Raw arrays** of fragment lengths saved as `.npy` (store range default `0..750` bp)
  — can be skipped with `--no-raw-arrays`.

• **NEW (compact histograms)**: `--save-hist-tsv` writes **1-bp histograms** with
  **count** and **CPM** for each BAM (and for each state if `--bed-states` is used).
  Defaults:
  - Histogram range = **plot range** if specified, else the **store range**.
  - Files are **gzip-compressed** (`.tsv.gz`) unless you pass `--no-hist-compress`.

• **FLD plots** across inputs on both **linear CPM** and **log10 CPM** axes.
  - **Colors**: categorical **Inferno** darkest→lightest **in input order**.
  - **Font size** via `--font-size` (points); **line thickness** via `--line-width`.

• **Medians & modes** are computed in a user-specified stats range (defaults to store
  range), displayed in legends and exported in a **stats TSV**. Optional **units per
  BAM** (e.g., enzyme U) are recorded and used for labels.

• **CPM normalization** uses counts in a **normalization range** (default = store range)
  computed from the **raw arrays** (independent of the plotted x-range).

• **Optional BED state stratification** via `pybedtools`:
  – For each BAM, overlay **all states** with distinct qualitative colors.  
  – For each state, overlay **all BAMs** (Inferno categorical again).  
  – Save per‑(BAM,state) raw arrays (`.npy`, unless `--no-raw-arrays`) and **TSV histograms** if requested.

• **Structured logging** (`run__<TS>.log`) and **YAML metadata** including versions,
  contig lists, arguments, and produced outputs.

Fragment definition
-------------------
We iterate over **read1** only to avoid double counting. A pair is considered if:
- paired, **properly paired**, primary (unless `--include-secondary/--include-supplementary`),  
- both mates mapped to the same reference,  
- `MAPQ >= --min-mapq`, and (by default) **not a duplicate**.  
The fragment length is `abs(TLEN)`; pairs with non-positive TLEN are ignored.

Font & line style toggles
-------------------------
- `--font-size <float>` sets a global base font size (default: 10.0 pt).  
- `--line-width <float>` sets the thickness of plotted curves (default: 1.8).  
All PDFs maintain **fonttype=42**.

Inputs
------
- One or more **sorted, indexed BAMs** (`--bams`). `.bai` or `.csi` must exist.  
- Optional **BED files** (`--bed-states`) for state stratification (e.g., ChromHMM).  
  Requires `pybedtools` and `bedtools` in `PATH`.

Outputs (within `<outdir>/fld_<TIMESTAMP>`)
-------------------------------------------
- `counts_only__<TS>.tsv` — eligible-pair counts per BAM (always produced).  
- Per BAM raw arrays: `"<label>__fraglens_<=<store_max>bp__<TS>.npy"` (unless `--no-raw-arrays`).  
- **NEW (if `--save-hist-tsv`)** per-BAM histograms
  `"<label>__hist_<HMIN>-<HMAX>bp__<TS>.tsv[.gz]"` with columns:
  `len_bp`, `count`, `cpm`.  
- Aggregate FLD plots: `FLD_linear__<TS>.pdf`, `FLD_log10__<TS>.pdf` (CPM; **fonttype=42**).  
- **Stats**: `stats__<TS>.tsv` — medians, modes, denominators, sampling info, array paths, MT counts/paths.  
- If `--export-chrM-bam`: `"<label>__chrM_alignments__<TS>.bam"` (+ `.bam.bai` if nonempty).  
- If `--bed-states`:  
  • `"<label>__fragments_for_states__<TS>.bed"` (sampled fragments, if states used);  
  • Per-(BAM,state) arrays + **TSV histograms** if requested;  
  • Per-BAM and per-state overlay plots;  
  • `stats_states__<TS>.tsv`.  
- `metadata__<TS>.yaml` and `run__<TS>.log`.

Normalization, ranges, and statistics
-------------------------------------
- **Storage range**: `--store-min-frag .. --store-max-frag` (default `0..750`) controls
  which lengths are kept in memory/`.npy`.  
- **Plot range**: `--plot-min-frag .. --plot-max-frag` controls x-axis.  
- **Histogram range**: `--hist-min-frag .. --hist-max-frag` (default = plot range if set,
  else store range).  
- **Normalization range**: `--norm-min-frag .. --norm-max-frag` defines the CPM denominator
  computed from the **raw array**.  
- **Stats range**: `--stat-min-frag .. --stat-max-frag` controls median/mode.

Performance notes
-----------------
- Streaming iteration with `pysam.fetch(until_eof=True)`; downsampling is per-pair
  Bernoulli with seed (`--seed`).  
- Multiprocessing across BAMs and (BAM,state) tasks.

Nonstandard Python packages
---------------------------
`pysam`, `numpy`, `matplotlib`, `pandas`, `pybedtools` (optional), `PyYAML`

Configuration file (optional)
-----------------------------
`--config` accepts **YAML or JSON** with keys matching CLI arguments. CLI values
**override** config values.

Example usage
-------------
1) **Write compact histograms only (skip raw arrays)**:
    python bam_insert_size_fld.py \
        --bams A.bam B.bam C.bam \
        --save-hist-tsv --no-raw-arrays \
        --plot-min-frag 30 --plot-max-frag 250 \
        --outdir results_fld

2) **Standard run; auto-downsample to shallowest; 0–750 store; 60–500 plot; 90–250 stats;**
   **+ histogram TSVs (gzip), larger fonts and thicker lines**:
    python bam_insert_size_fld.py \
        --bams A.bam B.bam C.bam \
        --labels "4U,20U,100U" --units 4,20,100 \
        --downsample-to-min \
        --store-min-frag 0 --store-max-frag 750 \
        --plot-min-frag 60 --plot-max-frag 500 \
        --stat-min-frag 90 --stat-max-frag 250 \
        --save-hist-tsv \
        --font-size 12 --line-width 2.2 \
        --threads 8 \
        --outdir results_fld

3) **With ChromHMM states; export chrM BAMs; hist TSVs**:
    python bam_insert_size_fld.py \
        --bams A.bam B.bam \
        --bed-states Enh.bed,Tss.bed,Repr.bed \
        --state-labels Enhancer,TSS,Repressed \
        --export-chrM-bam \
        --save-hist-tsv \
        --outdir results_fld

CLI reference additions
-----------------------
**NEW:** `--save-hist-tsv`, `--hist-min-frag`, `--hist-max-frag`,
`--no-hist-compress`, `--no-raw-arrays`  
(Existing: see `-h`.)

Reproducibility & provenance
----------------------------
All outputs live in `fld_<TIMESTAMP>`; every file in that folder shares `<TS>`.
YAML `metadata__<TS>.yaml` and `run__<TS>.log` record args, versions, contig lists,
and outputs.

================================================================================
"""
from __future__ import annotations

import os
import sys
import re
import argparse
import textwrap
from datetime import datetime
import socket
import platform
import json
import random
import math
import multiprocessing as mp
from functools import partial
from typing import List, Tuple, Optional, Dict, Set

# Third-party
import numpy as np
import pandas as pd
import matplotlib
matplotlib.use("Agg")
import matplotlib.pyplot as plt
from matplotlib.ticker import MaxNLocator
# PDF compatibility (ALWAYS)
matplotlib.rcParams["pdf.fonttype"] = 42
matplotlib.rcParams["ps.fonttype"]  = 42

try:
    import yaml
    HAVE_YAML = True
except Exception:
    HAVE_YAML = False

try:
    import pysam
    pysam.set_verbosity(0)
except Exception as e:
    print(f"[FATAL] pysam is required: {e}", file=sys.stderr)
    sys.exit(2)

# pybedtools is optional unless --bed-states is used
try:
    import pybedtools  # noqa: F401
    HAVE_PYBEDTOOLS = True
except Exception:
    HAVE_PYBEDTOOLS = False

__VERSION__ = "1.4.0"


# -----------------------
# Utility/helper methods
# -----------------------

def timestamp() -> str:
    return datetime.now().strftime("%Y%m%d_%H%M%S")


def ensure_dir(path: str) -> None:
    os.makedirs(path, exist_ok=True)


def infer_index_path(bam: str) -> Optional[str]:
    bai = bam + ".bai"
    csi = bam + ".csi"
    if os.path.exists(bai):
        return bai
    if os.path.exists(csi):
        return csi
    base = os.path.basename(bam)
    dirname = os.path.dirname(bam)
    for idx in (base + ".bai", base + ".csi"):
        p = os.path.join(dirname, idx)
        if os.path.exists(p):
            return p
    return None


def check_bams(bams: List[str]) -> None:
    if not bams:
        raise ValueError("No BAMs provided.")
    for b in bams:
        if not os.path.exists(b):
            raise FileNotFoundError(f"BAM not found: {b}")
        idx = infer_index_path(b)
        if idx is None:
            raise FileNotFoundError(f"Index (.bai/.csi) not found for: {b}")


def setup_logging(outdir: str, ts: str):
    import logging
    log_path = os.path.join(outdir, f"run__{ts}.log")
    logger = logging.getLogger("fld")
    logger.setLevel(logging.INFO)
    fh = logging.FileHandler(log_path)
    fh.setLevel(logging.INFO)
    ch = logging.StreamHandler(sys.stdout)
    ch.setLevel(logging.INFO)
    fmt = logging.Formatter("%(asctime)s | %(levelname)s | %(message)s")
    fh.setFormatter(fmt)
    ch.setFormatter(fmt)
    if not logger.handlers:
        logger.addHandler(fh)
        logger.addHandler(ch)
    return logger


def get_inferno_colors(n: int) -> List[Tuple[float, float, float, float]]:
    """Return n Inferno colors from darkest to lightest in order."""
    cmap = plt.get_cmap("inferno")
    vals = np.linspace(0.05, 0.95, num=n)
    return [cmap(v) for v in vals]


def get_distinct_state_colors(n: int) -> List[Tuple[float, float, float, float]]:
    """Return n distinct qualitative colors for state overlays."""
    base = plt.get_cmap("tab20")
    vals = [base(i) for i in range(base.N)]
    if n <= len(vals):
        return vals[:n]
    hsv = plt.get_cmap("hsv")
    return [hsv(i / n) for i in range(n)]


def write_yaml(path: str, data: dict):
    if not HAVE_YAML:
        with open(path.replace(".yaml", ".json"), "w") as fh:
            json.dump(data, fh, indent=2)
        return
    with open(path, "w") as fh:
        yaml.safe_dump(data, fh, sort_keys=False)


def save_tsv(path: str, df: pd.DataFrame, compress: bool = False):
    if compress:
        df.to_csv(path if path.endswith(".gz") else path + ".gz",
                  sep="\t", index=False, compression="gzip")
    else:
        df.to_csv(path, sep="\t", index=False)


def integer_bin_edges(xmin: int, xmax: int) -> np.ndarray:
    """1-bp bins centered on integers [xmin, xmax]."""
    return np.arange(xmin - 0.5, xmax + 0.5 + 1, 1, dtype=float)


def as_uint16_array(values: List[int]) -> np.ndarray:
    if len(values) == 0:
        return np.array([], dtype=np.uint16)
    arr = np.asarray(values, dtype=np.uint32)
    arr[arr > np.iinfo(np.uint16).max] = np.iinfo(np.uint16).max
    return arr.astype(np.uint16, copy=False)


def safe_median(arr: np.ndarray) -> float:
    if arr.size == 0:
        return float("nan")
    return float(np.median(arr))


def compute_mode_from_hist(bin_centers: np.ndarray, counts: np.ndarray) -> float:
    if counts.sum() == 0:
        return float("nan")
    return float(bin_centers[np.argmax(counts)])


def fragment_coords_from_read(r) -> Optional[Tuple[str, int, int, int]]:
    """Return (chrom, start, end, length) from a read1 of a proper pair, or None."""
    if r.is_unmapped or r.mate_is_unmapped:
        return None
    if r.reference_id != r.next_reference_id:
        return None
    tlen = abs(r.template_length)
    if tlen <= 0:
        return None
    start = min(r.reference_start, r.next_reference_start)
    end = start + tlen
    return (r.reference_name, start, end, tlen)


def apply_plot_style(base_font_size: float, line_width: float) -> None:
    """Set matplotlib rcParams for font sizes and line width (pdf/ps fonttype kept at 42)."""
    fs = float(base_font_size)
    lw = float(line_width)
    rc = matplotlib.rcParams
    rc["font.size"] = fs
    rc["axes.titlesize"] = fs + 2
    rc["axes.labelsize"] = fs + 1
    rc["xtick.labelsize"] = fs
    rc["ytick.labelsize"] = fs
    rc["legend.fontsize"] = max(fs - 1, 6)
    rc["lines.linewidth"] = lw


# -----------------------
# Contig-selection logic
# -----------------------

def derive_contig_sets_for_bam(
    bam_path: str,
    contig_regex: str,
    include_sex_chroms: bool,
    include_all_contigs: bool,
    mt_names: List[str],
) -> Tuple[Set[str], Set[str], List[str]]:
    """
    Returns:
      allowed_autosomes (set), present_mt_contigs (set), all_reference_names (list)
    """
    pat = re.compile(contig_regex)
    with pysam.AlignmentFile(bam_path, "rb") as bam:
        refs = list(bam.references)

    if include_all_contigs:
        allowed = set(refs)
    else:
        allowed = {r for r in refs if pat.match(r)}
        if include_sex_chroms:
            # Add X/Y as present in header (with or without 'chr' prefix)
            for base in ("X", "Y"):
                for form in (base, f"chr{base}", base.lower(), f"chr{base.lower()}"):
                    if form in refs:
                        allowed.add(form)

    mt_lower = {m.strip().lower() for m in mt_names if m.strip()}
    present_mt = {r for r in refs if r.lower() in mt_lower}

    return allowed, present_mt, refs


# -----------------------------------
# Counting and fragment collection
# -----------------------------------

def count_eligible_pairs(
    bam_path: str,
    min_mapq: int,
    include_dups: bool,
    include_secondary: bool,
    include_supplementary: bool,
    allowed_refs: Optional[Set[str]],
) -> int:
    n = 0
    with pysam.AlignmentFile(bam_path, "rb") as bam:
        for r in bam.fetch(until_eof=True):
            if not r.is_read1:
                continue
            if not r.is_paired or not r.is_proper_pair:
                continue
            if r.is_unmapped or r.mate_is_unmapped:
                continue
            if allowed_refs is not None and r.reference_name not in allowed_refs:
                continue
            if not include_dups and r.is_duplicate:
                continue
            if not include_secondary and r.is_secondary:
                continue
            if not include_supplementary and r.is_supplementary:
                continue
            if r.mapping_quality < min_mapq:
                continue
            if abs(r.template_length) <= 0:
                continue
            n += 1
    return n


def export_chrM_alignments(
    bam_path: str,
    out_bam: str,
    mt_contigs: Set[str],
    min_mapq: int,
    include_dups: bool,
    include_secondary: bool,
    include_supplementary: bool,
) -> int:
    """
    Write all alignments mapping to mitochondrial contigs into out_bam.
    Returns number of alignments written. If >0, a .bai index is created.
    """
    if not mt_contigs:
        return 0
    written = 0
    with pysam.AlignmentFile(bam_path, "rb") as bam, \
         pysam.AlignmentFile(out_bam, "wb", template=bam) as w:
        for mt in mt_contigs:
            try:
                for r in bam.fetch(mt):
                    if not include_dups and r.is_duplicate:
                        continue
                    if not include_secondary and r.is_secondary:
                        continue
                    if not include_supplementary and r.is_supplementary:
                        continue
                    if r.is_unmapped:
                        continue
                    if r.mapping_quality < min_mapq:
                        continue
                    w.write(r)
                    written += 1
            except ValueError:
                continue
    if written > 0:
        try:
            pysam.index(out_bam)
        except Exception:
            pass
    return written


def collect_fragments_for_bam(
    bam_path: str,
    sample_goal: Optional[int],
    eligible_count: int,
    rng_seed: int,
    min_mapq: int,
    include_dups: bool,
    include_secondary: bool,
    include_supplementary: bool,
    store_min_frag: int,
    store_max_frag: int,
    write_fragment_bed: bool,
    fragment_bed_path: Optional[str],
    allowed_refs: Optional[Set[str]],
    export_chrM_bam: bool,
    mt_contigs: Set[str],
    chrM_bam_path: Optional[str],
) -> Dict:
    rng = random.Random(rng_seed)
    if sample_goal is None or sample_goal >= eligible_count:
        samp_frac = 1.0
    else:
        samp_frac = max(0.0, float(sample_goal) / float(eligible_count + 1e-12))

    kept_lengths: List[int] = []
    sampled = 0
    written_bed = 0

    if write_fragment_bed and not fragment_bed_path:
        raise ValueError("fragment_bed_path must be provided when write_fragment_bed=True")

    bed_fh = open(fragment_bed_path, "w") if write_fragment_bed else None

    with pysam.AlignmentFile(bam_path, "rb") as bam:
        for r in bam.fetch(until_eof=True):
            if not r.is_read1:
                continue
            if not r.is_paired or not r.is_proper_pair:
                continue
            if r.is_unmapped or r.mate_is_unmapped:
                continue
            if allowed_refs is not None and r.reference_name not in allowed_refs:
                continue
            if not include_dups and r.is_duplicate:
                continue
            if not include_secondary and r.is_secondary:
                continue
            if not include_supplementary and r.is_supplementary:
                continue
            if r.mapping_quality < min_mapq:
                continue

            if samp_frac < 1.0 and rng.random() >= samp_frac:
                continue

            coords = fragment_coords_from_read(r)
            if coords is None:
                continue
            chrom, start, end, tlen = coords

            sampled += 1

            if store_min_frag <= tlen <= store_max_frag:
                kept_lengths.append(tlen)
                if bed_fh is not None:
                    bed_fh.write(f"{chrom}\t{start}\t{end}\t{tlen}\n")
                    written_bed += 1

    if bed_fh is not None:
        bed_fh.close()

    chrM_wrote = 0
    if export_chrM_bam and mt_contigs and chrM_bam_path:
        chrM_wrote = export_chrM_alignments(
            bam_path=bam_path,
            out_bam=chrM_bam_path,
            mt_contigs=mt_contigs,
            min_mapq=min_mapq,
            include_dups=include_dups,
            include_secondary=include_secondary,
            include_supplementary=include_supplementary,
        )

    return {
        "bam": bam_path,
        "eligible": eligible_count,
        "sampling_fraction": samp_frac,
        "sampled_pairs": sampled,
        "kept_lengths": as_uint16_array(kept_lengths),
        "fragment_bed_path": fragment_bed_path if write_fragment_bed else None,
        "fragment_bed_lines": written_bed if write_fragment_bed else 0,
        "chrM_bam_path": chrM_bam_path or "",
        "chrM_alignments_written": int(chrM_wrote),
    }


# --------------------
# Plotting primitives
# --------------------

def _fmt_val(val: float, ndigits: int = 1, nan_str: str = "NA") -> str:
    if isinstance(val, float) and (math.isnan(val) or math.isinf(val)):
        return nan_str
    return f"{val:.{ndigits}f}"


def make_fld_plot(
    out_pdf: str,
    title: str,
    series_list: List[Dict],
    xmin: int,
    xmax: int,
    norm_min: int,
    norm_max: int,
    ylog: bool,
    inferno_colors: List[Tuple[float, float, float, float]],
    line_width: float,
):
    fig = plt.figure(figsize=(9, 5.5))
    ax = fig.add_subplot(111)

    for i, s in enumerate(series_list):
        ax.plot(
            s["bin_centers"],
            s["cpm_counts"],
            label=f"{s['label']} | N={s['n_norm']:,} | "
                  f"median={_fmt_val(s['median'])} bp | "
                  f"mode={_fmt_val(s['mode'], ndigits=0)} bp",
            lw=line_width,
            color=inferno_colors[i % len(inferno_colors)],
        )

    ax.set_xlim(xmin, xmax)
    if ylog:
        ax.set_yscale("log")
        ax.yaxis.set_major_locator(MaxNLocator(nbins=8, prune=None))
        ax.yaxis.set_minor_locator(MaxNLocator(nbins=20, prune=None))
        ax.set_ylabel("Counts per million (CPM), log10")
    else:
        ax.set_ylabel("Counts per million (CPM)")

    ax.set_xlabel("Insert size (bp)")
    ax.grid(True, which="both", alpha=0.25, linestyle="--")
    ax.set_title(title)
    ax.axvspan(norm_min, norm_max, color="k", alpha=0.04, lw=0)
    ax.legend(loc="best", frameon=False)
    fig.tight_layout()
    fig.savefig(out_pdf, dpi=300, bbox_inches="tight")
    plt.close(fig)


# ------------------
# BED state helpers
# ------------------

def intersect_lengths_with_state(
    fragments_bed_path: str,
    state_bed_path: str,
) -> np.ndarray:
    if not HAVE_PYBEDTOOLS:
        raise RuntimeError("pybedtools is required for --bed-states")

    import pybedtools  # local import
    frag_bt = pybedtools.BedTool(fragments_bed_path)
    state_bt = pybedtools.BedTool(state_bed_path)

    out = frag_bt.intersect(state_bt, wa=True, u=True)
    lens: List[int] = []
    for it in out:
        try:
            tlen = int(it[3])
            lens.append(tlen)
        except Exception:
            continue
    return as_uint16_array(lens)


# -----------------------
# Configuration handling
# -----------------------

def apply_config_defaults(parser: argparse.ArgumentParser, args: argparse.Namespace) -> argparse.Namespace:
    """If --config is supplied, merge in YAML/JSON values where CLI kept defaults."""
    cfg_path = getattr(args, "config", None)
    if not cfg_path:
        return args
    if not os.path.exists(cfg_path):
        raise FileNotFoundError(f"Config file not found: {cfg_path}")

    try:
        if cfg_path.lower().endswith((".yaml", ".yml")) and HAVE_YAML:
            with open(cfg_path, "r") as fh:
                cfg = yaml.safe_load(fh) or {}
        else:
            with open(cfg_path, "r") as fh:
                cfg = json.load(fh)
    except Exception as e:
        raise RuntimeError(f"Failed to parse config file {cfg_path}: {e}")

    for k, v in (cfg.items() if isinstance(cfg, dict) else []):
        if not hasattr(args, k):
            continue
        default = parser.get_default(k)
        current = getattr(args, k)
        if current == default and v is not None:
            setattr(args, k, v)
    return args


# ---------------
# Main entrypoint
# ---------------

def main():
    ap = argparse.ArgumentParser(
        prog="bam_insert_size_fld.py",
        formatter_class=argparse.RawDescriptionHelpFormatter,
        description=textwrap.dedent(
            """\
            Insert-size (fragment length) distributions (FLDs) from paired-end BAMs.

            Default behavior:
              • Use only autosomes (main scaffolds matching ^(chr)?[0-9]+$), excluding X/Y.
              • Exclude non-primary random/decoy/aux scrappy contigs by pattern.
              • Optionally export mitochondrial alignments (chrM/MT) to separate BAM.
              • NEW: optional compact TSV histograms with counts and CPM.
            """
        ),
    )

    # Required I/O
    ap.add_argument("--bams", nargs="+", required=True, help="Input sorted, indexed BAMs.")
    ap.add_argument("--outdir", required=True, help="Base output directory (a timestamped subfolder will be created).")

    # Labels & units
    ap.add_argument("--labels", type=str, default=None,
                    help="Comma-separated labels for BAMs (same order). Defaults to BAM basenames.")
    ap.add_argument("--units", type=str, default=None,
                    help="Comma-separated numeric 'Units' per BAM (same order).")

    # Counting / sampling
    ap.add_argument("--count-only", action="store_true", help="Only count eligible pairs per BAM and exit.")
    ap.add_argument("--downsample-to-min", action="store_true",
                    help="Downsample all BAMs to the shallowest eligible count across inputs.")
    ap.add_argument("--goal-reads", type=int, default=None,
                    help="Downsample each BAM to at most this many eligible pairs (overrides --downsample-to-min).")
    ap.add_argument("--seed", type=int, default=13, help="RNG seed for downsampling.")

    # Filters
    ap.add_argument("--min-mapq", type=int, default=30, help="Minimum MAPQ for reads (default: 30).")
    ap.add_argument("--include-duplicates", action="store_true", help="Include duplicate-marked reads (default: exclude).")
    ap.add_argument("--include-secondary", action="store_true", help="Include secondary alignments (default: exclude).")
    ap.add_argument("--include-supplementary", action="store_true", help="Include supplementary alignments (default: exclude).")

    # Ranges
    ap.add_argument("--store-min-frag", type=int, default=0, help="Minimum fragment length to store (default: 0).")
    ap.add_argument("--store-max-frag", type=int, default=750, help="Maximum fragment length to store (default: 750).")
    ap.add_argument("--plot-min-frag", type=int, default=None, help="x-axis min for plots (default: store-min-frag).")
    ap.add_argument("--plot-max-frag", type=int, default=None, help="x-axis max for plots (default: store-max-frag).")
    ap.add_argument("--norm-min-frag", type=int, default=None, help="CPM normalization min (default: store-min-frag).")
    ap.add_argument("--norm-max-frag", type=int, default=None, help="CPM normalization max (default: store-max-frag).")
    ap.add_argument("--stat-min-frag", type=int, default=None, help="Stats (median/mode) min (default: store-min-frag).")
    ap.add_argument("--stat-max-frag", type=int, default=None, help="Stats (median/mode) max (default: store-max-frag).")

    # Histogram TSV options (NEW)
    ap.add_argument("--save-hist-tsv", action="store_true",
                    help="Write per-BAM (and per-state) 1-bp histogram TSVs with columns len_bp,count,cpm.")
    ap.add_argument("--hist-min-frag", type=int, default=None,
                    help="Histogram min length (default: plot-min if set, else store-min).")
    ap.add_argument("--hist-max-frag", type=int, default=None,
                    help="Histogram max length (default: plot-max if set, else store-max).")
    ap.add_argument("--no-hist-compress", action="store_true",
                    help="Do NOT gzip-compress histogram TSVs (default: compress).")
    ap.add_argument("--no-raw-arrays", action="store_true",
                    help="Skip writing .npy raw arrays to disk (still computed in memory).")

    # Contig selection (autosomes-only by default)
    ap.add_argument("--contig-regex", type=str, default=r"^(chr)?[0-9]+$",
                    help="Regex for allowed contigs (default keeps main autosomes only).")
    ap.add_argument("--include-sex-chroms", action="store_true",
                    help="Include X and Y in addition to autosomes (default: exclude).")
    ap.add_argument("--include-all-contigs", action="store_true",
                    help="Ignore regex and include all contigs (default: False).")

    # Mitochondrial export
    ap.add_argument("--export-chrM-bam", action="store_true",
                    help="Write a separate BAM with all alignments mapping to mitochondrial contigs.")
    ap.add_argument("--mt-contigs", type=str, default="chrM,MT",
                    help="Comma-separated contig names considered mitochondrial (default: chrM,MT).")

    # BED states
    ap.add_argument("--bed-states", type=str, default=None,
                    help="Comma-separated BED files for state intersections (e.g., ChromHMM states).")
    ap.add_argument("--state-labels", type=str, default=None,
                    help="Comma-separated labels for states (same order as --bed-states).")

    # Plot styling toggles
    ap.add_argument("--font-size", type=float, default=10.0,
                    help="Base font size (points) for all text (default: 10.0).")
    ap.add_argument("--line-width", type=float, default=1.8,
                    help="Line width for plotted curves (default: 1.8).")

    # Misc
    ap.add_argument("--threads", type=int, default=max(1, mp.cpu_count() // 2),
                    help="Parallel worker processes (default: half of CPUs).")
    ap.add_argument("--no-plots", action="store_true", help="Compute arrays/stats only; skip plotting.")
    ap.add_argument("--config", type=str, default=None, help="Optional YAML/JSON config file (CLI overrides config).")

    args = ap.parse_args()
    args = apply_config_defaults(ap, args)

    # Validate BAMs & indices
    check_bams(args.bams)

    # Labels & Units
    if args.labels:
        labels = [x.strip() for x in args.labels.split(",")]
        if len(labels) != len(args.bams):
            raise ValueError("--labels length must match number of BAMs.")
    else:
        labels = [os.path.splitext(os.path.basename(b))[0] for b in args.bams]

    units = None
    if args.units:
        try:
            units = [float(x.strip()) for x in args.units.split(",")]
        except Exception:
            raise ValueError("--units must be comma-separated numeric values.")
        if len(units) != len(args.bams):
            raise ValueError("--units length must match number of BAMs.")

    # States
    state_paths = None
    state_labels = None
    if args.bed_states:
        if not HAVE_PYBEDTOOLS:
            raise RuntimeError("pybedtools/bedtools are required when --bed-states is given.")
        state_paths = [x.strip() for x in args.bed_states.split(",")]
        for p in state_paths:
            if not os.path.exists(p):
                raise FileNotFoundError(f"State BED not found: {p}")
        if args.state_labels:
            state_labels = [x.strip() for x in args.state_labels.split(",")]
            if len(state_labels) != len(state_paths):
                raise ValueError("--state-labels length must match --bed-states.")
        else:
            state_labels = [os.path.splitext(os.path.basename(p))[0] for p in state_paths]

    # Ranges defaults/validation
    plot_min = args.plot_min_frag if args.plot_min_frag is not None else args.store_min_frag
    plot_max = args.plot_max_frag if args.plot_max_frag is not None else args.store_max_frag
    norm_min = args.norm_min_frag if args.norm_min_frag is not None else args.store_min_frag
    norm_max = args.norm_max_frag if args.norm_max_frag is not None else args.store_max_frag
    stat_min = args.stat_min_frag if args.stat_min_frag is not None else args.store_min_frag
    stat_max = args.stat_max_frag if args.stat_max_frag is not None else args.store_max_frag

    hist_min = args.hist_min_frag if args.hist_min_frag is not None else plot_min
    hist_max = args.hist_max_frag if args.hist_max_frag is not None else plot_max
    hist_compress = not args.no_hist_compress

    if not (0 <= plot_min <= plot_max):
        raise ValueError("Invalid plot range.")
    if not (0 <= norm_min <= norm_max):
        raise ValueError("Invalid normalization range.")
    if not (0 <= stat_min <= stat_max):
        raise ValueError("Invalid stats range.")
    if not (args.store_min_frag <= args.store_max_frag):
        raise ValueError("Invalid store range.")
    if not (0 <= hist_min <= hist_max):
        raise ValueError("Invalid histogram range.")

    # Output directory (timestamped subfolder)
    ts = timestamp()
    base_outdir = os.path.abspath(args.outdir)
    outdir = os.path.join(base_outdir, f"fld_{ts}")
    ensure_dir(outdir)

    logger = setup_logging(outdir, ts)
    logger.info(f"bam_insert_size_fld.py v{__VERSION__} | TS={ts}")
    logger.info(f"Base output dir: {base_outdir}")
    logger.info(f"Timestamped dir: {outdir}")
    logger.info(f"BAMs: {', '.join(args.bams)}")

    # Apply plot style toggles
    apply_plot_style(base_font_size=args.font_size, line_width=args.line_width)

    # Environment/metadata
    meta = {
        "script": "bam_insert_size_fld.py",
        "version": __VERSION__,
        "timestamp": ts,
        "host": socket.gethostname(),
        "platform": platform.platform(),
        "python": sys.version.replace("\n", " "),
        "packages": {
            "numpy": np.__version__,
            "pandas": pd.__version__,
            "matplotlib": matplotlib.__version__,
            "pysam": pysam.__version__,
            "pybedtools": "available" if HAVE_PYBEDTOOLS else "not_available",
            "pyyaml": "available" if HAVE_YAML else "not_available",
        },
        "args": vars(args),
        "contigs": {},
    }

    # Prepare per-BAM allowed contigs and MT contigs
    mt_names = [x.strip() for x in args.mt_contigs.split(",") if x.strip()]
    allowed_per_bam: List[Set[str]] = []
    mt_present_per_bam: List[Set[str]] = []
    all_refs_per_bam: List[List[str]] = []

    for i, b in enumerate(args.bams):
        allowed, mt_present, refs = derive_contig_sets_for_bam(
            bam_path=b,
            contig_regex=args.contig_regex,
            include_sex_chroms=args.include_sex_chroms,
            include_all_contigs=args.include_all_contigs,
            mt_names=mt_names,
        )
        allowed_per_bam.append(allowed)
        mt_present_per_bam.append(mt_present)
        all_refs_per_bam.append(refs)
        meta["contigs"][labels[i]] = {
            "all_references": refs,
            "allowed_used_for_FLD": sorted(list(allowed)),
            "mt_present": sorted(list(mt_present)),
        }
        logger.info(f"[{labels[i]}] Allowed contigs used for FLD: {len(allowed)} (e.g., {', '.join(sorted(list(allowed))[:8])}{'...' if len(allowed)>8 else ''})")

    # -----------------
    # Phase 1: Counting
    # -----------------
    logger.info("Counting eligible read pairs per BAM (respecting contig filter)...")
    with mp.Pool(processes=max(1, args.threads)) as pool:
        eligible_counts = pool.starmap(
            count_eligible_pairs,
            [
                (
                    args.bams[i],
                    args.min_mapq,
                    args.include_duplicates,
                    args.include_secondary,
                    args.include_supplementary,
                    allowed_per_bam[i],
                )
                for i in range(len(args.bams))
            ],
        )

    counts_df = pd.DataFrame({"bam": args.bams, "label": labels, "eligible_pairs": eligible_counts})
    counts_tsv = os.path.join(outdir, f"counts_only__{ts}.tsv")
    save_tsv(counts_tsv, counts_df)
    logger.info(f"Wrote: {counts_tsv}")

    if args.count_only:
        meta["outputs"] = {"counts_only": counts_tsv}
        write_yaml(os.path.join(outdir, f"metadata__{ts}.yaml"), meta)
        logger.info("Count-only requested; exiting.")
        return

    # Determine sample goals
    if args.goal_reads is not None:
        goal = int(args.goal_reads)
        logger.info(f"Downsampling to goal reads: {goal:,}")
        sample_goals = [min(c, goal) for c in eligible_counts]
    elif args.downsample_to_min:
        minc = int(min(eligible_counts))
        logger.info(f"Downsampling to shallowest eligible count across inputs: {minc:,}")
        sample_goals = [minc for _ in eligible_counts]
    else:
        sample_goals = [None for _ in eligible_counts]  # no downsampling

    # -----------------------
    # Phase 2: Collect arrays
    # -----------------------
    logger.info("Collecting fragment lengths (respecting contig filter)...")
    collect_fn = partial(
        collect_fragments_for_bam,
        rng_seed=args.seed,
        min_mapq=args.min_mapq,
        include_dups=args.include_duplicates,
        include_secondary=args.include_secondary,
        include_supplementary=args.include_supplementary,
        store_min_frag=args.store_min_frag,
        store_max_frag=args.store_max_frag,
        write_fragment_bed=bool(state_paths),
        export_chrM_bam=bool(args.export_chrM_bam),
    )

    # Per-file fragment BED path (if states) and chrM BAM path (if requested)
    fragment_beds = []
    chrM_bams = []
    for i, b in enumerate(args.bams):
        bed_path = os.path.join(outdir, f"{labels[i]}__fragments_for_states__{ts}.bed") if state_paths else None
        fragment_beds.append(bed_path)
        if args.export_chrM_bam and mt_present_per_bam[i]:
            chrM_path = os.path.join(outdir, f"{labels[i]}__chrM_alignments__{ts}.bam")
        else:
            chrM_path = None
        chrM_bams.append(chrM_path)

    tasks = []
    for i, b in enumerate(args.bams):
        tasks.append(
            {
                "bam_path": b,
                "sample_goal": sample_goals[i],
                "eligible_count": eligible_counts[i],
                "fragment_bed_path": fragment_beds[i],
                "allowed_refs": allowed_per_bam[i],
                "mt_contigs": mt_present_per_bam[i],
                "chrM_bam_path": chrM_bams[i],
            }
        )

    results = []
    with mp.Pool(processes=max(1, args.threads)) as pool:
        asyncs = [pool.apply_async(collect_fn, kwds=t) for t in tasks]
        for a in asyncs:
            results.append(a.get())

    # Save .npy raw arrays (optional)
    raw_outputs = []
    if not args.no_raw_arrays:
        for i, res in enumerate(results):
            arr = res["kept_lengths"]
            npy_path = os.path.join(outdir, f"{labels[i]}__fraglens_<=_{args.store_max_frag}bp__{ts}.npy".replace("<=_", "<="))
            np.save(npy_path, arr)
            res["raw_array_path"] = npy_path
            raw_outputs.append(npy_path)
            logger.info(f"[{labels[i]}] Stored {arr.size:,} fraglengths to: {npy_path}")
    else:
        for res in results:
            res["raw_array_path"] = ""  # placeholder in stats

    # -----------------------------------
    # Build global plots, stats & hist TSV
    # -----------------------------------
    bin_edges = integer_bin_edges(plot_min, plot_max)
    bin_centers = np.arange(plot_min, plot_max + 1, dtype=int)
    hist_edges = integer_bin_edges(hist_min, hist_max)
    hist_centers = np.arange(hist_min, hist_max + 1, dtype=int)

    inferno_cols = get_inferno_colors(len(args.bams))

    stats_rows = []
    series_for_plots = []
    hist_tsvs = []

    for i, res in enumerate(results):
        arr = res["kept_lengths"]

        # Histogram for plotting range
        arr_plot = arr[(arr >= plot_min) & (arr <= plot_max)]
        counts_plot = np.histogram(arr_plot, bins=bin_edges)[0]

        # Denominator from RAW array over norm range (independent of plot range)
        denom = int(((arr >= norm_min) & (arr <= norm_max)).sum())
        cpm_plot = (counts_plot / denom * 1e6) if denom > 0 else np.zeros_like(counts_plot, dtype=float)

        # Stats in stats range
        arr_stat = arr[(arr >= stat_min) & (arr <= stat_max)]
        med = safe_median(arr_stat)
        stat_edges = integer_bin_edges(stat_min, stat_max)
        stat_centers = np.arange(stat_min, stat_max + 1, dtype=int)
        counts_stat = np.histogram(arr_stat, bins=stat_edges)[0]
        mode_val = compute_mode_from_hist(stat_centers, counts_stat)

        label = labels[i]
        if units:
            label = f"{label} ({units[i]}U)"

        series_for_plots.append({
            "label": label,
            "bin_centers": bin_centers,
            "cpm_counts": cpm_plot,
            "median": med,
            "mode": mode_val,
            "n_norm": int(denom),
        })

        # Save histogram TSV if requested
        if args.save_hist_tsv:
            mask_hist = (arr >= hist_min) & (arr <= hist_max)
            counts_hist = np.histogram(arr[mask_hist], bins=hist_edges)[0]
            cpm_hist = (counts_hist / denom * 1e6) if denom > 0 else np.zeros_like(counts_hist, dtype=float)
            df_hist = pd.DataFrame({"len_bp": hist_centers, "count": counts_hist, "cpm": cpm_hist})
            hist_path = os.path.join(outdir, f"{labels[i]}__hist_{hist_min}-{hist_max}bp__{ts}.tsv")
            save_tsv(hist_path, df_hist, compress=hist_compress)
            hist_tsvs.append(hist_path if not hist_compress else (hist_path + ".gz"))
            logger.info(f"[{labels[i]}] Wrote histogram TSV: {hist_tsvs[-1]}")

        stats_rows.append({
            "bam": args.bams[i],
            "label": labels[i],
            "units": (units[i] if units else ""),
            "eligible_pairs": int(res["eligible"]),
            "sample_goal": (sample_goals[i] if sample_goals[i] is not None else ""),
            "sampling_fraction": float(res["sampling_fraction"]),
            "sampled_pairs": int(res["sampled_pairs"]),
            "stored_fraglens": int(len(res["kept_lengths"])),
            "store_min_frag": args.store_min_frag,
            "store_max_frag": args.store_max_frag,
            "plot_min_frag": plot_min,
            "plot_max_frag": plot_max,
            "hist_min_frag": hist_min,
            "hist_max_frag": hist_max,
            "norm_min_frag": norm_min,
            "norm_max_frag": norm_max,
            "stat_min_frag": stat_min,
            "stat_max_frag": stat_max,
            "n_in_norm_range": int(denom),
            "median_bp": round(med, 3) if not math.isnan(med) else "",
            "mode_bp": round(mode_val, 3) if not math.isnan(mode_val) else "",
            "raw_array_npy": res["raw_array_path"],
            "fragment_bed_path": (res["fragment_bed_path"] or ""),
            "chrM_alignments_written": int(res.get("chrM_alignments_written", 0)),
            "chrM_bam_path": res.get("chrM_bam_path", ""),
        })

    stats_df = pd.DataFrame(stats_rows)
    stats_tsv = os.path.join(outdir, f"stats__{ts}.tsv")
    save_tsv(stats_tsv, stats_df)
    logger.info(f"Wrote stats: {stats_tsv}")

    outputs = {
        "raw_arrays": raw_outputs,
        "hist_tsvs": hist_tsvs,
        "stats": stats_tsv,
        "counts": counts_tsv,
    }

    if not args.no_plots:
        # Linear CPM
        lin_pdf = os.path.join(outdir, f"FLD_linear__{ts}.pdf")
        make_fld_plot(
            out_pdf=lin_pdf,
            title="Insert-size distributions (linear CPM)",
            series_list=series_for_plots,
            xmin=plot_min,
            xmax=plot_max,
            norm_min=norm_min,
            norm_max=norm_max,
            ylog=False,
            inferno_colors=inferno_cols,
            line_width=args.line_width,
        )
        logger.info(f"Wrote: {lin_pdf}")

        # Log10 CPM
        log_pdf = os.path.join(outdir, f"FLD_log10__{ts}.pdf")
        make_fld_plot(
            out_pdf=log_pdf,
            title="Insert-size distributions (log10 CPM)",
            series_list=series_for_plots,
            xmin=plot_min,
            xmax=plot_max,
            norm_min=norm_min,
            norm_max=norm_max,
            ylog=True,
            inferno_colors=inferno_cols,
            line_width=args.line_width,
        )
        logger.info(f"Wrote: {log_pdf}")

        outputs["plots_linear"] = lin_pdf
        outputs["plots_log"] = log_pdf

    # -----------------------------------------
    # Optional: BED state intersections & plots
    # -----------------------------------------
    if state_paths:
        logger.info("Running BED state intersections via pybedtools...")
        state_cols = get_distinct_state_colors(len(state_paths))

        def _task_intersect(bam_idx: int, state_idx: int):
            st_lab = state_labels[state_idx]
            frag_bed = results[bam_idx]["fragment_bed_path"]
            arr = intersect_lengths_with_state(frag_bed, state_paths[state_idx])
            # Save array if requested
            npy_path = ""
            if not args.no_raw_arrays:
                npy_path = os.path.join(
                    outdir,
                    f"{labels[bam_idx]}__state-{st_lab}__fraglens_<=_{args.store_max_frag}bp__{ts}.npy".replace("<=_", "<="),
                )
                np.save(npy_path, arr)
            # Save histogram TSV if requested
            hist_path = ""
            if args.save_hist_tsv:
                counts_hist = np.histogram(arr[(arr >= hist_min) & (arr <= hist_max)], bins=hist_edges)[0]
                denom = int(((arr >= norm_min) & (arr <= norm_max)).sum())
                cpm_hist = (counts_hist / denom * 1e6) if denom > 0 else np.zeros_like(counts_hist, dtype=float)
                df_hist = pd.DataFrame({"len_bp": hist_centers, "count": counts_hist, "cpm": cpm_hist})
                hist_path = os.path.join(outdir, f"{labels[bam_idx]}__state-{st_lab}__hist_{hist_min}-{hist_max}bp__{ts}.tsv")
                save_tsv(hist_path, df_hist, compress=hist_compress)
                if hist_compress:
                    hist_path = hist_path + ".gz"
            return (bam_idx, state_idx, npy_path, hist_path, arr.size)

        pair_jobs = [(i, j) for i in range(len(args.bams)) for j in range(len(state_paths))]
        per_state_rows = []
        per_bam_state_arrays: Dict[Tuple[int, int], str] = {}
        per_bam_state_hists: Dict[Tuple[int, int], str] = {}

        with mp.Pool(processes=max(1, args.threads)) as pool:
            inter_results = pool.starmap(_task_intersect, pair_jobs)

        for (bi, sj, npy_path, hist_path, nsz) in inter_results:
            if npy_path:
                per_bam_state_arrays[(bi, sj)] = npy_path
            if hist_path:
                per_bam_state_hists[(bi, sj)] = hist_path
            per_state_rows.append({
                "bam": args.bams[bi],
                "label": labels[bi],
                "units": (units[bi] if units else ""),
                "state": state_labels[sj],
                "stored_fraglens_state": int(nsz),
                "state_array_npy": npy_path,
                "state_hist_tsv": hist_path,
            })

        if per_state_rows:
            per_state_df = pd.DataFrame(per_state_rows)
            st_tsv = os.path.join(outdir, f"stats_states__{ts}.tsv")
            save_tsv(st_tsv, per_state_df)
            outputs["stats_states"] = st_tsv
            logger.info(f"Wrote: {st_tsv}")

        if not args.no_plots:
            # Per-BAM: overlay all states
            for bi in range(len(args.bams)):
                lab = labels[bi]
                series = []
                # Build arrays for plotting (load from file if saved; otherwise compute on-the-fly)
                for sj in range(len(state_paths)):
                    if (bi, sj) in per_bam_state_arrays and per_bam_state_arrays[(bi, sj)]:
                        arr = np.load(per_bam_state_arrays[(bi, sj)])
                    else:
                        # If arrays not saved, recompute quickly from BED intersections
                        arr = intersect_lengths_with_state(results[bi]["fragment_bed_path"], state_paths[sj])

                    counts = np.histogram(arr[(arr >= plot_min) & (arr <= plot_max)], bins=bin_edges)[0]
                    denom = int(((arr >= norm_min) & (arr <= norm_max)).sum())
                    cpm = (counts / denom * 1e6) if denom > 0 else np.zeros_like(counts, dtype=float)
                    arr_stat = arr[(arr >= stat_min) & (arr <= stat_max)]
                    med = safe_median(arr_stat)
                    stat_edges_loc = integer_bin_edges(stat_min, stat_max)
                    stat_centers_loc = np.arange(stat_min, stat_max + 1, dtype=int)
                    counts_stat = np.histogram(arr_stat, bins=stat_edges_loc)[0]
                    mode_val = compute_mode_from_hist(stat_centers_loc, counts_stat)
                    series.append({
                        "label": f"{state_labels[sj]} | N={int(denom):,} | median={_fmt_val(med)} | mode={_fmt_val(mode_val, 0)}",
                        "bin_centers": bin_centers,
                        "cpm_counts": cpm,
                        "median": med,
                        "mode": mode_val,
                        "n_norm": int(denom),
                    })

                def _plot_states_for_bam(out_pdf: str, ylog: bool):
                    fig = plt.figure(figsize=(9, 5.5))
                    ax = fig.add_subplot(111)
                    state_cols = get_distinct_state_colors(len(series))
                    for k, s in enumerate(series):
                        ax.plot(s["bin_centers"], s["cpm_counts"], lw=args.line_width,
                                color=state_cols[k % len(state_cols)], label=s["label"])
                    ax.set_xlim(plot_min, plot_max)
                    ax.set_yscale("log" if ylog else "linear")
                    ax.set_ylabel("Counts per million (CPM), log10" if ylog else "Counts per million (CPM)")
                    ax.set_xlabel("Insert size (bp)")
                    ax.grid(True, which="both", alpha=0.25, linestyle="--")
                    ax.set_title(f"{lab}: FLDs by state")
                    ax.axvspan(norm_min, norm_max, color="k", alpha=0.04, lw=0)
                    ax.legend(loc="best", frameon=False)
                    fig.tight_layout()
                    fig.savefig(out_pdf, dpi=300, bbox_inches="tight")
                    plt.close(fig)

                out_lin = os.path.join(outdir, f"FLD_states_linear__{lab}__{ts}.pdf")
                _plot_states_for_bam(out_lin, ylog=False)
                logger.info(f"Wrote: {out_lin}")
                out_log = os.path.join(outdir, f"FLD_states_log10__{lab}__{ts}.pdf")
                _plot_states_for_bam(out_log, ylog=True)
                logger.info(f"Wrote: {out_log}")

            # Per-state: overlay all BAMs
            cols = inferno_cols
            for sj in range(len(state_paths)):
                st_lab = state_labels[sj]
                series = []
                for bi in range(len(args.bams)):
                    if (bi, sj) in per_bam_state_arrays and per_bam_state_arrays[(bi, sj)]:
                        arr = np.load(per_bam_state_arrays[(bi, sj)])
                    else:
                        arr = intersect_lengths_with_state(results[bi]["fragment_bed_path"], state_paths[sj])

                    counts = np.histogram(arr[(arr >= plot_min) & (arr <= plot_max)], bins=bin_edges)[0]
                    denom = int(((arr >= norm_min) & (arr <= norm_max)).sum())
                    cpm = (counts / denom * 1e6) if denom > 0 else np.zeros_like(counts, dtype=float)
                    arr_stat = arr[(arr >= stat_min) & (arr <= stat_max)]
                    med = safe_median(arr_stat)
                    stat_edges_loc = integer_bin_edges(stat_min, stat_max)
                    stat_centers_loc = np.arange(stat_min, stat_max + 1, dtype=int)
                    counts_stat = np.histogram(arr_stat, bins=stat_edges_loc)[0]
                    mode_val = compute_mode_from_hist(stat_centers_loc, counts_stat)
                    label = labels[bi]
                    if units:
                        label = f"{label} ({units[bi]}U)"
                    series.append({
                        "label": f"{label} | N={int(denom):,} | median={_fmt_val(med)} | mode={_fmt_val(mode_val, 0)}",
                        "bin_centers": bin_centers,
                        "cpm_counts": cpm,
                        "median": med,
                        "mode": mode_val,
                        "n_norm": int(denom),
                    })

                def _plot_bams_for_state(out_pdf: str, ylog: bool):
                    fig = plt.figure(figsize=(9, 5.5))
                    ax = fig.add_subplot(111)
                    for k, s in enumerate(series):
                        ax.plot(s["bin_centers"], s["cpm_counts"], lw=args.line_width,
                                color=cols[k % len(cols)], label=s["label"])
                    ax.set_xlim(plot_min, plot_max)
                    ax.set_yscale("log" if ylog else "linear")
                    ax.set_ylabel("Counts per million (CPM), log10" if ylog else "Counts per million (CPM)")
                    ax.set_xlabel("Insert size (bp)")
                    ax.grid(True, which="both", alpha=0.25, linestyle="--")
                    ax.set_title(f"State: {st_lab} — FLDs across BAMs")
                    ax.axvspan(norm_min, norm_max, color="k", alpha=0.04, lw=0)
                    ax.legend(loc="best", frameon=False)
                    fig.tight_layout()
                    fig.savefig(out_pdf, dpi=300, bbox_inches="tight")
                    plt.close(fig)

                out_lin = os.path.join(outdir, f"FLD_by_state_linear__{st_lab}__{ts}.pdf")
                _plot_bams_for_state(out_lin, ylog=False)
                logger.info(f"Wrote: {out_lin}")
                out_log = os.path.join(outdir, f"FLD_by_state_log10__{st_lab}__{ts}.pdf")
                _plot_bams_for_state(out_log, ylog=True)
                logger.info(f"Wrote: {out_log}")

    # ----------------
    # Write metadata
    # ----------------
    outputs.update({
        "chrM_bams": [r.get("chrM_bam_path", "") for r in results] if args.export_chrM_bam else [],
    })
    meta["outputs"] = outputs
    write_yaml(os.path.join(outdir, f"metadata__{ts}.yaml"), meta)
    logger.info("Done.")


if __name__ == "__main__":
    try:
        main()
    except KeyboardInterrupt:
        print("\n[Interrupted] Exiting.", file=sys.stderr)
        sys.exit(130)
    except Exception as e:
        print(f"[FATAL] {e}", file=sys.stderr)
        sys.exit(1)
```
