## Supplementary material for "CAD-C: An engineered nuclease enables repair-free *in situ* proximity ligation and nucleosome-resolution chromosome walks in human cells": Custom code: bamfragend_GM12878_CHOMP_4U_v1_1.sh

```
#!/bin/bash
#SBATCH --job-name=GC_bal_bam
#SBATCH --output=%x_%j.out
#SBATCH --error=%x_%j.err
#SBATCH --nodes=1
#SBATCH --ntasks=1
#SBATCH --cpus-per-task=8

# Load Python (or your relevant environment)
# module load python/3.8

# Paths to script, BAM, FASTA, and output directory
SCRIPT="/rugpfs/fs0/risc_lab/scratch/jsoroczyns/CAD/CAD-C/scripts/2025_03_28/GCbal_bam_frag_and_upstream_bases_v1_20250328.py"
BAM="/rugpfs/fs0/risc_lab/scratch/jhickling/CAD/CAD-C/GM_CADC/GM_CAD_Titration/CHOMP_GM12878_4U_TR2_S8_001.trim.st.all.blft.qft.rmdup.bam"
FASTA="/rugpfs/fs0/risc_lab/store/risc_data/downloaded/hg38/genome/Sequence/WholeGenomeFasta/genome.fa"
OUTDIR="/rugpfs/fs0/risc_lab/scratch/jsoroczyns/CAD/CHOMP/2025_Frag_End_Composition_Analyses/"

# Optional: Activate a virtual env if needed
# source /path/to/venv/bin/activate

# Run the Python script with desired arguments
python "$SCRIPT" \
    -b "$BAM" \
    -f "$FASTA" \
    -o "$OUTDIR" \
    --chrom "chr1" \
    --bin-size 1000 \
    --flank 50 \
    -n 1000000 \
    --debug

# Optional: Deactivate virtual env
# deactivate
```
