## Supplementary material for "CAD-C: An engineered nuclease enables repair-free *in situ* proximity ligation and nucleosome-resolution chromosome walks in human cells": Custom code: bigwig_GC_cov_v7_wmeta_20250329.py

```
#!/usr/bin/env python3
"""
GC Bias Analysis Tool (Exact 1kb Bins, No Partials) - Filter-First Version
==========================================================================

This version filters zero-coverage and raw-coverage outlier bins BEFORE computing
the global mean coverage. Then it normalizes coverage only for the remaining bins.

Steps:
  1) Create exact 1kb bins (no partials).
  2) Compute GC fraction for each bin.
  3) Extract raw coverage for each bin from BigWig.
  4) Filter out bins with coverage <= 0 or NaN (zero coverage).
  5) Filter out bins outside the 1% and 99% quantiles (based on *raw* coverage).
  6) Compute mean coverage on the final set of bins.
  7) Compute norm_cov = coverage / mean_coverage for that final set.
  8) Group by GC, plot, etc.

All zero-coverage and outlier bins are saved separately and are NOT included
in the final normalization or plots.

Author: Your Name
Date: 2025-03-27
"""

import os
import sys
import argparse
import logging
import datetime
import json
import math

import numpy as np
import pandas as pd
import matplotlib
matplotlib.use('Agg')  # Safe for cluster environment without DISPLAY
matplotlib.rcParams['pdf.fonttype'] = 42  # Embed fonts in PDF (Type 42)
import matplotlib.pyplot as plt

import bioframe          # For GC calculation
import pyBigWig          # For coverage extraction
from pyfaidx import Fasta  # For FASTA loading with bioframe.frac_gc


def parse_args():
    parser = argparse.ArgumentParser(
        description="Filter-first GC/coverage analysis: remove zero/outlier bins, then compute mean coverage."
    )
    parser.add_argument("-b", "--bigwig", required=True,
                        help="Input BigWig file (.bw or .bigwig)")
    parser.add_argument("-f", "--fasta", required=True,
                        help="Reference genome FASTA file")
    parser.add_argument("--binsize", type=int, default=1000,
                        help="Bin size in base pairs (default: 1000)")
    parser.add_argument("--include-all-chroms", action="store_true",
                        help="Include all chromosomes/contigs (default: only autosomes chr1-22)")
    parser.add_argument("-o", "--outdir", default="output",
                        help="Output directory (default: ./output)")
    parser.add_argument("--log", default="INFO",
                        help="Logging level (default: INFO)")
    return parser.parse_args()


def setup_logging(log_level):
    logging.basicConfig(format='%(asctime)s %(levelname)s: %(message)s',
                        level=log_level)


def load_reference_fasta(fasta_path):
    try:
        # Load FASTA with pyfaidx
        fasta = Fasta(fasta_path, as_raw=True)
        return fasta
    except Exception as e:
        logging.error(f"Error loading FASTA file: {fasta_path} -- {str(e)}")
        sys.exit(1)


def load_bigwig(bigwig_path):
    try:
        bw = pyBigWig.open(bigwig_path)
        return bw
    except Exception as e:
        logging.error(f"Error opening BigWig file: {bigwig_path} -- {str(e)}")
        sys.exit(1)


def get_chromsizes(bw, include_all):
    chroms_dict = bw.chroms()
    if not include_all:
        autosomes = [f"chr{i}" for i in range(1, 23)]
        chroms_dict = {c: sz for c, sz in chroms_dict.items() if c in autosomes}
    return chroms_dict


def bin_genome_exact_1kb(chromsizes, binsize):
    """
    Manually create exact 1 kb bins, ignoring any partial bin at the end.
    """
    all_bins = []
    for chrom, size in chromsizes.items():
        n_full = size // binsize
        if n_full == 0:
            continue
        starts = np.arange(0, n_full * binsize, binsize, dtype=np.int64)
        ends = starts + binsize
        df = pd.DataFrame({'chrom': chrom, 'start': starts, 'end': ends})
        all_bins.append(df)

    if not all_bins:
        logging.error("No bins could be generated (check chromsizes?).")
        sys.exit(1)

    bins_df = pd.concat(all_bins, ignore_index=True)
    return bins_df


def calculate_gc(bins_df, fasta):
    try:
        bins_gc = bioframe.frac_gc(bins_df, fasta, mapped_only=True, return_input=True)
        return bins_gc
    except Exception as e:
        logging.error(f"Error calculating GC content: {str(e)}")
        sys.exit(1)


def extract_coverage_single_call(bw, bins_df, binsize):
    """
    For each chromosome, call bw.stats() once with nBins = # of bins.
    """
    coverage_dfs = []
    for chrom, sub_df in bins_df.groupby('chrom'):
        n_chr_bins = len(sub_df)
        if n_chr_bins == 0:
            continue

        # coverage range is from 0..(binsize * n_chr_bins)
        start_coord = 0
        end_coord = n_chr_bins * binsize
        try:
            cov_vals = bw.stats(chrom, start_coord, end_coord,
                                type='mean', nBins=n_chr_bins, exact=True)
        except Exception as e:
            logging.warning(f"Error fetching stats for {chrom}: {str(e)}")
            cov_vals = [0.0] * n_chr_bins
        
        # Clean up
        cov_vals = [
            0.0 if (v is None or (isinstance(v, float) and math.isnan(v))) else float(v)
            for v in cov_vals
        ]
        sub_df = sub_df.copy()
        sub_df['coverage'] = cov_vals
        coverage_dfs.append(sub_df)

    covered_df = pd.concat(coverage_dfs, ignore_index=True)
    return covered_df


def filter_coverage_zero_outliers(bins_df):
    """
    1) Separate out zero (<= 0) coverage bins.
    2) Among the rest, find 1%/99% quantiles based on *raw coverage*.
    3) Filter out bins outside that range.
    4) Return final filtered bins, plus the zero-coverage set and outlier set.

    Note: This is done on RAW coverage (not normalized).
    """

    # Bins with zero coverage
    zero_mask = (bins_df['coverage'] <= 0) | (bins_df['coverage'].isna())
    bins_zero_cov = bins_df[zero_mask].copy()
    bins_filtered = bins_df[~zero_mask].copy()

    # Outlier thresholds on raw coverage
    lower_thr = bins_filtered['coverage'].quantile(0.01)
    upper_thr = bins_filtered['coverage'].quantile(0.99)
    outlier_mask = (bins_filtered['coverage'] < lower_thr) | (bins_filtered['coverage'] > upper_thr)
    bins_outliers = bins_filtered[outlier_mask].copy()

    # Final bins that remain
    bins_final = bins_filtered[~outlier_mask].copy()

    logging.info(f"Zero coverage bins: {bins_zero_cov.shape[0]}")
    logging.info(f"Outlier coverage bins: {bins_outliers.shape[0]}")
    logging.info(f"Remaining bins: {bins_final.shape[0]}")
    logging.info(f"Outlier thresholds (raw coverage): lower={lower_thr:.4f}, upper={upper_thr:.4f}")

    return bins_final, bins_zero_cov, bins_outliers, lower_thr, upper_thr


def compute_normalized_coverage_after_filter(bins_final):
    """
    Compute mean coverage for the "final" bins that passed filtering,
    then create a new column 'norm_cov' in that final DataFrame.
    """
    if bins_final.empty:
        logging.error("No bins remain after filtering; cannot compute mean coverage.")
        sys.exit(1)

    mean_cov = bins_final['coverage'].mean()
    logging.info(f"Mean coverage (after filter) = {mean_cov:.4f}")

    bins_final['norm_cov'] = bins_final['coverage'] / mean_cov

    # Basic stats
    desc = bins_final['norm_cov'].describe()
    logging.info(f"Normalized coverage stats:\n{desc}")

    return bins_final, mean_cov


def group_by_gc(bins_df, num_groups=100):
    """
    Sort bins by GC, split into quantiles, compute summary stats.
    Expects bins_df to have 'norm_cov' and 'GC'.
    """
    if bins_df.empty:
        logging.warning("No bins to group. Returning empty group_df.")
        return pd.DataFrame()

    bins_sorted = bins_df.sort_values('GC').reset_index(drop=True)
    total_bins = len(bins_sorted)
    group_size = total_bins // num_groups if num_groups > 0 else total_bins

    group_stats = []
    for g in range(num_groups):
        start_idx = g * group_size
        if g == num_groups - 1:
            group_data = bins_sorted.iloc[start_idx:]
        else:
            group_data = bins_sorted.iloc[start_idx:start_idx + group_size]

        if group_data.empty:
            continue

        gc_vals = group_data['GC'] * 100.0
        cov_vals = group_data['norm_cov']
        group_stat = {
            'group': g,
            'count': len(group_data),
            'gc_min': gc_vals.min(),
            'gc_max': gc_vals.max(),
            'gc_mean': gc_vals.mean(),
            'cov_median': np.median(cov_vals),
            'cov_Q1': np.percentile(cov_vals, 25),
            'cov_Q3': np.percentile(cov_vals, 75)
        }
        group_stat['cov_IQR'] = group_stat['cov_Q3'] - group_stat['cov_Q1']
        group_stats.append(group_stat)

    group_df = pd.DataFrame(group_stats)
    return group_df


def plot_gc_bias(group_df, output_pdf, bw_filename):
    """
    Plot coverage vs GC in linear (left panel) and log2 (right panel) scale.
    Uses norm_cov from the group stats (median + IQR).
    """
    if group_df.empty:
        logging.warning("No GC groups to plot. Skipping.")
        return

    fig, axs = plt.subplots(1, 2, figsize=(14, 6))

    # -- Linear scale
    ax = axs[0]
    for _, row in group_df.iterrows():
        # IQR bar
        ax.bar(row['gc_min'],
               row['cov_IQR'],
               width=row['gc_max'] - row['gc_min'],
               bottom=row['cov_Q1'],
               align='edge',
               color='skyblue',
               edgecolor='none')
        # median line
        ax.hlines(row['cov_median'], row['gc_min'], row['gc_max'],
                  colors='blue', linewidth=1)
    ax.axhline(1.0, color='black', linestyle=':', label="Mean coverage = 1.0")
    ax.set_xlabel("GC Content (%)")
    ax.set_ylabel("Normalized Coverage (Linear)")
    ax.set_title(f"Coverage vs. GC (Linear Scale)\n(Input file: {bw_filename})")
    ax.legend()

    # -- Log2 scale
    ax2 = axs[1]
    eps = 1e-6
    for _, row in group_df.iterrows():
        q1_log = np.log2(row['cov_Q1'] + eps)
        q3_log = np.log2(row['cov_Q3'] + eps)
        med_log = np.log2(row['cov_median'] + eps)
        ax2.bar(row['gc_min'],
                q3_log - q1_log,
                width=row['gc_max'] - row['gc_min'],
                bottom=q1_log,
                align='edge',
                color='lightgreen',
                edgecolor='none')
        ax2.hlines(med_log, row['gc_min'], row['gc_max'],
                   colors='green', linewidth=1)
    ax2.axhline(0, color='black', linestyle=':', label="Log2(1) = 0")
    ax2.set_xlabel("GC Content (%)")
    ax2.set_ylabel("Log2 Normalized Coverage")
    ax2.set_title(f"Coverage vs. GC (Log2 Scale)\n(Input file: {bw_filename})")
    ax2.legend()

    plt.tight_layout()
    plt.savefig(output_pdf)
    plt.close()
    logging.info(f"GC bias plot saved as {output_pdf}")


def compute_gc_histogram(bins_df, output_pdf_hist, bw_filename):
    """
    Simple histogram of GC distribution in the final bins.
    """
    if bins_df.empty:
        logging.warning("No bins for GC histogram. Skipping.")
        return

    fig, ax = plt.subplots(figsize=(6, 4))
    gc_percent = bins_df['GC'] * 100.0
    ax.hist(gc_percent, bins=50, color='gray', edgecolor='black')
    ax.set_xlabel("GC Content (%)")
    ax.set_ylabel("Number of Bins")
    ax.set_title(f"GC Content Distribution (1kb Bins, Filtered)\n(Input file: {bw_filename})")
    plt.tight_layout()
    plt.savefig(output_pdf_hist)
    plt.close()
    logging.info(f"GC histogram saved as {output_pdf_hist}")


def main():
    args = parse_args()
    setup_logging(args.log)

    os.makedirs(args.outdir, exist_ok=True)
    timestamp = datetime.datetime.now().strftime("%Y%m%d_%H%M%S")
    logging.info("Starting GC Bias Analysis (Filter-First Approach)")

    # Capture the full input file name (with extension) for output naming & titles
    bw_filename = os.path.basename(args.bigwig)

    # 1) Load data
    fasta = load_reference_fasta(args.fasta)
    bw = load_bigwig(args.bigwig)
    chromsizes = get_chromsizes(bw, args.include_all_chroms)
    logging.info(f"Using {len(chromsizes)} chromosomes for analysis.")

    # 2) Generate exact 1 kb bins
    bins_df = bin_genome_exact_1kb(chromsizes, args.binsize)
    logging.info(f"Generated {len(bins_df)} x {args.binsize} bp bins.")

    # 3) GC content
    bins_df = calculate_gc(bins_df, fasta)
    logging.info("Calculated GC for each bin.")

    # 4) Raw coverage
    bins_df = extract_coverage_single_call(bw, bins_df, args.binsize)
    bw.close()
    logging.info("Extracted coverage for each bin from BigWig.")

    # 5) Filter zeros & outliers on raw coverage
    final_bins, bins_zero_cov, bins_outliers, low_thr, high_thr = filter_coverage_zero_outliers(bins_df)

    # 6) Compute mean coverage AFTER filtering
    final_bins, mean_cov = compute_normalized_coverage_after_filter(final_bins)

    # 7) Save data
    raw_bins_tsv = os.path.join(args.outdir, f"bins_data_raw_{bw_filename}_{timestamp}.tsv")
    bins_df.to_csv(raw_bins_tsv, sep='\t', index=False)
    logging.info(f"Raw bins (unfiltered) saved to: {raw_bins_tsv}")

    zero_cov_tsv = os.path.join(args.outdir, f"bins_zero_cov_{bw_filename}_{timestamp}.tsv")
    bins_zero_cov.to_csv(zero_cov_tsv, sep='\t', index=False)
    logging.info(f"Bins with zero coverage saved to: {zero_cov_tsv}")

    outliers_tsv = os.path.join(args.outdir, f"bins_outliers_{bw_filename}_{timestamp}.tsv")
    bins_outliers.to_csv(outliers_tsv, sep='\t', index=False)
    logging.info(f"Bins with coverage <{low_thr} or >{high_thr} saved to: {outliers_tsv}")

    final_tsv = os.path.join(args.outdir, f"bins_filtered_{bw_filename}_{timestamp}.tsv")
    final_bins.to_csv(final_tsv, sep='\t', index=False)
    logging.info(f"Final filtered bins (with norm_cov) saved to: {final_tsv}")

    # 8) Group by GC and plot
    group_df = group_by_gc(final_bins, num_groups=100)
    gc_groups_tsv = os.path.join(args.outdir, f"gc_groups_{bw_filename}_{timestamp}.tsv")
    group_df.to_csv(gc_groups_tsv, sep='\t', index=False)
    logging.info(f"GC group statistics saved to: {gc_groups_tsv}")

    plot_pdf = os.path.join(args.outdir, f"GC_bias_plot_{bw_filename}_{timestamp}.pdf")
    plot_gc_bias(group_df, plot_pdf, bw_filename)

    hist_pdf = os.path.join(args.outdir, f"GC_histogram_{bw_filename}_{timestamp}.pdf")
    compute_gc_histogram(final_bins, hist_pdf, bw_filename)

    # 9) Summary
    summary = {
        "timestamp": timestamp,
        "bigwig": os.path.abspath(args.bigwig),
        "fasta": os.path.abspath(args.fasta),
        "binsize": args.binsize,
        "include_all_chroms": args.include_all_chroms,
        "n_raw_bins": len(bins_df),
        "n_zero_cov_bins": len(bins_zero_cov),
        "n_outlier_bins": len(bins_outliers),
        "n_final_bins": len(final_bins),
        "mean_coverage_after_filter": mean_cov,
        "low_coverage_threshold": low_thr,
        "high_coverage_threshold": high_thr,
        "raw_bins_tsv": os.path.abspath(raw_bins_tsv),
        "bins_zero_cov_tsv": os.path.abspath(zero_cov_tsv),
        "bins_outliers_tsv": os.path.abspath(outliers_tsv),
        "final_bins_tsv": os.path.abspath(final_tsv),
        "gc_groups_tsv": os.path.abspath(gc_groups_tsv),
        "GC_bias_plot_pdf": os.path.abspath(plot_pdf),
        "GC_histogram_pdf": os.path.abspath(hist_pdf),
    }

    summary_file = os.path.join(args.outdir, f"analysis_summary_{bw_filename}_{timestamp}.json")
    with open(summary_file, 'w') as f:
        json.dump(summary, f, indent=4)
    logging.info(f"Summary saved to: {summary_file}")

    logging.info("Analysis complete (filter-first approach, mean coverage computed after filtering).")


if __name__ == "__main__":
    main()
```
