## Supplementary material for "CAD-C: An engineered nuclease enables repair-free *in situ* proximity ligation and nucleosome-resolution chromosome walks in human cells": Custom code: correlate_bigwigs_v3_WITHseaborn_20250331.py

```
#!/usr/bin/env python3
# -*- coding: utf-8 -*-
#
# Script: bigwig_correlation_analysis.py
# Description: Computes pairwise correlations between multiple BigWig files
#              over genomic bins and produces visualizations (pair plots and heatmaps).
#              Supports restricting analysis to specific chromosomes or regions,
#              filtering by BED features with extension, normalization options,
#              and multiprocessing. Now allows custom file aliases, a font-scale
#              option, and uses a shaded bivariate KDE plot when --plot-type=kde.

import os
import json
import math
import argparse
import numpy as np
import pandas as pd
import seaborn as sns
import matplotlib
matplotlib.use('Agg')  # For non-interactive backend (useful on clusters)
import matplotlib.pyplot as plt
from joblib import Parallel, delayed
import pyBigWig
from datetime import datetime

# Use Type 42 fonts in PDF (avoid Type 3)
matplotlib.rcParams['pdf.fonttype'] = 42
matplotlib.rcParams['ps.fonttype'] = 42

def main():
    parser = argparse.ArgumentParser(
        description='Compute pairwise correlations between BigWig files over genomic bins and generate plots.')
    parser.add_argument('bigwigs', nargs='+', 
                        help='Paths to input BigWig files (two or more).')
    parser.add_argument('--aliases', nargs='*', default=[],
                        help='Optional custom aliases for the BigWig files, in the same order. '
                             'If omitted, file basenames are used. Example: --aliases sampleA sampleB ...')
    parser.add_argument('-b', '--binsize', type=int, default=100,
                        help='Genomic bin size in base pairs (default: 100).')
    parser.add_argument('--chroms', dest='chroms_file',
                        help='TSV file of chromosomes to include (with lengths). If not provided, '
                             'use human autosomes chr1-22 by default.')
    parser.add_argument('--bed', dest='bed_file',
                        help='Optional BED file to filter regions (e.g. ChromHMM states or point features).')
    parser.add_argument('--bed-extension', type=int, default=0,
                        help='Extend BED intervals by this many base pairs on each side (default: 0).')
    parser.add_argument('--norm', dest='norm_method', 
                        choices=['none', 'mean', 'zscore'], default='mean',
                        help='Normalization method: none, mean, or zscore (default: mean).')
    parser.add_argument('--include-zeros', action='store_true', default=False,
                        help='Include bins with zero coverage (default: exclude them).')
    parser.add_argument('--trim-percent', type=float, default=1.0,
                        help='Percent of bins to trim from lower & upper tails of distribution (default: 1).')
    parser.add_argument('--metrics', nargs='+', 
                        choices=['pearson', 'spearman', 'kendall'],
                        default=['pearson', 'spearman', 'kendall'],
                        help='Correlation metrics to compute (default: all three).')
    parser.add_argument('--threads', '-p', type=int, default=-1,
                        help='Number of parallel threads/processes (default: all cores).')
    parser.add_argument('--plot-type', 
                        choices=['scatter', 'hex', 'kde'], default='hex',
                        help='Pairwise plot style: scatter, hexbin, or KDE (default: hex).')
    parser.add_argument('--font-scale', type=float, default=1.2,
                        help='Scale all font sizes by this factor (default: 1.2).')
    parser.add_argument('--outdir', default=None,
                        help='Output directory name. If omitted, a timestamped dir is created.')

    args = parser.parse_args()

    # Create output directory
    timestamp = datetime.now().strftime('%Y%m%d_%H%M%S')
    if args.outdir:
        base_out = args.outdir
    else:
        base_out = 'bigwig_correlation_output'
    outdir = f"{base_out}_{timestamp}"
    os.makedirs(outdir, exist_ok=True)

    # Store run parameters in metadata
    metadata = {
        'timestamp': timestamp,
        'bigwigs': args.bigwigs,
        'aliases': args.aliases,
        'binsize': args.binsize,
        'chroms_file': args.chroms_file,
        'bed_file': args.bed_file,
        'bed_extension': args.bed_extension,
        'norm_method': args.norm_method,
        'include_zeros': args.include_zeros,
        'trim_percent': args.trim_percent,
        'metrics': args.metrics,
        'plot_type': args.plot_type,
        'font_scale': args.font_scale,
        'outdir': outdir
    }

    # Log summary
    print(f"[INFO] BigWig files: {args.bigwigs}")
    if args.aliases:
        print(f"[INFO] Custom aliases: {args.aliases}")
    print(f"[INFO] Bin size: {args.binsize} bp")
    if args.chroms_file:
        print(f"[INFO] Chromosomes from: {args.chroms_file}")
    else:
        print(f"[INFO] Default chromosomes: chr1-22")
    if args.bed_file:
        print(f"[INFO] BED file: {args.bed_file} (extension: {args.bed_extension} bp)")
    print(f"[INFO] Normalization: {args.norm_method}")
    print(f"[INFO] Include zeros: {args.include_zeros}")
    print(f"[INFO] Trim percent: {args.trim_percent}%")
    print(f"[INFO] Correlation metrics: {args.metrics}")
    print(f"[INFO] Pairwise plot type: {args.plot_type}")
    print(f"[INFO] Font scale: {args.font_scale}")
    thread_msg = str(args.threads) if args.threads != -1 else "all available"
    print(f"[INFO] Using up to {thread_msg} threads.")
    print(f"[INFO] Output directory: {outdir}")

    # Attempt to open BigWigs
    bw_refs = []
    try:
        for bw_path in args.bigwigs:
            bw_refs.append(pyBigWig.open(bw_path))
    except Exception as e:
        print(f"[ERROR] Failed to open BigWig file: {e}")
        return

    if not bw_refs:
        print("[ERROR] No BigWig references opened.")
        return

    first_bw = bw_refs[0]

    # Determine chromosomes to analyze
    allowed_chroms = []
    chrom_lengths = {}
    if args.chroms_file:
        with open(args.chroms_file, 'r') as cf:
            for line in cf:
                line = line.strip()
                if not line or line.startswith('#'):
                    continue
                parts = line.split()
                name = parts[0]
                length = None
                if len(parts) > 1:
                    try:
                        length = int(parts[1])
                    except ValueError:
                        length = None
                allowed_chroms.append(name)
                if length is not None:
                    chrom_lengths[name] = length
    else:
        allowed_chroms = [f"chr{i}" for i in range(1, 23)]

    bw_chrom_dict = first_bw.chroms()
    for chrom in list(allowed_chroms):
        if chrom not in bw_chrom_dict:
            print(f"[WARN] Chrom {chrom} not in first BigWig, skipping.")
            allowed_chroms.remove(chrom)
            if chrom in chrom_lengths:
                chrom_lengths.pop(chrom, None)
        else:
            if chrom not in chrom_lengths:
                chrom_lengths[chrom] = bw_chrom_dict[chrom]

    if not allowed_chroms:
        print("[ERROR] No allowed chromosomes remain after checking.")
        for bw in bw_refs:
            bw.close()
        return

    # If BED file is provided, parse/merge intervals
    bed_intervals = {}
    if args.bed_file:
        print(f"[INFO] Reading BED: {args.bed_file}")
        with open(args.bed_file, 'r') as bf:
            for line in bf:
                line = line.strip()
                if not line or line.startswith('#'):
                    continue
                parts = line.split()
                if len(parts) < 3:
                    continue
                chrom = parts[0]
                start = int(parts[1])
                end = int(parts[2])
                if chrom not in allowed_chroms:
                    continue
                start_ext = max(0, start - args.bed_extension)
                chrom_len = chrom_lengths.get(chrom, 0)
                end_ext = min(chrom_len, end + args.bed_extension)
                if end_ext <= start_ext:
                    continue
                if chrom not in bed_intervals:
                    bed_intervals[chrom] = []
                bed_intervals[chrom].append((start_ext, end_ext))

        # Merge intervals
        for chrom, intervals in bed_intervals.items():
            intervals.sort(key=lambda x: x[0])
            merged = []
            cstart, cend = intervals[0]
            for (s, e) in intervals[1:]:
                if s <= cend:
                    if e > cend:
                        cend = e
                else:
                    merged.append((cstart, cend))
                    cstart, cend = s, e
            merged.append((cstart, cend))
            bed_intervals[chrom] = merged

        # Restrict allowed_chroms
        allowed_chroms = [c for c in allowed_chroms if c in bed_intervals]
        if not allowed_chroms:
            print("[WARN] BED filtering removed all chromosomes.")
            for bw in bw_refs:
                bw.close()
            return

    # Close BigWig references in main
    for bw in bw_refs:
        bw.close()

    # Prepare tasks
    tasks = []
    for chrom in allowed_chroms:
        if args.bed_file:
            intervals = bed_intervals[chrom]
            if not intervals:
                continue
        else:
            intervals = [(0, chrom_lengths[chrom])]
        tasks.append((chrom, intervals))

    def process_chrom(chrom, intervals, bigwig_paths):
        bws = [pyBigWig.open(p) for p in bigwig_paths]
        chrom_data = [[] for _ in bigwig_paths]

        for (start, end) in intervals:
            region_len = end - start
            if region_len <= 0:
                continue
            bin_count = math.ceil(region_len / args.binsize)
            for i, bw in enumerate(bws):
                try:
                    vals = bw.stats(chrom, start, end, nBins=bin_count, type='mean')
                except:
                    vals = [None]*bin_count
                if vals is None:
                    vals = [None]*bin_count
                chrom_data[i].extend([
                    0.0 if (v is None or math.isnan(v)) else v 
                    for v in vals
                ])
        for bw in bws:
            bw.close()
        return chrom_data

    print(f"[INFO] Extracting binned values from {len(args.bigwigs)} BigWigs.")
    n_jobs = args.threads if args.threads != -1 else -1
    results = Parallel(n_jobs=n_jobs, prefer='threads')(
        delayed(process_chrom)(chrom, intervals, args.bigwigs) 
        for chrom, intervals in tasks
    )

    track_values = [[] for _ in args.bigwigs]
    for chrom_data in results:
        for i, vals in enumerate(chrom_data):
            track_values[i].extend(vals)

    track_values = [np.array(vals, dtype=float) for vals in track_values]
    num_bins = len(track_values[0]) if track_values else 0
    if num_bins == 0:
        print("[ERROR] No bins left after filtering.")
        return
    print(f"[INFO] Collected values for {num_bins} bins.")

    # Zero-coverage & outlier trimming
    exclude_zeros = not args.include_zeros
    trim_frac = args.trim_percent / 100.0
    mask = np.ones(num_bins, dtype=bool)

    if exclude_zeros:
        zero_mask = np.zeros(num_bins, dtype=bool)
        for arr in track_values:
            zero_mask |= (arr == 0)
        mask &= ~zero_mask

    if trim_frac > 0.0:
        for arr in track_values:
            arr_sub = arr[arr != 0] if exclude_zeros else arr
            if arr_sub.size == 0:
                continue
            low_val = np.quantile(arr_sub, trim_frac)
            high_val = np.quantile(arr_sub, 1 - trim_frac)
            mask &= (arr >= low_val) & (arr <= high_val)

    if not mask.all():
        filtered_out = np.count_nonzero(~mask)
        print(f"[INFO] Filtering out {filtered_out} bins of {num_bins}.")
        track_values = [arr[mask] for arr in track_values]
        num_bins = len(track_values[0])
    else:
        print("[INFO] No bins removed by zero/outlier filters.")

    # Normalization
    def normalize_array(arr, method):
        if method == 'none':
            return arr.copy()
        elif method == 'mean':
            mean_val = arr.mean()
            return arr if mean_val == 0 else arr / mean_val
        elif method == 'zscore':
            mean_val = arr.mean()
            std_val = arr.std(ddof=0)
            return np.zeros_like(arr) if std_val == 0 else (arr - mean_val) / std_val
        else:
            return arr.copy()

    # If user provided aliases and length matches, use them; else fallback to basenames
    use_aliases = (len(args.aliases) == len(args.bigwigs))
    if not use_aliases and args.aliases:
        print("[WARN] Number of aliases != number of BigWigs; ignoring aliases.")

    normalized_values = [normalize_array(a, args.norm_method) for a in track_values]
    if use_aliases:
        track_names = args.aliases
    else:
        # Fallback: use basenames (and suffix duplicates if any)
        name_counts = {}
        track_names = []
        for bw_path in args.bigwigs:
            base = os.path.basename(bw_path)
            if base not in name_counts:
                name_counts[base] = 1
                track_names.append(base)
            else:
                name_counts[base] += 1
                track_names.append(f"{base}_#{name_counts[base]}")

    data_df = pd.DataFrame({tn: normalized_values[i] for i, tn in enumerate(track_names)})

    # Correlations
    corr_matrices = {}
    if 'pearson' in args.metrics:
        corr_matrices['pearson'] = data_df.corr(method='pearson').values
    if 'spearman' in args.metrics:
        corr_matrices['spearman'] = data_df.corr(method='spearman').values
    if 'kendall' in args.metrics:
        corr_matrices['kendall'] = data_df.corr(method='kendall').values

    # Save correlation in metadata
    metadata['track_names'] = track_names
    for metric, matrix in corr_matrices.items():
        metadata[f'{metric}_correlation_matrix'] = matrix.tolist()

    # Write metadata to JSON
    meta_path = os.path.join(outdir, 'metadata.json')
    with open(meta_path, 'w') as mf:
        json.dump(metadata, mf, indent=4)
    print(f"[INFO] Wrote metadata to {meta_path}")

    # Aesthetic setup: dynamic font scale
    sns.set_style('ticks')
    sns.set_context('talk', font_scale=args.font_scale)

    # Pairwise plot
    print("[INFO] Generating pairwise plot...")
    pairplot_path = os.path.join(outdir, 'pairplot.pdf')
    plot_type = args.plot_type.lower()

    if plot_type == 'scatter':
        # Use scatter for off-diagonal
        plot_kws = {}
        if num_bins > 50000:
            plot_kws['s'] = 10
            plot_kws['alpha'] = 0.3
            plot_kws['rasterized'] = True
        g = sns.pairplot(
            data_df, kind='scatter', diag_kind='hist',
            plot_kws=plot_kws, diag_kws={'bins': 50, 'color': 'gray'}
        )
        plt.suptitle('Pairwise Scatter Plot', y=1.02)

    elif plot_type == 'kde':
        # Custom approach to get "spraypaint" 2D shading
        g = sns.PairGrid(data_df)
        # Diagonal: 1D KDE with fill
        g.map_diag(sns.kdeplot, fill=True, color='gray')
        # Off-diagonal: 2D KDE with fill
        def bivariate_kde(x, y, **kwargs):
            sns.kdeplot(
                x=x, y=y, fill=True, cmap='Blues', 
                levels=15, thresh=0, alpha=0.8, ax=plt.gca()
            )
        g.map_offdiag(bivariate_kde)
        plt.suptitle('Pairwise KDE Density Plot', y=1.02)

    else:  # 'hex'
        g = sns.PairGrid(data_df)
        def hexbin_func(x, y, color, **kwargs):
            ax = plt.gca()
            cmap = plt.cm.get_cmap('Blues')
            ax.hexbin(x, y, gridsize=50, cmap=cmap, mincnt=1, bins='log', rasterized=True)
        g.map_diag(sns.histplot, color='gray', bins=50)
        g.map_offdiag(hexbin_func)
        plt.suptitle('Pairwise Hexbin Density Plot', y=1.02)

    g.fig.savefig(pairplot_path, format='pdf', bbox_inches='tight')
    plt.close(g.fig)
    print(f"[INFO] Saved pairwise plot as {pairplot_path}")

    # Correlation heatmaps
    print("[INFO] Generating correlation matrix heatmaps...")
    for metric, matrix in corr_matrices.items():
        heatmap_path = os.path.join(outdir, f'correlation_matrix_{metric}.pdf')
        plt.figure(figsize=(6, 6))
        ax = sns.heatmap(
            matrix, xticklabels=track_names, yticklabels=track_names,
            annot=True, fmt='.2f', square=True, cmap='coolwarm', 
            vmin=-1, vmax=1
        )
        plt.xticks(rotation=90)
        plt.yticks(rotation=0)
        plt.title(f'{metric.capitalize()} Correlation')
        plt.tight_layout()
        plt.savefig(heatmap_path, format='pdf')
        plt.close()
        print(f"[INFO] Saved {metric} heatmap to {heatmap_path}")

    print("[INFO] Analysis complete. Outputs in:", outdir)

if __name__ == '__main__':
    main()
```
