## Supplementary material for "CAD-C: An engineered nuclease enables repair-free *in situ* proximity ligation and nucleosome-resolution chromosome walks in human cells": Custom code: GCbal_bam_frag_and_upstream_bases_v1_20250328.py

```
#!/usr/bin/env python3
"""
GC-Balanced Sampling from Coordinate-Sorted, Indexed BAM
(single-end or paired-end), extracting upstream & fragment sequences
====================================================================

What it does:
-------------
1) Verifies the BAM is coordinate-sorted & has an accessible index.
2) Computes GC bins on a specified chromosome (default: chr1).
3) Iterates reads in that chromosome:
   - Single-end:
       * Fragment length = read.query_length
       * Lowest coord = read.reference_start
   - Paired-end:
       * Must be properly paired & mapped
       * Fragment length = abs(read.template_length)
       * Lowest coord = min(mate1.start, mate2.start)
         (we detect mate info from the read if it's read1, or handle read2 analogously)
4) Skips fragments if length < flank or out-of-bounds.
5) GC-bin reservoir sampling ~n_sample across 100 GC quantiles.
6) Extracts:
   - upstream: [lowestCoord - flank : lowestCoord)
   - fragment: [lowestCoord : lowestCoord + flank)
   => concatenates => 2*flank bp
7) Outputs:
   - A gzipped TSV with read metadata + sequence
   - A nucleotide_frequency.tsv, labeling positions -flank..-1 and +1..+flank
   - A PDF stacked bar of composition (fonttype=42)
   - A summary JSON

Author: Your Name
Date: 2025-xx-xx
"""

import argparse
import os
import sys
import gzip
import random
import math
import json
from datetime import datetime
import logging

import pysam
import numpy as np
import pandas as pd
from pyfaidx import Fasta
import matplotlib
matplotlib.use('Agg')  # no X11 display needed
matplotlib.rcParams['pdf.fonttype'] = 42
import matplotlib.pyplot as plt

#############################################################
# LOGGING SETUP
#############################################################
# We'll log info & above to stdout with timestamps. SLURM captures stdout in the .out file.
logging.basicConfig(
    stream=sys.stdout,
    level=logging.INFO,
    format='%(asctime)s [%(levelname)s] %(message)s',
    datefmt='%Y-%m-%d %H:%M:%S'
)

#############################################################
# 1) GC QUANTILE CALCULATION
#############################################################

def compute_gc_quantiles(fasta, chrom, bin_size, n_quantiles=100):
    logging.info(f"Computing GC quantiles on {chrom} with bin_size={bin_size} ...")
    seq = fasta[chrom][:].seq
    seq_length = len(seq)
    seq_array = np.frombuffer(seq.encode('ascii'), dtype=np.uint8)
    gc_mask = ((seq_array == ord('G')) | (seq_array == ord('C')) |
               (seq_array == ord('g')) | (seq_array == ord('c')))

    full_bins = seq_length // bin_size
    remainder = seq_length % bin_size

    if full_bins > 0:
        trimmed_length = full_bins * bin_size
        gc_mask_full = gc_mask[:trimmed_length]
        gc_counts_full = gc_mask_full.reshape(full_bins, bin_size).sum(axis=1)
    else:
        gc_counts_full = np.array([], dtype=int)

    if remainder > 0:
        last_bin_start = full_bins * bin_size
        gc_count_last = gc_mask[last_bin_start:].sum()
        gc_counts = np.concatenate([gc_counts_full, [gc_count_last]])
        bin_lengths = np.concatenate([
            np.full(full_bins, bin_size),
            [remainder]
        ])
    else:
        gc_counts = gc_counts_full
        bin_lengths = (np.full(full_bins, bin_size) if full_bins>0 else np.array([], dtype=int))

    gc_percents = gc_counts / bin_lengths * 100.0

    try:
        quantile_labels = list(range(1, n_quantiles+1))
        quantiles = pd.qcut(gc_percents, q=n_quantiles, labels=quantile_labels)
    except Exception as e:
        logging.warning(f"pd.qcut failed: {e}. Falling back to manual grouping.")
        sorted_idx = np.argsort(gc_percents)
        quantiles = np.empty_like(gc_percents, dtype=int)
        group_size = math.ceil(len(gc_percents)/n_quantiles) if len(gc_percents)>0 else 0
        for qi in range(n_quantiles):
            start_i = qi*group_size
            end_i   = min((qi+1)*group_size, len(sorted_idx))
            idx_grp = sorted_idx[start_i:end_i]
            quantiles[idx_grp] = qi+1

    logging.info("Finished computing GC bin assignments.")
    return gc_percents, quantiles

#############################################################
# 2) OPEN & CHECK BAM
#############################################################

def open_and_check_bam(bam_path):
    """
    1) Ensure we can open the file in read-binary mode.
    2) Check that it has a coordinate-sorted header: HD: SO=coordinate
    3) Check that a .bai (or CSI) index is present & usable.
    4) Return the pysam.AlignmentFile object if all checks pass.
    """
    if not os.path.exists(bam_path):
        raise FileNotFoundError(f"BAM file '{bam_path}' not found.")

    logging.info(f"Opening BAM: {bam_path} ...")
    sam = pysam.AlignmentFile(bam_path, 'rb')

    # 2) Check sorted
    hd = sam.header.get('HD', {})
    sort_order = hd.get('SO', None)
    if sort_order != 'coordinate':
        sam.close()
        raise ValueError(f"BAM '{bam_path}' is not coordinate-sorted (SO={sort_order}).")

    # 3) Check index availability
    # We'll do a small fetch test: if there's no index, we get an error.
    try:
        # Try reading a small region from the first reference. If no index, this fails.
        first_ref = sam.get_reference_name(0) if sam.nreferences > 0 else None
        if first_ref:
            _ = list(sam.fetch(first_ref, 0, 10))  # read an arbitrary small region
    except ValueError as e:
        sam.close()
        raise ValueError(f"No valid BAM index found for '{bam_path}'. Error: {e}")

    logging.info("BAM is coordinate-sorted and index is accessible.")
    return sam

#############################################################
# 3) SINGLE-PASS RESERVOIR SAMPLING FOR GC BINS
#############################################################

def sample_bam_entries_by_quantile(
    bam_path, chrom, quantile_array, bin_size, n_sample, flank, debug=False
):
    """
    Iterate reads in 'chrom' via sam.fetch(chrom).
    - If single-end: we treat read.query_length as fragment len,
      lowestCoord = read.reference_start.
    - If paired-end: check read.is_paired, read.is_proper_pair, read.is_read1, etc.
      * fragment_len = abs(read.template_length)
      * lowestCoord = min( startOfThisRead, next_reference_start ), skip if mate unmapped
    - skip if fragment_len < flank
    - reservoir-sample into GC bin using lowestCoord.

    Return the reservoir of selected records (with minimal metadata),
    plus total_chrom_count, selected_count.
    """
    from collections import defaultdict

    quantile_size = 100
    reservoirs = {q: [] for q in range(1, quantile_size+1)}
    counts = defaultdict(int)

    target_per_q = n_sample // quantile_size
    remainder = n_sample % quantile_size
    target_counts = {q: target_per_q for q in range(1, quantile_size+1)}
    q_i = 1
    while remainder>0 and q_i<=quantile_size:
        target_counts[q_i]+=1
        remainder-=1
        q_i+=1

    total_pairs_chrom = 0
    selected_total = 0

    sam = open_and_check_bam(bam_path)

    logging.info(f"Scanning reads on chromosome {chrom} ...")
    for read in sam.fetch(chrom):
        if read.is_secondary or read.is_supplementary:
            continue
        if read.is_unmapped:
            continue

        # Distinguish single-end vs. paired-end:
        if not read.is_paired:
            # single-end logic:
            frag_len = read.query_length
            if frag_len < flank:
                continue
            # lowestCoord
            lowest = read.reference_start  # leftmost aligned base
        else:
            # paired-end logic
            # only process read1 to avoid duplicates
            if not read.is_read1:
                continue
            if not read.is_proper_pair:
                continue
            mate_unmapped = read.mate_is_unmapped
            if mate_unmapped:
                continue

            start1 = read.reference_start
            start2 = read.next_reference_start
            lowest = start1 if (start1 < start2) else start2
            frag_len = abs(read.template_length)
            if frag_len < flank:
                continue

        # compute bin
        bin_idx = (lowest - 1)//bin_size
        if bin_idx<0 or bin_idx>=len(quantile_array):
            continue
        q_label = int(quantile_array[bin_idx])

        total_pairs_chrom += 1
        counts[q_label]+=1

        # minimal metadata
        # name, isPaired, isRead1, chrom, lowestCoord, frag_len
        # (pos1 or reference_start if single-end?),  plus isReverse, next_reference_start, etc. if we want
        rec_info = [
            read.query_name or "unknown",
            read.is_paired,
            read.is_read1,
            chrom,
            lowest,
            frag_len,
            read.reference_start,
            read.next_reference_start if read.is_paired else -1,
            read.is_reverse,
            read.mate_is_reverse if read.is_paired else False
        ]

        # reservoir sampling
        if len(reservoirs[q_label]) < target_counts[q_label]:
            reservoirs[q_label].append(rec_info)
            selected_total+=1
        else:
            r = random.randint(0, counts[q_label]-1)
            if r < target_counts[q_label]:
                reservoirs[q_label][r] = rec_info

    sam.close()
    logging.info(f"Done scanning. total_chrom={total_pairs_chrom}, selected_total={selected_total}")

    # combine & possibly truncate
    selected_records = []
    for q in range(1, quantile_size+1):
        selected_records.extend(reservoirs[q])
    if len(selected_records)>n_sample:
        selected_records = selected_records[:n_sample]
        selected_total = n_sample
    else:
        selected_total = len(selected_records)

    random.shuffle(selected_records)
    return selected_records, total_pairs_chrom, selected_total

#############################################################
# 4) EXTRACT UPSTREAM + FRAGMENT SEQ
#############################################################

def extract_2x_flank_sequence(rec_info, fasta, flank, debug=False):
    """
    rec_info:
      [
        read_name, isPaired, isRead1,
        chrom, lowest, frag_len,
        start1, start2, isReverse1, isReverse2
      ]
    We want [lowest-flank, lowest) + [lowest, lowest+flank)
    skip if out of bounds
    """
    (rname, isPaired, isRead1, chrom, lowestCoord, frag_len,
     start1, start2, isRev1, isRev2) = rec_info

    if lowestCoord < flank:
        # near chromosome start
        return ""

    chrom_len = len(fasta[chrom])
    if (lowestCoord + flank) > chrom_len:
        return ""

    # upstream
    startA = lowestCoord - flank
    endA   = lowestCoord
    seqA = fasta[chrom][startA:endA].seq

    # fragment portion
    startB = lowestCoord
    endB   = lowestCoord + flank
    seqB = fasta[chrom][startB:endB].seq

    combined = seqA + seqB
    if len(combined) != 2*flank:
        return ""
    return combined

#############################################################
# 5) MAIN
#############################################################

def main():
    parser = argparse.ArgumentParser(
        description="GC-balanced sampling from a coordinate-sorted, indexed BAM, extracting upstream + fragment sequences."
    )
    parser.add_argument("-b","--bam", required=True, help="Coordinate-sorted, indexed BAM file.")
    parser.add_argument("-f","--fasta", required=True, help="Reference genome FASTA (indexed).")
    parser.add_argument("-o","--output-dir", help="Output directory. Timestamp appended if not provided.")
    parser.add_argument("--chrom", default="chr1", help="Chromosome to compute GC quantiles on (default=chr1).")
    parser.add_argument("--bin-size", type=int, default=1000, help="Bin size for GC. (default=1000)")
    parser.add_argument("--flank", type=int, default=50, help="Flank length. (default=50)")
    parser.add_argument("-n","--num-junctions", type=int, default=1000000,
                        help="Number of fragments to sample (default=1,000,000).")
    parser.add_argument("--seed", type=int, help="Random seed for reproducibility.")
    parser.add_argument("--debug", action="store_true", help="Enable debug messages.")
    args = parser.parse_args()

    if args.seed is not None:
        random.seed(args.seed)

    # Maybe set logging to DEBUG if --debug
    if args.debug:
        logging.getLogger().setLevel(logging.DEBUG)

    bam_path = args.bam
    fasta_path = args.fasta
    out_dir = args.output_dir
    chrom = args.chrom
    bin_size = args.bin_size
    flank = args.flank
    n_sample = args.num_junctions

    timestamp = datetime.now().strftime("%Y%m%d_%H%M%S")

    # Derive stub from BAM name
    bam_name = os.path.basename(bam_path)
    if bam_name.endswith(".bam"):
        bam_name = bam_name[:-4]

    if not out_dir:
        out_dir = f"gc_bal_{bam_name}_{timestamp}"
    else:
        if not out_dir.endswith(timestamp):
            out_dir = f"{out_dir}_{bam_name}_{timestamp}"
    os.makedirs(out_dir, exist_ok=True)

    logging.info(f"Loading reference FASTA: {fasta_path}")
    fasta = Fasta(fasta_path, sequence_always_upper=True)

    # 1) compute GC quantiles
    gc_percents, quantiles = compute_gc_quantiles(fasta, chrom, bin_size, 100)
    quant_file = os.path.join(out_dir, f"{chrom}_gc_bins_{bam_name}_{timestamp}.tsv")
    logging.info(f"Writing GC bin file to {quant_file}")
    with open(quant_file,"w") as qf:
        qf.write("bin_start\tbin_end\tGC_percent\tquantile\n")
        for i,(gcval, qval) in enumerate(zip(gc_percents, quantiles)):
            start_bin = i*bin_size+1
            end_bin   = min((i+1)*bin_size, len(fasta[chrom]))
            qf.write(f"{start_bin}\t{end_bin}\t{gcval:.2f}\t{int(qval)}\n")

    # 2) sample from BAM
    logging.info("Starting GC-balanced sampling from BAM.")
    selected_records, total_chrom, selected_total = sample_bam_entries_by_quantile(
        bam_path, chrom, quantiles, bin_size, n_sample, flank, debug=args.debug
    )

    # Prepare composition arrays
    seq_length = 2*flank
    base_to_idx = [-1]*128
    for b, idx in zip("ACGT",[0,1,2,3]):
        base_to_idx[ord(b)] = idx
        base_to_idx[ord(b.lower())] = idx

    freq = [[0]*seq_length for _ in range(4)]

    # 3) Write out a gz TSV
    seq_tsv_path = os.path.join(out_dir, f"fragments_{bam_name}_{timestamp}.tsv.gz")
    logging.info(f"Writing final fragment sequences to {seq_tsv_path}")
    gz_out = gzip.open(seq_tsv_path,"wt")
    # We'll define columns for the metadata + final sequence
    header_cols = [
        "read_name","is_paired","is_read1","chrom","lowestCoord","frag_len",
        "read_start1","read_start2","is_rev1","is_rev2","sequence"
    ]
    gz_out.write("\t".join(header_cols)+"\n")

    skipped_bounds = 0
    for rec_info in selected_records:
        seq = extract_2x_flank_sequence(rec_info, fasta, flank, debug=args.debug)
        if not seq:
            skipped_bounds +=1
            continue

        # Update freq
        for i,ch in enumerate(seq):
            code = ord(ch)
            if code<128:
                idx = base_to_idx[code]
                if idx!=-1:
                    freq[idx][i]+=1

        # rec_info => [rname, isPaired, isRead1, chrom, lowest, frag_len, start1, start2, isRev1, isRev2]
        # we add final 'sequence'
        row = rec_info + [seq]
        row_str = "\t".join(map(str,row))
        gz_out.write(row_str+"\n")

    gz_out.close()
    output_count = selected_total - skipped_bounds

    # 4) Write composition table
    freq_file = os.path.join(out_dir, f"nucleotide_frequency_{bam_name}_{timestamp}.tsv")
    logging.info(f"Writing composition table to {freq_file}")
    with open(freq_file,"w") as ff:
        ff.write("position\tA\tC\tG\tT\n")
        # left half => -flank..-1
        for i in range(flank):
            pos_label = i - flank
            ff.write(f"{pos_label}\t{freq[0][i]}\t{freq[1][i]}\t{freq[2][i]}\t{freq[3][i]}\n")
        # right half => +1..+flank
        for i in range(flank, seq_length):
            pos_label = i - flank + 1
            ff.write(f"{pos_label}\t{freq[0][i]}\t{freq[1][i]}\t{freq[2][i]}\t{freq[3][i]}\n")

    # 5) Make a stacked bar PDF
    comp_pdf_path = os.path.join(out_dir, f"base_composition_{bam_name}_{timestamp}.pdf")
    total_seqs = output_count if output_count>0 else 1
    freq_array = np.array(freq)
    fractions = freq_array / total_seqs

    x_positions = list(range(-flank, 0)) + list(range(1, flank+1))

    logging.info(f"Creating PDF plot {comp_pdf_path}")
    plt.figure(figsize=(12,6))
    bottom = np.zeros(seq_length)
    colors = {'A':'green','T':'red','G':'orange','C':'blue'}
    base_order = ['A','G','C','T']

    for base in base_order:
        idx = {'A':0,'C':1,'G':2,'T':3}[base]
        plt.bar(x_positions, fractions[idx], bottom=bottom,
                color=colors[base], width=1.0, label=base)
        bottom += fractions[idx]

    plt.axvline(x=0, color='black', linewidth=1)
    plt.xlabel("Position relative to fragment's lowest coord (cut site)")
    plt.ylabel("Fraction of base")
    plt.title(f"Upstream + Fragment Composition for '{bam_name}'")
    plt.legend(title="Base", loc="upper right")
    plt.tight_layout()
    plt.savefig(comp_pdf_path)
    plt.close()

    # 6) Summary JSON
    summary = {
        "bam_file": os.path.basename(args.bam),
        "bam_stub": bam_name,
        "fasta_file": os.path.basename(args.fasta),
        "chromosome": chrom,
        "bin_size": bin_size,
        "num_quantiles": 100,
        "flank_length": flank,
        "requested_fragments": n_sample,
        f"total_reads_on_{chrom}": total_chrom,
        "selected_fragments": selected_total,
        "skipped_out_of_bounds": skipped_bounds,
        "final_written_fragments": output_count,
        "timestamp": timestamp,
        "output_files": {
            "fragments_tsv_gz": os.path.basename(seq_tsv_path),
            "nucleotide_frequency_tsv": os.path.basename(freq_file),
            "composition_plot_pdf": os.path.basename(comp_pdf_path),
            "gc_quantiles_tsv": os.path.basename(quant_file)
        }
    }

    summary_path = os.path.join(out_dir, f"summary_{bam_name}_{timestamp}.json")
    logging.info(f"Writing summary JSON to {summary_path}")
    with open(summary_path,"w") as js:
        json.dump(summary, js, indent=4)

    logging.info("All done!")

if __name__=="__main__":
    main()
```
