## Supplementary material for "CAD-C: An engineered nuclease enables repair-free *in situ* proximity ligation and nucleosome-resolution chromosome walks in human cells": Custom code: MACC_K562_Mieczkowski_downsampletomin_FLDreplot_v2_20251113.sh

```
#!/bin/bash
#SBATCH --job-name=GM12878_FLD_hist
#SBATCH --output=GM12878_FLD_hist.%j.out
#SBATCH --error=GM12878_FLD_hist.%j.err
#SBATCH --ntasks=1
#SBATCH --cpus-per-task=8

set -euo pipefail

#############################
# Project-specific paths
#############################
BAM_DIR="/rugpfs/fs0/risc_lab/scratch/jhickling/Microc_benchmark/MACC/fastq2bam/MACC_K562_MNasedigbams/"
OUT_BASE="/rugpfs/fs0/risc_lab/scratch/jsoroczyns/CAD/CAD-C/scripts/2025_11_13/FLD_plots_2025"
mkdir -p "${OUT_BASE}"

# Path to your patched Python script (must be v1.4.0+ with histogram options)
SCRIPT="/rugpfs/fs0/risc_lab/scratch/jsoroczyns/CAD/CAD-C/scripts/2025_09_27/bam_insert_size_fld_v2_20250927.py"

#############################
# Environment
#############################
# Adjust to your cluster setup
# source ~/.bashrc 2>/dev/null || true
# conda activate open2c_v3

THREADS="${SLURM_CPUS_PER_TASK:-8}"

#############################
# Run (autosomes-only by default)
# - Plot x-axis: 30..250 bp
# - Histograms also 30..250 bp
# - Downsample all BAMs to the shallowest input
# - Save ONLY compact histogram TSVs (gzip) — no .npy arrays
# - Font size 20; lines 2× thicker (3.6)
#############################
python "${SCRIPT}" \
  --bams \
    "${BAM_DIR}/SRR3211679_S1_001.trim.st.all.blft.qft.rmdup.bam" \
    "${BAM_DIR}/SRR3211680_S1_001.trim.st.all.blft.qft.rmdup.bam" \
    "${BAM_DIR}/SRR3211681_S2_001.trim.st.all.blft.qft.rmdup.bam" \
    "${BAM_DIR}/SRR3211682_S3_001.trim.st.all.blft.qft.rmdup.bam" \
  --labels "5.4U MNase,20.6U MNase, 79.2U MNase, 304U MNase" \
  --units 1,2,3,4\
  --plot-min-frag 30 --plot-max-frag 250 \
  --hist-min-frag 30 --hist-max-frag 250 \
  --save-hist-tsv --no-raw-arrays \
  --font-size 20 --line-width 3.6 \
  --downsample-to-min \
  --threads "${THREADS}" \
  --outdir "${OUT_BASE}"

# Optional: to include chrX/chrY, add: --include-sex-chroms
# Optional: to change stats band, add e.g.: --stat-min-frag 90 --stat-max-frag 250
# Optional: to export mitochondrial alignments, add: --export-chrM-bam --mt-contigs chrM,MT
```
