## Supplementary material for "CAD-C: An engineered nuclease enables repair-free *in situ* proximity ligation and nucleosome-resolution chromosome walks in human cells": Custom code: smart_replot_GCcov_tsvs_v1_20250330.py

```
#!/usr/bin/env python3
"""
Plot GC bias from multiple groups of coverage data, using pre-normalized coverage values.
We directly compute log2(cov) from the provided cov_median, cov_Q1, cov_Q3 (no extra normalization).

Each group:
  - Has one or more TSV files with these columns:
      gc_mean, cov_median, cov_Q1, cov_Q3
  - Possibly a label for each file (or auto-derived from filename).
  - A Seaborn color palette (light→dark).

Options:
  - Single combined panel (all groups on one plot) vs. multiple subplots (one group/panel).
  - Y-scale: 'log2' or 'linear' (the data is still log2, but 'linear' changes axis labeling).
  - Automatic or manual y-limits; optional symmetry around 0 (useful if data is truly log2).
  - Writes a PDF with timestamp in the filename + a metadata file.

Example (two groups, two files each):

  python plot_gc_bias_multi_groups.py \
    --num-groups 2 \
    --group1-files mnase_5.tsv mnase_20.tsv \
    --group1-labels "5 U" "20 U" \
    --palette-group1 "rocket_r" \
    --group2-files chomp_1.tsv chomp_10.tsv \
    --group2-labels "1 µL" "10 µL" \
    --palette-group2 "mako_r" \
    --subplot-mode separate \
    --y-scale log2 \
    --symmetrical \
    --ymin-auto --ymax-auto \
    --output GC_bias_multi_panel.pdf

Author: Your Name
Date: 2025-03-30
"""

import argparse
import os
import sys
import pandas as pd
import numpy as np
import matplotlib
matplotlib.use('Agg')  # For headless environments (clusters)
import matplotlib.pyplot as plt
import seaborn as sns
from datetime import datetime


###############################################################################
# 1) LOADING & LOG2 TRANSFORMATION (NO ADDITIONAL NORMALIZATION)
###############################################################################

def load_and_log2_transform(file_path, label=None):
    """
    Reads a TSV with columns [gc_mean, cov_median, cov_Q1, cov_Q3].
    - Checks required columns.
    - log2-transform each coverage column (clip at 1e-12 to avoid log2(<=0) issues).
    - Attach 'label' (derived from filename if not provided).

    Returns a DataFrame with columns:
      gc_mean, log2_cov_median, log2_cov_Q1, log2_cov_Q3, label
    """
    df = pd.read_csv(file_path, sep='\t')
    required_cols = ['gc_mean', 'cov_median', 'cov_Q1', 'cov_Q3']
    for c in required_cols:
        if c not in df.columns:
            raise ValueError(f"[ERROR] {file_path} is missing required column '{c}'.")

    # Auto-label from filename if user didn't provide one
    if label is None:
        label = os.path.splitext(os.path.basename(file_path))[0]

    # Clip coverage to avoid log2(<= 0)
    df['cov_median'] = df['cov_median'].clip(lower=1e-12)
    df['cov_Q1']     = df['cov_Q1'].clip(lower=1e-12)
    df['cov_Q3']     = df['cov_Q3'].clip(lower=1e-12)

    # Direct log2 transform (no additional normalization)
    df['log2_cov_median'] = np.log2(df['cov_median'])
    df['log2_cov_Q1']     = np.log2(df['cov_Q1'])
    df['log2_cov_Q3']     = np.log2(df['cov_Q3'])

    df['label'] = label

    return df[['gc_mean', 'log2_cov_median', 'log2_cov_Q1', 'log2_cov_Q3', 'label']]


###############################################################################
# 2) HELPER: COMPUTE GLOBAL Y-LIMITS
###############################################################################

def compute_y_lims(df_list):
    """
    Given a list of DataFrames (each with 'log2_cov_Q1' & 'log2_cov_Q3'),
    return the global min and max across Q1/Q3 of all DataFrames.
    """
    if not df_list:
        # If no data, fallback
        return -1.0, 1.0

    combined_q1 = []
    combined_q3 = []
    for df in df_list:
        combined_q1.extend(df['log2_cov_Q1'].values)
        combined_q3.extend(df['log2_cov_Q3'].values)

    y_min = min(combined_q1 + combined_q3)
    y_max = max(combined_q1 + combined_q3)
    return (y_min, y_max)


###############################################################################
# 3) PLOTTING LOGIC
###############################################################################

def plot_dataframes(ax, df_list, palette, y_scale='log2', xlim=(0,100)):
    """
    Plots multiple lines (each from df_list) on a single Axes 'ax':
      - Each df must have columns: gc_mean, log2_cov_median, log2_cov_Q1, log2_cov_Q3, label
      - 'palette' is a list of colors, length >= len(df_list).
      - 'y_scale' is either 'log2' or 'linear' => how we label the axis
         (the data is always stored in log2 already).
      - 'xlim' default = (0,100), typical GC range.

    For each df:
      - Sort by gc_mean => line from left to right
      - ax.plot(... log2_cov_median ...)
      - ax.fill_between(... log2_cov_Q1..Q3 ...)
    """
    for i, df in enumerate(df_list):
        df_sorted = df.sort_values('gc_mean')
        color = palette[i]
        lbl = df_sorted['label'].iloc[0]

        # Plot the median line
        ax.plot(df_sorted['gc_mean'], df_sorted['log2_cov_median'],
                label=lbl, color=color, linewidth=2.5)

        # Fill the ribbon (IQR)
        ax.fill_between(df_sorted['gc_mean'],
                        df_sorted['log2_cov_Q1'],
                        df_sorted['log2_cov_Q3'],
                        color=color, alpha=0.25)

    ax.axhline(0, linestyle='dotted', color='black', linewidth=1)
    ax.set_xlim(*xlim)

    if y_scale == 'log2':
        ax.set_ylabel("log₂(Coverage)", weight='bold')
    else:
        # The data is still in log2, but user wants a "linear" axis label 
        # => possibly a mismatch. We'll label for clarity:
        ax.set_ylabel("Coverage (log2 data)", weight='bold')


###############################################################################
# 4) MAIN: ARGUMENT PARSING & COORDINATING EVERYTHING
###############################################################################

def main():
    parser = argparse.ArgumentParser(
        description="Plot GC bias from multiple groups of pre-normalized coverage data (direct log2 transform)."
    )

    # Number of groups
    parser.add_argument('--num-groups', type=int, required=True,
                        help='Number of groups to plot (e.g. MNase, CHOMP, etc.).')

    # Subplot mode
    parser.add_argument('--subplot-mode', choices=['combined','separate'], default='combined',
                        help="If 'combined', all groups overlaid in one panel. If 'separate', each group in its own subplot.")

    # Y-scale & Y-limits
    parser.add_argument('--y-scale', choices=['log2','linear'], default='log2',
                        help="How to label/interpret the y-axis. Data is always log2 internally.")
    parser.add_argument('--ymin', type=float, default=-2.0,
                        help="Manual minimum y-limit (if not auto).")
    parser.add_argument('--ymax', type=float, default=2.0,
                        help="Manual maximum y-limit (if not auto).")
    parser.add_argument('--ymin-auto', action='store_true',
                        help="Compute y-min from data.")
    parser.add_argument('--ymax-auto', action='store_true',
                        help="Compute y-max from data.")
    parser.add_argument('--symmetrical', action='store_true',
                        help="Make y-limits symmetrical about 0 if in log2 scale.")

    # Output PDF
    parser.add_argument('--output', default='gc_bias_plot.pdf',
                        help="Base name for output PDF (timestamp appended).")

    args, unknown = parser.parse_known_args()

    # We'll parse group-based arguments from 'unknown' dynamically.
    # For each group i in [1..num_groups], we look for:
    #   --group{i}-files file1 file2 ...
    #   --group{i}-labels label1 label2 ...
    #   --palette-group{i} paletteName
    group_info = []
    for g in range(1, args.num_groups + 1):
        group_id = f"group{g}"

        files_key   = f"--{group_id}-files"
        labels_key  = f"--{group_id}-labels"
        palette_key = f"--palette-{group_id}"

        gf = []
        gl = []
        gp = "rocket_r"  # default palette
        i = 0
        while i < len(unknown):
            item = unknown[i]
            if item == files_key:
                j = i+1
                while j < len(unknown) and not unknown[j].startswith("--"):
                    gf.append(unknown[j])
                    j += 1
                del unknown[i:j]
                continue
            elif item == labels_key:
                j = i+1
                temp_labels = []
                while j < len(unknown) and not unknown[j].startswith("--"):
                    temp_labels.append(unknown[j])
                    j += 1
                del unknown[i:j]
                gl = temp_labels
                continue
            elif item == palette_key:
                if i+1 < len(unknown) and not unknown[i+1].startswith("--"):
                    gp = unknown[i+1]
                    del unknown[i:i+2]
                else:
                    del unknown[i]
                continue
            else:
                i += 1

        group_info.append({
            'name':  group_id,
            'files': gf,
            'labels': gl,
            'palette': gp
        })

    # Validate group data
    for g in group_info:
        if not g['files']:
            print(f"[ERROR] No files provided for {g['name']}.", file=sys.stderr)
            sys.exit(1)
        if len(g['labels']) > len(g['files']):
            print(f"[ERROR] More labels than files for {g['name']}.", file=sys.stderr)
            sys.exit(1)

    # Load data
    all_dfs_per_group = []
    for g in group_info:
        # Possibly fill missing labels with None => auto-inferred
        needed_auto = len(g['files']) - len(g['labels'])
        g['labels'].extend([None]*needed_auto)

        dfs = []
        for (fpath, lbl) in zip(g['files'], g['labels']):
            df = load_and_log2_transform(fpath, lbl)
            dfs.append(df)
        all_dfs_per_group.append(dfs)

    # Flatten all data for global y-limit calculation if needed
    flattened = []
    for dfs in all_dfs_per_group:
        flattened.extend(dfs)

    y_min, y_max = args.ymin, args.ymax
    if args.ymin_auto or args.ymax_auto:
        auto_lo, auto_hi = compute_y_lims(flattened)
        if args.ymin_auto:
            y_min = auto_lo
        if args.ymax_auto:
            y_max = auto_hi

    if args.symmetrical and args.y_scale == 'log2':
        biggest = max(abs(y_min), abs(y_max))
        y_min, y_max = -biggest, biggest

    # Set up plot style
    sns.set_style("whitegrid")
    plt.rcParams['pdf.fonttype'] = 42
    plt.rc('font', size=14)
    plt.rc('axes', titlesize=16, labelsize=14)
    plt.rc('xtick', labelsize=12)
    plt.rc('ytick', labelsize=12)
    plt.rc('legend', fontsize=12)

    # Plot
    num_groups = args.num_groups
    if args.subplot_mode == 'separate':
        fig, axes = plt.subplots(nrows=num_groups, ncols=1,
                                 figsize=(8, 4*num_groups),
                                 sharex=True)
        if num_groups == 1:
            axes = [axes]  # unify into list for single subplot

        for i, group in enumerate(group_info):
            dfs = all_dfs_per_group[i]
            pal = sns.color_palette(group['palette'], n_colors=len(dfs))

            ax = axes[i]
            plot_dataframes(ax, dfs, pal, y_scale=args.y_scale)
            ax.set_ylim(y_min, y_max)
            ax.set_title(f"{group['name']}", weight='bold')
            ax.legend(frameon=False)
            ax.set_xlabel("GC Content (%)", weight='bold')

        plt.tight_layout()

    else:
        # single combined panel
        fig, ax = plt.subplots(figsize=(8,6))
        for i, group in enumerate(group_info):
            dfs = all_dfs_per_group[i]
            pal = sns.color_palette(group['palette'], n_colors=len(dfs))
            plot_dataframes(ax, dfs, pal, y_scale=args.y_scale)
        ax.set_ylim(y_min, y_max)
        ax.legend(frameon=False)
        ax.set_xlabel("GC Content (%)", weight='bold')
        ax.set_title("Combined GC Bias Plot", weight='bold')
        plt.tight_layout()

    # Output with timestamp
    timestamp = datetime.now().strftime("%Y%m%d_%H%M%S")
    base_out, ext_out = os.path.splitext(args.output)
    if not ext_out:
        ext_out = ".pdf"
    final_out = f"{base_out}_{timestamp}{ext_out}"
    plt.savefig(final_out, dpi=300)
    print(f"[INFO] Plot saved to {final_out}")

    # Write a metadata file
    meta_out = f"{base_out}_metadata_{timestamp}.txt"
    with open(meta_out, 'w') as m:
        m.write(f"Timestamp: {timestamp}\n")
        m.write(f"Subplot Mode: {args.subplot_mode}\n")
        m.write(f"Y-scale: {args.y_scale}\n")
        m.write(f"Y-limits: [{y_min}, {y_max}], symmetrical={args.symmetrical}\n")
        m.write(f"Output PDF: {final_out}\n\n")
        m.write("Groups:\n")
        for g, dfs in zip(group_info, all_dfs_per_group):
            m.write(f"  {g['name']} => palette={g['palette']}\n")
            for df, fpath in zip(dfs, g['files']):
                lbl = df['label'].iloc[0]
                m.write(f"    - file={fpath}, label={lbl}\n")

    print(f"[INFO] Metadata written to {meta_out}")


if __name__ == "__main__":
    main()
```
