## Supplementary material for "CAD-C: An engineered nuclease enables repair-free *in situ* proximity ligation and nucleosome-resolution chromosome walks in human cells": Custom code: GC_bal_fragment1_only_v3_wmetadata_20250804_fixed.py

```
#!/usr/bin/env python3
###############################################################################
# Script:        GC_bal_fragment1_only_v3_wmetadata_20250804_fixed.py
# Author:        Jan Soroczynski, Risca Lab, The Rockefeller University
# Date:          2025_08_04
#
# Purpose
# -------
# Sample Hi‑C/Micro‑C junctions from a *.pairs(.gz)* file in a GC‑balanced
# manner and extract **exactly the first nucleotide at the ligation junction
# plus (flank‑1) downstream bases of Fragment 1**.  Coordinates are handled
# according to the pairtools specification:
#
#   * `pos1` is **1‑based inclusive** and already marks the junction base when
#     files were generated with `pairtools parse2 --report-position junction`.
#   * For `strand1 '+'` the slice therefore starts at `pos1‑1` (0‑based) and
#     spans `flank` nucleotides downstream.
#   * For `strand1 '-'` the slice is `[pos1‑flank, pos1)` in 1‑based space,
#     which converts to `[pos1‑flank‑1, pos1‑1]` 0‑based; the result is
#     reverse‑complemented so the output is oriented 5′→3′ away from the
#     junction, matching the ‘+’‑strand case.
#
# All other functionality (GC‑quantile reservoir sampling, frequency matrix,
# plots, summary JSON, timestamped filenames) is unchanged from v3 (2025‑03‑29).
#
# Inputs
# ------
#   • *.pairs(.gz)* generated with `--report-position junction`
#   • Indexed reference FASTA (pyfaidx compatible)
#
# Outputs
# -------
#   • junction_sequences_<stub>_<file>_<timestamp>.tsv.gz
#   • <chrom>_gc_bins_quantiles_<stub>_<file>_<timestamp>.tsv
#   • nucleotide_frequency_<stub>_<file>_<timestamp>.tsv
#   • base_composition_<stub>_<file>_<timestamp>.pdf
#   • summary_<stub>_<file>_<timestamp>.json
#
# Example
# -------
#   python GC_bal_fragment1_only_v3_wmetadata_20250804_fixed.py \
#       -i sample.pairs.gz -f hg38.fa -o ./outdir -n 1000000 --chrom chr1
#
# CHANGELOG (2025‑08‑04)
# ----------------------
#   • **Fixed off‑by‑one error** in `extract_fragment1_strand`: coordinates are
#     now converted from 1‑based inclusive to 0‑based half‑open before slicing.
#   • Added detailed README header and maintained full original logic.
###############################################################################

import argparse
import os
import gzip
import random
import math
import json
from datetime import datetime
import pandas as pd
import numpy as np
from pyfaidx import Fasta

import matplotlib
matplotlib.use('Agg')
matplotlib.rcParams['pdf.fonttype'] = 42
import matplotlib.pyplot as plt


# --------------------------- GC‑quantile utilities --------------------------- #

def compute_gc_quantiles(fasta, chrom, bin_size, n_quantiles):
    seq = fasta[chrom][:].seq
    seq_length = len(seq)
    seq_array = np.frombuffer(seq.encode('ascii'), dtype=np.uint8)
    gc_mask = ((seq_array == ord('G')) | (seq_array == ord('C')) |
               (seq_array == ord('g')) | (seq_array == ord('c')))

    full_bins = seq_length // bin_size
    remainder = seq_length % bin_size

    if full_bins > 0:
        trimmed_length = full_bins * bin_size
        gc_counts_full = gc_mask[:trimmed_length]\
            .reshape(full_bins, bin_size).sum(axis=1)
    else:
        gc_counts_full = np.array([], dtype=int)

    if remainder > 0:
        gc_count_last = gc_mask[-remainder:].sum()
        gc_counts = np.concatenate([gc_counts_full, [gc_count_last]])
        bin_lengths = np.concatenate(
            [np.full(full_bins, bin_size), [remainder]]
        )
    else:
        gc_counts = gc_counts_full
        bin_lengths = np.full(full_bins, bin_size) if full_bins else \
            np.array([], dtype=int)

    gc_percents = gc_counts / bin_lengths * 100.0

    try:
        labels = list(range(1, n_quantiles + 1))
        quantiles = pd.qcut(gc_percents, q=n_quantiles, labels=labels)
    except Exception:  # fallback if ties prevent qcut
        sorted_idx = np.argsort(gc_percents)
        quantiles = np.empty_like(gc_percents, dtype=int)
        group = math.ceil(len(gc_percents) / n_quantiles) if len(gc_percents)\
            else 0
        for qi in range(n_quantiles):
            idx = sorted_idx[qi*group: (qi+1)*group]
            quantiles[idx] = qi + 1
    return gc_percents, quantiles


# ------------------------ Reservoir‑sampling by GC bin ----------------------- #

def sample_junctions_by_quantile(pairs_path, chrom, quantile_array, bin_size,
                                 n_sample, debug=False):
    quantile_size = 100
    reservoirs = {q: [] for q in range(1, quantile_size + 1)}
    counts = {q: 0 for q in range(1, quantile_size + 1)}

    tgt_base = n_sample // quantile_size
    remainder = n_sample % quantile_size
    targets = {q: tgt_base + (1 if q <= remainder else 0)
               for q in range(1, quantile_size + 1)}

    total_selected = 0
    total_chrom = 0
    offset = None
    opener = gzip.open if pairs_path.endswith('.gz') else open

    with opener(pairs_path, 'rt') as fh:
        first = None
        for raw in fh:
            if not raw.startswith('#'):
                first = raw
                break
        if first is None:
            return [], 0, 0, 0

        off = 0 if first.split()[0].startswith('chr') or \
                 first.split()[0].isdigit() else 1
        offset = off

        def handle(line):
            nonlocal total_selected, total_chrom
            cols = line.rstrip().split()
            if cols[0+off] != chrom:
                return
            pos1 = int(cols[1+off])
            bin_idx = (pos1 - 1) // bin_size
            if not (0 <= bin_idx < len(quantile_array)):
                return
            qlab = int(quantile_array[bin_idx])
            total_chrom += 1
            counts[qlab] += 1
            if len(reservoirs[qlab]) < targets[qlab]:
                reservoirs[qlab].append(cols)
                total_selected += 1
            else:
                r = random.randint(0, counts[qlab]-1)
                if r < targets[qlab]:
                    reservoirs[qlab][r] = cols

        handle(first)
        for ln in fh:
            handle(ln)

    selected = [item for q in reservoirs for item in reservoirs[q]]
    random.shuffle(selected)
    return selected[:n_sample], total_chrom, min(len(selected), n_sample), offset


# ------------------ Junction‑flank extraction (FIXED INDEX) ------------------ #

def extract_fragment1_strand(cols, offset, fasta, flank, debug=False):
    """
    Return the 5′→3′ sequence of length *flank* starting at the ligation
    junction nucleotide of Fragment 1.

    For '+' : slice = [pos1‑1, pos1‑1+flank)  (0‑based half‑open)
    For '-' : slice = [pos1‑flank, pos1) in 1‑based →
               [pos1‑flank‑1, pos1‑1] 0‑based, then reverse‑complement.
    """
    chrom = cols[0 + offset]
    pos1_1b = int(cols[1 + offset])          # 1‑based inclusive
    strand = cols[4 + offset]
    chrom_len = len(fasta[chrom])

    if strand == '+':
        start0 = pos1_1b - 1                 # convert to 0‑based
        end0 = start0 + flank
    else:
        end0 = pos1_1b - 1                   # 0‑based inclusive of junction
        start0 = end0 - flank + 1            # inclusive start
        # Convert to half‑open for pyfaidx slice
        start0, end0 = start0, end0 + 1

    if start0 < 0 or end0 > chrom_len:
        if debug:
            print(f"[DEBUG] Out‑of‑bounds: {chrom}:{start0}-{end0} strand {strand}")
        return ""

    seq = fasta[chrom][start0:end0].seq
    if strand == '-':
        seq = seq.translate(str.maketrans("ACGTacgtNn", "TGCAtgcaNn"))[::-1]

    if len(seq) != flank:
        if debug:
            print(f"[DEBUG] Unexpected length {len(seq)} (expected {flank})")
        return ""
    return seq


# ------------------------------ Main programme ------------------------------ #

def main():
    parser = argparse.ArgumentParser(
        description=("GC‑balanced single‑fragment extraction (junction‑anchored) "
                     "with timestamped outputs."))
    parser.add_argument("-i", "--input", required=True,
                        help="Input .pairs(.gz) file.")
    parser.add_argument("-f", "--fasta", required=True,
                        help="Indexed reference FASTA.")
    parser.add_argument("-o", "--output-dir",
                        help="Output directory (timestamp appended if omitted).")
    parser.add_argument("-n", "--num-junctions", type=int, default=1000000,
                        help="Fragments to sample (default 1 000 000).")
    parser.add_argument("--chrom", default="chr1",
                        help="Chromosome for GC binning (default chr1).")
    parser.add_argument("--bin-size", type=int, default=1000,
                        help="Bin size for GC (default 1 kb).")
    parser.add_argument("--flank", type=int, default=50,
                        help="Length of fragment to extract (default 50).")
    parser.add_argument("--seed", type=int,
                        help="Random seed.")
    parser.add_argument("--debug", action="store_true",
                        help="Verbose debug output.")
    args = parser.parse_args()

    if args.seed is not None:
        random.seed(args.seed)

    timestamp = datetime.now().strftime("%Y%m%d_%H%M%S")
    pairs_basename = os.path.basename(args.input).replace(".pairs.gz", "")\
                                                .replace(".pairs", "")
    pairs_filename = os.path.basename(args.input)
    out_dir = args.output_dir or f"fragment1_rc_{pairs_basename}_{timestamp}"
    if args.output_dir and not args.output_dir.endswith(timestamp):
        out_dir = f"{args.output_dir}_{pairs_basename}_{timestamp}"
    os.makedirs(out_dir, exist_ok=True)

    fasta = Fasta(args.fasta, sequence_always_upper=True)

    # 1) GC quantiles
    gc_perc, quant = compute_gc_quantiles(fasta, args.chrom,
                                          args.bin_size, 100)
    quant_file = os.path.join(
        out_dir,
        f"{args.chrom}_gc_bins_quantiles_{pairs_basename}_"
        f"{pairs_filename}_{timestamp}.tsv")
    with open(quant_file, "w") as qf:
        qf.write("bin_start\tbin_end\tGC_percent\tquantile\n")
        for i, (g, q) in enumerate(zip(gc_perc, quant)):
            s = i*args.bin_size + 1
            e = min((i+1)*args.bin_size, len(fasta[args.chrom]))
            qf.write(f"{s}\t{e}\t{g:.2f}\t{int(q)}\n")

    # 2) Sample junctions
    sel_pairs, total_chr, sel_total, offset = \
        sample_junctions_by_quantile(
            args.input, args.chrom, quant, args.bin_size,
            args.num_junctions, debug=args.debug)

    # 3) Prepare nucleotide frequency matrix
    seq_len = args.flank
    base2idx = {b: i for i, b in enumerate("ACGT")}
    freq = np.zeros((4, seq_len), dtype=int)

    # 4) Output sequence TSV
    seq_tsv = os.path.join(
        out_dir,
        f"junction_sequences_{pairs_basename}_{pairs_filename}_{timestamp}.tsv.gz")
    gz_out = gzip.open(seq_tsv, "wt")

    # Header
    header = ["sequence"]
    if offset == 1:
        header.append("readID")
    header += ["chrom1", "pos1", "chrom2", "pos2", "strand1", "strand2"]
    if sel_pairs:
        nf = len(sel_pairs[0])
        extra_cols = ["pair_type", "mapq1", "mapq2"]
        start = 8 if offset == 1 else 7
        header += extra_cols[:max(0, nf-start)]
    gz_out.write("\t".join(header) + "\n")

    skipped = 0
    for cols in sel_pairs:
        seq = extract_fragment1_strand(
            cols, offset, fasta, args.flank, debug=args.debug)
        if not seq:
            skipped += 1
            continue
        for i, ch in enumerate(seq):
            if ch in base2idx:
                freq[base2idx[ch], i] += 1
        out_line = ([seq] + cols) if offset == 0 else \
            ([seq, cols[0]] + cols[1:])
        gz_out.write("\t".join(map(str, out_line)) + "\n")
    gz_out.close()
    written = sel_total - skipped

    # 5) Nucleotide‑frequency table
    freq_file = os.path.join(
        out_dir,
        f"nucleotide_frequency_{pairs_basename}_{pairs_filename}_{timestamp}.tsv")
    with open(freq_file, "w") as ff:
        ff.write("position\tA\tC\tG\tT\n")
        for i in range(seq_len):
            ff.write(f"{i+1}\t{freq[0,i]}\t{freq[1,i]}\t"
                     f"{freq[2,i]}\t{freq[3,i]}\n")

    # 6) Composition plot
    plot_pdf = os.path.join(
        out_dir,
        f"base_composition_{pairs_basename}_{pairs_filename}_{timestamp}.pdf")
    total = max(written, 1)
    frac = freq / total
    x = np.arange(1, seq_len+1)
    color = {'A': 'green', 'G': 'orange', 'C': 'blue', 'T': 'red'}
    order = ['A', 'G', 'C', 'T']
    plt.figure(figsize=(12, 6))
    bottom = np.zeros(seq_len)
    for base in order:
        idx = base2idx[base]
        plt.bar(x, frac[idx], bottom=bottom, color=color[base],
                width=1.0, label=base)
        bottom += frac[idx]
    plt.xlabel("Position in Fragment1 (1‑based from junction)")
    plt.ylabel("Fraction")
    plt.title(f"Base composition for '{pairs_basename}' ({pairs_filename})")
    plt.legend(title="Base", loc="upper right")
    plt.tight_layout()
    plt.savefig(plot_pdf)
    plt.close()

    # 7) Summary JSON
    summary = {
        "chromosome": args.chrom,
        "bin_size": args.bin_size,
        "num_quantiles": 100,
        "flank_length": args.flank,
        "pairs_file": pairs_filename,
        "pairs_stub": pairs_basename,
        "fasta_file": os.path.basename(args.fasta),
        "output_directory": os.path.basename(out_dir),
        f"total_pairs_on_{args.chrom}": total_chr,
        "requested_junctions": args.num_junctions,
        "sampled_fragments_written": written,
        "skipped_truncated": skipped,
        "timestamp": timestamp,
        "output_files": {
            "junction_sequences_tsv_gz": os.path.basename(seq_tsv),
            "gc_quantiles_tsv": os.path.basename(quant_file),
            "nucleotide_frequency_tsv": os.path.basename(freq_file),
            "composition_plot_pdf": os.path.basename(plot_pdf)
        }
    }
    with open(os.path.join(
            out_dir,
            f"summary_{pairs_basename}_{pairs_filename}_{timestamp}.json"), "w") as js:
        json.dump(summary, js, indent=4)

    if args.debug:
        print(json.dumps(summary, indent=2))


if __name__ == "__main__":
    main()
```
