## Supplementary material for "CAD-C: An engineered nuclease enables repair-free *in situ* proximity ligation and nucleosome-resolution chromosome walks in human cells": Custom code: pairs2bigwig_v5_20250819.py

```
#!/usr/bin/env python3
# -*- coding: utf-8 -*-
"""
pairs2bigwig_v4_FIXED_cli_compat.py  ·  2025-08-XX

Convert a pairtools .pairs(.gz) file (generated with
  --report-position junction
and typically
  --report-orientation junction
) into a single–base‑resolution BigWig that marks the exact ligation‑junction
nucleotide for each anchor.

Key fixes vs. 2025‑03‑25 "v4":
  • Removed the strand‑dependent ±1 bp shift. We now treat pos1/pos2 as the
    junction base (1‑based inclusive) and only convert to 0‑based half‑open
    when emitting BigWig (start = pos − 1, end = pos).
  • CLI is unchanged from the original v4.

Pipeline:
  1) Parse #chromsize header lines to get chromosome lengths.
  2) Optionally filter to autosomes (chr1..chr22), and compute an adjusted
     effective genome size that excludes --exclude-chr from depth scaling.
  3) Stream the body in chunks, count (chrom, pos) for both anchors exactly
     at pos1/pos2 (no shifting).
  4) Write a sorted intermediate file (version sort on chrom, numeric on pos),
     merge identical sites, scale to ~1x coverage (RPGC-like), and emit BigWig.

Inputs/assumptions:
  • .pairs(.gz) with #chromsize lines in the header.
  • pos1/pos2 are 1-based inclusive nucleotides at the junction (from parse2
    --report-position junction). If your file was generated differently,
    this script will still run but the genomic position semantics will differ.

Outputs:
  • <stub>_<YYYYMMDD_HHMMSS>.bw  (single‑bp BigWig)

Author: Jan Soroczynski (Risca Lab) — bug‑fix/cleanup & notes by request.
License: MIT
"""

import argparse
import gzip
import sys
import os
import logging
from collections import Counter
from multiprocessing import Pool
import tempfile
import subprocess
import datetime

try:
    import pandas as pd
    import numpy as np
    import pyBigWig
except ImportError as e:
    print(f"ERROR: Required library not found - {e}. "
          f"Please install pandas, numpy, and pyBigWig.", file=sys.stderr)
    sys.exit(1)


# ──────────────────────────────────────────────────────────────────────────
# CLI (unchanged)
# ──────────────────────────────────────────────────────────────────────────
def parse_args():
    parser = argparse.ArgumentParser(
        description=("Convert a pairtools .pairs.gz file to a single‑basepair "
                     "BigWig coverage track with an RPGC-like normalization and "
                     "optional autosome filtering. Assumes parse2 "
                     "'--report-position junction'."))
    parser.add_argument("-i", "--input", required=True,
                        help="Path to input .pairs(.gz) file from pairtools.")
    parser.add_argument("-o", "--output",
                        help=("Output .bigwig filename or directory. "
                              "If a file is provided, a timestamp is appended "
                              "before the extension. If a directory is provided, "
                              "the output file is named after the input file "
                              "with a timestamp."))
    parser.add_argument("-p", "--processors", type=int,
                        default=int(os.getenv("SLURM_CPUS_PER_TASK", 1)),
                        help=("Number of processes for parallel chunk processing "
                              "(default: 1, or $SLURM_CPUS_PER_TASK if set)."))
    parser.add_argument("--effective-size", type=int, default=2913022398,
                        help=("Effective genome size for 1x coverage normalization "
                              "(default: 2913022398 for hg38). "
                              "NOTE: This value may be overridden if "
                              "--only-autosomes is used or after filtering."))
    parser.add_argument("--exclude-chr", default="chrX",
                        help=("Chromosome to exclude from the normalization depth "
                              "calculation (default: chrX). Coverage is still "
                              "written for this chromosome if present, but it "
                              "doesn't affect the scaling factor."))
    parser.add_argument("--chunksize", type=int, default=500000,
                        help="Number of rows to process per chunk (default: 500000).")
    parser.add_argument("--only-autosomes", action="store_true",
                        help=("If set, keep only chr1..chr22 in both coverage "
                              "and effective genome size (others excluded)."))
    # External sort tuning
    parser.add_argument("--sort-parallel", action="store_true",
                        help="Use '--parallel=N' in 'sort' with N = --processors.")
    parser.add_argument("--sort-buffer-size", default=None,
                        help="Pass '--buffer-size=<value>' (e.g. 4G) to 'sort'.")
    return parser.parse_args()


# ──────────────────────────────────────────────────────────────────────────
# Chrom size filtering + adjusted effective size (same behavior)
# ──────────────────────────────────────────────────────────────────────────
def _adjust_chrom_sizes_and_filter(chrom_sizes, args):
    """
    Filter chromosome sizes based on --only-autosomes, then compute the final
    effective genome size by summing the filtered lengths and subtracting the
    length of --exclude-chr if present.

    Returns:
      filtered_chrom_sizes: list[(chrom, length)]
      adjusted_effective_size: int
    """
    chrom_dict = dict(chrom_sizes)

    if args.only_autosomes:
        desired = [f"chr{i}" for i in range(1, 23)]
        filtered = [(c, chrom_dict[c]) for c in desired if c in chrom_dict]
    else:
        filtered = chrom_sizes

    if not filtered:
        return [], 0

    total_len = sum(L for _, L in filtered)
    excluded_len = next((L for c, L in filtered if c == args.exclude_chr), 0)
    adjusted = total_len - excluded_len
    if adjusted < 0:
        logging.warning("Adjusted effective size < 0; clamping to 0.")
        adjusted = 0
    return filtered, adjusted


# ──────────────────────────────────────────────────────────────────────────
# Chunk processing: count junction bases exactly at pos1/pos2
# ──────────────────────────────────────────────────────────────────────────
def process_chunk(df, chrom_lengths):
    """
    Count (chrom, pos) for both anchors. Assumes pos1/pos2 are 1‑based inclusive
    nucleotides at the ligation junction (parse2 --report-position junction).
    NO strand‑dependent shift is applied here.
    """
    counts = Counter()

    # Arrays (categories are okay; we only test membership in chrom_lengths)
    chrom1 = df['chr1'].values
    chrom2 = df['chr2'].values
    pos1 = df['pos1'].values
    pos2 = df['pos2'].values

    # First anchor
    for c, p in zip(chrom1, pos1):
        if c in chrom_lengths and 1 <= p <= chrom_lengths[c]:
            counts[(c, int(p))] += 1
    # Second anchor
    for c, p in zip(chrom2, pos2):
        if c in chrom_lengths and 1 <= p <= chrom_lengths[c]:
            counts[(c, int(p))] += 1

    return counts


# ──────────────────────────────────────────────────────────────────────────
# Main
# ──────────────────────────────────────────────────────────────────────────
def main():
    args = parse_args()
    logging.basicConfig(format="%(asctime)s [%(levelname)s] %(message)s",
                        level=logging.INFO)
    infile = args.input

    # Timestamped output name
    ts = datetime.datetime.now().strftime("%Y%m%d_%H%M%S")
    if args.output:
        if os.path.isdir(args.output):
            stub = os.path.basename(os.path.splitext(os.path.splitext(infile)[0])[0])
            outfile = os.path.join(args.output, f"{stub}_{ts}.bw")
        else:
            base, ext = os.path.splitext(args.output)
            outfile = f"{base}_{ts}{ext if ext else '.bw'}"
    else:
        stub = os.path.basename(os.path.splitext(os.path.splitext(infile)[0])[0])
        outfile = os.path.join(os.path.dirname(infile), f"{stub}_{ts}.bw")

    logging.info(f"Input file: {infile}")
    logging.info(f"Output BigWig: {outfile}")
    logging.info(f"Processors: {max(1, args.processors)}")
    logging.info(f"Chunk size: {args.chunksize}")
    logging.info(f"Exclude chromosome for normalization: {args.exclude_chr}")

    # 1) Read #chromsize header
    raw_chrom_sizes = []
    try:
        opener = gzip.open if infile.endswith(".gz") else open
        with opener(infile, 'rt') as f:
            for line in f:
                if not line.startswith('#'):
                    break
                if line.startswith('#chromsize'):
                    parts = line.strip().split()
                    if len(parts) >= 3:
                        chrom = parts[1]
                        size = int(parts[2])
                        raw_chrom_sizes.append((chrom, size))
    except Exception as e:
        logging.error(f"Failed to read input file header: {e}")
        sys.exit(1)

    if not raw_chrom_sizes:
        logging.error("No #chromsize lines found in header; cannot build BigWig.")
        sys.exit(1)

    # 2) Filter chromosomes + compute adjusted effective size
    filtered_chrom_sizes, eff_size = _adjust_chrom_sizes_and_filter(raw_chrom_sizes, args)
    if not filtered_chrom_sizes or eff_size <= 0:
        logging.error("No valid chromosomes remain or adjusted effective size is 0.")
        sys.exit(1)

    chrom_lengths = {c: L for c, L in filtered_chrom_sizes}
    logging.info(f"Parsed {len(raw_chrom_sizes)} chroms; using {len(filtered_chrom_sizes)} "
                 f"after filtering. Adjusted effective size = {eff_size}.")

    # 3) Temporary file for coverage lines
    tmp_fd, tmp_path = tempfile.mkstemp(prefix="junction_cov_", suffix=".txt")
    tmp_fh = os.fdopen(tmp_fd, 'w')
    total_non_excl = 0

    # 4) Stream the body in chunks
    try:
        reader = pd.read_csv(
            infile, sep='\t', comment='#', header=None, chunksize=args.chunksize,
            # Keep the same columns as original v4 (we ignore strands here)
            usecols=[1, 2, 3, 4, 5, 6],
            names=['chr1', 'pos1', 'chr2', 'pos2', 'strand1', 'strand2'],
            dtype={'chr1': 'category', 'chr2': 'category'}
        )
    except Exception as e:
        logging.error(f"Failed to open/parse input file: {e}")
        sys.exit(1)

    def _write_counts(counter):
        nonlocal total_non_excl
        for (chrom, pos), count in counter.items():
            # Write 1-based position (we will convert to 0-based when creating BW)
            tmp_fh.write(f"{chrom}\t{pos}\t{count}\n")
            if chrom != args.exclude_chr:
                total_non_excl += count

    logging.info("Processing chunks…")
    nproc = max(1, args.processors)
    if nproc > 1:
        pool = Pool(processes=nproc)
        futures = []
        for chunk in reader:
            # ensure integer dtype for positions
            chunk['pos1'] = pd.to_numeric(chunk['pos1'], downcast='integer')
            chunk['pos2'] = pd.to_numeric(chunk['pos2'], downcast='integer')
            futures.append(pool.apply_async(process_chunk, (chunk[['chr1', 'pos1', 'chr2', 'pos2']], chrom_lengths)))
        pool.close()
        pool.join()
        for f in futures:
            _write_counts(f.get())
    else:
        for chunk in reader:
            chunk['pos1'] = pd.to_numeric(chunk['pos1'], downcast='integer')
            chunk['pos2'] = pd.to_numeric(chunk['pos2'], downcast='integer')
            cnt = process_chunk(chunk[['chr1', 'pos1', 'chr2', 'pos2']], chrom_lengths)
            _write_counts(cnt)

    tmp_fh.close()
    logging.info(f"Intermediate coverage file written: {tmp_path}")

    # 5) Compute RPGC-like normalization factor
    if total_non_excl == 0:
        logging.warning(f"No anchors outside {args.exclude_chr}; using scale=1.")
        scale = 1.0
    else:
        scale = eff_size / float(total_non_excl)
    logging.info(f"Total non-{args.exclude_chr} anchors = {total_non_excl}; scale = {scale:.6f}")

    # 6) External sort (version sort chrom, numeric pos)
    sorted_path = tmp_path + ".sorted"
    sort_cmd = ["sort", "-k1,1V", "-k2,2n", tmp_path, "-o", sorted_path]
    if args.sort_parallel:
        sort_cmd.insert(1, f"--parallel={nproc}")
    if args.sort_buffer_size:
        sort_cmd.insert(1, f"--buffer-size={args.sort_buffer_size}")
    logging.info("Sorting with: %s", ' '.join(sort_cmd))
    try:
        subprocess.run(sort_cmd, check=True)
    except subprocess.CalledProcessError as e:
        logging.error(f"Sorting intermediate file failed: {e}")
        sys.exit(1)
    finally:
        try:
            os.remove(tmp_path)
        except OSError:
            pass

    # 7) Create BigWig and write merged, scaled coverage
    try:
        bw = pyBigWig.open(outfile, "w")
        bw.addHeader(filtered_chrom_sizes)
    except Exception as e:
        logging.error(f"Failed to create BigWig {outfile}: {e}")
        sys.exit(1)

    logging.info("Merging coverage and writing BigWig…")
    current_chr = None
    current_pos = None
    current_sum = 0
    batch_chr = None
    batch_positions = []
    batch_values = []
    BATCH_THRESHOLD = 100000

    def flush_batch(chrom):
        nonlocal batch_positions, batch_values
        if not batch_positions:
            return
        starts = [p - 1 for p in batch_positions]  # 0-based
        ends = batch_positions                      # 1-based end = start+1
        values = [v * scale for v in batch_values]
        try:
            bw.addEntries([chrom] * len(starts), starts, ends=ends, values=values)
        except Exception as e:
            logging.error(f"pyBigWig write error on {chrom}: {e}")
            raise
        batch_positions.clear()
        batch_values.clear()

    with open(sorted_path, 'r') as sf:
        for line in sf:
            parts = line.rstrip().split("\t")
            if len(parts) != 3:
                continue
            chrom, pos_str, count_str = parts
            if chrom not in chrom_lengths:
                continue
            pos = int(pos_str)
            count = int(count_str)

            # merge identical (chrom, pos) rows
            if current_chr is None:
                current_chr, current_pos, current_sum = chrom, pos, count
                batch_chr = chrom
                continue

            if chrom == current_chr and pos == current_pos:
                current_sum += count
                continue

            # push merged record for previous site
            batch_positions.append(current_pos)
            batch_values.append(current_sum)

            # flush by chromosome change or large batch
            if chrom != current_chr or len(batch_positions) >= BATCH_THRESHOLD:
                flush_batch(current_chr)
                batch_chr = chrom

            current_chr, current_pos, current_sum = chrom, pos, count

        # flush last site
        if current_chr is not None:
            batch_positions.append(current_pos)
            batch_values.append(current_sum)
            flush_batch(current_chr)

    try:
        os.remove(sorted_path)
    except OSError:
        pass
    bw.close()
    logging.info(f"BigWig file created: {outfile}")


if __name__ == "__main__":
    main()
```
