## Supplementary material for "CAD-C: An engineered nuclease enables repair-free *in situ* proximity ligation and nucleosome-resolution chromosome walks in human cells": Custom code: Extended_Data_Fig5bc_plot_insulation_correlation.py

```
#!/usr/bin/env python3
# -*- coding: utf-8 -*-
import argparse, os, math
import numpy as np, pandas as pd
from matplotlib.backends.backend_pdf import PdfPages
import matplotlib.pyplot as plt

CAND_VALCOLS = ["log2_insulation_score_500000"]

def read_one(path, value_col=None, chroms=None):
    df = pd.read_csv(path, sep="\t", comment="#")
    if value_col is None:
        for c in CAND_VALCOLS:
            if c in df.columns: value_col = c; break
    if value_col is None or value_col not in df.columns:
        raise ValueError(f"Value column not found in {path}. Try --value-col")
    for c in ["chrom","start","end"]:
        if c not in df.columns: raise ValueError(f"{path} missing column '{c}'")
    df = df[["chrom","start","end", value_col]].copy()
    df.rename(columns={value_col:"ins"}, inplace=True)
    if chroms is not None: df = df[df["chrom"].isin(chroms)].copy()
    return df

def merge_on_bins(dfs, labels):
    key = ["chrom","start","end"]
    base = None
    for lab, df in zip(labels, dfs):
        df2 = df.copy(); df2.columns = key + [lab]
        base = df2 if base is None else base.merge(df2, on=key, how="inner")
    return base

def mask_nonfinite(df, labels):
    m = np.isfinite(df[labels]).all(axis=1)
    return df.loc[m].reset_index(drop=True)

def corr_weighted_by_chrom(df, a, b, method="spearman"):
    vals=[]
    for chrom, g in df.groupby("chrom", sort=False):
        if len(g)>=10:
            r = g[a].corr(g[b], method=method)
            if pd.isna(r): continue
            vals.append((len(g), r))
    if not vals: return math.nan
    n = np.array([v[0] for v in vals], float)
    r = np.array([v[1] for v in vals], float)
    w = n / n.sum()
    return float((w*r).sum())

def corr_simple(df, a, b, method="spearman"):
    return float(df[a].corr(df[b], method=method))

def build_corr_mats(df, labels, weighted=False):
    n=len(labels)
    pear=np.eye(n, dtype=float); spear=np.eye(n, dtype=float)
    for i in range(n):
        for j in range(i+1,n):
            a,b=labels[i],labels[j]
            if weighted:
                rp=corr_weighted_by_chrom(df,a,b,"pearson")
                rs=corr_weighted_by_chrom(df,a,b,"spearman")
            else:
                rp=corr_simple(df,a,b,"pearson")
                rs=corr_simple(df,a,b,"spearman")
            pear[i,j]=pear[j,i]=rp; spear[i,j]=spear[j,i]=rs
    return pear,spear

def plot_heatmap(ax, M, labels, title):
    im=ax.imshow(M, vmin=0, vmax=1)
    ax.set_title(title)
    ax.set_xticks(range(len(labels))); ax.set_xticklabels(labels, rotation=45, ha="right")
    ax.set_yticks(range(len(labels))); ax.set_yticklabels(labels)
    for i in range(len(labels)):
        for j in range(len(labels)):
            ax.text(j,i,f"{M[i,j]:.2f}", ha="center", va="center", fontsize=8)
    plt.colorbar(im, ax=ax, fraction=0.046, pad=0.04)

def make_table_page(pdf, labels, pear, spear, subtitle):
    fig,ax=plt.subplots(figsize=(10,6)); ax.axis('off')
    lines=[f"Pairwise correlations ({subtitle})",""]
    lines.append("Pearson:")
    header="       "+"  ".join([f"{l:>10}" for l in labels]); lines.append(header)
    for i,li in enumerate(labels):
        row=f"{li:>6} "+"  ".join([f"{pear[i,j]:>10.3f}" for j in range(len(labels))]); lines.append(row)
    lines.append(""); lines.append("Spearman:"); lines.append(header)
    for i,li in enumerate(labels):
        row=f"{li:>6} "+"  ".join([f"{spear[i,j]:>10.3f}" for j in range(len(labels))]); lines.append(row)
    ax.text(0.01,0.99,"\n".join(lines), va="top", family="monospace")
    pdf.savefig(fig, bbox_inches="tight"); plt.close(fig)

def main():
    ap=argparse.ArgumentParser(description="Insulation correlations → editable PDF & CSVs")
    ap.add_argument("--files", nargs="+", required=True, help="Insulation TSV/bedGraph files (>=2)")
    ap.add_argument("--labels", nargs="+", help="Labels for samples (same length as files)")
    ap.add_argument("--value-col", default=None, help="Insulation value column name")
    ap.add_argument("--chroms", nargs="+", default=None, help="Restrict to these chroms")
    ap.add_argument("--mask-nonfinite", action="store_true", help="Drop bins with any non-finite value")
    ap.add_argument("--by-chrom-weighted", action="store_true", help="Per-chrom correlations weighted by bin count")
    ap.add_argument("--out-pdf", default="insulation_correlations.pdf", help="Output PDF (vector)")
    ap.add_argument("--out-prefix", default="insulation_correlations", help="Prefix for CSV outputs")
    args=ap.parse_args()

    files=args.files
    labels=args.labels if args.labels else [os.path.basename(f).split('.')[0] for f in files]
    if len(labels)!=len(files): ap.error("labels length must match files length")

    dfs=[read_one(p, value_col=args.value_col, chroms=args.chroms) for p in files]
    merged=merge_on_bins(dfs, labels)
    if args.mask_nonfinite: merged=mask_nonfinite(merged, labels)

    pear,spear = build_corr_mats(merged, labels, weighted=args.by_chrom_weighted)
    pd.DataFrame(pear,index=labels,columns=labels).to_csv(f"{args.out_prefix}.pearson.csv")
    pd.DataFrame(spear,index=labels,columns=labels).to_csv(f"{args.out_prefix}.spearman.csv")

    with PdfPages(args.out_pdf) as pdf:
        fig,ax=plt.subplots(figsize=(6,5))
        plot_heatmap(ax, pear, labels, "Pearson correlation"+(" (weighted)" if args.by_chrom_weighted else ""))
        pdf.savefig(fig, bbox_inches="tight"); plt.close(fig)
        fig,ax=plt.subplots(figsize=(6,5))
        plot_heatmap(ax, spear, labels, "Spearman correlation"+(" (weighted)" if args.by_chrom_weighted else ""))
        pdf.savefig(fig, bbox_inches="tight"); plt.close(fig)
        make_table_page(pdf, labels, pear, spear, "Pearson & Spearman")

    print(f"[OK] PDF: {args.out_pdf}")
    print(f"[OK] CSV: {args.out_prefix}.pearson.csv, {args.out_prefix}.spearman.csv")

if __name__=="__main__": main()
```
