## Supplementary material for "CAD-C: An engineered nuclease enables repair-free *in situ* proximity ligation and nucleosome-resolution chromosome walks in human cells": Custom code: Extended_Data_Fig5de_call_loop_mustache.sh

```
#!/bin/bash
# Parse input arguments
while getopts i:o: flag
do
    case "${flag}" in
        i) input_folder=${OPTARG};;
        o) output_dir=${OPTARG};;
    esac
done

# Check if input folder and output directory are provided
if [ -z "$input_folder" ] || [ -z "$output_dir" ]; then
    echo "Usage: $0 -i <input_folder> -o <output_directory>"
    exit 1
fi

# Define the resolutions
resolutions=(2500 5000 10000)

# Create output directories for each tool if they don't exist
mkdir -p ${output_dir}/mustache

# Loop through all .mcool files in the input folder
for input_file in ${input_folder}/*.mcool; do
    for res in "${resolutions[@]}"; do
        echo "Processing ${input_file} at resolution ${res}bp..."
        # Mustache
        python /lustre/fs4/risc_lab/scratch/wzuo/tools/mustache/mustache/mustache.py -f "${input_file}" -o "${output_dir}/mustache/$(basename ${input_file})_mustache_loops_${res}.bedpe" -st 0.8 -pt 0.1 -r ${res} -p 14
        echo "Done with ${input_file} at resolution ${res}bp with Mustache."
    done
done

echo "Loop calling completed for all .mcool files and resolutions."
```
