## Supplementary material for "CAD-C: An engineered nuclease enables repair-free *in situ* proximity ligation and nucleosome-resolution chromosome walks in human cells": Custom code: Extended_Data_Fig6b_loop_anchor_pileup_with_HiChIP_data.sh

```
#!/bin/bash
#SBATCH --job-name=HiC_Processing
#SBATCH --output=hic_processing_%j.log
#SBATCH --cpus-per-task=12
#SBATCH --mem=32G
#SBATCH --time=24:00:00
#SBATCH --partition=hpc_l40_a

# Load necessary modules or activate your conda environment
# module load cooltools
source /lustre/fs4/risc_lab/scratch/wzuo/tools/conda/etc/profile.d/conda.sh
conda activate coolpup

GM12878_CADC_MicroC_5kb='/rugpfs/fs0/risc_lab/scratch/wzuo/project/CAD/GM12878_dataset/loops/loop_2500/GM12878_CADC_MicroC_2500_loop_5kb_slop2.bedpe'
GM12878_CADC_only_5kb='/rugpfs/fs0/risc_lab/scratch/wzuo/project/CAD/GM12878_dataset/loops/loop_2500/GM12878_CADC_only_2500_loop_5kb_slop2.bedpe'
GM12878_MicroC_only_5kb='/rugpfs/fs0/risc_lab/scratch/wzuo/project/CAD/GM12878_dataset/loops/loop_2500/GM12878_MicroC_only_2500_loop_5kb_slop2.bedpe'

coolpup.py /rugpfs/fs0/risc_lab/scratch/wzuo/project/CAD/dataset/GM_HiChIP/GM_HiChIP_cohesin_5kb.cool ${GM12878_CADC_MicroC_5kb} --unbalanced --pad 250 --nshifts 10 --mindist 500000 --maxdist 2000000 --outname GM12878_CADC_MicroC_5kb_HiChIP_cohesin.txt --n_proc 6 
coolpup.py /rugpfs/fs0/risc_lab/scratch/wzuo/project/CAD/dataset/GM_HiChIP/GM_HiChIP_cohesin_5kb.cool ${GM12878_CADC_only_5kb} --unbalanced --pad 250 --nshifts 10 --mindist 500000 --maxdist 2000000 --outname GM12878_CADC_only_5kb_HiChIP_cohesin.txt --n_proc 6 
coolpup.py /rugpfs/fs0/risc_lab/scratch/wzuo/project/CAD/dataset/GM_HiChIP/GM_HiChIP_cohesin_5kb.cool ${GM12878_MicroC_only_5kb} --unbalanced --pad 250 --nshifts 10 --mindist 500000 --maxdist 2000000 --outname GM12878_MicroC_only_5kb_HiChIP_cohesin.txt --n_proc 6 

coolpup.py /rugpfs/fs0/risc_lab/scratch/wzuo/project/CAD/dataset/GM_HiChIP/GM_HiChIP_H3K27Ac_5kb.cool ${GM12878_CADC_MicroC_5kb} --unbalanced --pad 250 --nshifts 10 --mindist 500000 --maxdist 2000000 --outname GM12878_CADC_MicroC_5kb_HiChIP_H3K27Ac.txt --n_proc 6 
coolpup.py /rugpfs/fs0/risc_lab/scratch/wzuo/project/CAD/dataset/GM_HiChIP/GM_HiChIP_H3K27Ac_5kb.cool ${GM12878_CADC_only_5kb} --unbalanced --pad 250 --nshifts 10 --mindist 500000 --maxdist 2000000 --outname GM12878_CADC_only_5kb_HiChIP_H3K27Ac.txt --n_proc 6 
coolpup.py /rugpfs/fs0/risc_lab/scratch/wzuo/project/CAD/dataset/GM_HiChIP/GM_HiChIP_H3K27Ac_5kb.cool ${GM12878_MicroC_only_5kb} --unbalanced --pad 250 --nshifts 10 --mindist 500000 --maxdist 2000000 --outname GM12878_MicroC_only_5kb_HiChIP_H3K27Ac.txt --n_proc 6 
```
