## Supplementary material for "CAD-C: An engineered nuclease enables repair-free *in situ* proximity ligation and nucleosome-resolution chromosome walks in human cells": Custom code: Extended_Data_Fig7a_loop_anchor_pileup_GM_Dovetail.sh

```
#!/bin/bash
#SBATCH --job-name=HiC_Processing
#SBATCH --output=hic_processing_%j.log
#SBATCH --cpus-per-task=12
#SBATCH --mem=32G
#SBATCH --time=24:00:00
#SBATCH --partition=hpc_a10_a

source /programs/biogrids.shrc
cores=14

input_dir="/rugpfs/fs0/risc_lab/scratch/wzuo/project/CAD/loops_subgroup/bed/"
output_dir="/rugpfs/fs0/risc_lab/scratch/wzuo/project/CAD/pile/loop_anchor/merge"

GM_both="$input_dir/GM12878_CADC_MicroC_both_10kb_slop.bed"
GM_CO="$input_dir/GM12878_CADC_only_10kb_slop.bed"
GM_MO="$input_dir/GM12878_MicroC_only_10kb_slop.bed"

GM_ATAC='/rugpfs/fs0/risc_lab/scratch/wzuo/project/CAD/dataset/bw/GM_ATAC_ENCFF667MDI.bigWig' 
GM_CTCF='/rugpfs/fs0/risc_lab/scratch/wzuo/project/CAD/dataset/bw/GM_CTCF_ENCFF680XUD.bigWig'
GM_CREB1='/rugpfs/fs0/risc_lab/scratch/wzuo/project/CAD/dataset/bw/GM_CREB1_ENCFF906KGM.bigWig'
GM_H3K27ac='/rugpfs/fs0/risc_lab/scratch/wzuo/project/CAD/dataset/bw/GM_H3K27ac_ENCFF798KYP.bigWig'
GM_H3K4me3='/rugpfs/fs0/risc_lab/scratch/wzuo/project/CAD/dataset/bw/GM_H3K4me3_ENCFF012DMX.bigWig'
GM_POLR2A='/rugpfs/fs0/risc_lab/scratch/wzuo/project/CAD/dataset/bw/GM_POLR2A_ENCFF322DRU.bigWig'
GM_Rad21='/rugpfs/fs0/risc_lab/scratch/wzuo/project/CAD/dataset/bw/GM_Rad21_ENCFF407HJR.bigWig'
GM_RELB='/rugpfs/fs0/risc_lab/scratch/wzuo/project/CAD/dataset/bw/GM_RELB_ENCFF936FAN.bigWig'
GM_PML='/rugpfs/fs0/risc_lab/scratch/wzuo/project/CAD/dataset/bw/GM_PML_ENCFF644HXX.bigWig' 
GM_minus='/rugpfs/fs0/risc_lab/scratch/wzuo/project/CAD/dataset/bw/GSM1480326_GM12878_GROseq_minus_liftover.bw' 
GM_plus='/rugpfs/fs0/risc_lab/scratch/wzuo/project/CAD/dataset/bw/GSM1480326_GM12878_GROseq_plus_liftover.bw'
GM_H2AZ='/rugpfs/fs0/risc_lab/scratch/wzuo/project/CAD/dataset/bw/GM_H2AZ_ENCFF270HCF.bigWig' 
GM_minus='/rugpfs/fs0/risc_lab/scratch/wzuo/project/CAD/dataset/bw/GSM1480326_GM12878_GROseq_plus_liftover.bw' 
GM_plus='/rugpfs/fs0/risc_lab/scratch/wzuo/project/CAD/dataset/bw/conversion/GSM1480326_GM12878_GROseq_minus_liftover_modified_sorted.bw'
GM_merge='/rugpfs/fs0/risc_lab/scratch/wzuo/project/CAD/dataset/bw/conversion/GSM1480326_GM12878_GROseq_liftover_merge_sorted.bw'

computeMatrix reference-point --referencePoint center -S ${GM_H2AZ} -R ${GM_both} ${GM_CO} ${GM_MO} -o "$output_dir/GM_H2AZ_pile.mat.gz" -b 25000 -a 25000 -bs 100 --skipZeros -p $cores
plotHeatmap -m "$output_dir/GM_H2AZ_pile.mat.gz" -out "$output_dir/GM_H2AZ_pile.heatmap.pdf" --perGroup --colorMap "inferno" --sortUsing sum --dpi 300 -T "" --startLabel ""  --endLabel "" --legendLocation "upper-right" --heatmapWidth 12 
plotProfile -m "$output_dir/GM_H2AZ_pile.mat.gz" -out "$output_dir/GM_H2AZ_pile.profile.pdf" --perGroup --dpi 300 -T "" --startLabel ""  --endLabel "" --legendLocation "upper-right" --plotHeight 10 --plotWidth 12

computeMatrix reference-point --referencePoint center -S ${GM_minus} -R ${GM_both} ${GM_CO} ${GM_MO} -o "$output_dir/GM_minus_pile.mat.gz" -b 25000 -a 25000 -bs 100 --skipZeros -p $cores
plotHeatmap -m "$output_dir/GM_minus_pile.mat.gz" -out "$output_dir/GM_minus_pile.heatmap.pdf" --perGroup --zMin -3 --zMax -2 --colorMap "inferno" --sortUsing sum --dpi 300 -T "" --startLabel ""  --endLabel "" --legendLocation "upper-right" --heatmapWidth 12 
plotProfile -m "$output_dir/GM_minus_pile.mat.gz" -out "$output_dir/GM_minus_pile.profile.pdf" --perGroup --dpi 300 -T "" --startLabel ""  --endLabel "" --legendLocation "upper-right" --plotHeight 10 --plotWidth 12

computeMatrix reference-point --referencePoint center -S ${GM_plus} -R ${GM_both} ${GM_CO} ${GM_MO} -o "$output_dir/GM_plus_pile.mat.gz" -b 25000 -a 25000 -bs 100 --skipZeros -p $cores
plotHeatmap -m "$output_dir/GM_plus_pile.mat.gz" -out "$output_dir/GM_plus_pile.heatmap.pdf" --perGroup --zMin 1.5 --zMax 2 --colorMap "inferno" --sortUsing sum --dpi 300 -T "" --startLabel ""  --endLabel "" --legendLocation "upper-right" --heatmapWidth 12 
plotProfile -m "$output_dir/GM_plus_pile.mat.gz" -out "$output_dir/GM_plus_pile.profile.pdf" --perGroup --dpi 300 -T "" --startLabel ""  --endLabel "" --legendLocation "upper-right" --plotHeight 10 --plotWidth 12

computeMatrix reference-point --referencePoint center -S ${GM_ATAC} -R ${GM_both} ${GM_CO} ${GM_MO} -o "$output_dir/GM_ATAC_pile.mat.gz" -b 25000 -a 25000 -bs 100 --skipZeros -p $cores
plotHeatmap -m "$output_dir/GM_ATAC_pile.mat.gz" -out "$output_dir/GM_ATAC_pile.heatmap.pdf" --perGroup --zMin 0 --zMax 35 --colorMap "inferno" --sortUsing sum --dpi 300 -T "" --startLabel ""  --endLabel "" --legendLocation "upper-right" --heatmapWidth 12 
plotProfile -m "$output_dir/GM_ATAC_pile.mat.gz" -out "$output_dir/GM_ATAC_pile.profile.pdf" --perGroup --dpi 300 -T "" --startLabel ""  --endLabel "" --legendLocation "upper-right" --plotHeight 10 --plotWidth 12

computeMatrix reference-point --referencePoint center -S ${GM_CREB1} -R ${GM_both} ${GM_CO} ${GM_MO} -o "$output_dir/GM_CREB1_pile.mat.gz" -b 25000 -a 25000 -bs 100 --skipZeros -p $cores
plotHeatmap -m "$output_dir/GM_CREB1_pile.mat.gz" -out "$output_dir/GM_CREB1_pile.heatmap.pdf" --perGroup --zMin 0.3 --zMax 1 --colorMap "inferno" --sortUsing sum --dpi 300 -T "" --startLabel ""  --endLabel "" --legendLocation "upper-right" --heatmapWidth 12 
plotProfile -m "$output_dir/GM_CREB1_pile.mat.gz" -out "$output_dir/GM_CREB1_pile.profile.pdf" --perGroup --dpi 300 -T "" --startLabel ""  --endLabel "" --legendLocation "upper-right" --plotHeight 10 --plotWidth 12

computeMatrix reference-point --referencePoint center -S ${GM_CTCF} -R ${GM_both} ${GM_CO} ${GM_MO} -o "$output_dir/GM_CTCF_pile.mat.gz" -b 25000 -a 25000 -bs 100 --skipZeros -p $cores
plotHeatmap -m "$output_dir/GM_CTCF_pile.mat.gz" -out "$output_dir/GM_CTCF_pile.heatmap.pdf" --perGroup --zMin 0 --zMax 1.5 --colorMap "inferno" --sortUsing sum --dpi 300 -T "" --startLabel ""  --endLabel "" --legendLocation "upper-right" --heatmapWidth 12 
plotProfile -m "$output_dir/GM_CTCF_pile.mat.gz" -out "$output_dir/GM_CTCF_pile.profile.pdf" --perGroup --dpi 300 -T "" --startLabel ""  --endLabel "" --legendLocation "upper-right" --plotHeight 10 --plotWidth 12

computeMatrix reference-point --referencePoint center -S ${GM_H3K27ac} -R ${GM_both} ${GM_CO} ${GM_MO} -o "$output_dir/GM_H3K27ac_pile.mat.gz" -b 25000 -a 25000 -bs 100 --skipZeros -p $cores
plotHeatmap -m "$output_dir/GM_H3K27ac_pile.mat.gz" -out "$output_dir/GM_H3K27ac_pile.heatmap.pdf" --perGroup --zMin 0 --zMax 10 --colorMap "inferno" --sortUsing sum --dpi 300 -T "" --startLabel ""  --endLabel "" --legendLocation "upper-right" --heatmapWidth 12 
plotProfile -m "$output_dir/GM_H3K27ac_pile.mat.gz" -out "$output_dir/GM_H3K27ac_pile.profile.pdf" --perGroup --dpi 300 -T "" --startLabel ""  --endLabel "" --legendLocation "upper-right" --plotHeight 10 --plotWidth 12

computeMatrix reference-point --referencePoint center -S ${GM_H3K4me3} -R ${GM_both} ${GM_CO} ${GM_MO} -o "$output_dir/GM_H3K4me3_pile.mat.gz" -b 25000 -a 25000 -bs 100 --skipZeros -p $cores
plotHeatmap -m "$output_dir/GM_H3K4me3_pile.mat.gz" -out "$output_dir/GM_H3K4me3_pile.heatmap.pdf" --perGroup --zMin 0 --zMax 6 --colorMap "inferno" --sortUsing sum --dpi 300 -T "" --startLabel ""  --endLabel "" --legendLocation "upper-right" --heatmapWidth 12 
plotProfile -m "$output_dir/GM_H3K4me3_pile.mat.gz" -out "$output_dir/GM_H3K4me3_pile.profile.pdf" --perGroup --dpi 300 -T "" --startLabel ""  --endLabel "" --legendLocation "upper-right" --plotHeight 10 --plotWidth 12

computeMatrix reference-point --referencePoint center -S ${GM_POLR2A} -R ${GM_both} ${GM_CO} ${GM_MO} -o "$output_dir/GM_POLR2A_pile.mat.gz" -b 25000 -a 25000 -bs 100 --skipZeros -p $cores
plotHeatmap -m "$output_dir/GM_POLR2A_pile.mat.gz" -out "$output_dir/GM_POLR2A_pile.heatmap.pdf" --perGroup --zMin 0 --zMax 6 --colorMap "inferno" --sortUsing sum --dpi 300 -T "" --startLabel ""  --endLabel "" --legendLocation "upper-right" --heatmapWidth 12 
plotProfile -m "$output_dir/GM_POLR2A_pile.mat.gz" -out "$output_dir/GM_POLR2A_pile.profile.pdf" --perGroup --dpi 300 -T "" --startLabel ""  --endLabel "" --legendLocation "upper-right" --plotHeight 10 --plotWidth 12

computeMatrix reference-point --referencePoint center -S ${GM_Rad21} -R ${GM_both} ${GM_CO} ${GM_MO} -o "$output_dir/GM_Rad21_pile.mat.gz" -b 25000 -a 25000 -bs 100 --skipZeros -p $cores
plotHeatmap -m "$output_dir/GM_Rad21_pile.mat.gz" -out "$output_dir/GM_Rad21_pile.heatmap.pdf" --perGroup --zMin 0 --zMax 1 --colorMap "inferno" --sortUsing sum --dpi 300 -T "" --startLabel ""  --endLabel "" --legendLocation "upper-right" --heatmapWidth 12 
plotProfile -m "$output_dir/GM_Rad21_pile.mat.gz" -out "$output_dir/GM_Rad21_pile.profile.pdf" --perGroup --dpi 300 -T "" --startLabel ""  --endLabel "" --legendLocation "upper-right" --plotHeight 10 --plotWidth 12

computeMatrix reference-point --referencePoint center -S ${GM_RELB} -R ${GM_both} ${GM_CO} ${GM_MO} -o "$output_dir/GM_RELB_pile.mat.gz" -b 25000 -a 25000 -bs 100 --skipZeros -p $cores
plotHeatmap -m "$output_dir/GM_RELB_pile.mat.gz" -out "$output_dir/GM_RELB_pile.heatmap.pdf" --perGroup --zMin 0 --zMax 6 --colorMap "inferno" --sortUsing sum --dpi 300 -T "" --startLabel ""  --endLabel "" --legendLocation "upper-right" --heatmapWidth 12 
plotProfile -m "$output_dir/GM_RELB_pile.mat.gz" -out "$output_dir/GM_RELB_pile.profile.pdf" --perGroup --dpi 300 -T "" --startLabel ""  --endLabel "" --legendLocation "upper-right" --plotHeight 10 --plotWidth 12

computeMatrix reference-point --referencePoint center -S ${GM_PML} -R ${GM_both} ${GM_CO} ${GM_MO} -o "$output_dir/GM_PML_pile.mat.gz" -b 25000 -a 25000 -bs 100 --skipZeros -p $cores
plotHeatmap -m "$output_dir/GM_PML_pile.mat.gz" -out "$output_dir/GM_PML_pile.heatmap.pdf" --perGroup --colorMap "inferno" --sortUsing sum --dpi 300 -T "" --startLabel ""  --endLabel "" --legendLocation "upper-right" --heatmapWidth 12 
plotProfile -m "$output_dir/GM_PML_pile.mat.gz" -out "$output_dir/GM_PML_pile.profile.pdf" --perGroup --dpi 300 -T "" --startLabel ""  --endLabel "" --legendLocation "upper-right" --plotHeight 10 --plotWidth 12

computeMatrix reference-point --referencePoint center -S ${GM_minus} -R ${GM_MO} ${GM_both} ${GM_CO} -o "$output_dir/GM_minus_pile2.mat.gz" -b 25000 -a 25000 -bs 100 --skipZeros -p $cores
plotHeatmap -m "$output_dir/GM_minus_pile2.mat.gz" -out "$output_dir/GM_minus_pile2.heatmap.pdf" --perGroup --colorMap "inferno_r" --sortUsing sum --dpi 300 -T "" --startLabel ""  --endLabel "" --legendLocation "upper-right" --heatmapWidth 12 
plotProfile -m "$output_dir/GM_minus_pile2.mat.gz" -out "$output_dir/GM_minus_pile2.profile.pdf" --perGroup --dpi 300 -T "" --startLabel ""  --endLabel "" --legendLocation "upper-right" --plotHeight 10 --plotWidth 12

computeMatrix reference-point --referencePoint center -S ${GM_plus} -R ${GM_MO} ${GM_both} ${GM_CO} -o "$output_dir/GM_plus_pile2.mat.gz" -b 25000 -a 25000 -bs 100 --skipZeros -p $cores
plotHeatmap -m "$output_dir/GM_plus_pile2.mat.gz" -out "$output_dir/GM_plus_pile2.heatmap.pdf" --zMin 1.6 --zMax 1.8 --perGroup --colorMap "inferno" --sortUsing sum --dpi 300 -T "" --startLabel ""  --endLabel "" --legendLocation "upper-right" --heatmapWidth 12 
plotProfile -m "$output_dir/GM_plus_pile2.mat.gz" -out "$output_dir/GM_plus_pile2.profile.pdf" --perGroup --dpi 300 -T "" --startLabel ""  --endLabel "" --legendLocation "upper-right" --plotHeight 10 --plotWidth 12

computeMatrix reference-point --referencePoint center -S ${GM_merge} -R ${GM_MO} ${GM_both} ${GM_CO} -o "$output_dir/GM_merge_pile2.mat.gz" -b 25000 -a 25000 -bs 100 --skipZeros -p $cores
plotHeatmap -m "$output_dir/GM_merge_pile2.mat.gz" -out "$output_dir/GM_merge_pile2.heatmap.pdf" --perGroup --colorMap "inferno" --sortUsing sum --dpi 300 -T "" --startLabel ""  --endLabel "" --legendLocation "upper-right" --heatmapWidth 12 
plotProfile -m "$output_dir/GM_merge_pile2.mat.gz" -out "$output_dir/GM_merge_pile2.profile.pdf" --perGroup --dpi 300 -T "" --startLabel ""  --endLabel "" --legendLocation "upper-right" --plotHeight 10 --plotWidth 12
```
