## Supplementary material for "CAD-C: An engineered nuclease enables repair-free *in situ* proximity ligation and nucleosome-resolution chromosome walks in human cells": Custom code: fraglen_by_AB_v2_test_1_submit.sh

```
#!/bin/bash
#
#SBATCH --job-name=fragmat         # Job name
#SBATCH --output=fragmat_%j.out    # Standard output (%j expands to jobId)
#SBATCH --error=fragmat_%j.err     # Standard error (%j expands to jobId)
#SBATCH --cpus-per-task=4          # Number of CPU cores

# (Optional) Load modules or activate your environment if needed
# module load python/3.9
# or
# source activate <your_conda_env>

# Paths
SCRIPT_PATH="/rugpfs/fs0/risc_lab/scratch/jsoroczyns/CAD/CAD-C/scripts/2025_06_04/fraglen_matrix_by_AB_v2_20250604.py"
INPUT_FILE="/rugpfs/fs0/risc_lab/scratch/jsoroczyns/CAD/CADwalk/CADwalk_E1_GM12878_2x250/results_20250207_0034_nonexpanded/ABC_walk_processing_20250209/20250214_Bfilt_flattened_parse2_UU_direct_ABC_complete_consequent12/filtzerolenB_flip_sort_dedup_split_20250214/merged_all/20250312_cis_intersect_ChromHMM_v2/merged_all_flipped_sorted.cis.ChromHMM_v2_fraglen_seps.tsv.gz"
OUTPUT_DIR="/rugpfs/fs0/risc_lab/scratch/jsoroczyns/CAD/CADwalk/CADwalk_E1_GM12878_2x250/results_20250207_0034_nonexpanded/ABC_walk_processing_20250209/20250214_Bfilt_flattened_parse2_UU_direct_ABC_complete_consequent12/filtzerolenB_flip_sort_dedup_split_20250214/merged_all/FragLen_Analyses_2025"

# Example parameters for your script:
# (Adjust or add as needed; these flags depend on your specific script arguments)
X_MIN=0
X_MAX=2000
X_BINS=400
Y_MIN=50
Y_MAX=250
Y_BINS=100
CHUNKSIZE=5000000

echo "Starting job on `date`"

python "$SCRIPT_PATH" \
  --input "$INPUT_FILE" \
  --outdir "$OUTPUT_DIR" \
  --x-min "$X_MIN" \
  --x-max "$X_MAX" \
  --xbins "$X_BINS" \
  --y-min "$Y_MIN" \
  --y-max "$Y_MAX" \
  --ybins "$Y_BINS" \
  --chunksize "$CHUNKSIZE" \

echo "Job finished on `date`"
```
