## Supplementary material for "CAD-C: An engineered nuclease enables repair-free *in situ* proximity ligation and nucleosome-resolution chromosome walks in human cells": Custom code: fraglen_matrix_by_AB_v2_20250604.py

```
#!/usr/bin/env python3
"""
Single-pass script to generate 2D heatmaps of:
  X-axis: AB distance (bp)
  Y-axis: fragment length (bp) for A/B/C

We skip leading '#' lines, parse the header from the first non-comment line,
locate columns "AB_dist", "A_len", "B_len", "C_len", then bin each row
(A, B, C) in a single pass.

Outputs:
  1) A_counts_<TIMESTAMP>.npy
  2) B_counts_<TIMESTAMP>.npy
  3) C_counts_<TIMESTAMP>.npy
  4) meta_<TIMESTAMP>.json
  5) heatmaps_<TIMESTAMP>.pdf (with colormap "inferno")

Usage:
  python fraglen_matrix_by_AB.py \\
    --input /path/to/data.tsv.gz \\
    --outdir /path/to/output \\
    --x-min 0 --x-max 2000 --xbins 100 \\
    --y-min 0 --y-max 1000 --ybins 100 \\
    --chunksize 1000000

No medians are computed. 
If needed, you can calculate medians separately using a different script/tool.
"""

import argparse
import gzip
import json
from pathlib import Path
from time import time
from datetime import datetime

import numpy as np

import matplotlib
matplotlib.rcParams['pdf.fonttype'] = 42
matplotlib.rcParams['ps.fonttype'] = 42

import matplotlib.pyplot as plt
from matplotlib.colors import Normalize


# ------------------------------------------------------------
# Helper Functions
# ------------------------------------------------------------

def parse_cli():
    parser = argparse.ArgumentParser(
        description="Single-pass: AB distance vs. fragment length heatmaps (skip leading # lines).",
        formatter_class=argparse.ArgumentDefaultsHelpFormatter
    )
    parser.add_argument("--input", required=True, help="Path to input .tsv.gz")
    parser.add_argument("--outdir", required=True, help="Directory in which to place results")

    # X-axis
    parser.add_argument("--x-min", type=float, default=0.0, help="Min AB distance")
    parser.add_argument("--x-max", type=float, default=2000.0, help="Max AB distance")
    parser.add_argument("--xbins", type=int, default=100, help="Number of bins on X-axis")

    # Y-axis
    parser.add_argument("--y-min", type=float, default=0.0, help="Min fragment length [bp]")
    parser.add_argument("--y-max", type=float, default=1000.0, help="Max fragment length [bp]")
    parser.add_argument("--ybins", type=int, default=100, help="Number of bins on Y-axis")

    parser.add_argument("--chunksize", type=int, default=1_000_000,
                        help="Number of lines to read per chunk from the .tsv.gz")
    return parser.parse_args()


def open_skipping_comments(path):
    """
    Open the .tsv.gz, skip leading lines starting with '#',
    yield the first non-comment line as the header, then subsequent lines as data.
    """
    f = gzip.open(path, "rt")
    header = None
    for line in f:
        if not line.startswith("#"):
            # This is the header line
            header = line.strip().split("\t")
            yield ("HEADER", header)
            break
    # Now yield the rest as data lines
    for line in f:
        yield ("DATA", line)
    f.close()


def chunk_generator(path, chunksize):
    """
    Yields: 
      - ( 'HEADER', [colnames] ) once
      - Then ('DATA', [line1, line2, ...]) in chunks of up to `chunksize`.
    """
    generator = open_skipping_comments(path)
    which, first = next(generator)  # This is the header
    yield (which, first)

    chunk = []
    for (lbl, line) in generator:
        chunk.append(line)
        if len(chunk) >= chunksize:
            yield ("DATA", chunk)
            chunk = []
    if chunk:
        yield ("DATA", chunk)


def find_col_indices(header):
    """
    Return a dict with indices of "AB_dist", "A_len", "B_len", "C_len".
    Raise error if missing.
    """
    needed = ["AB_dist", "A_len", "B_len", "C_len"]
    missing = [col for col in needed if col not in header]
    if missing:
        raise ValueError(f"ERROR: Missing columns {missing} in header.")
    return {
        "AB": header.index("AB_dist"),
        "A_len": header.index("A_len"),
        "B_len": header.index("B_len"),
        "C_len": header.index("C_len"),
    }


def bin_edges(lo, hi, n):
    """Return n+1 linearly spaced bin edges from lo to hi inclusive."""
    return np.linspace(lo, hi, n + 1)


def update_counts(arr_ab, arr_len, x_edges, y_edges, counts):
    """
    Increment 2D integer array `counts` in-place for AB distance vs fragment length.
    Returns the number of increments.
    """
    # Filter by x-range
    mask = (arr_ab >= x_edges[0]) & (arr_ab <= x_edges[-1])
    if not mask.any():
        return 0

    ab_vals = arr_ab[mask]
    frag_vals = arr_len[mask]

    x_idx = np.searchsorted(x_edges, ab_vals, side="right") - 1

    # Clip fragment lengths to y-range
    frag_vals_clipped = frag_vals.clip(y_edges[0], y_edges[-1])
    y_idx = np.searchsorted(y_edges, frag_vals_clipped, side="right") - 1

    valid = (
        (x_idx >= 0) & (x_idx < len(x_edges) - 1) &
        (y_idx >= 0) & (y_idx < len(y_edges) - 1)
    )
    np.add.at(counts, (y_idx[valid], x_idx[valid]), 1)
    return valid.sum()


# ------------------------------------------------------------
# Main
# ------------------------------------------------------------

def main():
    args = parse_cli()
    t0 = time()

    # Create timestamped output directory
    timestamp_str = datetime.now().strftime("%Y%m%d_%H%M%S")
    outdir = Path(args.outdir) / timestamp_str
    outdir.mkdir(parents=True, exist_ok=True)

    # Prepare bin edges
    x_edges = bin_edges(args.x_min, args.x_max, args.xbins)
    y_edges = bin_edges(args.y_min, args.y_max, args.ybins)

    # Prepare 2D counts for A, B, C
    counts_dict = {
        "A": np.zeros((args.ybins, args.xbins), dtype=np.int64),
        "B": np.zeros((args.ybins, args.xbins), dtype=np.int64),
        "C": np.zeros((args.ybins, args.xbins), dtype=np.int64),
    }

    # Chunk generator
    cg = chunk_generator(args.input, args.chunksize)
    which, header = next(cg)
    col_idx = find_col_indices(header)

    lines_processed = 0
    increments_binned = 0

    for (lbl, chunk) in cg:
        if lbl != "DATA":
            continue
        block = np.array([line.split("\t") for line in chunk])

        ab_arr = block[:, col_idx["AB"]].astype(float)

        # Update for A
        increments_binned += update_counts(
            ab_arr, block[:, col_idx["A_len"]].astype(float),
            x_edges, y_edges, counts_dict["A"]
        )
        # Update for B
        increments_binned += update_counts(
            ab_arr, block[:, col_idx["B_len"]].astype(float),
            x_edges, y_edges, counts_dict["B"]
        )
        # Update for C
        increments_binned += update_counts(
            ab_arr, block[:, col_idx["C_len"]].astype(float),
            x_edges, y_edges, counts_dict["C"]
        )

        lines_processed += len(chunk)

    # Save the 2D matrices
    suffix = f"_{timestamp_str}"
    np.save(outdir / f"A_counts{suffix}.npy", counts_dict["A"])
    np.save(outdir / f"B_counts{suffix}.npy", counts_dict["B"])
    np.save(outdir / f"C_counts{suffix}.npy", counts_dict["C"])

    # Save metadata
    meta = {
        "input_file": str(args.input),
        "timestamp": timestamp_str,
        "lines_processed": lines_processed,
        "increments_binned": increments_binned,
        "chunksize": args.chunksize,
        "x_min": args.x_min,
        "x_max": args.x_max,
        "xbins": args.xbins,
        "y_min": args.y_min,
        "y_max": args.y_max,
        "ybins": args.ybins,
        "created_epoch_s": time()
    }
    meta_path = outdir / f"meta{suffix}.json"
    meta_path.write_text(json.dumps(meta, indent=2))

    # Plot
    import matplotlib.pyplot as plt
    from matplotlib.colors import Normalize

    fig, axes = plt.subplots(3, 1, figsize=(6, 9), sharex=True, sharey=True)
    cmap = plt.cm.inferno.copy()
    cmap.set_under("white")

    for ax, frag in zip(axes, ["A", "B", "C"]):
        arr = counts_dict[frag]
        vmax = arr.max() if arr.max() > 0 else 1
        pcm = ax.pcolormesh(
            x_edges,
            y_edges,
            arr,
            cmap=cmap,
            norm=Normalize(vmin=1, vmax=vmax)
        )
        ax.set_xlim(args.x_min, args.x_max)
        ax.set_ylim(args.y_min, args.y_max)
        ax.set_ylabel(f"{frag} length (bp)")
        ax.set_title(f"{frag} fragment")

    axes[-1].set_xlabel("AB distance (bp)")
    cbar = fig.colorbar(pcm, ax=axes, location="right", pad=0.03, shrink=0.8)
    cbar.set_label("Count per bin")

    fig.suptitle("AB distance vs. Fragment Length", y=0.95)
    fig.tight_layout(rect=[0, 0, 0.94, 0.92])

    pdf_path = outdir / f"heatmaps{suffix}.pdf"
    fig.savefig(pdf_path, bbox_inches="tight")

    # Done
    elapsed = (time() - t0) / 60
    print(f"[INFO] Completed in {elapsed:.1f} min.")
    print(f"[INFO] Lines processed: {lines_processed:,}")
    print(f"[INFO] Increments binned: {increments_binned:,}")
    print(f"[INFO] Outputs saved in: {outdir}")


if __name__ == "__main__":
    main()
```
