## Supplementary material for "CAD-C: An engineered nuclease enables repair-free *in situ* proximity ligation and nucleosome-resolution chromosome walks in human cells": Custom code: replot_ABClens_by_ABsep_wMarginals_1_20250605.py

```
#!/usr/bin/env python3
"""
heatmaps_with_marginals.py

Generate three log10-scaled AB-distance × fragment-length heat-maps
(with top & right marginal histograms) from pre-binned .npy matrices.

Author : Jan Soroczynski (Risca Lab, The Rockefeller University)
Date   : 2025-06-05
License: MIT

----------------------------------------------------------------------
INPUTS
----------------------------------------------------------------------
  • Three NumPy *.npy* files — A_counts, B_counts, C_counts
    (shape = [Y_BINS, X_BINS] = [100, 400] for the default binning)

CLI usage example
-----------------
  python heatmaps_with_marginals.py \
         --A A_counts_20250604_154313.npy \
         --B B_counts_20250604_154313.npy \
         --C C_counts_20250604_154313.npy \
         --out heatmaps_with_marginals.pdf

Dependencies
------------
  numpy ≥ 1.26
  matplotlib ≥ 3.9
"""

from __future__ import annotations
import argparse, numpy as np, matplotlib.pyplot as plt
from matplotlib.colors import LogNorm
from matplotlib.gridspec import GridSpec
from pathlib import Path

# ────────────────────────────────────────────────────────────────────
# Bin-edge constants (edit if different in future runs)
X_MIN, X_MAX, X_BINS = 0, 2000, 400
Y_MIN, Y_MAX, Y_BINS = 50, 250, 100
EDGES_X = np.linspace(X_MIN, X_MAX, X_BINS + 1)
EDGES_Y = np.linspace(Y_MIN, Y_MAX, Y_BINS + 1)

# ────────────────────────────────────────────────────────────────────
def cli() -> argparse.Namespace:
    p = argparse.ArgumentParser(
        formatter_class=argparse.ArgumentDefaultsHelpFormatter,
        description="Plot three heat-maps with marginal histograms."
    )
    p.add_argument("--A", required=True, help="A_counts.npy")
    p.add_argument("--B", required=True, help="B_counts.npy")
    p.add_argument("--C", required=True, help="C_counts.npy")
    p.add_argument("--out", default="heatmaps_with_marginals.pdf",
                   help="Output PDF filename")
    return p.parse_args()

# ────────────────────────────────────────────────────────────────────
def main() -> None:
    args = cli()

    mats = [np.load(args.A), np.load(args.B), np.load(args.C)]
    titles = ["A fragment", "B fragment", "C fragment"]

    # log10 colour scale across all three matrices
    nonzero = np.concatenate([m[m > 0] for m in mats])
    norm = LogNorm(vmin=nonzero.min(), vmax=nonzero.max())

    plt.rcParams["pdf.fonttype"] = 42
    plt.rcParams["ps.fonttype"]  = 42

    fig = plt.figure(figsize=(10, 10))
    outer = GridSpec(3, 2,
                     width_ratios=[4, 1],
                     height_ratios=[1, 1, 1],
                     wspace=0.05, hspace=0.4)

    for r, (mat, title) in enumerate(zip(mats, titles)):
        # Sub-GridSpec in the left cell: [top marginal, heat-map]
        inner = outer[r, 0].subgridspec(2, 1, height_ratios=[1, 4], hspace=0.05)

        # ── top marginal (column sum) ───────────────────────────────
        ax_col = fig.add_subplot(inner[0, 0])
        ax_col.bar(EDGES_X[:-1], mat.sum(axis=0), width=np.diff(EDGES_X),
                   align="edge", color="grey")
        ax_col.set_xlim(X_MIN, X_MAX)
        ax_col.set_xticks([])
        ax_col.set_ylabel("Count", fontsize=8)
        ax_col.set_title(title, pad=2)
        ax_col.grid(True, linestyle="--", alpha=0.3)

        # ── heat-map ────────────────────────────────────────────────
        ax_heat = fig.add_subplot(inner[1, 0], sharex=ax_col)
        pcm = ax_heat.pcolormesh(EDGES_X, EDGES_Y,
                                 np.ma.masked_where(mat == 0, mat),
                                 cmap="inferno", norm=norm, shading="auto")
        ax_heat.set_ylabel("Fragment length (bp)")
        if r == 2:
            ax_heat.set_xlabel("AB distance (bp)")
        else:
            ax_heat.set_xticks([])

        ax_heat.set_yticks(np.arange(Y_MIN, Y_MAX + 1, 50))
        ax_heat.grid(True, linestyle="--", alpha=0.3)

        # ── right marginal (row sum) ───────────────────────────────
        ax_row = fig.add_subplot(outer[r, 1], sharey=ax_heat)
        ax_row.barh(EDGES_Y[:-1], mat.sum(axis=1), height=np.diff(EDGES_Y),
                    align="edge", color="grey")
        ax_row.set_xlabel("Count", fontsize=8)
        ax_row.set_yticks([])
        ax_row.grid(True, linestyle="--", alpha=0.3)

    # shared colour-bar
    cax = fig.add_axes([0.92, 0.1, 0.02, 0.8])
    fig.colorbar(pcm, cax=cax, label="Count (log₁₀ scale)")

    fig.tight_layout(rect=[0, 0, 0.9, 1])
    Path(args.out).parent.mkdir(parents=True, exist_ok=True)
    fig.savefig(args.out, bbox_inches="tight")
    print(f"PDF written → {args.out}")

# ────────────────────────────────────────────────────────────────────
if __name__ == "__main__":
    main()
```
