## Supplementary material for "CAD-C: An engineered nuclease enables repair-free *in situ* proximity ligation and nucleosome-resolution chromosome walks in human cells": Custom code: replot_fraglen_by_AB_20250606.py

```
RE-PLOT FRAGMENT-LENGTH × AB-DISTANCE HEAT-MAPS
================================================

This stand-alone CLI script rebuilds the three-panel heat-map from the
`.npy` count matrices produced by **`fraglen_matrix_by_AB_v3*.py`**.
It *reads* bin definitions from the accompanying JSON meta-file, so the
plot is always guaranteed to match the generator’s parameters.

-----------------------------------------------------------------------
CONTENTS
-----------------------------------------------------------------------
1.  Features
2.  Installation & required packages
3.  Usage + examples
4.  CLI arguments
5.  Script source  (indented four spaces — copy verbatim to a `.py` file)

-----------------------------------------------------------------------
1.  FEATURES
-----------------------------------------------------------------------
• Fully self-contained Python 3.x                                          
• Robust argument parsing (`argparse`)                                    
• Reads `x_min/x_max/xbins` & `y_min/y_max/ybins` from meta JSON          
• Auto-discovers `A_counts*.npy`, `B_counts*.npy`, `C_counts*.npy`         
  in the same directory as the meta-file — or accepts explicit paths      
• Log-scaled or linear colour-map                                         
• Illustrator-friendly output (`pdf.fonttype = ps.fonttype = 42`)          
• Saves **both** `.pdf` and `.png` into a unique timestamped sub-folder   
• Structured logging to stdout + log-file                                 
• Embedded version metadata in PDF (`fig.text`)                            
• Author attribution exactly as requested                                 

-----------------------------------------------------------------------
2.  INSTALLATION & REQUIRED PACKAGES
-----------------------------------------------------------------------
The script depends only on NumPy, Matplotlib ≥ 3.6, and Python’s
standard library (`argparse`, `logging`, `pathlib`, `json`, `datetime`).

Recommended installation (conda):

    conda create -n plot_env python=3.11 numpy matplotlib
    conda activate plot_env

-----------------------------------------------------------------------
3.  USAGE & EXAMPLES
-----------------------------------------------------------------------
Basic use when all files are in the same directory:

    python replot_fraglen_heatmaps.py \
        --meta /path/to/results/meta_20250605_151924.json

Specify matrices explicitly & choose linear colour-scale:

    python replot_fraglen_heatmaps.py \
        --meta /.../meta.json \
        --A /.../A_counts.npy --B /.../B_counts.npy --C /.../C_counts.npy \
        --norm linear

-----------------------------------------------------------------------
4.  CLI ARGUMENTS (run with -h for full help)
-----------------------------------------------------------------------
--meta            Path to the meta_*.json produced by the generator  [REQ]
--A/--B/--C       Explicit paths to the A/B/C .npy matrices (optional)
--outdir          Directory where the timestamped sub-folder is made
--colormap        Matplotlib cmap name            [default: inferno]
--norm            linear | log                    [default: log]
--yticks          Comma-separated list of y-tick positions (bp)
--xlabel / --ylabel  Override axis labels
--dpi             PNG resolution                  [default: 300]

-----------------------------------------------------------------------
5.  SCRIPT SOURCE  (save as replot_fraglen_heatmaps.py)
--------------------------------------------------------------------
    #!/usr/bin/env python3
    """
    ────────────────────────────────────────────────────────────────────────────
    REPLOT_FRAGLEN_HEATMAPS.PY
    ────────────────────────────────────────────────────────────────────────────
    Re-plot A/B/C fragment-length × AB-distance heat-maps from .npy matrices
    using bin definitions stored in the accompanying meta JSON.
    
    AUTHOR        Jan Soroczynski, Risca Lab, The Rockefeller University
    VERSION       1.0.0
    DATE          2025-06-06
    LICENSE       MIT
    ────────────────────────────────────────────────────────────────────────────
    """

    import argparse, json, logging, sys
    from datetime import datetime
    from pathlib import Path

    import numpy as np
    import matplotlib.pyplot as plt
    from matplotlib.colors import LogNorm

    # ────────────────────────────────────────────────────────────────────────
    # CLI PARSER
    # ────────────────────────────────────────────────────────────────────────
    def build_parser() -> argparse.ArgumentParser:
        p = argparse.ArgumentParser(
            description="Re-plot fragment-length × AB-distance heat-maps "
                        "from generator .npy output.")
        p.add_argument("--meta", required=True,
                       help="Path to meta_*.json written by the generator.")
        p.add_argument("--A", help="Path to A_counts_*.npy")
        p.add_argument("--B", help="Path to B_counts_*.npy")
        p.add_argument("--C", help="Path to C_counts_*.npy")
        p.add_argument("--outdir", default=".",
                       help="Base directory for results (default: cwd).")
        p.add_argument("--colormap", default="inferno",
                       help="Matplotlib colour-map (default: inferno).")
        p.add_argument("--norm", choices=["log", "linear"], default="log",
                       help="Colour normalisation (default: log).")
        p.add_argument("--yticks",
                       help="Comma-separated custom y-tick positions (bp).")
        p.add_argument("--xlabel", default="AB distance (bp)",
                       help="Override x-axis label.")
        p.add_argument("--ylabel", default="Fragment length (bp)",
                       help="Override y-axis label.")
        p.add_argument("--dpi", type=int, default=300,
                       help="PNG resolution (default: 300).")
        return p

    # ────────────────────────────────────────────────────────────────────────
    # UTILITIES
    # ────────────────────────────────────────────────────────────────────────
    def discover_npy(meta_path: Path, explicit: str | None, letter: str) -> Path:
        if explicit:
            return Path(explicit)
        candidates = list(meta_path.parent.glob(f"{letter}_counts_*.npy"))
        if not candidates:
            raise FileNotFoundError(f"No {letter}_counts_*.npy next to {meta_path}")
        if len(candidates) > 1:
            raise RuntimeError(f"Multiple {letter}_counts_*.npy found; "
                               "specify one with --{letter}")
        return candidates[0]

    # ────────────────────────────────────────────────────────────────────────
    # MAIN
    # ────────────────────────────────────────────────────────────────────────
    def main(argv: list[str] | None = None) -> None:
        args = build_parser().parse_args(argv)

        meta_path = Path(args.meta).expanduser().resolve()
        with meta_path.open() as fh:
            meta = json.load(fh)

        # Logging
        timestamp = datetime.now().strftime("%Y%m%d_%H%M%S")
        out_base = Path(args.outdir).expanduser().resolve()
        out_dir  = out_base / f"replot_heatmaps_{timestamp}"
        out_dir.mkdir(parents=True, exist_ok=True)

        log_path = out_dir / "replot.log"
        logging.basicConfig(
            level=logging.INFO,
            format="%(asctime)s  %(levelname)s  %(message)s",
            handlers=[logging.FileHandler(log_path), logging.StreamHandler(sys.stdout)]
        )
        logging.info("Output directory: %s", out_dir)

        # Locate matrices
        A_path = discover_npy(meta_path, args.A, "A")
        B_path = discover_npy(meta_path, args.B, "B")
        C_path = discover_npy(meta_path, args.C, "C")
        logging.info("A matrix: %s", A_path.name)
        logging.info("B matrix: %s", B_path.name)
        logging.info("C matrix: %s", C_path.name)

        A = np.load(A_path)
        B = np.load(B_path)
        C = np.load(C_path)

        # Derive bin edges
        x_min, x_max, xbins = meta["x_min"], meta["x_max"], meta["xbins"]
        y_min, y_max, ybins = meta["y_min"], meta["y_max"], meta["ybins"]
        edges_x = np.linspace(x_min, x_max, xbins + 1, dtype=np.float32)
        edges_y = np.linspace(y_min, y_max, ybins + 1, dtype=np.float32)
        logging.info("Bin size X: %.2f bp, Y: %.2f bp",
                     (x_max - x_min)/xbins, (y_max - y_min)/ybins)

        # Matplotlib params for Illustrator
        plt.rcParams["pdf.fonttype"] = 42
        plt.rcParams["ps.fonttype"]  = 42

        fig, axes = plt.subplots(3, 1, figsize=(7, 9), sharex=True, sharey=True)
        cmap = plt.get_cmap(args.colormap)
        cmap.set_under("white")

        norm = LogNorm(vmin=1,
                       vmax=max(A.max(), B.max(), C.max())) if args.norm == "log" else None

        titles = ("A fragment", "B fragment", "C fragment")
        for ax, data, ttl in zip(axes, (A, B, C), titles):
            pcm = ax.pcolormesh(edges_x, edges_y, data, cmap=cmap,
                                norm=norm, shading="auto")
            ax.set_title(ttl, fontsize=11)
            ax.set_ylabel(args.ylabel)
            yticks = ([int(v) for v in args.yticks.split(",")]
                      if args.yticks else [30, 100, 200, 300, 400, 500])
            ax.set_yticks(yticks)
            ax.set_yticklabels([f"{v}" for v in yticks])
            ax.grid(False)

        axes[-1].set_xlabel(args.xlabel)

        # Colour-bar
        fig.subplots_adjust(right=0.86)
        cbar_ax = fig.add_axes([0.88, 0.15, 0.03, 0.7])
        fig.colorbar(pcm, cax=cbar_ax, label="Count / bin")

        # Metadata footer
        fig.text(0.02, 0.01,
                 f"replot_fraglen_heatmaps.py v{meta.get('version', '1.0.0')}  "
                 f"generated {timestamp}",
                 fontsize=6, alpha=0.6)

        fig.suptitle("AB distance vs. fragment length", y=0.965, fontsize=13)
        fig.tight_layout(rect=[0, 0, 0.86, 0.94])

        # Save
        pdf_path = out_dir / "heatmaps.pdf"
        png_path = out_dir / "heatmaps.png"
        fig.savefig(pdf_path, bbox_inches="tight")
        fig.savefig(png_path, bbox_inches="tight", dpi=args.dpi)
        logging.info("Saved %s and %s", pdf_path.name, png_path.name)
        logging.info("All done ✨")


    if __name__ == "__main__":
        main()
```
