## Supplementary material for "CAD-C: An engineered nuclease enables repair-free *in situ* proximity ligation and nucleosome-resolution chromosome walks in human cells": Custom code: full_100quantiles_cor_compartment_20250617.r

```
#!/usr/bin/env Rscript

# ------------------- LIBRARIES -------------------
suppressPackageStartupMessages({
  library(data.table)
  library(dplyr)
  library(ggplot2)
  library(patchwork)
})

# ------------------- OUTPUT DIRECTORY -------------------
output_dir <- "Correlation_byCompartment"
dir.create(output_dir, showWarnings = FALSE, recursive = TRUE)

# ------------------- FILE PATH -------------------
full_reference_path <- full_reference_path <- "/lustre/fs4/risc_lab/scratch/landerson01/20250205_CAD_2x250bp_GM_analysis/Compartment/Cis/merged_all_flipped_sorted.cis.compt_annotated.tsv.gz_distances_compartment.tsv.gz"

# ------------------- LOAD DATA -------------------
# Define compartment mapping
compartment_map <- function(x) {
  case_when(
    x == "cad_GM12878_A_compartments_mergedBins.bed" ~ "A",
    x == "cad_GM12878_B_compartments_mergedBins.bed" ~ "B",
    TRUE ~ NA_character_
  )
}
# Read and preprocess absolute distances
message("Reading walk data (AB and BC distances only)...")
data <- fread(full_reference_path, select = c("A_start", "B_start", "C_start", "AB_distance", "BC_distance", "A_intersect")) %>%
  mutate(AB_distance = abs(AB_distance), BC_distance = abs(BC_distance)) %>%
  mutate(A_frag = compartment_map(A_intersect))

# ------------------- SPLIT INTO WALK TYPES (using A_start, B_start, C_start) -------------------
message("Classifying walks as linear, nonlinear, and diagonal...")

data <- data %>%
  mutate(
    linear = A_start < B_start & B_start < C_start,
    diagonal = abs(log10(AB_distance) - log10(BC_distance)) < 0.2,
    walk_class = case_when(
      linear ~ "linear",
      !linear ~ "nonlinear"
    )
  ) %>%
  mutate(
    walk_class = ifelse(diagonal, "diagonal", walk_class)
  )

# ------------------- CLEAN AND BIN -------------------
message("Filtering and binning AB distances into quantiles...")
data <- data[!is.na(AB_distance) & !is.na(BC_distance) & !is.na(A_frag)]
ab_breaks <- quantile(data$AB_distance, probs = seq(0, 1, 0.01), na.rm = TRUE)
message("AB quantile bin edges:")
print(ab_breaks)
data$AB_bin <- cut(data$AB_distance, breaks = ab_breaks, include.lowest = TRUE, labels = FALSE)

# ------------------- COMPUTE OVERALL CORRELATION -------------------
message("Calculating overall Spearman and Pearson correlation...")
spearman_val <- cor(data$AB_distance, data$BC_distance, method = "spearman")
pearson_val <- cor(data$AB_distance, data$BC_distance, method = "pearson")

# ------------------- COMPUTE CORRELATION PER DECILE & BIN LABELS FOR EACH WALK TYPE -------------------
message("Calculating Spearman and Pearson correlation per AB decile...")
cor_by_decile <- data %>%
  filter(!is.na(walk_class) & !is.na(A_frag)) %>%
  group_by(walk_class, A_frag, AB_bin) %>%
  summarise(
    bin_label = paste0("(", format(min(AB_distance), scientific = FALSE, big.mark=","), " - ", format(max(AB_distance), scientific = FALSE, big.mark=","), ")"),
    bin_count = n(),
    spearman_rho = cor(AB_distance, BC_distance, method = "spearman", use = "complete.obs"),
    pearson_r = cor(AB_distance, BC_distance, method = "pearson", use = "complete.obs"),
    .groups = "drop"
  )

# ------------------- LOGGING DISTANCE RANGES AND OVERALL CORRELATION -------------------

# Export Spearman correlation by quantile to CSV
write.csv(cor_by_decile %>% select(walk_class, A_frag, AB_bin, bin_label, bin_count, spearman_rho),
          file = file.path(output_dir, "spearman_correlation_by_quantile.csv"),
          row.names = FALSE)
message("Logging quantile distance ranges and calculating overall correlation per walk_class...")
quantile_ranges <- cor_by_decile %>%
  select(walk_class, A_frag, AB_bin, bin_label, bin_count) %>%
  arrange(walk_class, AB_bin)
print(quantile_ranges)

walk_class_summary <- data %>%
  group_by(walk_class, A_frag) %>%
  summarise(
    n = n(),
    spearman_rho = cor(AB_distance, BC_distance, method = "spearman"),
    pearson_r = cor(AB_distance, BC_distance, method = "pearson"),
    .groups = "drop"
  )

print(walk_class_summary)

# ------------------- SCATTERPLOTS OF CORRELATION BY DECILE AND BIN COUNT -------------------
message("Generating scatterplots of Spearman and Pearson correlation by A_frag only (no walk_class stratification)...")
cor_by_afrag <- data %>%
  filter(!is.na(A_frag)) %>%
  group_by(A_frag, AB_bin) %>%
  summarise(
    bin_label = paste0("(", format(min(AB_distance), scientific = FALSE, big.mark=","), " - ", format(max(AB_distance), scientific = FALSE, big.mark=","), ")"),
    bin_count = n(),
    spearman_rho = cor(AB_distance, BC_distance, method = "spearman", use = "complete.obs"),
    pearson_r = cor(AB_distance, BC_distance, method = "pearson", use = "complete.obs"),
    .groups = "drop"
  )

pdf(file.path(output_dir, "correlation_by_afrag_spearman.pdf"), width = 10, height = 8)

p_all_spear <- ggplot(cor_by_afrag, aes(x = as.numeric(AB_bin), y = spearman_rho)) +
  geom_point(size = 2) +
  geom_line(group = 1) +
  facet_wrap(~ A_frag, ncol = 1, scales = "free_y") +
  labs(title = "Spearman Correlation by AB Distance Quantile (Stratified by A_frag)") +
  xlab("AB Distance Quantile") + ylab("Spearman Rho") + theme_minimal(base_size = 13)

print(p_all_spear)
dev.off()

pdf(file.path(output_dir, "correlation_by_afrag_pearson_only.pdf"), width = 10, height = 8)

p_all_pear <- ggplot(cor_by_afrag, aes(x = as.numeric(AB_bin), y = pearson_r)) +
  geom_point(size = 2) +
  geom_line(group = 1) +
  facet_wrap(~ A_frag, ncol = 1, scales = "free_y") +
  labs(title = "Pearson Correlation by AB Distance Quantile (Stratified by A_frag)") +
  theme_minimal(base_size = 13)

print(p_all_pear)
dev.off()

pdf(file.path(output_dir, "correlation_by_walkclass_and_afrag_spearman.pdf"), width = 10, height = 8)

message("Generating scatterplots of Spearman correlation by walk_class and A_frag...")
p1 <- ggplot(cor_by_decile, aes(x = as.numeric(AB_bin), y = spearman_rho)) +
  geom_point(size = 2) +
  geom_line(group = 1) +
  facet_wrap(~ walk_class + A_frag, ncol = 2, scales = "free_y") +
  labs(title = "Spearman Correlation by AB Distance Quantile (Faceted by Walk Type and A_compartment)") +
  theme_minimal(base_size = 13)
print(p1)
dev.off()

pdf(file.path(output_dir, "correlation_by_walkclass_and_afrag_pearson.pdf"), width = 10, height = 8)
p2 <- ggplot(cor_by_decile, aes(x = as.numeric(AB_bin), y = pearson_r)) +
  geom_point(size = 2) +
  geom_line(group = 1) +
  facet_wrap(~ walk_class + A_frag, ncol = 2, scales = "free_y") +
  labs(title = "Pearson Correlation by AB Distance Quantile (Faceted by Walk Type and A_compartment)") +
  theme_minimal(base_size = 13)
print(p2)
dev.off()

pdf(file.path(output_dir, "correlation_by_walkclass_and_afrag_pearson.pdf"), width = 10, height = 8)
print(p2)
dev.off()

# Additional page with bin sizes
p3 <- ggplot(cor_by_decile, aes(x = as.numeric(AB_bin), y = bin_count)) +
  geom_col() +
  facet_wrap(~ walk_class, ncol = 1, scales = "free_y") +
  labs(title = "Number of Walks per AB Distance Quantile",
       x = "AB Distance Quantile", y = "Count of Walks") +
  theme_minimal(base_size = 13)

print(p3)

# Additional page with distance range per quantile
p4 <- ggplot(cor_by_decile, aes(x = as.numeric(AB_bin), y = bin_count, label = bin_label)) +
  geom_text(size = 3) +
  facet_wrap(~ walk_class, ncol = 1, scales = "free_y") +
  labs(title = "AB Distance Ranges per Quantile Bin",
       x = "AB Distance Quantile", y = "Count of Walks") +
  theme_minimal(base_size = 13)

print(p4)
dev.off()

message("Done.")
```
