## Supplementary material for "CAD-C: An engineered nuclease enables repair-free *in situ* proximity ligation and nucleosome-resolution chromosome walks in human cells": Custom code: readme.txt

Correlation - by - Geometry
Spearman correlation calculated for linear walks (A_start < B_start < C_start), diagonal walks (|log AB distance|-|log BC distance| < 0.2), and nonlinear walks masked by the diagonal signal. Correlation calculated in 100 bins from short AB distance to long AB distance.

Spearman Correlation of Stepwise Walks Genomic separations AB_distance and BC_distance on the same walk were tested for correlation by walk classes defined by genomic geometry in R. Walks with missing AB_distance or BC_distance were removed. Diagonal walks are defined by similarity in magnitude (|log10(AB) − log10(BC)| < 0.2). Off-diagonal walks were classified either as linear if fragment start coordinates were monotonic along the chromosome (α_start < β_start < γ_start ), otherwise they were defined as nonlinear. This analysis thus masks the diagonal from off-diagonal walks. Spearman correlation was calculated using stats::cor() within 100 quantile bins of AB_distance (x-axis; 1–100) separately for each walk class using cut() with breakpoints from quantile(AB_distance, probs = seq(0,1,0.01)).For each bin, the number of walks and the corresponding AB distance range (min–max within bin) was recorded. 
