## Supplementary material for "CAD-C: An engineered nuclease enables repair-free *in situ* proximity ligation and nucleosome-resolution chromosome walks in human cells": Custom code: v10p6_bedpe_2oppCTCF_zAWB1kb_zWPI1kb_zACd_zBEI_20250927.sh

```
#!/bin/bash
#SBATCH -J metacov_bedpe_scaled_four_individual
#SBATCH -o logs/metacov_bedpe_scaled_four_individual.%j.out
#SBATCH -e logs/metacov_bedpe_scaled_four_individual.%j.err
#SBATCH -c 8

set -euo pipefail

# --- Script (v10p6) ---
SCRIPT=/rugpfs/fs0/risc_lab/scratch/jsoroczyns/CAD/CAD-C/scripts/2025_09_27/smart_bigwig_over_bed_v10p6_20250927.py

# --- BEDPE (pair‑scaled) ---
# If your BEDPE lives next to the "as_bed" version, this is the typical path:
BEDPE=/rugpfs/fs0/risc_lab/scratch/jsoroczyns/CAD/CAD-C/Mustache_filtering_CTCF_2025/bedpe_truncated_with_strand_20250405/GM12878_deepmerge_igdig1_500.mcool_mustache_loops_2500_20250405143456_2_opp_20250405_234802_trunc_stranded.bedpe
# If this file does not exist on your system, set BEDPE= to the correct path and re-run.

# --- BigWigs (four tracks) ---
BW1=/rugpfs/fs0/risc_lab/scratch/jsoroczyns/CAD/CADwalk/CADwalk_E1_GM12878_2x250/results_20250207_0034_nonexpanded/ABC_walk_processing_20250209/20250214_Bfilt_flattened_parse2_UU_direct_ABC_complete_consequent12/filtzerolenB_flip_sort_dedup_split_20250214/merged_all/WPI_AWB_analyses_2025/wpi_awb_20250927_210158/Z_AWB_1kb.bw
BW2=/rugpfs/fs0/risc_lab/scratch/jsoroczyns/CAD/CADwalk/CADwalk_E1_GM12878_2x250/results_20250207_0034_nonexpanded/ABC_walk_processing_20250209/20250214_Bfilt_flattened_parse2_UU_direct_ABC_complete_consequent12/filtzerolenB_flip_sort_dedup_split_20250214/merged_all/WPI_AWB_analyses_2025/wpi_awb_20250927_210158/Z_WPI_1kb.bw
BW3=/rugpfs/fs0/risc_lab/scratch/jsoroczyns/CAD/CADwalk/CADwalk_E1_GM12878_2x250/results_20250207_0034_nonexpanded/ABC_walk_processing_20250209/20250214_Bfilt_flattened_parse2_UU_direct_ABC_complete_consequent12/filtzerolenB_flip_sort_dedup_split_20250214/merged_all/BEI_analyses_2025/bei_20250925_211641/bigwig_20250925_220622/bei_20250925_220622_Z_ACd.bw
BW4=/rugpfs/fs0/risc_lab/scratch/jsoroczyns/CAD/CADwalk/CADwalk_E1_GM12878_2x250/results_20250207_0034_nonexpanded/ABC_walk_processing_20250209/20250214_Bfilt_flattened_parse2_UU_direct_ABC_complete_consequent12/filtzerolenB_flip_sort_dedup_split_20250214/merged_all/BEI_analyses_2025/bei_20250925_211641/bigwig_20250925_220622/bei_20250925_220622_Z_BEI.bw

# --- Output directory (unchanged per your request) ---
OUTDIR=/rugpfs/fs0/risc_lab/scratch/jsoroczyns/CAD/outputs/metacov/three_individual_scaled
mkdir -p "$OUTDIR" logs

# --- Sanity checks ---
[[ -f "$SCRIPT" ]] || { echo "[ERROR] v10p6 script not found: $SCRIPT" >&2; exit 1; }
[[ -f "$BEDPE"  ]] || { echo "[ERROR] BEDPE not found: $BEDPE" >&2; exit 1; }
[[ -f "$BW1"    ]] || { echo "[ERROR] BigWig not found: $BW1" >&2; exit 1; }
[[ -f "$BW2"    ]] || { echo "[ERROR] BigWig not found: $BW2" >&2; exit 1; }
[[ -f "$BW3"    ]] || { echo "[ERROR] BigWig not found: $BW3" >&2; exit 1; }
[[ -f "$BW4"    ]] || { echo "[ERROR] BigWig not found: $BW4" >&2; exit 1; }

NPROC=${SLURM_CPUS_PER_TASK:-8}

# Labels for legend and force‑zero selection (use basenames)
LBL1=$(basename "$BW1")
LBL2=$(basename "$BW2")
LBL3=$(basename "$BW3")
LBL4=$(basename "$BW4")

# Build args (pileup over BEDPE, pair‑scaled; 4 rows × 1 column grid = one panel per BigWig)
ARGS=(
  pileup
  -B "$BEDPE"
  -w "$BW1" "$BW2" "$BW3" "$BW4"
  -o "$OUTDIR"
  -p "$NPROC"

  # Pair‑scaled summary for loops (BEDPE)
  --bedpe-main-bins 200
  --bedpe-flank-bins 100
  --bedpe-flank-left 0.5
  --bedpe-flank-right 0.5
  # Also emit anchor midpoint pileups if you want (commented by default)
  # --anchors-separate
  # --respect-bedpe-strand

  # Grid: rows=bigwigs, cols=datasets (here just 1 BEDPE) → 4×1 layout
  --grid
  --grid-rows-axis bigwig
  --lock-y none

  # Z‑score tracks: detect as zero‑centered and force symmetric y (exactly as in v10p5 example)
  --zero-centered-detect auto
  --force-zero-centered "$LBL1" "$LBL2" "$LBL3" "$LBL4"

  # Labels; colors default to Matplotlib cycle
  --bw-labels "$LBL1" "$LBL2" "$LBL3" "$LBL4"

  # Smoothing off (keep average TSVs raw and plots unsmoothed, per your earlier job)
  --no-savgol

  # Plotting
  --plot-format pdf
  --fonttype 42
  --font-size 11 --title-size 13 --label-size 12 --tick-size 10 --legend-size 10
)

python "$SCRIPT" "${ARGS[@]}"

# Outputs:
#   • coverage_matrix_* (pair‑scaled) and coverage_average_profile_* TSVs
#   • metaplot_* PDFs per combination
#   • A 4×1 grid PDF comparing Z_AWB_1kb, Z_WPI_1kb, Z_ACd, Z_BEI over the same BEDPE set
#   • Full provenance in run_metadata_<timestamp>.txt under $OUTDIR
```
