## Supplementary material for "CAD-C: An engineered nuclease enables repair-free *in situ* proximity ligation and nucleosome-resolution chromosome walks in human cells": Custom code: GM12878_Ps_E2_P1_NNUEFS_smallmedium_dedup_20250505.sh

```
#!/bin/bash
#SBATCH --job-name=PsCurve_v2
#SBATCH --output=PsCurve_v2_%j.out
#SBATCH --error=PsCurve_v2_%j.err
#SBATCH --cpus-per-task=24          # Number of threads


# If needed, load any required modules or activate a conda environment here.
# e.g. module load python/3.9
# or source activate hic_env

# Export or define the full paths of the .mcool files as environment variables:
export MCOOL_1="/rugpfs/fs0/risc_lab/scratch/jsoroczyns/CAD/CAD-C/CADC_E2_P1_Novogene/20240822_pairtools_pipe_out/results_20240822/CADC_GM12878_1_manual_merge_20240911/cool_output/CADC_GM12878_1_merged_dedup_20240822_500_bal_500N_igdiag1.mcool"
export MCOOL_2="/rugpfs/fs0/risc_lab/scratch/jsoroczyns/CAD/CAD-C/CADC_E2_P3_NNUEFS/20241128_pairtools_pipe_out/results_20241128/20241130_dedup_NNUEFS_merged/cooler_out_20250410_155412/CADC_E2_NNUEFS_S1234_dedup_merge_20241201_500_bal_500N_igdiag1.mcool"
export MCOOL_3="/rugpfs/fs0/risc_lab/scratch/jsoroczyns/CAD/CADwalk/CADwalk_E1_GM12878_2x250/results_20250207_0034_nonexpanded/small_medium_dedup_merge_20250208/cooler_out_20250208_165259/small_medium_nonexpanded_20250207_0034_dedup_merge_20250208_500_bal_500N_igdiag1.mcool"

# Run the v2 script with the 6 .mcool paths, passing aliases exactly.
python /rugpfs/fs0/risc_lab/scratch/jsoroczyns/CAD/CAD-C/scripts/2025_04_16/mcool_Ps_curve_v2_20250416.py \
  --mcool "$MCOOL_1" "$MCOOL_2" "$MCOOL_3"\
  --aliases \
    "CAD-C GM12878 P1 sonicated (dedup)" \
    "CAD-C GM12878 P2 NNUEFS (dedup)" \
    "CADwalk GM12878 2x250 Small+Medium (dedup) " \
  --binsize 500 \
  --threads 24 \
  --output-dir /rugpfs/fs0/risc_lab/scratch/jsoroczyns/CAD/CAD-C/Ps_curves_2025/

# End of SBATCH  
```
