## Supplementary material for "CAD-C: An engineered nuclease enables repair-free *in situ* proximity ligation and nucleosome-resolution chromosome walks in human cells": Custom code: Ps_curve_replot_v1_20250416.py

```
#!/usr/bin/env python3
"""
Ps_curve_replot.py
==================
Author:  Jan Soroczynski, Risca Lab, The Rockefeller University
Date:    2025-04-19
Version: 1.2.5  • Binning+Smoothing on full curve, then apply x-range, matching April-11 script
License: MIT

DESCRIPTION
-----------
Re‑plot genome‑wide P(s) curves from one or more TSV files that have
'dist' and 'ps' columns (e.g., from the original April‑11 pipeline).

**Default**: uses exactly the same binning (nbins=200) and Savitzky–Golay (window=7)
as the April‑11 script, binning & smoothing the entire curve first,
then applying the user’s x‑range (–x-min/–x-max).

To override binning or smoothing, pass:
  • --nbins  <int>  (0=skip binning) 
  • --savgol <int>  (0=skip smoothing)

ORDER OF ACTIONS
----------------
  1) Load the entire (dist, ps) data from each TSV.
  2) Bin in log‑space (default 200 bins).
  3) Apply Savitzky–Golay smoothing (default 7).
  4) THEN apply the x‑range filter to the smoothed bin centers.
  5) Compute log10 slope, plot log(P(s)) vs. log(dist) plus slope.

USAGE
-----
  python Ps_curve_replot.py \
      --tsv file1.tsv file2.tsv \
      --aliases Sample1 Sample2 \
      [--nbins 200] [--savgol 7] \
      [--x-min 500] [--x-max 1e8] \
      [--y-min ...] [--y-max ...] \
      --output-dir results_dir

OUTPUT
------
Creates a timestamped subfolder in --output-dir:
  • Ps_replot_<ISO8601>/Ps_replot_<ISO8601>.png
  • Ps_replot_<ISO8601>/Ps_replot_<ISO8601>.pdf
  • Ps_replot_<ISO8601>/metadata.yaml
"""

import argparse
import hashlib
import sys
from datetime import datetime, UTC
from pathlib import Path
from typing import Sequence

import matplotlib.pyplot as plt
import numpy as np
import pandas as pd
import yaml
from scipy.signal import savgol_filter


def geometric_bin_average(dist: np.ndarray, ps: np.ndarray, nbins: int):
    """
    Bin 'dist' and 'ps' into nbins equal log-width bins over the full range
    and return (centers, means).
    """
    edges = np.logspace(np.log10(dist.min()), np.log10(dist.max()), nbins + 1)
    idx   = np.digitize(dist, edges) - 1
    means = np.full(nbins, np.nan)
    for i in range(nbins):
        in_bin = (idx == i)
        if np.any(in_bin):
            means[i] = ps[in_bin].mean()
    # Geometric midpoint for bin i
    centers = np.sqrt(edges[:-1] * edges[1:])
    valid   = ~np.isnan(means)
    return centers[valid], means[valid]


def log_slope(x: np.ndarray, y: np.ndarray) -> np.ndarray:
    """
    Return the slope d[log10(y)] / d[log10(x)] by finite difference.
    """
    return np.gradient(np.log10(y), np.log10(x))


def get_colors(n: int) -> list[str]:
    """
    Provide a color palette for n lines. If n=2, use a high-contrast pair.
    """
    if n == 2:
        return ["#22B14C", "#C000C0"]
    try:
        import seaborn as sns
        return sns.color_palette("mako", n).as_hex()
    except ImportError:
        # fallback to default rc cycle
        return plt.rcParams["axes.prop_cycle"].by_key()["color"][:n]


def parse_cli(argv: Sequence[str]):
    """
    CLI: we default nbins=200, savgol=7, x_min=500, x_max=1e8
    to replicate the original April‑11 pipeline.
    """
    p = argparse.ArgumentParser(description="Bin+Smooth full P(s), then mask x-range, replicating April-11.")
    p.add_argument("--tsv",        nargs="+", required=True,
                   help="TSV files containing columns 'dist' and 'ps' (or synonyms).")
    p.add_argument("--aliases",    nargs="+", required=True,
                   help="Legend labels, same count as --tsv.")
    p.add_argument("--x-min",      type=float, default=5e2,
                   help="Lower bound for the final plotted x-range. Default=500.")
    p.add_argument("--x-max",      type=float, default=1e8,
                   help="Upper bound for the final plotted x-range. Default=1e8.")
    p.add_argument("--nbins",      type=int,   default=200,
                   help="Log-space bins (0=skip binning). Default=200 (April-11).")
    p.add_argument("--savgol",     type=int,   default=7,
                   help="S-G window (odd≥5) to smooth. 0=skip. Default=7 (April-11).")
    p.add_argument("--y-min",      type=float,
                   help="Manual lower limit on P(s) axis. Auto if omitted.")
    p.add_argument("--y-max",      type=float,
                   help="Manual upper limit on P(s) axis. Auto if omitted.")
    p.add_argument("--output-dir", required=True,
                   help="Parent directory for a Ps_replot_<timestamp> folder of outputs.")
    p.add_argument("--dpi",        type=int,   default=300,
                   help="PNG DPI. Default=300.")
    return p.parse_args(argv)


def main(argv: Sequence[str] | None = None):
    args = parse_cli(argv or sys.argv[1:])
    if len(args.tsv) != len(args.aliases):
        sys.exit("[ERROR] The number of --tsv files differs from --aliases.")

    # Create output folder
    timestamp_utc = datetime.now(UTC).strftime("%Y%m%dT%H%M%SZ")
    out_dir = Path(args.output_dir).expanduser().resolve() / f"Ps_replot_{timestamp_utc}"
    out_dir.mkdir(parents=True, exist_ok=False)

    # Minimal matplot styling
    plt.rcParams.update({
        "axes.grid": True,
        "grid.linestyle": "--",
        "grid.linewidth": 0.5,
        "grid.alpha": 0.3,
    })

    fig, (ax_ps, ax_sl) = plt.subplots(2, 1, figsize=(6, 7), sharex=True, height_ratios=[2, 1])
    ax_ps.set_axisbelow(True)
    ax_sl.set_axisbelow(True)

    colors = get_colors(len(args.tsv))
    global_min_ps = np.inf
    global_min_slope = np.inf

    # Process each dataset
    for idx, (tsv_file, alias, color) in enumerate(zip(args.tsv, args.aliases, colors)):
        # 1) Load + standardize columns
        df = pd.read_csv(tsv_file, sep="\t", compression="infer", comment="#", header=0)
        df.columns = (
            df.columns.astype(str)
              .str.strip()
              .str.lower()
              .str.replace(r"^distance(_bp|s)?$", "dist", regex=True)
              .str.replace(
                  r"^(ps|p\(s\)|contact(_frequency|_freq)?|balanced\.avg(\.smoothed\.agg)?)$",
                  "ps", regex=True
              )
        )
        if "dist" not in df.columns and df.index.name is None:
            df = df.reset_index().rename(columns={"index": "dist"})
        if not {"dist", "ps"}.issubset(df.columns):
            sys.exit(f"[ERROR] {alias}: Missing 'dist' or 'ps' in {tsv_file}.")

        dist = df["dist"].values
        ps   = df["ps"].values
        if dist.size < 2:
            print(f"[WARNING] {alias}: only {dist.size} points total. Skipping.")
            continue

        # 2) Bin on FULL curve
        if args.nbins > 0:
            centers, ps_proc = geometric_bin_average(dist, ps, args.nbins)
        else:
            centers, ps_proc = dist, ps
        if centers.size < 2:
            print(f"[WARNING] {alias}: after binning, <2 points remain. Skipping.")
            continue

        # 3) Smooth on FULL 200-bin curve
        if (args.savgol >= 5) and (args.savgol % 2 == 1) and (args.savgol < len(ps_proc)):
            ps_proc = savgol_filter(ps_proc, args.savgol, polyorder=3, mode="interp")

        # 4) NOW apply x-range mask
        mask = (centers >= args.x_min) & (centers <= args.x_max)
        centers, ps_proc = centers[mask], ps_proc[mask]
        if centers.size < 2:
            print(f"[WARNING] {alias}: after x-range, <2 points. Skipping.")
            continue

        # 5) Compute slope, track global minima
        slope_vals = log_slope(centers, ps_proc)
        global_min_ps     = min(global_min_ps, ps_proc.min())
        global_min_slope  = min(global_min_slope, slope_vals.min())

        # Plot top: loglog(P(s))
        ax_ps.loglog(centers, ps_proc, "-", lw=1.5, color=color, label=alias, zorder=2+idx)
        # Plot bottom: slope
        ax_sl.semilogx(centers, slope_vals, "-", lw=1.5, color=color, zorder=2+idx)

    # X-limits
    ax_ps.set_xlim(args.x_min, args.x_max)

    # Y-limits for P(s) panel
    if args.y_min is None and np.isfinite(global_min_ps) and global_min_ps > 0:
        # one decade below global min
        auto_y_min = 10 ** (np.floor(np.log10(global_min_ps)) - 1)
        args.y_min = auto_y_min
    if args.y_max is not None:
        ax_ps.set_ylim(args.y_min, args.y_max)
    else:
        ax_ps.set_ylim(bottom=args.y_min)

    # Slope panel: bottom=lowest slope, top=0
    if np.isfinite(global_min_slope):
        ax_sl.set_ylim(global_min_slope, 0)
    else:
        ax_sl.set_ylim(-3, 0)

    ax_ps.set_ylabel("P(s)")
    ax_sl.set_ylabel("Slope")
    ax_sl.set_xlabel("Separation (bp)")
    ax_ps.legend(frameon=False, fontsize="small")
    fig.tight_layout(h_pad=0.15)

    # Save figure (PNG + PDF)
    png_path = out_dir / f"Ps_replot_{timestamp_utc}.png"
    pdf_path = out_dir / f"Ps_replot_{timestamp_utc}.pdf"
    fig.savefig(png_path, dpi=args.dpi)
    fig.savefig(pdf_path, dpi=args.dpi, backend="pdf", metadata={"Title": pdf_path.stem})
    print(f"[INFO] Plots → {png_path}\n             {pdf_path}")

    # Write metadata
    meta = {
        "timestamp_utc": timestamp_utc,
        "inputs": dict(zip(args.aliases, map(str, args.tsv))),
        "x_min":   args.x_min,
        "x_max":   args.x_max,
        "y_min":   float(args.y_min) if args.y_min is not None else None,
        "y_max":   float(args.y_max) if args.y_max is not None else None,
        "nbins":   args.nbins,
        "savgol":  args.savgol,
        "png":     str(png_path),
        "pdf":     str(pdf_path),
        "python":  sys.version.split()[0],
        "numpy":   np.__version__,
        "pandas":  pd.__version__,
        "matplotlib": plt.matplotlib.__version__,
        "script_sha256": hashlib.sha256(Path(__file__).read_bytes()).hexdigest(),
    }
    meta_file = out_dir / "metadata.yaml"
    with open(meta_file, "w") as f:
        yaml.safe_dump(meta, f, sort_keys=False)
    print(f"[INFO] Metadata → {meta_file}")


if __name__ == "__main__":
    main()
```
