## Supplementary material for "CAD-C: An engineered nuclease enables repair-free *in situ* proximity ligation and nucleosome-resolution chromosome walks in human cells": Custom code: Ps_curve_replot_v1_RPE_set_1_2_20250416.sh

```
#!/bin/bash
#SBATCH --job-name=Ps_replot_RPE_20250416
#SBATCH --output=Ps_replot_RPE_20250416_%j.out
#SBATCH --error=Ps_replot_RPE_20250416_%j.err
#SBATCH --cpus-per-task=2
set -euo pipefail
set -x                                    # ⇦ prints every command

# ── Paths ────────────────────────────────────────────────────────────────────
ORIG_DIR="/rugpfs/fs0/risc_lab/scratch/jsoroczyns/CAD/CAD-C/Ps_curves_2025/20250416_134340"
REPLOT_SCRIPT="/rugpfs/fs0/risc_lab/scratch/jsoroczyns/CAD/CAD-C/scripts/2025_04_16/Ps_curve_replot_v1_20250416.py"

TSV_DAR="$ORIG_DIR/Darrow2016_20241125_dedup_final_merged_500_bal_500N_igdiag2_Ps.tsv"
TSV_AO="$ORIG_DIR/AO_MicroC_WT12and34_20241126_dedup_final_merge_500_bal_500N_igdiag1_Ps.tsv"
TSV_CAD="$ORIG_DIR/CADC_E1_P2andP3_20241126_dedup_final_merged_500_bal_500N_igdiag1_Ps.tsv"

OUT_ROOT="$ORIG_DIR/Replots"
mkdir -p "$OUT_ROOT"

# ── Run ──────────────────────────────────────────────────────────────────────
python "$REPLOT_SCRIPT" \
  --tsv "$TSV_DAR" "$TSV_AO" "$TSV_CAD" \
  --aliases "Darrow 2016 in situ Hi-C" "AO Micro-C (4xBRs)" "CAD-C" \
  --output-dir "$OUT_ROOT" \
  --x-min 500 --x-max 1e8        \
  --dpi 300

# Copy exit status explicitly so SLURM records it
exit $?
```
