## Supplementary material for "CAD-C: An engineered nuclease enables repair-free *in situ* proximity ligation and nucleosome-resolution chromosome walks in human cells": Custom code: Ps_curve_v1_RPE_set_1_1_20250416.sh

```
#!/bin/bash
#SBATCH --job-name=ps_curve_RPE_set_1_1     # Job name (shown by squeue)
#SBATCH --output=ps_curve_RPE_set_1_1_%j.out # Standard output (%j = JobID)
#SBATCH --error=ps_curve_RPE_set_1_1_%j.err  # Standard error
#SBATCH --cpus-per-task=16                    # Number of CPU cores requested

# ---------------------------------------------------------------------------
# 1) Activate your environment with `cooler`, `cooltools`, etc.
#    (Adjust these lines to match your HPC setup.)
#
# module load python/3.11
# source activate hic-env
# ---------------------------------------------------------------------------

# 2) Path to the P(s) Python driver (unchanged)
SCRIPT_PATH="/rugpfs/fs0/risc_lab/scratch/jsoroczyns/CAD/CAD-C/scripts/2025_04_11/mcool_Ps_curve_v1_20250411.py"

# 3) Input .mcool files (RPE‑1 datasets)
MCOOL_1="/rugpfs/fs0/risc_lab/scratch/jsoroczyns/CAD/CAD-C/Public_Datasets/Darrow_2016/20241125_merged_pairs/20241126_igdiag2_coolers/Darrow2016_20241125_dedup_final_merged_500_bal_500N_igdiag2.mcool"
MCOOL_2="/rugpfs/fs0/risc_lab/scratch/jsoroczyns/CAD/CAD-C/AO_MicroC_WT12and34_merge/20241126_igdiag1_coolers/AO_MicroC_WT12and34_20241126_dedup_final_merge_500_bal_500N_igdiag1.mcool"
MCOOL_3="/rugpfs/fs0/risc_lab/scratch/jsoroczyns/CAD/CAD-C/CADC_E1_P2andP3_merge/20241126_igdiag1_coolers/CADC_E1_P2andP3_20241126_dedup_final_merged_500_bal_500N_igdiag1.mcool"

# 4) Matching aliases (appear in the legend & filenames)
ALIAS_1="Darrow 2016 in situ Hi-C"
ALIAS_2="AO Micro-C (4xBRs)"
ALIAS_3="CAD-C"

# 5) General options
OUTDIR="/rugpfs/fs0/risc_lab/scratch/jsoroczyns/CAD/CAD-C/Ps_curves_2025"  # Will auto‑timestamp inside
GENOME_BUILD="hg38"                                                         # Passed straight through
THREADS=16                                                                    # Matches --cpus-per-task

# ---------------------------------------------------------------------------
# 6) Run the script
#    - --mcool lists all .mcool files in order
#    - --aliases lists the user‑friendly names (must match count & order)
#    - --threads sets nproc for cooltools.expected_cis
#    - --output-dir is where the driver will create its timestamped subfolder
# ---------------------------------------------------------------------------
python "$SCRIPT_PATH" \
  --mcool "$MCOOL_1" "$MCOOL_2" "$MCOOL_3" \
  --aliases "$ALIAS_1" "$ALIAS_2" "$ALIAS_3" \
  --threads "$THREADS" \
  --genome-build "$GENOME_BUILD" \
  --output-dir "$OUTDIR"

# Notes:
#  • If all three .mcool files share 500 bp resolution, the script will use 500 bp.
#    Override with `--binsize 1000` (for example) if desired.
#  • Adjust SBATCH memory/time directives above as appropriate for your cluster.
#  • The driver auto‑generates a unique timestamped subfolder within $OUTDIR,
#    ensuring each run’s outputs remain separate.
```
