## Supplementary material for "CAD-C: An engineered nuclease enables repair-free *in situ* proximity ligation and nucleosome-resolution chromosome walks in human cells": Custom code: Fig2b_compartment_strength.py

```
# compartment strength

import numpy as np

s = np.load("GM12878_CADC_500_igdig1_600m_10kb_saddle_E1.saddledump.npz")
s = np.load("GM12878_Dovetail_MicroC_500_igdig1_600m_10kb_saddle_E1.saddledump.npz")
s = np.load("GM12878_Hi-C_500_igdig1_600m_10kb_saddle_E1.saddledump.npz")
s = np.load("GM12878_microc_500_igdig1_600m_10kb_saddle_E1.saddledump.npz")
s = np.load("GM12878_OmniC_500_igdig1_600m_10kb_saddle_E1.saddledump.npz")

S = s["saddlecounts"]   # saddle matrix

# top/bottom 10%， 5 bins
k = 5
top_idx = np.arange(52-k, 52)
bot_idx = np.arange(0, k)
AA = S[np.ix_(top_idx, top_idx)].mean()
BB = S[np.ix_(bot_idx, bot_idx)].mean()
AB = S[np.ix_(top_idx, bot_idx)].mean()
comp_strength = (AA + BB) / (2 * AB)
print(comp_strength)
# GM12878_CADC: 2.7918696362564237
# GM12878_Dovetail_MicroC: 2.0454068655284225
# GM12878_Hi-C: 1.6253021849917817
# GM12878_microc: 2.714499824539436
# GM12878_OmniC: 1.90208231972275

# ≈1.0：almost no compartment
# 1.2–1.5：weak compartment
# 1.5–3.0：typical differentiated cells
# >3：very strong compartment

# 
s = np.load("RPE1_CADC_500_igdig1_500m_expected_10kb_saddle_E1.saddledump.npz")
s = np.load("RPE1_microc_500_igdig1_500m_expected_10kb_saddle_E1.saddledump.npz")
s = np.load("RPE1_Darrow2016_500_igdig1_500m_expected_10kb_saddle_E1.saddledump.npz")

S = s["saddlecounts"]   # saddle matrix

# top/bottom 10%， 5 bins
k = 5
top_idx = np.arange(52-k, 52)
bot_idx = np.arange(0, k)
AA = S[np.ix_(top_idx, top_idx)].mean()
BB = S[np.ix_(bot_idx, bot_idx)].mean()
AB = S[np.ix_(top_idx, bot_idx)].mean()
comp_strength = (AA + BB) / (2 * AB)
print(comp_strength)

# RPE1_CADC: 5.3276968278571735
# RPE1_microc: 2.8437051271382416
# RPE1_Darrow2016: 1.8888176157477983
```
