## Supplementary material for "CAD-C: An engineered nuclease enables repair-free *in situ* proximity ligation and nucleosome-resolution chromosome walks in human cells": Custom code: Fig2d_boundary_strength.sh

```
#!/bin/bash
#SBATCH --job-name=HiC_Processing
#SBATCH --output=hic_processing_%j.log
#SBATCH --cpus-per-task=8
#SBATCH --mem=32G
#SBATCH --time=24:00:00
#SBATCH --partition=hpc_a10_a

source /programs/biogrids.shrc
cores=8

# bed files for boundary groups
GM12878_CAD_DMicroC_common='/rugpfs/fs0/risc_lab/scratch/wzuo/project/CAD/insulations/20251116_insulation/GM12878_CAD_DMicroC_common_10kb.bed'
GM12878_CAD_DMicroC_CAD_only='/rugpfs/fs0/risc_lab/scratch/wzuo/project/CAD/insulations/20251116_insulation/GM12878_CAD_DMicroC_CAD_only_10kb.bed'
GM12878_CAD_DMicroC_DMicroC_only='/rugpfs/fs0/risc_lab/scratch/wzuo/project/CAD/insulations/20251116_insulation/GM12878_CAD_DMicroC_DMicroC_only_10kb.bed'

GM12878_CAD_HiC_common='/rugpfs/fs0/risc_lab/scratch/wzuo/project/CAD/insulations/20251116_insulation/GM12878_CAD_HiC_common_10kb.bed'
GM12878_CAD_HiC_CAD_only='/rugpfs/fs0/risc_lab/scratch/wzuo/project/CAD/insulations/20251116_insulation/GM12878_CAD_HiC_CAD_only_10kb.bed'
GM12878_CAD_HiC_HiC_only='/rugpfs/fs0/risc_lab/scratch/wzuo/project/CAD/insulations/20251116_insulation/GM12878_CAD_HiC_HiC_only_10kb.bed'

GM12878_CAD_MicroC_common='/rugpfs/fs0/risc_lab/scratch/wzuo/project/CAD/insulations/20251116_insulation/GM12878_CAD_MicroC_common_10kb.bed'
GM12878_CAD_MicroC_CAD_only='/rugpfs/fs0/risc_lab/scratch/wzuo/project/CAD/insulations/20251116_insulation/GM12878_CAD_MicroC_CAD_only_10kb.bed'
GM12878_CAD_MicroC_MicroC_only='/rugpfs/fs0/risc_lab/scratch/wzuo/project/CAD/insulations/20251116_insulation/GM12878_CAD_MicroC_MicroC_only_10kb.bed'

GM12878_CAD_OmniC_common='/rugpfs/fs0/risc_lab/scratch/wzuo/project/CAD/insulations/20251116_insulation/GM12878_CAD_OmniC_common_10kb.bed'
GM12878_CAD_OmniC_CAD_only='/rugpfs/fs0/risc_lab/scratch/wzuo/project/CAD/insulations/20251116_insulation/GM12878_CAD_OmniC_CAD_only_10kb.bed'
GM12878_CAD_OmniC_OmniC_only='/rugpfs/fs0/risc_lab/scratch/wzuo/project/CAD/insulations/20251116_insulation/GM12878_CAD_OmniC_OmniC_only_10kb.bed'

RPE1_CAD_HiC_common='/rugpfs/fs0/risc_lab/scratch/wzuo/project/CAD/insulations/20251116_insulation/RPE1_CAD_HiC_common_10kb.bed'
RPE1_CAD_HiC_CAD_only='/rugpfs/fs0/risc_lab/scratch/wzuo/project/CAD/insulations/20251116_insulation/RPE1_CAD_HiC_CAD_only_10kb.bed'
RPE1_CAD_HiC_HiC_only='/rugpfs/fs0/risc_lab/scratch/wzuo/project/CAD/insulations/20251116_insulation/RPE1_CAD_HiC_HiC_only_10kb.bed'

RPE1_CAD_MicroC_common='/rugpfs/fs0/risc_lab/scratch/wzuo/project/CAD/insulations/20251116_insulation/RPE1_CAD_MicroC_common_10kb.bed'
RPE1_CAD_MicroC_CAD_only='/rugpfs/fs0/risc_lab/scratch/wzuo/project/CAD/insulations/20251116_insulation/RPE1_CAD_MicroC_CAD_only_10kb.bed'
RPE1_CAD_MicroC_MicroC_only='/rugpfs/fs0/risc_lab/scratch/wzuo/project/CAD/insulations/20251116_insulation/RPE1_CAD_MicroC_MicroC_only_10kb.bed'

# bigwg files
GM12878_CADC="/rugpfs/fs0/risc_lab/scratch/wzuo/project/CAD/insulations/20251116_insulation/GM12878_CADC_10kb_TAD_sorted_fixed.bw"
GM12878_microc="/rugpfs/fs0/risc_lab/scratch/wzuo/project/CAD/insulations/20251116_insulation/GM12878_microc_10kb_TAD_sorted_fixed.bw"
GM12878_HiC="/rugpfs/fs0/risc_lab/scratch/wzuo/project/CAD/insulations/20251116_insulation/GM12878_Hi-C_10kb_TAD_sorted_fixed.bw"
GM12878_OmniC="/rugpfs/fs0/risc_lab/scratch/wzuo/project/CAD/insulations/20251116_insulation/GM12878_OmniC_10kb_TAD_sorted_fixed.bw"
GM12878_Dovetail_MicroC="/rugpfs/fs0/risc_lab/scratch/wzuo/project/CAD/insulations/20251116_insulation/GM12878_Dovetail_MicroC_10kb_TAD_sorted_fixed.bw"    

RPE1_CADC="/rugpfs/fs0/risc_lab/scratch/wzuo/project/CAD/insulations/20251116_insulation/RPE1_CADC_10kb_TAD_sorted_fixed.bw"
RPE1_microc="/rugpfs/fs0/risc_lab/scratch/wzuo/project/CAD/insulations/20251116_insulation/RPE1_microc_10kb_TAD_sorted_fixed.bw"
RPE1_HiC="/rugpfs/fs0/risc_lab/scratch/wzuo/project/CAD/insulations/20251116_insulation/RPE1_Darrow2016_10kb_TAD_sorted_fixed.bw"

computeMatrix reference-point --referencePoint center -S ${RPE1_CADC} ${RPE1_microc} -R ${RPE1_CAD_MicroC_MicroC_only} ${RPE1_CAD_MicroC_common} ${RPE1_CAD_MicroC_CAD_only} -o RPE1_CAD_MicroC_TAD.gz  -a 200000 -b 200000 -bs 10000
plotProfile -m RPE1_CAD_MicroC_TAD.gz --perGroup -out RPE1_CAD_MicroC_TAD.pdf

computeMatrix reference-point --referencePoint center -S ${RPE1_CADC} ${RPE1_HiC} -R ${RPE1_CAD_HiC_HiC_only} ${RPE1_CAD_HiC_common} ${RPE1_CAD_HiC_CAD_only} -o RPE1_CAD_HiC_TAD.gz  -a 200000 -b 200000 -bs 10000
plotProfile -m RPE1_CAD_HiC_TAD.gz --perGroup -out RPE1_CAD_HiC_TAD.pdf

computeMatrix reference-point --referencePoint center -S ${GM12878_CADC} ${GM12878_microc} -R ${GM12878_CAD_MicroC_MicroC_only} ${GM12878_CAD_MicroC_common} ${GM12878_CAD_MicroC_CAD_only} -o GM12878_CAD_MicroC_TAD.gz  -a 200000 -b 200000 -bs 10000
plotProfile -m GM12878_CAD_MicroC_TAD.gz --perGroup -out GM12878_CAD_MicroC_TAD.pdf

computeMatrix reference-point --referencePoint center -S ${GM12878_CADC} ${GM12878_HiC} -R ${GM12878_CAD_HiC_HiC_only} ${GM12878_CAD_HiC_common} ${GM12878_CAD_HiC_CAD_only} -o GM12878_CAD_HiC_TAD.gz  -a 200000 -b 200000 -bs 10000
plotProfile -m GM12878_CAD_HiC_TAD.gz --perGroup -out GM12878_CAD_HiC_TAD.pdf

computeMatrix reference-point --referencePoint center -S ${GM12878_CADC} ${GM12878_OmniC} -R ${GM12878_CAD_OmniC_OmniC_only} ${GM12878_CAD_OmniC_common} ${GM12878_CAD_OmniC_CAD_only} -o GM12878_CAD_OmniC_TAD.gz  -a 200000 -b 200000 -bs 10000
plotProfile -m GM12878_CAD_OmniC_TAD.gz --perGroup -out GM12878_CAD_OmniC_TAD.pdf

computeMatrix reference-point --referencePoint center -S ${GM12878_CADC} ${GM12878_Dovetail_MicroC} -R ${GM12878_CAD_DMicroC_DMicroC_only} ${GM12878_CAD_DMicroC_common} ${GM12878_CAD_DMicroC_CAD_only} -o GM12878_CAD_DMicroC_TAD.gz  -a 200000 -b 200000 -bs 10000
plotProfile -m GM12878_CAD_DMicroC_TAD.gz --perGroup -out GM12878_CAD_DMicroC_TAD.pdf
```
