## Supplementary material for "CAD-C: An engineered nuclease enables repair-free *in situ* proximity ligation and nucleosome-resolution chromosome walks in human cells": Custom code: Fig2i_loop_pileup_GM12878.sh

```
#!/bin/bash
#SBATCH --job-name=Pileup
#SBATCH --output=Pileup_%j.log
#SBATCH --cpus-per-task=14
#SBATCH --mem=32G
#SBATCH --time=24:00:00
#SBATCH --partition=hpc_l40_a

#conda activate coolpup
source ~/.bashrc_soft2
conda activate open2c_v3

# the first batch of the downsampled data
GM_CAD='/rugpfs/fs0/risc_lab/scratch/wzuo/project/CAD/GM12878_dataset/subsample/GM12878_CADC_500_igdig1_4b.mcool'
GM_MicroC='/rugpfs/fs0/risc_lab/scratch/wzuo/project/CAD/GM12878_dataset/subsample/GM12878_microc_500_igdig1_4b.mcool'

GM12878_CADC_MicroC_5kb='/rugpfs/fs0/risc_lab/scratch/wzuo/project/CAD/GM12878_dataset/loops/loop_2500/GM12878_CADC_MicroC_2500_loop_5kb_slop2.bedpe'
GM12878_CADC_only_5kb='/rugpfs/fs0/risc_lab/scratch/wzuo/project/CAD/GM12878_dataset/loops/loop_2500/GM12878_CADC_only_2500_loop_5kb_slop2.bedpe'
GM12878_MicroC_only_5kb='/rugpfs/fs0/risc_lab/scratch/wzuo/project/CAD/GM12878_dataset/loops/loop_2500//GM12878_MicroC_only_2500_loop_5kb_slop2.bedpe'

python /rugpfs/fs0/risc_lab/scratch/wzuo/CAD/loops_subgroup/loops_pileup/pileup_wrapa_coolpuppy_v13_20250929.py -c ${GM_CAD} ${GM_MicroC} -r ${GM12878_MicroC_only_5kb} ${GM12878_CADC_MicroC_5kb} ${GM12878_CADC_only_5kb} \
    -o GM_CAD_MicroC_2500_loop_5kb_slop_pileup --binsize 1000 --flank 40000 --cooler-labels "GM_CAD" "GM_MicroC" --region-labels "MicroC_only" "Common" "CAD_only" --threads 14
```
