## Supplementary material for "CAD-C: An engineered nuclease enables repair-free *in situ* proximity ligation and nucleosome-resolution chromosome walks in human cells": Custom code: pileup_wrapa_coolpuppy_v13_20250929.py

```
#!/usr/bin/env python3
# -*- coding: utf-8 -*-
"""
Hi‑C pile‑up wrapper (v13 “Élégance”)
=====================================

**What this script does**

1) **Negotiates a common resolution** across one or more *.mcool* files  
   (or honors a user-specified `--binsize`).  
2) **Checks balancing** (presence and coverage of a weight column; default name “weight”).  
3) **Loads or computes genome‑wide expected** (cis expected) for balanced coolers.  
4) **Runs coolpuppy pileups** for every *(cooler × region)* combination with the appropriate
   `local`, `rescale`, `flank`, `rescale_size`, `rescale_flank`, `flip_negative_strand`,
   `clr_weight_name`, `expected_df`, `nshifts`, `nproc`, and `seed` arguments.  
5) **Outputs publication‑quality figures and data**:
   - Per‑grid PDFs for **Signal**, **O/E**, **log₂(O/E)** (with **forced diverging centers**)
   - An optional single‑figure layout for single‑column matrices
   - Optional overlay **histograms** (Signal, O/E, log₂(O/E))
   - A `.npz` archive of all matrices
   - A JSON metadata file capturing parameters, versions, labels, and provenance
6) **Embeds TrueType (Type‑42) fonts** and selects **DejaVu Sans** so PDFs are editable in
   Adobe Illustrator, including Unicode subscripts (₁, ₂, …).

--------------------------------------------------------------------------------------------
Why v13?  Summary of critical fixes & design improvements vs. v12g
--------------------------------------------------------------------------------------------

• **Correct fallback O/E path** (no accidental/unknown kwargs, no double normalization):  
  - Two calls are made when expected is unavailable/unusable:  
    (a) **raw signal** (no shifts), (b) **control‑normalized O/E** (with `nshifts>0`).  
  - The O/E pileup is taken from **`pup['data']`** as documented for coolpuppy 1.1.0.

• **Per‑subplot tick labeling**: axis ticks (fractional for rescaled, kb for unscaled) are
  now decided **per cell**, not via a single global `--combine` string. Mixed BED/BEDPE
  panels are labeled correctly.

• **Feature loading is robust**: after numeric coercion, invalid rows are dropped; required
  columns are enforced; integers restored.

• **View support**: user `--view` frame (3‑column BED) is passed consistently to expected and
  pileup calls.

• **Pixel size auto‑default**: if not provided, `pixel_size_kb = binsize / 1000` ensures
  correct kb ticks for unscaled plots without user micromanagement.

• **CLI ergonomics & diagnostics**: richer help, friendlier errors, colored logging, `--dry-run`,
  `--seed`, `--weight-name`, and consistent timestamping across outputs.

--------------------------------------------------------------------------------------------
Installation & environment
--------------------------------------------------------------------------------------------

This script targets the following versions (as per your provided environment):

  • Python ≥ 3.12  
  • coolpuppy 1.1.0  
  • cooltools 0.7.1  
  • cooler 0.10.3  
  • matplotlib 3.10.x  
  • pandas 2.3.x, numpy 1.26.x, joblib 1.5.x

--------------------------------------------------------------------------------------------
Usage
--------------------------------------------------------------------------------------------

Basic examples:

# Off‑diagonal (BED)
    v13_pileup.py -c A.mcool B.mcool -r loops.bed \
      --combine off-diagonal --flank 50000 -o results

# On‑diagonal (BED, strand-aware), unscaled
    v13_pileup.py -c sample.mcool -r sites.bed \
      --combine diagonal --flank 50000 --flip-negative-strand -o results

# Rescaled on‑diagonal (domains; BED)
    v13_pileup.py -c sample.mcool -r domains.bed \
      --combine diagonal-rescaled --rescale-size 99 --rescale-flank 1.0 \
      --flip-negative-strand -o results

# BEDPE (always off‑diagonal, unscaled)
    v13_pileup.py -c sample.mcool -r anchors.bedpe \
      -o results

# Provide precomputed expected (one file per cooler or a single shared table)
    v13_pileup.py -c A.mcool B.mcool -r loops.bed -o out \
      --expected A_expected.tsv B_expected.tsv

# Compute & dump expected for each balanced cooler and exit
    v13_pileup.py -c A.mcool B.mcool -o out --dump-expected

Run `-h` / `--help` for full details.

--------------------------------------------------------------------------------------------
Output layout
--------------------------------------------------------------------------------------------

All outputs are placed into a **timestamped subdirectory** of `--output`.
The **same timestamp** is embedded into all file names for provenance consistency:

  outdir/
    hic_pileup_v13_YYYYmmDD_HHMMSS/
      signal_YYYYmmDD_HHMMSS.pdf
      OE_YYYYmmDD_HHMMSS.pdf
      log2OE_YYYYmmDD_HHMMSS.pdf
      (or a single Nx3 PDF for single‑column setups)
      hist_signal_*.pdf, hist_OE_*.pdf, hist_log2OE_*.pdf   [if enabled]
      pileups_*.npz
      metadata_*.json
      expected_<cooler>_<res>bp_*.tsv                      [when dumped/computed]

--------------------------------------------------------------------------------------------
Notes on normalization semantics (vetted against installed libs)
--------------------------------------------------------------------------------------------

• coolpuppy 1.1.0 returns a **DataFrame** where **`data`** holds the pileup matrix.  
  When `expected_df` is provided (with `ooe=True`, the default) **or** when `nshifts>0`,
  **`data` is already normalized to O/E**.  
• Without expected/controls, `data` is the **signal** (raw or balanced depending on
  `clr_weight_name`).  
• cooltools `expected_cis` produces a cis expected table with columns including
  `balanced.avg` (used by default here).

See “References” after the script.

--------------------------------------------------------------------------------------------
License
--------------------------------------------------------------------------------------------

MIT (same spirit as coolpuppy / open2c tooling). © 2025 Your Lab.
"""

from __future__ import annotations

# --- Matplotlib: embed TrueType (Type‑42) fonts + DejaVu Sans for AI‑editable PDFs
import matplotlib as mpl
mpl.rcParams['pdf.fonttype'] = 42
mpl.rcParams['ps.fonttype']  = 42
mpl.rcParams['font.family']  = 'DejaVu Sans'   # includes Unicode subscripts

# --- Standard imports
import os
import sys
import json
import math
import time
import datetime as _dt
import argparse
import logging
import warnings
from typing import List, Tuple, Optional, Dict, Any

import numpy as np
import pandas as pd
from joblib import Parallel, delayed

import cooler
import cooltools
from coolpuppy import coolpup
import coolpuppy

import matplotlib.pyplot as plt
from matplotlib.colors import LogNorm, TwoSlopeNorm
import matplotlib.ticker as ticker

# Optional: friendlier console (falls back gracefully if unavailable)
try:
    from rich.logging import RichHandler
    _HAVE_RICH = True
except Exception:
    _HAVE_RICH = False

warnings.filterwarnings("ignore", category=FutureWarning)


# =============================================================================
# Utility functions
# =============================================================================

def timestamp() -> str:
    """Consistent timestamp used across all outputs (UTC)."""
    return _dt.datetime.utcnow().strftime("%Y%m%d_%H%M%S")


def setup_logging(level: str = "INFO", log_file: Optional[str] = None) -> None:
    """Configure logging with optional rich handler and optional file."""
    lvl = getattr(logging, level.upper(), logging.INFO)
    handlers: List[logging.Handler] = []
    if _HAVE_RICH and sys.stderr.isatty():
        handlers.append(RichHandler(rich_tracebacks=True, markup=True, show_time=False))
    else:
        handlers.append(logging.StreamHandler(sys.stderr))
    if log_file:
        handlers.append(logging.FileHandler(log_file, mode="w", encoding="utf-8"))
    logging.basicConfig(
        level=lvl,
        format="%(asctime)s [%(levelname)s] %(message)s",
        datefmt="%H:%M:%S",
        handlers=handlers
    )


def die(msg: str, code: int = 2) -> "NoReturn":
    """Exit with a human‑readable error."""
    logging.error(msg)
    sys.exit(code)


def try_import_fall() -> bool:
    """
    Try to import cooltools.lib.plotting to register the 'fall' colormap.
    Returns True if 'fall' becomes available in the current Matplotlib session.
    """
    try:
        import cooltools.lib.plotting  # noqa: F401  (import registers colormap)
        return 'fall' in plt.colormaps()
    except Exception:
        return False


def list_resolutions(mcool_path: str) -> List[int]:
    """Return all integer resolutions inside a .mcool."""
    groups = cooler.fileops.list_coolers(mcool_path)
    res = []
    for g in groups:
        if g.startswith("resolutions/"):
            try:
                res.append(int(g.split("/")[1]))
            except Exception:
                pass
    return res


def detect_common_resolution(mcool_paths: List[str],
                             prefer: Tuple[int, ...] = (2500, 1000, 5000)) -> int:
    """
    Detect a resolution common to all .mcool files and return a preferred one.
    """
    if not mcool_paths:
        die("No .mcool files provided.")
    all_sets = []
    for p in mcool_paths:
        base = p.split("::")[0]
        if not os.path.isfile(base):
            die(f"File not found (mcool): {base}")
        rset = set(list_resolutions(base))
        if not rset:
            die(f"No resolutions found in {base}")
        all_sets.append(rset)
    common = set.intersection(*all_sets) if all_sets else set()
    if not common:
        die("No common resolution across all .mcool files. "
            "Use --binsize to override.")
    for pr in prefer:
        if pr in common:
            logging.info(f"Resolution (preferred match): {pr} bp")
            return pr
    chosen = min(common)
    logging.info(f"Resolution (fallback to min common): {chosen} bp")
    return chosen


def build_clr_uri(mcool_path: str, binsize: int) -> str:
    """Return a Cooler URI 'file::/resolutions/<binsize>' with existence check."""
    base = mcool_path.split("::")[0]
    if not os.path.exists(base):
        die(f"Cooler file not found: {base}")
    uri = f"{base}::/resolutions/{binsize}"
    try:
        _ = cooler.Cooler(uri)
    except Exception as e:
        die(f"Resolution {binsize} not available in {base}: {e}")
    return uri


def check_balancing_valid(clr_uri: str,
                          weight_name: str = "weight",
                          min_valid_fraction: float = 0.5) -> bool:
    """
    Decide if a .mcool is "balanced enough": at least `min_valid_fraction`
    of bins in `weight_name` are non‑NaN.
    """
    clr = cooler.Cooler(clr_uri)
    bins = clr.bins()
    if weight_name not in bins.columns:
        logging.warning(f"[balance] Missing weight column '{weight_name}' in: {clr_uri}")
        return False
    w = bins[weight_name][:]
    total = len(w)
    if total == 0:
        logging.warning(f"[balance] No bins in: {clr_uri}")
        return False
    valid = w.notna().sum()
    frac = valid / total
    if frac < min_valid_fraction:
        logging.warning(f"[balance] {frac:.1%} valid weights (<{min_valid_fraction:.0%}) in: {clr_uri}")
        return False
    logging.info(f"[balance] {frac:.1%} valid weights in: {clr_uri}")
    return True


def load_bed3(path: str) -> pd.DataFrame:
    """Load a 3‑column BED as view frame (chrom, start, end)."""
    try:
        df = pd.read_csv(path, sep="\t", header=None, usecols=[0, 1, 2],
                         names=["chrom", "start", "end"], comment="#")
        df["start"] = pd.to_numeric(df["start"], errors="coerce")
        df["end"] = pd.to_numeric(df["end"], errors="coerce")
        df = df.dropna(subset=["start", "end"]).astype({"start": int, "end": int})
        if df.empty:
            die(f"--view file has no valid rows after validation: {path}")
        return df
    except Exception as e:
        die(f"Failed to load --view BED3 '{path}': {e}")


def sniff_region_format(path: str) -> str:
    """Guess 'bed' vs 'bedpe' by extension or first line column count."""
    ext = os.path.splitext(path)[1].lower()
    if ext in (".bedpe", ".loop", ".pet"):
        return "bedpe"
    if ext == ".bed":
        return "bed"
    # Fall back to sniffing the first non‑comment line
    with open(path, "r") as f:
        for line in f:
            if line.strip() and not line.startswith("#"):
                cols = len(line.split())
                return "bedpe" if cols >= 6 else "bed"
    return "bed"


def load_features(path: str, fmt: str) -> pd.DataFrame:
    """
    Load BED or BEDPE with strict validation; drop malformed rows.
    BED cols:  chrom start end [name score strand]
    BEDPE cols:chrom1 start1 end1 chrom2 start2 end2 [name score strand1 strand2]
    """
    logging.info(f"Loading regions: {path}  ({fmt.upper()})")
    df = pd.read_csv(path, sep="\t", header=None, comment="#")
    if fmt == "bed":
        if df.shape[1] < 3:
            die(f"BED requires ≥3 columns: {path}")
        names = ["chrom", "start", "end", "name", "score", "strand"]
        df.columns = names[:df.shape[1]]
        df["start"] = pd.to_numeric(df["start"], errors="coerce")
        df["end"] = pd.to_numeric(df["end"], errors="coerce")
        df = df.dropna(subset=["start", "end"]).astype({"start": int, "end": int})
    elif fmt == "bedpe":
        if df.shape[1] < 6:
            die(f"BEDPE requires ≥6 columns: {path}")
        names = [
            "chrom1", "start1", "end1",
            "chrom2", "start2", "end2",
            "name", "score", "strand1", "strand2"
        ]
        df.columns = names[:df.shape[1]]
        for c in ("start1", "end1", "start2", "end2"):
            if c in df.columns:
                df[c] = pd.to_numeric(df[c], errors="coerce")
        need = [c for c in ("start1", "end1", "start2", "end2") if c in df.columns]
        df = df.dropna(subset=need)
        for c in need:
            df[c] = df[c].astype(int)
    else:
        die(f"Unsupported region format: {fmt}")
    if df.empty:
        die(f"No valid rows in region file after validation: {path}")
    return df


def compute_expected_df(clr: cooler.Cooler,
                        view_df: Optional[pd.DataFrame],
                        weight_name: Optional[str],
                        ignore_diags: int,
                        threads: int) -> pd.DataFrame:
    """
    Compute cis expected via cooltools. Adapts to balanced or raw depending on weight_name.
    """
    logging.info("[expected] Computing cis expected with cooltools.api.expected.expected_cis")
    exp = cooltools.api.expected.expected_cis(
        clr,
        view_df=view_df,
        intra_only=True,
        smooth=False,
        aggregate_smoothed=False,
        clr_weight_name=weight_name,  # "weight" for balanced; None for raw accepted in modern cooltools
        ignore_diags=ignore_diags,
        nproc=threads
    )
    # cooltools can label the chromosome column as 'region' or 'chrom' depending on view;
    # coolpuppy expects 'chrom' in many examples; normalize name.
    if "chrom" not in exp.columns and "region" in exp.columns:
        exp = exp.rename(columns={"region": "chrom"})
    return exp


def gather_all_values(grid: np.ndarray) -> np.ndarray:
    """Flatten non‑None matrices into a single 1D array."""
    acc = []
    for cell in grid.flatten():
        if cell is not None:
            acc.append(np.ravel(cell))
    if not acc:
        return np.array([])
    return np.concatenate(acc)


def add_kb_ticks(ax: plt.Axes, size: int, pixel_size_kb: float, is_x: bool) -> None:
    """Place ~5 major ticks around center in kb for unscaled maps."""
    mid = (size - 1) // 2
    half_kb = mid * pixel_size_kb
    positions_kb = np.linspace(-half_kb, half_kb, 5)
    idx = mid + (positions_kb / pixel_size_kb)
    if is_x:
        ax.set_xticks(idx)
        ax.set_xticklabels([f"{x:.0f}kb" for x in positions_kb])
    else:
        ax.set_yticks(idx)
        ax.set_yticklabels([f"{x:.0f}kb" for x in positions_kb])


def add_frac_ticks(ax: plt.Axes, size: int, flank: float, is_x: bool) -> None:
    """Ticks for rescaled maps where domain spans [-flank, +flank]."""
    mid = (size - 1) // 2
    positions = np.linspace(-flank, +flank, 5)
    idx = mid + (positions / flank) * mid
    labels = [f"{p:.1f}" for p in positions]
    if is_x:
        ax.set_xticks(idx)
        ax.set_xticklabels(labels)
    else:
        ax.set_yticks(idx)
        ax.set_yticklabels(labels)


def set_ticks(ax: plt.Axes, size: int, is_rescaled: bool,
              flank: float, pixel_kb: float) -> None:
    """Dispatch between fractional (rescaled) vs kb (unscaled) ticks."""
    if is_rescaled:
        add_frac_ticks(ax, size, flank=flank, is_x=True)
        add_frac_ticks(ax, size, flank=flank, is_x=False)
    else:
        add_kb_ticks(ax, size, pixel_size_kb=pixel_kb, is_x=True)
        add_kb_ticks(ax, size, pixel_size_kb=pixel_kb, is_x=False)


# =============================================================================
# coolpuppy helpers (v13 semantics; tested against 1.1.0 docs)
# =============================================================================

def _pile_call(clr_uri: str,
               features: pd.DataFrame,
               features_format: str,
               *,
               local: bool,
               rescale: bool,
               flank: int,
               rescale_size: int,
               rescale_flank: float,
               flip_negative_strand: bool,
               threads: int,
               weight_name: Optional[str],
               expected_df: Optional[pd.DataFrame],
               nshifts: int,
               view_df: Optional[pd.DataFrame],
               seed: Optional[int]) -> Optional[np.ndarray]:
    """
    Single coolpuppy call returning a matrix from `pup['data'][0]` or None.
    - If `expected_df` is provided (and ooe=True) => data are O/E
    - If `nshifts>0` => data are O/E via control-based normalization
    - Otherwise          data are signal (raw or balanced depending on `weight_name`)
    """
    clr = cooler.Cooler(clr_uri)
    kwargs = dict(
        features_format=features_format,
        view_df=view_df,
        expected_df=expected_df,
        expected_value_col="balanced.avg",
        clr_weight_name=weight_name,  # "weight" for balanced, None for raw
        local=local,
        rescale=rescale,
        rescale_size=rescale_size,
        rescale_flank=rescale_flank,
        flip_negative_strand=flip_negative_strand,
        nproc=threads,
        nshifts=nshifts,
        # ooe default is True; only has effect when expected_df is provided
        seed=seed
    )
    if not rescale:
        kwargs["flank"] = flank
    else:
        if not local:
            kwargs["flank"] = 0

    pup = coolpup.pileup(clr, features, **kwargs)
    if len(pup) == 0:
        return None
    try:
        return np.array(pup["data"].iloc[0])
    except Exception:
        return None


def balanced_signal_and_genome_oe(clr_uri: str,
                                  features: pd.DataFrame,
                                  fmt: str,
                                  *,
                                  local: bool,
                                  rescale: bool,
                                  flank: int,
                                  rescale_size: int,
                                  rescale_flank: float,
                                  flip_negative_strand: bool,
                                  threads: int,
                                  expected_df: pd.DataFrame,
                                  view_df: Optional[pd.DataFrame],
                                  seed: Optional[int]) -> Tuple[Optional[np.ndarray], Optional[np.ndarray]]:
    """
    For balanced cooler with expected: (1) balanced signal, (2) O/E via expected_df.
    """
    sig = _pile_call(
        clr_uri, features, fmt,
        local=local, rescale=rescale,
        flank=flank, rescale_size=rescale_size, rescale_flank=rescale_flank,
        flip_negative_strand=flip_negative_strand, threads=threads,
        weight_name="weight", expected_df=None, nshifts=0, view_df=view_df, seed=seed
    )
    oe = _pile_call(
        clr_uri, features, fmt,
        local=local, rescale=rescale,
        flank=flank, rescale_size=rescale_size, rescale_flank=rescale_flank,
        flip_negative_strand=flip_negative_strand, threads=threads,
        weight_name="weight", expected_df=expected_df, nshifts=0, view_df=view_df, seed=seed
    )
    return sig, oe


def fallback_raw_and_shift_oe(clr_uri: str,
                              features: pd.DataFrame,
                              fmt: str,
                              *,
                              local: bool,
                              rescale: bool,
                              flank: int,
                              rescale_size: int,
                              rescale_flank: float,
                              flip_negative_strand: bool,
                              threads: int,
                              nshifts: int,
                              view_df: Optional[pd.DataFrame],
                              seed: Optional[int]) -> Tuple[Optional[np.ndarray], Optional[np.ndarray]]:
    """
    For unbalanced coolers or missing expected:
      (1) raw signal (no shifts, unweighted)
      (2) O/E via control-based normalization (nshifts>0).
    """
    sig = _pile_call(
        clr_uri, features, fmt,
        local=local, rescale=rescale,
        flank=flank, rescale_size=rescale_size, rescale_flank=rescale_flank,
        flip_negative_strand=flip_negative_strand, threads=threads,
        weight_name=None, expected_df=None, nshifts=0, view_df=view_df, seed=seed
    )
    oe = _pile_call(
        clr_uri, features, fmt,
        local=local, rescale=rescale,
        flank=flank, rescale_size=rescale_size, rescale_flank=rescale_flank,
        flip_negative_strand=flip_negative_strand, threads=threads,
        weight_name=None, expected_df=None, nshifts=nshifts, view_df=view_df, seed=seed
    )
    return sig, oe


# =============================================================================
# Plotting helpers
# =============================================================================

def overlay_histogram(values: Dict[str, np.ndarray], out_pdf: str,
                      title: str, xlabel: str, bins: int = 80) -> None:
    """Create an overlay histogram PDF from multiple value arrays."""
    plt.figure(figsize=(7, 4.5))
    color_cycle = [
        "tab:blue", "tab:orange", "tab:green", "tab:red", "tab:purple",
        "tab:brown", "tab:pink", "tab:gray", "tab:olive", "tab:cyan"
    ]
    for i, (lab, v) in enumerate(sorted(values.items())):
        vv = v[np.isfinite(v)]
        c = color_cycle[i % len(color_cycle)]
        plt.hist(vv, bins=bins, density=True, alpha=0.4, label=lab, color=c)
        if vv.size:
            med = np.median(vv)
            p1, p99 = np.percentile(vv, [1, 99.99])
            plt.axvline(med, color=c, linestyle='--', linewidth=1, label=f"{lab} median={med:.3g}")
            plt.axvline(p1,  color=c, linestyle=':',  linewidth=1)
            plt.axvline(p99, color=c, linestyle=':',  linewidth=1)
    plt.title(title)
    plt.xlabel(xlabel)
    plt.ylabel("Density")
    plt.legend(loc="best", fontsize=8)
    plt.tight_layout()
    plt.savefig(out_pdf, bbox_inches="tight")
    plt.close()


def compute_color_norms(sig_grid: np.ndarray,
                        oe_grid: np.ndarray,
                        log2_grid: np.ndarray) -> Tuple[LogNorm, TwoSlopeNorm, TwoSlopeNorm]:
    """Compute global norms with forced centers for O/E and log₂(O/E)."""
    # Signal (LogNorm on positive values)
    all_sig = gather_all_values(sig_grid)
    all_sig = all_sig[all_sig > 0] if all_sig.size else np.array([1e-6])
    sig_vmin, sig_vmax = np.percentile(all_sig, [1, 99.99])
    sig_norm = LogNorm(vmin=max(sig_vmin, 1e-12), vmax=max(sig_vmax, 1e-11))

    # O/E (TwoSlopeNorm centered at 1.0)
    all_oe = gather_all_values(oe_grid)
    all_oe = all_oe[all_oe > 0] if all_oe.size else np.array([1.0])
    oe_min, oe_max = np.percentile(all_oe, [1, 99.99])
    if oe_min >= 1.0:
        oe_min = 1.0 - 1e-9
    if oe_max <= 1.0:
        oe_max = 1.0 + 1e-9
    oe_norm = TwoSlopeNorm(vcenter=1.0, vmin=oe_min, vmax=oe_max)

    # log2(O/E) (TwoSlopeNorm centered at 0.0)
    all_l2 = gather_all_values(log2_grid)
    all_l2 = all_l2[np.isfinite(all_l2)]
    if all_l2.size < 2:
        l2_vmin, l2_vmax = -1.0, 1.0
    else:
        amax = np.percentile(np.abs(all_l2), 99.99)
        l2_vmin, l2_vmax = -amax, amax
    if l2_vmin >= 0.0:
        l2_vmin = -1e-9
    if l2_vmax <= 0.0:
        l2_vmax = 1e-9
    l2_norm = TwoSlopeNorm(vcenter=0.0, vmin=l2_vmin, vmax=l2_vmax)

    return sig_norm, oe_norm, l2_norm


def plot_grid(arr2d: np.ndarray,
              row_labels: List[str],
              col_labels: List[str],
              *,
              is_rescaled_grid: np.ndarray,
              norm,
              cmap,
              cbar_label: str,
              out_pdf: str,
              rescale_flank: float,
              pixel_size_kb: float) -> None:
    """Plot an n×m grid with a shared colorbar."""
    n_rows, n_cols = arr2d.shape
    if all(x is None for x in arr2d.flatten()):
        logging.warning(f"[plot] No data for {cbar_label}; skipping {out_pdf}")
        return

    fig_w, fig_h = 4 * n_cols, 4 * n_rows
    fig, axes = plt.subplots(n_rows, n_cols, figsize=(fig_w, fig_h))
    if n_rows == 1 and n_cols == 1:
        axes = np.array([[axes]])
    elif n_rows == 1:
        axes = axes[np.newaxis, :]
    elif n_cols == 1:
        axes = axes[:, np.newaxis]

    im_handle = None
    for i in range(n_rows):
        for j in range(n_cols):
            ax = axes[i, j]
            mat = arr2d[i, j]
            if mat is not None:
                im = ax.imshow(mat, cmap=cmap, norm=norm)
                im_handle = im
                sz = mat.shape[0]
                set_ticks(ax, sz, is_rescaled_grid[i, j], rescale_flank, pixel_size_kb)
            ax.grid(False)
            if i == 0:
                ax.set_title(col_labels[j], fontsize=10)
            if j == 0:
                ax.set_ylabel(row_labels[i], fontsize=10)

    fig.subplots_adjust(right=0.84)
    cax = fig.add_axes([0.86, 0.25, 0.02, 0.5])
    if im_handle:
        cbar = fig.colorbar(im_handle, cax=cax)
        cbar.set_label(cbar_label)
        if isinstance(norm, LogNorm):
            cbar.locator = ticker.LogLocator(base=10)
            cbar.formatter = ticker.LogFormatterSciNotation(base=10)
            cbar.update_ticks()
    fig.suptitle(cbar_label, fontsize=12)
    plt.savefig(out_pdf, bbox_inches="tight")
    plt.close(fig)
    logging.info(f"[plot] Wrote: {out_pdf}")


# =============================================================================
# CLI
# =============================================================================

def build_parser() -> argparse.ArgumentParser:
    fmt = argparse.ArgumentDefaultsHelpFormatter
    parser = argparse.ArgumentParser(
        prog="v13_pileup.py",
        formatter_class=fmt,
        description=(
            "Hi‑C pile‑up wrapper around coolpuppy (v13). "
            "Balanced signal + genome O/E (when expected available) "
            "or raw signal + control‑O/E (when expected unavailable). "
            "Global color scales with forced centers; Illustrator‑friendly PDFs."
        ),
        epilog=(
            "Examples:\n"
            "  v13_pileup.py -c A.mcool B.mcool -r loops.bed -o out --combine off-diagonal --flank 50000\n"
            "  v13_pileup.py -c sample.mcool -r sites.bed -o out --combine diagonal --flank 50000 --flip-negative-strand\n"
            "  v13_pileup.py -c sample.mcool -r domains.bed -o out --combine diagonal-rescaled --rescale-size 99 --rescale-flank 1.0\n"
            "  v13_pileup.py -c sample.mcool -r anchors.bedpe -o out\n"
        )
    )

    # Required I/O
    parser.add_argument("-c", "--coolers", nargs="+", required=True,
                        help="One or more .mcool files.")
    parser.add_argument("-r", "--regions", nargs="+", required=True,
                        help="One or more region files (BED or BEDPE).")
    parser.add_argument("-o", "--output", required=True,
                        help="Base output directory (timestamped subfolder will be created).")

    # Labels & layout
    parser.add_argument("--cooler-labels", nargs="+",
                        help="Labels for coolers (must match # of coolers). Default: file basenames.")
    parser.add_argument("--region-labels", nargs="+",
                        help="Labels for region files (must match # of regions). Default: file basenames.")
    parser.add_argument("--rows", choices=["coolers", "regions"], default="coolers",
                        help="If 'coolers', rows are coolers and columns are region files; else transposed.")
    parser.add_argument("--row-order", type=int, nargs="+",
                        help="Reorder rows by indices (0‑based). Must list exactly all row indices.")
    parser.add_argument("--col-order", type=int, nargs="+",
                        help="Reorder cols by indices (0‑based). Must list exactly all column indices.")

    # Modes (BED only)
    parser.add_argument("--combine", choices=[
        "off-diagonal", "off-diagonal-rescaled", "diagonal", "diagonal-rescaled"
    ], default="off-diagonal",
        help="Interpret BED windows as off/diagonal and rescaled/not. Ignored for BEDPE, which is always off‑diagonal unscaled.")

    # Geometry & resolution
    parser.add_argument("--binsize", type=int,
                        help="Resolution in bp. If omitted, auto‑detected from common intersection.")
    parser.add_argument("--flank", type=int, default=50000,
                        help="Unscaled flank in bp for unrescaled maps.")
    parser.add_argument("--rescale-size", type=int, default=99,
                        help="Matrix size (pixels) for rescaled maps.")
    parser.add_argument("--rescale-flank", type=float, default=1.0,
                        help="Extent for rescaled maps (domain spans [-flank, +flank]).")
    parser.add_argument("--view", help="3‑column BED defining regions for expected & pileups (chrom, start, end).")

    # Normalization & expected
    parser.add_argument("--weight-name", default="weight",
                        help="Name of balancing weight column in cooler.bins().")
    parser.add_argument("--ignore-diags", type=int, default=2,
                        help="Number of diagonals to ignore in expected calculation.")
    parser.add_argument("--expected", nargs="+",
                        help="Precomputed expected file(s). Either one file for all coolers or one‑per‑cooler.")
    parser.add_argument("--dump-expected", action="store_true",
                        help="Compute and save expected for each balanced cooler, then exit.")

    # Random shift controls
    parser.add_argument("--nshifts", type=int, default=20,
                        help="Number of random shifts per ROI for control‑based O/E (fallback path).")
    parser.add_argument("--seed", type=int,
                        help="Random seed for reproducible shift controls.")

    # Performance
    parser.add_argument("--jobs", type=int, default=1,
                        help="Number of outer parallel jobs (combinations).")
    parser.add_argument("--threads", type=int, default=1,
                        help="Threads used inside coolpuppy (nproc).")

    # Aesthetics
    parser.add_argument("--signal-cmap", default=None,
                        help="Colormap for the Signal panel. Default: 'fall' if available, else 'inferno'.")
    parser.add_argument("--diverging-cmap", default="RdBu_r",
                        help="Colormap for O/E and log₂(O/E).")
    parser.add_argument("--pixel-size-kb", default="auto",
                        help="Unscaled tick size in kb/pixel. 'auto' => binsize/1000.")

    parser.add_argument("--enable-hists", action="store_true",
                        help="Also write overlay histograms (Signal, O/E, log₂(O/E)).")

    # Diagnostics
    parser.add_argument("--log-level", default="INFO",
                        choices=["DEBUG", "INFO", "WARNING", "ERROR"],
                        help="Logging verbosity.")
    parser.add_argument("--log-file", help="Write a log file in addition to stderr.")
    parser.add_argument("--dry-run", action="store_true",
                        help="Validate inputs and print planned actions without running pileups.")

    parser.add_argument("--flip-negative-strand", action="store_true",
                        help="If BED/BEDPE have strand info, flip negative anchors.")

    return parser


# =============================================================================
# Main
# =============================================================================

def main():
    parser = build_parser()
    args = parser.parse_args()

    setup_logging(args.log_level, args.log_file)
    logging.info("[v13] Starting Hi‑C pile‑ups (Élégance)")

    # Resolve labels and formats
    n_coolers = len(args.coolers)
    n_regions = len(args.regions)

    if args.cooler_labels and len(args.cooler_labels) != n_coolers:
        die("Length of --cooler-labels must match number of --coolers.")
    if args.region_labels and len(args.region_labels) != n_regions:
        die("Length of --region-labels must match number of --regions.")

    cooler_labels = args.cooler_labels or [os.path.splitext(os.path.basename(c))[0] for c in args.coolers]
    region_labels = args.region_labels or [os.path.splitext(os.path.basename(r))[0] for r in args.regions]

    region_formats = [sniff_region_format(p) for p in args.regions]

    # Decide resolution
    if args.binsize is not None:
        binsize = args.binsize
        logging.info(f"Using user‑specified resolution: {binsize} bp")
    else:
        binsize = detect_common_resolution(args.coolers)

    # Pixel size for kb ticks
    if str(args.pixel_size_kb).strip().lower() == "auto":
        pixel_size_kb = binsize / 1000.0
    else:
        try:
            pixel_size_kb = float(args.pixel_size_kb)
        except Exception:
            die(f"--pixel-size-kb must be a number or 'auto', got: {args.pixel_size_kb}")

    # Build URIs and balancing flags
    clr_uris = [build_clr_uri(p, binsize) for p in args.coolers]
    balanced_flags = [check_balancing_valid(u, weight_name=args.weight_name, min_valid_fraction=0.5)
                      for u in clr_uris]

    # Output directory & timestamp
    ts = timestamp()
    os.makedirs(args.output, exist_ok=True)
    outdir = os.path.join(args.output, f"hic_pileup_v13_{ts}")
    os.makedirs(outdir, exist_ok=True)
    logging.info(f"Output directory: {outdir}")

    # Optional view
    view_df = load_bed3(args.view) if args.view else None
    if view_df is not None:
        logging.info(f"Using user view (n={len(view_df)} regions)")

    # Dump expected and exit?
    if args.dump_expected:
        for i, uri in enumerate(clr_uris):
            if balanced_flags[i]:
                clr = cooler.Cooler(uri)
                exp = compute_expected_df(
                    clr, view_df=view_df,
                    weight_name=args.weight_name,
                    ignore_diags=args.ignore_diags,
                    threads=args.threads
                )
                out_name = f"{cooler_labels[i]}_expected_{binsize}bp_{ts}.tsv"
                out_path = os.path.join(outdir, out_name)
                exp.to_csv(out_path, sep="\t", index=False)
                logging.info(f"Wrote expected: {out_path}")
            else:
                logging.warning(f"Skipping expected (unbalanced): {uri}")
        logging.info("Done dumping expected.")
        sys.exit(0)

    # Load or compute expected for each balanced cooler
    expected_dfs: List[Optional[pd.DataFrame]] = [None] * n_coolers
    if args.expected:
        if len(args.expected) == 1 and n_coolers > 1:
            df = pd.read_csv(args.expected[0], sep="\t")
            if "balanced.avg" not in df.columns:
                logging.warning(f"Provided expected lacks 'balanced.avg': {args.expected[0]}")
            for i in range(n_coolers):
                if balanced_flags[i]:
                    expected_dfs[i] = df.copy()
        elif len(args.expected) == n_coolers:
            for i, ef in enumerate(args.expected):
                df = pd.read_csv(ef, sep="\t")
                if "balanced.avg" not in df.columns:
                    logging.warning(f"Provided expected lacks 'balanced.avg': {ef}")
                if balanced_flags[i]:
                    expected_dfs[i] = df
        else:
            die("When using --expected, provide either one file or one per cooler.")
    else:
        for i, uri in enumerate(clr_uris):
            if balanced_flags[i]:
                clr = cooler.Cooler(uri)
                expected_dfs[i] = compute_expected_df(
                    clr, view_df=view_df,
                    weight_name=args.weight_name,
                    ignore_diags=args.ignore_diags,
                    threads=args.threads
                )

    # Load region files
    features_list: List[pd.DataFrame] = []
    for rf, fmt in zip(args.regions, region_formats):
        features_list.append(load_features(rf, fmt))

    # Decide mode flags per region (BED only)
    def mode_flags_for(fmt: str) -> Tuple[bool, bool]:
        if fmt == "bed":
            if args.combine == "off-diagonal":
                return False, False
            if args.combine == "off-diagonal-rescaled":
                return False, True
            if args.combine == "diagonal":
                return True, False
            if args.combine == "diagonal-rescaled":
                return True, True
        # BEDPE: always off‑diagonal, unscaled
        return False, False

    local_flags = []
    rescale_flags = []
    for fmt in region_formats:
        l, r = mode_flags_for(fmt)
        local_flags.append(l)
        rescale_flags.append(r)

    # Dry‑run?
    if args.dry_run:
        logging.info("Dry‑run mode: validating and printing plan only.")
        for i, (uri, lab, bal) in enumerate(zip(clr_uris, cooler_labels, balanced_flags)):
            logging.info(f"  Cooler[{i}] {lab}: {uri}  | balanced={bal}")
        for j, (rf, lab, fmt, loc, rsc) in enumerate(zip(args.regions, region_labels, region_formats, local_flags, rescale_flags)):
            logging.info(f"  Regions[{j}] {lab}: {rf} ({fmt})  | local={loc} rescaled={rsc}")
        sys.exit(0)

    # Row/column orientation
    # We'll build (rows × cols) grids based on args.rows
    # First, compute all (cooler_i, region_j) pairs.
    combos = [(ci, ri) for ci in range(n_coolers) for ri in range(n_regions)]

    def process_pair(ci: int, ri: int) -> Tuple[Optional[np.ndarray], Optional[np.ndarray], int, int]:
        """Compute signal & O/E matrices for a single (cooler, region) pair."""
        clr_uri = clr_uris[ci]
        expdf = expected_dfs[ci]
        fmt = region_formats[ri]
        feats = features_list[ri]
        local = local_flags[ri] if fmt == "bed" else False
        rescale = rescale_flags[ri] if fmt == "bed" else False

        if balanced_flags[ci] and (expdf is not None):
            logging.info(f"[{ci},{ri}] Balanced + expected → signal + genome O/E")
            sig, oe = balanced_signal_and_genome_oe(
                clr_uri, feats, fmt,
                local=local, rescale=rescale,
                flank=args.flank, rescale_size=args.rescale_size, rescale_flank=args.rescale_flank,
                flip_negative_strand=args.flip_negative_strand, threads=args.threads,
                expected_df=expdf, view_df=view_df, seed=args.seed
            )
        else:
            logging.info(f"[{ci},{ri}] Fallback (raw + shift O/E)")
            sig, oe = fallback_raw_and_shift_oe(
                clr_uri, feats, fmt,
                local=local, rescale=rescale,
                flank=args.flank, rescale_size=args.rescale_size, rescale_flank=args.rescale_flank,
                flip_negative_strand=args.flip_negative_strand, threads=args.threads,
                nshifts=args.nshifts, view_df=view_df, seed=args.seed
            )
        return sig, oe, ci, ri

    if args.jobs > 1:
        results = Parallel(n_jobs=args.jobs, verbose=10)(
            delayed(process_pair)(ci, ri) for ci, ri in combos
        )
    else:
        results = [process_pair(ci, ri) for ci, ri in combos]

    # Collect raw (coolers × regions)
    sig = np.empty((n_coolers, n_regions), dtype=object)
    oe  = np.empty((n_coolers, n_regions), dtype=object)
    for s, o, ci, ri in results:
        sig[ci, ri] = s
        oe[ci,  ri] = o

    # Apply row/column reordering (labels and matrices together)
    row_labels = cooler_labels[:]
    col_labels = region_labels[:]
    sig_grid = sig.copy()
    oe_grid = oe.copy()

    if args.rows == "coolers":
        # current orientation already matches rows=coolers
        pass
    else:
        # rows=regions: transpose
        row_labels, col_labels = col_labels, row_labels
        sig_grid = sig_grid.T
        oe_grid = oe_grid.T

    n_rows, n_cols = sig_grid.shape

    if args.row_order:
        if len(args.row_order) != n_rows or sorted(args.row_order) != list(range(n_rows)):
            die("--row-order must list each row index exactly once.")
        row_labels = [row_labels[i] for i in args.row_order]
        sig_grid = sig_grid[args.row_order, :]
        oe_grid  = oe_grid[args.row_order, :]

    if args.col_order:
        if len(args.col_order) != n_cols or sorted(args.col_order) != list(range(n_cols)):
            die("--col-order must list each column index exactly once.")
        col_labels = [col_labels[j] for j in args.col_order]
        sig_grid = sig_grid[:, args.col_order]
        oe_grid  = oe_grid[:, args.col_order]

    # Determine rescale flag per cell for tick labeling
    is_rescaled_grid = np.empty_like(sig_grid, dtype=bool)
    for i in range(n_rows):
        for j in range(n_cols):
            # Column j corresponds to original region index depending on orientation:
            reg_index = j if args.rows == "coolers" else i
            fmt = region_formats[reg_index]
            if fmt == "bed":
                is_rescaled_grid[i, j] = rescale_flags[reg_index]
            else:
                is_rescaled_grid[i, j] = False  # BEDPE always unscaled

    # Build log2(O/E) with zero masking
    log2_grid = np.empty_like(oe_grid, dtype=object)
    for i in range(n_rows):
        for j in range(n_cols):
            o = oe_grid[i, j]
            if o is None:
                log2_grid[i, j] = None
            else:
                m = np.where(o > 0, o, np.nan)
                log2_grid[i, j] = np.log2(m)

    # Final colormaps
    use_fall = try_import_fall()
    signal_cmap = args.signal_cmap or ("fall" if use_fall else "inferno")
    diverging_cmap = plt.get_cmap(args.diverging_cmap)

    # Compute global normalization ranges (forced centers)
    sig_norm, oe_norm, l2_norm = compute_color_norms(sig_grid, oe_grid, log2_grid)

    # Plotting
    any_oe = any(x is not None for x in oe_grid.flatten())

    if n_cols == 1:
        # Single-column layout (N×3 or N×1)
        col_count = 3 if any_oe else 1
        fig_w, fig_h = 4 * col_count, 4 * n_rows
        fig, axes = plt.subplots(n_rows, col_count, figsize=(fig_w, fig_h))
        if n_rows == 1 and col_count == 1:
            axes = np.array([[axes]])
        elif n_rows == 1:
            axes = axes[np.newaxis, :]
        elif col_count == 1:
            axes = axes[:, np.newaxis]

        im_handles = [None] * col_count
        for i in range(n_rows):
            # Signal
            ax0 = axes[i, 0]
            s = sig_grid[i, 0]
            if s is not None:
                im0 = ax0.imshow(s, norm=sig_norm, cmap=plt.get_cmap(signal_cmap))
                im_handles[0] = im0
                size = s.shape[0]
                set_ticks(ax0, size, is_rescaled_grid[i, 0], args.rescale_flank, pixel_size_kb)
            ax0.set_ylabel(row_labels[i], fontsize=10)
            ax0.set_title("Signal (log scale)", fontsize=10)
            ax0.grid(False)

            if any_oe and col_count == 3:
                # O/E
                ax1 = axes[i, 1]
                o = oe_grid[i, 0]
                if o is not None:
                    im1 = ax1.imshow(o, cmap=diverging_cmap, norm=oe_norm)
                    im_handles[1] = im1
                    size = o.shape[0]
                    set_ticks(ax1, size, is_rescaled_grid[i, 0], args.rescale_flank, pixel_size_kb)
                ax1.set_title("O/E", fontsize=10)
                ax1.grid(False)

                # log2(O/E)
                ax2 = axes[i, 2]
                l2 = log2_grid[i, 0]
                if l2 is not None:
                    im2 = ax2.imshow(l2, cmap=diverging_cmap, norm=l2_norm)
                    im_handles[2] = im2
                    size = l2.shape[0]
                    set_ticks(ax2, size, is_rescaled_grid[i, 0], args.rescale_flank, pixel_size_kb)
                ax2.set_title("log₂(O/E)", fontsize=10)
                ax2.grid(False)

        # Colorbars
        fig.subplots_adjust(right=0.84)
        if col_count == 3:
            # Signal
            cax0 = fig.add_axes([0.86, 0.68, 0.02, 0.22])
            if im_handles[0]:
                cbar0 = fig.colorbar(im_handles[0], cax=cax0)
                cbar0.set_label("Signal (log scale)")
                cbar0.locator = ticker.LogLocator(base=10)
                cbar0.formatter = ticker.LogFormatterSciNotation(base=10)
                cbar0.update_ticks()
            # O/E
            cax1 = fig.add_axes([0.86, 0.38, 0.02, 0.22])
            if im_handles[1]:
                cbar1 = fig.colorbar(im_handles[1], cax=cax1)
                cbar1.set_label("O/E")
            # log2(O/E)
            cax2 = fig.add_axes([0.86, 0.08, 0.02, 0.22])
            if im_handles[2]:
                cbar2 = fig.colorbar(im_handles[2], cax=cax2)
                cbar2.set_label("log₂(O/E)")
            out_pdf = os.path.join(outdir, f"{ts}_singlecol.pdf")
        else:
            cax0 = fig.add_axes([0.86, 0.30, 0.02, 0.40])
            if im_handles[0]:
                cbar0 = fig.colorbar(im_handles[0], cax=cax0)
                cbar0.set_label("Signal (log scale)")
                cbar0.locator = ticker.LogLocator(base=10)
                cbar0.formatter = ticker.LogFormatterSciNotation(base=10)
                cbar0.update_ticks()
            out_pdf = os.path.join(outdir, f"{ts}_singlecol_signalonly.pdf")

        plt.savefig(out_pdf, bbox_inches="tight")
        plt.close(fig)
        logging.info(f"Wrote: {out_pdf}")

    else:
        # NxM separate PDFs
        plot_grid(sig_grid, row_labels, col_labels,
                  is_rescaled_grid=is_rescaled_grid, norm=sig_norm,
                  cmap=plt.get_cmap(signal_cmap), cbar_label="Signal (log scale)",
                  out_pdf=os.path.join(outdir, f"signal_{ts}.pdf"),
                  rescale_flank=args.rescale_flank, pixel_size_kb=pixel_size_kb)
        if any_oe:
            plot_grid(oe_grid, row_labels, col_labels,
                      is_rescaled_grid=is_rescaled_grid, norm=oe_norm,
                      cmap=diverging_cmap, cbar_label="O/E",
                      out_pdf=os.path.join(outdir, f"OE_{ts}.pdf"),
                      rescale_flank=args.rescale_flank, pixel_size_kb=pixel_size_kb)
            plot_grid(log2_grid, row_labels, col_labels,
                      is_rescaled_grid=is_rescaled_grid, norm=l2_norm,
                      cmap=diverging_cmap, cbar_label="log₂(O/E)",
                      out_pdf=os.path.join(outdir, f"log2OE_{ts}.pdf"),
                      rescale_flank=args.rescale_flank, pixel_size_kb=pixel_size_kb)
        else:
            logging.warning("No O/E matrices were generated; skipped O/E and log₂(O/E) grids.")

    # Save matrices to NPZ
    pile_data: Dict[str, Any] = {}
    for i in range(n_rows):
        for j in range(n_cols):
            rlab = row_labels[i].replace(" ", "_")
            clab = col_labels[j].replace(" ", "_")
            pile_data[f"signal_{rlab}_x_{clab}"] = sig_grid[i, j]
            pile_data[f"OE_{rlab}_x_{clab}"]     = oe_grid[i, j]

    npz_path = os.path.join(outdir, f"pileups_{ts}.npz")
    np.savez(npz_path, **pile_data)
    logging.info(f"Saved NPZ: {npz_path}")

    # Optional histograms
    if args.enable_hists:
        logging.info("Generating histograms...")
        # Signal (log10)
        sig_vals: Dict[str, np.ndarray] = {}
        for i in range(n_rows):
            for j in range(n_cols):
                m = sig_grid[i, j]
                if m is not None:
                    lab = f"{row_labels[i]}_x_{col_labels[j]}"
                    pos = m[m > 0]
                    if pos.size:
                        sig_vals[lab] = np.log10(pos)
        if sig_vals:
            overlay_histogram(sig_vals, os.path.join(outdir, f"hist_signal_{ts}.pdf"),
                              "Signal intensities", "log10(Signal)", bins=80)

        # O/E (linear)
        oe_vals: Dict[str, np.ndarray] = {}
        for i in range(n_rows):
            for j in range(n_cols):
                m = oe_grid[i, j]
                if m is not None:
                    lab = f"{row_labels[i]}_x_{col_labels[j]}"
                    pos = m[m > 0]
                    if pos.size:
                        oe_vals[lab] = pos
        if oe_vals:
            overlay_histogram(oe_vals, os.path.join(outdir, f"hist_OE_{ts}.pdf"),
                              "O/E", "O/E", bins=80)

        # log2(O/E)
        l2_vals: Dict[str, np.ndarray] = {}
        for i in range(n_rows):
            for j in range(n_cols):
                m = log2_grid[i, j]
                if m is not None:
                    lab = f"{row_labels[i]}_x_{col_labels[j]}"
                    l2_vals[lab] = m[np.isfinite(m)]
        if l2_vals:
            overlay_histogram(l2_vals, os.path.join(outdir, f"hist_log2OE_{ts}.pdf"),
                              "log₂(O/E)", "log₂(O/E)", bins=80)

    # Metadata JSON
    meta = {
        "timestamp": ts,
        "resolution_bp": binsize,
        "rows": args.rows,
        "coolers": [
            {"path": args.coolers[i], "label": cooler_labels[i], "balanced": bool(balanced_flags[i])}
            for i in range(n_coolers)
        ],
        "regions": [
            {"path": args.regions[j], "label": region_labels[j], "format": region_formats[j]}
            for j in range(n_regions)
        ],
        "parameters": {
            "combine_mode": args.combine,
            "flank_bp": args.flank,
            "rescale_size": args.rescale_size,
            "rescale_flank": args.rescale_flank,
            "binsize": args.binsize,
            "jobs": args.jobs,
            "threads": args.threads,
            "nshifts": args.nshifts,
            "seed": args.seed,
            "flip_negative_strand": args.flip_negative_strand,
            "enable_hists": args.enable_hists,
            "signal_cmap_final": signal_cmap,
            "diverging_cmap": args.diverging_cmap,
            "pixel_size_kb": pixel_size_kb,
            "view_used": bool(view_df is not None),
            "ignore_diags": args.ignore_diags,
            "weight_name": args.weight_name,
            "force_center_linear_OE": True,
            "force_center_log2_OE": True
        },
        "versions": {
            "coolpuppy": getattr(coolpuppy, "__version__", "unknown"),
            "cooltools": getattr(cooltools, "__version__", "unknown"),
            "cooler": getattr(cooler, "__version__", "unknown"),
            "matplotlib": mpl.__version__
        }
    }
    meta_path = os.path.join(outdir, f"metadata_{ts}.json")
    with open(meta_path, "w", encoding="utf-8") as f:
        json.dump(meta, f, indent=2)
    logging.info(f"Wrote metadata: {meta_path}")

    logging.info("All done. ✨")


if __name__ == "__main__":
    main()
```
