## Supplementary material for "CAD-C: An engineered nuclease enables repair-free *in situ* proximity ligation and nucleosome-resolution chromosome walks in human cells": Custom code: annotate_bedpe_by_ChromHMM_v1_20250421.py

```
# =============================================================================
#  Anchor‑ChromHMM Overlap & Base‑Pair Enrichment Calculator
#  Author: Jan Soroczynski, Risca Lab, The Rockefeller University
#  Date: 2025_04_20
#  Version: 0.1.0
# =============================================================================
#
#  INTRODUCTION
#  ------------
#  This fully‑production script ingests one BEDPE file of chromatin‑loop
#  anchors (e.g. Mustache loop calls) and a ChromHMM segmentation
#  (either one BED per state or a single combined BED with a state label
#  column).  It:
#
#    1. Converts both anchors of every BEDPE line into an “anchor” BED.
#    2. Optionally normalises all anchor intervals to the widest anchor
#       length present (centre‑expansion logic), if   --normalize‑width
#       is requested.  If widths vary and normalisation is *not*
#       requested, the script aborts with an informative error.
#    3. Computes per‑anchor, per‑state base‑pair overlaps using
#       bedtools intersect ‑wao via pybedtools.
#    4. Calculates genome‑wide base‑pair coverage of each ChromHMM
#       state.
#    5. Outputs:
#         • anchor_state_overlaps.tsv.gz     – every overlap record
#         • state_enrichment.tsv             – per‑state log₂ enrichment
#         • metadata.json                    – full run metadata
#         • run.log                          – timestamped execution log
#       all housed in   <outdir>/<timestamp>/
#
#  EXPECTED INPUTS
#  ---------------
#    --bedpe FILE            : bgzipped or plain BEDPE (columns ≥ 6).
#    --chromhmm‑files FILE(s) : one BED per state   *OR*
#    --chromhmm‑combined FILE : combined BED with state in 4th column.
#    --chrom‑sizes FILE       : optional two‑col chrom sizes; used to
#                               clip expanded anchors to chromosome bounds.
#
#  OPTIONAL ARGUMENTS
#  ------------------
#    --normalize‑width        : expand all anchors to the widest anchor.
#    --threads N              : multiprocessing workers (default = 1).
#    --outdir DIR             : parent directory for outputs.
#
#  OUTPUTS
#  -------
#    <timestamp>/anchor_state_overlaps.tsv.gz
#    <timestamp>/state_enrichment.tsv
#    <timestamp>/metadata.json
#    <timestamp>/run.log
#
#  EXAMPLE USAGE
#  -------------
#    anchor_chromHMM_enrich.py \
#      --bedpe loops_2.5kb.bedpe.gz \
#      --chromhmm‑files states/E??_*.bed \
#      --outdir results \
#      --threads 4 \
#      --normalize‑width
#
#  REQUIRED THIRD‑PARTY PYTHON PACKAGES
#  ------------------------------------
#    • pybedtools   (>=0.9.0)   –  wraps bedtools for Python
#    • pandas       (>=2.0)
#    • numpy        (>=1.25)
#
#  ENVIRONMENT
#  -----------
#    The script expects GNU bedtools ≥ 2.30 on $PATH and, if bgzip
#    BEDPE is supplied, pigz/tabix for decompression.  It runs under
#    Python ≥ 3.9 and automatically records package versions.
#
# =============================================================================
"""
Production‑grade anchor‑ChromHMM enrichment calculator.

See header for full documentation.
"""
from __future__ import annotations
import argparse
import hashlib
import json
import logging
import multiprocessing as mp
import os
import platform
import shutil
import sys
import tempfile
from datetime import datetime
from pathlib import Path
from typing import Dict, List, Tuple

import numpy as np
import pandas as pd
import pybedtools

# --------------------------------------------------------------------------- #
#  Helper functions
# --------------------------------------------------------------------------- #

def make_timestamp() -> str:
    return datetime.now().strftime("%Y%m%d_%H%M%S")


def sha256_of_file(fp: Path) -> str:
    h = hashlib.sha256()
    with open(fp, "rb") as f:
        for chunk in iter(lambda: f.read(8192), b""):
            h.update(chunk)
    return h.hexdigest()


def read_chrom_sizes(chrom_sizes_fp: Path | None) -> Dict[str, int] | None:
    if chrom_sizes_fp is None:
        return None
    sizes = {}
    with open(chrom_sizes_fp) as fh:
        for line in fh:
            chrom, size = line.rstrip().split()[:2]
            sizes[chrom] = int(size)
    return sizes


def anchor_bed_from_bedpe(bedpe_fp: Path, chrom_sizes: Dict[str, int] | None,
                          normalize_width: bool) -> Tuple[pybedtools.BedTool, int]:
    """Return BedTool of anchors and canonical anchor length."""
    cols = ["chrom1", "start1", "end1", "chrom2", "start2", "end2"]
    df = pd.read_table(bedpe_fp, usecols=range(6), names=cols, comment="#")
    df["width1"] = df["end1"] - df["start1"]
    df["width2"] = df["end2"] - df["start2"]

    unique_widths = pd.concat([df["width1"], df["width2"]]).unique()
    if len(unique_widths) > 1 and not normalize_width:
        raise ValueError(
            f"Variable anchor widths detected ({unique_widths}). "
            "Re‑run with --normalize‑width to equalise intervals."
        )

    anchor_len = int(max(unique_widths))  # canonical width
    records = []
    for idx, row in df.iterrows():
        for side in (1, 2):
            chrom = row[f"chrom{side}"]
            start = int(row[f"start{side}"])
            end = int(row[f"end{side}"])
            if normalize_width and (end - start) != anchor_len:
                mid = (start + end) // 2
                half = anchor_len // 2
                start = mid - half
                end = mid + half
                if chrom_sizes:
                    start = max(start, 0)
                    end = min(end, chrom_sizes.get(chrom, end))
            records.append((chrom, start, end, f"anchor_{idx}_{side}"))
    anchors_df = pd.DataFrame(records, columns=["chrom", "start", "end", "name"])
    return pybedtools.BedTool.from_dataframe(anchors_df), anchor_len


def load_chromhmm_bed(chromhmm_files: List[Path] | None,
                      combined_fp: Path | None) -> Tuple[pybedtools.BedTool, Dict[str, int]]:
    """Return BedTool of ChromHMM segments and dict of per‑state genome bp."""
    if chromhmm_files and combined_fp:
        raise ValueError("Provide either --chromhmm‑files or --chromhmm‑combined, not both.")
    if chromhmm_files:
        records = []
        state_bp = {}
        for fp in chromhmm_files:
            state = Path(fp).stem
            bed = pd.read_table(fp, header=None, usecols=[0, 1, 2], names=["chrom", "start", "end"])
            bed["state"] = state
            bp = (bed["end"] - bed["start"]).sum()
            state_bp[state] = int(bp)
            records.append(bed)
        segs_df = pd.concat(records, ignore_index=True)
    elif combined_fp:
        cols = ["chrom", "start", "end", "state"]
        segs_df = pd.read_table(combined_fp, header=None, usecols=range(4), names=cols)
        state_bp = segs_df.assign(length=lambda d: d["end"] - d["start"]).groupby("state")["length"].sum().astype(int).to_dict()
    else:
        raise ValueError("You must supply ChromHMM segmentation via --chromhmm‑files or --chromhmm‑combined.")
    return pybedtools.BedTool.from_dataframe(segs_df), state_bp


def intersect_chunk(chunk_bed: pybedtools.BedTool, chromhmm_bt: pybedtools.BedTool) -> pd.DataFrame:
    tmp = chunk_bed.intersect(chromhmm_bt, wao=True)
    cols = ["aChrom", "aStart", "aEnd", "aName",
            "bChrom", "bStart", "bEnd", "state", "overlap"]
    df = tmp.to_dataframe(names=cols, dtype={"overlap": int})
    df = df[df["overlap"] > 0]
    return df[["aName", "state", "overlap"]]


def compute_overlaps(anchors_bt: pybedtools.BedTool,
                     chromhmm_bt: pybedtools.BedTool,
                     threads: int) -> pd.DataFrame:
    """Intersect anchors vs. ChromHMM with optional multiprocessing."""
    # If BedTool has chunkit() (pybedtools ≥ 0.9.0) → fast path
    if hasattr(anchors_bt, "chunkit"):
        if threads <= 1:
            return intersect_chunk(anchors_bt, chromhmm_bt)
        with tempfile.TemporaryDirectory() as td:
            chunk_paths = anchors_bt.chunkit(threads, temp_dir=td)
            with mp.Pool(processes=threads) as pool:
                dfs = pool.starmap(
                    intersect_chunk,
                    [(pybedtools.BedTool(cp), chromhmm_bt) for cp in chunk_paths]
                )
        return pd.concat(dfs, ignore_index=True)

    # ------------------------------------------------------------------
    # Fallback for pybedtools < 0.9.0  (no .chunkit)
    # ------------------------------------------------------------------
    logging.warning(
        "pybedtools %s lacks chunkit(); falling back to single‑core intersect.",
        pybedtools.__version__,
    )
    # --- SIMPLE SERIAL PATH ---
    return intersect_chunk(anchors_bt, chromhmm_bt)

    # --- OPTIONAL DIY PARALLEL PATH ---
    # Uncomment this block if you truly need multiprocessing but cannot
    # upgrade pybedtools.  It writes a temp BED, splits it by line count,
    # and processes chunks in parallel.
    """
    if threads <= 1:
        return intersect_chunk(anchors_bt, chromhmm_bt)

    with tempfile.TemporaryDirectory() as td:
        anchor_path = anchors_bt.saveas(td + "/anchors.bed").fn
        # Count lines to size chunks
        with open(anchor_path) as fh:
            total = sum(1 for _ in fh)
        per_chunk = (total + threads - 1) // threads

        chunk_paths = []
        with open(anchor_path) as fh:
            for i in range(threads):
                out_fp = os.path.join(td, f"chunk_{i}.bed")
                with open(out_fp, "w") as out:
                    for _ in range(per_chunk):
                        line = fh.readline()
                        if not line:
                            break
                        out.write(line)
                if os.path.getsize(out_fp) > 0:
                    chunk_paths.append(out_fp)

        with mp.Pool(processes=threads) as pool:
            dfs = pool.starmap(
                intersect_chunk,
                [(pybedtools.BedTool(cp), chromhmm_bt) for cp in chunk_paths]
            )
    return pd.concat(dfs, ignore_index=True)
    """


def summarise_enrichment(overlaps_df: pd.DataFrame, state_bp: Dict[str, int],
                         total_anchor_bp: int) -> pd.DataFrame:
    obs = overlaps_df.groupby("state")["overlap"].sum()
    exp_frac = pd.Series(state_bp, name="state_bp") / sum(state_bp.values())
    observed_frac = obs / total_anchor_bp
    enrich = np.log2(observed_frac / exp_frac)
    out = pd.concat([obs.rename("observed_bp"),
                     pd.Series(state_bp, name="genome_bp"),
                     observed_frac.rename("observed_frac"),
                     exp_frac.rename("expected_frac"),
                     enrich.rename("log2_enrichment")], axis=1).reset_index().rename(columns={"index": "state"})
    out = out.sort_values("log2_enrichment", ascending=False)
    return out


# --------------------------------------------------------------------------- #
#  Main
# --------------------------------------------------------------------------- #

def main(argv: List[str] | None = None) -> None:
    parser = argparse.ArgumentParser(
        description="Anchor–ChromHMM base‑pair enrichment calculator",
        formatter_class=argparse.ArgumentDefaultsHelpFormatter,
    )
    parser.add_argument("--bedpe", required=True, type=Path, help="Loop anchors BEDPE")
    group = parser.add_mutually_exclusive_group(required=True)
    group.add_argument("--chromhmm-files", nargs="+", type=Path, help="One BED per ChromHMM state")
    group.add_argument("--chromhmm-combined", type=Path, help="Combined ChromHMM BED (state in 4th col)")
    parser.add_argument("--chrom-sizes", type=Path, help="Chromosome sizes TSV")
    parser.add_argument("--normalize-width", action="store_true",
                        help="Expand all anchors to widest interval detected")
    parser.add_argument("--threads", type=int, default=1)
    parser.add_argument("--outdir", type=Path, default=Path("anchor_chromhmm_enrich_out"))
    args = parser.parse_args(argv)

    ts = make_timestamp()
    out_root = args.outdir / ts
    out_root.mkdir(parents=True, exist_ok=True)

    # --------------------------------------------------------------------- #
    #  Logging
    # --------------------------------------------------------------------- #
    log_fp = out_root / f"run_{ts}.log"
    logging.basicConfig(
        level=logging.INFO,
        format="%(asctime)s [%(levelname)s] %(message)s",
        handlers=[
            logging.FileHandler(log_fp),
            logging.StreamHandler(sys.stdout),
        ],
    )
    logger = logging.getLogger(__name__)
    logger.info("♻️  Anchor‑ChromHMM enrichment run started.")

    chrom_sizes = read_chrom_sizes(args.chrom_sizes)

    logger.info("Reading BEDPE and constructing anchor BED …")
    anchors_bt, anchor_len = anchor_bed_from_bedpe(
        args.bedpe, chrom_sizes, args.normalize_width
    )
    total_anchor_bp = anchor_len * len(anchors_bt)
    logger.info(f"Anchors: {len(anchors_bt):,d}  •  Canonical width: {anchor_len} bp")

    logger.info("Loading ChromHMM segmentation …")
    chromhmm_bt, state_bp = load_chromhmm_bed(
        args.chromhmm_files, args.chromhmm_combined
    )
    logger.info(f"ChromHMM states detected: {len(state_bp)}")

    logger.info("Intersecting anchors with ChromHMM states …")
    overlaps_df = compute_overlaps(anchors_bt, chromhmm_bt, args.threads)
    overlaps_out = out_root / f"anchor_state_overlaps_{ts}.tsv.gz"
    overlaps_df.to_csv(overlaps_out, sep="\t", index=False, compression="gzip")
    logger.info(f"Anchor‑state overlap table written: {overlaps_out}")

    logger.info("Computing base‑pair enrichments …")
    enrich_df = summarise_enrichment(overlaps_df, state_bp, total_anchor_bp)
    enrich_out = out_root / f"state_enrichment_{ts}.tsv"
    enrich_df.to_csv(enrich_out, sep="\t", index=False)
    logger.info(f"Enrichment summary written: {enrich_out}")

    # ------------------------------------------------------------------ #
    #  Metadata
    # ------------------------------------------------------------------ #
    meta = {
        "timestamp": ts,
        "script": Path(__file__).name,
        "script_sha256": sha256_of_file(Path(__file__)),
        "bedpe": str(args.bedpe.resolve()),
        "chromhmm_files": [str(p) for p in (args.chromhmm_files or [])],
        "chromhmm_combined": str(args.chromhmm_combined) if args.chromhmm_combined else None,
        "chrom_sizes": str(args.chrom_sizes) if args.chrom_sizes else None,
        "normalize_width": args.normalize_width,
        "anchor_width_bp": anchor_len,
        "threads": args.threads,
        "packages": {
            "pybedtools": pybedtools.__version__,
            "pandas": pd.__version__,
            "numpy": np.__version__,
        },
        "platform": {
            "python": sys.version,
            "system": platform.platform(),
        },
    }
    meta_fp = out_root / f"metadata_{ts}.json"
    with open(meta_fp, "w") as fh:
        json.dump(meta, fh, indent=2)
    logger.info(f"Metadata written: {meta_fp}")

    logger.info("🎉  Run completed successfully.")


if __name__ == "__main__":
    try:
        main()
    except Exception as e:
        logging.exception("Fatal error: %s", e)
        sys.exit(1)
```
