## Supplementary material for "CAD-C: An engineered nuclease enables repair-free *in situ* proximity ligation and nucleosome-resolution chromosome walks in human cells": Custom code: loopmodel_final_pairs_script_WITH_MAKO.py

```
import numpy as np
import matplotlib.pyplot as plt
import seaborn as sns
from matplotlib.colors import LogNorm, BoundaryNorm, ListedColormap
import matplotlib.cm as cm
from matplotlib import rcParams
from matplotlib.ticker import NullLocator

# Settings for Illustrator-safe PDF
rcParams['pdf.fonttype'] = 42

# Input data and model definitions
s = np.logspace(0, 8, 2000)
rho_eff_values = np.array([
    4e-2, 2e-2, 1e-2,
    4e-3, 2e-3, 1e-3,
    4e-4, 2e-4, 1e-4,
    4e-5, 2e-5, 1e-5,
    4e-6, 2e-6, 1e-6
])
colors = plt.get_cmap("mako")(np.linspace(0, 1, len(rho_eff_values)))
colors_sorted = list(colors)
rho_sorted_indices = np.argsort(rho_eff_values)[::-1]
rho_eff_values_sorted = rho_eff_values[rho_sorted_indices]
colors_sorted = [colors[i] for i in rho_sorted_indices]

alpha = 1.0
gamma_s = s**(-alpha)
s_min, s_max = 1.0, 1e8
Gamma_s = (np.log(s) - np.log(s_min)) / (np.log(s_max) - np.log(s_min))
loop_center = 200_000
loop_sigma = 1_000
f_loop = 0.10

def loop_state_pdf(s, center, sigma):
    norm = 1 / (sigma * np.sqrt(2 * np.pi))
    return norm * np.exp(-0.5 * ((s - center) / sigma)**2)

loop_pdf = loop_state_pdf(s, loop_center, loop_sigma)
loop_pdf /= np.trapz(loop_pdf, s)

# Extract zoom region
s_zoom_mask = (s >= 1.90e5) & (s <= 2.10e5)
s_zoom = s[s_zoom_mask]
pobs_lines, oe_lines = [], []
pobs_min, pobs_max = np.inf, -np.inf

for rho_eff in rho_eff_values_sorted:
    gating_double = 1 - np.exp(-rho_eff * s) * (1 + rho_eff * s)
    dGamma_ds = 1 / ((np.log(s_max) - np.log(s_min)) * s)
    dgate_ds = (rho_eff**2) * s * np.exp(-rho_eff * s)
    P_base = Gamma_s * dgate_ds + gating_double * dGamma_ds
    gating_single_loop = 1 - np.exp(-rho_eff * 1e3)
    P_loop = loop_pdf * gating_single_loop
    P_obs = (1 - f_loop) * P_base + f_loop * P_loop
    OE_ratio = P_obs / P_base
    pobs_zoom = P_obs[s_zoom_mask]
    oe_zoom = OE_ratio[s_zoom_mask]
    pobs_lines.append(pobs_zoom)
    oe_lines.append(oe_zoom)
    pobs_min = min(pobs_min, np.min(pobs_zoom))
    pobs_max = max(pobs_max, np.max(pobs_zoom))

ymin = 10 ** (np.floor(np.log10(pobs_min)))
ymax = 10 ** (np.ceil(np.log10(pobs_max)))

# === Pair 1: Gating Functions ===
fig, (ax_g1, ax_g2) = plt.subplots(2, 1, figsize=(8, 6), sharex=True)
for i, (rho_eff, color) in reversed(list(enumerate(zip(rho_eff_values_sorted, colors_sorted)))):
    ax_g1.semilogx(s, 1 - np.exp(-rho_eff * s) * (1 + rho_eff * s), color=color, lw=1.5)
    ax_g2.semilogx(s, 1 - np.exp(-rho_eff * s), color=color, lw=1.5)

ax_g1.set_title('Double-anchor Gating Function')
ax_g1.set_ylabel('Probability')
ax_g2.set_title('Single-anchor Gating Function')
ax_g2.set_ylabel('Probability')
ax_g2.set_xlabel('Genomic separation (bp)')
for ax in [ax_g1, ax_g2]:
    ax.set_xlim(1e1, 1e8)
    ax.set_xticks([10**i for i in range(1, 9)])
    ax.get_xaxis().set_major_formatter(plt.FuncFormatter(lambda x, _: f'$10^{{{int(np.log10(x))}}}$'))
    ax.grid(True, which='both', ls='--', alpha=0.3)
fig.subplots_adjust(right=0.87)
cbar_ax = fig.add_axes([0.89, 0.2, 0.02, 0.6])
cmap = ListedColormap(colors_sorted[::-1])
norm = LogNorm(vmin=rho_eff_values_sorted.min(), vmax=rho_eff_values_sorted.max())
cbar = fig.colorbar(cm.ScalarMappable(norm=norm, cmap=cmap), cax=cbar_ax)
cbar.set_label(r'$\rho_{\mathrm{eff}}$', rotation=270, labelpad=15)
cbar.set_ticks(rho_eff_values_sorted)
cbar.ax.set_yticklabels([f"{v:.0e}" for v in rho_eff_values_sorted])
fig.savefig("pair1_gating_with_colorbar.pdf", bbox_inches='tight')
fig.savefig("pair1_gating_with_colorbar.png", dpi=300, bbox_inches='tight', transparent=True)

# === Pair 2: P_obs and O/E ===
fig, (ax_pobs, ax_oe) = plt.subplots(2, 1, figsize=(8, 10), sharex=True, gridspec_kw={"hspace": 0.35})
x_ticks_log = [10**i for i in range(1, 9)]
x_ticklabels_log = [f'$10^{i}$' for i in range(1, 9)]

ax_pobs.loglog(s, gamma_s, color='grey', alpha=0.5, lw=2)
for i, (rho_eff, color) in reversed(list(enumerate(zip(rho_eff_values_sorted, colors_sorted)))):
    gating_double = 1 - np.exp(-rho_eff * s) * (1 + rho_eff * s)
    dGamma_ds = 1 / ((np.log(s_max) - np.log(s_min)) * s)
    dgate_ds = (rho_eff**2) * s * np.exp(-rho_eff * s)
    P_base = Gamma_s * dgate_ds + gating_double * dGamma_ds
    gating_single_loop = 1 - np.exp(-rho_eff * 1e3)
    P_loop = loop_pdf * gating_single_loop
    P_obs = (1 - f_loop) * P_base + f_loop * P_loop
    ax_pobs.loglog(s, P_obs, color=color, lw=1.5)

ax_pobs.set_title('Observed Probability Distribution')
ax_pobs.set_ylabel(r'$P_{\mathrm{obs}}(s)$')
ax_pobs.set_xlim(1e1, 1e8)
ax_pobs.set_xticks(x_ticks_log)
ax_pobs.set_xticklabels(x_ticklabels_log)
ax_pobs.tick_params(axis='x', rotation=45, labelbottom=True)
ax_pobs.grid(True, which='both', ls='--', alpha=0.3)

for i, (rho_eff, color) in reversed(list(enumerate(zip(rho_eff_values_sorted, colors_sorted)))):
    gating_double = 1 - np.exp(-rho_eff * s) * (1 + rho_eff * s)
    dGamma_ds = 1 / ((np.log(s_max) - np.log(s_min)) * s)
    dgate_ds = (rho_eff**2) * s * np.exp(-rho_eff * s)
    P_base = Gamma_s * dgate_ds + gating_double * dGamma_ds
    gating_single_loop = 1 - np.exp(-rho_eff * 1e3)
    P_loop = loop_pdf * gating_single_loop
    P_obs = (1 - f_loop) * P_base + f_loop * P_loop
    OE_ratio = P_obs / P_base
    ax_oe.semilogx(s, OE_ratio, color=color, lw=1.5)

ax_oe.set_title('Observed/Expected (O/E) Ratio')
ax_oe.set_xlabel('Genomic separation (bp)')
ax_oe.set_ylabel('O/E ratio')
ax_oe.set_xlim(1e1, 1e8)
ax_oe.set_xticks(x_ticks_log)
ax_oe.set_xticklabels(x_ticklabels_log)
ax_oe.tick_params(axis='x', rotation=45)
ax_oe.grid(True, which='both', ls='--', alpha=0.3)

fig.subplots_adjust(right=0.87)
cbar_ax = fig.add_axes([0.89, 0.2, 0.02, 0.6])
cbar = fig.colorbar(cm.ScalarMappable(norm=norm, cmap=cmap), cax=cbar_ax)
cbar.set_label(r'$\rho_{\mathrm{eff}}$', rotation=270, labelpad=15)
cbar.set_ticks(rho_eff_values_sorted)
cbar.ax.set_yticklabels([f"{v:.0e}" for v in rho_eff_values_sorted])
fig.savefig("pair2_ps_and_oe_xticks_on_top_plot_fixed.pdf", bbox_inches='tight')
fig.savefig("pair2_ps_and_oe_xticks_on_top_plot_fixed.png", dpi=300, bbox_inches='tight', transparent=True)

# === Pair 3: Zoomed Loop Region ===
fig, (ax_zoom_pobs, ax_zoom_oe) = plt.subplots(2, 1, figsize=(8, 6), sharex=True, gridspec_kw={"height_ratios": [1, 1]})
xticks_kb = [190000, 195000, 200000, 205000, 210000]
xticklabels_kb = ['190 kb', '195 kb', '200 kb', '205 kb', '210 kb']

for i, color in reversed(list(enumerate(colors_sorted))):
    ax_zoom_pobs.semilogx(s_zoom, pobs_lines[i], color=color, lw=1.5)
ax_zoom_pobs.set_yscale('log')
ax_zoom_pobs.set_xlim(1.90e5, 2.10e5)
ax_zoom_pobs.set_ylim(ymin, ymax)
ax_zoom_pobs.set_title(r'Zoomed $P_{\mathrm{obs}}(s)$ (190–210 kb)')
ax_zoom_pobs.set_ylabel(r'$P_{\mathrm{obs}}(s)$')
ax_zoom_pobs.set_xticks(xticks_kb)
ax_zoom_pobs.set_xticklabels(xticklabels_kb)
ax_zoom_pobs.tick_params(axis='x', rotation=45)
ax_zoom_pobs.grid(True, which='both', ls='--', alpha=0.3)
ax_zoom_pobs.xaxis.set_minor_locator(NullLocator())

for i, color in reversed(list(enumerate(colors_sorted))):
    ax_zoom_oe.semilogx(s_zoom, oe_lines[i], color=color, lw=1.5)
ax_zoom_oe.set_xlim(1.90e5, 2.10e5)
ax_zoom_oe.set_title('Zoomed O/E Ratio (190–210 kb)')
ax_zoom_oe.set_xlabel('Genomic separation (bp)')
ax_zoom_oe.set_ylabel('O/E ratio')
ax_zoom_oe.set_xticks(xticks_kb)
ax_zoom_oe.set_xticklabels(xticklabels_kb)
ax_zoom_oe.tick_params(axis='x', rotation=45)
ax_zoom_oe.grid(True, which='both', ls='--', alpha=0.3)
ax_zoom_oe.xaxis.set_minor_locator(NullLocator())

fig.subplots_adjust(right=0.87)
cbar_ax = fig.add_axes([0.89, 0.2, 0.02, 0.6])
cbar = fig.colorbar(cm.ScalarMappable(norm=norm, cmap=cmap), cax=cbar_ax)
cbar.set_label(r'$\rho_{\mathrm{eff}}$', rotation=270, labelpad=15)
cbar.set_ticks(rho_eff_values_sorted)
cbar.ax.set_yticklabels([f"{v:.0e}" for v in rho_eff_values_sorted])
fig.savefig("pair3_zoomed_final_cleaned.pdf", bbox_inches='tight')
fig.savefig("pair3_zoomed_final_cleaned.png", dpi=300, bbox_inches='tight', transparent=True)
```
