## Supplementary material for "CAD-C: An engineered nuclease enables repair-free *in situ* proximity ligation and nucleosome-resolution chromosome walks in human cells": Custom code: submit_three_loop_sets_intersect_wbed_upset_v4_test1_20250923.sh

```
#!/bin/bash
#==============================================================================
#  Loop–Feature Intersect & Cross‑Set UpSet + Stats (A, A∩B, B)
#  SBATCH wrapper
#------------------------------------------------------------------------------
#  This submits one SLURM job that launches:
#    loop_feature_intersect_and_upset_v1.py
#
#  You provide:
#    • FEATURE_BED  : generic feature BED (e.g., ENCODE RAMPAGE TSS, narrowPeak, etc.)
#    • BEDPE_A      : Method A loops (e.g., Micro‑C only)
#    • BEDPE_COMMON : Common A∩B loops (detected by both; precomputed outside)
#    • BEDPE_B      : Method B loops (e.g., CAD‑C only)
#  Optionally:
#    • MASTER_BED   : the full feature universe (same format as FEATURE_BED) to add “Neither”
#    • WINDOW_HALF_WIDTH : to use ±W bp windows around anchor midpoints (else original widths)
#    • CHROM_SIZES  : chrom sizes TSV for safe window clipping and sorting
#    • LABEL_*      : friendly labels for plots/filenames (defaults = BEDPE stems)
#
#  Outputs land under:  ${OUTDIR}/<timestamp>/
#
#  Requirements on the compute node:
#    - python ≥ 3.9
#    - bedtools ≥ 2.30 on $PATH
#    - Python pkgs: pybedtools, pandas, numpy, matplotlib (SciPy optional)
#
#==============================================================================

#SBATCH --job-name=loopFeatUpSet
#SBATCH --partition=risc
#SBATCH --ntasks=1
#SBATCH --cpus-per-task=8
#SBATCH --output=loopFeatUpSet_%j.out
#SBATCH --error=loopFeatUpSet_%j.err
# (Add time/memory directives if your cluster requires them)

set -euo pipefail

# ----------------------------- USER EDITABLES ------------------------------ #

# Path to the Python script you saved from our conversation:
SCRIPT="/rugpfs/fs0/risc_lab/scratch/jsoroczyns/CAD/CAD-C/scripts/2025_09_23/three_loop_sets_intersect_wbed_upset_v4_20250923.py"

# Feature BED (e.g., RAMPAGE TSS, narrowPeak, etc.)
FEATURE_BED="/rugpfs/fs0/risc_lab/scratch/jsoroczyns/CAD/CAD-C/Ligation_Junction_Coverage_2025/GM12878_reference_beds/ENCFF591KXZ.bed"         # <—— EDIT

# Optional: alias used in output names/plots (defaults to basename of FEATURE_BED)
FEATURE_ALIAS="RAMPAGE_TSS"                     # e.g., "RAMPAGE_TSS" or "narrowPeak"; can leave empty

# Three BEDPE inputs (A, A∩B, B)
BEDPE_A="/rugpfs/fs0/risc_lab/scratch/wzuo/project/CAD/GM12878_dataset/loops/loop_2500/GM12878_MicroC_only_2500_loop_5kb_slop.bedpe"          # <—— EDIT
BEDPE_COMMON="/rugpfs/fs0/risc_lab/scratch/wzuo/project/CAD/GM12878_dataset/loops/loop_2500/GM12878_CADC_MicroC_2500_loop_5kb_slop.bedpe"     # <—— EDIT
BEDPE_B="/rugpfs/fs0/risc_lab/scratch/wzuo/project/CAD/GM12878_dataset/loops/loop_2500/GM12878_CADC_only_2500_loop_5kb_slop.bedpe"            # <—— EDIT

# Optional friendly labels for plots/paths (defaults = BEDPE stem if left blank)
LABEL_A="Micro-C only"
LABEL_COMMON="Common (both)"
LABEL_B="CAD-C only"

# Optional: master feature BED to enable “Neither” category in UpSet + absolute stats
MASTER_BED="/rugpfs/fs0/risc_lab/scratch/jsoroczyns/CAD/CAD-C/Ligation_Junction_Coverage_2025/GM12878_reference_beds/ENCFF591KXZ.bed"           # leave empty "" to skip

# Optional: windowed‑anchor mode (±W bp around anchor midpoints). Leave empty to use original widths.
WINDOW_HALF_WIDTH=""                             # e.g., 1500  (empty == use original BEDPE anchor widths)

# Optional: chrom sizes for window clipping & chromosome ordering
CHROM_SIZES="/rugpfs/fs0/risc_lab/scratch/jsoroczyns/CAD/CAD-C/CADC_E2_P1_Novogene/20240320_pt_flip_out/hg38.chrom.sizes"         # leave empty "" if not using windowing

# Output parent directory
OUTDIR="/rugpfs/fs0/risc_lab/scratch/jsoroczyns/CAD/CAD-C/TSS_anchor_intersects_2025"  # <—— EDIT or keep

# Threads (defaults to SLURM_CPUS_PER_TASK)
THREADS="${SLURM_CPUS_PER_TASK:-8}"

# Optional conda environment activation (uncomment & edit if you use conda)
# CONDA_ENV="open2c_v3"
# source "$(conda info --base)/etc/profile.d/conda.sh"
# conda activate "${CONDA_ENV}"

# Optional module loads (uncomment if needed on your cluster)
# module load bedtools/2.30.0

# ----------------------------- ARG ASSEMBLY -------------------------------- #

ARGS=( --feature-bed "${FEATURE_BED}"
       --bedpe-a "${BEDPE_A}"
       --bedpe-common "${BEDPE_COMMON}"
       --bedpe-b "${BEDPE_B}"
       --threads "${THREADS}"
       --outdir "${OUTDIR}"
)

# Optional flags
if [[ -n "${FEATURE_ALIAS}" ]]; then
  ARGS+=( --feature-alias "${FEATURE_ALIAS}" )
fi
if [[ -n "${LABEL_A}" ]]; then
  ARGS+=( --label-a "${LABEL_A}" )
fi
if [[ -n "${LABEL_COMMON}" ]]; then
  ARGS+=( --label-common "${LABEL_COMMON}" )
fi
if [[ -n "${LABEL_B}" ]]; then
  ARGS+=( --label-b "${LABEL_B}" )
fi
if [[ -n "${MASTER_BED}" ]]; then
  ARGS+=( --master-bed "${MASTER_BED}" )
fi
if [[ -n "${WINDOW_HALF_WIDTH}" ]]; then
  ARGS+=( --window-half-width "${WINDOW_HALF_WIDTH}" )
  if [[ -n "${CHROM_SIZES}" ]]; then
    ARGS+=( --chrom-sizes "${CHROM_SIZES}" )
  else
    echo "[WARN] WINDOW_HALF_WIDTH set, but CHROM_SIZES is empty. Windows will not be clipped to chrom bounds."
  fi
fi

# ------------------------------- EXECUTION --------------------------------- #

echo "[$(date +%F' '%T)] Starting loop_feature_intersect_and_upset_v1.py"
echo "python ${SCRIPT} \\"
for a in "${ARGS[@]}"; do printf "  %q \\\n" "$a"; done
echo

python "${SCRIPT}" "${ARGS[@]}"

echo "[$(date +%F' '%T)] Job completed."
```
