## Supplementary material for "CAD-C: An engineered nuclease enables repair-free *in situ* proximity ligation and nucleosome-resolution chromosome walks in human cells": Custom code: three_loop_sets_intersect_wbed_upset_v4_20250923.py

```
#!/usr/bin/env python3
# -*- coding: utf-8 -*-
"""
Loop–Feature Intersect & Cross‑Set UpSet + Stats (BED ↔ BEDPE) — Combined Pipeline
==================================================================================

Author: Jan Soroczynski, Risca Lab, The Rockefeller University
Date:   2025-09-23
Version: 4.0.0

OVERVIEW
--------
This script performs two stages:

STAGE 1 — Per‑set feature↔anchor intersects (3 BEDPE sets):
    Inputs:
      • One generic BED of genomic features (“feature BED”; e.g., ENCODE RAMPAGE TSS,
        ENCODE narrowPeak, etc.). Not hard‑coded to TSS; supports .bed or .bed.gz.
      • Three BEDPE loop sets prepared upstream:
          (A)  Method A specific         (e.g., Micro‑C only)
          (AB) Common A∩B (same loops)   (e.g., both)
          (B)  Method B specific         (e.g., CAD‑C only)
        NOTE: This script **does not** compute loop intersections; it assumes A, AB, B were
              already produced elsewhere.

    What STAGE 1 does for **each** BEDPE:
      1) Build anchor intervals from BEDPE:
         • Option A (default): use original anchor intervals from BEDPE.
         • Option B: windowed anchors: center on anchor midpoint and expand ±W bp
           (clip to chrom sizes if provided).
      2) Intersect anchors vs features (feature BED) using bedtools via pybedtools.
         Intersection uses `-wao` and keeps hits with **overlap > 0**.
         The feature BED is represented as a safe 4‑column BED [chrom, start, end, feat_id],
         so bedtools does not choke on quoted/space‑rich fields.
      3) Output per‑set artifacts (exact input lines preserved; no extra columns in BED/BEDPE):
         • **bed/**    features_intersect__<alias>__<setLabel>__<ts>.bed[.gz]
                          – de‑duplicated exact feature lines that hit ≥1 anchor in this set.
         • **bedpe/**  (FOUR loop subsets; all are exact original loop lines)
                          – loops_nohit__<setLabel>__<ts>.bedpe        (0 anchors hit)
                          – loops_onehit__<setLabel>__<ts>.bedpe       (exactly 1 anchor hit)
                          – loops_bothhit__<setLabel>__<ts>.bedpe      (both anchors hit)
                          – loops_anyhit__<setLabel>__<ts>.bedpe       (≥1 anchor hit; one ∪ both)
         • **tables/** pairs__<alias>__<setLabel>__<ts>.tsv.gz
                       classification__<setLabel>__<ts>.tsv
                       anchor_summary__<alias>__<setLabel>__<ts>.tsv
                          – provenance, loop 0/1/2 counts, anchor‑centric hit metrics.
         • **metadata/** JSON with parameters, counts, and file paths.
         • **logs/**   per‑set run log (clear category counts and semantics).

      4) Statistics **per set** (anchor‑centric):
         • #loops with 0/1/2 anchors hit (multiple features per anchor count as **1** hit).
         • #anchors, #anchors with ≥1 feature, **anchor hit rate**.
         • **Unique features per 10k anchors** and **per Mb of anchor DNA** (median anchor width).

STAGE 2 — Cross‑set UpSet + Stats across the three sets:
    Inputs:
      • The three per‑set **feature sets** from Stage 1 (A, AB, B) represented by exact
        feature lines (identity = full line MD5).
      • OPTIONAL: a **master feature BED** (e.g., ENCODE RAMPAGE TSS) to define the universe
        so we can compute the **“Neither”** category (master minus union).

    What STAGE 2 outputs:
      • **cross_set/**:
          - summary.tsv – full 7 Venn cells; totals; plus master totals if provided.
          - combo_counts_union.tsv
          - combo_counts_with_neither.tsv (if master provided)
          - **features_*.bed[.gz]** for **all 8 categories**:
              A_only, AB_only, B_only, A∩AB_only, A∩B_only, AB∩B_only, A∩AB∩B, Neither
              (each BED contains the exact feature lines for that category).
          - UpSet figures (**black bars, filled/hollow dots, PDF fonttype=42**):
              upset_union.[png|pdf]
              upset_with_neither.[png|pdf] (if master provided)
          - McNemar / exact binomial stats tables:
              stats_union_mcnemar.tsv      – among union (detected by ≥1)
              stats_absolute_mcnemar.tsv   – vs master universe (includes “Neither”)

DEFAULTS & COMPRESSION
----------------------
• **Default**: feature BED outputs are **plain .bed** (no compression).
  Pass **--compress-bed** to write them as `.bed.gz`.
• Large provenance tables (pairs__*.tsv.gz) remain gzipped by default.

REQUIREMENTS
------------
Python ≥ 3.9
bedtools ≥ 2.30 on $PATH
Python packages: pybedtools (≥0.9.0 recommended), pandas (≥2.0), numpy (≥1.25), matplotlib (≥3).
SciPy optional for exact binomial; a reliable normal‑approx fallback is provided if absent.

NOTES
-----
• Exact input lines for feature BED and loop BEDPE are preserved in outputs.
• Feature identity = MD5(full canonical line) 16‑char prefix (safe for intersects).
• Intersections require **overlap > 0** bp (no minimum size).
• For loop/anchor propensity, multiple features within an anchor count as **1 anchor hit**.
• STAGE 2 “union‑only” McNemar test uses: A_detected = A ∪ AB, B_detected = B ∪ AB.
• PDF outputs use **fonttype=42** to embed TrueType fonts.

CHANGES IN 4.0.0
----------------
• **BREAKING**: removed the legacy alias `loops_withhit__*.bedpe`.  
  The script now writes **only**: `loops_nohit`, `loops_onehit`, `loops_bothhit`, `loops_anyhit`.
• Logs/metadata/outputs contain only those four categories (no alias anywhere).
"""

from __future__ import annotations

import argparse
import gzip
import hashlib
import json
import logging
import math
import multiprocessing as mp
import sys
import tempfile
from dataclasses import dataclass
from datetime import datetime
from pathlib import Path
from typing import Dict, Iterable, List, Optional, Tuple

import numpy as np
import pandas as pd
import pybedtools

# =============================== Utilities ================================ #

def ts_now() -> str:
    return datetime.now().strftime("%Y%m%d_%H%M%S")

def ensure_dir(p: Path) -> None:
    p.mkdir(parents=True, exist_ok=True)

def sha256_of_file(fp: Path) -> str:
    h = hashlib.sha256()
    with open(fp, "rb") as f:
        for chunk in iter(lambda: f.read(1 << 16), b""):
            h.update(chunk)
    return h.hexdigest()

def read_chrom_sizes(fp: Optional[Path]) -> Optional[Dict[str, int]]:
    if fp is None:
        return None
    sizes: Dict[str, int] = {}
    with open(fp) as fh:
        for line in fh:
            if not line.strip() or line.startswith("#"):
                continue
            chrom, size = line.rstrip().split()[:2]
            sizes[chrom] = int(size)
    return sizes

def open_maybe_gzip_read(path: Path):
    suf = "".join(path.suffixes).lower()
    if suf.endswith(".gz") or suf.endswith(".bgz") or suf.endswith(".bgzip"):
        return gzip.open(path, "rt")
    return open(path, "r")

def canonicalize_bed_line(line: str) -> str:
    parts = line.rstrip("\n\r").rstrip().split("\t")
    return "\t".join(parts)

def parse_bed_line(line: str) -> Optional[Tuple[str,int,int,str]]:
    if not line or line.startswith("#"):
        return None
    parts = line.split("\t")
    if len(parts) < 3:
        return None
    try:
        chrom = parts[0]
        start = int(parts[1])
        end   = int(parts[2])
    except ValueError:
        return None
    rest = "\t".join(parts[3:]) if len(parts) > 3 else ""
    return chrom, start, end, rest

def natural_chrom_key(chrom: str) -> Tuple[int, str]:
    c = chrom[3:] if chrom.startswith("chr") else chrom
    mapping = {"X": 23, "Y": 24, "M": 25, "MT": 25}
    if c in mapping:
        return (mapping[c], chrom)
    try:
        return (int(c), chrom)
    except ValueError:
        return (1000, chrom)

# =============================== Feature BED =============================== #

@dataclass
class FeatureData:
    path: Path
    alias: str
    lines: List[str]                      # exact lines (non‑comment)
    rows: List[Tuple[str,int,int,str]]    # (chrom,start,end,rest)
    ids:  List[str]                       # 16‑char md5 per exact line

def read_feature_bed(path: Path, alias: Optional[str]) -> FeatureData:
    lines: List[str] = []
    rows:  List[Tuple[str,int,int,str]] = []
    with open_maybe_gzip_read(path) as fh:
        for raw in fh:
            raw = raw.rstrip("\n\r")
            if not raw or raw.startswith("#"):
                continue
            canon = canonicalize_bed_line(raw)
            parsed = parse_bed_line(canon)
            if parsed is None:
                continue
            lines.append(canon)
            rows.append(parsed)
    ids = [hashlib.md5(l.encode("utf-8")).hexdigest()[:16] for l in lines]
    alias2 = alias if (alias and alias.strip()) else path.stem
    return FeatureData(path=path, alias=alias2, lines=lines, rows=rows, ids=ids)

def feature_bedtool(fd: FeatureData) -> Tuple[pybedtools.BedTool, Dict[str,str], Dict[str,Tuple[str,int,int]]]:
    df = pd.DataFrame({
        "chrom": [r[0] for r in fd.rows],
        "start": [r[1] for r in fd.rows],
        "end":   [r[2] for r in fd.rows],
        "feat_id": fd.ids,
    })
    bt = pybedtools.BedTool.from_dataframe(df)
    id_to_line  = dict(zip(fd.ids, fd.lines))
    id_to_coord = dict(zip(fd.ids, [(r[0],r[1],r[2]) for r in fd.rows]))
    return bt, id_to_line, id_to_coord

# =============================== BEDPE & Anchors =========================== #

@dataclass
class Anchors:
    bedtool: pybedtools.BedTool
    names: List[str]        # "anchor_<loopidx>_<1|2>"
    loop_count: int
    bedpe_lines: List[str]  # exact original lines for all loops (non‑comment)

def read_bedpe_with_raw(bedpe_fp: Path) -> Tuple[pd.DataFrame, List[str]]:
    raw_lines: List[str] = []
    with open(bedpe_fp) as fh:
        for line in fh:
            if not line.strip() or line.startswith("#"):
                continue
            raw_lines.append(line.rstrip("\n\r"))
    if not raw_lines:
        raise ValueError(f"{bedpe_fp} appears empty.")
    df = pd.read_table(
        bedpe_fp, comment="#", header=None, usecols=[0,1,2,3,4,5],
        dtype={0:str,1:np.int64,2:np.int64,3:str,4:np.int64,5:np.int64}, engine="c"
    )
    return df, raw_lines

def bedpe_to_anchors_original(bedpe_fp: Path) -> Anchors:
    df, raw = read_bedpe_with_raw(bedpe_fp)
    recs, names = [], []
    for i, r in df.iterrows():
        for side, (chrom, start, end) in enumerate([(r[0], int(r[1]), int(r[2])),
                                                    (r[3], int(r[4]), int(r[5]))], start=1):
            name = f"anchor_{i}_{side}"
            names.append(name)
            recs.append((chrom, start, end, name))
    bt = pybedtools.BedTool.from_dataframe(pd.DataFrame(recs, columns=["chrom","start","end","name"]))
    return Anchors(bedtool=bt, names=names, loop_count=df.shape[0], bedpe_lines=raw)

def bedpe_to_anchors_windowed(bedpe_fp: Path, half_width: int, chrom_sizes: Optional[Dict[str,int]]) -> Anchors:
    df, raw = read_bedpe_with_raw(bedpe_fp)
    recs, names = [], []
    for i, r in df.iterrows():
        for side, (chrom, start, end) in enumerate([(r[0], int(r[1]), int(r[2])),
                                                    (r[3], int(r[4]), int(r[5]))], start=1):
            mid = (start + end) // 2
            s = mid - half_width
            e = mid + half_width
            if chrom_sizes:
                s = max(0, s)
                e = min(chrom_sizes.get(chrom, e), e)
            name = f"anchor_{i}_{side}"
            names.append(name)
            recs.append((chrom, s, e, name))
    bt = pybedtools.BedTool.from_dataframe(pd.DataFrame(recs, columns=["chrom","start","end","name"]))
    return Anchors(bedtool=bt, names=names, loop_count=df.shape[0], bedpe_lines=raw)

# =============================== Intersections ============================= #

def intersect_anchors_features(
    anchors_bt: pybedtools.BedTool,
    feat_bt: pybedtools.BedTool,
    threads: int,
    logger: logging.Logger,
) -> pd.DataFrame:
    """
    Return dataframe columns:
      aChrom aStart aEnd aName  bChrom bStart bEnd feat_id overlap
    (read as strings; keep overlap>0; cast numeric after filtering)
    """
    def run(bt: pybedtools.BedTool) -> pd.DataFrame:
        tmp = bt.intersect(feat_bt, wao=True)
        cols = ["aChrom","aStart","aEnd","aName","bChrom","bStart","bEnd","feat_id","overlap"]
        df = tmp.to_dataframe(names=cols, dtype=str)
        df["overlap"] = pd.to_numeric(df["overlap"], errors="coerce").fillna(0).astype(int)
        df = df[df["overlap"] > 0].copy()
        if not df.empty:
            df["bStart"] = pd.to_numeric(df["bStart"], errors="coerce").astype(int)
            df["bEnd"]   = pd.to_numeric(df["bEnd"],   errors="coerce").astype(int)
        return df

    if hasattr(anchors_bt, "chunkit") and threads > 1:
        with tempfile.TemporaryDirectory() as td:
            chunks = anchors_bt.chunkit(threads, temp_dir=td)
            with mp.Pool(processes=threads) as pool:
                dfs = pool.map(lambda p: run(pybedtools.BedTool(p)), chunks)
        return pd.concat(dfs, ignore_index=True) if dfs else pd.DataFrame(
            columns=["aChrom","aStart","aEnd","aName","bChrom","bStart","bEnd","feat_id","overlap"]
        )
    else:
        if not hasattr(anchors_bt, "chunkit"):
            logger.warning("pybedtools %s lacks chunkit(); falling back to single‑core.", pybedtools.__version__)
        return run(anchors_bt)

# =============================== Per‑set processing ======================== #

@dataclass
class SetOutputs:
    label: str
    base_dir: Path
    features_bed: Path
    loops_nohit_bedpe: Path
    loops_one_bedpe: Path
    loops_both_bedpe: Path
    loops_any_bedpe: Path
    pairs_tsv_gz: Path
    class_tsv: Path
    anchor_summary_tsv: Path
    feat_ids: set
    log_path: Path

def write_feature_bed_lines(lines: List[str], out_base: Path, compress_bed: bool) -> Path:
    """Write feature lines to out_base with .bed or .bed.gz suffix depending on compress_bed."""
    fp = out_base.with_suffix(out_base.suffix + ".gz") if compress_bed else out_base
    ensure_dir(fp.parent)
    if compress_bed:
        with gzip.open(fp, "wt") as out:
            for line in lines:
                out.write(line + "\n")
    else:
        with open(fp, "w") as out:
            for line in lines:
                out.write(line + "\n")
    return fp

def process_one_set(
    label: str,
    bedpe_fp: Path,
    feat_data: FeatureData,
    feat_bt: pybedtools.BedTool,
    id_to_line: Dict[str,str],
    id_to_coord: Dict[str,Tuple[str,int,int]],
    chrom_sizes: Optional[Dict[str,int]],
    window_half_width: Optional[int],
    threads: int,
    out_root: Path,
    ts: str,
    compress_bed: bool,
    logger_parent: logging.Logger,
) -> SetOutputs:
    set_dir = out_root / "per_set" / label
    ensure_dir(set_dir)
    log_dir = set_dir / "logs"; ensure_dir(log_dir)
    log_path = log_dir / f"run__{label}__{ts}.log"
    logger = logging.getLogger(f"loop_feat.{label}")
    fh = logging.FileHandler(log_path); fh.setFormatter(logging.Formatter("%(asctime)s [%(levelname)s] %(message)s"))
    logger.addHandler(fh); logger.setLevel(logger_parent.level); logger.propagate = True

    logger.info("▶ Processing set: %s", label)
    logger.info("Feature alias: %s | Feature BED: %s", feat_data.alias, feat_data.path)
    logger.info("BEDPE: %s", bedpe_fp)

    # Anchors
    anchors = (bedpe_to_anchors_windowed if window_half_width is not None else bedpe_to_anchors_original)(
        bedpe_fp, window_half_width, chrom_sizes) if window_half_width is not None else bedpe_to_anchors_original(bedpe_fp)
    if window_half_width is not None:
        logger.info("Anchor mode: WINDOW ±%d bp around midpoints.", window_half_width)
    else:
        logger.info("Anchor mode: ORIGINAL widths from BEDPE.")
    anchors_total = anchors.loop_count * 2
    logger.info("Loops=%d  Anchors=%d", anchors.loop_count, anchors_total)

    # Intersect
    inter = intersect_anchors_features(anchors.bedtool, feat_bt, threads, logger)
    inter["anchor_id"] = inter["aName"].astype(str)
    inter["which_anchor"] = inter["aName"].str.rsplit("_", n=1, expand=True)[1].astype(int)
    inter["feat_line"] = inter["feat_id"].map(id_to_line)
    parts = inter["feat_line"].str.split("\t", n=3, expand=True)
    inter["bChrom"] = parts[0]; inter["bStart"] = parts[1].astype(int); inter["bEnd"] = parts[2].astype(int)
    inter["bRest"]  = parts[3].fillna("")

    # Pairs table
    tables_dir = set_dir / "tables"; ensure_dir(tables_dir)
    pairs_tsv_gz = tables_dir / f"pairs__{feat_data.alias}__{label}__{ts}.tsv.gz"
    inter[["anchor_id","which_anchor","bChrom","bStart","bEnd","bRest","feat_id","overlap","feat_line"]].to_csv(
        pairs_tsv_gz, sep="\t", index=False, compression="gzip"
    )
    logger.info("Pairs table written: %s (rows=%d)", pairs_tsv_gz, len(inter))

    # Feature BED (exact lines; de‑duplicate; coordinate‑sorted)
    feat_ids = list(dict.fromkeys(inter["feat_id"])) if not inter.empty else []
    def sortkey(tid: str):
        if tid in id_to_coord:
            chrom,s,e = id_to_coord[tid]
            if chrom_sizes:
                order = {c:i for i,c in enumerate(chrom_sizes.keys())}
                return (order.get(chrom, 10**9), s, e)
            else:
                return (natural_chrom_key(chrom), s, e)
        return ((1000, "chrZZZ"), 10**12, 10**12)
    feat_ids_sorted = sorted(feat_ids, key=sortkey)
    feat_lines_sorted = [id_to_line[i] for i in feat_ids_sorted]
    bed_dir = set_dir / "bed"; ensure_dir(bed_dir)
    features_bed = write_feature_bed_lines(
        feat_lines_sorted,
        out_base=(bed_dir / f"features_intersect__{feat_data.alias}__{label}__{ts}.bed"),
        compress_bed=compress_bed
    )
    logger.info("Intersected feature BED written: %s (n=%d unique features)", features_bed, len(feat_ids_sorted))

    # Loop classification: 0/1/2 anchors hit (anchor is binary: ≥1 feature = hit)
    anchor_hit = set(inter["anchor_id"].unique().tolist())
    def a(i, side): return f"anchor_{i}_{side}"
    class_rows = []
    loops_no_idx, loops_one_idx, loops_both_idx = [], [], []
    for i in range(anchors.loop_count):
        h1 = int(a(i,1) in anchor_hit); h2 = int(a(i,2) in anchor_hit)
        n_hit_anchors = h1 + h2
        if n_hit_anchors == 0:
            cat = "none"; loops_no_idx.append(i)
        elif n_hit_anchors == 1:
            cat = "one";  loops_one_idx.append(i)
        else:
            cat = "both"; loops_both_idx.append(i)
        class_rows.append({"loop_index": i, "anchor1_hit": h1, "anchor2_hit": h2, "category": cat})

    class_df = pd.DataFrame(class_rows)
    class_tsv = tables_dir / f"classification__{label}__{ts}.tsv"
    class_df.to_csv(class_tsv, sep="\t", index=False)
    logger.info("Loop classification written: %s", class_tsv)

    # Loop BEDPE subsets (FOUR categories)
    bedpe_dir = set_dir / "bedpe"; ensure_dir(bedpe_dir)
    def write_subset(idxs: List[int], name: str) -> Path:
        fp = bedpe_dir / f"{name}__{label}__{ts}.bedpe"
        with open(fp, "w") as out:
            for idx in idxs:
                out.write(anchors.bedpe_lines[idx] + "\n")
        return fp

    loops_any_idx = sorted(loops_one_idx + loops_both_idx)
    loops_nohit_bedpe   = write_subset(loops_no_idx,   "loops_nohit")     # 0 anchors hit
    loops_one_bedpe     = write_subset(loops_one_idx,  "loops_onehit")    # 1 anchor hit
    loops_both_bedpe    = write_subset(loops_both_idx, "loops_bothhit")   # 2 anchors hit
    loops_any_bedpe     = write_subset(loops_any_idx,  "loops_anyhit")    # ≥1 anchor hit

    logger.info("Loop BEDPE subsets written: nohit=%d, onehit=%d, bothhit=%d, anyhit(≥1)=%d",
                len(loops_no_idx), len(loops_one_idx), len(loops_both_idx), len(loops_any_idx))

    # Anchor‑centric summary
    anchors_with_hit = len(anchor_hit)
    anchor_hit_rate = anchors_with_hit / anchors_total if anchors_total else 0.0
    df6 = pd.read_table(bedpe_fp, comment="#", header=None, usecols=[1,2,4,5])
    widths = pd.concat([(df6[2]-df6[1]), (df6[5]-df6[4])], ignore_index=True).astype(int)
    median_w = float(widths.median()) if len(widths) else float("nan")
    features_per_10k_anchors = len(feat_ids_sorted) / anchors_total * 10000 if anchors_total else float("nan")
    features_per_Mb_anchor   = len(feat_ids_sorted) / (anchors_total * median_w) * 1e6 if anchors_total and median_w>0 else float("nan")

    anchor_summary = pd.DataFrame([{
        "set": label,
        "feature_alias": feat_data.alias,
        "loops_total": anchors.loop_count,
        "anchors_total": anchors_total,
        "anchors_with_≥1_feature": anchors_with_hit,
        "anchor_hit_rate": anchor_hit_rate,
        "unique_features_in_set": len(feat_ids_sorted),
        "unique_features_per_10k_anchors": features_per_10k_anchors,
        "unique_features_per_Mb_anchor": features_per_Mb_anchor,
        "median_anchor_width_bp_from_BEDPE": median_w,
        # four loop categories
        "loops_nohit":   len(loops_no_idx),
        "loops_onehit":  len(loops_one_idx),
        "loops_bothhit": len(loops_both_idx),
        "loops_anyhit":  len(loops_any_idx),
    }])
    anchor_summary_tsv = tables_dir / f"anchor_summary__{feat_data.alias}__{label}__{ts}.tsv"
    anchor_summary.to_csv(anchor_summary_tsv, sep="\t", index=False)
    logger.info("Anchor summary written: %s", anchor_summary_tsv)

    # Metadata
    meta_dir = set_dir / "metadata"; ensure_dir(meta_dir)
    meta = {
        "timestamp": ts,
        "script": Path(__file__).name,
        "version": "4.0.0",
        "params": {
            "label": label,
            "feature_alias": feat_data.alias,
            "feature_bed": str(feat_data.path.resolve()),
            "bedpe": str(bedpe_fp.resolve()),
            "window_half_width": window_half_width,
            "threads": threads,
            "compress_bed": compress_bed,
        },
        "hashes": {
            "feature_bed_sha256": sha256_of_file(feat_data.path),
            "bedpe_sha256": sha256_of_file(bedpe_fp),
        },
        "counts": {
            "loops_total": anchors.loop_count,
            "anchors_total": anchors_total,
            "pairs": int(len(inter)),
            "unique_features": int(len(feat_ids_sorted)),
            "loops_nohit":   len(loops_no_idx),
            "loops_onehit":  len(loops_one_idx),
            "loops_bothhit": len(loops_both_idx),
            "loops_anyhit":  len(loops_any_idx),
        },
        "outputs": {
            "features_bed": str(features_bed),
            "loops_nohit_bedpe":   str(loops_nohit_bedpe),
            "loops_one_bedpe":     str(loops_one_bedpe),
            "loops_both_bedpe":    str(loops_both_bedpe),
            "loops_any_bedpe":     str(loops_any_bedpe),
            "pairs_tsv_gz": str(pairs_tsv_gz),
            "classification_tsv": str(class_tsv),
            "anchor_summary_tsv": str(anchor_summary_tsv),
            "log": str(log_path),
        },
    }
    with open(meta_dir / f"metadata__{label}__{ts}.json", "w") as fh:
        json.dump(meta, fh, indent=2)
    logger.info("Metadata written.")

    return SetOutputs(
        label=label, base_dir=set_dir, features_bed=features_bed,
        loops_nohit_bedpe=loops_nohit_bedpe,
        loops_one_bedpe=loops_one_bedpe,
        loops_both_bedpe=loops_both_bedpe,
        loops_any_bedpe=loops_any_bedpe,
        pairs_tsv_gz=pairs_tsv_gz, class_tsv=class_tsv, anchor_summary_tsv=anchor_summary_tsv,
        feat_ids=set(feat_ids_sorted), log_path=log_path,
    )

# =============================== Cross‑set (UpSet + Stats) ================= #

def try_import_scipy() -> bool:
    try:
        import scipy.stats as st  # noqa
        return True
    except Exception:
        return False

def binom_one_two_sided(a: int, b: int) -> Tuple[float, float]:
    n = a + b
    if n == 0:
        return float("nan"), float("nan")
    if try_import_scipy():
        import scipy.stats as st
        one = float(st.binomtest(a, n, 0.5, alternative="greater").pvalue)
        two = float(st.binomtest(a, n, 0.5, alternative="two-sided").pvalue)
        return one, two
    # Normal approx with continuity correction
    mu = n * 0.5; sigma = math.sqrt(n * 0.25)
    z = (a - mu - 0.5) / sigma
    from math import erf, sqrt
    sf = 0.5 * (1.0 - erf(z / sqrt(2)))
    two = min(1.0, 2.0 * min(sf, 1.0 - sf))
    return sf, two

def wilson_ci(p: float, n: int) -> Tuple[float, float]:
    if n == 0: return float("nan"), float("nan")
    z = 1.959963984540054
    denom = 1 + z*z/n
    center = (p + z*z/(2*n)) / denom
    half = z * math.sqrt(p*(1-p)/n + z*z/(4*n*n)) / denom
    return max(0.0, center - half), min(1.0, center + half)

def odds_from_p(p: float) -> float:
    if p <= 0: return 0.0
    if p >= 1: return float("inf")
    return p / (1 - p)

def write_kv_tsv(rows: List[Tuple[str, int | str]], fp: Path, header=("metric","value")) -> None:
    ensure_dir(fp.parent)
    with open(fp, "w") as out:
        out.write("\t".join(header) + "\n")
        for k, v in rows:
            out.write(f"{k}\t{v}\n")

def plot_upset(combos: List[Tuple[int,int,int,int]],
               include_neither: bool,
               out_png: Path, out_pdf: Path,
               title: str,
               set_labels: Tuple[str,str,str]) -> None:
    import matplotlib
    import matplotlib.pyplot as plt
    matplotlib.rcParams["pdf.fonttype"] = 42
    matplotlib.rcParams["ps.fonttype"]  = 42
    base_fs = 16
    matplotlib.rcParams["font.size"] = base_fs
    matplotlib.rcParams["axes.titlesize"] = base_fs+2
    matplotlib.rcParams["axes.labelsize"] = base_fs
    matplotlib.rcParams["xtick.labelsize"] = base_fs-1
    matplotlib.rcParams["ytick.labelsize"] = base_fs-1

    # sort combos by count desc; append Neither last if requested
    filtered = []
    neither_row = None
    for m,c,d,n in combos:
        if (m,c,d) == (0,0,0):
            neither_row = (m,c,d,n)
        else:
            filtered.append((m,c,d,n))
    filtered.sort(key=lambda x: x[3], reverse=True)
    if include_neither and neither_row is not None:
        filtered.append(neither_row)

    labels = []; values = []; matrix = []
    for m,c,d,n in filtered:
        on = [m,c,d]
        lab = " | ".join([lbl for lbl,flag in zip(set_labels,on) if flag]) if any(on) else "Neither"
        labels.append(lab); values.append(n); matrix.append(on)
    matrix = list(zip(*matrix)) if matrix else [[],[],[]]
    N = len(values)

    fig_w = max(10, min(24, 0.9 * max(N, 10)))
    fig = plt.figure(figsize=(fig_w, 8), facecolor="white")

    ax_bar = fig.add_axes([0.08, 0.55, 0.84, 0.38])
    ax_bar.bar(range(N), values, color="black")
    ax_bar.set_ylabel("Feature count")
    ax_bar.set_xticks(range(N)); ax_bar.set_xticklabels([])
    for i, v in enumerate(values):
        ax_bar.text(i, v, f"{v:,}", ha="center", va="bottom", fontsize=base_fs-2)
    ax_bar.set_title(title)

    ax_mat = fig.add_axes([0.08, 0.20, 0.84, 0.28])
    ax_mat.set_xlim(-0.5, N-0.5 if N>0 else 0.5); ax_mat.set_ylim(-0.5, 2.5)
    ax_mat.set_yticks([0,1,2]); ax_mat.set_yticklabels(list(set_labels))
    ax_mat.set_xticks(range(N)); ax_mat.set_xticklabels(labels, rotation=30, ha="right")
    for col in range(N):
        for row in range(3):
            if matrix[row][col] == 1:
                ax_mat.plot(col, row, 'o', markersize=10, color="black")
            else:
                ax_mat.plot(col, row, 'o', mfc='none', mec='black', markersize=10)
    for ax in (ax_bar, ax_mat):
        ax.grid(False)
        for s in ax.spines.values(): s.set_visible(True)
    ensure_dir(out_png.parent)
    fig.savefig(out_png, dpi=300, bbox_inches="tight")
    fig.savefig(out_pdf, bbox_inches="tight")
    plt.close(fig)

# =============================== CLI / Main ================================ #

def parse_args(argv: Optional[List[str]] = None) -> argparse.Namespace:
    p = argparse.ArgumentParser(
        description="Loop–Feature intersect (BED↔BEDPE) + cross‑set UpSet & stats (A, A∩B, B).",
        formatter_class=argparse.ArgumentDefaultsHelpFormatter,
    )
    p.add_argument("--feature-bed", required=True, type=Path, help="Generic feature BED (.bed or .bed.gz).")
    p.add_argument("--feature-alias", type=str, default=None, help="Short alias (used in file names).")
    p.add_argument("--bedpe-a", required=True, type=Path, help="Method A BEDPE.")
    p.add_argument("--bedpe-common", required=True, type=Path, help="Common A∩B BEDPE.")
    p.add_argument("--bedpe-b", required=True, type=Path, help="Method B BEDPE.")
    p.add_argument("--label-a", type=str, default=None, help="Label for Method A (plot/paths).")
    p.add_argument("--label-common", type=str, default=None, help="Label for Common (plot/paths).")
    p.add_argument("--label-b", type=str, default=None, help="Label for Method B (plot/paths).")
    p.add_argument("--master-bed", type=Path, help="Master feature BED (.bed or .bed.gz) for 'Neither' & absolute stats.")
    p.add_argument("--window-half-width", type=int, default=None,
                   help="If set, build windowed anchors centered at midpoints (±W bp).")
    p.add_argument("--chrom-sizes", type=Path, help="Chrom sizes for window clipping and sorting.")
    p.add_argument("--threads", type=int, default=1)
    p.add_argument("--outdir", type=Path, default=Path("loop_feature_intersect_out"))
    p.add_argument("--compress-bed", action="store_true",
                   help="Write feature BED outputs as .bed.gz (default: plain .bed).")
    p.add_argument("--log-level", default="INFO", choices=["DEBUG","INFO","WARNING","ERROR"])
    return p.parse_args(argv)

def make_logger(out_root: Path, level: str) -> logging.Logger:
    ensure_dir(out_root)
    log_fp = out_root / f"run__{ts_now()}.log"
    logging.basicConfig(
        level=getattr(logging, level),
        format="%(asctime)s [%(levelname)s] %(message)s",
        handlers=[logging.FileHandler(log_fp), logging.StreamHandler(sys.stdout)],
    )
    logger = logging.getLogger("loop_feat_combined")
    logger.info("Log file: %s", log_fp)
    return logger

def main(argv: Optional[List[str]] = None) -> int:
    args = parse_args(argv)
    ts = ts_now()
    out_root = args.outdir / ts
    logger = make_logger(out_root, args.log_level)
    logger.info("♻️  Combined loop–feature pipeline started. Version 4.0.0")

    chrom_sizes = read_chrom_sizes(args.chrom_sizes)

    # Stage 1: per‑set intersects
    feat = read_feature_bed(args.feature_bed, args.feature_alias)
    feat_bt, id_to_line, id_to_coord = feature_bedtool(feat)
    label_a      = args.label_a      or args.bedpe_a.stem
    label_common = args.label_common or args.bedpe_common.stem
    label_b      = args.label_b      or args.bedpe_b.stem

    setA = process_one_set(
        label=label_a, bedpe_fp=args.bedpe_a, feat_data=feat, feat_bt=feat_bt,
        id_to_line=id_to_line, id_to_coord=id_to_coord, chrom_sizes=chrom_sizes,
        window_half_width=args.window_half_width, threads=args.threads,
        out_root=out_root, ts=ts, compress_bed=args.compress_bed, logger_parent=logger,
    )
    setAB = process_one_set(
        label=label_common, bedpe_fp=args.bedpe_common, feat_data=feat, feat_bt=feat_bt,
        id_to_line=id_to_line, id_to_coord=id_to_coord, chrom_sizes=chrom_sizes,
        window_half_width=args.window_half_width, threads=args.threads,
        out_root=out_root, ts=ts, compress_bed=args.compress_bed, logger_parent=logger,
    )
    setB = process_one_set(
        label=label_b, bedpe_fp=args.bedpe_b, feat_data=feat, feat_bt=feat_bt,
        id_to_line=id_to_line, id_to_coord=id_to_coord, chrom_sizes=chrom_sizes,
        window_half_width=args.window_half_width, threads=args.threads,
        out_root=out_root, ts=ts, compress_bed=args.compress_bed, logger_parent=logger,
    )

    # Stage 2: cross‑set analysis (features)
    cross_dir = out_root / "cross_set"; ensure_dir(cross_dir)
    A, AB, B = setA.feat_ids, setAB.feat_ids, setB.feat_ids
    union = A | AB | B

    # 7 disjoint Venn cells
    only_A   = A  - (AB | B)
    only_AB  = AB - (A  | B)
    only_B   = B  - (A  | AB)
    A_AB     = (A & AB) - B
    A_B      = (A & B)  - AB
    AB_B     = (AB & B) - A
    A_AB_B   =  A & AB & B
    cells = [only_A, only_AB, only_B, A_AB, A_B, AB_B, A_AB_B]
    assert sum(len(s) for s in cells) == len(union), "Sum of 7 cells must equal union size."

    # Optional master to compute “Neither”
    neither_ids: Optional[set] = None
    master_total: Optional[int] = None
    master_ids: Optional[set] = None
    master_id_to_line: Dict[str,str] = {}
    master_id_to_coord: Dict[str,Tuple[str,int,int]] = {}
    if args.master_bed and args.master_bed.exists():
        master_fd = read_feature_bed(args.master_bed, alias=None)
        master_id_to_line = dict(zip(master_fd.ids, master_fd.lines))
        master_id_to_coord = dict(zip(master_fd.ids, [(r[0],r[1],r[2]) for r in master_fd.rows]))
        master_ids = set(master_fd.ids)
        master_total = len(master_ids)
        neither_ids = master_ids - union
        logger.info("Master present: total=%d  Neither=%d", master_total, len(neither_ids))
    else:
        logger.warning("No master BED provided or not found. Skipping 'Neither' and absolute stats.")

    # Write summary & combo counts
    summary_rows = [
        ("label_A", label_a), ("label_common", label_common), ("label_B", label_b),
        ("feature_alias", feat.alias),
        ("A_total", len(A)), ("AB_total", len(AB)), ("B_total", len(B)),
        ("union_total", len(union)),
        ("venn_A_only", len(only_A)),
        ("venn_AB_only", len(only_AB)),
        ("venn_B_only", len(only_B)),
        ("venn_A_and_AB_only", len(A_AB)),
        ("venn_A_and_B_only", len(A_B)),
        ("venn_AB_and_B_only", len(AB_B)),
        ("venn_all_three", len(A_AB_B)),
    ]
    if master_total is not None:
        summary_rows.extend([("master_total", master_total), ("neither_total", len(neither_ids))])
    write_kv_tsv(summary_rows, cross_dir / "summary.tsv")

    combos_union = [
        (1,0,0, len(only_A)),
        (0,1,0, len(only_AB)),
        (0,0,1, len(only_B)),
        (1,1,0, len(A_AB)),
        (1,0,1, len(A_B)),
        (0,1,1, len(AB_B)),
        (1,1,1, len(A_AB_B)),
    ]
    with open(cross_dir / "combo_counts_union.tsv", "w") as out:
        out.write("in_A\tin_AB\tin_B\tcount\n")
        for m,c,d,n in combos_union: out.write(f"{m}\t{c}\t{d}\t{n}\n")
    if master_total is not None and neither_ids is not None:
        combos_with_neither = combos_union + [(0,0,0, len(neither_ids))]
        with open(cross_dir / "combo_counts_with_neither.tsv", "w") as out:
            out.write("in_A\tin_AB\tin_B\tcount\n")
            for m,c,d,n in combos_with_neither: out.write(f"{m}\t{c}\t{d}\t{n}\n")

    # Helper for Stage‑2 feature BED writing that respects --compress-bed and master fallback
    def write_feat_bed_from_ids(idset: set, name: str) -> Path:
        # choose coordinates for sorting: prefer Stage‑1 mapping; fallback to master mapping
        def coord_of(tid: str) -> Tuple[str,int,int]:
            if tid in id_to_coord: return id_to_coord[tid]
            if tid in master_id_to_coord: return master_id_to_coord[tid]
            return ("chrZZZ", 10**12, 10**12)
        ids_sorted = sorted(
            idset, key=lambda tid: (natural_chrom_key(coord_of(tid)[0]), coord_of(tid)[1], coord_of(tid)[2])
        )
        # choose exact line source: prefer Stage‑1; fallback to master
        lines: List[str] = []
        for tid in ids_sorted:
            if tid in id_to_line:
                lines.append(id_to_line[tid])
            elif tid in master_id_to_line:
                lines.append(master_id_to_line[tid])
            else:
                continue
        base = cross_dir / f"features_{name}__{feat.alias}__{ts}.bed"
        return write_feature_bed_lines(lines, base, compress_bed=args.compress_bed)

    _ = write_feat_bed_from_ids(only_A,  "A_only")
    _ = write_feat_bed_from_ids(only_AB, "AB_only")
    _ = write_feat_bed_from_ids(only_B,  "B_only")
    _ = write_feat_bed_from_ids(A_AB,    "A_and_AB_only")
    _ = write_feat_bed_from_ids(A_B,     "A_and_B_only")
    _ = write_feat_bed_from_ids(AB_B,    "AB_and_B_only")
    _ = write_feat_bed_from_ids(A_AB_B,  "A_and_AB_and_B")
    if master_total is not None and neither_ids is not None:
        _ = write_feat_bed_from_ids(neither_ids, "Neither")

    # ---------------------- McNemar / binomial tests ----------------------
    A_detected = A | AB
    B_detected = B | AB
    A_only_u   = len(A_detected - B_detected)
    B_only_u   = len(B_detected - A_detected)
    Both_u     = len(A_detected & B_detected)
    assert A_only_u + B_only_u + Both_u == len(union), "Union mismatch in stats."

    p_one_u, p_two_u = binom_one_two_sided(B_only_u, A_only_u)  # one‑sided tests B>A
    n_u = A_only_u + B_only_u
    p_hat_u = (B_only_u / n_u) if n_u > 0 else float("nan")
    lo_u, hi_u = wilson_ci(p_hat_u, n_u) if n_u>0 else (float("nan"), float("nan"))
    OR_u, ORlo_u, ORhi_u = (odds_from_p(p_hat_u), odds_from_p(lo_u), odds_from_p(hi_u)) if n_u>0 else (float("nan"),)*3

    stats_union_rows = [
        ("scope",                    "Union‑only (detected by ≥1)"),
        (f"{label_b}_only",         B_only_u),
        (f"{label_a}_only",         A_only_u),
        ("Both",                    Both_u),
        ("Union_total",             len(union)),
        (f"one_sided_p({label_b}>{label_a})", f"{p_one_u:.3e}"),
        ("two_sided_p",             f"{p_two_u:.3e}"),
        (f"OR_hat ({label_b}/{label_a})", f"{OR_u:.3f}"),
        ("OR_95%_CI_low",           f"{ORlo_u:.3f}"),
        ("OR_95%_CI_high",          f"{ORhi_u:.3f}"),
    ]
    write_kv_tsv(stats_union_rows, cross_dir / "stats_union_mcnemar.tsv")
    logger.info("INTERPRETATION (union): %s vs %s among detected features — %s-only=%d, %s-only=%d; one‑sided p=%s; OR≈%s.",
                label_b, label_a, label_b, B_only_u, label_a, A_only_u, f"{p_one_u:.3e}", f"{OR_u:.2f}")

    # Absolute vs master (if available)
    if master_total is not None and master_ids is not None:
        B_only_abs   = len((B_detected - A_detected) & master_ids)
        A_only_abs   = len((A_detected - B_detected) & master_ids)
        Both_abs     = len((A_detected & B_detected) & master_ids)
        Neither_abs  = len(master_ids - (A_detected | B_detected))
        assert B_only_abs + A_only_abs + Both_abs + Neither_abs == len(master_ids)

        p_one_a, p_two_a = binom_one_two_sided(B_only_abs, A_only_abs)
        n_a = B_only_abs + A_only_abs
        p_hat_a = (B_only_abs / n_a) if n_a>0 else float("nan")
        lo_a, hi_a = wilson_ci(p_hat_a, n_a) if n_a>0 else (float("nan"), float("nan"))
        OR_a, ORlo_a, ORhi_a = (odds_from_p(p_hat_a), odds_from_p(lo_a), odds_from_p(hi_a)) if n_a>0 else (float("nan"),)*3

        stats_abs_rows = [
            ("scope",                    "Absolute vs master"),
            ("N_master",                 len(master_ids)),
            (f"{label_b}_only",         B_only_abs),
            (f"{label_a}_only",         A_only_abs),
            ("Both",                    Both_abs),
            ("Neither",                 Neither_abs),
            (f"one_sided_p({label_b}>{label_a})", f"{p_one_a:.3e}"),
            ("two_sided_p",             f"{p_two_a:.3e}"),
            (f"OR_hat ({label_b}/{label_a})", f"{OR_a:.3f}"),
            ("OR_95%_CI_low",           f"{ORlo_a:.3f}"),
            ("OR_95%_CI_high",          f"{ORhi_a:.3f}"),
        ]
        write_kv_tsv(stats_abs_rows, cross_dir / "stats_absolute_mcnemar.tsv")
        logger.info("INTERPRETATION (absolute): %s vs %s over master — %s-only=%d, %s-only=%d; one‑sided p=%s; OR≈%s.",
                    label_b, label_a, label_b, B_only_abs, label_a, A_only_abs, f"{p_one_a:.3e}", f"{OR_a:.2f}")

    # ------------------------------ UpSet plots ----------------------------
    try:
        plot_upset(
            combos=combos_union, include_neither=False,
            out_png=cross_dir / "upset_union.png", out_pdf=cross_dir / "upset_union.pdf",
            title=f"UpSet (Union only): {feat.alias}",
            set_labels=(label_a, label_common, label_b),
        )
        logger.info("UpSet (union) written.")
    except Exception as e:
        logger.exception("Failed to plot UpSet (union): %s", e)
    if master_total is not None and neither_ids is not None:
        try:
            combos_with_neither = combos_union + [(0,0,0, len(neither_ids))]
            plot_upset(
                combos=combos_with_neither, include_neither=True,
                out_png=cross_dir / "upset_with_neither.png", out_pdf=cross_dir / "upset_with_neither.pdf",
                title=f"UpSet (With Neither): {feat.alias}",
                set_labels=(label_a, label_common, label_b),
            )
            logger.info("UpSet (with neither) written.")
        except Exception as e:
            logger.exception("Failed to plot UpSet (with neither): %s", e)

    logger.info("🎉  Completed. Outputs in: %s", out_root)
    print(f"[OK] Outputs: {out_root}")
    return 0

if __name__ == "__main__":
    try:
        sys.exit(main(sys.argv[1:]))
    except Exception as e:
        logging.exception("Fatal error: %s", e)
        sys.exit(1)
```
