## Supplementary material for "CAD-C: An engineered nuclease enables repair-free *in situ* proximity ligation and nucleosome-resolution chromosome walks in human cells": Custom code: flip_and_combine_plot_vplot_matrices_1_20250308.py

```
#!/usr/bin/env python3

"""
flip_sum_vplot.py

Example usage:
  python flip_sum_vplot.py \
    --plus_vplot plus_strand/vplot_matrix.tsv \
    --minus_vplot minus_strand/vplot_matrix.tsv \
    --window_bp 1500 \
    --ymax_fraglen 200 \
    --output_dir final_flipped_summed_vplot

What it does:
  1) Reads plus_vplot => shape=(y_bins, x_bins)
  2) Reads minus_vplot => shape=(y_bins, x_bins)
  3) Flips minus horizontally => np.fliplr
  4) Sums with plus => combined
  5) Writes vplot_matrix_combined.tsv
  6) Plots combined v-plot in a single PDF, with x in [-window_bp..+window_bp], y in [0..ymax_fraglen]
"""

import os
import sys
import argparse
import datetime
import logging
import numpy as np
import matplotlib
matplotlib.use("Agg")
import matplotlib.pyplot as plt
import seaborn as sns

logging.basicConfig(level=logging.INFO, format="%(asctime)s %(levelname)s: %(message)s")

def parse_args():
    p = argparse.ArgumentParser(description="Flip minus-strand v-plot matrix in x, sum with plus, write combined .tsv + PDF.")
    p.add_argument("--plus_vplot", required=True,
                   help="Path to the plus-strand vplot_matrix.tsv (2D ASCII).")
    p.add_argument("--minus_vplot", required=True,
                   help="Path to the minus-strand vplot_matrix.tsv (2D ASCII).")
    p.add_argument("--window_bp", type=int, default=1000,
                   help="Half-range in x => x in [-window_bp..+window_bp].")
    p.add_argument("--ymax_fraglen", type=int, default=200,
                   help="Height in y => y in [0..ymax_fraglen]. Must match the shape of the matrix.")
    p.add_argument("--output_dir", default=".",
                   help="Directory to write combined v-plot matrix + PDF.")
    return p.parse_args()

def read_vplot_tsv(filepath):
    """
    Reads an ASCII 2D v-plot matrix => (y_rows, x_cols).
    Each line => one row. Space-separated floats.
    Returns np.array shape=(y,x).
    """
    arr=[]
    with open(filepath,"r") as f:
        for line in f:
            line=line.strip()
            if not line:
                continue
            parts=line.split()
            rowvals= [float(x) for x in parts]
            arr.append(rowvals)
    vmat= np.array(arr)
    return vmat

def write_vplot_tsv(filepath, vmat):
    """
    Writes 2D np.array => lines of space-separated floats.
    """
    with open(filepath,"w") as w:
        for row in vmat:
            line_data= " ".join(
                str(int(x)) if x.is_integer() else f"{x:.2f}"
                for x in row
            )
            w.write(line_data + "\n")
    logging.info(f"Wrote combined vplot => {filepath}")

def flip_vplot_x(vmat):
    """
    Flip horizontally => vmat[:, ::-1].
    Returns a new array.
    """
    return np.fliplr(vmat)

def main():
    args= parse_args()

    ts= datetime.datetime.now().strftime("%Y%m%d_%H%M%S")
    out_sub= f"flip_sum_vplot_{ts}"
    out_dir= os.path.join(args.output_dir, out_sub)
    os.makedirs(out_dir, exist_ok=True)

    # write metadata
    meta_path= os.path.join(out_dir, "metadata.txt")
    with open(meta_path,"w") as mf:
        mf.write(f"Timestamp: {ts}\n")
        mf.write(f"plus_vplot: {args.plus_vplot}\n")
        mf.write(f"minus_vplot: {args.minus_vplot}\n")
        mf.write(f"window_bp: {args.window_bp}\n")
        mf.write(f"ymax_fraglen: {args.ymax_fraglen}\n")
        mf.write(f"output_dir: {out_dir}\n")

    logging.info(f"Output => {out_dir}")
    logging.info(f"Reading plus vplot => {args.plus_vplot}")
    vm_plus= read_vplot_tsv(args.plus_vplot)

    logging.info(f"Reading minus vplot => {args.minus_vplot}")
    vm_minus= read_vplot_tsv(args.minus_vplot)

    # shape check
    if vm_plus.shape != vm_minus.shape:
        logging.error(f"Shape mismatch: plus={vm_plus.shape}, minus={vm_minus.shape}")
        sys.exit(1)

    logging.info("Flipping minus horizontally..")
    vm_flip= flip_vplot_x(vm_minus)

    logging.info("Summing plus + flipped minus..")
    vm_comb= vm_plus + vm_flip

    # write out
    comb_tsv= os.path.join(out_dir, "vplot_matrix_combined.tsv")
    write_vplot_tsv(comb_tsv, vm_comb)

    # now plot
    logging.info("Plotting combined v-plot => PDF.")
    pdf_path= os.path.join(out_dir, "vplot_combined.pdf")

    plt.rcParams["pdf.fonttype"] = 42
    plt.rcParams["ps.fonttype"]  = 42

    fig= plt.figure(figsize=(10,6))
    # shape => (y_bins, x_bins)
    y_bins, x_bins= vm_comb.shape

    cmap= sns.color_palette("mako_r", as_cmap=True)
    vmax= vm_comb.max() if vm_comb.max()>0 else 1

    # We'll define extent => x in [-window_bp.. +window_bp], y in [0..ymax_fraglen]
    x_min= -args.window_bp
    x_max= +args.window_bp
    y_min= 0
    y_max= args.ymax_fraglen

    ax= fig.add_subplot(111)
    im= ax.imshow(
        vm_comb,
        origin='lower',
        cmap=cmap,
        aspect='auto',
        extent=[x_min,x_max,y_min,y_max],
        vmin=0, vmax=vmax,
        interpolation='none'
    )
    cb= fig.colorbar(im, ax=ax, fraction=0.046, pad=0.04)
    ax.set_title("Combined V-plot (plus + flipped minus)")
    ax.set_xlabel("Midpoint dist (bp)")
    ax.set_ylabel("Fragment length (bp)")
    ax.set_xlim(x_min, x_max)
    ax.set_ylim(y_min, y_max)

    plt.tight_layout()
    plt.savefig(pdf_path, dpi=300)
    plt.close(fig)

    logging.info(f"Done. Wrote combined v-plot => {comb_tsv}")
    logging.info(f"Saved PDF => {pdf_path}")

if __name__=="__main__":
    main()
```
