## Supplementary material for "CAD-C: An engineered nuclease enables repair-free *in situ* proximity ligation and nucleosome-resolution chromosome walks in human cells": Custom code: generate_and_plot_vplot_alone_1_20250308.py

```
#!/usr/bin/env python3

"""
vplot_generation_from_walks.py

Example usage:
  python vplot_generation_from_walks.py \
    --walks_file my_abc_3frag.tsv.gz \
    --bed_file motif_intervals.bed \
    --window_bp 1500 \
    --chunk_size 1000000 \
    --ymax_fraglen 200 \
    --output_dir vplot_out

Description:
  1) Read BED, expand ±window_bp => "motif windows".
  2) Read .tsv.gz in chunk_size lines. For each chunk, parse (A,B,C).
  3) Overlap with motif windows => for each overlap:
       - dist_mid = midpoint - motif_center
       - length = fragment_end - fragment_start
       - fill [y_bin, x_bin] => vplot_matrix[y_bin, x_bin]
     ignoring fragments with length>ymax_fraglen
  4) Write final vplot_matrix.tsv (ASCII 2D).
  5) Plot it in a PDF if desired.
"""

import os
import sys
import argparse
import datetime
import logging
import gzip
import numpy as np
import pandas as pd
import polars as pl
import pyranges as pr
import matplotlib
matplotlib.use("Agg")
import matplotlib.pyplot as plt
import seaborn as sns

logging.basicConfig(level=logging.INFO, format="%(asctime)s %(levelname)s: %(message)s")

def parse_args():
    p = argparse.ArgumentParser()
    p.add_argument("--walks_file", required=True,
                   help="3-fragment .tsv.gz (A_chrom, A_start, A_end, B..., C...).")
    p.add_argument("--bed_file", required=True,
                   help="BED intervals for motifs. We'll expand ±window_bp => motif windows.")
    p.add_argument("--window_bp", type=int, default=1000,
                   help="± window around motif center for overlap (default=1000).")
    p.add_argument("--ymax_fraglen", type=int, default=200,
                   help="Max fragment length to keep in V-plot (default=200). Others are ignored.")
    p.add_argument("--chunk_size", type=int, default=500000,
                   help="Lines per chunk from walks_file.")
    p.add_argument("--output_dir", default=".",
                   help="Where to write the vplot_matrix.tsv and pdf.")
    return p.parse_args()

def expand_bed_motifs(bed_file, window_bp):
    """
    Read bed => for each line => center => expand ±window_bp => [Chromosome, Start, End].
    """
    motifs=[]
    with open(bed_file,"r") as f:
        for line in f:
            line=line.strip()
            if not line or line.startswith("#"):
                continue
            parts=line.split("\t")
            chrom= parts[0]
            st= int(parts[1])
            en= int(parts[2])
            center= (st+en)//2
            wstart= max(center - window_bp, 0)
            wend= center + window_bp
            motifs.append([chrom, wstart, wend])
    df= pd.DataFrame(motifs, columns=["Chromosome","Start","End"])
    return df

def write_vplot_tsv(filepath, vmat):
    with open(filepath,"w") as w:
        for row in vmat:
            line_data= " ".join(str(int(x)) if x.is_integer() else f"{x:.2f}" for x in row)
            w.write(line_data+"\n")
    logging.info(f"Wrote vplot => {filepath}")

def main():
    args= parse_args()

    ts= datetime.datetime.now().strftime("%Y%m%d_%H%M%S")
    out_dir= os.path.join(args.output_dir, f"vplot_{ts}")
    os.makedirs(out_dir, exist_ok=True)

    meta_file= os.path.join(out_dir, "metadata.txt")
    with open(meta_file,"w") as mf:
        mf.write(f"Timestamp: {ts}\n")
        mf.write(f"walks_file: {args.walks_file}\n")
        mf.write(f"bed_file: {args.bed_file}\n")
        mf.write(f"window_bp: {args.window_bp}\n")
        mf.write(f"ymax_fraglen: {args.ymax_fraglen}\n")
        mf.write(f"chunk_size: {args.chunk_size}\n")

    logging.info(f"Output => {out_dir}")
    logging.info("Reading & expanding bed => motif windows..")
    motif_df= expand_bed_motifs(args.bed_file, args.window_bp)
    motif_py= pr.PyRanges(motif_df)

    # We'll define x_bins => 2*window_bp+1 => but let's do a direct approach: we won't store coverage, only v-plot
    # We'll define pass1 => find max length if user doesn't set? But user sets a fixed y=0..ymax_fraglen => easy.

    # Let's define x_bins => 2*args.window_bp+1 or so, but for a more flexible approach, we can do "bin_size"? 
    # Actually, we might do the approach: x dimension => (2*window_bp)//bin_size + 1. 
    # We can do that or do a single bin per bp? We'll do the simpler approach => 1 bin per bp => x_bins=2*window_bp+1. 
    # Then dist= [-window_bp..+window_bp].
    x_bins= 2*args.window_bp + 1
    y_bins= args.ymax_fraglen
    vmat= np.zeros((y_bins, x_bins), dtype=float)

    def dist_to_xidx(dist):
        # dist in [-window_bp.. +window_bp]
        shift= dist+ args.window_bp
        if shift<0 or shift>= x_bins:
            return -1
        return int(shift)

    logging.info(f"Building vplot matrix => y in [0..{args.ymax_fraglen-1}], x in [-{args.window_bp}..+{args.window_bp}] => shape=({y_bins},{x_bins})")

    colnames= [
        "readID","A_chrom","A_start","A_end","A_strand",
        "B_chrom","B_start","B_end","B_strand",
        "C_chrom","C_start","C_end","C_strand"
    ]
    lines_buffer=[]
    lines_read=0

    def process_chunk(df_chunk: pl.DataFrame):
        if df_chunk.is_empty():
            return
        rows=[]
        for i in range(len(df_chunk)):
            A_ch= df_chunk["A_chrom"][i]
            A_st= int(df_chunk["A_start"][i])
            A_en= int(df_chunk["A_end"][i])
            if A_st>A_en: A_st,A_en= A_en,A_st

            B_ch= df_chunk["B_chrom"][i]
            B_st= int(df_chunk["B_start"][i])
            B_en= int(df_chunk["B_end"][i])
            if B_st>B_en: B_st,B_en= B_en,B_st

            C_ch= df_chunk["C_chrom"][i]
            C_st= int(df_chunk["C_start"][i])
            C_en= int(df_chunk["C_end"][i])
            if C_st>C_en: C_st,C_en= C_en,C_st

            # store all fragments
            rows.append([A_ch, A_st, A_en])
            rows.append([B_ch, B_st, B_en])
            rows.append([C_ch, C_st, C_en])

        frag_df= pd.DataFrame(rows, columns=["Chromosome","Start","End"])
        frag_py= pr.PyRanges(frag_df)
        overlap_py= frag_py.join(motif_py)
        overlap_df= overlap_py.df
        # overlap => columns [Chromosome, Start, End, Start_b, End_b, ...]
        for row in overlap_df.itertuples(index=False):
            fstart= getattr(row, "Start")
            fend=   getattr(row, "End")
            start_b= getattr(row, "Start_b")
            end_b=   getattr(row, "End_b")
            center= (start_b+ end_b)//2

            length= abs(fend- fstart)
            if length>= args.ymax_fraglen:
                continue
            # y bin => length
            ybin= length
            dist_mid= ((fstart + fend)//2) - center
            xbin= dist_to_xidx(dist_mid)
            if xbin>=0:
                vmat[ybin, xbin]+=1

    logging.info("Pass => reading walks file in chunks, filling vplot matrix..")
    with gzip.open(args.walks_file,"rt") as f:
        header_line= next(f).strip().split("\t")
        if len(header_line)< len(colnames):
            logging.error("Walks file has fewer columns than expected.")
            sys.exit(1)

        for line in f:
            line=line.strip()
            if not line: continue
            lines_buffer.append(line.split("\t"))
            lines_read+=1
            if lines_read== args.chunk_size:
                df_chunk= pl.DataFrame(lines_buffer, schema=colnames, orient="row")
                lines_buffer=[]
                lines_read=0
                process_chunk(df_chunk)
        if lines_buffer:
            df_chunk= pl.DataFrame(lines_buffer, schema=colnames, orient="row")
            process_chunk(df_chunk)

    # done => write vplot
    vplot_path= os.path.join(out_dir, "vplot_matrix.tsv")
    with open(vplot_path,"w") as w:
        for row in vmat:
            line_data= " ".join(str(int(x)) if x.is_integer() else f"{x:.2f}" for x in row)
            w.write(line_data+"\n")
    logging.info(f"Wrote vplot => {vplot_path}")

    # optional: plot
    pdf_path= os.path.join(out_dir, "vplot.pdf")
    logging.info(f"Plotting => {pdf_path}")

    plt.rcParams["pdf.fonttype"]=42
    plt.rcParams["ps.fonttype"]=42

    fig, ax= plt.subplots(figsize=(10,6))
    cmap= sns.color_palette("mako_r", as_cmap=True)
    vmax= vmat.max() if vmat.max()>0 else 1
    # x in [-window_bp..+window_bp], y in [0..ymax_fraglen]
    im= ax.imshow(
        vmat,
        origin='lower',
        cmap=cmap,
        aspect='auto',
        extent=[-args.window_bp, +args.window_bp, 0, args.ymax_fraglen],
        vmin=0, vmax=vmax,
        interpolation='none'
    )
    cb= plt.colorbar(im, ax=ax, fraction=0.046, pad=0.04)
    ax.set_title(f"V-plot from {args.walks_file}\n±{args.window_bp}, y=0..{args.ymax_fraglen-1}")
    ax.set_xlabel("Midpoint dist (bp)")
    ax.set_ylabel("Fragment length (bp)")
    plt.tight_layout()
    plt.savefig(pdf_path, dpi=300)
    plt.close(fig)

    logging.info(f"Done. V-plot => {out_dir}")

if __name__=="__main__":
    main()
```
