## Supplementary material for "CAD-C: An engineered nuclease enables repair-free *in situ* proximity ligation and nucleosome-resolution chromosome walks in human cells": Custom code: submit_append_bed_annotated_walks_with_fraglen_AB_BC_AC_midpointsep_v2_20250312.sh

```
#!/bin/bash
#SBATCH --job-name=fraglen
#SBATCH --output=fraglen_%j.log
#SBATCH --error=fraglen_%j.err
#SBATCH --cpus-per-task=4


# module load python/3.10  # if needed

INPUT_GZ="/rugpfs/fs0/risc_lab/scratch/jsoroczyns/CAD/CADwalk/CADwalk_E1_GM12878_2x250/results_20250207_0034_nonexpanded/ABC_walk_processing_20250209/20250214_Bfilt_flattened_parse2_UU_direct_ABC_complete_consequent12/filtzerolenB_flip_sort_dedup_split_20250214/merged_all/20250312_cis_intersect_ChromHMM_v2/merged_all_flipped_sorted.cis.ChromHMM_v2.tsv.gz"
OUTPUT_GZ="/rugpfs/fs0/risc_lab/scratch/jsoroczyns/CAD/CADwalk/CADwalk_E1_GM12878_2x250/results_20250207_0034_nonexpanded/ABC_walk_processing_20250209/20250214_Bfilt_flattened_parse2_UU_direct_ABC_complete_consequent12/filtzerolenB_flip_sort_dedup_split_20250214/merged_all/20250312_cis_intersect_ChromHMM_v2/merged_all_flipped_sorted.cis.ChromHMM_v2_fraglen_seps.tsv.gz"
CHUNKSIZE=5000000

gunzip -c "$INPUT_GZ" | \
  python /rugpfs/fs0/risc_lab/scratch/jsoroczyns/CAD/CAD-C/scripts/2025_03_12/append_bed_annotated_walks_with_fraglen_AB_BC_AC_midpointsep_v2_20250312.py \
    --output "$OUTPUT_GZ" \
    --chunksize "$CHUNKSIZE"
```
