## Supplementary material for "CAD-C: An engineered nuclease enables repair-free *in situ* proximity ligation and nucleosome-resolution chromosome walks in human cells": Custom code: submit_cis_fraglen_hist_by_bed_v2_20250312.sh

```
#!/bin/bash
#SBATCH --job-name=frag_lens
#SBATCH --output=frag_lens_%j.out
#SBATCH --error=frag_lens_%j.err

# Activate any needed environment or modules, if required
# e.g. module load python/3.10

# Define paths/args:
INPUT_GZ="/rugpfs/fs0/risc_lab/scratch/jsoroczyns/CAD/CADwalk/CADwalk_E1_GM12878_2x250/results_20250207_0034_nonexpanded/ABC_walk_processing_20250209/20250214_Bfilt_flattened_parse2_UU_direct_ABC_complete_consequent12/filtzerolenB_flip_sort_dedup_split_20250214/merged_all/20250312_cis_intersect_ChromHMM_v2/merged_all_flipped_sorted.cis.ChromHMM_v2_fraglen_seps.tsv.gz"

SCRIPT="/rugpfs/fs0/risc_lab/scratch/jsoroczyns/CAD/CAD-C/scripts/2025_03_12/parse_frag_lens_by_bed_v2_20250312.py"

# Adjust the output CSV and bed_map file paths as you like:
HIST_OUT="/rugpfs/fs0/risc_lab/scratch/jsoroczyns/CAD/CADwalk/CADwalk_E1_GM12878_2x250/results_20250207_0034_nonexpanded/ABC_walk_processing_20250209/20250214_Bfilt_flattened_parse2_UU_direct_ABC_complete_consequent12/filtzerolenB_flip_sort_dedup_split_20250214/merged_all/20250312_cis_intersect_ChromHMM_v2/histogram_output.csv"
BED_MAP_OUT="/rugpfs/fs0/risc_lab/scratch/jsoroczyns/CAD/CADwalk/CADwalk_E1_GM12878_2x250/results_20250207_0034_nonexpanded/ABC_walk_processing_20250209/20250214_Bfilt_flattened_parse2_UU_direct_ABC_complete_consequent12/filtzerolenB_flip_sort_dedup_split_20250214/merged_all/20250312_cis_intersect_ChromHMM_v2/bed_map_info.txt"

# Run the pipeline:
gunzip -c "$INPUT_GZ" | \
python "$SCRIPT" \
  --hist-out "$HIST_OUT" \
  --bed-map-out "$BED_MAP_OUT" \
  --retain-comments \
  --chunksize 5000000
```
