## Supplementary material for "CAD-C: An engineered nuclease enables repair-free *in situ* proximity ligation and nucleosome-resolution chromosome walks in human cells": Custom code: submit_stewam_PBCw_v7_test1_20251017.sh

```
#!/bin/bash
#SBATCH --job-name=cadwalk_v6
#SBATCH --output=%x_%j.out
#SBATCH --error=%x_%j.err
#SBATCH --cpus-per-task=1

# Paths
OUTDIR=/rugpfs/fs0/risc_lab/scratch/jsoroczyns/CAD/CADwalk/CADwalk_E1_GM12878/bwamem2_out_20250408/local_parse2_out/analyses_20251017/
P1=/rugpfs/fs0/risc_lab/scratch/jsoroczyns/CAD/CADwalk/CADwalk_E1_GM12878/bwamem2_out_20250408/local_parse2_out/bc1001_parse2_20250408.pairs.gz
P2=/rugpfs/fs0/risc_lab/scratch/jsoroczyns/CAD/CADwalk/CADwalk_E1_GM12878/bwamem2_out_20250408/local_parse2_out/bc1002_parse2_20250408.pairs.gz
SCRIPT=/rugpfs/fs0/risc_lab/scratch/jsoroczyns/CAD/CAD-C/scripts/2025_10_17/streaming_PBCADwalk_analyses_v7_20251017.py

set -euo pipefail
export OMP_NUM_THREADS=${SLURM_CPUS_PER_TASK:-1}
export OPENBLAS_NUM_THREADS=${SLURM_CPUS_PER_TASK:-1}
export MKL_NUM_THREADS=${SLURM_CPUS_PER_TASK:-1}
export NUMEXPR_NUM_THREADS=${SLURM_CPUS_PER_TASK:-1}

echo "[INFO] Host: $(hostname)"
echo "[INFO] When: $(date)"
echo "[INFO] Script: $SCRIPT"
echo "[INFO] Outdir: $OUTDIR"
echo

python "$SCRIPT" -o "$OUTDIR" \
  --pairs "$P1" "$P2" \
  --label bc1001 bc1002 \
  --small-hist-bin 5 --small-hist-max 2000 \
  --nonadj-reservoir 2000000 \
  --frag-ecdf-max 1000

echo "[INFO] Done at $(date)"
```
