## Supplementary material for "CAD-C: An engineered nuclease enables repair-free *in situ* proximity ligation and nucleosome-resolution chromosome walks in human cells": Custom code: v8p1_pack_plotFLDs_v1_20251020.py

```
#!/usr/bin/env python3
# -*- coding: utf-8 -*-
"""
fraglen_qc_runlength_v81.py
---------------------------
Purpose
  QC the fragment length distribution by *run length* using a CADpack v8.1 NPZ.

What it produces (exactly as shown in prior figures)
  1) Individual histograms (one figure per stratum): run_L = 1, 2, 3, ≥4
     • Range 0–250 bp, unfilled binned histogram (bin width = 3 bp)
     • x-ticks every 25 bp
     • PDF font type = 42 (AI editable)
     • File names: fraglen_hist_runL_1/2/3/ge4.(png|pdf)

  2) Combined overlay plots (single figure each)
     • Unfilled step histograms for strata + a dark-grey “ALL” series
     • Palette: mako-inspired categorical (manual hex), ALL in dark grey
     • (A) y-axis = raw counts → fraglen_overlay_counts.(png|pdf)
     • (B) y-axis = counts per million (CPM) → fraglen_overlay_cpm.(png|pdf)
     • Legend includes the fragment **counts (N)** actually used (0–250 bp)

Inputs
  • v8.1 NPZ with keys:
      - run_L (per run), frag_run_idx (per fragment->run)
      - frag_len_bp  (preferred), or fallback: frag_ref_len_choice
    (All other fields are not required for this QC.)

Defaults & Notes
  • All runs are included (cis-confined and trans-exploring).
  • Only fragments with length > 0 and within [0, 250] bp are plotted.
  • Bin width, xmax, and xtick step are configurable via CLI flags.

Usage
  python fraglen_qc_runlength_v81.py \
    --npz /path/to/cadpack_bc1001_bc1002_v8_1_mterm_14352915.npz \
    --outdir ./fraglen_qc \
    --binw 3 --xmax 250 --xtick 25

Outputs
  ./fraglen_qc/
    ├─ fraglen_hist_runL_1.(png|pdf)
    ├─ fraglen_hist_runL_2.(png|pdf)
    ├─ fraglen_hist_runL_3.(png|pdf)
    ├─ fraglen_hist_runL_ge4.(png|pdf)
    ├─ fraglen_overlay_counts.(png|pdf)
    ├─ fraglen_overlay_cpm.(png|pdf)
    └─ fraglen_counts_summary.tsv    # N per stratum and within 0–250
"""

import os
import argparse
import numpy as np
import matplotlib
matplotlib.use("Agg")  # non-interactive backend
import matplotlib.pyplot as plt
import pandas as pd


def parse_args():
    ap = argparse.ArgumentParser(formatter_class=argparse.ArgumentDefaultsHelpFormatter)
    ap.add_argument("--npz", required=True, help="Path to CADpack v8.1 NPZ")
    ap.add_argument("--outdir", default=".", help="Directory for outputs")
    ap.add_argument("--binw", type=int, default=3, help="Bin width (bp)")
    ap.add_argument("--xmax", type=int, default=250, help="Max x (bp) for plotting")
    ap.add_argument("--xtick", type=int, default=25, help="x-axis tick interval (bp)")
    return ap.parse_args()


def main():
    args = parse_args()
    os.makedirs(args.outdir, exist_ok=True)

    # Illustrator-friendly fonts
    plt.rcParams["pdf.fonttype"] = 42
    plt.rcParams["ps.fonttype"]  = 42

    Z = np.load(args.npz, allow_pickle=False, mmap_mode=None)

    # Required arrays
    run_L  = Z["run_L"].astype(np.int64)              # per-run
    fr_run = Z["frag_run_idx"].astype(np.int64)       # per-fragment -> run id

    # Preferred length; fallback if unified missing
    if "frag_len_bp" in Z.files:
        frag_len = Z["frag_len_bp"].astype(np.int64)
    else:
        frag_len = Z["frag_ref_len_choice"].astype(np.int64)

    # Map fragments to run length
    frag_runlen = run_L[fr_run]

    # Keep valid lengths and in-range for plotting
    valid = frag_len > 0
    frag_len = frag_len[valid]
    frag_runlen = frag_runlen[valid]

    in_range = (frag_len >= 0) & (frag_len <= args.xmax)
    frag_len = frag_len[in_range]
    frag_runlen = frag_runlen[in_range]

    # Strata masks
    strata = {
        "run_L=1":  (frag_runlen == 1),
        "run_L=2":  (frag_runlen == 2),
        "run_L=3":  (frag_runlen == 3),
        "run_L≥4":  (frag_runlen >= 4),
    }

    # Bins
    bins = np.arange(0, args.xmax + args.binw, args.binw)

    # Summary counts
    rows = []
    for lab, m in strata.items():
        lens = frag_len[m]
        rows.append({"stratum_run_L": lab,
                     "N_frag_0_{}bp".format(args.xmax): int(lens.size)})
    # add ALL
    rows.append({"stratum_run_L": "ALL",
                 "N_frag_0_{}bp".format(args.xmax): int(frag_len.size)})
    summary_df = pd.DataFrame(rows)
    summary_tsv = os.path.join(args.outdir, "fraglen_counts_summary.tsv")
    summary_df.to_csv(summary_tsv, sep="\t", index=False)

    # -------- (1) Individual histograms, one per stratum (unfilled bins) --------
    for lab, m in strata.items():
        lens = frag_len[m]
        fig = plt.figure(figsize=(7.6, 4.8))
        ax = fig.add_subplot(111)
        # Unfilled binned histogram — draw as step to match prior behavior
        counts, edges = np.histogram(lens, bins=bins)
        ax.step(edges[:-1], counts, where="post", color="#2e6e8e", linewidth=1.8)
        ax.set_xlim(0, args.xmax)
        ax.set_xlabel("Fragment length (bp)")
        ax.set_ylabel("Count")
        ax.set_title(f"Fragment length histogram — {lab}\n(bin = {args.binw} bp, range 0–{args.xmax} bp)")
        ax.set_xticks(np.arange(0, args.xmax+1, args.xtick))
        # Remove gridlines on the plot area (per request)
        ax.grid(False)
        fig.tight_layout()
        tag = "ge4" if lab.endswith("≥4") else lab.split("=")[-1]
        png = os.path.join(args.outdir, f"fraglen_hist_runL_{tag}.png")
        pdf = os.path.join(args.outdir, f"fraglen_hist_runL_{tag}.pdf")
        fig.savefig(png, dpi=160)
        fig.savefig(pdf)
        plt.close(fig)

    # -------- (2) Combined overlay: strata + ALL (counts) --------
    # Palette (mako-inspired categorical) + ALL = dark grey
    palette = {
        "run_L=1":  "#0b1026",
        "run_L=2":  "#1d3b6a",
        "run_L=3":  "#2e6e8e",
        "run_L≥4":  "#66b7c1",
        "ALL":      "#444444",
    }

    def compute_counts_by(mask):
        return np.histogram(frag_len[mask], bins=bins)[0]

    series = {}
    Ns = {}
    series["ALL"] = np.histogram(frag_len, bins=bins)[0]
    Ns["ALL"] = int(frag_len.size)
    for lab, m in strata.items():
        series[lab] = compute_counts_by(m)
        Ns[lab] = int(frag_len[m].size)

    def plot_overlay(counts_dict, Ns_dict, ylabel, out_png, out_pdf):
        fig = plt.figure(figsize=(8.8, 5.2))
        ax = fig.add_subplot(111)
        # ALL first (dark grey)
        ax.step(bins[:-1], counts_dict["ALL"], where="post",
                color=palette["ALL"], linewidth=2.0,
                label=f"ALL (N={Ns_dict['ALL']:,})")
        # Strata overlays
        for lab in ["run_L=1","run_L=2","run_L=3","run_L≥4"]:
            ax.step(bins[:-1], counts_dict[lab], where="post",
                    color=palette[lab], linewidth=1.8,
                    label=f"{lab} (N={Ns_dict[lab]:,})")
        ax.set_xlim(0, args.xmax)
        ax.set_xlabel("Fragment length (bp)")
        ax.set_ylabel(ylabel)
        ax.set_title(f"Fragment length distribution by run length (bin = {args.binw} bp, range 0–{args.xmax} bp)")
        # No gridlines
        ax.grid(False)
        ax.set_xticks(np.arange(0, args.xmax+1, args.xtick))
        ax.legend(frameon=False, ncol=2)
        fig.tight_layout()
        fig.savefig(out_png, dpi=160)
        fig.savefig(out_pdf)
        plt.close(fig)

    # (A) raw counts
    out_counts_png = os.path.join(args.outdir, "fraglen_overlay_counts.png")
    out_counts_pdf = os.path.join(args.outdir, "fraglen_overlay_counts.pdf")
    plot_overlay(series, Ns, "Count", out_counts_png, out_counts_pdf)

    # (B) counts per million (CPM)
    series_cpm = {}
    for lab, cnts in series.items():
        N = Ns[lab]
        series_cpm[lab] = (cnts.astype(float) / N * 1_000_000.0) if N > 0 else np.zeros_like(cnts, float)

    out_cpm_png = os.path.join(args.outdir, "fraglen_overlay_cpm.png")
    out_cpm_pdf = os.path.join(args.outdir, "fraglen_overlay_cpm.pdf")
    plot_overlay(series_cpm, Ns, "Count per million fragments", out_cpm_png, out_cpm_pdf)

    print(f"[OK] Wrote individual histograms and overlays to: {args.outdir}")
    print(f"[OK] Summary counts: {os.path.basename(summary_tsv)}")


if __name__ == "__main__":
    main()
```
