## Supplementary material for "CAD-C: An engineered nuclease enables repair-free *in situ* proximity ligation and nucleosome-resolution chromosome walks in human cells": Custom code: vis_ABC_position_merged_FLD_by_bed_chromHMM_colorscheme_v5_ridge_20250312.py

```
#!/usr/bin/env python3

import pandas as pd
import matplotlib.pyplot as plt
import datetime
import numpy as np
import os
import argparse
import matplotlib

# Ensure proper PDF font for Adobe Illustrator compatibility
matplotlib.rcParams['pdf.fonttype'] = 42
matplotlib.rcParams['ps.fonttype'] = 42

# ✅ Argument Parser for User-Controlled Parameters
parser = argparse.ArgumentParser(description="Generate filled histograms and ridge plots for fragment length distributions.")
parser.add_argument("--input", type=str, required=True, help="Path to the input CSV file.")
parser.add_argument("--scale", type=float, default=4.0, help="Scaling factor for ridge plot height (default: 4.0).")
parser.add_argument("--offset", type=float, default=0.03, help="Offset between ridge plots (default: 0.03).")

args = parser.parse_args()

# ✅ Load the CSV file
df = pd.read_csv(args.input)

# ✅ Ensure expected columns exist
required_columns = {"position", "bed_label", "length", "count"}
if not required_columns.issubset(df.columns):
    raise ValueError(f"Missing expected columns in CSV. Found: {df.columns}")

# ✅ Define corrected mapping from full ChromHMM file names to short labels
bed_label_remap = {
    "ChromHMM_ActiveEnhancer_merged.bed": "ActiveEnhancer",
    "ChromHMM_WeakEnhancer_merged.bed": "WeakEnhancer",
    "ChromHMM_PromoterActive_merged.bed": "PromoterActive",
    "ChromHMM_TxStrong_merged.bed": "TxStrong",
    "ChromHMM_TxWeak_merged.bed": "TxWeak",
    "ChromHMM_ReprPC.bed": "ReprPC",
    "ChromHMM_Het.bed": "Het",
    "ChromHMM_ZNF_Rpts.bed": "ZNF_Rpts",
    "ChromHMM_DNase.bed": "DNase",
    "ChromHMM_PromBiv.bed": "PromBiv",
    "ChromHMM_PromP.bed": "PromP"
}

# ✅ Helper function to extract short labels from file paths
def extract_bed_label(path):
    if pd.isna(path):
        return "NA"
    filename = os.path.basename(path)  # Extract filename
    return bed_label_remap.get(filename, "NA")  # Map to short label

# ✅ Apply label extraction
df["short_label"] = df["bed_label"].apply(extract_bed_label)

# ✅ Aggregate counts by (short_label, length)
df_summed = df.groupby(["short_label", "length"], as_index=False)["count"].sum()

# ✅ Restrict to lengths between 30 and 300 bp
df_summed = df_summed[(df_summed["length"] >= 30) & (df_summed["length"] <= 300)]

# ✅ Normalize counts per short_label
df_summed["normalized_count"] = df_summed.groupby("short_label")["count"].transform(lambda x: x / x.sum())
df_summed["normalized_count"] = df_summed["normalized_count"].fillna(0)  # Replace NaN with 0

# ✅ Define colors for labels
short_label_colors = {
    "ActiveEnhancer": (240/255, 180/255, 90/255),
    "DNase": (250/255, 230/255, 100/255),
    "Het": (90/255, 120/255, 180/255),
    "PromBiv": (180/255, 120/255, 150/255),
    "PromP": (180/255, 120/255, 150/255),
    "PromoterActive": (230/255, 60/255, 50/255),
    "ReprPC": (130/255, 130/255, 130/255),
    "TxStrong": (30/255, 140/255, 50/255),
    "TxWeak": (190/255, 200/255, 80/255),
    "WeakEnhancer": (250/255, 230/255, 100/255),
    "ZNF_Rpts": (90/255, 120/255, 180/255),
    "NA": (0.5, 0.5, 0.5)
}

# ✅ Generate unique timestamp
timestamp = datetime.datetime.now().strftime("%Y%m%d_%H%M")

# ✅ Function to create and save filled step plots
def plot_distribution(y_scale, output_file):
    plt.figure(figsize=(8, 5))
    all_labels = sorted(df_summed["short_label"].unique())

    for label in all_labels:
        subset = df_summed[df_summed["short_label"] == label].sort_values("length")

        if not subset.empty:
            color = short_label_colors.get(label, (0, 0, 0))  # Default to black if not found
            plt.fill_between(subset["length"], subset["normalized_count"], step="pre", color=color, alpha=0.5)

    plt.xlabel("Fragment length (bp) (30-300)")
    plt.ylabel("Normalized frequency")
    plt.title("Fragment Length Distribution by ChromHMM Merged States\n(Filled Hist, Summed A/B/C)")
    plt.legend(fontsize="x-small", loc="upper right")
    plt.xlim(30, 300)

    if y_scale == "log":
        plt.yscale("log")

    plt.savefig(output_file, bbox_inches="tight")
    plt.close()
    print(f"Saved plot: {output_file}")

# ✅ Save filled plots for Linear and Log10 y-axis
plot_distribution(y_scale="linear", output_file=f"fraglen_hist_filled_linear_{timestamp}.pdf")
plot_distribution(y_scale="log", output_file=f"fraglen_hist_filled_log10_{timestamp}.pdf")

# ✅ Function to generate Ridge Plot with customizable height and spacing
def ridge_plot(output_file, offset=args.offset, scale=args.scale):
    plt.figure(figsize=(8, 8))
    all_labels = sorted(df_summed["short_label"].unique(), reverse=True)  # Reverse for better ordering
    y_offset = 0

    for label in all_labels:
        subset = df_summed[df_summed["short_label"] == label].sort_values("length")

        if not subset.empty:
            color = short_label_colors.get(label, (0, 0, 0))
            
            # Scale height of FLDs
            plt.fill_between(
                subset["length"],
                y_offset + subset["normalized_count"] * scale,  # Taller FLDs
                y_offset,
                step="pre",
                color=color,
                alpha=0.6
            )
            
            # Labels
            plt.text(
                310,  # Adjust x-position
                y_offset + max(subset["normalized_count"]) * scale / 2,  # Center label
                label,
                fontsize=10,
                va="center"
            )

            y_offset += offset  # Adjust spacing between ridges

    plt.xlabel("Fragment length (bp) (30-300)")
    plt.ylabel("Stacked normalized counts")
    plt.title("Ridge Plot: Fragment Length Distribution by ChromHMM States")
    plt.xlim(30, 300)
    plt.yticks([])

    plt.savefig(output_file, bbox_inches="tight")
    plt.close()
    print(f"Saved ridge plot: {output_file}")

# ✅ Save Ridge Plot with user-defined height and spacing
ridge_plot(output_file=f"fraglen_ridgeplot_{timestamp}.pdf")
```
