## Supplementary material for "CAD-C: An engineered nuclease enables repair-free *in situ* proximity ligation and nucleosome-resolution chromosome walks in human cells": Custom code: ABC_calculatedistance.R

```
#!/usr/bin/env Rscript

# Load necessary libraries
library(dplyr)
library(tidyr)
library(ggplot2)

# Parse command-line arguments
args <- commandArgs(trailingOnly = TRUE)
if (length(args) < 2) {
  stop("Usage: Rscript compute_genomic_distances.R <input_tsv> <output_directory>")
}

# Read input and output parameters
input_file <- args[1]
output_dir <- args[2]

distance_data <- read.delim(input_file, header = TRUE, sep = "\t")

# Compute absolute genomic distances
distance <- distance_data %>%
  mutate(
    AB_distance = abs(B_start - A_start),
    BC_distance = abs(C_start - B_start),
    AC_distance = abs(C_start - A_start)
  )

# Save processed data
output_file <- file.path(output_dir, gsub(".tsv$", "_distances.tsv", basename(input_file)))
write.table(distance, output_file, sep = "\t", quote = FALSE, row.names = FALSE)

# Log10 transform the data
distance_long <- distance %>%
  pivot_longer(cols = c(AB_distance, BC_distance, AC_distance), 
               names_to = "Distance_Type", values_to = "Distance")

# Plot violin plot
a <- ggplot(distance_long, aes(x = Distance_Type, y = log10(Distance), fill = Distance_Type)) +
  geom_violin(alpha = 0.7) +
  geom_boxplot(width = 0.1, color = "black", alpha = 0.5) + 
  theme_minimal() +
  scale_y_continuous(name = "Log10 Distance (bp)") +
  labs(title = "Genomic Distance Violin Plot", x = "Distance Type") +
  theme(legend.position = "none")

# Save violin plot
ggsave(file.path(output_dir, "Genomic_Distance_Violin_Plot.pdf"), a)

# Frequency polygon plot
b <- ggplot(distance_long, aes(x = log10(Distance), fill = Distance_Type)) +
  geom_freqpoly(binwidth = 0.01) +
  theme_minimal() +
  facet_wrap(~Distance_Type, nrow=2)

# Save frequency plot
ggsave(file.path(output_dir, "GenomeWide-distance-log10-frequency.pdf"), b)

# Compute summary statistics
summary_stats <- distance_long %>%
  group_by(Distance_Type) %>%
  summarise(
    Mean = mean(Distance, na.rm = TRUE),
    Median = median(Distance, na.rm = TRUE),
    Mode = as.numeric(names(sort(table(Distance), decreasing = TRUE)[1])),
    SD = sd(Distance, na.rm = TRUE),
    Min = min(Distance, na.rm = TRUE),
    Max = max(Distance, na.rm = TRUE),
    Count = n()
  )

# Save summary statistics
ggsave(file.path(output_dir, "Distance_Summary_Stats.tsv"), summary_stats, sep = "\t", quote = FALSE, row.names = FALSE)

cat("Distance calculations and visualizations completed. Outputs saved to:", output_dir, "\n")
```
