## Supplementary material for "CAD-C: An engineered nuclease enables repair-free *in situ* proximity ligation and nucleosome-resolution chromosome walks in human cells": Custom code: Compartment_Density_byStart_20250609.r

```
#!/usr/bin/env Rscript

# Load necessary libraries
library(data.table)
library(dplyr)
library(tidyr)
library(ggplot2)
library(scales)

# Parse command-line arguments
args <- commandArgs(trailingOnly = TRUE)
if (length(args) < 2) {
  stop("Usage: Rscript compute_genomic_distances.R <input_tsv> <output_directory>")
}

# Read input and output parameters
input_file <- args[1]
output_dir <- args[2]
file_base <- sub("\\.tsv$", "", basename(input_file))

# Import .tsv as dataframe using fread (skip malformed header lines)
data <- fread(input_file, sep = "\t", skip = 2, data.table = FALSE)

# Assign column names
colnames(data) <- c("readID", "A_chrom", "A_start", "A_end", "A_strand",
                    "B_chrom", "B_start", "B_end", "B_strand",
                    "C_chrom", "C_start", "C_end", "C_strand",
                    "A_intersect", "B_intersect", "C_intersect",
                    "AB_distance", "BC_distance", "AC_distance", "Compartment_Class")

# Define compartment file names
compartment_A <- "cad_GM12878_A_compartments_mergedBins.bed"
compartment_B <- "cad_GM12878_B_compartments_mergedBins.bed"

# Re-assign Compartment_Class based on A_intersect
data <- data %>%
  mutate(
    Compartment_Class = case_when(
      is.na(A_intersect) ~ NA_character_,
      A_intersect == compartment_A ~ "A Start",
      A_intersect == compartment_B ~ "B Start",
      TRUE ~ "Other Start"
    )
  )

data_filtered <- data %>% filter(Compartment_Class %in% c("A Start", "B Start"))

# Summary statistics
summary_stats <- data %>%
  group_by(Compartment_Class) %>%
  summarise(
    Mean = mean(AC_distance, na.rm = TRUE),
    Median = median(AC_distance, na.rm = TRUE),
    SD = sd(AC_distance, na.rm = TRUE),
    Min = min(AC_distance, na.rm = TRUE),
    Max = max(AC_distance, na.rm = TRUE),
    Count = n()
  )
write.table(summary_stats, file.path(output_dir, paste0(file_base, "_Distance_Summary_Stats.tsv")),
            sep = "\t", quote = FALSE, row.names = FALSE)


# Density Plots
for (dist in c("AC", "AB", "BC")) {
  dist_col <- paste0(dist, "_distance")
  p <- ggplot(data_filtered, aes(x = log10(.data[[dist_col]]), fill = Compartment_Class)) +
    geom_density(alpha = 0.5) +
    theme_minimal() +
    scale_x_continuous(name = paste("Log10", dist, "Distance (bp)")) +
    scale_y_continuous(name = "Density") +
    labs(title = paste(dist, "Distance Density (Start A vs B) -", file_base), fill = "Start Compartment") +
    theme(legend.position = "right")
  ggsave(file.path(output_dir, paste0(file_base, "_", dist, "_Distance_Density_Compartment.pdf")), p)
}

# Bin definitions and plots for AC, AB, and BC
bin_labels <- c("Bin 1: <10kb", "Bin 2: >10kb, <5.5Mbp", "Bin 3: >5.5Mb, < 1Gbp")
bin_ranges <- list(c(1.5, 4), c(4, 6.5), c(6.5, 9.5))

for (dist in c("AC", "AB", "BC")) {
  dist_col <- paste0(dist, "_distance")
  bin_col <- paste0(dist, "_bin")
  
  data_filtered <- data_filtered %>%
    mutate(!!bin_col := case_when(
      log10(.data[[dist_col]]) >= bin_ranges[[1]][1] & log10(.data[[dist_col]]) < bin_ranges[[1]][2] ~ bin_labels[1],
      log10(.data[[dist_col]]) >= bin_ranges[[2]][1] & log10(.data[[dist_col]]) < bin_ranges[[2]][2] ~ bin_labels[2],
      log10(.data[[dist_col]]) >= bin_ranges[[3]][1] & log10(.data[[dist_col]]) < bin_ranges[[3]][2] ~ bin_labels[3]
    ))

  density_df <- data_filtered %>%
    filter(!is.na(.data[[bin_col]])) %>%
    group_by(Compartment_Class, .data[[bin_col]]) %>%
    summarise(count = n(), .groups = "drop") %>%
    group_by(Compartment_Class) %>%
    mutate(density = count / sum(count))

  p <- ggplot(density_df, aes(x = Compartment_Class, y = density, fill = .data[[bin_col]])) +
    geom_bar(stat = "identity", position = "fill", width = 0.4) + 
    theme_minimal() +
    scale_y_continuous(name = "Normalized Density") +
    scale_fill_manual(values = c("sienna2", "hotpink1", "hotpink3")) +
    labs(title = paste(dist, "Span Proportion by Bin")) +
    theme(legend.position = "right", text = element_text(size = 14)) +
    geom_text(aes(label = scales::percent(density, accuracy = 2)), 
              position = position_fill(vjust = 0.5), size = 5, color = "black")

  ggsave(file.path(output_dir, paste0(file_base, "_", dist, "_Span_Proportion_byBin_Compartment.pdf")), p)

  write.table(density_df, file.path(output_dir, paste0(file_base, "_", dist, "_compartment_density.tsv")), sep = "\t", quote = FALSE, row.names = FALSE)
}

# ------------------- AC, AB, BC DENSITY HISTOGRAMS -------------------
distance_types <- c("AC_distance", "AB_distance", "BC_distance")

for (dist in distance_types) {
  data_filtered <- data %>% filter(!is.na(.data[[dist]]))

  compartment_counts <- data_filtered %>%
    group_by(Compartment_Class) %>%
    summarise(n = n()) %>%
    mutate(label = paste0(Compartment_Class, "\nn = ", format(n, big.mark=",")),
           y_pos = seq(0.85, 0.65, length.out = n()) * 
             max(density(log10(data_filtered[[dist]]))$y, na.rm = TRUE))

  log_breaks <- seq(0, 10, by = 1)
  linear_breaks <- 10^log_breaks
  linear_labels <- label_number(scale_cut = cut_short_scale())(linear_breaks)

  p <- ggplot(data_filtered, aes(x = log10(.data[[dist]]), color = Compartment_Class, fill = Compartment_Class)) +
    geom_histogram(aes(y = ..density..), alpha = 0.5, position = "identity", bins = 100) +
    geom_density(alpha = .2) +
    theme_minimal() +
    geom_vline(xintercept = c(1.5, 4, 6.5, 9), linetype = 'dashed', color = 'black', size = 0.5) +
    labs(title = paste("Cis contacts chr1, Span of Walks (", dist, ") by Compartment"),
         x = paste("log10", dist, "(bp)"),
         y = "Density") +
    scale_x_continuous(
      breaks = log_breaks, 
      labels = log_breaks,
      sec.axis = sec_axis(~ 10^.x, name = paste(dist, "(bp)"), breaks = linear_breaks, labels = linear_labels)
    ) +
    geom_text(data = compartment_counts, 
              aes(x = max(log10(data_filtered[[dist]])) - 0.5, 
                  y = y_pos, 
                  label = label), 
              hjust = 0.8, inherit.aes = FALSE, size = 4)

  ggsave(file.path(output_dir, paste0("Binned_Distance_Density_Histogram_", dist, ".pdf")), p, width = 8, height = 5)
}  

# ------------------- AC, AB, BC DENSITY HISTOGRAMS -------------------
distance_types <- c("AC_distance", "AB_distance", "BC_distance")

for (dist in distance_types) {
  data_filtered <- data %>%
    filter(!is.na(.data[[dist]]), !is.na(Afrag))  # <- exclude NA starting fragments

  compartment_counts <- data_filtered %>%
    group_by(Compartment_Class) %>%
    summarise(n = n()) %>%
    mutate(label = paste0(Compartment_Class, "\nn = ", format(n, big.mark=",")),
           y_pos = seq(0.85, 0.65, length.out = n()) * 
             max(density(log10(data_filtered[[dist]]))$y, na.rm = TRUE))

  log_breaks <- seq(0, 10, by = 1)
  linear_breaks <- 10^log_breaks
  linear_labels <- label_number(scale_cut = cut_short_scale())(linear_breaks)

  p <- ggplot(data_filtered, aes(x = log10(.data[[dist]]), color = Compartment_Class, fill = Compartment_Class)) +
    geom_histogram(aes(y = ..density..), alpha = 0.5, position = "identity", bins = 100) +
    geom_density(alpha = .2) +
    theme_minimal() +
    geom_vline(xintercept = c(1.5, 4, 6.5, 9), linetype = 'dashed', color = 'black', size = 0.5) +
    labs(title = paste("Cis contacts chr1, Span of Walks (", dist, ") by Compartment"),
         x = paste("log10", dist, "(bp)"),
         y = "Density") +
    scale_x_continuous(
      breaks = log_breaks, 
      labels = log_breaks,
      sec.axis = sec_axis(~ 10^.x, name = paste(dist, "(bp)"), breaks = linear_breaks, labels = linear_labels)
    ) +
    geom_text(data = compartment_counts, 
              aes(x = max(log10(data_filtered[[dist]])) - 0.5, 
                  y = y_pos, 
                  label = label), 
              hjust = 0.8, inherit.aes = FALSE, size = 4)

  ggsave(file.path(output_dir, paste0("Binned_NoNA_Distance_Density_Histogram_", dist, ".pdf")), p, width = 8, height = 5)
}

cat("Distance calculations and visualizations completed for:", file_base, "\nOutputs saved to:", output_dir, "\n")
```
