## Supplementary material for "CAD-C: An engineered nuclease enables repair-free *in situ* proximity ligation and nucleosome-resolution chromosome walks in human cells": Custom code: readme.txt

Proportion of walks starting in A or B compartment, defined by A_intersect annotation. 

From the cis ABC flttened TSV, use ABC_calculatedistance.R to calculate the absolute distance btwn alpha and gamma — creates extra columns AB_distance", "BC_distance", "AC_distance"

Plotting with Compartment_Density_byStart_20250609.r

Step 1 vs Step 2 Density Hexbin We plotted cis-chromosomal α–β–γ walk step-lengths AB_distance (step 1; x-axis) versus BC_distance (step 2; y-axis) using ggplot2::geom_hex(bins = 100). Distances are shown in base pairs (bp). For walk-type–stratified plots, rows were filtered to retain complete cases (non-missing AB_distance, BC_distance, and AC_distance) and to exclude walks with undefined compartment labels (walk_type containing “NA”), where walk_type was defined from A/B compartment assignments of fragments A, B, and C. To make hexbin intensities comparable across panels, each walk contributed equal weight w = 1/N so that the total weight sums to 1 within each plotted panel (all-walk plot: N = nrow(data); per-walk-type plots: N = nrow(walk_data)). Both axes were displayed on log10 scales (scale_x_log10 and scale_y_log10). Hexbin fill used scale_fill_viridis_c(option = “plasma”, trans = “log10”), applying a log10 color transform to the normalized per-bin weights. 
