## Supplementary material for "CAD-C: An engineered nuclease enables repair-free *in situ* proximity ligation and nucleosome-resolution chromosome walks in human cells": Custom code: data provenance note.rtf

Density hexbin of AB distance vs BC distance (step 1 vs step 2) of Cis chromosomal walks. File path plotting from: /lustre/fs4/risc_lab/scratch/landerson01/20250205_CAD_2x250bp_GM_analysis/Compartment/Cis/merged_all_flipped_sorted.cis.compt_annotated.tsv.gz_distances_compartment.tsv.gz

Figure 5G generated with this section of code
/ru-auth/local/home/landerson01/scripts/
#!/usr/bin/env Rscript

### ----------------- LIBRARIES -----------------
suppressPackageStartupMessages({
  library(data.table)
  library(dplyr)
  library(ggplot2)
  library(viridis)
  library(stringr)
  library(tidyr)  
})

### ----------------- INPUT ARGUMENTS -----------------
args <- commandArgs(trailingOnly = TRUE)
if (length(args) < 3) {
  stop("Usage: Rscript obs_exp_hexbin_runner.R <observed.tsv.gz> <expected.tsv.gz> <output_dir>")
}
obs_file <- args[1]
exp_file <- args[2]
output_dir <- args[3]
dir.create(output_dir, showWarnings = FALSE, recursive = TRUE)

### ----------------- GLOBAL SETTINGS -----------------
bins <- 100
x_limits <- c(1e2, 1e8)
y_limits <- c(1e2, 1e8)

### ----------------- FUNCTION: COMPUTE HEXBIN -----------------
compute_hexbin <- function(data, label) {
  before_n <- nrow(data)
  data <- data %>% filter(AB_distance > 0, BC_distance > 0)
  after_n <- nrow(data)
  message("Filtered ", before_n - after_n, " rows with non-positive distances (", label, ")")

  p <- ggplot(data, aes(x = AB_distance, y = BC_distance, weight = 1 / nrow(data))) +
    geom_hex(bins = bins) +
    scale_x_log10(limits = x_limits) +
    scale_y_log10(limits = y_limits)
