## Supplementary material for "CAD-C: An engineered nuclease enables repair-free *in situ* proximity ligation and nucleosome-resolution chromosome walks in human cells": Custom code: step1vsstep2_obsvsexp_hexbins_square.r

```
/ru-auth/local/home/landerson01/scripts/
#!/usr/bin/env Rscript

# ----------------- LIBRARIES -----------------
suppressPackageStartupMessages({
  library(data.table)
  library(dplyr)
  library(ggplot2)
  library(viridis)
  library(stringr)
  library(tidyr)  
})

# ----------------- INPUT ARGUMENTS -----------------
args <- commandArgs(trailingOnly = TRUE)
if (length(args) < 3) {
  stop("Usage: Rscript obs_exp_hexbin_runner.R <observed.tsv.gz> <expected.tsv.gz> <output_dir>")
}
obs_file <- args[1]
exp_file <- args[2]
output_dir <- args[3]
dir.create(output_dir, showWarnings = FALSE, recursive = TRUE)

# ----------------- GLOBAL SETTINGS -----------------
bins <- 100
x_limits <- c(1e2, 1e8)
y_limits <- c(1e2, 1e8)

# ----------------- FUNCTION: COMPUTE HEXBIN -----------------
compute_hexbin <- function(data, label) {
  before_n <- nrow(data)
  data <- data %>% filter(AB_distance > 0, BC_distance > 0)
  after_n <- nrow(data)
  message("Filtered ", before_n - after_n, " rows with non-positive distances (", label, ")")

  p <- ggplot(data, aes(x = AB_distance, y = BC_distance, weight = 1 / nrow(data))) +
    geom_hex(bins = bins) +
    scale_x_log10(limits = x_limits) +
    scale_y_log10(limits = y_limits)

  hex_data <- ggplot_build(p)$data[[1]] %>%
    select(x, y, count, density) %>%
    mutate(
      AB_distance = 10^x,  # back-transform bin center
      BC_distance = 10^y
    )

  # Export hexbin data as TSV
  hex_file <- file.path(output_dir, paste0(label, "_hexbin_data.tsv"))
  fwrite(hex_data, hex_file, sep = "\t")
  message("Saved hexbin table to: ", hex_file)

  return(hex_data)
}

# ----------------- LOAD AND PROCESS OBSERVED -----------------
message("Reading observed (real) data...")
obs_data <- fread(obs_file)
obs_label <- str_remove(tools::file_path_sans_ext(basename(obs_file)), "\\.tsv\\.gz$|\\.tsv$")
obs_hex <- compute_hexbin(obs_data, obs_label)

# ----------------- LOAD AND PROCESS EXPECTED -----------------
message("Reading expected (synthetic) data...")
exp_data <- fread(exp_file)
exp_label <- str_remove(tools::file_path_sans_ext(basename(exp_file)), "\\.tsv\\.gz$|\\.tsv$")
exp_hex <- compute_hexbin(exp_data, exp_label)

# ----------------- MERGE AND CALCULATE OBS/EXP -----------------
merged <- full_join(obs_hex, exp_hex, by = c("x", "y"), suffix = c("_obs", "_exp")) %>%
  replace_na(list(density_obs = 0, density_exp = 0)) %>%
  mutate(
    obs_exp_ratio = (density_obs + 1e-6) / (density_exp + 1e-6),
    log2_obs_exp = log2(obs_exp_ratio)
  )

# ----------------- EXPORT MERGED DATA -----------------
merged_file <- file.path(output_dir, "merged_obs_exp_hexbin.tsv")
fwrite(merged, merged_file, sep = "\t")
message("Saved merged obs/exp table to: ", merged_file)

# ----------------- PLOT OBS/EXP -----------------
message("Plotting log2(Obs/Exp) heatmap...")

plot <- ggplot(merged, aes(x = AB_distance_obs, y = BC_distance_obs, fill = log2_obs_exp)) +
  geom_tile() +
  scale_x_log10(limits = x_limits) +
  scale_y_log10(limits = y_limits) +
  scale_fill_gradient2(
    low = "blue", mid = "white", high = "red", midpoint = 0,
    limits = c(-2, 2), oob = scales::squish
  ) +
  labs(
    title = "Log2(Observed / Expected) AB vs BC",
    x = "AB Distance (bp)",
    y = "BC Distance (bp)",
    fill = "log2(Obs/Exp)"
  ) +
  theme_minimal(base_size = 12)

ggsave(file.path(output_dir, "obs_exp_log2_heatmap.pdf"), plot, width = 6, height = 6, device = cairo_pdf)

message("Done! Output saved to ", output_dir)
```
