## Supplementary material for "CAD-C: An engineered nuclease enables repair-free *in situ* proximity ligation and nucleosome-resolution chromosome walks in human cells": Custom code: data provenance note.rtf

Log₂ enrichment of ChromHMM states in in nonlinear (top) vs linear walks (bottom). Density of Alpha, Beta, Gamma fragments by ChromHMM annotation divided by all other walks position+annotation.
Path plotting from: /lustre/fs4/risc_lab/scratch/landerson01/20250205_CAD_2x250bp_GM_analysis/ChromHMM/Cis/merged_all_flipped_sorted.cis.ChromHMM_v2.tsv.gz_distances_compartment.tsv.gz


Output plots:
/ru-auth/local/home/landerson01/scratch/20250205_CAD_2x250bp_GM_analysis/ChromHMM/Cis/Fragment_Distribution_20250528

Note that the figure was edited in illustrator, raw output pdf is /ru-auth/local/home/landerson01/scratch/20250205_CAD_2x250bp_GM_analysis/ChromHMM/Cis/Fragment_Distribution_20250528/diagonal_over_offdiag_delta_log2_barplot.pdf

ChromHMM Annotation and Per-Fragment Enrichment Each fragment in the cis-chromosomal two-start α-β-γ data was annotated by GM12878 imputed ChromHMM state. See Methods section “Fragment‑lengths, midpoints, and pairwise separations appended to ChromHMM‑annotated ABC walks.” Diagonal walks are defined by similarity in magnitude (|log10(AB) − log10(BC)| < 0.2), and all remaining walks were classified as other. For each position (α, β, or γ), fragments were counted by ChromHMM state and converted to percent frequencies. The full cis-chromosomal ChromHMM walk set was used as a baseline reference to calculate per state, per position log2 enrichment of diagonal and other walks. Diagonal log2 enrichment (over full data) was divided by “other” log2 enrichment (over full data) to visualize differences in per-position ChromHMM state. 
