## Supplementary material for "CAD-C: An engineered nuclease enables repair-free *in situ* proximity ligation and nucleosome-resolution chromosome walks in human cells": Custom code: Fragment_Distribution_20250528.r

```
#!/usr/bin/env Rscript

# ------------------- LIBRARIES -------------------
library(data.table)
library(dplyr)
library(ggplot2)
library(tidyr)
library(forcats)
library(readr)

# ------------------- CONFIG -------------------
ChromHMM_colors <- c(
  "DNase" = "#FFC107", "ActiveEnhancer_merged" = "#FF5722", "PromBiv" = "#4B0082",
  "PromP" = "#D291BC", "PromoterActive_merged" = "#F39C12", "ReprPC" = "gray",
  "ZNF_Rpts" = "#5DADE2", "Het" = "#8E44AD", "WeakEnhancer_merged" = "#FDFD96",
  "TxStrong_merged" = "#2ECC71", "TxWeak_merged" = "#90EE90"
)
my_levels <- c(
  "PromoterActive_merged", "TxWeak_merged", "ActiveEnhancer_merged",
  "PromBiv", "WeakEnhancer_merged", "DNase", "PromP",
  "ReprPC", "TxStrong_merged", "ZNF_Rpts", "Het"
)

output_dir <- "Fragment_Distribution_20250528"
dir.create(output_dir, showWarnings = FALSE)

file <- "/lustre/fs4/risc_lab/scratch/landerson01/20250205_CAD_2x250bp_GM_analysis/ChromHMM/Cis/merged_all_flipped_sorted.cis.ChromHMM_v2.tsv.gz_distances_compartment.tsv.gz"
data <- fread(file)

# ------------------- SUBSETS -------------------
message("Filtering walk types...")
full_data <- data
message("Total full_data rows: ", nrow(full_data))
linear_data <- full_data %>% filter(A_start < B_start & B_start < C_start)
message("Linear data rows: ", nrow(linear_data))
nonlinear_data <- anti_join(full_data, linear_data, by = "readID")
message("Nonlinear data rows: ", nrow(nonlinear_data))

diagonal_data <- nonlinear_data %>%
  mutate(diagonal_zone = abs(log10(AB_distance) - log10(BC_distance)) < 0.2) %>%
  filter(diagonal_zone)
message("Diagonal data rows: ", nrow(diagonal_data))

nonlinear_off_diagonal <- anti_join(nonlinear_data, diagonal_data, by = "readID")
message("Nonlinear off-diagonal data rows: ", nrow(nonlinear_off_diagonal))

# ------------------- HELPER FUNCTIONS -------------------
get_percent_table <- function(df, label) {
  df %>%
    select(A_intersect, B_intersect, C_intersect) %>%
    pivot_longer(cols = everything(), names_to = "Position", values_to = "Annotation") %>%
    filter(!is.na(Annotation)) %>%
    count(Position, Annotation) %>%
    group_by(Position) %>%
    mutate(percent = (n / sum(n)) * 100, group = label) %>%
    ungroup()
}

compute_log2_enrichment <- function(table, label) {
  table %>%
    left_join(reference_table, by = c("Position", "Annotation")) %>%
    mutate(
      log2_enrichment = ifelse(!is.na(ref_percent), log2(percent / ref_percent), NA),
      group = label
    )
}

plot_log2_enrichment <- function(df, file_base) {
  p <- ggplot(df, aes(x = fct_relevel(Annotation, my_levels), y = log2_enrichment, fill = Position)) +
    geom_col(position = "dodge") +
    geom_hline(yintercept = 0, linetype = "dashed") +
    labs(
      title = paste("log2(Obs / Exp) Enrichment:", unique(df$group)),
      x = "ChromHMM State",
      y = "log2(Obs / Exp)"
    ) +
    theme_minimal(base_size = 12) +
    theme(axis.text.x = element_text(angle = 45, hjust = 1))

  ggsave(file.path(output_dir, paste0(file_base, "_log2enrichment_plot.pdf")), plot = p, width = 8, height = 5)
  write_csv(df, file.path(output_dir, paste0(file_base, "_log2enrichment_data.csv")))
  message("Saved log2 enrichment plot and data for: ", file_base)
}

plot_log2_delta <- function(df1, df2, label1, label2, tag) {
  delta_data <- full_join(df1, df2, by = c("Position", "Annotation"), suffix = c(".x", ".y")) %>%
    mutate(delta_log2 = get("log2_enrichment.x", .) - get("log2_enrichment.y", .))

  # Barplot
  p1 <- ggplot(delta_data, aes(x = fct_relevel(Annotation, my_levels), y = delta_log2, fill = Position)) +
    geom_col(position = "dodge") +
    geom_hline(yintercept = 0, linetype = "dashed") +
    labs(
      title = paste("log2(Obs/Exp) Delta:", label1, "vs", label2),
      x = "ChromHMM State",
      y = paste("Delta log2(Obs/Exp):", label1, "-", label2)
    ) +
    theme_minimal() +
    theme(axis.text.x = element_text(angle = 45, hjust = 1))

  ggsave(file.path(output_dir, paste0(tag, "_delta_log2_barplot.pdf")), plot = p1, width = 8, height = 5)

  # Heatmap
  p2 <- ggplot(delta_data, aes(x = fct_relevel(Annotation, my_levels), y = Position, fill = delta_log2)) +
    geom_tile(color = "white") +
    scale_fill_distiller(palette = "RdYlBu", direction = -1, name = "Delta log2") +
    labs(
      title = paste("Heatmap of log2(Obs/Exp) Delta:", label1, "vs", label2),
      x = "ChromHMM State",
      y = "Fragment Position"
    ) +
    theme_minimal() +
    theme(axis.text.x = element_text(angle = 45, hjust = 1))

  ggsave(file.path(output_dir, paste0(tag, "_delta_log2_heatmap.pdf")), plot = p2, width = 10, height = 5)

  # Save data
  write_csv(delta_data, file.path(output_dir, paste0(tag, "_delta_log2_table.csv")))
  message("Saved delta log2 enrichment plots and table for: ", tag)
}

# ------------------- CALCULATIONS -------------------
reference_table_full <- get_percent_table(full_data, "Full") %>%
  select(Position, Annotation, ref_percent = percent)
reference_table <- reference_table_full

linear_table    <- get_percent_table(linear_data, "Linear")
nonlinear_table <- get_percent_table(nonlinear_data, "Nonlinear")
offdiag_table   <- get_percent_table(nonlinear_off_diagonal, "Nonlinear_OffDiagonal")
diagonal_table  <- get_percent_table(diagonal_data, "Diagonal")


log2_linear    <- compute_log2_enrichment(linear_table, "Linear")
log2_nonlinear <- compute_log2_enrichment(nonlinear_table, "Nonlinear")
log2_offdiag   <- compute_log2_enrichment(offdiag_table, "Nonlinear_OffDiagonal")
log2_diagonal  <- compute_log2_enrichment(diagonal_table, "Diagonal")

plot_log2_enrichment(log2_linear,    "log2enrichment_linear")
plot_log2_enrichment(log2_nonlinear, "log2enrichment_nonlinear")
plot_log2_enrichment(log2_offdiag,   "log2enrichment_offdiag")
plot_log2_enrichment(log2_diagonal,  "log2enrichment_diagonal")

# Log2 delta comparisons
plot_log2_delta(log2_nonlinear, log2_linear, "Nonlinear", "Linear", "nonlinear_vs_linear")
plot_log2_delta(log2_diagonal, log2_offdiag, "Diagonal", "Nonlinear_OffDiagonal", "diagonal_vs_offdiag")

# Export combined tables
bind_rows(log2_nonlinear, log2_linear) %>%
  write_csv(file.path(output_dir, "log2_linear_nonlinear_combined.csv"))

bind_rows(log2_offdiag, log2_diagonal) %>%
  write_csv(file.path(output_dir, "log2_offdiag_diagonal_combined.csv"))

# Plot stacked bar of walk types
walktype_counts <- tibble(
  Category = c("Linear", "Nonlinear (OffDiag)", "Nonlinear (Diagonal)"),
  Count = c(nrow(linear_data), nrow(nonlinear_off_diagonal), nrow(diagonal_data))
) %>%
  mutate(
    Group = ifelse(Category == "Linear", "Linear", "Nonlinear"),
    Category = factor(Category, levels = c("Linear", "Nonlinear (OffDiag)", "Nonlinear (Diagonal)"))
  )

walktype_counts <- walktype_counts %>%
  mutate(Proportion = Count / sum(Count))

p_stack <- ggplot(walktype_counts, aes(x = Group, y = Proportion, fill = Category)) +
  geom_bar(stat = "identity") +
  geom_text(aes(label = scales::percent(Proportion, accuracy = 0.1)),
            position = position_stack(vjust = 0.5), size = 3.5) +
  labs(
    title = "Distribution of Walk Types",
    x = "Walk Type",
    y = "Proportion"
  ) +
  theme_minimal(base_size = 12)

ggsave(file.path(output_dir, "walktype_stacked_barplot.pdf"), plot = p_stack, width = 5, height = 4)

# ------------------- Diagonal vs Other Analysis -------------------
message("Extracting diagonal vs other from full dataset")
full_diagonal <- full_data %>%
  mutate(diagonal_zone = abs(log10(AB_distance) - log10(BC_distance)) < 0.2) %>%
  filter(diagonal_zone)

other_walks <- anti_join(full_data, full_diagonal, by = "readID")

message("Full diagonal: ", nrow(full_diagonal))
message("Other walks: ", nrow(other_walks))

full_diagonal_table <- get_percent_table(full_diagonal, "Full_Diagonal")
other_walks_table   <- get_percent_table(other_walks, "Other")

log2_full_diagonal <- compute_log2_enrichment(full_diagonal_table, "Full_Diagonal")
log2_other_walks   <- compute_log2_enrichment(other_walks_table, "Other")


log2_reference <- reference_table_full %>%
  mutate(
    log2_enrichment = 0,
    group = "Full_Reference"
  )
plot_log2_delta(log2_full_diagonal, log2_reference, "Full_Diagonal", "Full_Reference", "full_diagonal_vs_full")
plot_log2_delta(log2_other_walks, log2_reference, "Other", "Full_Reference", "other_vs_full")
plot_log2_delta(log2_full_diagonal, log2_other_walks, "Full_Diagonal", "Other", "full_diagonal_vs_other")

bind_rows(log2_full_diagonal, log2_other_walks) %>%
  write_csv(file.path(output_dir, "log2_full_diagonal_vs_other_combined.csv"))

# Additional: Diagonal over Off-Diagonal
plot_log2_delta(log2_diagonal, log2_offdiag, "Diagonal", "Nonlinear_OffDiagonal", "diagonal_over_offdiag")

# ------------------- Diagonal vs Other Analysis -------------------
message("Extracting diagonal vs other from full dataset")
full_diagonal <- full_data %>%
  mutate(diagonal_zone = abs(log10(AB_distance) - log10(BC_distance)) < 0.2) %>%
  filter(diagonal_zone)

other_walks <- anti_join(full_data, full_diagonal, by = "readID")

message("Full diagonal: ", nrow(full_diagonal))
message("Other walks: ", nrow(other_walks))

full_diagonal_table <- get_percent_table(full_diagonal, "Full_Diagonal")
other_walks_table   <- get_percent_table(other_walks, "Other")

log2_full_diagonal <- compute_log2_enrichment(full_diagonal_table, "Full_Diagonal")
log2_other_walks   <- compute_log2_enrichment(other_walks_table, "Other")


log2_reference <- reference_table_full %>%
  mutate(
    log2_enrichment = 0,
    group = "Full_Reference"
  )
plot_log2_delta(log2_full_diagonal, log2_reference, "Full_Diagonal", "Full_Reference", "full_diagonal_vs_full")
plot_log2_delta(log2_other_walks, log2_reference, "Other", "Full_Reference", "other_vs_full")
plot_log2_delta(log2_full_diagonal, log2_other_walks, "Full_Diagonal", "Other", "full_diagonal_vs_other")

bind_rows(log2_full_diagonal, log2_other_walks) %>%
  write_csv(file.path(output_dir, "log2_full_diagonal_vs_other_combined.csv"))

# Additional: Diagonal over Off-Diagonal
plot_log2_delta(log2_diagonal, log2_offdiag, "Diagonal", "Nonlinear_OffDiagonal", "diagonal_over_offdiag")

# Additional: Full Diagonal over Other
# Instead of log2 difference, compute direct enrichment ratio
full_diagonal_over_other <- full_join(log2_full_diagonal, log2_other_walks, by = c("Position", "Annotation"), suffix = c(".diag", ".other")) %>%
  mutate(
    enrichment_ratio = percent.diag / percent.other,
    log2_enrichment_ratio = log2(enrichment_ratio)
  )

# Barplot of enrichment ratio
p_ratio_bar <- ggplot(full_diagonal_over_other, aes(x = fct_relevel(Annotation, my_levels), y = enrichment_ratio, fill = Position)) +
  geom_col(position = "dodge") +
  geom_hline(yintercept = 1, linetype = "dashed") +
  labs(
    title = "Enrichment Ratio: Full Diagonal / Other",
    x = "ChromHMM State",
    y = "Enrichment Ratio"
  ) +
  theme_minimal() +
  theme(axis.text.x = element_text(angle = 45, hjust = 1))

ggsave(file.path(output_dir, "full_diagonal_over_other_ratio_barplot.pdf"), plot = p_ratio_bar, width = 8, height = 5)

# Heatmap of log2 enrichment ratio
p_ratio_heat <- ggplot(full_diagonal_over_other, aes(x = fct_relevel(Annotation, my_levels), y = Position, fill = log2_enrichment_ratio)) +
  geom_tile(color = "white") +
  scale_fill_distiller(palette = "RdYlBu", direction = 1, name = "log2(Diag / Other)") +
  labs(
    title = "Heatmap of Enrichment Ratio: Full Diagonal / Other",
    x = "ChromHMM State",
    y = "Fragment Position"
  ) +
  theme_minimal() +
  theme(axis.text.x = element_text(angle = 45, hjust = 1))

ggsave(file.path(output_dir, "full_diagonal_over_other_ratio_heatmap.pdf"), plot = p_ratio_heat, width = 10, height = 5)

write_csv(full_diagonal_over_other, file.path(output_dir, "full_diagonal_over_other_ratio_table.csv"))
message("Saved log2 enrichment ratio heatmap for: full_diagonal_over_other")
ggsave(file.path(output_dir, "full_diagonal_over_other_log2ratio_barplot.pdf"), plot = ggplot(full_diagonal_over_other, aes(x = fct_relevel(Annotation, my_levels), y = log2_enrichment_ratio, fill = Position)) +
  geom_col(position = "dodge") +
  geom_hline(yintercept = 0, linetype = "dashed") +
  labs(
    title = "Log2 Enrichment Ratio: Full Diagonal / Other",
    x = "ChromHMM State",
    y = "log2(Enrichment Ratio)"
  ) +
  theme_minimal() +
  theme(axis.text.x = element_text(angle = 45, hjust = 1)), width = 8, height = 5)

message("Saved log2 enrichment ratio plots and table for: full_diagonal_over_other")

# ------------------- Nonlinear over Linear Enrichment Ratio -------------------
message("Calculating enrichment ratio: Nonlinear / Linear")

nonlinear_over_linear <- full_join(log2_nonlinear, log2_linear, by = c("Position", "Annotation"), suffix = c(".nonlinear", ".linear")) %>%
  mutate(
    enrichment_ratio = percent.nonlinear / percent.linear,
    log2_enrichment_ratio = log2(enrichment_ratio)
  )

# Barplot: Enrichment Ratio (Nonlinear / Linear)
p_nl_l_bar <- ggplot(nonlinear_over_linear, aes(x = fct_relevel(Annotation, my_levels), y = enrichment_ratio, fill = Position)) +
  geom_col(position = "dodge") +
  geom_hline(yintercept = 1, linetype = "dashed") +
  labs(
    title = "Enrichment Ratio: Nonlinear / Linear",
    x = "ChromHMM State",
    y = "Enrichment Ratio"
  ) +
  theme_minimal() +
  theme(axis.text.x = element_text(angle = 45, hjust = 1))

ggsave(file.path(output_dir, "nonlinear_over_linear_ratio_barplot.pdf"), plot = p_nl_l_bar, width = 8, height = 5)

# Heatmap: log2(Nonlinear / Linear)
p_nl_l_heat <- ggplot(nonlinear_over_linear, aes(x = fct_relevel(Annotation, my_levels), y = Position, fill = log2_enrichment_ratio)) +
  geom_tile(color = "white") +
  scale_fill_distiller(palette = "RdYlBu", direction = 1, name = "log2(NL / L)") +
  labs(
    title = "Heatmap of Enrichment Ratio: Nonlinear / Linear",
    x = "ChromHMM State",
    y = "Fragment Position"
  ) +
  theme_minimal() +
  theme(axis.text.x = element_text(angle = 45, hjust = 1))

ggsave(file.path(output_dir, "nonlinear_over_linear_ratio_heatmap.pdf"), plot = p_nl_l_heat, width = 10, height = 5)

# Barplot: log2(Nonlinear / Linear)
p_nl_l_log2_bar <- ggplot(nonlinear_over_linear, aes(x = fct_relevel(Annotation, my_levels), y = log2_enrichment_ratio, fill = Position)) +
  geom_col(position = "dodge") +
  geom_hline(yintercept = 0, linetype = "dashed") +
  labs(
    title = "Log2 Enrichment Ratio: Nonlinear / Linear",
    x = "ChromHMM State",
    y = "log2(Enrichment Ratio)"
  ) +
  theme_minimal() +
  theme(axis.text.x = element_text(angle = 45, hjust = 1))

ggsave(file.path(output_dir, "nonlinear_over_linear_log2ratio_barplot.pdf"), plot = p_nl_l_log2_bar, width = 8, height = 5)

# Save data
write_csv(nonlinear_over_linear, file.path(output_dir, "nonlinear_over_linear_ratio_table.csv"))
message("Saved enrichment ratio plots and table for: nonlinear_over_linear")
```
