## Supplementary material for "CAD-C: An engineered nuclease enables repair-free *in situ* proximity ligation and nucleosome-resolution chromosome walks in human cells": Custom code: submit_chromHMM_fragmentdist_barplots_20250528.sh

```
#!/bin/bash
#SBATCH -n 1
#SBATCH -c 30
#SBATCH --verbose
#SBATCH -J fragments
#SBATCH -e %x-%j-%a.err
#SBATCH -o %x-%j-%a.out


# Activate Conda environment
source /ru-auth/local/home/risc_soft2/miniconda3/etc/profile.d/conda.sh
conda activate lauren

# Run R script with specified input and output directory
Rscript /lustre/fs4/risc_lab/scratch/landerson01/20250205_CAD_2x250bp_GM_analysis/ChromHMM/Cis/Fragment_Distribution_20250528.r
```
