## Supplementary material for "CAD-C: An engineered nuclease enables repair-free *in situ* proximity ligation and nucleosome-resolution chromosome walks in human cells": Custom code: replot_Dmax_J_M_by_cisrunlen_v1_GPT_20251020.py

```
#!/usr/bin/env python3
# -*- coding: utf-8 -*-
"""
Dmax overlay (J or M) — true semilog x-axis with preserved log-space density
============================================================================

Author: Jan Soroczynski (Risca Lab, The Rockefeller University)
Date   : 2025_10_21
Version: 2.0.0

Overview
--------
This script plots overlaid *Dmax* distributions from a CADpack **v9** NPZ for chosen
run-length strata (e.g., L==1, L==2, L==3, L>=4). It reproduces the exact style we
agreed on:

  • **Normalization**: density per log10(bp) (area = 1 in log-space).
  • **Axis**: true log-x with decade major ticks and 2–9 minor ticks.
  • **Styling**: bottom ticks only, outward tick direction, clean frame (no top/right
    spines), Illustrator-friendly **fonttype=42**, and larger fonts by default.

It supports Dmax computed in either **J** (junction) space or **M** (midpoint) space.
For CADpack v9 NPZs, both `run_Dmax_J` and `run_Dmax_M` should be present; for safety
the script can re-compute Dmax_J from step endpoints if the run-level array is absent.

Selection / Gating
------------------
• By default, the script restricts to **cis-confined, single-chromosome** runs:
  every step is cis and all steps lie on the same chromosome. Disable with
  `--no-cis-only` if you truly need everything.

• Strata are flexible, provided as boolean expressions on run length `L`:
  e.g. `--strata "L==1;L==2;L==3;L>=4"` (matching `--labels`).

Inputs
------
• `--npz`  : path to CADpack **v9** NPZ (merged tech reps or otherwise).
• The NPZ must contain at minimum:
  - run tables: `run_n_steps`, `run_first_step_row`
  - step tables: `chrom1_id`, `chrom2_id`, `pos1`, `pos2`, `step_is_cis`
  - run Dmax: `run_Dmax_J` or `run_Dmax_M` (or both)
  If `run_Dmax_J` is missing, the script recomputes it per run as the maximum span
  of cis step endpoints within the run’s chromosome.

Outputs
-------
• **PDF** figure (fonttype=42), bottom semilog ticks only.
• **CSV** stats table with per-stratum:
  `metric,label,n,median_bp,p25_bp,p75_bp,mean_log10_bp`.

Usage (examples)
----------------
# Dmax_J with standard strata and color palette, larger fonts
python dmax_overlay_cli_v2.py \
  --npz cadpack_bc1001_bc1002_v9_14422039.npz \
  --metric J \
  --outpdf DmaxJ_overlay.pdf \
  --stats  DmaxJ_stats.csv \
  --strata "L==1;L==2;L==3;L>=4" \
  --labels "run_len = 1;run_len = 2;run_len = 3;run_len ≥ 4" \
  --colors "#3b0f70,#5c1e9b,#2878b5,#2aa198" \
  --xmin 1e2 --xmax 1e9 --bin-dex 0.1 \
  --font-base 28

# Dmax_M with custom strata (leave default colors)
python dmax_overlay_cli_v2.py \
  --npz cadpack_bc1001_bc1002_v9_14422039.npz \
  --metric M \
  --outpdf DmaxM_overlay.pdf \
  --stats  DmaxM_stats.csv \
  --strata "L==1;L==2;L>=3" \
  --font-base 28

CLI Reference
-------------
--npz         Path to CADpack v9 NPZ (required)
--metric      Which Dmax to plot: J | M   (default: J)
--outpdf      Output PDF path             (required)
--stats       Output CSV path             (required)
--strata      Semicolon-separated boolean expressions on L
              (default: "L==1;L==2;L==3;L>=4")
--labels      Matching semicolon-separated legend labels
              (default: "run_len = 1;run_len = 2;run_len = 3;run_len ≥ 4")
--colors      Comma-separated colors (hex or names); if omitted, Matplotlib cycle
--xmin        X-min in bp (default 1e2)
--xmax        X-max in bp (default 1e9)
--bin-dex     Bin width in decades for log-space histogram (default 0.1)
--font-base   Base font size for labels/ticks/legend (default 26)
--cis-only    Keep cis-confined single-chrom runs (default ON)
--no-cis-only Disable the cis/same-chrom restriction

Methodological Notes (important)
--------------------------------
• We histogram **log10(Dmax)** with equal-width bins and `density=True`, which yields a
  probability density per *dex* (per log10 bp). To show real semilog ticks while
  preserving that normalization, we plot **stairs** in **D space** using edges
  `10**(log10_edges)` on a log-x axis. This preserves “area = 1 in log-space.”

• Axis & ticks: major decades + minor ticks (2..9 per decade), ticks drawn outward,
  labels only on the bottom axis (no top “floating” ticks).

Dependencies
------------
Python ≥3.8, NumPy, Matplotlib.

Reproducibility
---------------
All plotting rcParams (fonts, fonttype, tick sizes) are set inside the script based on
`--font-base`. PDFs embed TrueType (fonttype=42) for Illustrator-friendly text.

License
-------
MIT — include attribution above.

"""

import argparse
import csv
import math
import os
from typing import List, Tuple

import numpy as np
import matplotlib
import matplotlib.pyplot as plt
from matplotlib.ticker import LogLocator, LogFormatterMathtext


def _update_fonts(base: float) -> None:
    """Set Illustrator-friendly font embedding and scaled text sizes."""
    matplotlib.rcParams.update(
        {
            "pdf.fonttype": 42,
            "ps.fonttype": 42,
            "axes.unicode_minus": False,
            "font.size": base * 1.00,
            "axes.labelsize": base * 1.00,
            "axes.titlesize": base * 1.15,
            "xtick.labelsize": base * 0.90,
            "ytick.labelsize": base * 0.90,
            "legend.fontsize": base * 0.90,
        }
    )


def _parse_strata(exprs: str, Lvals: np.ndarray) -> List[np.ndarray]:
    """
    Parse semicolon-separated boolean expressions in L (e.g., "L==1;L>=4")
    and return boolean masks for each expression.
    """
    masks: List[np.ndarray] = []
    for expr in exprs.split(";"):
        expr = expr.strip()
        if not expr:
            continue
        if expr.startswith("L=="):
            v = int(expr[3:])
            masks.append(Lvals == v)
        elif expr.startswith("L>="):
            v = int(expr[3:])
            masks.append(Lvals >= v)
        elif expr.startswith("L<="):
            v = int(expr[3:])
            masks.append(Lvals <= v)
        elif expr.startswith("L>"):
            v = int(expr[2:])
            masks.append(Lvals > v)
        elif expr.startswith("L<"):
            v = int(expr[2:])
            masks.append(Lvals < v)
        else:
            raise ValueError(f"Unsupported stratum expression: {expr}")
    return masks


def _cis_singlechrom_mask(
    chrom1_id: np.ndarray,
    chrom2_id: np.ndarray,
    step_is_cis: np.ndarray,
    run_first: np.ndarray,
    run_n: np.ndarray,
) -> np.ndarray:
    """True for runs where all steps are cis and all steps share a single chromosome."""
    keep = np.zeros(run_n.size, dtype=bool)
    for i in range(run_n.size):
        s0 = int(run_first[i])
        n = int(run_n[i])
        if n <= 0:
            continue
        S = slice(s0, s0 + n)
        if not step_is_cis[S].all():
            continue
        chr0 = chrom1_id[s0]
        if np.all(chrom1_id[S] == chr0) and np.all(chrom2_id[S] == chr0):
            keep[i] = True
    return keep


def _recompute_dmax_j_if_missing(npz: dict) -> np.ndarray:
    """
    If run_Dmax_J is absent, recompute per run as the maximum span across cis steps
    within each run’s chromosome using step endpoints (pos1/pos2).
    """
    chrom1_id = npz["chrom1_id"].astype(np.int32)
    chrom2_id = npz["chrom2_id"].astype(np.int32)
    pos1 = npz["pos1"].astype(np.int64)
    pos2 = npz["pos2"].astype(np.int64)
    step_is_cis = npz["step_is_cis"].astype(bool)
    run_first = npz["run_first_step_row"]
    run_n = npz["run_n_steps"]

    out = np.zeros(run_n.size, dtype=np.float64)
    for i in range(run_n.size):
        s0 = int(run_first[i])
        n = int(run_n[i])
        if n <= 0:
            continue
        S = slice(s0, s0 + n)
        mask = step_is_cis[S]
        if not mask.any():
            continue
        dmax = 0.0
        # For safety, compute per run without chrom split (all steps are same chrom if gated).
        coords = np.concatenate([pos1[S][mask], pos2[S][mask]])
        if coords.size:
            dmax = float(coords.max() - coords.min())
        out[i] = dmax
    return out


def plot_dmax_overlay(
    *,
    npz_path: str,
    metric: str,
    outpdf: str,
    stats_csv: str,
    strata_expr: str,
    labels: List[str],
    colors: List[str],
    xmin_bp: float,
    xmax_bp: float,
    bin_dex: float,
    font_base: float,
    cis_only: bool,
) -> None:
    """Core plotting + stats writer."""
    _update_fonts(font_base)

    Z = np.load(npz_path, allow_pickle=True)

    # Select metric (with safe fallback for J).
    if metric.upper() == "M":
        if "run_Dmax_M" not in Z:
            raise SystemExit("run_Dmax_M not found in NPZ.")
        Dmax = Z["run_Dmax_M"].astype(np.float64)
    else:
        if "run_Dmax_J" in Z:
            Dmax = Z["run_Dmax_J"].astype(np.float64)
        else:
            Dmax = _recompute_dmax_j_if_missing(Z)

    L = Z["run_n_steps"]
    keep = np.ones(L.size, dtype=bool)
    if cis_only:
        keep = _cis_singlechrom_mask(
            Z["chrom1_id"].astype(np.int32),
            Z["chrom2_id"].astype(np.int32),
            Z["step_is_cis"].astype(bool),
            Z["run_first_step_row"],
            Z["run_n_steps"],
        )

    Lvals = L[keep]
    Dvals = Dmax[keep].astype(np.float64)

    # Build strata
    masks = _parse_strata(strata_expr, Lvals)
    if len(labels) != len(masks):
        raise SystemExit("Number of --labels must match number of --strata")
    use_colors = len(colors) == len(masks)

    # Log-space histogram bins
    xmin_dex = math.log10(xmin_bp)
    xmax_dex = math.log10(xmax_bp)
    bins_log = np.arange(xmin_dex, xmax_dex + bin_dex, bin_dex)
    edges_D = 10 ** bins_log

    # Figure
    fig = plt.figure(figsize=(14.0, 6.6), constrained_layout=True)
    ax = fig.add_subplot(1, 1, 1)
    ax.set_xscale("log", base=10)

    ymax = 0.0
    # Draw each stratum
    for k, (mask, lab) in enumerate(zip(masks, labels)):
        data = Dvals[mask]
        data = data[np.isfinite(data) & (data > 0)]
        n = data.size
        if n == 0:
            continue
        dens, _ = np.histogram(np.log10(data), bins=bins_log, density=True)
        col = colors[k] if use_colors else None
        ax.step(
            edges_D[:-1],
            dens,
            where="post",
            color=col,
            linewidth=3.0,
            label=f"{lab} (n={n:,})",
        )
        ymax = max(ymax, float(np.nanmax(dens)))

    # Semilog ticks (bottom only)
    ax.set_xlim(xmin_bp, xmax_bp)
    ax.xaxis.set_major_locator(LogLocator(base=10.0, numticks=8))
    ax.xaxis.set_major_formatter(LogFormatterMathtext(base=10.0))
    ax.xaxis.set_minor_locator(LogLocator(base=10.0, subs=np.arange(2, 10) * 0.1, numticks=80))
    ax.tick_params(axis="x", which="both", direction="out", top=False, bottom=True, labeltop=False)
    ax.tick_params(axis="y", which="both", direction="out")

    ax.set_ylim(0, ymax * 1.08 if ymax > 0 else 1)
    ax.set_ylabel("density per log$_{10}$ bp (area = 1 in log-space)")
    ax.set_xlabel(f"Max within-run separation, Dmax$_{metric.upper()}$ (bp)")
    ax.set_title("Overlay with semilog ticks — equal log-space bins, density per log$_{10}$ bp")
    ax.legend(frameon=False, loc="upper right", ncol=2)
    ax.grid(False)
    for spine in ("top", "right"):
        ax.spines[spine].set_visible(False)

    fig.savefig(outpdf, dpi=300)
    plt.close(fig)

    # Stats CSV
    with open(stats_csv, "w", newline="") as fh:
        w = csv.writer(fh)
        w.writerow(["metric", "label", "n", "median_bp", "p25_bp", "p75_bp", "mean_log10_bp"])
        for mask, lab in zip(masks, labels):
            data = Dvals[mask]
            data = data[np.isfinite(data) & (data > 0)]
            if data.size == 0:
                w.writerow([metric.upper(), lab, 0, "", "", "", ""])
                continue
            p25, med, p75 = np.percentile(data, [25, 50, 75])
            mean_log10 = float(np.mean(np.log10(data)))
            w.writerow([metric.upper(), lab, int(data.size), float(med), float(p25), float(p75), mean_log10])


def _parse_labels(s: str) -> List[str]:
    return [x.strip() for x in s.split(";") if x.strip()]


def _parse_colors(s: str) -> List[str]:
    return [x.strip() for x in s.split(",") if x.strip()]


def main() -> None:
    ap = argparse.ArgumentParser(
        description="Dmax overlay with true semilog ticks and preserved log-space density (CADpack v9)."
    )
    ap.add_argument("--npz", required=True, help="Path to CADpack v9 NPZ.")
    ap.add_argument("--metric", choices=["J", "M"], default="J", help="Which Dmax to plot (default: J).")
    ap.add_argument("--outpdf", required=True, help="Output PDF filename.")
    ap.add_argument("--stats", required=True, help="Output CSV for per-stratum stats.")
    ap.add_argument(
        "--strata", default="L==1;L==2;L==3;L>=4", help='Semicolon-separated strata in L (e.g., "L==1;L>=4").'
    )
    ap.add_argument(
        "--labels",
        default="run_len = 1;run_len = 2;run_len = 3;run_len ≥ 4",
        help="Semicolon-separated legend labels (must match --strata).",
    )
    ap.add_argument(
        "--colors",
        default="#3b0f70,#5c1e9b,#2878b5,#2aa198",
        help="Comma-separated colors (hex or names). If count mismatches strata, Matplotlib cycle is used.",
    )
    ap.add_argument("--xmin", type=float, default=1e2, help="X-axis min in bp (default 1e2).")
    ap.add_argument("--xmax", type=float, default=1e9, help="X-axis max in bp (default 1e9).")
    ap.add_argument("--bin-dex", type=float, default=0.1, help="Bin width in decades for log-space histogram.")
    ap.add_argument("--font-base", type=float, default=26, help="Base font size for labels/ticks/legend.")
    ap.add_argument("--cis-only", action="store_true", default=True, help="Keep cis-confined single-chrom runs (default).")
    ap.add_argument("--no-cis-only", dest="cis_only", action="store_false")

    args = ap.parse_args()

    labels = _parse_labels(args.labels)
    colors = _parse_colors(args.colors)

    plot_dmax_overlay(
        npz_path=args.npz,
        metric=args.metric,
        outpdf=args.outpdf,
        stats_csv=args.stats,
        strata_expr=args.strata,
        labels=labels,
        colors=colors,
        xmin_bp=args.xmin,
        xmax_bp=args.xmax,
        bin_dex=args.bin_dex,
        font_base=args.font_base,
        cis_only=args.cis_only,
    )


if __name__ == "__main__":
    main()
```
