## Supplementary material for "CAD-C: An engineered nuclease enables repair-free *in situ* proximity ligation and nucleosome-resolution chromosome walks in human cells": Custom code: merged_dedup_vs_final_replot_summary_GPT_v3_20251107.py

```
#!/usr/bin/env python3
# -*- coding: utf-8 -*-
from __future__ import annotations
import argparse, glob, hashlib, io, json, math, os, re, sys
from dataclasses import dataclass
from datetime import datetime
from pathlib import Path
from typing import Dict, Iterable, List, Tuple, Optional, Union
from zipfile import ZipFile, BadZipFile

import numpy as np
import pandas as pd
import matplotlib
matplotlib.rcParams["pdf.fonttype"] = 42          # Illustrator‑friendly (TrueType)
matplotlib.rcParams["ps.fonttype"]  = 42
matplotlib.rcParams["text.usetex"]  = False
matplotlib.rcParams["axes.grid"]    = False       # no gridlines
import matplotlib.pyplot as plt
from matplotlib.patches import Rectangle

# Colors (match your v4 CLI)
COL_LT1   = "#c6dbef"   # <1 kb (cis)
COL_1_20  = "#6baed6"   # 1–20 kb (cis)
COL_GT20  = "#2171b5"   # >20 kb (cis)
COL_TRANS = "#d81b60"   # trans
COL_RET   = "#2ca02c"   # retained UU
COL_REM   = "#e0e0e0"   # remainder

# Default display order (can be overridden by --order)
PRIORITY_ORDER = ["ao_microc","e1_p2andp3","gm12878_1_merged","k562_1_merged","darrow2016","gm12878_deepmerge"]

@dataclass
class StatsRow:
    file: str; stage: str; key: str
    UU: float; cis_lt1kb: float; cis_1_20kb: float; cis_gt20kb: float; trans: float
    cis_total: float; cistrans_total: float; cis_pct: float

def timestamp_now(): return datetime.now().strftime("%Y_%m_%d_%H%M%S")

def build_outdir(preferred: Optional[str], exemplar: Optional[str], prefix: str) -> Path:
    ts = timestamp_now()
    if preferred: out = Path(preferred).expanduser().resolve()
    else:
        base = Path(exemplar).expanduser().resolve().parent if exemplar else Path.cwd()
        out = base / f"qc_pairs_{prefix}_{ts}"
    out.mkdir(parents=True, exist_ok=True)
    return out

def is_appledouble(name: str) -> bool:
    base = Path(name).name
    return base.startswith("._") or "/._" in name

def canonical_key(fname: str) -> str:
    base = Path(fname).name
    stem = base.split("_stats", 1)[0] if "_stats" in base else Path(base).stem
    toks = re.split(r"[._\-]+|_", stem)
    kept: List[str] = []
    for t in toks:
        if not t: continue
        tl = t.lower()
        if tl in {"dedup","dedupfinal","rededup","final","stats"}: continue
        if tl == "dedup_final": continue
        if t == "CADC": continue
        if re.fullmatch(r"\d{6,}", t): continue
        kept.append(tl)
    kept = [("darrow2016" if k.startswith("darrow2016") else k) for k in kept]
    return "_".join(kept)

def parse_kv_text(text: str) -> Dict[str, str]:
    d: Dict[str, str] = {}
    for ln in text.splitlines():
        if not ln.strip() or ln.startswith("#"): continue
        parts = ln.rstrip("\n").split("\t")
        if len(parts) >= 2: d[parts[0]] = parts[1]
    return d

def fnum(v) -> float:
    if v is None or v == "": return float("nan")
    try: return float(v)
    except Exception:
        try: return float(int(v))
        except Exception: return float("nan")

def read_stats_from_text(text: str):
    kv = parse_kv_text(text)
    UU    = fnum(kv.get("pair_types/UU"))
    cis   = fnum(kv.get("cis"))
    cis1k = fnum(kv.get("cis_1kb+")) if kv.get("cis_1kb+") is not None else fnum(kv.get("cis_1kbp+"))
    cis20 = fnum(kv.get("cis_20kb+"))
    trans = fnum(kv.get("trans"))
    lt1k = cis - cis1k if (math.isfinite(cis) and math.isfinite(cis1k)) else float("nan")
    mid  = cis1k - cis20 if (math.isfinite(cis1k) and math.isfinite(cis20)) else float("nan")
    gt20 = cis20
    ctot = (0 if not math.isfinite(lt1k) else lt1k) + (0 if not math.isfinite(mid) else mid) + (0 if not math.isfinite(gt20) else gt20)
    cistrans_total = ctot + (0 if not math.isfinite(trans) else trans)
    cis_pct = (100.0 * ctot / cistrans_total) if cistrans_total > 0 else float("nan")
    return UU, lt1k, mid, gt20, trans, ctot, cistrans_total, cis_pct

def infer_stage_from_filename_or_text(fname: str, text: str) -> str:
    bl = fname.lower(); tl = text.lower()
    if ("dedup_final" in bl) or ("dedupfinal" in bl) or ("dedup_final" in tl) or ("dedupfinal" in tl): return "dedup_final"
    if re.search(r"(?<!re)dedup(?:$|[^a-z0-9])", bl) and ("dedup_final" not in bl): return "dedup"
    if re.search(r"(?<!re)dedup(?:$|[^a-z0-9])", tl) and ("dedup_final" not in tl): return "dedup"
    return "dedup_final"

def sha256_file(path: Union[str, Path]) -> str:
    try:
        import hashlib
        h=hashlib.sha256()
        with open(path,"rb") as fh:
            for chunk in iter(lambda: fh.read(1024*1024), b""): h.update(chunk)
        return h.hexdigest()
    except Exception: return ""

def collect_inputs(args):
    srcs=[]
    for g in args.input_glob:
        for p in glob.glob(g): srcs.append(("file", p))
    for p in args.inputs: srcs.append(("file", p))
    if args.inputs_file:
        with open(args.inputs_file,"r") as fh:
            for ln in fh:
                ln=ln.strip()
                if ln: srcs.append(("file", ln))
    if args.input_zip: srcs.append(("zip", args.input_zip))
    return srcs

@dataclass
class Row(StatsRow): pass

def load_all_stats(args):
    alias_rules=[]
    for rule in args.key_alias:
        if "=" not in rule: continue
        pat, alias = rule.split("=",1)
        alias_rules.append((re.compile(pat), alias))
    def aliased(basename, key):
        for rx, alias in alias_rules:
            if rx.search(basename): return alias
        return key
    rows=[]
    for src_type, payload in collect_inputs(args):
        if src_type=="file":
            p=Path(payload).expanduser()
            if not p.exists(): continue
            base=p.name
            if base in args.omit_file or is_appledouble(base): continue
            text=p.read_text(encoding="utf-8", errors="replace")
            UU,lt1k,mid,gt20,trans,ctot,cistrans_total,cis_pct=read_stats_from_text(text)
            key=aliased(base, canonical_key(base))
            rows.append(Row(str(p.resolve()), infer_stage_from_filename_or_text(base,text), key,
                            UU,lt1k,mid,gt20,trans,ctot,cistrans_total,cis_pct))
        else:
            zp=Path(payload).expanduser()
            try:
                with ZipFile(zp,"r") as z:
                    for name in z.namelist():
                        if not name.lower().endswith(".tsv"): continue
                        base=Path(name).name
                        if base in args.omit_file or is_appledouble(base): continue
                        raw=z.read(name).decode("utf-8", errors="replace")
                        UU,lt1k,mid,gt20,trans,ctot,cistrans_total,cis_pct=read_stats_from_text(raw)
                        key=aliased(base, canonical_key(base))
                        rows.append(Row(f"{str(zp.resolve())}::{name}", infer_stage_from_filename_or_text(base,raw), key,
                                        UU,lt1k,mid,gt20,trans,ctot,cistrans_total,cis_pct))
            except BadZipFile:
                print(f"[WARN] Not a valid zip: {zp}", file=sys.stderr)
    return rows

def filter_rows_like_last_stable(rows, allow_all_e1=False, keep_rpe1_1=False):
    out=[]
    for r in rows:
        base=Path(r.file).name
        if (not keep_rpe1_1) and ("CADC_RPE1_1" in base): continue
        if (("CADC_E1_" in base) and ("CADC_E1_P2andP3_" not in base) and (not allow_all_e1)): continue
        out.append(r)
    return out

def build_pairs(rows):
    by_key={}
    for r in rows: by_key.setdefault(r.key, []).append(r)
    pairs=[]
    for key,lst in by_key.items():
        dedups=[x for x in lst if x.stage=="dedup"]; finals=[x for x in lst if x.stage=="dedup_final"]
        if not dedups or not finals: continue
        de=sorted(dedups, key=lambda x: (x.cistrans_total if math.isfinite(x.cistrans_total) else -1), reverse=True)[0]
        df=sorted(finals, key=lambda x: (x.cistrans_total if math.isfinite(x.cistrans_total) else -1), reverse=True)[0]
        pairs.append((key,de,df))
    return pairs

def order_pairs(pairs, custom_order=None):
    order=custom_order or PRIORITY_ORDER
    def ok(k):
        for i,tag in enumerate(order):
            if tag in k: return (i,k)
        return (len(order)+1, k)
    return sorted(pairs, key=lambda t: ok(t[0]))

def style_axes(ax):
    ax.grid(False); ax.yaxis.grid(False); ax.xaxis.grid(False)
    ax.tick_params(axis="both", which="major", direction="out", length=8, width=1.8, labelsize=11)
    ax.spines["top"].set_visible(False); ax.spines["right"].set_visible(False)
    ax.spines["left"].set_linewidth(1.8); ax.spines["bottom"].set_linewidth(1.8)

def plot_cistrans_paired_stacked(outdir, prefix, pairs):
    if not pairs: return
    labels=[]; segs={"cis_lt1kb":[], "cis_1_20kb":[], "cis_gt20kb":[], "trans":[]}
    for _,de,df in pairs:
        for row in (de,df):
            tot=row.cistrans_total if math.isfinite(row.cistrans_total) and row.cistrans_total>0 else float("nan")
            for k in segs:
                v=getattr(row,k); pct=(100.0*float(v)/tot) if (math.isfinite(v) and math.isfinite(tot) and tot>0) else 0.0
                segs[k].append(pct)
            labels.append(Path(row.file).name)
    n_pairs=len(pairs); n_bars=2*n_pairs; width=0.86; within_gap=0.30; between_gap=1.10
    xs=[]; x=0.0
    for _ in range(n_pairs): xs.append(x); xs.append(x+width+within_gap); x+=(2*width+within_gap+between_gap)
    fig_w=max(10.0, 0.90*n_bars+6.5); fig_h=7.4
    fig, ax = plt.subplots(figsize=(fig_w, fig_h))
    bottom=np.zeros(n_bars)
    order=[("cis_lt1kb",COL_LT1,"<1 kb (cis)"), ("cis_1_20kb",COL_1_20,"1–20 kb (cis)"),
           ("cis_gt20kb",COL_GT20,">20 kb (cis)"), ("trans",COL_TRANS,"trans")]
    for seg,color,lab in order:
        vals=np.nan_to_num(np.array(segs[seg], dtype=float), nan=0.0)
        ax.bar(xs, vals, width=width, bottom=bottom, color=color, linewidth=0, label=lab)
        for xi,v,b in zip(xs,vals,bottom):
            if v>=1.0: ax.text(xi+width/2.0, b+v/2.0, f"{v:.1f}%", ha="center", va="center", fontsize=10)
        bottom+=vals
    ax.set_xticks([xi+width/2.0 for xi in xs]); ax.set_xticklabels(labels, rotation=35, ha="right", fontsize=10)
    ax.set_ylim(0,100); ax.set_ylabel("Composition (%)", fontsize=13)
    ax.legend([Rectangle((0,0),1,1,color=c) for _,c,_ in order], [lab for _,_,lab in order],
              loc="center left", bbox_to_anchor=(1.01,0.5), frameon=False, fontsize=11)
    ax.set_title("cis distance strata + trans — dedup vs dedup_final (paired, normalized)", fontsize=14, pad=10); style_axes(ax)
    p_png=outdir / f"{prefix}_cistrans_paired_stacked.png"; p_pdf=outdir / f"{prefix}_cistrans_paired_stacked.pdf"
    plt.tight_layout(rect=[0.04,0.04,0.86,0.98]); plt.savefig(p_png, dpi=200); plt.savefig(p_pdf); plt.close(fig)

def plot_uu_retention(outdir, prefix, pairs):
    if not pairs: return
    labels=[]; retained=[]; remainder=[]
    for _,de,df in pairs:
        r=float("nan")
        if (math.isfinite(de.UU) and de.UU>0 and math.isfinite(df.UU) and df.UU>=0): r=max(0.0, min(100.0, 100.0*df.UU/de.UU))
        if not math.isfinite(r): r=0.0
        retained.append(r); remainder.append(100.0-r); labels.append(Path(df.file).name)
    n=len(labels); x=np.arange(n); width=0.86
    fig_w=max(8.0, 0.90*n+6.5); fig_h=6.8
    fig, ax = plt.subplots(figsize=(fig_w, fig_h))
    ax.bar(x,retained,width=width,color=COL_RET,linewidth=0,label="Retained UU (Pairs with Ligation Junction Observed)")
    ax.bar(x,remainder,width=width,bottom=retained,color=COL_REM,linewidth=0,label="Remainder to 100%")
    for xi,r in zip(x,retained):
        if r>=1.0: ax.text(xi, r/2.0, f"{r:.1f}%", ha="center", va="center", fontsize=11, color="black")
    ax.set_xticks(x); ax.set_xticklabels(labels, rotation=35, ha="right", fontsize=10)
    ax.set_ylim(0,100); ax.set_ylabel("Retention vs. dedup UU (%)", fontsize=13)
    ax.legend(loc="center left", bbox_to_anchor=(1.01,0.5), frameon=False, fontsize=11)
    ax.set_title("UU retention — dedup_final vs dedup (baseline = dedup UU)", fontsize=14, pad=10); style_axes(ax)
    p_png=outdir / f"{prefix}_uu_retention_vs_dedup.png"; p_pdf=outdir / f"{prefix}_uu_retention_vs_dedup.pdf"
    plt.tight_layout(rect=[0.04,0.04,0.86,0.98]); plt.savefig(p_png, dpi=200); plt.savefig(p_pdf); plt.close(fig)

def _to_Int64(x):
    if x is None or not math.isfinite(x): return pd.NA
    try: return int(round(float(x)))
    except Exception: return pd.NA

def export_master_csv_ordered(outdir, prefix, rows, ordered_pairs):
    # Build master to match the plot order exactly: for each pair → [dedup, dedup_final]
    ordered_rows = []
    for k, de, df in ordered_pairs:
        ordered_rows.append(de)
        ordered_rows.append(df)
    # Any remaining rows (should not occur) appended at the end
    seen = {id(r) for r in ordered_rows}
    for r in rows:
        if id(r) not in seen: ordered_rows.append(r)
    cols=["file","stage","key","UU","cis_lt1kb","cis_1_20kb","cis_gt20kb","trans","cis_total","cistrans_total","cis_pct"]
    df=pd.DataFrame([{c:getattr(r,c) for c in cols} for r in ordered_rows])
    for c in ["UU","cis_lt1kb","cis_1_20kb","cis_gt20kb","trans","cis_total","cistrans_total"]:
        df[c]=df[c].map(_to_Int64)
    p=outdir / f"{prefix}_master.csv"; df.to_csv(p, index=False); return p

def export_pairs_summary_csv(outdir, prefix, pairs):
    recs=[]
    for idx,(key,de,df) in enumerate(pairs, start=1):
        uu_ret=float("nan")
        if (math.isfinite(de.UU) and de.UU>0 and math.isfinite(df.UU) and df.UU>=0): uu_ret=max(0.0, min(100.0, 100.0*df.UU/de.UU))
        recs.append({"pair_order":idx,"pair_key":key,"dedup_file":Path(de.file).name,"dedup_final_file":Path(df.file).name,
                     "UU_dedup":_to_Int64(de.UU),
                     "dedup_cis_lt1kb":_to_Int64(de.cis_lt1kb),"dedup_cis_1_20kb":_to_Int64(de.cis_1_20kb),"dedup_cis_gt20kb":_to_Int64(de.cis_gt20kb),"dedup_trans":_to_Int64(de.trans),
                     "dedup_cis_total":_to_Int64(de.cis_total),"dedup_cistrans_total":_to_Int64(de.cistrans_total),"dedup_cis_pct":de.cis_pct,
                     "UU_dedup_final":_to_Int64(df.UU),
                     "dedup_final_cis_lt1kb":_to_Int64(df.cis_lt1kb),"dedup_final_cis_1_20kb":_to_Int64(df.cis_1_20kb),"dedup_final_cis_gt20kb":_to_Int64(df.cis_gt20kb),"dedup_final_trans":_to_Int64(df.trans),
                     "dedup_final_cis_total":_to_Int64(df.cis_total),"dedup_final_cistrans_total":_to_Int64(df.cistrans_total),"dedup_final_cis_pct":df.cis_pct,
                     "UU_retention_pct_vs_dedup":uu_ret})
    p=outdir / f"{prefix}_pairs_summary.csv"; pd.DataFrame(recs).to_csv(p, index=False); return p

def export_provenance(outdir, prefix, rows, args_ns):
    prov_rows=[]
    for r in rows:
        if "::" in r.file: abs_path=r.file; sha=""; size=""; mtime=""
        else:
            p=Path(r.file); abs_path=str(p.resolve())
            try:
                import hashlib
                sha=hashlib.sha256(open(p,'rb').read()).hexdigest(); st=p.stat()
                size=st.st_size; mtime=datetime.fromtimestamp(st.st_mtime).isoformat()
            except Exception: sha=""; size=""; mtime=""
        prov_rows.append({"file":abs_path,"stage":r.stage,"key":r.key,"sha256":sha,"filesize_bytes":size,"mtime_iso":mtime})
    p_csv=outdir / f"{prefix}_provenance.csv"; pd.DataFrame(prov_rows).to_csv(p_csv, index=False)
    env={"script":Path(sys.argv[0]).name,"version":"v3.0.0","timestamp":timestamp_now(),
         "python":sys.version,"pandas":pd.__version__,"numpy":np.__version__,"matplotlib":matplotlib.__version__,
         "rcParams":{"pdf.fonttype":matplotlib.rcParams["pdf.fonttype"],"ps.fonttype":matplotlib.rcParams["ps.fonttype"],"axes.grid":matplotlib.rcParams["axes.grid"]},
         "args":vars(args_ns)}
    p_json=outdir / f"{prefix}_args_env.json"; p_json.write_text(json.dumps(env, indent=2)); return p_csv,p_json

def main():
    ap=argparse.ArgumentParser(description="Pairtools stats → paired cistrans plot, UU retention, CSVs, provenance.")
    ap.add_argument("--inputs", action="append", default=[])
    ap.add_argument("--inputs-file", default=None)
    ap.add_argument("--input-glob", action="append", default=[])
    ap.add_argument("--input-zip", default=None)
    ap.add_argument("--omit-file", action="append", default=[])
    ap.add_argument("--key-alias", action="append", default=[])
    ap.add_argument("--allow-all-e1", action="store_true")
    ap.add_argument("--keep-rpe1-1", action="store_true")
    ap.add_argument("--order", default="")
    ap.add_argument("--prefix", default="CADC_summary")
    ap.add_argument("--outdir", default=None)
    ap.add_argument("--zip", action="store_true")
    args=ap.parse_args()

    rows=load_all_stats(args)
    if not rows:
        print("No inputs found.", file=sys.stderr); sys.exit(2)

    exemplar=None
    for r in rows:
        if "::" not in r.file: exemplar=r.file; break
    outdir=build_outdir(args.outdir, exemplar, args.prefix)

    rows=filter_rows_like_last_stable(rows, allow_all_e1=args.allow_all_e1, keep_rpe1_1=args.keep_rpe1_1)
    pairs=build_pairs(rows)
    if not pairs:
        print("No dedup ↔ dedup_final pairs detected after filtering.", file=sys.stderr); sys.exit(3)

    custom_order=[t.strip() for t in args.order.split(",")] if args.order.strip() else None
    pairs=order_pairs(pairs, custom_order)

    # v3: MASTER is ordered by couples to match the plots
    master_csv=export_master_csv_ordered(outdir, args.prefix, rows, pairs)
    pairs_csv=export_pairs_summary_csv(outdir, args.prefix, pairs)
    prov_csv, env_json=export_provenance(outdir, args.prefix, rows, args)

    plot_cistrans_paired_stacked(outdir, args.prefix, pairs)
    plot_uu_retention(outdir, args.prefix, pairs)

    if args.zip:
        import shutil; shutil.make_archive(str(outdir), "zip", root_dir=outdir)
        print("Zipped:", f"{str(outdir)}.zip")

    print("Done. Outputs in:", outdir)
    print("CSVs:", master_csv, pairs_csv, prov_csv)
    print("Env:", env_json)

if __name__=="__main__":
    main()
```
