## Supplementary material for "CAD-C: An engineered nuclease enables repair-free *in situ* proximity ligation and nucleosome-resolution chromosome walks in human cells": Custom code: pairtools_qc_cli_v4_20251031.py

```
#!/usr/bin/env python3
# -*- coding: utf-8 -*-
"""
pairtools_qc_cli_v4.py
======================

Purpose
-------
Produce compact, publication‑ready QC plots and tables from **pairtools dedup stats TSV files**.
Focus: replicate-aware summaries that are faithful to *pairtools* outputs and easy to drop into
LaTeX/Illustrator and GEO submissions.

Key outputs (per group of inputs)
---------------------------------
1) **Stacked bar (pair types)** — UU / U×(M|N) / (M|N)×(M|N); individual TRs + pooled “Merged”.
2) **% duplicates** — per‑TR filled dots + a **weighted mean** line/label (weighted by total pairs).
3) **Estimated complexity** — per‑TR dots on **log10 y‑axis** with ticks like `10⁹`, and the **arithmetic mean**
   labeled in scientific notation (e.g., `2.65 × 10⁸`).
4) **Stacked bar (cis distance strata + trans)** — <1 kb, 1–20 kb, >20 kb (light→dark blues), trans (magenta);
   individual TRs + pooled “Merged”.
5) **Pies (area ∝ UU unique reads)** — per‑TR and for pooled “Merged”, wedges = cis bins + trans; with a
   true‑area size legend (10M…1B) and right‑hand counts/% for the pooled “Merged”.
6) **Row‑of‑four combined** (pairtypes → %dups → complexity → cis/trans) as:
   - PNG (raster preview) and
   - **Vector PDF** stitched from the four panel PDFs **without rasterization** (Type 42 preserved).

Tables
------
- `<prefix>_master.csv` — sample, path, pair types counts/%, duplicates, complexity, cis/trans counts.
- `<prefix>_cistrans_summary.csv` — compact per‑sample cis/trans counts with percentages.
- `<prefix>_provenance.csv` — path, SHA‑256 (optional), file size bytes (GEO‑friendly provenance).

Illustrator & LaTeX
-------------------
- All PDFs are vector; **no mathtext**; **pdf.fonttype=42** and **ps.fonttype=42**.
- Each PDF contains a **copyable textbox** listing **absolute input paths**, placed below the axes and away
  from the plotting area. Paste directly from Illustrator.
- Consistent design: outward ticks only; uniform axes line widths; **no gridlines over bars**;
  legends sit outside; merged bar/pie labeled with side‑percentages.

Inputs
------
- Any number of pairtools **dedup** stats TSVs:
  * `--input-glob "/path/*dedup*_stats_*.tsv"`
  * `--inputs /path/a.tsv --inputs /path/b.tsv` (repeatable)
  * `--inputs-file filelist.txt` (one TSV path per line)
- Sample IDs:
  * If a filename matches `NNUEFS_(S#)_` the `S#` label is used; otherwise the basename is used.

Output directory
----------------
By default: `qc_<common-basename>_<YYYYmmdd_HHMMSS>` in the directory of the first input file.
Or provide `--outdir /path/to/out`. Use `--zip` to create a zipped archive of that directory on completion.

Assumptions & details
---------------------
- Pair types: UU; U×(M|N) = NU+UN+MU+UM; (M|N)×(M|N) = MM+MN+NM+NN.
- % duplicates: read from `total_dups`, `total_nodups` → weighted mean across replicates.
- **Complexity**: pulls from `summary/complexity_naive` if present, then other reasonable keys containing
  “complex”; plotted on log10 scale. Reported mean is the **arithmetic** mean of per‑TR estimates.
- cis/trans post‑dedup bins:
  - `cis`, `cis_1kb+` (or `cis_1kbp+`), `cis_20kb+`, `trans`
  - We derive `<1 kb = cis - cis_1kb+`, `1–20 kb = cis_1kb+ - cis_20kb+`, `>20 kb = cis_20kb+`.
  - “Merged” percentages come from **absolute sums across TRs**.

Dependencies
------------
- Python 3.8+; `numpy`, `pandas`, `matplotlib`
- Optional: `Pillow` (PNG row stitch preview), `PyPDF2>=2.0` (vector PDF row stitch)
  Install: `pip install numpy pandas matplotlib pillow PyPDF2`

Usage examples
--------------
# Run on four TRs and include checksums in CSVs
python pairtools_qc_cli_v3.py \
  --prefix GM12878_CADwalk_2x250 --checksum \
  --inputs /data/medium_TR1_dedup_stats.tsv \
  --inputs /data/medium_TR2_dedup_stats.tsv \
  --inputs /data/small_TR1_dedup_stats.tsv \
  --inputs /data/small_TR2_dedup_stats.tsv

# Glob + automatic output dir + ZIP the directory
python pairtools_qc_cli_v3.py \
  --input-glob "/data/GM12878/*dedup*_stats_*.tsv" \
  --prefix GM12878_run --zip

Verification tips
-----------------
- Open PDFs in Illustrator: axis text/ticks should be editable (TrueType).
- `pdffonts file.pdf` should report *type: TrueType* (not Type 3).
"""

from __future__ import annotations

import argparse
import glob
import hashlib
import math
import os
import re
import sys
import textwrap
import shutil
from pathlib import Path
from typing import Dict, List

import numpy as np
import pandas as pd
import matplotlib
matplotlib.rcParams["pdf.fonttype"] = 42
matplotlib.rcParams["ps.fonttype"]  = 42
matplotlib.rcParams["text.usetex"]  = False
import matplotlib.pyplot as plt
from matplotlib.patches import Circle

# Optional imaging lib for PNG stitching (preview)
try:
    from PIL import Image
    PIL_OK = True
except Exception:
    PIL_OK = False

# ---------------------------
# Style (locked to your preferences)
# ---------------------------
AX_LINEWIDTH   = 1.8
MAJOR_TICK_LEN = 8.0
AX_LABEL_FS    = 16
TICK_LABEL_FS  = 14
LEGEND_FS      = 12
DOT_SIZE       = 110
DEEP_RED       = "#b22222"

COL_UU   = "#2ca02c"  # UU (green)
COL_UxMN = "#8DAE8C"  # U×(M|N) (grey‑green)
COL_MN   = "#7f7f7f"  # (M|N)×(M|N) (grey)

COL_LT1   = "#c6dbef"  # <1 kb (cis) light blue
COL_1_20  = "#6baed6"  # 1–20 kb (cis) mid blue
COL_GT20  = "#2171b5"  # >20 kb (cis) dark blue
COL_TRANS = "#d81b60"  # trans (magenta)

# Unicode superscripts helpers (no mathtext anywhere)
SUP = str.maketrans("0123456789-+", "⁰¹²³⁴⁵⁶⁷⁸⁹⁻⁺")
def to_sup(k:int)->str:
    return f"10{str(k).translate(SUP)}"

def sci_txt(val: float) -> str:
    if not (isinstance(val, (int, float)) and math.isfinite(val)) or val <= 0:
        return "NA"
    exp = int(math.floor(math.log10(val)))
    mant = val/(10.0**exp)
    return f"{mant:.2f} × 10{str(exp).translate(SUP)}"

# ---------------------------
# Helpers
# ---------------------------
def read_kv_tsv(path: str) -> Dict[str, str]:
    """Read a 2‑column TSV as a dict: key \t value."""
    d: Dict[str, str] = {}
    with open(path, "r") as f:
        for ln in f:
            if not ln.strip() or ln.startswith("#"):
                continue
            parts = ln.rstrip("\n").split("\t")
            key = parts[0]
            val = parts[1] if len(parts) > 1 else ""
            d[key] = val
    return d

def parse_sample_from_name(fname: str) -> str:
    """Use NNUEFS_(S#)_ label if present; else fallback to filename stem."""
    m = re.search(r"NNUEFS_(S\d+)_", fname)
    return m.group(1) if m else Path(fname).stem

def num0(v, default=0.0) -> float:
    """Robust numeric cast with fallback to default."""
    if v is None or v == "":
        return float(default)
    try:
        return float(int(v))
    except Exception:
        try:
            return float(v)
        except Exception:
            return float(default)

def common_stem(strings: List[str]) -> str:
    """Common prefix across stems; trimmed of trailing separators."""
    if not strings:
        return "group"
    prefix = os.path.commonprefix([Path(s).stem for s in strings])
    prefix = re.sub(r"[_\-]+$", "", prefix)
    return prefix or "group"

def timestamp_now() -> str:
    from datetime import datetime
    return datetime.now().strftime("%Y%m%d_%H%M%S")

def build_auto_outdir(files: List[str]) -> Path:
    stem = common_stem(files)
    ts = timestamp_now()
    base = Path(files[0]).resolve().parent if files else Path.cwd()
    return base / f"qc_{stem}_{ts}"

def wrap_paths_for_textbox(paths: List[str], max_chars=92) -> str:
    lines = ["Inputs:"]
    for p in paths:
        absp = str(Path(p).resolve())
        for chunk in textwrap.wrap(absp, width=max_chars):
            lines.append(chunk)
    return "\n".join(lines)

def add_input_textbox(fig, inputs_text: str):
    # Bottom‑right, outside plotting areas
    fig.text(0.995, 0.02, inputs_text, ha="right", va="bottom",
             fontsize=8, color="black",
             bbox=dict(boxstyle="round,pad=0.35", fc="white",
                       ec="#666666", lw=0.8, alpha=0.9))

def get_complexity_from_kv(kv: dict) -> float:
    """Preferred key, then fallbacks that include 'complex'."""
    preferred = [
        "summary/complexity_naive",
        "summary/complexity",
        "summary/complexity_ml",
        "complexity",
        "library_complexity",
    ]
    for key in preferred:
        if key in kv and kv[key]:
            try:
                val = float(kv[key])
                if val > 0:
                    return val
            except Exception:
                pass
    for k, v in kv.items():
        if "complex" in k.lower():
            try:
                val = float(v)
                if val > 0:
                    return val
            except Exception:
                continue
    return float("nan")

# ---------------------------
# Data loader
# ---------------------------
def load_dedup_stats(paths: List[str]) -> pd.DataFrame:
    rows = []
    for p in paths:
        kv = read_kv_tsv(p)
        sample = parse_sample_from_name(os.path.basename(p))

        # Pair types
        UU = num0(kv.get("pair_types/UU"))
        Ux = sum(num0(kv.get(k)) for k in ["pair_types/NU","pair_types/UN","pair_types/MU","pair_types/UM"])
        MN = sum(num0(kv.get(k)) for k in ["pair_types/MM","pair_types/MN","pair_types/NM","pair_types/NN"])

        # Duplicates
        dups   = num0(kv.get("total_dups"))
        nodups = num0(kv.get("total_nodups"))
        dup_pct = 100.0 * dups / (dups + nodups) if (dups + nodups) > 0 else float("nan")

        # Complexity
        complexity = get_complexity_from_kv(kv)

        # cis/trans (post‑dedup bins)
        cis   = num0(kv.get("cis"))
        cis1k = num0(kv.get("cis_1kb+")) if kv.get("cis_1kb+") is not None else num0(kv.get("cis_1kbp+"))
        cis20 = num0(kv.get("cis_20kb+"))
        trans = num0(kv.get("trans"))
        lt1k = cis - cis1k
        mid  = cis1k - cis20
        gt20 = cis20
        total = lt1k + mid + gt20 + trans

        tot_pairtypes = UU + Ux + MN
        UU_pct   = 100.0 * UU / tot_pairtypes if tot_pairtypes>0 else float("nan")
        UxMN_pct = 100.0 * Ux / tot_pairtypes if tot_pairtypes>0 else float("nan")
        MN_pct   = 100.0 * MN / tot_pairtypes if tot_pairtypes>0 else float("nan")

        rows.append({
            "sample": sample, "path": p,
            "UU_n": UU, "UxMN_n": Ux, "MNonly_n": MN, "pairtypes_total": tot_pairtypes,
            "UU_pct": UU_pct, "UxMN_pct": UxMN_pct, "MNonly_pct": MN_pct,
            "dups": dups, "nodups": nodups, "dup_pct": dup_pct,
            "complexity": complexity,
            "cis_lt1kb_n": lt1k, "cis_1_20kb_n": mid, "cis_gt20kb_n": gt20, "trans_n": trans,
            "cistrans_total": total
        })

    df = pd.DataFrame(rows)

    # Sort by S# if present
    def _k(s):
        m = re.search(r"S(\d+)", s)
        return int(m.group(1)) if m else 1_000_000
    df = df.sort_values("sample", key=lambda c: c.map(_k)).reset_index(drop=True)
    return df

# ---------------------------
# Plot helpers
# ---------------------------
def style_ax(ax):
    ax.tick_params(axis="both", which="major",
                   length=MAJOR_TICK_LEN, labelsize=TICK_LABEL_FS,
                   width=AX_LINEWIDTH, direction="out")
    for sp in ("top","right"):
        ax.spines[sp].set_visible(False)
    for sp in ("left","bottom"):
        ax.spines[sp].set_linewidth(AX_LINEWIDTH)
    ax.grid(False)
    ax.yaxis.grid(False)

# ---------------------------
# Plot: pair types stacked
# ---------------------------
def plot_pairtypes(df: pd.DataFrame, out_png: str, out_pdf: str):
    sdf = df.copy()
    n = len(sdf); x_tr = np.arange(n); width = 0.90
    fig, ax = plt.subplots(figsize=(6.0, 7.2), constrained_layout=False)
    bottom = np.zeros(n)
    for key, col in [("UU_pct",COL_UU), ("UxMN_pct",COL_UxMN), ("MNonly_pct",COL_MN)]:
        vals = np.nan_to_num(sdf[key].to_numpy(), nan=0.0)
        ax.bar(x_tr, vals, width=width, bottom=bottom, color=col, linewidth=0)
        bottom += vals

    # Pooled merged bar
    merged_counts = sdf[["UU_n","UxMN_n","MNonly_n"]].sum(skipna=True)
    m_tot = merged_counts.sum()
    m_pct = [100*merged_counts["UU_n"]/m_tot, 100*merged_counts["UxMN_n"]/m_tot, 100*merged_counts["MNonly_n"]/m_tot]
    x_m = n + 1.6; bottom_m = 0.0
    for val, col in zip(m_pct, [COL_UU,COL_UxMN,COL_MN]):
        ax.bar([x_m], [val], width=width, bottom=[bottom_m], color=col, linewidth=0)
        bottom_m += val
    # side labels on merged
    y0=0.0
    for val in m_pct:
        yc = y0 + val/2
        if val>0.2:
            ax.text(x_m + width/2 + 0.20, yc, f"{val:.1f}%", ha="left", va="center", fontsize=12)
        y0 += val

    xticks = list(np.arange(n)) + [x_m]
    ax.set_xticks(xticks)
    labs = sdf["sample"].tolist() + ["Merged"]
    xt = ax.set_xticklabels(labs, fontsize=TICK_LABEL_FS)
    for i,t in enumerate(xt):
        if i < n:
            t.set_rotation(45); t.set_ha("right")
        else:
            t.set_fontweight("bold")
    ax.set_xlim(-0.6, x_m + 1.8); ax.set_ylim(0, 100)
    ax.set_ylabel("Composition (%)", fontsize=AX_LABEL_FS)
    ax.set_title("UU / U×(M|N) / (M|N)×(M|N) (weighted 'Merged')", fontsize=AX_LABEL_FS, pad=8)
    style_ax(ax)

    handles = [plt.Rectangle((0,0),1,1,color=c) for c in (COL_UU,COL_UxMN,COL_MN)]
    labels  = ["UU","U×(M|N)","(M|N)×(M|N)"]
    ax.legend(handles, labels, loc="center left", bbox_to_anchor=(1.02,0.5),
              frameon=False, prop={"size":LEGEND_FS})

    try:
        add_input_textbox(fig, wrap_paths_for_textbox(df["path"].tolist()))
    except Exception:
        pass

    plt.subplots_adjust(left=0.12, right=0.84, top=0.96, bottom=0.18)
    plt.savefig(out_png, dpi=200); plt.savefig(out_pdf); plt.close(fig)

# ---------------------------
# Plot: duplicates (%)
# ---------------------------
def plot_dups(df: pd.DataFrame, out_png: str, out_pdf: str):
    sdf = df.copy()
    w = 100.0 * sdf["dups"].sum() / (sdf["dups"].sum()+sdf["nodups"].sum())
    n = len(sdf); x = np.linspace(-0.06, 0.06, n) if n>1 else np.array([0.0])

    fig, ax = plt.subplots(figsize=(2.6, 7.2))
    ax.scatter(x, sdf["dup_pct"], s=DOT_SIZE, marker="o", linewidths=0,
               edgecolors="none", color=DEEP_RED)
    ax.hlines(w, xmin=-0.20, xmax=0.20, linewidth=3.0, color=DEEP_RED)
    ax.text(1.02, w, f"{w:.2f}%", va="center", ha="left",
            fontsize=14, transform=ax.get_yaxis_transform(), clip_on=False)
    ax.set_ylim(0,100); ax.set_xlim(-0.35,0.60)
    ax.set_ylabel("% duplicates", fontsize=AX_LABEL_FS); ax.set_xticks([])
    style_ax(ax)
    for sp in ("top","right","bottom"): ax.spines[sp].set_visible(False)

    try:
        add_input_textbox(fig, wrap_paths_for_textbox(df["path"].tolist()))
    except Exception:
        pass

    plt.subplots_adjust(left=0.22, right=0.82, top=0.98, bottom=0.06)
    plt.savefig(out_png, dpi=200); plt.savefig(out_pdf); plt.close(fig)

# ---------------------------
# Plot: complexity (log10 axis)
# ---------------------------
def plot_complexity(df: pd.DataFrame, out_png: str, out_pdf: str):
    sdf = df.copy()
    vals = sdf["complexity"].astype(float).to_numpy()
    vals = vals[np.isfinite(vals) & (vals > 0)]
    fig, ax = plt.subplots(figsize=(2.8, 7.2))

    if len(vals) == 0:
        ax.text(0.5, 0.5, "No complexity values in stats",
                ha="center", va="center", fontsize=13, transform=ax.transAxes)
        ax.set_axis_off()
        try:
            add_input_textbox(fig, wrap_paths_for_textbox(df["path"].tolist()))
        except Exception:
            pass
        plt.savefig(out_png, dpi=200); plt.savefig(out_pdf); plt.close(fig)
        return

    mean_c = float(np.mean(vals))
    n = len(vals); x = np.linspace(-0.06, 0.06, n) if n>1 else np.array([0.0])

    cmin, cmax = float(np.min(vals)), float(np.max(vals))
    lo_pow = int(np.floor(np.log10(max(1.0, cmin))))
    hi_pow = int(np.ceil(np.log10(cmax)))
    if hi_pow <= lo_pow: hi_pow = lo_pow + 1
    yticks = [10.0**k for k in range(lo_pow, hi_pow+1)]

    ax.scatter(x, vals, s=DOT_SIZE, marker="o", linewidths=0,
               edgecolors="none", color=DEEP_RED)
    ax.hlines(mean_c, xmin=-0.20, xmax=0.20, linewidth=3.0, color=DEEP_RED)
    ax.text(1.02, mean_c, sci_txt(mean_c),
            transform=ax.get_yaxis_transform(), va="center", ha="left",
            fontsize=16, clip_on=False)

    ax.set_yscale("log"); ax.set_ylim(10.0**lo_pow, 10.0**hi_pow)
    ax.set_yticks(yticks)
    ax.set_yticklabels([to_sup(k) for k in range(lo_pow, hi_pow+1)],
                       fontsize=TICK_LABEL_FS)
    ax.tick_params(axis="y", which="major", length=MAJOR_TICK_LEN,
                   width=AX_LINEWIDTH, direction="out")
    ax.tick_params(axis="y", which="minor", length=0)
    ax.set_xlim(-0.45, 0.90); ax.set_xticks([])
    ax.set_ylabel("Estimated complexity — log10 (unique pairs)",
                  fontsize=AX_LABEL_FS)
    for sp in ("top","right","bottom"): ax.spines[sp].set_visible(False)
    ax.spines["left"].set_linewidth(AX_LINEWIDTH)

    try:
        add_input_textbox(fig, wrap_paths_for_textbox(df["path"].tolist()))
    except Exception:
        pass

    plt.subplots_adjust(left=0.52, right=0.80, top=0.98, bottom=0.06)
    plt.savefig(out_png, dpi=200); plt.savefig(out_pdf); plt.close(fig)

# ---------------------------
# Plot: cis distance strata + trans (stacked)
# ---------------------------
def plot_cistrans(df: pd.DataFrame, out_png: str, out_pdf: str):
    sdf = df.copy()
    n = len(sdf); x_tr = np.arange(n); width=0.90
    fig, ax = plt.subplots(figsize=(6.0, 7.2), constrained_layout=False)
    bottom = np.zeros(n)

    for key, col, lab in [("cis_lt1kb_n",COL_LT1,"<1 kb (cis)"),
                          ("cis_1_20kb_n",COL_1_20,"1–20 kb (cis)"),
                          ("cis_gt20kb_n",COL_GT20,">20 kb (cis)"),
                          ("trans_n",COL_TRANS,"trans")]:
        pct = 100.0 * np.nan_to_num(sdf[key].to_numpy(), nan=0.0) / \
              np.maximum(1.0, np.nan_to_num(sdf["cistrans_total"].to_numpy(), nan=0.0))
        ax.bar(x_tr, pct, width=width, bottom=bottom, color=col,
               align="center", linewidth=0, label=lab)
        bottom += pct

    # Merged
    counts = sdf[["cis_lt1kb_n","cis_1_20kb_n","cis_gt20kb_n","trans_n"]].sum(skipna=True)
    m_tot = counts.sum()
    x_m = n + 1.6; bottom_m = 0.0
    for key, col in zip(["cis_lt1kb_n","cis_1_20kb_n","cis_gt20kb_n","trans_n"],
                        [COL_LT1,COL_1_20,COL_GT20,COL_TRANS]):
        val = float(100.0 * counts[key] / m_tot) if m_tot>0 else 0.0
        ax.bar([x_m], [val], width=width, bottom=[bottom_m], color=col, linewidth=0)
        bottom_m += val

    # Side labels on merged
    y0=0.0
    for key in ["cis_lt1kb_n","cis_1_20kb_n","cis_gt20kb_n","trans_n"]:
        val = float(100.0 * counts[key] / m_tot) if m_tot>0 else 0.0
        yc=(y0+val/2.0)
        if val>0.2:
            ax.text(x_m + width/2 + 0.20, yc, f"{val:.1f}%", ha="left", va="center", fontsize=12)
        y0 += val

    xticks = list(np.arange(n)) + [x_m]
    ax.set_xticks(xticks)
    labs = sdf["sample"].tolist() + ["Merged"]
    xt = ax.set_xticklabels(labs, fontsize=TICK_LABEL_FS)
    for i,t in enumerate(xt):
        if i < n: t.set_rotation(45); t.set_ha("right")
        else: t.set_fontweight("bold")
    ax.set_xlim(-0.6, x_m + 1.8); ax.set_ylim(0, 100)
    ax.set_ylabel("Composition (%)", fontsize=AX_LABEL_FS)
    ax.set_title("cis distance strata + trans (weighted 'Merged')",
                 fontsize=AX_LABEL_FS, pad=8)
    style_ax(ax)

    ax.legend(loc="center left", bbox_to_anchor=(1.02,0.5),
              frameon=False, prop={"size":LEGEND_FS})

    try:
        add_input_textbox(fig, wrap_paths_for_textbox(df["path"].tolist()))
    except Exception:
        pass

    plt.subplots_adjust(left=0.12, right=0.84, top=0.96, bottom=0.18)
    plt.savefig(out_png, dpi=200); plt.savefig(out_pdf); plt.close(fig)

# ---------------------------
# Plot: pies (area ∝ UU)
# ---------------------------
def plot_pies(df: pd.DataFrame, out_png: str, out_pdf: str):
    sdf = df.copy()
    # Append pooled “Merged” pseudo-row
    pooled = {
        "sample": "Merged",
        "UU_n": float(sdf["UU_n"].sum()),
        "cis_lt1kb_n": float(sdf["cis_lt1kb_n"].sum()),
        "cis_1_20kb_n": float(sdf["cis_1_20kb_n"].sum()),
        "cis_gt20kb_n": float(sdf["cis_gt20kb_n"].sum()),
        "trans_n": float(sdf["trans_n"].sum()),
        "cistrans_total": float(sdf["cistrans_total"].sum()),
        "path": ""  # skip for textbox
    }
    sdf = pd.concat([sdf, pd.DataFrame([pooled])], ignore_index=True)

    n = len(sdf)
    uu_vals = sdf["UU_n"].fillna(0).to_numpy(float)
    uu_max  = max(uu_vals.max(), 1.0)
    R_MAX   = 0.58
    radii   = R_MAX * np.sqrt(uu_vals / uu_max)

    fig = plt.figure(constrained_layout=True, figsize=(1.9*(n+2.6), 6.6))
    gs  = fig.add_gridspec(2, n+2, height_ratios=[4.6, 1.5], width_ratios=[1]*n+[0.9, 1.7])

    # Top pies
    for i in range(n):
        ax = fig.add_subplot(gs[0,i])
        row = sdf.iloc[i]
        fr = [row["cis_lt1kb_n"], row["cis_1_20kb_n"], row["cis_gt20kb_n"], row["trans_n"]]
        tot = sum(v for v in fr if pd.notna(v))
        fr = [float(v)/tot if tot>0 else 0 for v in fr]
        ax.pie(fr, radius=radii[i],
               colors=[COL_LT1,COL_1_20,COL_GT20,COL_TRANS],
               startangle=90, counterclock=False)
        ax.set(aspect="equal"); ax.set_xticks([]); ax.set_yticks([])
        for sp in ax.spines.values(): sp.set_visible(False)
        ax.set_title(row.get("sample",""), fontsize=14, pad=6,
                     fontweight=("bold" if row.get("sample","")=="Merged" else "normal"))
        UUval = int(row["UU_n"]) if pd.notna(row["UU_n"]) else 0
        ax.text(0.5, -0.13, f"UU = {UUval:,}", transform=ax.transAxes,
                ha="center", va="top", fontsize=11)

    # Legend column
    leg_ax = fig.add_subplot(gs[0,n]); leg_ax.axis("off")
    for j,(lab,col) in enumerate([("<1 kb (cis)",COL_LT1),
                                  ("1–20 kb (cis)",COL_1_20),
                                  (">20 kb (cis)",COL_GT20),
                                  ("trans",COL_TRANS)]):
        y = 0.95 - j*0.18
        leg_ax.text(0.0, y, u"\u25A0", color=col, fontsize=16, va="center")
        leg_ax.text(0.10, y, lab, fontsize=12, va="center")

    # Pooled merged annotation column
    ann_ax = fig.add_subplot(gs[0,n+1]); ann_ax.axis("off")
    m = sdf.iloc[-1]; m_tot = float(m["cistrans_total"]) if pd.notna(m["cistrans_total"]) else 0.0
    def pct(n): return 100.0*n/m_tot if m_tot>0 else float("nan")
    pairs = [("<1 kb (cis)", m["cis_lt1kb_n"], pct(m["cis_lt1kb_n"]), COL_LT1),
             ("1–20 kb (cis)", m["cis_1_20kb_n"], pct(m["cis_1_20kb_n"]), COL_1_20),
             (">20 kb (cis)", m["cis_gt20kb_n"], pct(m["cis_gt20kb_n"]), COL_GT20),
             ("trans", m["trans_n"], pct(m["trans_n"]), COL_TRANS)]
    ann_ax.text(0.0, 1.00, "Merged (pooled)", fontsize=14, fontweight="bold", va="top")
    y = 0.86
    for lab,cnt,p, col in pairs:
        ann_ax.text(0.02, y, u"\u25A0", color=col, fontsize=14, va="center")
        ann_ax.text(0.10, y, f"{int(cnt):,}  ({p:.1f}%)", fontsize=12, va="center")
        y -= 0.18
    ann_ax.text(0.02, y-0.06, f"Total = {int(m_tot):,}", fontsize=12, va="top")

    # Size legend
    scale_ax = fig.add_subplot(gs[1,:])
    scale_ax.set_xlim(0,1); scale_ax.set_ylim(0,1)
    scale_ax.set_aspect("equal"); scale_ax.axis("off")
    k = R_MAX/np.sqrt(uu_max)
    ticks = [10_000_000, 50_000_000, 100_000_000, 500_000_000, 1_000_000_000]
    labs  = ["10M","50M","100M","500M","1B"]
    xs    = np.linspace(0.10, 0.90, len(ticks))
    for x, val, lab in zip(xs, ticks, labs):
        r = min(k*np.sqrt(val), 0.20)
        scale_ax.add_patch(Circle((x, 0.64), r, transform=scale_ax.transData,
                                  facecolor="#bdbdbd", edgecolor="none"))
        scale_ax.text(x, 0.20, lab, ha="center", va="top", fontsize=12,
                      transform=scale_ax.transData)

    try:
        paths = [p for p in df["path"].tolist() if p]
        add_input_textbox(fig, wrap_paths_for_textbox(paths))
    except Exception:
        pass

    fig.suptitle("Contact composition (dedup) — pie area ∝ UU (unique reads)",
                 fontsize=16, y=0.995)
    plt.savefig(out_png, dpi=200); plt.savefig(out_pdf); plt.close(fig)

# ---------------------------
# Stitchers (PNG preview + Vector PDF)
# ---------------------------
def stitch_row4_png(pngs: List[str], out_png: str):
    """Raster preview: paste PNGs side‑by‑side (kept only for convenience)."""
    if not PIL_OK or len(pngs)!=4:
        return
    imgs = [Image.open(p).convert("RGBA") for p in pngs]
    target_h = max(im.height for im in imgs)
    rs = []
    for im in imgs:
        if im.height != target_h:
            new_w = int(im.width * (target_h / im.height))
            im = im.resize((new_w, target_h), resample=Image.Resampling.BICUBIC)
        rs.append(im)
    total_w = sum(im.width for im in rs)
    combo = Image.new("RGBA", (total_w, target_h), (255,255,255,0))
    xoff = 0
    for im in rs:
        combo.paste(im, (xoff, 0)); xoff += im.width
    combo.convert("RGB").save(out_png, dpi=(200,200))

def stitch_row4_pdf_vector(pdf_paths: List[str], out_pdf: str):
    """Vector stitcher: merges first page of each PDF into one row **without** rasterizing."""
    try:
        from PyPDF2 import PdfReader, PdfWriter, Transformation
    except Exception:
        return  # PyPDF2 not available; skip vector combo

    if len(pdf_paths) != 4:
        return

    readers = [PdfReader(p) for p in pdf_paths]
    pages   = [r.pages[0] for r in readers]

    widths  = [float(pg.mediabox.right - pg.mediabox.left) for pg in pages]
    heights = [float(pg.mediabox.top - pg.mediabox.bottom) for pg in pages]
    target_h = max(heights)
    scales    = [target_h / h for h in heights]
    scaled_w  = [w*s for w,s in zip(widths, scales)]
    total_w   = sum(scaled_w)

    writer = PdfWriter()
    new_page = writer.add_blank_page(width=total_w, height=target_h)

    x = 0.0
    for pg, s, sw in zip(pages, scales, scaled_w):
        t = Transformation().scale(sx=s, sy=s).translate(tx=x, ty=0.0)
        new_page.merge_transformed_page(pg, t)
        x += sw

    writer.add_page(new_page)
    with open(out_pdf, "wb") as f:
        writer.write(f)

# ---------------------------
# CSV exports
# ---------------------------
def export_summary_tables(df: pd.DataFrame, outdir: Path, prefix: str, checksum: bool):
    csum = df[["sample","UU_n","cistrans_total","cis_lt1kb_n",
               "cis_1_20kb_n","cis_gt20kb_n","trans_n"]].copy()
    for coln, colp in [("cis_lt1kb_n","cis_lt1kb_pct"),
                       ("cis_1_20kb_n","cis_1_20kb_pct"),
                       ("cis_gt20kb_n","cis_gt20kb_pct"),
                       ("trans_n","trans_pct")]:
        csum[colp] = 100.0 * csum[coln] / csum["cistrans_total"]
    csum.to_csv(outdir / f"{prefix}_cistrans_summary.csv", index=False)

    master = df[[
        "sample","path",
        "UU_n","UxMN_n","MNonly_n","pairtypes_total","UU_pct","UxMN_pct","MNonly_pct",
        "dups","nodups","dup_pct",
        "complexity",
        "cistrans_total","cis_lt1kb_n","cis_1_20kb_n","cis_gt20kb_n","trans_n"
    ]].copy()
    if checksum:
        def sha(p):
            try:
                with open(p,"rb") as fh: return hashlib.sha256(fh.read()).hexdigest()
            except Exception: return ""
        master["sha256"] = master["path"].apply(sha)
    else:
        master["sha256"] = ""
    master.to_csv(outdir / f"{prefix}_master.csv", index=False)

    prov = master[["sample","path","sha256"]].copy()
    prov["filesize_bytes"] = prov["path"].apply(lambda p: os.path.getsize(p) if os.path.exists(p) else "")
    prov.to_csv(outdir / f"{prefix}_provenance.csv", index=False)

# ---------------------------
# Main
# ---------------------------
def main():
    ap = argparse.ArgumentParser(description="QC plots & tables for pairtools dedup stats.")
    ap.add_argument("--input-glob", default="")
    ap.add_argument("--inputs", action="append", help="add one or more input TSV paths; can repeat", default=[])
    ap.add_argument("--inputs-file", help="text file with one TSV path per line", default=None)
    ap.add_argument("--outdir", default=None)
    ap.add_argument("--prefix", default="QC")
    ap.add_argument("--no-pairtypes", action="store_true")
    ap.add_argument("--no-cistrans", action="store_true")
    ap.add_argument("--no-dups", action="store_true")
    ap.add_argument("--no-complexity", action="store_true")
    ap.add_argument("--no-pies", action="store_true")
    ap.add_argument("--no-row", action="store_true", help="skip stitched 4‑wide row panel")
    ap.add_argument("--checksum", action="store_true", help="include SHA‑256 in provenance/master CSVs")
    ap.add_argument("--zip", action="store_true", help="zip the output directory at the end")
    args = ap.parse_args()

    files = []
    if args.input_glob:
        files.extend(glob.glob(args.input_glob))
    files.extend(args.inputs)
    if args.inputs_file:
        with open(args.inputs_file, "r") as f:
            files.extend([ln.strip() for ln in f if ln.strip()])
    files = sorted(set(files))
    if not files:
        print("No input files found. Use --input-glob or --inputs/--inputs-file.", file=sys.stderr)
        sys.exit(2)

    outdir = Path(args.outdir) if args.outdir else build_auto_outdir(files)
    outdir.mkdir(parents=True, exist_ok=True)

    df = load_dedup_stats(files)
    export_summary_tables(df, outdir, args.prefix, checksum=args.checksum)

    # Plot paths
    p_pair = outdir / f"{args.prefix}_pairtypes_stacked.png"
    p_dup  = outdir / f"{args.prefix}_dups_dots.png"
    p_cx   = outdir / f"{args.prefix}_complexity_log10.png"
    p_ct   = outdir / f"{args.prefix}_cistrans_stacked.png"
    p_pies = outdir / f"{args.prefix}_pies_areaUU.png"

    # Generate panels
    if not args.no_pairtypes:
        plot_pairtypes(df, str(p_pair), str(p_pair.with_suffix(".pdf")))
    if not args.no_dups:
        plot_dups(df, str(p_dup), str(p_dup.with_suffix(".pdf")))
    if not args.no_complexity:
        plot_complexity(df, str(p_cx), str(p_cx.with_suffix(".pdf")))
    if not args.no_cistrans:
        plot_cistrans(df, str(p_ct), str(p_ct.with_suffix(".pdf")))
    if not args.no_pies:
        plot_pies(df, str(p_pies), str(p_pies.with_suffix(".pdf")))

    # Vector + PNG stitched row
    if not args.no_row and not any([args.no_pairtypes, args.no_dups, args.no_complexity, args.no_cistrans]):
        stitch_row4_png([str(p_pair), str(p_dup), str(p_cx), str(p_ct)],
                        str(outdir / f"{args.prefix}_row4_combined.png"))
        stitch_row4_pdf_vector([str(p_pair.with_suffix(".pdf")),
                                str(p_dup.with_suffix(".pdf")),
                                str(p_cx.with_suffix(".pdf")),
                                str(p_ct.with_suffix(".pdf"))],
                               str(outdir / f"{args.prefix}_row4_combined.pdf"))

    if args.zip:
        zip_path_base = str(outdir)
        shutil.make_archive(zip_path_base, 'zip', root_dir=outdir)
        print("Zipped:", f"{zip_path_base}.zip")

    print("Done. Outputs in", outdir)

if __name__ == "__main__":
    main()
```
