## Supplementary material for "CAD-C: An engineered nuclease enables repair-free *in situ* proximity ligation and nucleosome-resolution chromosome walks in human cells": Custom code: submit_Ps_dedup_GM12878deepmerge_RPE1P2andP3_K562E3_v2_8CPUs_20251110.sh

```
#!/bin/bash
#SBATCH --job-name=PsCurve_CADC_3cells
#SBATCH --output=PsCurve_CADC_3cells_%j.out
#SBATCH --error=PsCurve_CADC_3cells_%j.err
#SBATCH --cpus-per-task=8   # threads; no time/mem per request

# (Optional) activate your Open2C env if needed:
# source activate open2c_v3

# ---------- MCOOL paths (from your logs) ----------
export MCOOL_GM12878="/rugpfs/fs0/risc_lab/scratch/jsoroczyns/CAD/CAD-C/CADC_GM12878_deepmerge_20250205/cooler_out_20250205_023558/GM12878_deepmerge_20250205_011600_dedup_500_bal_500N_igdiag1.mcool"
export MCOOL_RPE1="/rugpfs/fs0/risc_lab/scratch/jsoroczyns/CAD/CAD-C/CADC_E1_P2andP3_merge/20241125_post_merge_rededup/20241127_rededup_igdiag1_coolers/CADC_E1_P2andP3_20241126_dedup_merged_rededup_500_bal_500N_igdiag1.mcool"
export MCOOL_K562="/rugpfs/fs0/risc_lab/scratch/jsoroczyns/CAD/CAD-C/CADC_E2_P2_E3_Novogene/FASTQS/CADC_E3_K562_BR1/20240822_pairtools_pipe_out/results_20240822/CADC_K562_1_manual_merge_20240917/merged_20240917/cool_output/CADC_K562_1_merged_dedup_20240822_500_bal_500N_igdiag1.mcool"

# Fail early if any path is missing
for f in "$MCOOL_GM12878" "$MCOOL_RPE1" "$MCOOL_K562"; do
  if [[ ! -f "$f" ]]; then
    echo "[ERROR] Missing file: $f" >&2
    exit 1
  fi
done

# ---------- Run P(s) v2 (overlay all three) ----------
python /rugpfs/fs0/risc_lab/scratch/jsoroczyns/CAD/CAD-C/scripts/2025_04_16/mcool_Ps_curve_v2_20250416.py \
  --mcool  "$MCOOL_GM12878" \
           "$MCOOL_RPE1" \
           "$MCOOL_K562" \
  --aliases \
    "CAD-C GM12878 E2 (Classic+Express+Direct dedup)" \
    "CAD-C RPE1 E1 (Classic, dedup)" \
    "CAD-C K562 E3 (Classic, dedup)" \
  --binsize 500 \
  --threads "${SLURM_CPUS_PER_TASK:-8}" \
  --output-dir /rugpfs/fs0/risc_lab/scratch/jsoroczyns/CAD/CAD-C/Ps_curves_2025/
```
