## Supplementary material for "CAD-C: An engineered nuclease enables repair-free *in situ* proximity ligation and nucleosome-resolution chromosome walks in human cells": Custom code: Fig2h_loop_pileup_RPE1.sh

```
#!/bin/bash
#SBATCH --job-name=Pileup
#SBATCH --output=Pileup_%j.log
#SBATCH --cpus-per-task=14
#SBATCH --mem=64G
#SBATCH --time=24:00:00
#SBATCH --partition=hpc_l40_a

#conda activate coolpup
source ~/.bashrc_soft2
conda activate open2c_v3

# the first batch of the downsampled data
RPE1_CAD='/rugpfs/fs0/risc_lab/scratch/wzuo/project/CAD/RPE1_dataset/subsample/RPE1_CADC_500_igdig1_1b.mcool'
RPE1_MicroC='/rugpfs/fs0/risc_lab/scratch/wzuo/project/CAD/RPE1_dataset/subsample/RPE1_microc_500_igdig1_1b.mcool'

RPE1_CADC_MicroC_5kb='/rugpfs/fs0/risc_lab/scratch/wzuo/project/CAD/RPE1_dataset/subsample/loops/mustache/RPE1_1b_2500/RPE1_CADC_MicroC_2500_loop_5kb_slop2.bedpe'
RPE1_CADC_only_5kb='/rugpfs/fs0/risc_lab/scratch/wzuo/project/CAD/RPE1_dataset/subsample/loops/mustache/RPE1_1b_2500/RPE1_CADC_only_2500_loop_5kb_slop2.bedpe'
RPE1_MicroC_only_5kb='/rugpfs/fs0/risc_lab/scratch/wzuo/project/CAD/RPE1_dataset/subsample/loops/mustache/RPE1_1b_2500/RPE1_MicroC_only_2500_loop_5kb_slop2.bedpe'

python /rugpfs/fs0/risc_lab/scratch/wzuo/CAD/loops_subgroup/loops_pileup/pileup_wrapa_coolpuppy_v13_20250929.py -c ${RPE1_CAD} ${RPE1_MicroC} -r ${RPE1_MicroC_only_5kb} ${RPE1_CADC_MicroC_5kb} ${RPE1_CADC_only_5kb} \
    -o RPE1_CAD_MicroC_2500_loop_5kb_slop_pileup --binsize 1000 --flank 40000 --cooler-labels "RPE1_CAD" "RPE1_MicroC" --region-labels "MicroC_only" "Common" "CAD_only" --threads 14
```
