## Supplementary material for "CAD-C: An engineered nuclease enables repair-free *in situ* proximity ligation and nucleosome-resolution chromosome walks in human cells": Custom code: Supplementary_Fig3a_RPE_loop_anchor_pileup.sh

```
#!/bin/bash
#SBATCH --job-name=HiC_Processing
#SBATCH --output=hic_processing_%j.log
#SBATCH --cpus-per-task=12
#SBATCH --mem=32G
#SBATCH --time=24:00:00
#SBATCH --partition=hpc_l40_a

source /programs/biogrids.shrc
cores=14

input_dir="/rugpfs/fs0/risc_lab/scratch/wzuo/project/CAD/RPE1_dataset/subsample/loops/mustache/RPE1_1b_2500"
output_dir="/rugpfs/fs0/risc_lab/scratch/wzuo/project/CAD/RPE1_dataset/subsample/loops/pile"

RPE1_both="$input_dir/RPE1_CADC_MicroC_2500_loop_5kb_slop.bed"
RPE1_CO="$input_dir/RPE1_CADC_only_2500_loop_5kb_slop.bed"
RPE1_MO="$input_dir/RPE1_MicroC_only_2500_loop_5kb_slop.bed"

RPE1_ATAC='/rugpfs/fs0/risc_lab/scratch/wzuo/project/CAD/dataset/bw/RPE1_ATAC.bw'
RPE1_CTCF_vr='/lustre/fs4/risc_lab/scratch/aou/cad_microc/Epigenetics/CTCF/RPE1_WT2_CnT_CTCF.filt.bw'
RPE1_H3K27Ac_vr='/rugpfs/fs0/risc_lab/scratch/wzuo/project/CAD/dataset/bw/RPE1_H3K27Ac.bw'
RPE1_H3K27me3_vr='/rugpfs/fs0/risc_lab/scratch/wzuo/project/CAD/dataset/bw/RPE1_H3K27me3.bw'
RPE1_H3K4me1_vr='/rugpfs/fs0/risc_lab/scratch/wzuo/project/CAD/dataset/bw/RPE1_H3K4me1.bw'
RPE1_H3K4me3_vr='/rugpfs/fs0/risc_lab/scratch/wzuo/project/CAD/dataset/bw/RPE1_H3K4me3.bw'
RPE1_Pol2ser5='/rugpfs/fs0/risc_lab/scratch/wzuo/project/CAD/dataset/bw/RPE1_Pol2ser5.bw'
RPE1_YY1='/rugpfs/fs0/risc_lab/scratch/wzuo/project/CAD/dataset/bw/RPE1_YY1.bw'
RPE1_CTCF='/rugpfs/fs0/risc_lab/scratch/wzuo/project/CAD/dataset/bw/GSE196727_RAW/GSM5899605_2017_034B_Ct_CTCF-bowtie2-hg38-raw-GR.100.bw'
RPE1_Rad21='/rugpfs/fs0/risc_lab/scratch/wzuo/project/CAD/dataset/bw/GSE196727_RAW/GSM5899607_2017_034B_Ct_Rad21-bowtie2-hg38-raw-GR.100.bw'
RPE1_CTCF2='/rugpfs/fs0/risc_lab/scratch/wzuo/project/CAD/dataset/bw/GSE196727_RAW/GSM5899621_2018_014A_015B_Ct_CTCF-bowtie2-hg38-raw-GR.100.bw'
RPE1_Rad212='/rugpfs/fs0/risc_lab/scratch/wzuo/project/CAD/dataset/bw/GSE196727_RAW/GSM5899617_2018_014A_Ct_Rad21-bowtie2-hg38-raw-GR.100.bw'
RPE1_BRD4m='/rugpfs/fs0/risc_lab/scratch/wzuo/project/CAD/dataset/bw/GSE196727_RAW/GSM5899623_2019_004B_JQ1_minus_BRD4_IP-bowtie2-hg38-raw-GR.100.bw'
RPE1_BRD4p='/rugpfs/fs0/risc_lab/scratch/wzuo/project/CAD/dataset/bw/GSE196727_RAW/GSM5899625_2019_004B_JQ1_plus_BRD4_IP-bowtie2-hg38-raw-GR.100.bw'
RPE1_H3K27ac='/rugpfs/fs0/risc_lab/scratch/wzuo/project/CAD/dataset/bw/GSE196727_RAW/GSM5899629_2021_010_H3K27ac_IP_Control-bowtie2-hg38-raw-GR.100.bw'
RPE1_H3K36me3='/rugpfs/fs0/risc_lab/scratch/wzuo/project/CAD/dataset/bw/GSE196727_RAW/GSM5899632_2021_010_H3K36me3_IP_Control-bowtie2-hg38-raw-GR.100.bw'
RPE1_H3K4me3='/rugpfs/fs0/risc_lab/scratch/wzuo/project/CAD/dataset/bw/GSE196727_RAW/GSM5899635_2021_010_H3K4me3_IP_Control-bowtie2-hg38-raw-GR.100.bw'
RPE1_H3K27me3='/rugpfs/fs0/risc_lab/scratch/wzuo/project/CAD/dataset/bw/GSE196727_RAW/GSM5899641_2021_023_H3K27me3_ab_IP_Control-bowtie2-hg38-raw-GR.100.bw'

computeMatrix reference-point --referencePoint center -S ${RPE1_ATAC} -R ${RPE1_MO} ${RPE1_both} ${RPE1_CO} -o $output_dir/RPE1_ATAC_pile.mat.gz -b 25000 -a 25000 -bs 100 --skipZeros -p $cores
plotHeatmap -m "$output_dir/RPE1_ATAC_pile.mat.gz" -out "$output_dir/RPE1_ATAC_pile.heatmap.pdf" --perGroup --colorMap "inferno" --sortUsing sum --dpi 300 -T "" --startLabel ""  --endLabel "" --legendLocation "upper-right" --heatmapWidth 12 
plotProfile -m "$output_dir/RPE1_ATAC_pile.mat.gz" -out "$output_dir/RPE1_ATAC_pile.profile.pdf" --perGroup --dpi 300 -T "" --startLabel ""  --endLabel "" --legendLocation "upper-right" --plotHeight 10 --plotWidth 12

computeMatrix reference-point --referencePoint center -S ${RPE1_CTCF_vr} -R ${RPE1_MO} ${RPE1_both} ${RPE1_CO} -o $output_dir/RPE1_CTCF_pile0.mat.gz -b 25000 -a 25000 -bs 100 --skipZeros -p $cores
plotHeatmap -m "$output_dir/RPE1_CTCF_pile0.mat.gz" -out "$output_dir/RPE1_CTCF_pile0.heatmap.pdf" --perGroup --colorMap "inferno" --sortUsing sum --dpi 300 -T "" --startLabel ""  --endLabel "" --legendLocation "upper-right" --heatmapWidth 12 
plotProfile -m "$output_dir/RPE1_CTCF_pile0.mat.gz" -out "$output_dir/RPE1_CTCF_pile0.profile.pdf" --perGroup --dpi 300 -T "" --startLabel ""  --endLabel "" --legendLocation "upper-right" --plotHeight 10 --plotWidth 12

computeMatrix reference-point --referencePoint center -S ${RPE1_H3K27Ac_vr} -R ${RPE1_MO} ${RPE1_both} ${RPE1_CO} -o $output_dir/RPE1_H3K27Ac_pile0.mat.gz -b 25000 -a 25000 -bs 100 --skipZeros -p $cores
plotHeatmap -m "$output_dir/RPE1_H3K27Ac_pile0.mat.gz" -out "$output_dir/RPE1_H3K27Ac_pile0.heatmap.pdf" --perGroup --colorMap "inferno" --sortUsing sum --dpi 300 -T "" --startLabel ""  --endLabel "" --legendLocation "upper-right" --heatmapWidth 12 
plotProfile -m "$output_dir/RPE1_H3K27Ac_pile0.mat.gz" -out "$output_dir/RPE1_H3K27Ac_pile0.profile.pdf" --perGroup --dpi 300 -T "" --startLabel ""  --endLabel "" --legendLocation "upper-right" --plotHeight 10 --plotWidth 12

computeMatrix reference-point --referencePoint center -S ${RPE1_H3K27me3_vr} -R ${RPE1_MO} ${RPE1_both} ${RPE1_CO} -o $output_dir/RPE1_H3K27me3_pile0.mat.gz -b 25000 -a 25000 -bs 100 --skipZeros -p $cores
plotHeatmap -m "$output_dir/RPE1_H3K27me3_pile0.mat.gz" -out "$output_dir/RPE1_H3K27me3_pile0.heatmap.pdf" --perGroup --colorMap "inferno" --sortUsing sum --dpi 300 -T "" --startLabel ""  --endLabel "" --legendLocation "upper-right" --heatmapWidth 12 
plotProfile -m "$output_dir/RPE1_H3K27me3_pile0.mat.gz" -out "$output_dir/RPE1_H3K27me3_pile0.profile.pdf" --perGroup --dpi 300 -T "" --startLabel ""  --endLabel "" --legendLocation "upper-right" --plotHeight 10 --plotWidth 12

computeMatrix reference-point --referencePoint center -S ${RPE1_H3K4me1_vr} -R ${RPE1_MO} ${RPE1_both} ${RPE1_CO} -o $output_dir/RPE1_H3K4me1_pile0.mat.gz -b 25000 -a 25000 -bs 100 --skipZeros -p $cores
plotHeatmap -m "$output_dir/RPE1_H3K4me1_pile0.mat.gz" -out "$output_dir/RPE1_H3K4me1_pile0.heatmap.pdf" --perGroup --colorMap "inferno" --sortUsing sum --dpi 300 -T "" --startLabel ""  --endLabel "" --legendLocation "upper-right" --heatmapWidth 12 
plotProfile -m "$output_dir/RPE1_H3K4me1_pile0.mat.gz" -out "$output_dir/RPE1_H3K4me1_pile0.profile.pdf" --perGroup --dpi 300 -T "" --startLabel ""  --endLabel "" --legendLocation "upper-right" --plotHeight 10 --plotWidth 12

computeMatrix reference-point --referencePoint center -S ${RPE1_H3K4me3_vr} -R ${RPE1_MO} ${RPE1_both} ${RPE1_CO} -o $output_dir/RPE1_H3K4me3_pile0.mat.gz -b 25000 -a 25000 -bs 100 --skipZeros -p $cores
plotHeatmap -m "$output_dir/RPE1_H3K4me3_pile0.mat.gz" -out "$output_dir/RPE1_H3K4me3_pile0.heatmap.pdf" --perGroup --colorMap "inferno" --sortUsing sum --dpi 300 -T "" --startLabel ""  --endLabel "" --legendLocation "upper-right" --heatmapWidth 12 
plotProfile -m "$output_dir/RPE1_H3K4me3_pile0.mat.gz" -out "$output_dir/RPE1_H3K4me3_pile0.profile.pdf" --perGroup --dpi 300 -T "" --startLabel ""  --endLabel "" --legendLocation "upper-right" --plotHeight 10 --plotWidth 12

computeMatrix reference-point --referencePoint center -S ${RPE1_Pol2ser5} -R ${RPE1_MO} ${RPE1_both} ${RPE1_CO} -o $output_dir/RPE1_Pol2ser5_pile.mat.gz -b 25000 -a 25000 -bs 100 --skipZeros -p $cores
plotHeatmap -m "$output_dir/RPE1_Pol2ser5_pile.mat.gz" -out "$output_dir/RPE1_Pol2ser5_pile.heatmap.pdf" --perGroup --colorMap "inferno" --sortUsing sum --dpi 300 -T "" --startLabel ""  --endLabel "" --legendLocation "upper-right" --heatmapWidth 12 
plotProfile -m "$output_dir/RPE1_Pol2ser5_pile.mat.gz" -out "$output_dir/RPE1_Pol2ser5_pile.profile.pdf" --perGroup --dpi 300 -T "" --startLabel ""  --endLabel "" --legendLocation "upper-right" --plotHeight 10 --plotWidth 12

computeMatrix reference-point --referencePoint center -S ${RPE1_YY1} -R ${RPE1_MO} ${RPE1_both} ${RPE1_CO} -o $output_dir/RPE1_YY1_pile.mat.gz -b 25000 -a 25000 -bs 100 --skipZeros -p $cores
plotHeatmap -m "$output_dir/RPE1_YY1_pile.mat.gz" -out "$output_dir/RPE1_YY1_pile.heatmap.pdf" --perGroup --colorMap "inferno" --sortUsing sum --dpi 300 -T "" --startLabel ""  --endLabel "" --legendLocation "upper-right" --heatmapWidth 12 
plotProfile -m "$output_dir/RPE1_YY1_pile.mat.gz" -out "$output_dir/RPE1_YY1_pile.profile.pdf" --perGroup --dpi 300 -T "" --startLabel ""  --endLabel "" --legendLocation "upper-right" --plotHeight 10 --plotWidth 12

computeMatrix reference-point --referencePoint center -S ${RPE1_CTCF} -R ${RPE1_MO} ${RPE1_both} ${RPE1_CO} -o $output_dir/RPE1_CTCF_pile.mat.gz -b 25000 -a 25000 -bs 100 --skipZeros -p $cores
plotHeatmap -m "$output_dir/RPE1_CTCF_pile.mat.gz" -out "$output_dir/RPE1_CTCF_pile.heatmap.pdf" --perGroup --colorMap "inferno" --sortUsing sum --dpi 300 -T "" --startLabel ""  --endLabel "" --legendLocation "upper-right" --heatmapWidth 12 
plotProfile -m "$output_dir/RPE1_CTCF_pile.mat.gz" -out "$output_dir/RPE1_CTCF_pile.profile.pdf" --perGroup --dpi 300 -T "" --startLabel ""  --endLabel "" --legendLocation "upper-right" --plotHeight 10 --plotWidth 12

computeMatrix reference-point --referencePoint center -S ${RPE1_Rad21} -R ${RPE1_MO} ${RPE1_both} ${RPE1_CO} -o $output_dir/RPE1_Rad21_pile.mat.gz -b 25000 -a 25000 -bs 100 --skipZeros -p $cores
plotHeatmap -m "$output_dir/RPE1_Rad21_pile.mat.gz" -out "$output_dir/RPE1_Rad21_pile.heatmap.pdf" --perGroup --colorMap "inferno" --sortUsing sum --dpi 300 -T "" --startLabel ""  --endLabel "" --legendLocation "upper-right" --heatmapWidth 12 
plotProfile -m "$output_dir/RPE1_Rad21_pile.mat.gz" -out "$output_dir/RPE1_Rad21_pile.profile.pdf" --perGroup --dpi 300 -T "" --startLabel ""  --endLabel "" --legendLocation "upper-right" --plotHeight 10 --plotWidth 12

computeMatrix reference-point --referencePoint center -S ${RPE1_CTCF2} -R ${RPE1_MO} ${RPE1_both} ${RPE1_CO} -o $output_dir/RPE1_CTCF2_pile.mat.gz -b 25000 -a 25000 -bs 100 --skipZeros -p $cores
plotHeatmap -m "$output_dir/RPE1_CTCF2_pile.mat.gz" -out "$output_dir/RPE1_CTCF2_pile.heatmap.pdf" --perGroup --colorMap "inferno" --sortUsing sum --dpi 300 -T "" --startLabel ""  --endLabel "" --legendLocation "upper-right" --heatmapWidth 12 
plotProfile -m "$output_dir/RPE1_CTCF2_pile.mat.gz" -out "$output_dir/RPE1_CTCF2_pile.profile.pdf" --perGroup --dpi 300 -T "" --startLabel ""  --endLabel "" --legendLocation "upper-right" --plotHeight 10 --plotWidth 12

computeMatrix reference-point --referencePoint center -S ${RPE1_Rad212} -R ${RPE1_MO} ${RPE1_both} ${RPE1_CO} -o $output_dir/RPE1_Rad212_pile.mat.gz -b 25000 -a 25000 -bs 100 --skipZeros -p $cores
plotHeatmap -m "$output_dir/RPE1_Rad212_pile.mat.gz" -out "$output_dir/RPE1_Rad212_pile.heatmap.pdf" --perGroup --colorMap "inferno" --sortUsing sum --dpi 300 -T "" --startLabel ""  --endLabel "" --legendLocation "upper-right" --heatmapWidth 12 
plotProfile -m "$output_dir/RPE1_Rad212_pile.mat.gz" -out "$output_dir/RPE1_Rad212_pile.profile.pdf" --perGroup --dpi 300 -T "" --startLabel ""  --endLabel "" --legendLocation "upper-right" --plotHeight 10 --plotWidth 12

computeMatrix reference-point --referencePoint center -S ${RPE1_BRD4m} -R ${RPE1_MO} ${RPE1_both} ${RPE1_CO} -o $output_dir/RPE1_BRD4m_pile.mat.gz -b 25000 -a 25000 -bs 100 --skipZeros -p $cores
plotHeatmap -m "$output_dir/RPE1_BRD4m_pile.mat.gz" -out "$output_dir/RPE1_BRD4m_pile.heatmap.pdf" --perGroup --colorMap "inferno" --sortUsing sum --dpi 300 -T "" --startLabel ""  --endLabel "" --legendLocation "upper-right" --heatmapWidth 12 
plotProfile -m "$output_dir/RPE1_BRD4m_pile.mat.gz" -out "$output_dir/RPE1_BRD4m_pile.profile.pdf" --perGroup --dpi 300 -T "" --startLabel ""  --endLabel "" --legendLocation "upper-right" --plotHeight 10 --plotWidth 12

computeMatrix reference-point --referencePoint center -S ${RPE1_BRD4p} -R ${RPE1_MO} ${RPE1_both} ${RPE1_CO} -o $output_dir/RPE1_BRD4p_pile.mat.gz -b 25000 -a 25000 -bs 100 --skipZeros -p $cores
plotHeatmap -m "$output_dir/RPE1_BRD4p_pile.mat.gz" -out "$output_dir/RPE1_BRD4p_pile.heatmap.pdf" --perGroup --colorMap "inferno" --sortUsing sum --dpi 300 -T "" --startLabel ""  --endLabel "" --legendLocation "upper-right" --heatmapWidth 12 
plotProfile -m "$output_dir/RPE1_BRD4p_pile.mat.gz" -out "$output_dir/RPE1_BRD4p_pile.profile.pdf" --perGroup --dpi 300 -T "" --startLabel ""  --endLabel "" --legendLocation "upper-right" --plotHeight 10 --plotWidth 12

computeMatrix reference-point --referencePoint center -S ${RPE1_H3K27ac} -R ${RPE1_MO} ${RPE1_both} ${RPE1_CO} -o $output_dir/RPE1_H3K27ac_pile.mat.gz -b 25000 -a 25000 -bs 100 --skipZeros -p $cores
plotHeatmap -m "$output_dir/RPE1_H3K27ac_pile.mat.gz" -out "$output_dir/RPE1_H3K27ac_pile.heatmap.pdf" --perGroup --colorMap "inferno" --sortUsing sum --dpi 300 -T "" --startLabel ""  --endLabel "" --legendLocation "upper-right" --heatmapWidth 12 
plotProfile -m "$output_dir/RPE1_H3K27ac_pile.mat.gz" -out "$output_dir/RPE1_H3K27ac_pile.profile.pdf" --perGroup --dpi 300 -T "" --startLabel ""  --endLabel "" --legendLocation "upper-right" --plotHeight 10 --plotWidth 12

computeMatrix reference-point --referencePoint center -S ${RPE1_H3K36me3} -R ${RPE1_MO} ${RPE1_both} ${RPE1_CO} -o $output_dir/RPE1_H3K36me3_pile.mat.gz -b 25000 -a 25000 -bs 100 --skipZeros -p $cores
plotHeatmap -m "$output_dir/RPE1_H3K36me3_pile.mat.gz" -out "$output_dir/RPE1_H3K36me3_pile.heatmap.pdf" --perGroup --colorMap "inferno" --sortUsing sum --dpi 300 -T "" --startLabel ""  --endLabel "" --legendLocation "upper-right" --heatmapWidth 12 
plotProfile -m "$output_dir/RPE1_H3K36me3_pile.mat.gz" -out "$output_dir/RPE1_H3K36me3_pile.profile.pdf" --perGroup --dpi 300 -T "" --startLabel ""  --endLabel "" --legendLocation "upper-right" --plotHeight 10 --plotWidth 12

computeMatrix reference-point --referencePoint center -S ${RPE1_H3K4me3} -R ${RPE1_MO} ${RPE1_both} ${RPE1_CO} -o $output_dir/RPE1_H3K4me3_pile.mat.gz -b 25000 -a 25000 -bs 100 --skipZeros -p $cores
plotHeatmap -m "$output_dir/RPE1_H3K4me3_pile.mat.gz" -out "$output_dir/RPE1_H3K4me3_pile.heatmap.pdf" --perGroup --colorMap "inferno" --sortUsing sum --dpi 300 -T "" --startLabel ""  --endLabel "" --legendLocation "upper-right" --heatmapWidth 12 
plotProfile -m "$output_dir/RPE1_H3K4me3_pile.mat.gz" -out "$output_dir/RPE1_H3K4me3_pile.profile.pdf" --perGroup --dpi 300 -T "" --startLabel ""  --endLabel "" --legendLocation "upper-right" --plotHeight 10 --plotWidth 12

computeMatrix reference-point --referencePoint center -S ${RPE1_H3K27me3} -R ${RPE1_MO} ${RPE1_both} ${RPE1_CO} -o $output_dir/RPE1_H3K27me3_pile.mat.gz -b 25000 -a 25000 -bs 100 --skipZeros -p $cores
plotHeatmap -m "$output_dir/RPE1_H3K27me3_pile.mat.gz" -out "$output_dir/RPE1_H3K27me3_pile.heatmap.pdf" --perGroup --colorMap "inferno" --sortUsing sum --dpi 300 -T "" --startLabel ""  --endLabel "" --legendLocation "upper-right" --heatmapWidth 12 
plotProfile -m "$output_dir/RPE1_H3K27me3_pile.mat.gz" -out "$output_dir/RPE1_H3K27me3_pile.profile.pdf" --perGroup --dpi 300 -T "" --startLabel ""  --endLabel "" --legendLocation "upper-right" --plotHeight 10 --plotWidth 12
```
