## Supplementary material for "CAD-C: An engineered nuclease enables repair-free *in situ* proximity ligation and nucleosome-resolution chromosome walks in human cells": Custom code: submit_Dmax_empirical_vs_resample_v2_20251021.sh

```
#!/bin/bash
#SBATCH -J dmax_v9
#SBATCH -o %x-%j.out
#SBATCH -e %x-%j.err

set -euo pipefail

# Path to your v9 NPZ (as provided)
NPZ="/rugpfs/fs0/risc_lab/scratch/jsoroczyns/CAD/CADwalk/CADwalk_E1_GM12878/bwamem2_out_20250408/local_parse2_out/analyses_20251020/cadpack_bc1001_bc1002_v9_14422039.npz"
SCRIPT="/rugpfs/fs0/risc_lab/scratch/jsoroczyns/CAD/CAD-C/scripts/2025_10_21/Dmax_empirical_vs_signedstep_resampling_null_Lstratified_v2_20251021.py"   # adjust if saved elsewhere

OUTDIR="${SCRATCH:-$PWD}/dmax_v9_$(date +%Y%m%d_%H%M%S)"
mkdir -p "$OUTDIR"

NULL_REPS=150000
FRAG_QLO=0.01
FRAG_QHI=0.99
LOG_MIN=2
LOG_MAX=9
BINS=60
SEED=13

echo "python: $(command -v python)"
python --version || true
echo "NPZ: $NPZ"
echo "OUTDIR: $OUTDIR"

# --- L=3, midpoints (primary) ---
OUTP="${OUTDIR}/v9_L3_mids"
srun python "$SCRIPT" \
  --npz "$NPZ" --L 3 --geometry mids \
  --null-reps "$NULL_REPS" \
  --frag-qlo "$FRAG_QLO" --frag-qhi "$FRAG_QHI" \
  --log-min "$LOG_MIN" --log-max "$LOG_MAX" --bins "$BINS" \
  --seed "$SEED" \
  --outprefix "$OUTP"

# --- L=4, midpoints (optional example) ---
OUTP="${OUTDIR}/v9_L4_mids"
srun python "$SCRIPT" \
  --npz "$NPZ" --L 4 --geometry mids \
  --null-reps "$NULL_REPS" \
  --frag-qlo "$FRAG_QLO" --frag-qhi "$FRAG_QHI" \
  --log-min "$LOG_MIN" --log-max "$LOG_MAX" --bins "$BINS" \
  --seed "$SEED" \
  --outprefix "$OUTP"

# --- Junctions (optional; uncomment if you want) ---
# OUTP="${OUTDIR}/v9_L3_junc"
# srun python "$SCRIPT" \
#   --npz "$NPZ" --L 3 --geometry junctions \
#   --null-reps "$NULL_REPS" \
#   --frag-qlo "$FRAG_QLO" --frag-qhi "$FRAG_QHI" \
#   --log-min "$LOG_MIN" --log-max "$LOG_MAX" --bins "$BINS" \
#   --seed "$SEED" \
#   --outprefix "$OUTP"

echo "Artifacts:"
ls -lh "$OUTDIR"/*
```
