## Supplementary material for "CAD-C: An engineered nuclease enables repair-free *in situ* proximity ligation and nucleosome-resolution chromosome walks in human cells": Custom code: datanote.rtf

Supp figure 5A plots the simulated splines used to pick synthetic data vs the real cadwalk data.
The code used to plot is within the script simulate_splines_AB_BC_ACpdfaccept_2.r

##this section below is the plotting for support figure 5A 
sample_log10dist_AB <- AB$sampler
sample_log10dist_BC <- BC$sampler
pdfAC <- AC$pdf_func
max_pdfAC <- AC$max_pdf

AB_plot <- ggplot() +
  geom_line(data = AB$dens_full_df, aes(x = x, y = y), color = "blue") +
  geom_smooth(data = AB$dens_sub_df, aes(x = x, y = y), method = "gam", formula = y ~ s(x, bs = "cr", k = 20), se = FALSE, color = "red") +
  theme_minimal() + labs(x = "Log10(AB_distance)", y = "Density")
BC_plot <- ggplot() +
  geom_line(data = BC$dens_full_df, aes(x = x, y = y), color = "blue") +
  geom_smooth(data = BC$dens_sub_df, aes(x = x, y = y), method = "gam", formula = y ~ s(x, bs = "cr", k = 20), se = FALSE, color = "red") +
  theme_minimal() + labs(x = "Log10(BC_distance)", y = "Density")
AC_plot <- ggplot() +
  geom_line(data = AC$dens_full_df, aes(x = x, y = y), color = "blue") +
  geom_smooth(data = AC$dens_sub_df, aes(x = x, y = y), method = "gam", formula = y ~ s(x, bs = "cr", k = 20), se = FALSE, color = "red") +
  theme_minimal() + labs(x = "Log10(AC_distance)", y = "Density")

combo_plot <- AB_plot / BC_plot / AC_plot
ggsave(splines_pdf, combo_plot, width = 7, height = 12)
