## Supplementary material for "CAD-C: An engineered nuclease enables repair-free *in situ* proximity ligation and nucleosome-resolution chromosome walks in human cells": Custom code: simulate_splines_AB_BC_ACpdfaccept_2.r

```
#!/usr/bin/env Rscript

# Usage:
#   Rscript simulate_splines_AB_AC_anyAnchor_acceptReject_BC.R <input_tsv> <output_dir>

suppressPackageStartupMessages({
  library(dplyr)
  library(ggplot2)
  library(mgcv)
  library(data.table)
  library(patchwork)
})

args <- commandArgs(trailingOnly = TRUE)
if (length(args) < 2) {
  stop("Usage: Rscript simulate_splines_AB_AC_anyAnchor_acceptReject_BC.R <input_tsv> <output_dir>")
}

input_file <- args[1]
output_dir <- args[2]

# Create output directory if it doesn't exist
dir.create(output_dir, showWarnings = FALSE, recursive = TRUE)

file_base <- sub("\\.tsv$", "", basename(input_file))
synthetic_tsv   <- file.path(output_dir, paste0(file_base, "_synthetic.tsv"))
splines_pdf      <- file.path(output_dir, paste0(file_base, "_splines.pdf"))
dist_compare_pdf <- file.path(output_dir, paste0(file_base, "_distanceCompare.pdf"))

message("Input TSV:   ", input_file)
message("Output dir:  ", output_dir)
message("Synthetic:   ", synthetic_tsv)
message("Splines PDF: ", splines_pdf)
message("Compare PDF: ", dist_compare_pdf)

chrom_sizes_file <- "/rugpfs/fs0/risc_lab/store/risc_data/downloaded/hg38/genome/chrom.sizes"
chrom_df <- read.table(chrom_sizes_file, header=FALSE, sep="\t", col.names=c("chr", "length"))
chrom_lengths <- setNames(chrom_df$length, chrom_df$chr)
message("Loaded ", length(chrom_lengths), " chromosomes from hg38 chrom.sizes.")

place_fragment <- function(chrom, anchor_pos, dist_bp, max_tries=10) {
  if (!chrom %in% names(chrom_lengths)) return(NA)
  chr_len <- chrom_lengths[[chrom]]
  for (i in seq_len(max_tries)) {
    sign <- sample(c(-1, 1), 1)
    new_pos <- anchor_pos + sign * dist_bp
    if (new_pos >= 1 && new_pos <= chr_len) return(new_pos)
  }
  return(NA)
}

data <- fread(input_file, sep="\t", header=TRUE) %>% as_tibble()

required_cols <- c("A_chrom", "A_start", "B_chrom", "B_start", "C_chrom", "C_start",
                   "AB_distance", "BC_distance", "AC_distance")
missing_cols <- setdiff(required_cols, names(data))
if (length(missing_cols) > 0) stop("Missing required columns: ", paste(missing_cols, collapse=", "))

build_spline <- function(dist_vec, dist_name) {
  dist_full <- dist_vec[!is.na(dist_vec) & dist_vec > 0]
  if (length(dist_full) < 2) stop(paste("Not enough valid", dist_name))
  dens_full <- density(log10(dist_full))
  dens_sub <- density(log10(dist_full[dist_full >= 100]))
  sub_df <- data.frame(x = dens_sub$x, y = dens_sub$y) %>% filter(y > 0)
  fit <- mgcv::gam(y ~ s(x, bs = "cr", k = 20), data = sub_df)
  x_grid <- seq(min(sub_df$x), max(sub_df$x), length.out = 500)
  pred <- predict(fit, newdata = data.frame(x = x_grid), type = "response")
  pred[pred < 0] <- 0
  dx <- x_grid[2] - x_grid[1]
  area <- sum(pred * dx)
  pdf_norm <- pred / area
  cdf_vals <- cumsum(pdf_norm) * dx
  sampler <- function(n) {
    out <- numeric(n)
    for (i in seq_len(n)) {
      u <- runif(1)
      idx <- which.min(abs(cdf_vals - u))
      out[i] <- x_grid[idx]
    }
    out
  }
  pdf_func <- function(dist_bp) {
    if (dist_bp <= 0) return(0)
    lx <- log10(dist_bp)
    if (lx < x_grid[1] || lx > x_grid[length(x_grid)]) return(0)
    idx <- which.min(abs(x_grid - lx))
    pdf_norm[idx]
  }
  list(dens_full_df = data.frame(x = dens_full$x, y = dens_full$y),
       dens_sub_df = sub_df,
       fit = fit,
       sampler = sampler,
       pdf_func = pdf_func,
       max_pdf = max(pdf_norm))
}

AB <- build_spline(data$AB_distance, "AB_distance")
BC <- build_spline(data$BC_distance, "BC_distance")
AC <- build_spline(data$AC_distance, "AC_distance")

sample_log10dist_AB <- AB$sampler
sample_log10dist_BC <- BC$sampler
pdfAC <- AC$pdf_func
max_pdfAC <- AC$max_pdf

AB_plot <- ggplot() +
  geom_line(data = AB$dens_full_df, aes(x = x, y = y), color = "blue") +
  geom_smooth(data = AB$dens_sub_df, aes(x = x, y = y), method = "gam", formula = y ~ s(x, bs = "cr", k = 20), se = FALSE, color = "red") +
  theme_minimal() + labs(x = "Log10(AB_distance)", y = "Density")
BC_plot <- ggplot() +
  geom_line(data = BC$dens_full_df, aes(x = x, y = y), color = "blue") +
  geom_smooth(data = BC$dens_sub_df, aes(x = x, y = y), method = "gam", formula = y ~ s(x, bs = "cr", k = 20), se = FALSE, color = "red") +
  theme_minimal() + labs(x = "Log10(BC_distance)", y = "Density")
AC_plot <- ggplot() +
  geom_line(data = AC$dens_full_df, aes(x = x, y = y), color = "blue") +
  geom_smooth(data = AC$dens_sub_df, aes(x = x, y = y), method = "gam", formula = y ~ s(x, bs = "cr", k = 20), se = FALSE, color = "red") +
  theme_minimal() + labs(x = "Log10(AC_distance)", y = "Density")

combo_plot <- AB_plot / BC_plot / AC_plot
ggsave(splines_pdf, combo_plot, width = 7, height = 12)

sample_one_triple <- function(chrom_i, anchorA, max_pdfAC, max_tries = 50) {
  for (attempt in seq_len(max_tries)) {
    distAB <- 10^sample_log10dist_AB(1)
    distBC <- 10^sample_log10dist_BC(1)
    B_pos <- place_fragment(chrom_i, anchorA, distAB)
    if (is.na(B_pos)) next
    C_pos <- place_fragment(chrom_i, B_pos, distBC)
    if (is.na(C_pos)) next
    AC_implied <- abs(C_pos - anchorA)
    val <- pdfAC(AC_implied)
    if (val <= 0) next
    if (runif(1, 0, max_pdfAC) <= val) {
      return(data.frame(
        A_chrom = chrom_i,
        A_start = anchorA,
        B_chrom = chrom_i,
        B_start = B_pos,
        C_chrom = chrom_i,
        C_start = C_pos,
        AB_distance = distAB,
        BC_distance = distBC,
        AC_distance = AC_implied
      ))
    }
  }
  return(NULL)
}

anchor_pool <- bind_rows(
  data.frame(chrom = data$A_chrom, pos = data$A_start),
  data.frame(chrom = data$B_chrom, pos = data$B_start),
  data.frame(chrom = data$C_chrom, pos = data$C_start)
) %>% distinct()

N <- 1e7
chunk_size <- 1e6
total_chunks <- ceiling(N / chunk_size)
accepted_total <- 0

for (chunk_idx in seq_len(total_chunks)) {
  message("Processing chunk ", chunk_idx, " of ", total_chunks, "...")
  this_N <- ifelse(chunk_idx < total_chunks, chunk_size, N - chunk_size * (chunk_idx - 1))
  anchors <- anchor_pool %>% slice_sample(n = this_N, replace = TRUE)
  synthetic_list <- vector("list", this_N)
  accepted <- 0

  for (i in seq_len(this_N)) {
    anchor_chrom <- anchors$chrom[i]
    anchor_pos <- anchors$pos[i]
    triple_row <- sample_one_triple(anchor_chrom, anchor_pos, max_pdfAC)
    if (!is.null(triple_row)) {
      synthetic_list[[i]] <- triple_row
      accepted <- accepted + 1
    }
  }

  synthetic_df <- bind_rows(synthetic_list)
  if (nrow(synthetic_df) > 0) {
    write.table(
      synthetic_df,
      synthetic_tsv,
      sep = "\t",
      quote = FALSE,
      row.names = FALSE,
      col.names = !(file.exists(synthetic_tsv) && chunk_idx > 1),
      append = (chunk_idx > 1)
    )
  }
  accepted_total <- accepted_total + accepted
  message("Chunk ", chunk_idx, ": accepted ", accepted, " rows.")
}

message("Total accepted rows: ", accepted_total)

if (file.exists(synthetic_tsv)) {
  synthetic_df <- fread(synthetic_tsv)
  message("Comparing AB, BC, AC distributions in log10 space...")

  real_dists <- data.frame(
    dist_type = rep(c("AB","BC","AC"), each=nrow(data)),
    distance = c(data$AB_distance, data$BC_distance, data$AC_distance),
    dataset  = "real"
  )

  syn_dists <- data.frame(
    dist_type = rep(c("AB","BC","AC"), each=nrow(synthetic_df)),
    distance = c(synthetic_df$AB_distance, synthetic_df$BC_distance, synthetic_df$AC_distance),
    dataset  = "synthetic"
  )

  combined_dists <- bind_rows(real_dists, syn_dists) %>% filter(distance > 0)

  p2 <- ggplot(combined_dists, aes(x=log10(distance), color=dataset)) +
    geom_density(linewidth=1) +
    facet_wrap(~dist_type, scales="free") +
    theme_minimal() +
    labs(
      title="AB, BC, AC Distance Comparison: Real vs. Synthetic",
      x="log10(distance) (bp)",
      y="Density"
    )

  message("Saving distance comparison plot to: ", dist_compare_pdf)
  ggsave(dist_compare_pdf, p2, width=10, height=4)

  message("\n--- Spearman Correlation for Real vs. Synthetic ---\n")

  real_mat <- data %>% select(AB_distance, BC_distance, AC_distance)
  real_corr <- cor(real_mat, method="spearman", use="complete.obs")
  syn_mat <- synthetic_df %>% select(AB_distance, BC_distance, AC_distance)
  syn_corr <- cor(syn_mat, method="spearman", use="complete.obs")

  cat("\n--- REAL correlation matrix (Spearman) ---\n")
  print(real_corr)
  cat("\n--- SYNTHETIC correlation matrix (Spearman) ---\n")
  print(syn_corr)
}

message("All done.")
```
