## Supplementary material for "CAD-C: An engineered nuclease enables repair-free *in situ* proximity ligation and nucleosome-resolution chromosome walks in human cells": Custom code: datanote.rtf

Supp figure 5B top is from synthetic model 1 code — comparing real data to simulated by AB/BC/AC distance. simulate_splines_AB_BC.r
Supp figure 5B top is from synthetic model 2 code - comparing real data to simulated by AB/BC/AC distance
simulate_splines_AB_BC_ACpdfaccept_2.r


##Plotting section from both scripts for Supp figure 5B comparisons:
if (file.exists(synthetic_tsv)) {
  synthetic_df <- fread(synthetic_tsv)
  message("Comparing AB, BC, AC distributions in log10 space...")

  real_dists <- data.frame(
    dist_type = rep(c("AB","BC","AC"), each=nrow(data)),
    distance = c(data$AB_distance, data$BC_distance, data$AC_distance),
    dataset  = "real"
  )

  syn_dists <- data.frame(
    dist_type = rep(c("AB","BC","AC"), each=nrow(synthetic_df)),
    distance = c(synthetic_df$AB_distance, synthetic_df$BC_distance, synthetic_df$AC_distance),
    dataset  = "synthetic"
  )

  combined_dists <- bind_rows(real_dists, syn_dists) %>% filter(distance > 0)

  p2 <- ggplot(combined_dists, aes(x=log10(distance), color=dataset)) +
    geom_density(linewidth=1) +
    facet_wrap(~dist_type, scales="free") +
    theme_minimal() +
    labs(
      title="AB, BC, AC Distance Comparison: Real vs. Synthetic",
      x="log10(distance) (bp)",
      y="Density"
    )

  message("Saving distance comparison plot to: ", dist_compare_pdf)
  ggsave(dist_compare_pdf, p2, width=10, height=4)
