## Supplementary material for "CAD-C: An engineered nuclease enables repair-free *in situ* proximity ligation and nucleosome-resolution chromosome walks in human cells": Custom code: ABCspearmancorr.r

```
#!/usr/bin/env Rscript

suppressPackageStartupMessages({
  library(dplyr)
  library(data.table)  # for fread
})

# Get command-line arguments
args <- commandArgs(trailingOnly = TRUE)

# If no argument is given, read from standard input
if (length(args) < 1) {
  cat("No file argument provided. Reading data from standard input...\n")
  data <- fread("stdin", sep="\t", header=TRUE) %>% as_tibble()
} else {
  data_file <- args[1]
  cat("Data file: ", data_file, "\n")
  data <- fread(data_file, sep="\t", header=TRUE) %>% as_tibble()
}

# Select the relevant columns (assumes they exist in the file)
mat <- data %>% select(AB_distance, BC_distance, AC_distance)

# Compute Spearman correlation matrix (ignoring rows with NA)
corr <- cor(mat, method="spearman", use="complete.obs")

cat("\n--- Correlation matrix (Spearman) ---\n")
print(corr)
```
