## Supplementary material for "CAD-C: An engineered nuclease enables repair-free *in situ* proximity ligation and nucleosome-resolution chromosome walks in human cells": Custom code: data note.rtf

For supp. Figure 5C, the output from simulate_splines_AB_BC_ACpdfaccept_2.r prints spearman correlation for synthetic model 2. 
For synthetic model 1, I  used ABCspearmancorr.r and sumbit_ABCspearmancorr.sh
ABCspearmancoor.r created Superman correlation matrix of AB/BC/AC separations from input data. 
The plots were generated in chatgpt window by JS from the output matrices 

Output from both plots (underlying matrices) printed here:
 Model 2:
https://ood01-hpc.rockefeller.edu/pun/sys/dashboard/files/fs//ru-auth/local/home/landerson01/scratch/20250205_CAD_2x250bp_GM_analysis/Synthetic/AB_BC_Spline_AcceptonAC/syntheticdata_7376827_4294967294.out

--- REAL correlation matrix (Spearman) ---
            AB_distance BC_distance AC_distance
AB_distance   1.0000000   0.1666458   0.4244194
BC_distance   0.1666458   1.0000000   0.4231854
AC_distance   0.4244194   0.4231854   1.0000000

--- SYNTHETIC correlation matrix (Spearman) --- Model 2 
            AB_distance BC_distance AC_distance
AB_distance  1.00000000  0.09673914   0.5911147
BC_distance  0.09673914  1.00000000   0.6309320
AC_distance  0.59111469  0.63093198   1.0000000

Model 1:

https://ood01-hpc.rockefeller.edu/pun/sys/dashboard/files/fs//ru-auth/local/home/landerson01/scratch/20250205_CAD_2x250bp_GM_analysis/Synthetic/AB_BC_Spline/corrchecksyntheticdata_7306735_4294967294.out

Real data file:       /rugpfs/fs0/risc_lab/scratch/landerson01/20250205_CAD_2x250bp_GM_analysis/Compartment/merged_all_flipped_sorted.cis.compt_annotated.tsv.gz_distances_compartment.tsv 
Synthetic data file:  /lustre/fs4/risc_lab/scratch/landerson01/20250205_CAD_2x250bp_GM_analysis/Synthetic/AB_BC_Spline/merged_all_flipped_sorted.cis.compt_annotated.tsv.gz_distances_compartment_synthetic.tsv 
Reading real data...
Reading synthetic data...

--- REAL correlation matrix (Spearman) ---
            AB_distance BC_distance AC_distance
AB_distance   1.0000000   0.1619293   0.4274486
BC_distance   0.1619293   1.0000000   0.4258913
AC_distance   0.4274486   0.4258913   1.0000000

--- SYNTHETIC correlation matrix (Spearman) ---
            AB_distance BC_distance AC_distance
AB_distance 1.000000000 0.005681023   0.5562167
BC_distance 0.005681023 1.000000000   0.6108210
AC_distance 0.556216748 0.610821016   1.0000000

Generating PDF:  real_vs_synthetic_cor.pdf 
null device 
          1 

Done. Created  real_vs_synthetic_cor.pdf  with two pages.
