## Supplementary material for "CAD-C: An engineered nuclease enables repair-free *in situ* proximity ligation and nucleosome-resolution chromosome walks in human cells": Custom code: sumbit_ABCspearmancorr.sh

```
#!/bin/bash
#SBATCH --job-name=spearmancorr
#SBATCH --output=spearmancorr_%A_%a.out
#SBATCH --error=spearmancorr_%A_%a.err
#SBATCH -n 1
#SBATCH -c 8


# Check that two arguments were passed
if [ "$#" -ne 2 ]; then
  echo "Usage: sbatch submit_ABCspearmancorr.sh <input_file>"
  exit 1
fi

# Assign command-line arguments
INPUT_FILE=$1

# Activate Conda environment
source /ru-auth/local/home/risc_soft2/miniconda3/etc/profile.d/conda.sh
conda activate lauren


# Pass the arguments to the Python script
Rscript /ru-auth/local/home/landerson01/scripts/ABCspearmancorr.r "$1" "$2"
```
