## Supplementary material for "CAD-C: An engineered nuclease enables repair-free *in situ* proximity ligation and nucleosome-resolution chromosome walks in human cells": Custom code: TXT.rtf

Supplementary figure 5D is step 1 vs step 2 hex bin for real data vs synthetic, plotted with  step1vstep2_input_20250430.r

Real cadwalk data output: /ru-auth/local/home/landerson01/scratch/20250205_CAD_2x250bp_GM_analysis/Compartment/Cis/Hexbin_AB_vs_BC_Limit1e8/merged_all_flipped_sorted.cis.compt_annotated.tsv.gz_distances_compartment_hexbin_AB_vs_BC.pdf

Synthetic model 1 output: /ru-auth/local/home/landerson01/scratch/20250205_CAD_2x250bp_GM_analysis/Synthetic/AB_BC_Spline/Hexbin_AB_vs_BC_Limit1e8/merged_all_flipped_sorted.cis.compt_annotated.tsv.gz_distances_compartment_synthetic_hexbin_AB_vs_BC.pdf

Synthetic model 2 output: /ru-auth/local/home/landerson01/scratch/20250205_CAD_2x250bp_GM_analysis/Synthetic/AB_BC_Spline_AcceptonAC/Synthetic_100M/Hexbin_AB_vs_BC_Limit1e8/merged_all_flipped_sorted.cis.compt_annotated.tsv.gz_distances_compartment.tsv.gz_synthetic_hexbin_AB_vs_BC.pdf
