## Supplementary material for "CAD-C: An engineered nuclease enables repair-free *in situ* proximity ligation and nucleosome-resolution chromosome walks in human cells": Custom code: step1vstep2_input_20250430.r

```
#!/usr/bin/env Rscript

# ------------------- SETUP -------------------
suppressPackageStartupMessages({
  library(data.table)
  library(dplyr)
  library(ggplot2)
  library(viridis)
  library(stringr)
})

# ------------------- PARSE ARGUMENT -------------------
args <- commandArgs(trailingOnly = TRUE)
if (length(args) < 1) {
  stop("Usage: Rscript plot_hexbin_AB_BC.R <input_file>")
}
input_file <- args[1]
output_dir <- "Hexbin_AB_vs_BC_Limit1e8"
dir.create(output_dir, showWarnings = FALSE, recursive = TRUE)

# Derive label for file
file_label <- tools::file_path_sans_ext(basename(input_file))
file_label <- str_remove(file_label, "\\.tsv\\.gz$|\\.tsv$")

# ------------------- LOAD DATA -------------------
message("Reading input file: ", input_file)
data <- tryCatch({
  fread(input_file, sep = "\t", header = TRUE, data.table = FALSE)
}, error = function(e) {
  stop("Failed to read input file: ", e$message)
})

# ------------------- GLOBAL PLOT SETTINGS -------------------
x_limits <- c(1e2, 1e8)
y_limits <- c(1e2, 1e8)
fill_limits <- c(1e-8, 1e-3)

theme_set(theme_minimal(base_size = 12))

# ------------------- PLOTTING -------------------
message("Generating plot for: ", file_label)

plot <- ggplot(data, aes(x = AB_distance, y = BC_distance, weight = 1 / nrow(data))) +
  geom_hex(bins = 100) +
  scale_fill_viridis_c(option = "plasma", trans = "log10", limits = fill_limits, oob = scales::squish) +
  scale_x_log10(limits = x_limits) +
  scale_y_log10(limits = y_limits) +
  labs(
    title = paste("Hexbin of AB vs BC Distances:", file_label),
    x = "AB Distance (bp)",
    y = "BC Distance (bp)",
    fill = "Density (log)"
  )

# Save plot as PDF
output_file <- file.path(output_dir, paste0(file_label, "_hexbin_AB_vs_BC.pdf"))
ggsave(output_file, plot = plot, width = 6, height = 5, useDingbats = FALSE)

message("Saved plot to: ", output_file)
```
