## Supplementary material for "CAD-C: An engineered nuclease enables repair-free *in situ* proximity ligation and nucleosome-resolution chromosome walks in human cells": Custom code: submitstep1vsstep2.sh

```
#!/bin/bash
#SBATCH -n 1
#SBATCH -c 20
#SBATCH --verbose
#SBATCH -J hexbins
#SBATCH -e %x-%j-%a.err
#SBATCH -o %x-%j-%a.out

# Check if correct number of arguments is provided
if [ "$#" -ne 1 ]; then
    echo "Usage: sbatch submitstep1vsstep2.sh <input_tsv> "
    exit 1
fi

# Assign command-line arguments
INPUT_FILE=$1

# Activate Conda environment
source /ru-auth/local/home/risc_soft2/miniconda3/etc/profile.d/conda.sh
conda activate lauren

# Run R script with specified input and output directory
Rscript /ru-auth/local/home/landerson01/scripts/step1vstep2_input_20250430.r "$INPUT_FILE"
```
