## Supplementary material for "CAD-C: An engineered nuclease enables repair-free *in situ* proximity ligation and nucleosome-resolution chromosome walks in human cells": Custom code: data note.rtf

Data location: /ru-auth/local/home/landerson01/scratch/20250205_CAD_2x250bp_GM_analysis/Synthetic

These scripts were used to generate two synthetic null genome models of two-step walks
Synthetic model 1: simulate_splines_AB_BC.r
Synthetic model 2: simulate_splines_AB_BC_ACpdfaccept_2.r

These scripts were used to annotate the data by compartment (not used in this version of paper but could be useful in future)
LA_classify_ABC_frags_by_bed_list.py
submit_annotate_compt_synthetic.sh
