## Supplementary material for "CAD-C: An engineered nuclease enables repair-free *in situ* proximity ligation and nucleosome-resolution chromosome walks in human cells": Custom code: LA_classify_ABC_frags_by_bed_list.py

```
#!/usr/bin/env python3

"""
Classify the A/B/C fragment START coordinates from an ABC-like TSV file
by intersecting them against multiple non-overlapping BED files.

Supports two input formats:
  - 'full': 13-column ABC format with readID, chrom/start/end/strand for A/B/C (default)
  - 'simple': 9-column format with just chrom/start for A/B/C and distances

Usage:
  python classify_fragments.py \
    --abc input.tsv.gz \
    --bed-dir bed_files/ \
    --out output.tsv.gz \
    --chunk-size 500000 \
    --input-format simple
"""

import argparse
import sys
import csv
import gzip
import os
from pathlib import Path
from pybedtools import BedTool
from collections import defaultdict

def combine_bed_files(bed_files):
    all_beds = []
    for i, bf in enumerate(bed_files):
        label = f"bed_{i}"
        bt = BedTool(bf)
        labeled = bt.each(lambda x: x.fields[:3] + [label]).saveas()
        all_beds.append(labeled)
    combined = all_beds[0]
    for bed_part in all_beds[1:]:
        combined = combined.cat(bed_part, postmerge=False)
    return combined.saveas()

def process_chunk(chunk_rows, annotated_bed, out_f, frag_label_counts, input_format):
    bed_lines = []
    for idx, row in enumerate(chunk_rows):
        if input_format == "full":
            A_chrom, A_start = row[1], int(row[2])
            B_chrom, B_start = row[5], int(row[6])
            C_chrom, C_start = row[9], int(row[10])
        elif input_format == "simple":
            A_chrom, A_start = row[0], int(float(row[1]))
            B_chrom, B_start = row[2], int(float(row[3]))
            C_chrom, C_start = row[4], int(float(row[5]))
        else:
            raise ValueError(f"Unknown input format: {input_format}")

        bed_lines.append(f"{A_chrom}\t{A_start}\t{A_start+1}\t{idx}\tA")
        bed_lines.append(f"{B_chrom}\t{B_start}\t{B_start+1}\t{idx}\tB")
        bed_lines.append(f"{C_chrom}\t{C_start}\t{C_start+1}\t{idx}\tC")

    chunk_bed = BedTool("\n".join(bed_lines) + "\n", from_string=True)
    intersected = chunk_bed.intersect(annotated_bed, wa=True, wb=True)

    label_map = {}
    for x in intersected:
        row_idx = int(x[3])
        frag = x[4]
        bed_label = x[8]
        label_map[(row_idx, frag)] = bed_label

    for idx, row in enumerate(chunk_rows):
        A_label = label_map.get((idx, "A"), "NA")
        B_label = label_map.get((idx, "B"), "NA")
        C_label = label_map.get((idx, "C"), "NA")

        frag_label_counts[("A", A_label)] += A_label != "NA"
        frag_label_counts[("B", B_label)] += B_label != "NA"
        frag_label_counts[("C", C_label)] += C_label != "NA"

        out_f.write("\t".join(map(str, row + [A_label, B_label, C_label])) + "\n")

def main():
    parser = argparse.ArgumentParser()
    parser.add_argument("--abc", required=True, help="Input ABC file (.tsv or .tsv.gz)")
    parser.add_argument("--bed-dir", required=True, help="Directory with .bed files")
    parser.add_argument("--out", default=os.path.join(os.getcwd(), "annotated_output.tsv.gz"), help="Output file")
    parser.add_argument("--chunk-size", type=int, default=100_000, help="Lines per chunk")
    parser.add_argument("--input-format", choices=["full", "simple"], default="full", help="Input format type")
    args = parser.parse_args()

    abc_path = Path(args.abc)
    opener = gzip.open if abc_path.suffix == ".gz" else open
    bed_dir = Path(args.bed_dir)
    out_path = Path(args.out)

    if not abc_path.exists():
        sys.exit(f"ERROR: File not found: {abc_path}")
    if not bed_dir.is_dir():
        sys.exit(f"ERROR: Invalid bed directory: {bed_dir}")

    bed_files = sorted(bed_dir.glob("*.bed"))
    if not bed_files:
        sys.exit(f"ERROR: No BED files found in: {bed_dir}")

    try:
        annotated_bed = combine_bed_files(bed_files)
    except Exception as e:
        sys.exit(f"ERROR combining BED files: {e}")

    with (gzip.open(out_path, "wt") if out_path.suffix == ".gz" else open(out_path, "w")) as out_f:
        for i, bf in enumerate(bed_files):
            out_f.write(f"# bed_index={i} => {bf}\n")

        frag_label_counts = defaultdict(int)
        chunk_rows = []
        total_lines = 0

        try:
            with opener(abc_path, "rt") as in_f:
                reader = csv.reader(in_f, delimiter="\t")
                header = next(reader)
                if args.input_format == "full" and len(header) < 13:
                    raise ValueError(f"Expected 13+ columns in 'full' format, got: {header}")
                if args.input_format == "simple" and len(header) < 6:
                    raise ValueError(f"Expected 6+ columns in 'simple' format, got: {header}")

                out_f.write("\t".join(header + ["A_bed", "B_bed", "C_bed"]) + "\n")

                for row in reader:
                    chunk_rows.append(row)
                    if len(chunk_rows) >= args.chunk_size:
                        process_chunk(chunk_rows, annotated_bed, out_f, frag_label_counts, args.input_format)
                        total_lines += len(chunk_rows)
                        chunk_rows = []
                if chunk_rows:
                    process_chunk(chunk_rows, annotated_bed, out_f, frag_label_counts, args.input_format)
                    total_lines += len(chunk_rows)

        except Exception as e:
            sys.exit(f"ERROR while reading or processing {abc_path}: {e}")

    print(f"Done. Processed {total_lines} rows. Output written to: {out_path}")
    if frag_label_counts:
        print("\nFragment label usage:")
        label_stats = defaultdict(lambda: {"A": 0, "B": 0, "C": 0})
        for (frag, label), count in frag_label_counts.items():
            label_stats[label][frag] += count
        for label in sorted(label_stats):
            a, b, c = label_stats[label]["A"], label_stats[label]["B"], label_stats[label]["C"]
            print(f"  {label}: A={a}, B={b}, C={c}, total={a+b+c}")
    else:
        print("No intersections found.")

if __name__ == "__main__":
    main()
```
