## Supplementary material for "CAD-C: An engineered nuclease enables repair-free *in situ* proximity ligation and nucleosome-resolution chromosome walks in human cells": Custom code: simulate_splines_AB_BC.r

```
#!/usr/bin/env Rscript

# Usage:
#   Rscript simulate_splines_AB_BC_anyAnchor.R <input_tsv> <output_dir>
#
# This script:
#  1) Hardcodes the hg38 chrom.sizes path.
#  2) Reads an input TSV with columns:
#       A_chrom, A_start, B_chrom, B_start, C_chrom, C_start,
#       AB_distance, BC_distance, AC_distance
#  3) Builds two separate cubic spline distributions for:
#       - AB_distance >= 100
#       - BC_distance >= 100
#     using 20 knots for the cubic spline fit.
#  4) Allows the anchor pool to come from *any* of (A, B, C) positions.
#  5) For each synthetic row:
#       - Sample AB distance => place B from the anchor
#       - Sample BC distance => place C from B
#       - Compute AC_distance = |C - anchor|
#  6) Writes outputs to <output_dir>:
#       <basename>_synthetic.tsv
#       <basename>_splines.pdf   (AB, BC univariate fits)
#       <basename>_distanceCompare.pdf (AB, BC, AC real vs. synthetic)

suppressPackageStartupMessages({
  library(dplyr)
  library(ggplot2)
  library(mgcv)        # for cubic regression spline
  library(data.table)  # for fread if you want faster I/O
  library(patchwork)   # to combine AB/BC plots into one PDF
})

args <- commandArgs(trailingOnly = TRUE)
if (length(args) < 2) {
  stop("Usage: Rscript simulate_splines_AB_BC_anyAnchor.R <input_tsv> <output_dir>")
}

input_file <- args[1]
output_dir <- args[2]

# Create output directory if it doesn't exist
dir.create(output_dir, showWarnings = FALSE, recursive = TRUE)

# Derive a base name from input TSV
file_base <- sub("\\.tsv$", "", basename(input_file))

# Construct output paths
synthetic_tsv   <- file.path(output_dir, paste0(file_base, "_synthetic.tsv"))
splines_pdf      <- file.path(output_dir, paste0(file_base, "_splines.pdf"))
dist_compare_pdf <- file.path(output_dir, paste0(file_base, "_distanceCompare.pdf"))

message("Input TSV:   ", input_file)
message("Output dir:  ", output_dir)
message("Synthetic:   ", synthetic_tsv)
message("Splines PDF: ", splines_pdf)
message("Compare PDF: ", dist_compare_pdf)

#----------------------------------------------------------------------
# Hardcode the hg38 chrom.sizes path
#----------------------------------------------------------------------
chrom_sizes_file <- "/rugpfs/fs0/risc_lab/store/risc_data/downloaded/hg38/genome/chrom.sizes"
message("Reading chrom.sizes from: ", chrom_sizes_file)

chrom_df <- read.table(chrom_sizes_file, header=FALSE, sep="\t",
                       col.names=c("chr","length"))
chrom_lengths <- setNames(chrom_df$length, chrom_df$chr)
message("Loaded ", length(chrom_lengths), " chromosomes from hg38 chrom.sizes.")

# Minimal boundary check function
place_fragment <- function(chrom, anchor_pos, dist_bp, max_tries=10) {
  if (!chrom %in% names(chrom_lengths)) {
    return(NA)  # unknown chromosome
  }
  chr_len <- chrom_lengths[[chrom]]
  for (i in seq_len(max_tries)) {
    sign <- sample(c(-1, 1), 1)
    new_pos <- anchor_pos + sign * dist_bp
    if (new_pos >= 1 && new_pos <= chr_len) {
      return(new_pos)
    }
  }
  return(NA)
}

#----------------------------------------------------------------------
# 1) Read the input data
#----------------------------------------------------------------------
message("Reading data with fread from: ", input_file)
data <- fread(input_file, sep="\t", header=TRUE) %>% as_tibble()

required_cols <- c("A_chrom", "A_start", "B_chrom", "B_start", "C_chrom", "C_start",
                   "AB_distance", "BC_distance", "AC_distance")
missing_cols <- setdiff(required_cols, names(data))
if (length(missing_cols) > 0) {
  stop("Missing required columns: ", paste(missing_cols, collapse=", "))
}

#----------------------------------------------------------------------
# 2) Build two separate spline distributions: AB, BC
#    We also produce "full" densities for reference.
#----------------------------------------------------------------------
# ---- AB_distance ----
AB_dist_full <- data$AB_distance
AB_dist_full <- AB_dist_full[!is.na(AB_dist_full) & AB_dist_full > 0]
if (length(AB_dist_full) < 2) {
  stop("Not enough valid AB_distance values to compute full AB density.")
}
dens_AB_full <- density(log10(AB_dist_full))
dens_AB_full_df <- data.frame(x = dens_AB_full$x, y = dens_AB_full$y)

AB_sub <- data %>% filter(!is.na(AB_distance) & AB_distance >= 100)
if (nrow(AB_sub) < 10) {
  stop("Not enough data with AB_distance >= 100 to fit AB spline.")
}
AB_dist_sub <- AB_sub$AB_distance
AB_dist_sub <- AB_dist_sub[AB_dist_sub > 0]
if (length(AB_dist_sub) < 2) {
  stop("Not enough valid AB_distance in subset for spline.")
}
dens_AB_sub <- density(log10(AB_dist_sub))
dens_AB_sub_df <- data.frame(x = dens_AB_sub$x, y = dens_AB_sub$y)

# Fit a cubic regression spline to the subset using 20 knots
AB_sub_df2 <- dens_AB_sub_df %>% filter(y > 0)
fit_AB <- mgcv::gam(y ~ s(x, bs = "cr", k = 20), data = AB_sub_df2)

x_grid_AB <- seq(min(AB_sub_df2$x), max(AB_sub_df2$x), length.out = 500)
pred_AB <- predict(fit_AB, newdata = data.frame(x = x_grid_AB), type = "response")
pred_AB[pred_AB < 0] <- 0

dx_AB <- x_grid_AB[2] - x_grid_AB[1]
area_AB <- sum(pred_AB * dx_AB)
pdf_norm_AB <- pred_AB / area_AB
cdf_vals_AB <- cumsum(pdf_norm_AB) * dx_AB

sample_log10dist_AB <- function(n) {
  out <- numeric(n)
  for (i in seq_len(n)) {
    u <- runif(1)
    idx <- which.min(abs(cdf_vals_AB - u))
    out[i] <- x_grid_AB[idx]
  }
  out
}

# ---- BC_distance ----
BC_dist_full <- data$BC_distance
BC_dist_full <- BC_dist_full[!is.na(BC_dist_full) & BC_dist_full > 0]
if (length(BC_dist_full) < 2) {
  stop("Not enough valid BC_distance values to compute full BC density.")
}
dens_BC_full <- density(log10(BC_dist_full))
dens_BC_full_df <- data.frame(x = dens_BC_full$x, y = dens_BC_full$y)

BC_sub <- data %>% filter(!is.na(BC_distance) & BC_distance >= 100)
if (nrow(BC_sub) < 10) {
  stop("Not enough data with BC_distance >= 100 to fit BC spline.")
}
BC_dist_sub <- BC_sub$BC_distance
BC_dist_sub <- BC_dist_sub[BC_dist_sub > 0]
if (length(BC_dist_sub) < 2) {
  stop("Not enough valid BC_distance in subset for spline.")
}
dens_BC_sub <- density(log10(BC_dist_sub))
dens_BC_sub_df <- data.frame(x = dens_BC_sub$x, y = dens_BC_sub$y)

BC_sub_df2 <- dens_BC_sub_df %>% filter(y > 0)
# Fit a cubic regression spline to the BC subset using 20 knots
fit_BC <- mgcv::gam(y ~ s(x, bs = "cr", k = 20), data = BC_sub_df2)

x_grid_BC <- seq(min(BC_sub_df2$x), max(BC_sub_df2$x), length.out = 500)
pred_BC <- predict(fit_BC, newdata = data.frame(x = x_grid_BC), type = "response")
pred_BC[pred_BC < 0] <- 0

dx_BC <- x_grid_BC[2] - x_grid_BC[1]
area_BC <- sum(pred_BC * dx_BC)
pdf_norm_BC <- pred_BC / area_BC
cdf_vals_BC <- cumsum(pdf_norm_BC) * dx_BC

sample_log10dist_BC <- function(n) {
  out <- numeric(n)
  for (i in seq_len(n)) {
    u <- runif(1)
    idx <- which.min(abs(cdf_vals_BC - u))
    out[i] <- x_grid_BC[idx]
  }
  out
}

#----------------------------------------------------------------------
# 3) Plot AB, BC univariate fits => single PDF with 2 panels (using k=20 in geom_smooth)
#----------------------------------------------------------------------
AB_plot <- ggplot() +
  geom_line(data = dens_AB_full_df, aes(x = x, y = y),
            color = "blue", linewidth = 1, alpha = 0.5) +
  geom_smooth(data = dens_AB_sub_df, aes(x = x, y = y),
              method = "gam", formula = y ~ s(x, bs = "cr", k = 20),
              se = FALSE, color = "red", linewidth = 1) +
  theme_minimal() +
  labs(
    x = "Log10(AB_distance) (bp)",
    y = "Density",
    title = "AB Distance: Full (blue) vs. Subset>=100 w/ Cubic Spline (red, k=20)"
  )

BC_plot <- ggplot() +
  geom_line(data = dens_BC_full_df, aes(x = x, y = y),
            color = "blue", linewidth = 1, alpha = 0.5) +
  geom_smooth(data = dens_BC_sub_df, aes(x = x, y = y),
              method = "gam", formula = y ~ s(x, bs = "cr", k = 20),
              se = FALSE, color = "red", linewidth = 1) +
  theme_minimal() +
  labs(
    x = "Log10(BC_distance) (bp)",
    y = "Density",
    title = "BC Distance: Full (blue) vs. Subset>=100 w/ Cubic Spline (red, k=20)"
  )

combo_plot <- AB_plot / BC_plot  # vertical stacking using patchwork
message("Saving AB, BC splines plot to: ", splines_pdf)
ggsave(splines_pdf, combo_plot, width = 7, height = 8)

#----------------------------------------------------------------------
# 4) Anchor from *any* of A, B, C => generate synthetic data
#----------------------------------------------------------------------
N <- 1000000
message("Generating ", N, " synthetic rows...")

# Create an anchor pool from A, B, and C positions
anchor_pool <- bind_rows(
  data.frame(chrom = data$A_chrom, pos = data$A_start),
  data.frame(chrom = data$B_chrom, pos = data$B_start),
  data.frame(chrom = data$C_chrom, pos = data$C_start)
) %>%
  distinct()

message("Anchor pool size: ", nrow(anchor_pool))

anchors <- anchor_pool %>% slice_sample(n = N, replace = TRUE)
anchors$B_pos <- NA
anchors$C_pos <- NA

for (i in seq_len(N)) {
  chrom_i <- anchors$chrom[i]
  anchorA <- anchors$pos[i]

  # 1) Sample AB distance
  distAB_log <- sample_log10dist_AB(1)
  distAB_bp  <- 10^distAB_log
  newB <- place_fragment(chrom_i, anchorA, distAB_bp)
  if (is.na(newB)) next
  anchors$B_pos[i] <- newB

  # 2) Sample BC distance
  distBC_log <- sample_log10dist_BC(1)
  distBC_bp  <- 10^distBC_log
  newC <- place_fragment(chrom_i, newB, distBC_bp)
  if (is.na(newC)) next
  anchors$C_pos[i] <- newC
}

synthetic_df <- anchors %>%
  mutate(
    AB_distance = abs(B_pos - pos),
    BC_distance = abs(C_pos - B_pos),
    AC_distance = abs(C_pos - pos),
    A_chrom     = chrom,
    A_start     = pos,
    B_chrom     = chrom,
    B_start     = B_pos,
    C_chrom     = chrom,
    C_start     = C_pos
  ) %>%
  select(
    A_chrom, A_start,
    B_chrom, B_start,
    C_chrom, C_start,
    AB_distance, BC_distance, AC_distance
  )

message("Writing synthetic data to: ", synthetic_tsv)
write.table(synthetic_df, synthetic_tsv, sep = "\t", quote = FALSE, row.names = FALSE)

#----------------------------------------------------------------------
# 5) Compare AB, BC, AC distributions (real vs. synthetic) and compute Spearman correlations
#----------------------------------------------------------------------
message("Comparing AB, BC, AC distributions in log10 space...")

real_dists <- data.frame(
  dist_type = rep(c("AB", "BC", "AC"), each = nrow(data)),
  distance  = c(data$AB_distance, data$BC_distance, data$AC_distance),
  dataset   = "real"
)

syn_dists <- data.frame(
  dist_type = rep(c("AB", "BC", "AC"), each = nrow(synthetic_df)),
  distance  = c(synthetic_df$AB_distance, synthetic_df$BC_distance, synthetic_df$AC_distance),
  dataset   = "synthetic"
)

combined_dists <- bind_rows(real_dists, syn_dists) %>%
  filter(distance > 0)

p2 <- ggplot(combined_dists, aes(x = log10(distance), color = dataset)) +
  geom_density(linewidth = 1) +
  facet_wrap(~dist_type, scales = "free") +
  theme_minimal() +
  labs(
    title = "AB, BC, AC Distance Comparison: Real vs. Synthetic",
    x = "log10(distance) (bp)",
    y = "Density"
  )

message("Saving distance comparison plot to: ", dist_compare_pdf)
ggsave(dist_compare_pdf, p2, width = 10, height = 4)

# Compute and print Spearman correlation matrices
message("\n--- Spearman Correlation for Real Data ---\n")
real_mat <- data %>% select(AB_distance, BC_distance, AC_distance)
real_corr <- cor(real_mat, method = "spearman", use = "complete.obs")
print(real_corr)

message("\n--- Spearman Correlation for Synthetic Data ---\n")
syn_mat <- synthetic_df %>% select(AB_distance, BC_distance, AC_distance)
syn_corr <- cor(syn_mat, method = "spearman", use = "complete.obs")
print(syn_corr)

message("All done.")
```
