## Supplementary material for "CAD-C: An engineered nuclease enables repair-free *in situ* proximity ligation and nucleosome-resolution chromosome walks in human cells": Custom code: submit_annotate_compt_synthetic.sh

```
#!/bin/bash
#SBATCH --job-name=annotate_walks
#SBATCH --output=annotate_walks_%j.out
#SBATCH --error=annotate_walks_%j.err
#SBATCH --cpus-per-task=8
#SBATCH --nodes=1
# Activate conda
source /ru-auth/local/home/risc_soft2/miniconda3/etc/profile.d/conda.sh
conda activate polars_env  # or whatever env has pybedtools

python /ru-auth/local/home/landerson01/scripts/LA_classify_ABC_frags_by_bed_list.py \
  --abc /rugpfs/fs0/risc_lab/scratch/landerson01/20250205_CAD_2x250bp_GM_analysis/Synthetic/AB_BC_Spline_AcceptonAC/merged_all_flipped_sorted.cis.compt_annotated.tsv.gz_distances_compartment_synthetic.tsv \
  --bed-dir /rugpfs/fs0/risc_lab/scratch/jsoroczyns/CAD/CADwalk/CADwalk_E1_GM12878_2x250/results_20250207_0034_nonexpanded/ABC_walk_processing_20250209/20250214_Bfilt_flattened_parse2_UU_direct_ABC_complete_consequent12/filtzerolenB_flip_sort_dedup_split_20250214/merged_all/GM12878_AO_A_B_compt_mergedBins/ \
  --out /rugpfs/fs0/risc_lab/scratch/landerson01/20250205_CAD_2x250bp_GM_analysis/Synthetic/AB_BC_Spline_AcceptonAC/merged_all_flipped_sorted.cis.compt_annotated.tsv.gz_distances_compartment_synthetic_annotated.tsv \
  --chunk-size 500000 \
  --input_format simple
```
