## Supplementary material for "CAD-C: An engineered nuclease enables repair-free *in situ* proximity ligation and nucleosome-resolution chromosome walks in human cells": Custom code: smart_bigwig_over_bed_v10p6_20250927.py

```
#!/usr/bin/env python3
# =============================================================================
# smart_bigwig_over_bed_v10p6_2025_09_26.py
#
# PURPOSE
#   A flexible, robust, general‑purpose meta‑coverage (pileup) engine for
#   “any .bigWig over any .BED/.BEDPE”, with publication‑quality plotting,
#   provenance, multiprocessing, and professional CLI help.
#
# WHAT'S NEW IN v10p6 (builds on v10p5/v10p4/v10p3/v10p2/v10p1/v10)
#   • Directional sign handling for zero‑centered / directional tracks:
#       - NEW: --flip-sign <selectors...>
#           Invert the signal (multiply by −1) for selected bigWigs globally.
#       - NEW: --flip-sign-if-minus <selectors...>
#           For selected bigWigs, invert the signal **only for minus‑strand
#           intervals** (BED/anchors). This aligns signed scores (e.g. DI,
#           directional Z) into a plus‑strand frame in addition to the usual
#           horizontal flip of the profile.
#       - NEW: --bedpe-pair-sign-from {none,anchor1,anchor2} (default: none)
#           In pair‑scaled BEDPE, use anchor1 or anchor2 strand (if present in
#           BEDPE) to decide minus‑strand sign inversion when the track is listed
#           in --flip-sign-if-minus.
#       - Selectors can be bigWig **labels** (via --bw-labels), **basenames**, or
#         **0‑based indices**; mix & match.
#       - Sign flipping is applied **before** fold‑diff normalization and plotting,
#         so matrices, average TSVs, and plots are coherent.
#
#   • v10p5 recap (retained):
#       - Overlay multiple bigWigs in a single axis per dataset: --overlay-bigwigs
#         with Illustrator‑style draw order via --overlay-order.
#       - Font size toggles: --font-size, --title-size, --label-size, --tick-size,
#         --legend-size; PDF export stays fonttype=42 (TrueType outlines).
#       - Savitzky–Golay smoothing consistently applied to QC and grid plots
#         (TSVs remain raw).
#
#   • v10p4 recap (retained):
#       - Multiprocessing fix (no local nested worker functions; safe pickling).
#
#   • v10p1 recap (retained):
#       - Zero‑centered track detection (robust) with overrides; symmetric y‑axes;
#         boxgrid uses mean signed signal for zero‑centered tracks.
#
# SCOPE & INPUTS
#   • BED: narrow/broad peaks, ChromHMM segments, junctions, TSSs (strand aware).
#   • BEDPE: loops/anchor pairs; intra‑chrom pairs for pair‑scaled mode.
#
# OUTPUTS (per BED or BEDPE × bigWig in `pileup`)
#   • coverage_matrix_*.tsv.gz   (rows = intervals or pairs; columns = positions)
#   • coverage_average_profile_*.tsv   (raw mean profile; smoothing is render‑time)
#   • metaplot_*.pdf             (QC plot per combination)
#   • Optional multi‑panel grid across datasets; with --overlay-bigwigs, all
#     bigWigs are drawn on the same y‑axis with a legend and controlled layering.
#
# BOXGRID SUBCOMMAND
#   • Heatmap (rows=datasets, cols=bigWigs):
#       - zero‑centered → mean signed signal;
#       - others → log2(mean / genome‑wide mean).
#     Palette: metallic blue–white–red, centered at 0. Supports sign‑flip options.
#
# RESOLUTION & ACCURACY
#   • Per‑base extraction for midpoint windows (pyBigWig.values).
#   • nBins “stats” extraction for downsampling and pair‑scaled mode.
#   • Chromosome‑edge clamping and zero‑padding.
#
# REQUIREMENTS
#   Python ≥ 3.9; numpy; matplotlib; pyBigWig; pybedtools; SciPy (Savgol);
#   PyYAML (optional) for YAML configs; pandas (optional). Conda recommended.
#
# AUTHOR & DATE
#   Jan Soroczynski, Risca Lab, The Rockefeller University
#   2025_09_26
#
# LICENSE
#   Apache-2.0 (adjust as needed).
# =============================================================================

import argparse
import sys
import os
import gzip
import math
import json
import re
import datetime
import logging
import platform
import multiprocessing
from dataclasses import dataclass, asdict
from typing import List, Optional, Dict, Any, Tuple

import numpy as np
import matplotlib
matplotlib.use('Agg')
import matplotlib.pyplot as plt

try:
    import pyBigWig
except ImportError as e:
    print("ERROR: pyBigWig is required. Install with conda/pip before running.", file=sys.stderr)
    sys.exit(1)

try:
    import pybedtools
except ImportError:
    print("ERROR: pybedtools is required. Install with conda/pip before running.", file=sys.stderr)
    sys.exit(1)

# Optional dependencies
try:
    from scipy.signal import savgol_filter
    _HAVE_SCIPY = True
except Exception:
    _HAVE_SCIPY = False

try:
    import yaml  # type: ignore
    _HAVE_YAML = True
except Exception:
    _HAVE_YAML = False

try:
    import importlib.metadata as importlib_metadata
except Exception:
    import importlib_metadata  # type: ignore


SCRIPT_VERSION = "v10p6_2025_09_26"
EPS = 1e-12  # numerical stability for ratios/logs


# ------------------------------- Utility helpers ------------------------------

def _now_timestamp() -> str:
    return datetime.datetime.now().strftime("%Y%m%d_%H%M%S")


def _setup_logging():
    logging.basicConfig(format="%(asctime)s [%(levelname)s] %(message)s",
                        level=logging.INFO)


def _pkg_version(pkg: str) -> Optional[str]:
    try:
        return importlib_metadata.version(pkg)
    except Exception:
        return None


def _safe_join_name(parts: List[str], maxlen: int = 60) -> str:
    base = "_PLUS_".join(os.path.basename(p) for p in parts)
    return base if len(base) <= maxlen else base[:maxlen] + "_trunc"


def _create_outdir(base_dir: str, left_parts: List[str], right_parts: List[str], ts: str) -> str:
    left = _safe_join_name(left_parts) if left_parts else "NA"
    right = _safe_join_name(right_parts) if right_parts else "NA"
    outdir_name = f"metacov_{left}_{right}_{ts}"
    outdir = os.path.join(base_dir, outdir_name)
    os.makedirs(outdir, exist_ok=True)
    return outdir


def _write_metadata(outdir: str, ts: str, args: argparse.Namespace, effective_cfg: Dict[str, Any],
                    extra: Optional[Dict[str, Any]] = None):
    meta_path = os.path.join(outdir, f"run_metadata_{ts}.txt")
    with open(meta_path, "w") as f:
        f.write(f"Script: smart_bigwig_over_bed_{SCRIPT_VERSION}.py\n")
        f.write(f"Timestamp: {ts}\n\n")

        f.write("Command-line arguments (raw):\n")
        for k, v in vars(args).items():
            f.write(f"  {k} = {v}\n")
        f.write("\nEffective configuration (after merging config + CLI):\n")
        f.write(json.dumps(effective_cfg, indent=2))
        f.write("\n\nEnvironment:\n")
        f.write(f"  Python: {sys.version}\n")
        f.write(f"  Platform: {platform.platform()}\n")
        f.write(f"  numpy: {_pkg_version('numpy')}\n")
        f.write(f"  scipy: {_pkg_version('scipy')}\n")
        f.write(f"  matplotlib: {_pkg_version('matplotlib')}\n")
        f.write(f"  pyBigWig: {_pkg_version('pyBigWig')}\n")
        f.write(f"  pybedtools: {_pkg_version('pybedtools')}\n")
        f.write(f"  pandas: {_pkg_version('pandas')}\n")
        if extra:
            f.write("\nExtra metadata:\n")
            f.write(json.dumps(extra, indent=2))
    logging.info(f"Metadata written: {meta_path}")


# ------------------------------- Data structures ------------------------------

class IntervalWithStrand:
    __slots__ = ("chrom", "start", "end", "strand")
    def __init__(self, chrom: str, start: int, end: int, strand: Optional[str] = None):
        self.chrom = chrom
        self.start = int(start)
        self.end   = int(end)
        self.strand = strand  # '+', '-', or None


@dataclass
class BedPeEntry:
    chrom1: str
    start1: int
    end1: int
    chrom2: str
    start2: int
    end2: int
    strand1: Optional[str] = None
    strand2: Optional[str] = None
    line_no: Optional[int] = None

    def anchor1(self) -> IntervalWithStrand:
        return IntervalWithStrand(self.chrom1, self.start1, self.end1, self.strand1)

    def anchor2(self) -> IntervalWithStrand:
        return IntervalWithStrand(self.chrom2, self.start2, self.end2, self.strand2)


# ------------------------------- BED utilities --------------------------------

def _interval_midpoint(start: int, end: int) -> int:
    return (start + end) // 2


def _detect_bed_strand_column(bed_path: str, max_lines: int = 50) -> int:
    """
    Intersect columns among first `max_lines` that contain only {+,-,.} and
    retain columns that observed at least one '+' or '-'. Return the smallest
    index or -1 if not found.
    """
    possible_cols = None
    plusminus_count: Dict[int, int] = {}
    checked = 0
    with open(bed_path, "r") as fh:
        for line in fh:
            if not line or line.startswith("#"):
                continue
            parts = line.rstrip("\n").split("\t")
            if len(parts) < 3:
                continue
            checked += 1
            cols_this = []
            for i, field in enumerate(parts):
                if field in ("+", "-", "."):
                    cols_this.append(i)
                    if field in ("+", "-"):
                        plusminus_count[i] = plusminus_count.get(i, 0) + 1
            if possible_cols is None:
                possible_cols = set(cols_this)
            else:
                possible_cols.intersection_update(cols_this)
            if checked >= max_lines:
                break
    if not possible_cols:
        return -1
    final = [c for c in possible_cols if plusminus_count.get(c, 0) > 0]
    return min(final) if final else -1


def read_bed_with_strand(bed_path: str) -> Tuple[List[IntervalWithStrand], Dict[str, int]]:
    strand_col = _detect_bed_strand_column(bed_path)
    plus = minus = nostrand = 0
    intervals: List[IntervalWithStrand] = []
    bt = pybedtools.BedTool(bed_path)
    for feat in bt:
        chrom = feat.chrom
        start = int(feat.start)
        end   = int(feat.end)
        s: Optional[str] = None
        if strand_col >= 0 and len(feat.fields) > strand_col:
            raw = feat.fields[strand_col]
            if raw == '+':
                s = '+'
                plus += 1
            elif raw == '-':
                s = '-'
                minus += 1
            else:
                nostrand += 1
        else:
            nostrand += 1
        intervals.append(IntervalWithStrand(chrom, start, end, s))
    stats = {"plus": plus, "minus": minus, "no_strand": nostrand, "total": len(intervals), "strand_col": strand_col}
    return intervals, stats


# ------------------------------ BEDPE utilities -------------------------------

@dataclass
class BedPeColumnSpec:
    c1: int
    s1: int
    e1: int
    c2: int
    s2: int
    e2: int
    strand1: Optional[int]
    strand2: Optional[int]


_chr_pat = re.compile(r"^(chr|Chr|CHR)[\w._-]*$")


def _is_int(x: str) -> bool:
    try:
        int(x)
        return True
    except Exception:
        return False


def _detect_bedpe_columns(bedpe_path: str, max_lines: int = 200) -> BedPeColumnSpec:
    """
    Heuristic detector for BEDPE columns. Falls back to standard 0..5.
    Also scans for two strand columns with '+'/'-'.
    """
    lines: List[List[str]] = []
    with open(bedpe_path, "r") as fh:
        for line in fh:
            if not line or line.startswith("#"):
                continue
            parts = line.rstrip("\n").split("\t")
            if len(parts) < 6:
                continue
            lines.append(parts)
            if len(lines) >= max_lines:
                break

    def _score_spec(spec: Tuple[int, int, int, int, int, int]) -> int:
        c1, s1, e1, c2, s2, e2 = spec
        ok = 0
        for p in lines:
            if len(p) <= max(c1, s1, e1, c2, s2, e2):
                continue
            try:
                if not (_is_int(p[s1]) and _is_int(p[e1]) and _is_int(p[s2]) and _is_int(p[e2])):
                    continue
                if int(p[s1]) < int(p[e1]) and int(p[s2]) < int(p[e2]):
                    ok += 1
            except Exception:
                continue
        return ok

    spec = (0, 1, 2, 3, 4, 5)
    score_std = _score_spec(spec)

    best_spec = spec
    best_score = score_std

    if best_score < max(5, int(0.6 * len(lines))):
        ncols = max(len(p) for p in lines) if lines else 6
        chrom_candidates = set()
        for i in range(min(12, ncols)):
            hits = sum(1 for p in lines if len(p) > i and (_chr_pat.match(p[i]) or p[i].lower().startswith("chr")))
            if hits >= max(3, int(0.3 * len(lines))):
                chrom_candidates.add(i)
        bin_specs: List[Tuple[int, int, int]] = []
        for c in chrom_candidates:
            for s in range(c + 1, min(c + 4, ncols - 1)):
                e = s + 1
                good = 0
                for p in lines:
                    if len(p) <= e:
                        continue
                    if _is_int(p[s]) and _is_int(p[e]) and (int(p[s]) < int(p[e])):
                        good += 1
                if good >= max(3, int(0.3 * len(lines))):
                    bin_specs.append((c, s, e))
        for (c1, s1, e1) in bin_specs:
            for (c2, s2, e2) in bin_specs:
                if c2 > e1:
                    sc = _score_spec((c1, s1, e1, c2, s2, e2))
                    if sc > best_score:
                        best_score = sc
                        best_spec = (c1, s1, e1, c2, s2, e2)

    strand_cols: List[int] = []
    ncols2 = max(len(p) for p in lines) if lines else 0
    for j in range(ncols2):
        pm_hits = sum(1 for p in lines if len(p) > j and p[j] in ("+", "-"))
        if pm_hits >= max(3, int(0.3 * len(lines))):
            strand_cols.append(j)
    strand1 = strand_cols[0] if len(strand_cols) >= 1 else None
    strand2 = strand_cols[1] if len(strand_cols) >= 2 else None

    c1, s1, e1, c2, s2, e2 = best_spec
    return BedPeColumnSpec(c1, s1, e1, c2, s2, e2, strand1, strand2)


def read_bedpe(bedpe_path: str,
               override_cols: Optional[Tuple[int, int, int, int, int, int]] = None,
               override_strand_cols: Optional[Tuple[Optional[int], Optional[int]]] = None
               ) -> Tuple[List[BedPeEntry], Dict[str, Any]]:
    spec = None
    if override_cols:
        spec = BedPeColumnSpec(*override_cols, strand1=None, strand2=None)
    else:
        spec = _detect_bedpe_columns(bedpe_path)

    if override_strand_cols:
        spec.strand1, spec.strand2 = override_strand_cols

    entries: List[BedPeEntry] = []
    skipped_short = 0
    with open(bedpe_path, "r") as fh:
        for ln, line in enumerate(fh, start=1):
            if not line or line.startswith("#"):
                continue
            parts = line.rstrip("\n").split("\t")
            try:
                c1, s1, e1 = parts[spec.c1], int(parts[spec.s1]), int(parts[spec.e1])
                c2, s2, e2 = parts[spec.c2], int(parts[spec.s2]), int(parts[spec.e2])
            except Exception:
                continue
            st1 = parts[spec.strand1] if (spec.strand1 is not None and spec.strand1 < len(parts)) else None
            st2 = parts[spec.strand2] if (spec.strand2 is not None and spec.strand2 < len(parts)) else None
            if e1 <= s1 or e2 <= s2:
                skipped_short += 1
                continue
            entries.append(BedPeEntry(c1, s1, e1, c2, s2, e2,
                                      st1 if st1 in ("+", "-") else None,
                                      st2 if st2 in ("+", "-") else None, ln))
    meta = {
        "bedpe_columns_used": asdict(spec),
        "bedpe_total_read": len(entries),
        "bedpe_skipped_nonpositive_length": skipped_short
    }
    return entries, meta


# ----------------------------- BigWig I/O helpers -----------------------------

def _bw_values_segment(bw, chrom: str, start_i: int, end_i: int) -> np.ndarray:
    vals = bw.values(chrom, start_i, end_i, numpy=True)
    if vals is None or len(vals) == 0:
        return np.zeros(max(0, end_i - start_i), dtype=float)
    return np.asarray(vals, dtype=float)


def bigwig_global_mean(bw_path: str) -> float:
    """
    Weighted genome‑wide mean from bigWig using pyBigWig.stats for each chrom.
    """
    bw = pyBigWig.open(bw_path)
    try:
        total_len = 0
        acc = 0.0
        for chrom, clen in bw.chroms().items():
            m = bw.stats(chrom, 0, int(clen), type="mean")[0]
            m = 0.0 if (m is None or not np.isfinite(m)) else float(m)
            total_len += int(clen)
            acc += m * int(clen)
        return (acc / total_len) if total_len > 0 else 0.0
    finally:
        bw.close()


# -------------------- Zero‑centered track detection (v10p1+) ------------------

def detect_zero_centered_bigwig(bw_path: str,
                                sample_bins_per_chrom: int = 200,
                                max_total_bins: int = 6000,
                                robust=True,
                                tol_center: float = 0.15) -> Dict[str, Any]:
    """
    Heuristically decide if a bigWig is zero‑centered (robust‑Z, DI, insulation).
    Returns dict with: zero_centered, signed, median (~center), mad (~sigma),
    min, max, n.
    """
    bw = pyBigWig.open(bw_path)
    vals: List[float] = []
    try:
        for chrom, L in bw.chroms().items():
            if L <= 0:
                continue
            nBins = min(sample_bins_per_chrom, int(max(1, L // max(1, L // sample_bins_per_chrom))))
            stats = bw.stats(chrom, 0, int(L), nBins=nBins, type="mean")
            for x in stats:
                if x is not None and np.isfinite(x):
                    vals.append(float(x))
            if len(vals) >= max_total_bins:
                break
    finally:
        bw.close()

    if len(vals) == 0:
        return {"zero_centered": False, "signed": False, "median": 0.0,
                "mad": 0.0, "min": 0.0, "max": 0.0, "n": 0}

    arr = np.array(vals, dtype=float)
    vmin, vmax = float(np.nanmin(arr)), float(np.nanmax(arr))
    signed = (vmin < 0.0) and (vmax > 0.0)

    if robust:
        med = float(np.nanmedian(arr))
        mad = float(np.nanmedian(np.abs(arr - med))) * 1.4826  # ≈ robust σ
        scale = max(mad, 1e-12)
        zeroish = (abs(med) <= tol_center * scale)
        return {"zero_centered": (signed and zeroish), "signed": signed,
                "median": med, "mad": mad, "min": vmin, "max": vmax, "n": len(arr)}
    else:
        mu = float(np.nanmean(arr)); sd = float(np.nanstd(arr))
        scale = max(sd, 1e-12)
        zeroish = (abs(mu) <= tol_center * scale)
        return {"zero_centered": (signed and zeroish), "signed": signed,
                "median": mu, "mad": sd, "min": vmin, "max": vmax, "n": len(arr)}


# ------------------------------ Coverage engines ------------------------------

def compute_midpoint_matrix(intervals: List[IntervalWithStrand],
                            bw_path: str, half_window: int,
                            smoothing_bp: int = 0,
                            read_mode: str = "values") -> np.ndarray:
    """
    Returns shape (N, 2*half_window); read_mode in {"values","stats"}.
    """
    bw = pyBigWig.open(bw_path)
    bw_chroms = bw.chroms()
    wsize = 2 * half_window
    out: List[np.ndarray] = []

    for iv in intervals:
        chrom, strand = iv.chrom, iv.strand
        if chrom not in bw_chroms:
            out.append(np.zeros(wsize, dtype=float)); continue

        chr_len = int(bw_chroms[chrom])
        mid = _interval_midpoint(iv.start, iv.end)
        s = max(0, min(chr_len, mid - half_window))
        e = max(0, min(chr_len, mid + half_window))
        if e <= s:
            out.append(np.zeros(wsize, dtype=float)); continue

        if read_mode == "stats":
            stats = bw.stats(chrom, int(s), int(e), nBins=wsize)
            arr = np.array([0.0 if (x is None or not np.isfinite(x)) else float(x) for x in stats], dtype=float)
        else:
            cov = _bw_values_segment(bw, chrom, int(s), int(e))
            arr = np.zeros(wsize, dtype=float)
            left_offset = int(s - (mid - half_window))
            left_offset = max(0, left_offset)
            right_idx = min(wsize, left_offset + len(cov))
            arr[left_offset:right_idx] = cov[:(right_idx - left_offset)]

        if smoothing_bp and smoothing_bp > 0:
            kernel = np.ones(smoothing_bp, dtype=float)
            arr = np.convolve(arr, kernel, mode="same") / smoothing_bp

        if strand == '-':
            arr = arr[::-1]

        out.append(arr)

    bw.close()
    return np.vstack(out) if out else np.zeros((0, wsize), dtype=float)


def compute_scaled_matrix(intervals: List[IntervalWithStrand],
                          bw_path: str, main_bins: int,
                          flank_bins: int = 0, use_flank: bool = False) -> np.ndarray:
    """
    BED scaled mode: mean per bin across main (and optional flanks ±50%).
    """
    bw = pyBigWig.open(bw_path)
    bw_chroms = bw.chroms()
    total_bins = (main_bins + 2*flank_bins) if use_flank else main_bins
    out: List[np.ndarray] = []

    def _bin_mean(c: str, a: float, b: float, clen: int) -> float:
        a = max(0, min(clen, a)); b = max(0, min(clen, b))
        ai, bi = int(math.floor(a)), int(math.ceil(b))
        if bi <= ai:
            return 0.0
        vals = _bw_values_segment(bw, c, ai, bi)
        return float(np.nanmean(vals)) if vals.size > 0 else 0.0

    for iv in intervals:
        chrom, strand = iv.chrom, iv.strand
        arr = np.zeros(total_bins, dtype=float)
        if chrom not in bw_chroms:
            out.append(arr); continue

        clen = int(bw_chroms[chrom])
        ivlen = max(1, iv.end - iv.start)
        idx = 0

        if use_flank:
            L = 0.5 * ivlen
            step = L / flank_bins if flank_bins > 0 else 0
            left_start = iv.start - L
            for i in range(flank_bins):
                arr[idx] = _bin_mean(chrom, left_start + i*step, left_start + (i+1)*step, clen); idx += 1

        main_start = max(0, min(clen, iv.start))
        main_end   = max(0, min(clen, iv.end))
        main_len   = max(0.0, main_end - main_start)
        if main_len > 0 and main_bins > 0:
            step = main_len / main_bins
            for i in range(main_bins):
                arr[idx] = _bin_mean(chrom, main_start + i*step, main_start + (i+1)*step, clen); idx += 1
        else:
            idx += main_bins  # zeros

        if use_flank:
            R = 0.5 * ivlen
            step = R / flank_bins if flank_bins > 0 else 0
            right_start = iv.end
            for i in range(flank_bins):
                arr[idx] = _bin_mean(chrom, right_start + i*step, right_start + (i+1)*step, clen); idx += 1

        if strand == '-':
            arr = arr[::-1]
        out.append(arr)

    bw.close()
    return np.vstack(out) if out else np.zeros((0, total_bins), dtype=float)


def compute_bedpe_pair_scaled_matrix(pairs: List[BedPeEntry],
                                     bw_path: str,
                                     main_bins: int,
                                     flank_frac_left: float = 0.5,
                                     flank_frac_right: float = 0.5,
                                     flank_bins: int = 100) -> Tuple[np.ndarray, int]:
    """
    For intra-chrom pairs only: rescale [mid1, mid2] to main_bins; add left/right
    flanks with lengths = flank_frac_* × (mid2-mid1), each divided into flank_bins.
    Extraction uses bigWig.stats(..., nBins=total_bins) for speed.
    Returns (matrix, skipped_interchrom_count).
    """
    bw = pyBigWig.open(bw_path)
    bw_chroms = bw.chroms()
    out: List[np.ndarray] = []
    skipped_interchrom = 0

    for entry in pairs:
        if entry.chrom1 != entry.chrom2:
            skipped_interchrom += 1
            continue
        chrom = entry.chrom1
        if chrom not in bw_chroms:
            total_bins = flank_bins + main_bins + flank_bins
            out.append(np.zeros(total_bins, dtype=float))
            continue

        clen = int(bw_chroms[chrom])
        mid1 = _interval_midpoint(entry.start1, entry.end1)
        mid2 = _interval_midpoint(entry.start2, entry.end2)
        if mid1 == mid2:
            total_bins = flank_bins + main_bins + flank_bins
            out.append(np.zeros(total_bins, dtype=float))
            continue

        left_mid, right_mid = (mid1, mid2) if mid1 < mid2 else (mid2, mid1)
        span = right_mid - left_mid
        L = int(round(flank_frac_left * span))
        R = int(round(flank_frac_right * span))
        start = max(0, left_mid - L)
        end   = min(clen, right_mid + R)
        total_bins = flank_bins + main_bins + flank_bins

        stats = bw.stats(chrom, start, end, nBins=total_bins)
        arr = np.array([0.0 if (x is None or not np.isfinite(x)) else float(x) for x in stats], dtype=float)
        out.append(arr)

    bw.close()
    return (np.vstack(out) if out else np.zeros((0, main_bins + 2*flank_bins), dtype=float),
            skipped_interchrom)


# ------------------------------ Fonts & plotting ------------------------------

def apply_font_rcparams(fonttype: int,
                        font_size: int,
                        title_size: int,
                        label_size: int,
                        tick_size: int,
                        legend_size: int):
    plt.rcParams['pdf.fonttype'] = fonttype
    plt.rcParams.update({
        'font.size': font_size,
        'axes.titlesize': title_size,
        'axes.labelsize': label_size,
        'xtick.labelsize': tick_size,
        'ytick.labelsize': tick_size,
        'legend.fontsize': legend_size
    })


def _ensure_odd_leq(value: int, L: int) -> int:
    w = min(value, L if L % 2 == 1 else L - 1)
    if w < 3:
        w = 3
    if w % 2 == 0:
        w -= 1
    return w


def savgol_safe(y: np.ndarray, enabled: bool, window: int, poly: int) -> np.ndarray:
    if not enabled or y.size == 0:
        return y
    if not _HAVE_SCIPY:
        logging.warning("SciPy not installed; skipping Savitzky–Golay smoothing.")
        return y
    L = len(y)
    w = _ensure_odd_leq(window, L)
    p = min(poly, w - 1)
    try:
        return savgol_filter(y, window_length=w, polyorder=p, mode="interp")
    except Exception:
        return savgol_filter(y, window_length=w, polyorder=p, mode="nearest")


def metallic_bwr_cmap():
    from matplotlib.colors import LinearSegmentedColormap
    colors = ["#1f3b99", "#3f67ff", "#b5c7ff", "#ffffff", "#ffb3b3", "#ff5757", "#b81e1e"]
    return LinearSegmentedColormap.from_list("metallic_bwr", colors, N=256)


def compute_y_limits_aggregated(panels: List[Dict[str, Any]], lock_y: str
                                ) -> Dict[Tuple[int, int], Tuple[float, float]]:
    # Aggregate all series per (row,col) and compute symmetric/global/row/column locks
    agg: Dict[Tuple[int, int], Dict[str, Any]] = {}
    for p in panels:
        key = (p["row"], p["col"])
        if key not in agg:
            agg[key] = {"vals": [], "sym": False}
        ys = []
        if p.get("y_raw") is not None:
            ys.append(p["y_raw"])
        if p.get("y_smooth") is not None:
            ys.append(p["y_smooth"])
        agg[key]["vals"].extend(ys)
        agg[key]["sym"] = agg[key]["sym"] or bool(p.get("symmetric_zero", False))

    per_panel_limits: Dict[Tuple[int, int], Tuple[float, float]] = {}
    per_panel_absmax: Dict[Tuple[int, int], float] = {}
    for key, info in agg.items():
        ys = info["vals"]
        if len(ys) == 0:
            ymin, ymax, amax = 0.0, 1.0, 1.0
        else:
            ymin = float(np.nanmin([np.nanmin(y) for y in ys]))
            ymax = float(np.nanmax([np.nanmax(y) for y in ys]))
            if not np.isfinite(ymin) or not np.isfinite(ymax) or ymin == ymax:
                ymin, ymax = 0.0, 1.0
            amax = float(max(abs(ymin), abs(ymax)))
            if not np.isfinite(amax) or amax == 0:
                amax = 1.0
        if info["sym"]:
            per_panel_limits[key] = (-amax, amax)
        else:
            per_panel_limits[key] = (ymin, ymax)
        per_panel_absmax[key] = amax

    if lock_y == "none":
        return per_panel_limits

    rows = sorted({r for (r, c) in per_panel_limits.keys()})
    cols = sorted({c for (r, c) in per_panel_limits.keys()})

    def scope_has_symmetric(scope: List[Tuple[int, int]]) -> bool:
        return any((per_panel_limits[s][0] == -per_panel_absmax[s] and per_panel_limits[s][1] == per_panel_absmax[s])
                   for s in scope)

    if lock_y == "global":
        scope = [(r, c) for r in rows for c in cols if (r, c) in per_panel_limits]
        if scope_has_symmetric(scope):
            gabs = max(per_panel_absmax[s] for s in scope)
            return {(r, c): (-gabs, gabs) for r in rows for c in cols if (r, c) in per_panel_limits}
        gmin = min(per_panel_limits[s][0] for s in scope)
        gmax = max(per_panel_limits[s][1] for s in scope)
        return {(r, c): (gmin, gmax) for r in rows for c in cols if (r, c) in per_panel_limits}

    if lock_y == "row":
        out: Dict[Tuple[int, int], Tuple[float, float]] = {}
        for r in rows:
            scope = [(r, c) for c in cols if (r, c) in per_panel_limits]
            if scope_has_symmetric(scope):
                rabs = max(per_panel_absmax[s] for s in scope)
                for c in cols:
                    if (r, c) in per_panel_limits:
                        out[(r, c)] = (-rabs, rabs)
            else:
                rmin = min(per_panel_limits[s][0] for s in scope)
                rmax = max(per_panel_limits[s][1] for s in scope)
                for c in cols:
                    if (r, c) in per_panel_limits:
                        out[(r, c)] = (rmin, rmax)
        return out

    if lock_y == "column":
        out: Dict[Tuple[int, int], Tuple[float, float]] = {}
        for c in cols:
            scope = [(r, c) for r in rows if (r, c) in per_panel_limits]
            if scope_has_symmetric(scope):
                cabs = max(per_panel_absmax[s] for s in scope)
                for r in rows:
                    if (r, c) in per_panel_limits:
                        out[(r, c)] = (-cabs, cabs)
            else:
                cmin = min(per_panel_limits[s][0] for s in scope)
                cmax = max(per_panel_limits[s][1] for s in scope)
                for r in rows:
                    if (r, c) in per_panel_limits:
                        out[(r, c)] = (cmin, cmax)
        return out

    return per_panel_limits


def plot_grid(grid_data: List[Dict[str, Any]], nrows: int, ncols: int,
              mode: str, out_base: str, plot_format: str,
              lock_y: str, fonttype: int, dpi: int,
              xlabel: str, ylabel: str,
              font_size: int, title_size: int, label_size: int, tick_size: int, legend_size: int):
    """
    grid_data items:
      row, col, x, y_raw (array or None), y_smooth (array or None),
      color, title, symmetric_zero (bool), raw_alpha, raw_lw, smooth_lw,
      legend_label (optional), layer (int; lower drawn first).
    Multiple entries with identical (row,col) are overlaid in that cell.
    """
    apply_font_rcparams(fonttype, font_size, title_size, label_size, tick_size, legend_size)

    grid_sorted = sorted(grid_data, key=lambda d: (d["row"], d["col"], d.get("layer", 0)))

    limits_map = compute_y_limits_aggregated(
        [{"row": d["row"], "col": d["col"], "y_raw": d.get("y_raw"), "y_smooth": d.get("y_smooth"),
          "symmetric_zero": d.get("symmetric_zero", False)} for d in grid_sorted],
        lock_y
    )

    fig, axes = plt.subplots(nrows, ncols, figsize=(4*ncols + 1, 3*nrows + 1), sharex=True)
    if nrows == 1 and ncols == 1:
        axes = np.array([[axes]])
    elif nrows == 1:
        axes = axes.reshape(1, -1)
    elif ncols == 1:
        axes = axes.reshape(-1, 1)

    legend_handles: Dict[Tuple[int, int], Dict[str, Any]] = {}

    for d in grid_sorted:
        ax = axes[d["row"], d["col"]]
        x = d["x"]; color = d["color"]; lh = None
        if d.get("y_raw") is not None:
            (lh_raw,) = ax.plot(x, d["y_raw"], alpha=d.get("raw_alpha", 0.3),
                                linewidth=d.get("raw_lw", 1.0), color=color, label=None)
            lh = lh_raw
        if d.get("y_smooth") is not None:
            (lh_sm,) = ax.plot(x, d["y_smooth"], linewidth=d.get("smooth_lw", 2.0),
                               color=color, label=None)
            lh = lh_sm
        ymin, ymax = limits_map[(d["row"], d["col"])]
        ax.set_ylim(ymin, ymax)
        if mode == "midpoint":
            ax.axvline(0, color="black", linestyle="--", linewidth=1)
        if not hasattr(ax, "_metacov_title_set"):
            ax.set_title(d.get("title", ""), fontsize=title_size)
            ax._metacov_title_set = True
        lab = d.get("legend_label", None)
        if lab and lh is not None:
            key = (d["row"], d["col"])
            legend_handles.setdefault(key, {})[lab] = lh

    for r in range(nrows):
        for c in range(ncols):
            ax = axes[r, c]
            if r == nrows - 1:
                ax.set_xlabel(xlabel)
            if c == 0:
                ax.set_ylabel(ylabel)
            key = (r, c)
            if key in legend_handles and len(legend_handles[key]) >= 2:
                labs = list(legend_handles[key].keys())
                hnds = [legend_handles[key][lab] for lab in labs]
                ax.legend(hnds, labs, loc="best")

    fig.tight_layout()
    if plot_format in ("pdf", "both"):
        fig.savefig(f"{out_base}.pdf")
        logging.info(f"Grid plot saved: {out_base}.pdf")
    if plot_format in ("png", "both"):
        fig.savefig(f"{out_base}.png", dpi=dpi)
        logging.info(f"Grid plot saved: {out_base}.png")
    plt.close(fig)


# ------------------------------ Config & parsing ------------------------------

def load_config(path: Optional[str]) -> Dict[str, Any]:
    if not path:
        return {}
    ext = os.path.splitext(path)[1].lower()
    with open(path, "r") as f:
        if ext in (".yaml", ".yml"):
            if not _HAVE_YAML:
                raise RuntimeError("PyYAML is not installed; cannot read YAML config.")
            return yaml.safe_load(f) or {}
        else:
            return json.load(f)


def merge_cli_cfg(cli_ns: argparse.Namespace, cfg: Dict[str, Any]) -> Dict[str, Any]:
    merged = dict(cfg)
    for k, v in vars(cli_ns).items():
        if v is None:
            continue
        if isinstance(v, list) and len(v) == 0:
            continue
        merged[k] = v
    return merged


def resolve_overlay_order(tokens: Optional[List[str]],
                          bigwigs: List[str],
                          bw_labels: List[str]) -> List[int]:
    n = len(bigwigs)
    remaining = list(range(n))
    order: List[int] = []
    if tokens:
        tks = [str(t).strip() for t in tokens if str(t).strip() != ""]
        for tok in tks:
            idx = None
            if re.fullmatch(r"\d+", tok):
                ii = int(tok)
                if 0 <= ii < n:
                    idx = ii
            if idx is None:
                for i, lab in enumerate(bw_labels):
                    if tok == lab and i in remaining:
                        idx = i; break
            if idx is None:
                for i, path in enumerate(bigwigs):
                    base = os.path.basename(path)
                    if tok == base and i in remaining:
                        idx = i; break
            if idx is not None and idx in remaining:
                order.append(idx); remaining.remove(idx)
    order.extend(remaining)
    return order


def resolve_track_selectors(tokens: Optional[List[str]],
                            bigwigs: List[str],
                            bw_labels: List[str]) -> set:
    """
    Map a list of selectors (labels, basenames, or 0-based indices) to a set
    of track indices.
    """
    sel = set()
    if not tokens:
        return sel
    n = len(bigwigs)
    for tok in tokens:
        if tok is None:
            continue
        s = str(tok).strip()
        if s == "":
            continue
        if re.fullmatch(r"\d+", s):
            i = int(s)
            if 0 <= i < n:
                sel.add(i)
            continue
        for i, lab in enumerate(bw_labels):
            if s == lab:
                sel.add(i)
        for i, path in enumerate(bigwigs):
            if s == os.path.basename(path):
                sel.add(i)
    return sel


# ------------------------------ Worker functions ------------------------------

def _process_bed_subset(subset: List[IntervalWithStrand],
                        bw_path: str,
                        mode: str,
                        window: int,
                        smoothing_bp: int,
                        read_mode: str,
                        scaled_bins: int,
                        flank_bins: int,
                        flank: bool,
                        flip_always: bool,
                        flip_on_minus: bool) -> np.ndarray:
    if mode == "midpoint":
        M = compute_midpoint_matrix(subset, bw_path, window, smoothing_bp=smoothing_bp, read_mode=read_mode)
    else:
        M = compute_scaled_matrix(subset, bw_path, scaled_bins, flank_bins, flank)
    if flip_always and M.size:
        M = -M
    if flip_on_minus and M.size:
        signs = np.array([(-1.0 if iv.strand == '-' else 1.0) for iv in subset], dtype=float)
        M = M * signs[:, None]
    return M


def _process_bedpe_pair_subset(subset_entries: List[BedPeEntry],
                               bw_path: str,
                               main_bins: int,
                               flank_left: float,
                               flank_right: float,
                               flank_bins: int,
                               flip_always: bool,
                               flip_on_minus: bool,
                               pair_sign_from: str  # "none" | "anchor1" | "anchor2"
                               ) -> Tuple[np.ndarray, int]:
    M, skipped = compute_bedpe_pair_scaled_matrix(subset_entries, bw_path,
                                                  main_bins=main_bins,
                                                  flank_frac_left=flank_left,
                                                  flank_frac_right=flank_right,
                                                  flank_bins=flank_bins)
    if M.size:
        if flip_always:
            M = -M
        if flip_on_minus and pair_sign_from != "none":
            if pair_sign_from == "anchor1":
                signs = np.array([(-1.0 if e.strand1 == '-' else 1.0) for e in subset_entries], dtype=float)
            else:
                signs = np.array([(-1.0 if e.strand2 == '-' else 1.0) for e in subset_entries], dtype=float)
            M = M * signs[:, None]
    return M, skipped


def _process_bedpe_anchor_subset(anchor_intervals: List[IntervalWithStrand],
                                 bw_path: str,
                                 window: int,
                                 smoothing_bp: int,
                                 read_mode: str,
                                 flip_always: bool,
                                 flip_on_minus: bool) -> np.ndarray:
    M = compute_midpoint_matrix(anchor_intervals, bw_path, window, smoothing_bp=smoothing_bp, read_mode=read_mode)
    if M.size:
        if flip_always:
            M = -M
        if flip_on_minus:
            signs = np.array([(-1.0 if iv.strand == '-' else 1.0) for iv in anchor_intervals], dtype=float)
            M = M * signs[:, None]
    return M


# --------------------------------- Subcommands --------------------------------

def build_parser() -> argparse.ArgumentParser:
    parser = argparse.ArgumentParser(
        description="BigWig over BED/BEDPE meta‑coverage with robust plotting, zero‑centered detection, overlay, layering, sign‑flip control, and provenance.",
        formatter_class=argparse.ArgumentDefaultsHelpFormatter
    )
    sub = parser.add_subparsers(dest="subcmd", required=True)

    # ---- pileup ----
    p = sub.add_parser("pileup", help="Compute coverage matrices, average profiles, and plots.")
    p.add_argument("-b", "--beds", nargs="*", default=[], help="BED files.")
    p.add_argument("-B", "--bedpes", nargs="*", default=[], help="BEDPE files.")
    p.add_argument("-w", "--bigwigs", nargs="+", required=True, help="bigWig files.")
    p.add_argument("-o", "--output-dir", required=True, help="Base output directory.")
    p.add_argument("-p", "--processors", type=int, default=1, help="Worker processes.")

    # BED modes
    p.add_argument("--mode", choices=["midpoint", "scaled"], default="midpoint", help="BED aggregation mode.")
    p.add_argument("--window", type=int, default=5000, help="Half-window for midpoint mode (bp).")
    p.add_argument("--smoothing", type=int, default=0, help="Legacy per‑interval boxcar smoothing width (bp).")
    p.add_argument("--read-mode", choices=["values", "stats"], default="values",
                   help="Extraction method for midpoint mode.")
    p.add_argument("--scaled-bins", type=int, default=100, help="Main bins for BED scaled mode.")
    p.add_argument("--flank-bins", type=int, default=50, help="Bins per flank for BED scaled mode when --flank.")
    p.add_argument("--flank", action="store_true", help="Include ±50%% flanks in BED scaled mode.")

    # BEDPE controls
    p.add_argument("--bedpe-main-bins", type=int, default=200, help="Main bins for BEDPE pair‑scaled mode.")
    p.add_argument("--bedpe-flank-bins", type=int, default=100, help="Bins per flank for BEDPE pair‑scaled mode.")
    p.add_argument("--bedpe-flank-left", type=float, default=0.5,
                   help="Left flank length as a fraction of pair span (mid2-mid1).")
    p.add_argument("--bedpe-flank-right", type=float, default=0.5,
                   help="Right flank length as a fraction of pair span (mid2-mid1).")
    p.add_argument("--anchors-separate", action="store_true",
                   help="Also compute anchor‑midpoint pileups for BEDPE anchors separately.")
    p.add_argument("--flip-anchor2", action="store_true",
                   help="Flip anchor2 midpoint windows so both anchors align to plus orientation (affects anchor strand and thus sign‑flip if enabled).")
    p.add_argument("--respect-bedpe-strand", action="store_true",
                   help="If strand columns exist in BEDPE, use them to orient midpoint windows (anchors) and to decide minus‑strand sign‑flip if requested.")
    p.add_argument("--bedpe-cols", nargs=6, type=int, metavar=("c1","s1","e1","c2","s2","e2"),
                   help="Override BEDPE column indices (0‑based).")
    p.add_argument("--bedpe-strand-cols", nargs=2, type=int, metavar=("strand1_col","strand2_col"),
                   help="Override BEDPE strand column indices (0‑based).")

    # Zero‑centered handling
    p.add_argument("--zero-centered-detect", choices=["auto", "off"], default="auto",
                   help="Auto‑detect zero‑centered tracks (robust Z/DI/log‑ratio).")
    p.add_argument("--force-zero-centered", nargs="*", default=None,
                   help="Labels or basenames of bigWigs to force as zero‑centered (override detection).")

    # NEW: Sign‑flip controls (per bigWig)
    p.add_argument("--flip-sign", nargs="*", default=None,
                   help="Selectors (labels/basenames/indices) of bigWigs to invert sign globally (multiply by −1).")
    p.add_argument("--flip-sign-if-minus", nargs="*", default=None,
                   help="Selectors (labels/basenames/indices) of bigWigs to invert sign only for minus‑strand intervals (BED and BEDPE anchors).")
    p.add_argument("--bedpe-pair-sign-from", choices=["none", "anchor1", "anchor2"], default="none",
                   help="In pair‑scaled BEDPE, if a bigWig is listed in --flip-sign-if-minus, use this anchor’s strand to decide per‑pair sign inversion.")

    # Overlay & layering
    p.add_argument("--grid", action="store_true", help="Build a comparison grid across datasets.")
    p.add_argument("--grid-rows-axis", choices=["bigwig", "bed"], default="bigwig",
                   help="Which dimension defines grid rows.")
    p.add_argument("--overlay-bigwigs", action="store_true",
                   help="Overlay all bigWigs in a single axis per dataset (requires --grid and --grid-rows-axis bed).")
    p.add_argument("--overlay-order", nargs="*", default=None,
                   help="Overlay draw order (first item drawn first/back). Accepts bigWig labels, basenames, or 0‑based indices. Unspecified tracks keep input order.")
    p.add_argument("--bw-labels", nargs="*", default=None, help="Labels for bigWigs (order matches inputs).")
    p.add_argument("--bw-colors", nargs="*", default=None, help="Colors for bigWigs (order matches inputs).")

    # Normalization & outputs
    p.add_argument("--fold-diff", action="store_true",
                   help="Divide coverage matrices by their global mean (per combo). Skipped for zero‑centered tracks.")
    p.add_argument("--no-write-matrix", dest="write_matrix", action="store_false",
                   help="Do not write the large per‑interval coverage matrix.")
    p.set_defaults(write_matrix=True)

    # Smoothing & plotting
    p.add_argument("--savgol", dest="savgol", action="store_true", help="Enable Savitzky–Golay smoothing (preferred).")
    p.add_argument("--no-savgol", dest="savgol", action="store_false", help="Disable Savitzky–Golay smoothing.")
    p.set_defaults(savgol=True)
    p.add_argument("--savgol-window", type=int, default=101, help="Savgol window length (odd; auto‑adjusted).")
    p.add_argument("--savgol-poly", type=int, default=3, help="Savgol polynomial order.")
    p.add_argument("--lock-y", choices=["none", "global", "row", "column"], default="none",
                   help="Y‑axis locking for grid.")
    p.add_argument("--raw-alpha", type=float, default=0.3, help="Alpha for raw trace.")
    p.add_argument("--raw-lw", type=float, default=1.0, help="Line width for raw trace.")
    p.add_argument("--smooth-lw", type=float, default=2.0, help="Line width for smoothed trace.")
    p.add_argument("--plot-format", choices=["pdf", "png", "both"], default="pdf", help="Grid/QC output format.")
    p.add_argument("--fonttype", type=int, default=42, help="PDF font embedding (42 for TrueType outlines).")

    # Font size toggles
    p.add_argument("--font-size", type=int, default=10, help="Base font size (ticks, general text).")
    p.add_argument("--title-size", type=int, default=10, help="Axes title font size.")
    p.add_argument("--label-size", type=int, default=10, help="Axis label font size.")
    p.add_argument("--tick-size", type=int, default=8,  help="Tick label font size.")
    p.add_argument("--legend-size", type=int, default=8, help="Legend font size.")
    p.add_argument("--dpi", type=int, default=300, help="PNG DPI.")

    p.add_argument("--color", default="magenta", help="Color for simple per‑combo QC plots.")
    p.add_argument("--config", help="YAML or JSON config file; CLI overrides config values.")
    p.add_argument("--example-config", help="Write an example config to PATH and exit.")

    # ---- boxgrid ----
    g = sub.add_parser("boxgrid", help="Produce a boxed heatmap of dataset×bigWig summaries.")
    g.add_argument("-b", "--beds", nargs="*", default=[], help="BED files.")
    g.add_argument("-B", "--bedpes", nargs="*", default=[], help="BEDPE files.")
    g.add_argument("-w", "--bigwigs", nargs="+", required=True, help="bigWig files.")
    g.add_argument("-o", "--output-dir", required=True, help="Base output directory.")
    g.add_argument("-p", "--processors", type=int, default=1, help="Worker processes.")
    g.add_argument("--mode", choices=["midpoint", "scaled", "bedpe_pair", "bedpe_anchors"], default="midpoint",
                   help="How to summarize each BED/BEDPE set.")
    g.add_argument("--window", type=int, default=5000, help="Half‑window for midpoint/anchors mode (bp).")
    g.add_argument("--scaled-bins", type=int, default=100, help="Main bins for BED scaled summary.")
    g.add_argument("--flank", action="store_true", help="Include ±50%% flanks for BED scaled summary.")
    g.add_argument("--flank-bins", type=int, default=50, help="Flank bins for BED scaled summary.")
    g.add_argument("--bedpe-main-bins", type=int, default=200, help="Main bins for BEDPE pair summary.")
    g.add_argument("--bedpe-flank-bins", type=int, default=100, help="Flank bins for BEDPE pair summary.")
    g.add_argument("--bedpe-flank-left", type=float, default=0.5, help="Left flank fraction for BEDPE pair.")
    g.add_argument("--bedpe-flank-right", type=float, default=0.5, help="Right flank fraction for BEDPE pair.")
    g.add_argument("--flip-anchor2", action="store_true", help="Flip anchor2 in anchors summary.")
    g.add_argument("--respect-bedpe-strand", action="store_true",
                   help="Use BEDPE strand columns when present (anchors summary).")
    g.add_argument("--bedpe-cols", nargs=6, type=int, metavar=("c1","s1","e1","c2","s2","e2"),
                   help="Override BEDPE column indices (0‑based).")
    g.add_argument("--bedpe-strand-cols", nargs=2, type=int, metavar=("strand1_col","strand2_col"),
                   help="Override BEDPE strand column indices (0‑based).")
    g.add_argument("--core-fraction", type=float, default=1.0,
                   help="For midpoint/anchors: fraction of window (centered) to average (0<frac≤1).")
    g.add_argument("--fonttype", type=int, default=42, help="PDF font embedding.")
    g.add_argument("--dpi", type=int, default=300, help="PNG DPI.")
    g.add_argument("--plot-format", choices=["pdf", "png", "both"], default="pdf", help="Output format for heatmap.")
    g.add_argument("--cbar-label", default="log2 fold over genome‑wide mean", help="Colorbar label.")
    g.add_argument("--zero-centered-detect", choices=["auto", "off"], default="auto",
                   help="Auto‑detect zero‑centered tracks for summary/scale.")
    g.add_argument("--force-zero-centered", nargs="*", default=None,
                   help="Labels or basenames of bigWigs to force as zero‑centered (override detection).")
    # Sign‑flip controls in boxgrid too (for consistency)
    g.add_argument("--flip-sign", nargs="*", default=None,
                   help="Selectors of bigWigs to invert sign globally.")
    g.add_argument("--flip-sign-if-minus", nargs="*", default=None,
                   help="Selectors of bigWigs to invert sign on minus‑strand intervals (BED/anchors).")
    g.add_argument("--bedpe-pair-sign-from", choices=["none", "anchor1", "anchor2"], default="none",
                   help="Pair‑scaled: source anchor strand for minus‑based sign inversion when requested.")
    # Optional font sizes for boxgrid
    g.add_argument("--font-size", type=int, default=10, help="Base font size.")
    g.add_argument("--title-size", type=int, default=10, help="Title font size.")
    g.add_argument("--label-size", type=int, default=10, help="Axis label font size.")
    g.add_argument("--tick-size", type=int, default=8,  help="Tick label font size.")
    g.add_argument("--legend-size", type=int, default=8, help="Legend font size (reserved).")

    return parser


# ----------------------------------- pileup -----------------------------------

def run_pileup(args: argparse.Namespace):
    # Example config writer
    if getattr(args, "example_config", None):
        example = {
            "beds": ["example1.bed"],
            "bedpes": ["example_pairs.bedpe"],
            "bigwigs": ["track1.bw", "track2.bw"],
            "output_dir": "./out",
            "mode": "midpoint",
            "window": 5000,
            "smoothing": 0,
            "read_mode": "values",
            "scaled_bins": 100,
            "flank_bins": 50,
            "flank": False,
            "bedpe_main_bins": 200,
            "bedpe_flank_bins": 100,
            "bedpe_flank_left": 0.5,
            "bedpe_flank_right": 0.5,
            "anchors_separate": False,
            "flip_anchor2": True,
            "respect_bedpe_strand": False,
            "zero_centered_detect": "auto",
            "force_zero_centered": [],
            "flip_sign": [],
            "flip_sign_if_minus": [],
            "bedpe_pair_sign_from": "none",
            "fold_diff": False,
            "write_matrix": True,
            "grid": True,
            "grid_rows_axis": "bed",
            "overlay_bigwigs": True,
            "overlay_order": [],
            "bw_labels": None,
            "bw_colors": None,
            "savgol": True,
            "savgol_window": 101,
            "savgol_poly": 3,
            "lock_y": "row",
            "raw_alpha": 0.3,
            "raw_lw": 1.0,
            "smooth_lw": 2.0,
            "plot_format": "pdf",
            "fonttype": 42,
            "font_size": 10,
            "title_size": 10,
            "label_size": 10,
            "tick_size": 8,
            "legend_size": 8,
            "dpi": 300
        }
        path = args.example_config
        ext = os.path.splitext(path)[1].lower()
        with open(path, "w") as f:
            if ext in (".yaml", ".yml"):
                if not _HAVE_YAML:
                    raise RuntimeError("PyYAML not installed; cannot write YAML.")
                yaml.safe_dump(example, f, sort_keys=False)
            else:
                json.dump(example, f, indent=2)
        print(f"Wrote example config to: {path}")
        return

    # Merge config if provided
    cfg = load_config(getattr(args, "config", None))
    merged = merge_cli_cfg(args, cfg)

    beds: List[str] = merged.get("beds", [])
    bedpes: List[str] = merged.get("bedpes", [])
    bigwigs: List[str] = merged.get("bigwigs", [])
    output_dir: str = merged.get("output_dir")

    if len(beds) == 0 and len(bedpes) == 0:
        raise SystemExit("ERROR: Provide at least one BED (-b) or BEDPE (-B) file.")

    ts = _now_timestamp()
    outdir = _create_outdir(output_dir, beds + bedpes, bigwigs, ts)

    # Prepare labels/colors for bigWigs
    bw_labels = merged.get("bw_labels", None)
    if bw_labels and len(bw_labels) != len(bigwigs):
        logging.warning("Number of --bw-labels does not match bigWigs; falling back to basenames.")
        bw_labels = [os.path.basename(bw) for bw in bigwigs]
    else:
        bw_labels = bw_labels or [os.path.basename(bw) for bw in bigwigs]

    bw_colors = merged.get("bw_colors", None)
    if bw_colors and len(bw_colors) != len(bigwigs):
        logging.warning("Number of --bw-colors does not match bigWigs; using Matplotlib defaults.")
        bw_colors = None

    # Zero‑centered detection / overrides
    if merged.get("zero_centered_detect", "auto") == "auto":
        zc_info = [detect_zero_centered_bigwig(bw) for bw in bigwigs]
        zero_centered_flags = [z["zero_centered"] for z in zc_info]
    else:
        zc_info = [{"zero_centered": False} for _ in bigwigs]
        zero_centered_flags = [False] * len(bigwigs)

    force_list = merged.get("force_zero_centered") or []
    for idx, bw in enumerate(bigwigs):
        base = os.path.basename(bw)
        label = bw_labels[idx] if idx < len(bw_labels) else base
        if base in force_list or label in force_list:
            zero_centered_flags[idx] = True

    # Sign‑flip selection per bigWig
    flip_always_set = resolve_track_selectors(merged.get("flip_sign"), bigwigs, bw_labels)
    flip_minus_set  = resolve_track_selectors(merged.get("flip_sign_if_minus"), bigwigs, bw_labels)
    pair_sign_from  = merged.get("bedpe_pair_sign_from", "none")

    # Metadata supplement (zero‑center & sign‑flip)
    extra_meta = {
        "zero_centered_detection": {
            bw_labels[i]: {"flag": zero_centered_flags[i], **(zc_info[i] if i < len(zc_info) else {})}
            for i in range(len(bigwigs))
        },
        "sign_flip": {
            bw_labels[i]: {
                "flip_sign_always": (i in flip_always_set),
                "flip_sign_if_minus": (i in flip_minus_set)
            } for i in range(len(bigwigs))
        },
        "bedpe_pair_sign_from": pair_sign_from
    }
    _write_metadata(outdir, ts, args, merged, extra=extra_meta)

    # Overlay setup
    overlay = bool(merged.get("overlay_bigwigs", False))
    if merged.get("grid_rows_axis", "bigwig") != "bed" and overlay:
        logging.warning("--overlay-bigwigs requires --grid and --grid-rows-axis bed; disabling overlay.")
        overlay = False

    # Determine grid geometry (rows × cols)
    all_datasets = beds + bedpes
    if merged.get("grid_rows_axis", "bigwig") == "bigwig":
        nrows, ncols = len(bigwigs), len(all_datasets)
    else:
        nrows, ncols = len(all_datasets), (1 if overlay else len(bigwigs))

    grid_cells: List[Dict[str, Any]] = []
    cpu = max(1, int(merged.get("processors", 1)))

    # Fonts (passed down to plotting)
    fonttype    = int(merged.get("fonttype", 42))
    font_size   = int(merged.get("font_size", 10))
    title_size  = int(merged.get("title_size", 10))
    label_size  = int(merged.get("label_size", 10))
    tick_size   = int(merged.get("tick_size", 8))
    legend_size = int(merged.get("legend_size", 8))

    # Resolve overlay draw order (first = back).
    draw_order = resolve_overlay_order(merged.get("overlay_order"), bigwigs, bw_labels)
    rank = {idx: pos for pos, idx in enumerate(draw_order)}

    # ---------------- BED processing ----------------
    bed_stats_all: Dict[str, Dict[str, int]] = {}
    for bed in beds:
        intervals, stats = read_bed_with_strand(bed)
        bed_stats_all[bed] = stats
        n = len(intervals)
        if n == 0:
            logging.warning(f"{bed}: no intervals; skipping.")
            continue

        chunk = (n + cpu - 1) // cpu
        subsets = [intervals[i:i+chunk] for i in range(0, n, chunk)]

        for wi, bw in enumerate(bigwigs):
            color = (bw_colors[wi] if (bw_colors and wi < len(bw_colors)) else
                     plt.rcParams['axes.prop_cycle'].by_key()['color'][wi % len(plt.rcParams['axes.prop_cycle'].by_key()['color'])])

            # Multiprocessing: top‑level worker (pickle‑safe)
            tasks = [(s, bw, merged.get("mode"), merged.get("window", 5000),
                      merged.get("smoothing", 0), merged.get("read_mode", "values"),
                      merged.get("scaled_bins", 100), merged.get("flank_bins", 50), merged.get("flank", False),
                      (wi in flip_always_set), (wi in flip_minus_set))
                     for s in subsets]
            mats = []
            if cpu > 1:
                with multiprocessing.Pool(processes=cpu) as pool:
                    mats = pool.starmap(_process_bed_subset, tasks)
            else:
                mats = [_process_bed_subset(*t) for t in tasks]

            cov = np.concatenate(mats, axis=0) if mats else np.zeros((0, 2*merged.get("window", 5000)), dtype=float)

            # fold-diff normalization (skip for zero-centered tracks)
            if merged.get("fold_diff", False):
                if zero_centered_flags[wi]:
                    logging.warning(f"{bw_labels[wi]} looks zero-centered; skipping --fold-diff for this track.")
                else:
                    gmean = np.nanmean(cov)
                    if (not np.isfinite(gmean)) or (abs(gmean) < 1e-6):
                        logging.warning("Detected near-zero or non-finite global mean; skipping fold-diff.")
                    else:
                        cov = cov / gmean

            bed_part = _safe_join_name([bed])
            bw_part  = _safe_join_name([bw])

            # Write matrix
            if merged.get("write_matrix", True):
                mat_path = os.path.join(outdir, f"coverage_matrix_{bed_part}_{bw_part}_{ts}.tsv.gz")
                if merged.get("mode") == "midpoint":
                    cols = [str(i - merged.get("window", 5000)) for i in range(2 * merged.get("window", 5000))]
                else:
                    if merged.get("flank", False):
                        cols = [f"leftFlank_{i}" for i in range(merged.get("flank_bins", 50))] + \
                               [f"main_{i}" for i in range(merged.get("scaled_bins", 100))] + \
                               [f"rightFlank_{i}" for i in range(merged.get("flank_bins", 50))]
                    else:
                        cols = [f"bin_{i}" for i in range(merged.get("scaled_bins", 100))]
                with gzip.open(mat_path, "wt") as tsv:
                    tsv.write("interval_id\t" + "\t".join(cols) + "\n")
                    for i, row in enumerate(cov):
                        tsv.write(f"interval_{i}\t" + "\t".join("nan" if not np.isfinite(v) else f"{v:.5f}" for v in row) + "\n")
                logging.info(f"Wrote matrix: {mat_path}")

            # Average profile (raw)
            avg = np.nanmean(cov, axis=0) if cov.size else np.array([], dtype=float)
            if merged.get("mode") == "midpoint":
                xvals = np.arange(-merged.get("window", 5000), merged.get("window", 5000), 1)
            else:
                total_bins = merged.get("scaled_bins", 100) + (2*merged.get("flank_bins", 50) if merged.get("flank", False) else 0)
                xvals = np.arange(total_bins)

            avg_tsv = os.path.join(outdir, f"coverage_average_profile_{bed_part}_{bw_part}_{ts}.tsv")
            with open(avg_tsv, "w") as f:
                if merged.get("mode") == "midpoint":
                    f.write("rel_bp\tcoverage\n")
                else:
                    f.write("bin_index\tcoverage\n")
                for x, y in zip(xvals, avg):
                    f.write(f"{int(x)}\t{('nan' if not np.isfinite(y) else f'{y:.5f}')}\n")
            logging.info(f"Wrote average profile: {avg_tsv}")

            # QC plot per combo (smoothed overlay)
            ylabel = ("Signal (zero-centered)" if zero_centered_flags[wi]
                      else ("Fold diff" if merged.get("fold_diff", False) else "Coverage"))
            y_smooth = savgol_safe(avg, merged.get("savgol", True), merged.get("savgol_window", 101), merged.get("savgol_poly", 3))
            apply_font_rcparams(fonttype, font_size, title_size, label_size, tick_size, legend_size)
            plt.figure(figsize=(6, 4))
            plt.plot(xvals, avg, label="Raw mean", alpha=merged.get("raw_alpha", 0.3), linewidth=merged.get("raw_lw", 1.0), color=color)
            if y_smooth.size:
                plt.plot(xvals, y_smooth, label=f"Savgol (w={merged.get('savgol_window',101)}, p={merged.get('savgol_poly',3)})",
                         linewidth=merged.get("smooth_lw", 2.0), color=color)
            if merged.get("mode") == "midpoint":
                plt.axvline(0, color="black", linestyle="--", linewidth=1)
                plt.xlabel("Position relative to midpoint (bp)")
            else:
                plt.xlabel("Scaled bins")
            if zero_centered_flags[wi]:
                amax = max(abs(np.nanmin(avg)), abs(np.nanmax(avg)))
                if np.isfinite(amax) and amax > 0:
                    plt.ylim(-amax, amax)
            plt.ylabel(ylabel)
            plt.title("Aggregated Coverage")
            plt.legend(loc="best")
            plt.tight_layout()
            qc_pdf = os.path.join(outdir, f"metaplot_{bed_part}_{bw_part}_{ts}.pdf")
            plt.savefig(qc_pdf); plt.close()
            logging.info(f"QC metaplot saved: {qc_pdf}")

            # Grid data
            if merged.get("grid", False):
                if merged.get("savgol", True) and avg.size:
                    y_smooth_grid = savgol_safe(avg, True, merged.get("savgol_window", 101), merged.get("savgol_poly", 3))
                else:
                    y_smooth_grid = None

                if merged.get("grid_rows_axis", "bigwig") == "bigwig" or not overlay:
                    if merged.get("grid_rows_axis", "bigwig") == "bigwig":
                        row, col = wi, (beds + bedpes).index(bed)
                        title = f"{bw_labels[wi]} × {os.path.basename(bed)}"
                        legend_label = None
                    else:
                        row, col = (beds + bedpes).index(bed), wi
                        title = f"{os.path.basename(bed)} × {bw_labels[wi]}"
                        legend_label = bw_labels[wi] if overlay else None
                else:
                    row, col = (beds + bedpes).index(bed), 0
                    title = f"{os.path.basename(bed)} (overlay of {len(bigwigs)} tracks)"
                    legend_label = bw_labels[wi]

                grid_cells.append({
                    "row": row, "col": col, "x": xvals,
                    "y_raw": avg.copy() if avg.size else None,
                    "y_smooth": y_smooth_grid,
                    "title": title,
                    "color": color,
                    "raw_alpha": merged.get("raw_alpha", 0.3),
                    "raw_lw": merged.get("raw_lw", 1.0),
                    "smooth_lw": merged.get("smooth_lw", 2.0),
                    "symmetric_zero": zero_centered_flags[wi],
                    "legend_label": legend_label,
                    "layer": rank.get(wi, wi)
                })

    # ---------------- BEDPE processing ----------------
    bedpe_meta_all = {}
    for bedpe in bedpes:
        override_cols = tuple(merged.get("bedpe_cols")) if merged.get("bedpe_cols") else None
        override_str  = tuple(merged.get("bedpe_strand_cols")) if merged.get("bedpe_strand_cols") else None
        entries, meta = read_bedpe(bedpe, override_cols, override_str)
        bedpe_meta_all[bedpe] = meta
        if len(entries) == 0:
            logging.warning(f"{bedpe}: no valid pairs found; skipping.")
            continue

        # --- Pair‑scaled ---
        for wi, bw in enumerate(bigwigs):
            color = (bw_colors[wi] if (bw_colors and wi < len(bw_colors)) else
                     plt.rcParams['axes.prop_cycle'].by_key()['color'][wi % len(plt.rcParams['axes.prop_cycle'].by_key()['color'])])

            # chunk entries
            n = len(entries)
            chunk = (n + cpu - 1) // cpu
            subsets = [entries[i:i+chunk] for i in range(0, n, chunk)]
            tasks = [(s, bw,
                      merged.get("bedpe_main_bins", 200),
                      merged.get("bedpe_flank_left", 0.5),
                      merged.get("bedpe_flank_right", 0.5),
                      merged.get("bedpe_flank_bins", 100),
                      (wi in flip_always_set),
                      (wi in flip_minus_set),
                      pair_sign_from)
                     for s in subsets]
            mats = []; skipped_sum = 0
            if cpu > 1:
                with multiprocessing.Pool(processes=cpu) as pool:
                    results = pool.starmap(_process_bedpe_pair_subset, tasks)
                for M, sk in results:
                    mats.append(M); skipped_sum += sk
            else:
                for t in tasks:
                    M, sk = _process_bedpe_pair_subset(*t)
                    mats.append(M); skipped_sum += sk

            matrix = np.concatenate(mats, axis=0) if mats else np.zeros((0, merged.get("bedpe_main_bins", 200) + 2*merged.get("bedpe_flank_bins", 100)), dtype=float)

            # fold-diff normalization
            if merged.get("fold_diff", False):
                if zero_centered_flags[wi]:
                    logging.warning(f"{bw_labels[wi]} looks zero-centered; skipping --fold-diff for this track.")
                else:
                    gmean = np.nanmean(matrix)
                    if (not np.isfinite(gmean)) or (abs(gmean) < 1e-6):
                        logging.warning("Detected near-zero or non-finite global mean; skipping fold-diff.")
                    else:
                        matrix = matrix / gmean

            bed_part = _safe_join_name([bedpe + "_PAIR"])
            bw_part  = _safe_join_name([bw])

            # Write matrix
            if merged.get("write_matrix", True):
                cols = [f"leftPairFlank_{i}" for i in range(merged.get("bedpe_flank_bins", 100))] + \
                       [f"pairMain_{i}" for i in range(merged.get("bedpe_main_bins", 200))] + \
                       [f"rightPairFlank_{i}" for i in range(merged.get("bedpe_flank_bins", 100))]
                mat_path = os.path.join(outdir, f"coverage_matrix_{bed_part}_{bw_part}_{ts}.tsv.gz")
                with gzip.open(mat_path, "wt") as tsv:
                    tsv.write("pair_id\t" + "\t".join(cols) + "\n")
                    for i, row in enumerate(matrix):
                        tsv.write(f"pair_{i}\t" + "\t".join("nan" if not np.isfinite(v) else f"{v:.5f}" for v in row) + "\n")
                logging.info(f"Wrote BEDPE pair‑scaled matrix: {mat_path} (skipped inter‑chrom: {skipped_sum})")

            # Average profile
            avg = np.nanmean(matrix, axis=0) if matrix.size else np.array([], dtype=float)
            xvals = np.arange(matrix.shape[1]) if matrix.size else np.array([], dtype=int)
            avg_tsv = os.path.join(outdir, f"coverage_average_profile_{bed_part}_{bw_part}_{ts}.tsv")
            with open(avg_tsv, "w") as f:
                f.write("bin_index\tcoverage\n")
                for x, y in zip(xvals, avg):
                    f.write(f"{int(x)}\t{('nan' if not np.isfinite(y) else f'{y:.5f}')}\n")
            logging.info(f"Wrote BEDPE pair‑scaled average profile: {avg_tsv}")
            # QC
            ylabel = ("Signal (zero-centered)" if zero_centered_flags[wi]
                      else ("Fold diff" if merged.get("fold_diff", False) else "Coverage"))
            y_smooth = savgol_safe(avg, merged.get("savgol", True), merged.get("savgol_window", 101), merged.get("savgol_poly", 3))
            apply_font_rcparams(fonttype, font_size, title_size, label_size, tick_size, legend_size)
            plt.figure(figsize=(6, 4))
            plt.plot(xvals, avg, label="Raw mean", alpha=merged.get("raw_alpha", 0.3), linewidth=merged.get("raw_lw", 1.0), color=color)
            if y_smooth.size:
                plt.plot(xvals, y_smooth, label=f"Savgol (w={merged.get('savgol_window',101)}, p={merged.get('savgol_poly',3)})",
                         linewidth=merged.get("smooth_lw", 2.0), color=color)
            plt.xlabel("Scaled bins")
            plt.ylabel(ylabel)
            plt.title("Aggregated Coverage")
            plt.legend(loc="best")
            plt.tight_layout()
            qc_pdf = os.path.join(outdir, f"metaplot_{bed_part}_{bw_part}_{ts}.pdf")
            plt.savefig(qc_pdf); plt.close()
            logging.info(f"QC metaplot saved: {qc_pdf}")

            # Grid data
            if merged.get("grid", False):
                if merged.get("savgol", True) and avg.size:
                    y_smooth_grid = savgol_safe(avg, True, merged.get("savgol_window", 101), merged.get("savgol_poly", 3))
                else:
                    y_smooth_grid = None
                if merged.get("grid_rows_axis", "bigwig") == "bigwig" or not overlay:
                    if merged.get("grid_rows_axis", "bigwig") == "bigwig":
                        row, col = wi, (beds + bedpes).index(bedpe)
                        title = f"{bw_labels[wi]} × {os.path.basename(bedpe)} (pair)"
                        legend_label = None
                    else:
                        row, col = (beds + bedpes).index(bedpe), wi
                        title = f"{os.path.basename(bedpe)} (pair) × {bw_labels[wi]}"
                        legend_label = bw_labels[wi] if overlay else None
                else:
                    row, col = (beds + bedpes).index(bedpe), 0
                    title = f"{os.path.basename(bedpe)} (pair; overlay of {len(bigwigs)} tracks)"
                    legend_label = bw_labels[wi]
                grid_cells.append({
                    "row": row, "col": col, "x": xvals,
                    "y_raw": avg.copy() if avg.size else None,
                    "y_smooth": y_smooth_grid,
                    "title": title, "color": color,
                    "raw_alpha": merged.get("raw_alpha", 0.3),
                    "raw_lw": merged.get("raw_lw", 1.0),
                    "smooth_lw": merged.get("smooth_lw", 2.0),
                    "symmetric_zero": zero_centered_flags[wi],
                    "legend_label": legend_label,
                    "layer": rank.get(wi, wi)
                })

        # --- Anchors (midpoint windows) ---
        if merged.get("anchors_separate", False):
            anchor1_list = [e.anchor1() for e in entries]
            anchor2_list = [e.anchor2() for e in entries]
            if not merged.get("respect_bedpe_strand", False):
                for iv in anchor1_list:
                    iv.strand = None
                for iv in anchor2_list:
                    iv.strand = '-' if merged.get("flip_anchor2", False) else None

            for wi, bw in enumerate(bigwigs):
                color = (bw_colors[wi] if (bw_colors and wi < len(bw_colors)) else
                         plt.rcParams['axes.prop_cycle'].by_key()['color'][wi % len(plt.rcParams['axes.prop_cycle'].by_key()['color'])])

                # chunk anchors
                n1 = len(anchor1_list); n2 = len(anchor2_list)
                chunk1 = (n1 + cpu - 1) // cpu; chunk2 = (n2 + cpu - 1) // cpu
                subsets1 = [anchor1_list[i:i+chunk1] for i in range(0, n1, chunk1)]
                subsets2 = [anchor2_list[i:i+chunk2] for i in range(0, n2, chunk2)]

                tasks1 = [(s, bw, merged.get("window", 5000), merged.get("smoothing", 0),
                           merged.get("read_mode", "values"), (wi in flip_always_set), (wi in flip_minus_set))
                          for s in subsets1]
                tasks2 = [(s, bw, merged.get("window", 5000), merged.get("smoothing", 0),
                           merged.get("read_mode", "values"), (wi in flip_always_set), (wi in flip_minus_set))
                          for s in subsets2]

                if cpu > 1:
                    with multiprocessing.Pool(processes=cpu) as pool:
                        mats1 = pool.starmap(_process_bedpe_anchor_subset, tasks1)
                    with multiprocessing.Pool(processes=cpu) as pool:
                        mats2 = pool.starmap(_process_bedpe_anchor_subset, tasks2)
                else:
                    mats1 = [_process_bedpe_anchor_subset(*t) for t in tasks1]
                    mats2 = [_process_bedpe_anchor_subset(*t) for t in tasks2]

                m1 = np.concatenate(mats1, axis=0) if mats1 else np.zeros((0, 2*merged.get("window", 5000)), dtype=float)
                m2 = np.concatenate(mats2, axis=0) if mats2 else np.zeros((0, 2*merged.get("window", 5000)), dtype=float)

                for tag, M in (("ANCHOR1", m1), ("ANCHOR2", m2)):
                    bed_part = _safe_join_name([bedpe + f"_{tag}"])
                    bw_part  = _safe_join_name([bw])

                    if merged.get("fold_diff", False):
                        if zero_centered_flags[wi]:
                            logging.warning(f"{bw_labels[wi]} looks zero-centered; skipping --fold-diff for this track.")
                        else:
                            gmean = np.nanmean(M)
                            if (not np.isfinite(gmean)) or (abs(gmean) < 1e-6):
                                logging.warning("Detected near-zero or non-finite global mean; skipping fold-diff.")
                            else:
                                M = M / gmean

                    if merged.get("write_matrix", True):
                        cols = [str(i - merged.get("window", 5000)) for i in range(2 * merged.get("window", 5000))]
                        mat_path = os.path.join(outdir, f"coverage_matrix_{bed_part}_{bw_part}_{ts}.tsv.gz")
                        with gzip.open(mat_path, "wt") as tsv:
                            tsv.write("interval_id\t" + "\t".join(cols) + "\n")
                            for i, row in enumerate(M):
                                tsv.write(f"interval_{i}\t" + "\t".join("nan" if not np.isfinite(v) else f"{v:.5f}" for v in row) + "\n")
                        logging.info(f"Wrote BEDPE {tag} midpoint matrix: {mat_path}")

                    avg = np.nanmean(M, axis=0) if M.size else np.array([], dtype=float)
                    xvals = np.arange(-merged.get("window", 5000), merged.get("window", 5000), 1)

                    avg_tsv = os.path.join(outdir, f"coverage_average_profile_{bed_part}_{bw_part}_{ts}.tsv")
                    with open(avg_tsv, "w") as f:
                        f.write("rel_bp\tcoverage\n")
                        for x, y in zip(xvals, avg):
                            f.write(f"{int(x)}\t{('nan' if not np.isfinite(y) else f'{y:.5f}')}\n")

                    ylabel = ("Signal (zero-centered)" if zero_centered_flags[wi] else ("Fold diff" if merged.get("fold_diff", False) else "Coverage"))
                    y_smooth = savgol_safe(avg, merged.get("savgol", True), merged.get("savgol_window", 101), merged.get("savgol_poly", 3))
                    apply_font_rcparams(fonttype, font_size, title_size, label_size, tick_size, legend_size)
                    plt.figure(figsize=(6, 4))
                    plt.plot(xvals, avg, label="Raw mean", alpha=merged.get("raw_alpha", 0.3), linewidth=merged.get("raw_lw", 1.0), color=color)
                    if y_smooth.size:
                        plt.plot(xvals, y_smooth, label=f"Savgol (w={merged.get('savgol_window',101)}, p={merged.get('savgol_poly',3)})",
                                 linewidth=merged.get("smooth_lw", 2.0), color=color)
                    plt.axvline(0, color="black", linestyle="--", linewidth=1)
                    plt.xlabel("Position relative to midpoint (bp)")
                    plt.ylabel(ylabel)
                    plt.title("Aggregated Coverage")
                    plt.legend(loc="best")
                    plt.tight_layout()
                    qc_pdf = os.path.join(outdir, f"metaplot_{bed_part}_{bw_part}_{ts}.pdf")
                    plt.savefig(qc_pdf); plt.close()
                    logging.info(f"QC metaplot saved: {qc_pdf}")

                    if merged.get("grid", False):
                        if merged.get("savgol", True) and avg.size:
                            y_smooth_grid = savgol_safe(avg, True, merged.get("savgol_window", 101), merged.get("savgol_poly", 3))
                        else:
                            y_smooth_grid = None
                        if merged.get("grid_rows_axis", "bigwig") == "bigwig" or not overlay:
                            if merged.get("grid_rows_axis", "bigwig") == "bigwig":
                                row, col = wi, (beds + bedpes).index(bedpe)
                                title = f"{bw_labels[wi]} × {os.path.basename(bedpe)} ({tag.lower()})"
                                legend_label = None
                            else:
                                row, col = (beds + bedpes).index(bedpe), wi
                                title = f"{os.path.basename(bedpe)} ({tag.lower()}) × {bw_labels[wi]}"
                                legend_label = f"{bw_labels[wi]} ({'A1' if tag=='ANCHOR1' else 'A2'})" if overlay else None
                        else:
                            row, col = (beds + bedpes).index(bedpe), 0
                            title = f"{os.path.basename(bedpe)} ({tag.lower()}; overlay of {len(bigwigs)} tracks)"
                            legend_label = f"{bw_labels[wi]} ({'A1' if tag=='ANCHOR1' else 'A2'})"
                        grid_cells.append({
                            "row": row, "col": col, "x": xvals,
                            "y_raw": avg.copy() if avg.size else None,
                            "y_smooth": y_smooth_grid,
                            "title": title, "color": color,
                            "raw_alpha": merged.get("raw_alpha", 0.3),
                            "raw_lw": merged.get("raw_lw", 1.0),
                            "smooth_lw": merged.get("smooth_lw", 2.0),
                            "symmetric_zero": zero_centered_flags[wi],
                            "legend_label": legend_label,
                            "layer": rank.get(wi, wi)
                        })

    # Render grid if requested
    if merged.get("grid", False) and len(grid_cells) > 0:
        if all(zero_centered_flags):
            ylabel = "Signal (zero-centered)"
        elif any(zero_centered_flags):
            ylabel = "Signal"
        else:
            ylabel = "Fold diff" if merged.get("fold_diff", False) else "Coverage"
        xlabel = "Position relative to midpoint (bp)" if merged.get("mode") == "midpoint" else "Scaled bins"
        grid_base = os.path.join(outdir, f"metaplot_grid_{nrows}x{ncols}_{ts}")
        plot_grid(grid_cells,
                  nrows=nrows, ncols=ncols,
                  mode=(merged.get("mode") if len(beds) > 0 else "scaled"),
                  out_base=grid_base,
                  plot_format=merged.get("plot_format", "pdf"),
                  lock_y=merged.get("lock_y", "none"),
                  fonttype=fonttype,
                  dpi=merged.get("dpi", 300),
                  xlabel=xlabel, ylabel=ylabel,
                  font_size=font_size, title_size=title_size, label_size=label_size, tick_size=tick_size, legend_size=legend_size)

    # Final metadata supplement
    extras = {"bed_stats": bed_stats_all, "bedpe_meta": bedpe_meta_all}
    _write_metadata(outdir, ts, args, merged, extra=extras)

    logging.info("pileup: Done.")


# ---------------------------------- boxgrid -----------------------------------

def summarize_set_over_bw(bw_path: str,
                          bed_intervals: Optional[List[IntervalWithStrand]] = None,
                          bed_mode: Optional[str] = None,
                          window: int = 5000,
                          scaled_bins: int = 100,
                          flank: bool = False,
                          flank_bins: int = 50,
                          bedpe_entries: Optional[List[BedPeEntry]] = None,
                          bedpe_mode: Optional[str] = None,  # "pair" or "anchors"
                          bedpe_main_bins: int = 200,
                          bedpe_flank_bins: int = 100,
                          bedpe_flank_left: float = 0.5,
                          bedpe_flank_right: float = 0.5,
                          flip_anchor2: bool = False,
                          respect_bedpe_strand: bool = False,
                          core_fraction: float = 1.0,
                          flip_sign_always: bool = False,
                          flip_sign_if_minus: bool = False,
                          pair_sign_from: str = "none") -> float:
    """
    Compute the mean signal for a dataset over a bigWig according to a chosen
    strategy. Applies sign‑flip options analogously to pileup.
    Returns the average signal (not yet transformed).
    """
    if bed_intervals is not None:
        if bed_mode == "midpoint":
            M = compute_midpoint_matrix(bed_intervals, bw_path, window, read_mode="stats")
            if flip_sign_always and M.size:
                M = -M
            if flip_sign_if_minus and M.size:
                signs = np.array([(-1.0 if iv.strand == '-' else 1.0) for iv in bed_intervals], dtype=float)
                M = M * signs[:, None]
            cols = M.shape[1]
            half = cols // 2
            core_half = max(1, int(round(core_fraction * half)))
            sel = slice(half - core_half, half + core_half)
            return float(np.nanmean(M[:, sel])) if M.size else 0.0
        elif bed_mode == "scaled":
            M = compute_scaled_matrix(bed_intervals, bw_path, scaled_bins, flank_bins, flank)
            if flip_sign_always and M.size:
                M = -M
            if flip_sign_if_minus and M.size:
                signs = np.array([(-1.0 if iv.strand == '-' else 1.0) for iv in bed_intervals], dtype=float)
                M = M * signs[:, None]
            if flank:
                sel = slice(flank_bins, flank_bins + scaled_bins)
                return float(np.nanmean(M[:, sel])) if M.size else 0.0
            else:
                return float(np.nanmean(M)) if M.size else 0.0

    if bedpe_entries is not None:
        if bedpe_mode == "pair":
            M, _ = compute_bedpe_pair_scaled_matrix(bedpe_entries, bw_path,
                                                    main_bins=bedpe_main_bins,
                                                    flank_frac_left=bedpe_flank_left,
                                                    flank_frac_right=bedpe_flank_right,
                                                    flank_bins=bedpe_flank_bins)
            if M.size:
                if flip_sign_always:
                    M = -M
                if flip_sign_if_minus and pair_sign_from != "none":
                    if pair_sign_from == "anchor1":
                        signs = np.array([(-1.0 if e.strand1 == '-' else 1.0) for e in bedpe_entries], dtype=float)
                    else:
                        signs = np.array([(-1.0 if e.strand2 == '-' else 1.0) for e in bedpe_entries], dtype=float)
                    M = M * signs[:, None]
            if M.size == 0:
                return 0.0
            sel = slice(bedpe_flank_bins, bedpe_flank_bins + bedpe_main_bins)
            return float(np.nanmean(M[:, sel]))
        elif bedpe_mode == "anchors":
            a1 = [e.anchor1() for e in bedpe_entries]
            a2 = [e.anchor2() for e in bedpe_entries]
            if not respect_bedpe_strand:
                for iv in a1:
                    iv.strand = None
                for iv in a2:
                    iv.strand = '-' if flip_anchor2 else None
            M1 = compute_midpoint_matrix(a1, bw_path, window, read_mode="stats")
            M2 = compute_midpoint_matrix(a2, bw_path, window, read_mode="stats")
            if flip_sign_always:
                M1 = -M1; M2 = -M2
            if flip_sign_if_minus:
                s1 = np.array([(-1.0 if iv.strand == '-' else 1.0) for iv in a1], dtype=float)
                s2 = np.array([(-1.0 if iv.strand == '-' else 1.0) for iv in a2], dtype=float)
                M1 = M1 * s1[:, None]; M2 = M2 * s2[:, None]
            return float(np.nanmean(np.concatenate([M1, M2], axis=0))) if (M1.size or M2.size) else 0.0

    return 0.0


def run_boxgrid(args: argparse.Namespace):
    ts = _now_timestamp()
    outdir = _create_outdir(args.output_dir, args.beds + args.bedpes, args.bigwigs, ts)
    _write_metadata(outdir, ts, args, vars(args))

    # Prepare datasets
    bed_sets: Dict[str, Tuple[List[IntervalWithStrand], Dict[str, int]]] = {}
    for b in args.beds:
        bed_sets[b] = read_bed_with_strand(b)
    bedpe_sets: Dict[str, Tuple[List[BedPeEntry], Dict[str, Any]]] = {}
    for bp in args.bedpes:
        override_cols = tuple(args.bedpe_cols) if args.bedpe_cols else None
        override_str  = tuple(args.bedpe_strand_cols) if args.bedpe_strand_cols else None
        bedpe_sets[bp] = read_bedpe(bp, override_cols, override_str)

    # Zero‑centered detection / overrides
    if args.zero_centered_detect == "auto":
        zc_info = [detect_zero_centered_bigwig(bw) for bw in args.bigwigs]
        zero_centered_flags = [z["zero_centered"] for z in zc_info]
    else:
        zc_info = [{"zero_centered": False} for _ in args.bigwigs]
        zero_centered_flags = [False] * len(args.bigwigs)
    labels = [os.path.basename(bw) for bw in args.bigwigs]
    force_list = args.force_zero_centered or []
    for idx, base in enumerate(labels):
        if base in force_list:
            zero_centered_flags[idx] = True

    # Sign‑flip selection per bigWig
    flip_always_set = resolve_track_selectors(args.flip_sign, args.bigwigs, labels)
    flip_minus_set  = resolve_track_selectors(args.flip_sign_if_minus, args.bigwigs, labels)
    pair_sign_from  = args.bedpe_pair_sign_from

    # Compute matrix [rows=datasets, cols=bigwigs] of summary values
    datasets = list(bed_sets.keys()) + list(bedpe_sets.keys())
    values = np.zeros((len(datasets), len(args.bigwigs)), dtype=float)

    # Precompute bigWig genome‑wide means for non‑zero‑centered tracks
    bw_means = [bigwig_global_mean(bw) for bw in args.bigwigs]

    for i, ds in enumerate(datasets):
        for j, bw in enumerate(args.bigwigs):
            if ds in bed_sets:
                intervals, _ = bed_sets[ds]
                mean_sig = summarize_set_over_bw(
                    bw_path=bw, bed_intervals=intervals,
                    bed_mode=("midpoint" if args.mode in ("midpoint", "bedpe_pair", "bedpe_anchors") else "scaled"),
                    window=args.window, scaled_bins=args.scaled_bins,
                    flank=args.flank, flank_bins=args.flank_bins,
                    core_fraction=args.core_fraction,
                    flip_sign_always=(j in flip_always_set),
                    flip_sign_if_minus=(j in flip_minus_set),
                    pair_sign_from="none"
                )
            else:
                entries, _ = bedpe_sets[ds]
                if args.mode == "bedpe_pair":
                    mean_sig = summarize_set_over_bw(
                        bw_path=bw, bedpe_entries=entries, bedpe_mode="pair",
                        bedpe_main_bins=args.bedpe_main_bins, bedpe_flank_bins=args.bedpe_flank_bins,
                        bedpe_flank_left=args.bedpe_flank_left, bedpe_flank_right=args.bedpe_flank_right,
                        flip_sign_always=(j in flip_always_set),
                        flip_sign_if_minus=(j in flip_minus_set),
                        pair_sign_from=pair_sign_from
                    )
                else:
                    mean_sig = summarize_set_over_bw(
                        bw_path=bw, bedpe_entries=entries, bedpe_mode="anchors",
                        window=args.window, flip_anchor2=args.flip_anchor2,
                        respect_bedpe_strand=args.respect_bedpe_strand,
                        flip_sign_always=(j in flip_always_set),
                        flip_sign_if_minus=(j in flip_minus_set),
                        pair_sign_from="none"
                    )

            if zero_centered_flags[j]:
                values[i, j] = mean_sig  # mean signed signal
            else:
                baseline = bw_means[j]
                if baseline <= 0 or not np.isfinite(baseline):
                    values[i, j] = np.nan
                else:
                    values[i, j] = math.log2((mean_sig + EPS) / (baseline + EPS))

    # Plot heatmap
    apply_font_rcparams(args.fonttype, args.font_size, args.title_size, args.label_size, args.tick_size, args.legend_size)
    cmap = metallic_bwr_cmap()
    vmax = float(np.nanmax(np.abs(values))) if values.size else 1.0
    vmax = max(vmax, 0.5)
    vmin = -vmax

    fig, ax = plt.subplots(figsize=(1.2*len(args.bigwigs) + 3, 0.5*len(datasets) + 2))
    im = ax.imshow(values, aspect="auto", cmap=cmap, vmin=vmin, vmax=vmax)
    ax.set_xticks(np.arange(len(args.bigwigs)))
    ax.set_xticklabels([os.path.basename(x) for x in args.bigwigs], rotation=45, ha="right")
    ax.set_yticks(np.arange(len(datasets)))
    ax.set_yticklabels([os.path.basename(x) for x in datasets])
    if all(zero_centered_flags):
        title = "mean signal (zero‑centered units)"
        cbar_label = "mean (zero‑centered)"
    elif any(zero_centered_flags):
        title = "mixed metrics (zero‑centered tracks show mean; others log2 fold)"
        cbar_label = "value"
    else:
        title = "log2 fold over genome‑wide mean"
        cbar_label = args.cbar_label
    ax.set_title(title, fontsize=args.title_size)
    cbar = fig.colorbar(im, ax=ax, fraction=0.046, pad=0.04)
    cbar.set_label(cbar_label)
    fig.tight_layout()

    base = os.path.join(outdir, f"boxgrid_{len(datasets)}x{len(args.bigwigs)}_{ts}")
    if args.plot_format in ("pdf", "both"):
        fig.savefig(f"{base}.pdf")
        logging.info(f"Boxgrid saved: {base}.pdf")
    if args.plot_format in ("png", "both"):
        fig.savefig(f"{base}.png", dpi=args.dpi)
        logging.info(f"Boxgrid saved: {base}.png")
    plt.close(fig)

    logging.info("boxgrid: Done.")


# ------------------------------------ main ------------------------------------

def main():
    _setup_logging()
    parser = build_parser()
    args = parser.parse_args()
    if args.subcmd == "pileup":
        run_pileup(args)
    elif args.subcmd == "boxgrid":
        run_boxgrid(args)
    else:
        raise SystemExit("Unknown subcommand.")


if __name__ == "__main__":
    main()
```
