## Supplementary material for "CAD-C: An engineered nuclease enables repair-free *in situ* proximity ligation and nucleosome-resolution chromosome walks in human cells": Custom code: v10p6_auto_ZAWBbin1_RAMPAGE_all_20250927.sh

```
#!/bin/bash
#SBATCH -J metacov_midpoint_1bw_1bed
#SBATCH -o metacov_midpoint_1bw_1bed.%j.out
#SBATCH -e metacov_midpoint_1bw_1bed.%j.err
#SBATCH -c 8

set -euo pipefail

# Paths
SCRIPT=/rugpfs/fs0/risc_lab/scratch/jsoroczyns/CAD/CAD-C/scripts/2025_09_27/smart_bigwig_over_bed_v10p6_20250927.py
BED=/rugpfs/fs0/risc_lab/scratch/jsoroczyns/CAD/CAD-C/Pileups_2025/GM12878_annotations/ENCFF591KXZ.bed
# AWB (bin1, robust-Z per chromosome)
BW=/lustre/fs4/risc_lab/scratch/jsoroczyns/CAD/CADwalk/CADwalk_E1_GM12878_2x250/results_20250207_0034_nonexpanded/ABC_walk_processing_20250209/20250214_Bfilt_flattened_parse2_UU_direct_ABC_complete_consequent12/filtzerolenB_flip_sort_dedup_split_20250214/merged_all/WPI_AWB_analyses_2025/wpi_awb_20250927_210158/Z_AWB_bin1.bw

OUTDIR=/rugpfs/fs0/risc_lab/scratch/jsoroczyns/CAD/CADwalk/CADwalk_E1_GM12878_2x250/results_20250207_0034_nonexpanded/ABC_walk_processing_20250209/20250214_Bfilt_flattened_parse2_UU_direct_ABC_complete_consequent12/filtzerolenB_flip_sort_dedup_split_20250214/merged_all/WPI_analyses_2025/wpi_20250926_190935/Z_WPI_BW_Pileups_2025/RAMPAGE_2500/
mkdir -p "$OUTDIR" logs

NPROC=${SLURM_CPUS_PER_TASK:-8}

# IMPORTANT: turn OFF zero-centering detection; do NOT force zero-centering.
python "$SCRIPT" pileup \
  -b "$BED" \
  -w "$BW" \
  -o "$OUTDIR" \
  -p "$NPROC" \
  --mode midpoint \
  --window 2500 \
  --zero-centered-detect off \
  --bw-labels Z_AWB_bin1 \
  --savgol --savgol-window 101 --savgol-poly 3 \
  --plot-format pdf \
  --fonttype 42 \
  --font-size 11 --title-size 13 --label-size 12 --tick-size 10 --legend-size 10
```
