## Supplementary material for "CAD-C: An engineered nuclease enables repair-free *in situ* proximity ligation and nucleosome-resolution chromosome walks in human cells": Custom code: v10p6_ZAWBbin1_NEWfromv5generator_Ibw_over_top5CTCF_2500_DONTflipZscore_if_CTCFstrandnegative_20250927.sh

```
#!/bin/bash
#SBATCH -J metacov_midpoint_1bw_1bed
#SBATCH -o metacov_midpoint_1bw_1bed.%j.out
#SBATCH -e metacov_midpoint_1bw_1bed.%j.err
#SBATCH -c 8

set -euo pipefail

# --- Environment (adapt to your cluster) ---
# module purge
# module load python/3.10
# If you use conda:
# source ~/miniconda3/etc/profile.d/conda.sh
# conda activate metacov

# --- Paths ---
# NOTE: updated to your new dated script with sign-flip support (v10p6).
SCRIPT=/rugpfs/fs0/risc_lab/scratch/jsoroczyns/CAD/CAD-C/scripts/2025_09_27/smart_bigwig_over_bed_v10p6_20250927.py
BED=/rugpfs/fs0/risc_lab/scratch/jsoroczyns/CAD/CHOMP/2025_CHOMP_bigwig_pileups/GM12878_bed_annotations/GM12878_top_5_percent_CTCF_motifs.bed
BW=/lustre/fs4/risc_lab/scratch/jsoroczyns/CAD/CADwalk/CADwalk_E1_GM12878_2x250/results_20250207_0034_nonexpanded/ABC_walk_processing_20250209/20250214_Bfilt_flattened_parse2_UU_direct_ABC_complete_consequent12/filtzerolenB_flip_sort_dedup_split_20250214/merged_all/WPI_AWB_analyses_2025/wpi_awb_20250927_210158/Z_AWB_bin1.bw    # robust Z-score (zero-centered, non-directional; DO NOT flip sign)

# Kept EXACTLY as you provided.
OUTDIR=/rugpfs/fs0/risc_lab/scratch/jsoroczyns/CAD/CADwalk/CADwalk_E1_GM12878_2x250/results_20250207_0034_nonexpanded/ABC_walk_processing_20250209/20250214_Bfilt_flattened_parse2_UU_direct_ABC_complete_consequent12/filtzerolenB_flip_sort_dedup_split_20250214/merged_all/WPI_analyses_2025/wpi_20250926_190935/Z_WPI_BW_Pileups_2025/top5CTCF_750bp
mkdir -p "$OUTDIR" logs

NPROC=${SLURM_CPUS_PER_TASK:-8}

# Label used for legend/overrides (force zero-centered, flip-selectors, etc.)
LABEL=ZSCORE

python "$SCRIPT" pileup \
  -b "$BED" \
  -w "$BW" \
  -o "$OUTDIR" \
  -p "$NPROC" \
  --mode midpoint \
  --window 2500 \
  --zero-centered-detect auto \
  --bw-labels "$LABEL" \
  --force-zero-centered "$LABEL" \
  --plot-format pdf \
  --fonttype 42 \
  --font-size 11 --title-size 13 --label-size 12 --tick-size 10 --legend-size 10

# Outputs:
#   - coverage_matrix_<bed>_<bw>_<ts>.tsv.gz
#   - coverage_average_profile_<bed>_<bw>_<ts>.tsv  (columns: rel_bp, coverage)
#   - metaplot_<bed>_<bw>_<ts>.pdf
# Metadata + env/packages written to run_metadata_<ts>.txt in a timestamped OUTDIR.
```
