## Supplementary material for "CAD-C: An engineered nuclease enables repair-free *in situ* proximity ligation and nucleosome-resolution chromosome walks in human cells": Custom code: WPI_AWB_allinone_v5_20250927.py

```
#!/usr/bin/env python3
# =========================================================================================
#  WPI_AWB_allinone_v4p1_20250927.py
#  -----------------------------------------------------------------------------------------
#  ALL-IN-ONE:
#    • Compute signed (WPI) and magnitude-only (AWB) bend metrics from 3-fragment CAD-walks.
#    • Emit both bin1 (B_mid sites; 1-bp sparse) and 1 kb binned (fixed-grid) tables.
#    • Per-chromosome robust-Z for: Z_WPI (signed) and Z_AWB (magnitude-only).
#    • BigWigs:
#         - bin1:  Z_WPI_bin1.bw   and   Z_AWB_bin1.bw        (alias: zWPI_bin1.bw → copy of Z_WPI_bin1.bw)
#         - 1 kb:  Z_WPI_1kb.bw    and   Z_AWB_1kb.bw        (aliases: 1kb_bin_zWPI.bw, 1kb_bin_AWB.bw)
#    • Correlation QC (1 kb): Z_WPI vs Z_AWB — PDF (fonttype=42), robust axes, Pearson/Spearman;
#      exact replotting data saved as .npz (2D histogram + edges + axes + stats).
#
#  Version : 4.1   •   Date : 2025-09-27
#  Author  : Jan Soroczynski (Risca Lab, The Rockefeller University)
#
#  WHAT'S NEW IN v4p1 (robust zero-centering)
#  -----------------------------------------
#  • Per-chrom winsorization before Z-scaling (--winsorize-q, default 0.005).
#  • Robust fallback when MAD is ~0 or below floor (--z-mad-floor, default 1e-9):
#      scale = IQR/1.349 (default), or STD, or 1 (none) via --z-fallback {iqr,std,none}.
#  • Records per-chrom median/scale and whether fallback was used into metadata.
#
#  KEY CONVENTIONS
#  ---------------
#  • WPI (Walk Polarity Index):  WPI = s * F ∈ [-1,1]
#       F = 1 - AC/(AB+BC)   (clamped to [0,1]; computed only where AB+BC>0)
#       s = sign( (A+C)/2 - B )   (+ upstream, 0 midpoint, − downstream)
#  • AWB (Absolute Walk Bend):  AWB = F ∈ [0,1]   (direction-agnostic bend magnitude)
#  • Robust Z per chromosome (no detrending):  Z = (x - median) / scale,
#       where scale = MAD*1.4826 (robust) or fallback as above.
#  • Coordinates:
#       - bin1 (sites): per-site means at B_mid (1-bp intervals [B_mid, B_mid+1))
#       - 1 kb bins:   aggregate B_mid sites into Start=k*bin_size, End=Start+bin_size
#
#  WHY DEFAULTS DISABLE COVERAGE FILTERS
#  -------------------------------------
#  BEI / Z_ACd aggregate three midpoints per walk (A,B,C). WPI/AWB aggregate only B (one per walk).
#  Using a default min_cov_1kb=50 would prune far too many WPI/AWB bins. Defaults keep all:
#      --min_cov_site=1   and   --min_cov_1kb=1
#
#  WHEN FILTERING HAPPENS (IMPORTANT FOR Z-SCORES)
#  -----------------------------------------------
#  Any coverage filtering (if thresholds >1) is applied AFTER per-site/per-bin mean aggregation
#  but BEFORE per-chrom robust Z. Thus only kept rows contribute to each chromosome’s
#  median/scale used for Z-scaling (“Signal (zero-centered)” convention).
#
#  INPUT
#  -----
#  A tab-delimited CAD-walk file (.tsv/.tsv.gz; header present; leading '#' comments OK)
#  with columns:  A_chrom, A_mid, B_mid, C_mid.  Walks are pre-oriented (A upstream of C).
#
#  OUTPUTS (timestamped subdir under --outdir)
#  -------------------------------------------
#    A) BIN1 (sites):
#       • wpi_awb_sites_<ts>.tsv.gz   (Chrom, Pos, WPI_mean_site, AWB_mean_site, n, Z_WPI_site, Z_AWB_site)
#       • Z_WPI_bin1.bw,  Z_AWB_bin1.bw,  zWPI_bin1.bw (copy)
#    B) 1 KB:
#       • wpi_awb_1kb_<ts>.tsv.gz     (Chrom, Start, End, WPI_mean_1kb, AWB_mean_1kb, n_walks, n_sites, Z_WPI_1kb, Z_AWB_1kb)
#       • Z_WPI_1kb.bw,  Z_AWB_1kb.bw,  1kb_bin_zWPI.bw, 1kb_bin_AWB.bw (copies)
#    C) Correlation (1 kb):
#       • corr_Z_WPI_vs_Z_AWB_1kb_<ts>.pdf / .npz
#       • corr_stats_<ts>.txt
#
#  CLI (new flags in v4p1 are marked ★)
#  ---
#    --walks          path/to/walks.tsv(.gz)                     [required]
#    --outdir         output root directory                      [required]
#    --fai            genome FASTA index (.fai)                  [required]
#    --bin_size       1000                                       [default]
#    --chunk_rows     8000000                                    [default]
#    --min_cov_site   1                                          [default: keep all]
#    --min_cov_1kb    1                                          [default: keep all]
#    --exclude_random                                       
#    --chrom          chrN
#    --region         chr:start-end
#    --z_axis_cap     10.0
#    --z_axis_q       0.995
#    --hexbin_bins    200
#    --plot_bin1
#    --bin1_sample    2000000
#    ★ --winsorize-q  0.005     # 0 disables winsorization
#    ★ --z-mad-floor  1e-9      # floor for MAD/scale to avoid blow-up
#    ★ --z-fallback   iqr       # {iqr,std,none} fallback scale if MAD<floor
#
#  DEPENDENCIES
#    numpy • pandas • pyBigWig • matplotlib (PDF fonttype=42)
#    optional: statsmodels (robust.mad), scipy (Pearson/Spearman)
# =========================================================================================
from __future__ import annotations
import argparse, gzip, json, logging, hashlib, sys, math
from datetime import datetime
from pathlib import Path
from typing import Dict, List, Tuple, Literal

import numpy as np
import pandas as pd

# Optional deps
try:
    import pyBigWig
except Exception:
    pyBigWig = None

# robust MAD (statsmodels optional)
try:
    import importlib.util
    _sm = importlib.util.find_spec("statsmodels")
    if _sm is not None:
        from statsmodels import robust
        def mad(a): return robust.mad(a, c=1.4826)  # already scaled to robust σ
    else:
        raise ImportError
except Exception:
    def mad(a, c: float = 1.4826) -> float:
        a = np.asarray(a, dtype=float)
        med = np.nanmedian(a)
        return c * np.nanmedian(np.abs(a - med))

# plotting
try:
    import matplotlib
    matplotlib.use("Agg")
    import matplotlib.pyplot as plt
    from matplotlib import rcParams
    rcParams["pdf.fonttype"] = 42
    rcParams["ps.fonttype"]  = 42
except Exception:
    plt = None

# correlations
try:
    import scipy.stats as ss
    HAVE_SCIPY = True
except Exception:
    HAVE_SCIPY = False

pd.options.mode.copy_on_write = True

# ---------------- helpers ------------------------------------------------------
def sha256_of_file(p: Path) -> str:
    h = hashlib.sha256(); h.update(p.read_bytes()); return h.hexdigest()

def parse_region(arg: str) -> Tuple[str, int, int]:
    chrom, rest = arg.split(":")
    start, end  = rest.replace(",", "").split("-")
    return chrom, int(float(start)), int(float(end))

def ensure_dir(p: Path): p.mkdir(parents=True, exist_ok=True)

def read_fai(path: str) -> Dict[str, int]:
    d: Dict[str, int] = {}
    with open(path) as fh:
        for line in fh:
            chrom, length, *_ = line.split()
            d[chrom] = int(length)
    return d

def write_bw_fixed(out_path: Path, chrom_sizes: Dict[str, int],
                   df: pd.DataFrame, col: str, bin_size: int):
    if pyBigWig is None:
        raise RuntimeError("pyBigWig not available")
    bw = pyBigWig.open(str(out_path), "w")
    header = list(chrom_sizes.items()); bw.addHeader(header)
    for chrom, chrom_len in header:
        sub = df[df["Chrom"] == chrom][["Start", col]].drop_duplicates("Start").sort_values("Start")
        if sub.empty: bw.addEntries([chrom],[0],[1],[0.0]); continue
        s = sub["Start"].astype(np.int64).values
        e = s + bin_size
        v = sub[col].astype(float).values
        m = (s>=0) & (e<=chrom_len) & np.isfinite(v)
        s,e,v = s[m], e[m], v[m]
        if len(v)==0: bw.addEntries([chrom],[0],[1],[0.0]); continue
        bw.addEntries([chrom]*len(v), s.tolist(), e.tolist(), v.tolist())
    bw.close()

def write_bw_sparse_sites(out_path: Path, chrom_sizes: Dict[str, int],
                          df: pd.DataFrame, col: str):
    if pyBigWig is None:
        raise RuntimeError("pyBigWig not available")
    bw = pyBigWig.open(str(out_path), "w")
    header = list(chrom_sizes.items()); bw.addHeader(header)
    for chrom, chrom_len in header:
        sub = df[df["Chrom"] == chrom][["Pos", col]].drop_duplicates("Pos")
        if sub.empty: bw.addEntries([chrom],[0],[1],[0.0]); continue
        s = sub["Pos"].astype(np.int64).values
        e = s + 1
        v = sub[col].astype(float).values
        m = (s>=0) & (e<=chrom_len) & np.isfinite(v)
        s,e,v = s[m], e[m], v[m]
        if len(v)==0: bw.addEntries([chrom],[0],[1],[0.0]); continue
        bw.addEntries([chrom]*len(v), s.tolist(), e.tolist(), v.tolist())
    bw.close()

# ---------------- robust zero-centering utilities (v4p1) ----------------------
def _winsorize_series(s: pd.Series, q: float) -> pd.Series:
    if q <= 0.0:  # disabled
        return s
    lo, hi = s.quantile(q), s.quantile(1.0 - q)
    return s.clip(lower=lo, upper=hi)

def _fallback_scale(s: pd.Series, method: Literal["iqr","std","none"]) -> float:
    if method == "iqr":
        q75, q25 = s.quantile(0.75), s.quantile(0.25)
        scale = float(q75 - q25) / 1.349 if np.isfinite(q75) and np.isfinite(q25) else np.nan
    elif method == "std":
        scale = float(s.std(ddof=0))
    else:
        scale = 1.0
    if not np.isfinite(scale) or scale == 0.0:
        scale = 1.0
    return scale

def robust_center_scale_per_chrom(df: pd.DataFrame, col: str, chrom_col: str,
                                  winsor_q: float, mad_floor: float,
                                  fallback: Literal["iqr","std","none"]="iqr"
                                  ) -> Tuple[pd.Series, Dict[str, Dict[str, float]]]:
    """
    Return Z-scaled series per chromosome + per-chrom stats.
    Steps per chrom:
      1) (optional) winsorize at [q, 1-q]
      2) center at median
      3) scale by MAD*1.4826; if < mad_floor or non-finite → fallback (IQR/1.349 by default)
    """
    z = pd.Series(index=df.index, dtype=float)
    stats: Dict[str, Dict[str, float]] = {}
    for chrom, sub in df.groupby(chrom_col, sort=False):
        s = pd.to_numeric(sub[col], errors="coerce")
        s = s[np.isfinite(s)]
        if s.empty:
            # leave NaNs
            stats[chrom] = {"n": 0, "median": np.nan, "mad": np.nan, "scale": np.nan, "fallback": 0.0}
            continue
        s_w = _winsorize_series(s, winsor_q)
        med = float(s_w.median())
        mad_val = float(mad(s_w.values))
        use_fallback = False
        if (not np.isfinite(mad_val)) or (mad_val < mad_floor):
            scale = _fallback_scale(s_w, fallback)
            use_fallback = True
        else:
            scale = mad_val
        if (not np.isfinite(scale)) or (scale < mad_floor):
            scale = 1.0
            use_fallback = True
        # write back Z = (winsorized - med)/scale, aligned to original group indices
        z.loc[s_w.index] = (s_w - med) / scale
        stats[chrom] = {"n": int(s_w.size), "median": med, "mad": mad_val, "scale": float(scale),
                        "fallback": 1.0 if use_fallback else 0.0}
    return z, stats

def merge_stats(prefix: str, stats: Dict[str, Dict[str, float]]) -> Dict[str, Dict[str, float]]:
    return {f"{prefix}::{chrom}": vals for chrom, vals in stats.items()}

# ---------------- main ---------------------------------------------------------
def main():
    ap = argparse.ArgumentParser("WPI/AWB all-in-one (bin1 + 1kb) with robust-Z, BigWigs, and correlation PDF",
                                 formatter_class=argparse.ArgumentDefaultsHelpFormatter)
    ap.add_argument("--walks", required=True, help="Input walks .tsv or .tsv.gz with A_chrom,A_mid,B_mid,C_mid")
    ap.add_argument("--outdir", required=True, help="Output root directory (timestamped subdir will be created)")
    ap.add_argument("--fai", required=True, help="Genome FASTA index (.fai)")
    ap.add_argument("--bin_size", type=int, default=1000, help="Bin size for fixed-grid outputs")
    ap.add_argument("--chunk_rows", type=int, default=8_000_000, help="Rows per stream chunk")
    ap.add_argument("--min_cov_site", type=int, default=1, help="Min #walks per site (bin1); 1 disables filtering")
    ap.add_argument("--min_cov_1kb", type=int, default=1, help="Min #walks per 1 kb bin; 1 disables filtering")
    ap.add_argument("--exclude_random", action="store_true", help="Exclude *_random, chrUn, EBV, KI/GL scaffolds")
    ap.add_argument("--chrom", default=None, help="Process a single chromosome (e.g., chr1)")
    ap.add_argument("--region", default=None, help="Region filter chr:start-end (overrides --chrom)")
    # correlation controls
    ap.add_argument("--z_axis_cap", type=float, default=10.0, help="Hard cap for correlation axes half-range")
    ap.add_argument("--z_axis_q", type=float, default=0.995, help="Quantile for symmetric axes (0.5%–99.5%)")
    ap.add_argument("--hexbin_bins", type=int, default=200, help="Resolution of 2D histogram used for plotting/npz")
    ap.add_argument("--plot_bin1", action="store_true", help="Also generate a sampled bin1 correlation PDF/NPZ")
    ap.add_argument("--bin1_sample", type=int, default=2_000_000, help="Max points for bin1 correlation")
    # v4p1 robust-centering controls
    ap.add_argument("--winsorize-q", type=float, default=0.005, help="Per-chrom winsorization tail fraction; 0 disables")
    ap.add_argument("--z-mad-floor", type=float, default=1e-9, help="Floor for MAD/scale to avoid division blow-up")
    ap.add_argument("--z-fallback", choices=["iqr","std","none"], default="iqr",
                    help="Fallback scale if MAD<floor (iqr/1.349, std, or none=1.0)")

    args = ap.parse_args()

    ts = datetime.now().strftime("%Y%m%d_%H%M%S")
    outdir = Path(args.outdir).expanduser().resolve() / f"wpi_awb_{ts}"
    ensure_dir(outdir)
    logging.basicConfig(filename=outdir / "run.log",
                        level=logging.INFO, format="%(asctime)s | %(levelname)s | %(message)s")
    logging.info("Parameters: %s", vars(args))

    # region/chrom filters
    region_chr = None; region_start = region_end = None
    if args.region:
        region_chr, region_start, region_end = parse_region(args.region)
    elif args.chrom:
        region_chr = args.chrom

    # temp partials
    pdir = outdir / "partials"; ensure_dir(pdir)
    site_partials: List[Path] = []; bin_partials: List[Path] = []

    usecols = ["A_chrom","A_mid","B_mid","C_mid"]
    path = Path(args.walks).expanduser()
    open_fn = gzip.open if path.suffix.endswith(".gz") else open

    # ----- Stream & per-chunk reduction -----
    with open_fn(path, "rt") as fh:
        reader = pd.read_csv(fh, sep="\t", comment="#", header=0,
                             usecols=usecols, chunksize=args.chunk_rows, low_memory=False)
        for cno, chunk in enumerate(reader, 1):
            if region_chr:
                chunk = chunk[chunk["A_chrom"] == region_chr]
            if region_start is not None:
                m = ((chunk["A_mid"] >= region_start) & (chunk["A_mid"] < region_end)) \
                    | ((chunk["B_mid"] >= region_start) & (chunk["B_mid"] < region_end)) \
                    | ((chunk["C_mid"] >= region_start) & (chunk["C_mid"] < region_end))
                chunk = chunk[m]
            if chunk.empty:
                logging.info("chunk %d: empty after filters", cno); continue

            for col in ("A_mid","B_mid","C_mid"):
                chunk[col] = pd.to_numeric(chunk[col], errors="coerce")
            chunk.dropna(subset=["A_mid","B_mid","C_mid"], inplace=True)
            if chunk.empty:
                logging.info("chunk %d: empty after numeric dropna", cno); continue
            chunk[["A_mid","B_mid","C_mid"]] = chunk[["A_mid","B_mid","C_mid"]].astype(np.int64)

            A = chunk["A_mid"].to_numpy(np.int64)
            B = chunk["B_mid"].to_numpy(np.int64)
            C = chunk["C_mid"].to_numpy(np.int64)

            AB = np.abs(B - A).astype(np.float64)
            BC = np.abs(C - B).astype(np.float64)
            AC = np.abs(C - A).astype(np.float64)
            denom = AB + BC

            # Compute F only where denom>0 to avoid divide warnings
            F = np.full_like(denom, np.nan, dtype=np.float64)
            mask_div = denom > 0
            F[mask_div] = 1.0 - (AC[mask_div] / denom[mask_div])
            np.clip(F, 0.0, 1.0, out=F)

            s = np.sign(((A + C) / 2.0) - B)
            WPI = s * F
            AWB = F

            mask = np.isfinite(WPI) & np.isfinite(AWB)
            if not mask.any():
                logging.info("chunk %d: all values non-finite", cno); continue

            # Site-level partial (Chrom=A_chrom, Pos=B_mid)
            df_site = pd.DataFrame({
                "Chrom": chunk["A_chrom"].to_numpy()[mask],
                "Pos":   B[mask].astype(np.int64),
                "sum_WPI": WPI[mask].astype(np.float64),
                "sum_AWB": AWB[mask].astype(np.float64),
                "n": np.ones(mask.sum(), dtype=np.int32)
            })
            df_site = df_site.groupby(["Chrom","Pos"], as_index=False).agg(
                sum_WPI=("sum_WPI","sum"), sum_AWB=("sum_AWB","sum"),
                n=("n","sum")
            )
            p_site = pdir / f"site_{cno:06d}.tsv.gz"
            df_site.to_csv(p_site, sep="\t", index=False, compression="gzip")
            site_partials.append(p_site)

            # 1 kb bin partial (Start = floor(Pos/bin_size)*bin_size)
            Start = (df_site["Pos"] // args.bin_size) * args.bin_size
            df_bin = pd.DataFrame({
                "Chrom": df_site["Chrom"],
                "Start": Start.astype(np.int64),
                "sum_WPI": df_site["sum_WPI"],
                "sum_AWB": df_site["sum_AWB"],
                "n_walks": df_site["n"],
                "n_sites": np.ones(len(df_site), dtype=np.int32)
            })
            df_bin = df_bin.groupby(["Chrom","Start"], as_index=False).agg(
                sum_WPI=("sum_WPI","sum"), sum_AWB=("sum_AWB","sum"),
                n_walks=("n_walks","sum"), n_sites=("n_sites","sum")
            )
            p_bin = pdir / f"bin_{cno:06d}.tsv.gz"
            df_bin.to_csv(p_bin, sep="\t", index=False, compression="gzip")
            bin_partials.append(p_bin)

            logging.info("chunk %d: wrote partials (sites=%d, bins=%d)", cno, len(df_site), len(df_bin))

    if not site_partials:
        print("[FATAL] No data after filtering; abort.", file=sys.stderr); sys.exit(2)

    # ----- Merge site-level partials -----
    logging.info("Merging %d site partials", len(site_partials))
    site_frames = [pd.read_csv(p, sep="\t") for p in site_partials]
    sites = (pd.concat(site_frames, ignore_index=True)
               .groupby(["Chrom","Pos"], as_index=False)
               .agg(sum_WPI=("sum_WPI","sum"), sum_AWB=("sum_AWB","sum"),
                    n=("n","sum")))
    sites["WPI_mean_site"] = sites["sum_WPI"] / sites["n"].replace(0, np.nan)
    sites["AWB_mean_site"] = sites["sum_AWB"] / sites["n"].replace(0, np.nan)

    # ----- Merge 1 kb partials -----
    logging.info("Merging %d bin partials", len(bin_partials))
    bin_frames = [pd.read_csv(p, sep="\t") for p in bin_partials]
    bins = (pd.concat(bin_frames, ignore_index=True)
              .groupby(["Chrom","Start"], as_index=False)
              .agg(sum_WPI=("sum_WPI","sum"), sum_AWB=("sum_AWB","sum"),
                   n_walks=("n_walks","sum"), n_sites=("n_sites","sum")))
    bins["End"] = bins["Start"] + args.bin_size
    bins["WPI_mean_1kb"] = bins["sum_WPI"] / bins["n_walks"].replace(0, np.nan)
    bins["AWB_mean_1kb"] = bins["sum_AWB"] / bins["n_walks"].replace(0, np.nan)

    # ----- Chromosome filters -----
    def apply_chr_filters(df, chr_col="Chrom"):
        out = df[df[chr_col].astype(str).str.startswith("chr")]
        if args.exclude_random:
            out = out[~out[chr_col].str.contains("_random|_KI|_GL|chrUn|EBV")]
        return out

    sites = apply_chr_filters(sites, "Chrom")
    bins  = apply_chr_filters(bins, "Chrom")

    # ----- Coverage filters (defaults: disabled; 1 == keep-all) ----------------
    before_sites = len(sites)
    if args.min_cov_site > 1:
        sites = sites[sites["n"] >= args.min_cov_site]
    logging.info("Site coverage filter (min_cov_site=%d) kept %d/%d", args.min_cov_site, len(sites), before_sites)

    before_bins = len(bins)
    if args.min_cov_1kb > 1:
        bins  = bins[bins["n_walks"] >= args.min_cov_1kb]
    logging.info("1 kb coverage filter (min_cov_1kb=%d) kept %d/%d", args.min_cov_1kb, len(bins), before_bins)

    # ----- Robust Z per chromosome (v4p1: winsorize + fallback) ----------------
    # Sites:
    z_wpi_site, stats_wpi_site = robust_center_scale_per_chrom(
        sites, "WPI_mean_site", "Chrom", args.winsorize_q, args.z_mad_floor, args.z_fallback
    )
    z_awb_site, stats_awb_site = robust_center_scale_per_chrom(
        sites, "AWB_mean_site", "Chrom", args.winsorize_q, args.z_mad_floor, args.z_fallback
    )
    sites["Z_WPI_site"] = z_wpi_site
    sites["Z_AWB_site"] = z_awb_site

    # 1 kb:
    z_wpi_1kb, stats_wpi_1kb = robust_center_scale_per_chrom(
        bins, "WPI_mean_1kb", "Chrom", args.winsorize_q, args.z_mad_floor, args.z_fallback
    )
    z_awb_1kb, stats_awb_1kb = robust_center_scale_per_chrom(
        bins, "AWB_mean_1kb", "Chrom", args.winsorize_q, args.z_mad_floor, args.z_fallback
    )
    bins["Z_WPI_1kb"] = z_wpi_1kb
    bins["Z_AWB_1kb"] = z_awb_1kb

    # ----- Write TSVs -----
    sites_tsv = outdir / f"wpi_awb_sites_{ts}.tsv.gz"
    sites[["Chrom","Pos","WPI_mean_site","AWB_mean_site","n","Z_WPI_site","Z_AWB_site"]].to_csv(
        sites_tsv, sep="\t", index=False, compression="gzip")

    bins_tsv  = outdir / f"wpi_awb_1kb_{ts}.tsv.gz"
    bins[["Chrom","Start","End","WPI_mean_1kb","AWB_mean_1kb","n_walks","n_sites","Z_WPI_1kb","Z_AWB_1kb"]].to_csv(
        bins_tsv, sep="\t", index=False, compression="gzip")

    # ----- BigWigs -----
    chrom_sizes = read_fai(args.fai)
    # bin1 (1-bp sparse)
    z_wpi_bin1_bw = outdir / "Z_WPI_bin1.bw"
    z_awb_bin1_bw = outdir / "Z_AWB_bin1.bw"
    write_bw_sparse_sites(z_wpi_bin1_bw, chrom_sizes, sites[["Chrom","Pos","Z_WPI_site"]], "Z_WPI_site")
    write_bw_sparse_sites(z_awb_bin1_bw, chrom_sizes, sites[["Chrom","Pos","Z_AWB_site"]], "Z_AWB_site")
    # aliases (copies)
    (outdir / "zWPI_bin1.bw").write_bytes(z_wpi_bin1_bw.read_bytes())

    # 1 kb fixed bins
    z_wpi_1kb_bw = outdir / "Z_WPI_1kb.bw"
    z_awb_1kb_bw = outdir / "Z_AWB_1kb.bw"
    write_bw_fixed(z_wpi_1kb_bw, chrom_sizes, bins[["Chrom","Start","Z_WPI_1kb"]], "Z_WPI_1kb", args.bin_size)
    write_bw_fixed(z_awb_1kb_bw, chrom_sizes, bins[["Chrom","Start","Z_AWB_1kb"]], "Z_AWB_1kb", args.bin_size)
    # aliases (copies)
    (outdir / "1kb_bin_zWPI.bw").write_bytes(z_wpi_1kb_bw.read_bytes())
    (outdir / "1kb_bin_AWB.bw").write_bytes(z_awb_1kb_bw.read_bytes())

    # ----- Correlation (1 kb bins; Z_WPI vs Z_AWB) -----
    def finite_pair(x: np.ndarray, y: np.ndarray) -> Tuple[np.ndarray, np.ndarray, np.ndarray]:
        m = np.isfinite(x) & np.isfinite(y)
        return x[m], y[m], m

    x_all, y_all, _ = finite_pair(bins["Z_AWB_1kb"].to_numpy(), bins["Z_WPI_1kb"].to_numpy())
    def symmetric_axis_from_quantiles(x: np.ndarray, q: float, cap: float, min_half: float = 3.0) -> float:
        if x.size == 0: return min_half
        x = x[np.isfinite(x)]
        if x.size == 0: return min_half
        lo, hi = np.quantile(x, 1.0-q), np.quantile(x, q)
        half = max(abs(lo), abs(hi))
        half = min(half, cap)
        return max(half, min_half)
    def corr_stats(x: np.ndarray, y: np.ndarray) -> Dict[str, float]:
        stats = {}
        if x.size == 0:
            return dict(n=0, pearson_r=np.nan, pearson_p=np.nan, spearman_rho=np.nan, spearman_p=np.nan)
        try:
            import scipy.stats as ss
            r, p = ss.pearsonr(x, y); stats["pearson_r"] = float(r); stats["pearson_p"] = float(p)
            rho, pp = ss.spearmanr(x, y, nan_policy="omit"); stats["spearman_rho"] = float(rho); stats["spearman_p"] = float(pp)
        except Exception:
            r = np.corrcoef(x, y)[0,1]; stats["pearson_r"] = float(r); stats["pearson_p"] = np.nan
            xr = pd.Series(x).rank().values; yr = pd.Series(y).rank().values
            rho = np.corrcoef(xr, yr)[0,1]; stats["spearman_rho"] = float(rho); stats["spearman_p"] = np.nan
        stats["n"] = int(x.size)
        return stats

    stats_full = corr_stats(x_all, y_all)
    halfx = symmetric_axis_from_quantiles(x_all, args.z_axis_q, args.z_axis_cap, min_half=3.0)
    halfy = symmetric_axis_from_quantiles(y_all, args.z_axis_q, args.z_axis_cap, min_half=3.0)
    x_clip = np.clip(x_all, -halfx, halfx); y_clip = np.clip(y_all, -halfy, halfy)
    stats_clip = corr_stats(x_clip, y_clip)

    def plot_corr_pdf(out_pdf: Path, x: np.ndarray, y: np.ndarray,
                      halfx: float, halfy: float, bins: int,
                      title: str, xlabel: str, ylabel: str,
                      stats: Dict[str, float]):
        if plt is None:
            return None, None, None
        H, xedges, yedges = np.histogram2d(
            np.clip(x, -halfx, halfx), np.clip(y, -halfy, halfy), bins=bins,
            range=[[-halfx, halfx], [-halfy, halfy]]
        )
        H = H.T
        fig = plt.figure(figsize=(6.5,5.5), dpi=120)
        ax = fig.add_subplot(111)
        im = ax.imshow(H, origin="lower",
                       extent=[xedges[0], xedges[-1], yedges[0], yedges[-1]],
                       aspect="auto", cmap="viridis")
        cbar = fig.colorbar(im, ax=ax, fraction=0.046, pad=0.04)
        cbar.set_label("count")
        ax.set_xlim(-halfx, halfx); ax.set_ylim(-halfy, halfy)
        ax.set_xlabel(xlabel); ax.set_ylabel(ylabel); ax.set_title(title)
        txt = f"N={stats.get('n',0):,}\nPearson r={stats.get('pearson_r',np.nan):.3f}"
        if not np.isnan(stats.get('pearson_p', np.nan)): txt += f" (p={stats.get('pearson_p',np.nan):.2e})"
        txt += f"\nSpearman ρ={stats.get('spearman_rho',np.nan):.3f}"
        if not np.isnan(stats.get('spearman_p',np.nan)): txt += f" (p={stats.get('spearman_p',np.nan):.2e})"
        ax.text(0.02, 0.98, txt, transform=ax.transAxes, va="top", ha="left",
                bbox=dict(facecolor="white", alpha=0.8, edgecolor="none"), fontsize=9)
        fig.tight_layout()
        fig.savefig(out_pdf, format="pdf")
        plt.close(fig)
        return H, xedges, yedges

    corr_pdf_1kb = outdir / f"corr_Z_WPI_vs_Z_AWB_1kb_{ts}.pdf"
    H1, xedges1, yedges1 = plot_corr_pdf(
        corr_pdf_1kb, x_all, y_all, halfx, halfy, args.hexbin_bins,
        title="Z_WPI vs Z_AWB (1 kb bins)",
        xlabel="Z_AWB (bend magnitude; robust-Z)",
        ylabel="Z_WPI (signed bend; robust-Z)",
        stats=stats_clip
    )
    corr_npz_1kb = outdir / f"corr_Z_WPI_vs_Z_AWB_1kb_{ts}.npz"
    if H1 is not None:
        np.savez(corr_npz_1kb, H=H1, xedges=xedges1, yedges=yedges1,
                 axes=np.array([halfx, halfy]),
                 stats_full=np.array(list(stats_full.values()), dtype=float),
                 stats_clip=np.array(list(stats_clip.values()), dtype=float),
                 stats_labels=np.array(list(stats_full.keys()), dtype=object))

    # Optional: sampled bin1 correlation
    if args.plot_bin1 and len(sites):
        x_b = sites["Z_AWB_site"].to_numpy(); y_b = sites["Z_WPI_site"].to_numpy()
        m = np.isfinite(x_b) & np.isfinite(y_b); x_b = x_b[m]; y_b = y_b[m]
        N = x_b.size
        if N > args.bin1_sample:
            rng = np.random.default_rng(42)
            idx = rng.choice(N, size=args.bin1_sample, replace=False)
            x_b = x_b[idx]; y_b = y_b[idx]
        stats_full_b = {"n": int(x_b.size)}
        halfx_b = halfy_b = max(3.0, min(args.z_axis_cap, np.nanpercentile(np.abs(np.concatenate([x_b, y_b])), 99.5)))
        corr_pdf_bin1 = outdir / f"corr_Z_WPI_vs_Z_AWB_bin1_{ts}.pdf"
        if plt is not None:
            H2, xedges2, yedges2 = np.histogram2d(
                np.clip(x_b, -halfx_b, halfx_b), np.clip(y_b, -halfy_b, halfy_b),
                bins=args.hexbin_bins,
                range=[[-halfx_b, halfx_b], [-halfy_b, halfy_b]]
            )
            H2 = H2.T
            fig = plt.figure(figsize=(6.5,5.5), dpi=120)
            ax = fig.add_subplot(111)
            im = ax.imshow(H2, origin="lower",
                           extent=[xedges2[0], xedges2[-1], yedges2[0], yedges2[-1]],
                           aspect="auto", cmap="viridis")
            fig.colorbar(im, ax=ax, fraction=0.046, pad=0.04).set_label("count")
            ax.set_xlim(-halfx_b, halfx_b); ax.set_ylim(-halfy_b, halfy_b)
            ax.set_xlabel("Z_AWB_site"); ax.set_ylabel("Z_WPI_site")
            ax.set_title(f"Z_WPI vs Z_AWB (bin1 sampled; N≤{args.bin1_sample:,})")
            fig.tight_layout(); fig.savefig(corr_pdf_bin1, format="pdf"); plt.close(fig)

    # ----- Metadata ------------------------------------------------------------
    meta = {
        "timestamp": ts,
        "walks_file": str(path),
        "outdir": str(outdir),
        "fai": str(Path(args.fai).resolve()),
        "bin_size": args.bin_size,
        "chunk_rows": args.chunk_rows,
        "min_cov_site": args.min_cov_site,
        "min_cov_1kb": args.min_cov_1kb,
        "exclude_random": bool(args.exclude_random),
        "filters": args.region or args.chrom or "genome",
        "metrics": {
            "WPI": "sign((A+C)/2 - B) * (1 - AC/(AB+BC)) ∈ [-1,1]",
            "AWB": "1 - AC/(AB+BC) ∈ [0,1]"
        },
        "zscore": {
            "type": "per-chrom robust Z with winsorization",
            "winsorize_q": args.winsorize_q,
            "mad_floor": args.z_mad_floor,
            "fallback": args.z_fallback,
            "per_chrom_stats": {
                **merge_stats("sites/WPI", stats_wpi_site),
                **merge_stats("sites/AWB", stats_awb_site),
                **merge_stats("1kb/WPI",  stats_wpi_1kb),
                **merge_stats("1kb/AWB",  stats_awb_1kb),
            }
        },
        "coverage_filter_stage": "AFTER per-site/per-bin mean aggregation, BEFORE robust-Z",
        "outputs": {
            "sites_tsv": str(sites_tsv), "bins_tsv": str(bins_tsv),
            "Z_WPI_bin1.bw": str(z_wpi_bin1_bw), "Z_AWB_bin1.bw": str(z_awb_bin1_bw),
            "Z_WPI_1kb.bw": str(z_wpi_1kb_bw), "Z_AWB_1kb.bw": str(z_awb_1kb_bw),
            "aliases": ["zWPI_bin1.bw", "1kb_bin_zWPI.bw", "1kb_bin_AWB.bw"]
        },
        "script_sha256": sha256_of_file(Path(__file__))
    }
    with open(outdir / f"metadata_{ts}.json", "w") as fh:
        json.dump(meta, fh, indent=2)

    # ----- Final prints -----
    print(f"[DONE] Sites TSV  → {sites_tsv}")
    print(f"[DONE] 1kb TSV    → {bins_tsv}")
    print(f"[DONE] BigWigs    → {z_wpi_bin1_bw}, {z_awb_bin1_bw}, {z_wpi_1kb_bw}, {z_awb_1kb_bw}")
    print(f"[DONE] Correlation→ {corr_pdf_1kb}, {corr_npz_1kb}")
    print(f"[DONE] Stage note → Coverage filters applied AFTER aggregation, BEFORE robust-Z.")
    print(f"[DONE] Z-centering→ winsorize_q={args.winsorize_q}, mad_floor={args.z_mad_floor}, fallback={args.z_fallback}")

if __name__ == "__main__":
    main()
```
