## Supplementary material for "CAD-C: An engineered nuclease enables repair-free *in situ* proximity ligation and nucleosome-resolution chromosome walks in human cells": Custom code: WPI_compute_and_bigwig_v1_20250926.py

```
#!/usr/bin/env python3
# =========================================================================================
#  WPI_compute_and_bigwig_v1_20250926.py
#  -----------------------------------------------------------------------------------------
#  Walk Polarity Index (WPI): per-walk, signed dog-leg amplitude with robust-Z BigWig export
#
#  Version : 1.0   •   Date : 2025-09-26
#  Author  : Jan Soroczynski (Risca Lab, The Rockefeller University)
#
#  README
#  ------
#  PURPOSE
#    Compute a signed "dog-leg" index for three-fragment CAD walks and export a genome-wide
#    BigWig without binning. For each walk with fragment midpoints A, B, C (bp; A upstream of C):
#
#      Fold fraction (magnitude):  F = 1 - AC / (AB + BC)       ∈ [0,1]
#      Polarity (direction)     :  s = sign( (A + C)/2 - B )    (+ upstream, − downstream)
#      Walk Polarity Index      :  WPI = s * F                  ∈ [-1,1]
#
#    WPI=0 for linear walks (A<B<C). |WPI|→1 for near-symmetric dog-legs; sign encodes whether
#    B lies upstream (+) or downstream (−) of the A–C midpoint.
#
#  INPUT
#    • A tab-delimited walks table (.tsv or .tsv.gz) with header (leading "#" comments allowed)
#      containing *named* columns:  A_chrom, A_mid, B_mid, C_mid
#    • Walks must be pre-oriented so that A is upstream of C (A_mid < C_mid).
#
#  OUTPUTS (written under: --outdir / wpi_<timestamp>/)
#    • wpi_sites_<ts>.tsv.gz
#        Chrom, Pos, WPI_mean, n, Z_WPI    # per-site (B_mid) mean WPI, coverage, robust-Z
#    • Z_WPI.bw
#        1-bp intervals [Pos, Pos+1) with per-chromosome robust Z of WPI_mean (no detrend)
#    • (QC) wpi_hist_<ts>.png, z_wpi_hist_<ts>.png, wpi_summary_<ts>.txt
#    • metadata_<ts>.json  (parameters, formulae, script checksum)
#    • run.log
#
#  CLI
#    --walks        path/to/walks.tsv(.gz)                 [required]
#    --outdir       output root directory                  [required]
#    --fai          genome FASTA index (.fai)              [required]
#    --chunk_rows   8000000                                [default]
#    --min_cov      5   # per-site minimum n before Z/bw   [default 5]
#    --exclude_random         # drop *_random, chrUn, EBV, KI/GL   [flag]
#    --chrom        chr1       # optional chromosome filter
#    --region       chr3:5e6-8e6  # optional region (overrides --chrom)
#
#  IMPLEMENTATION NOTES
#    • Streaming: reads the walks file in chunks, computes per-walk WPI, reduces each chunk to
#      per-site partial sums (sum_WPI, n), writes gzipped partials, then merges all partials.
#    • Aggregation: per-site mean WPI (Chrom=A_chrom, Pos=B_mid). Per-chrom robust Z is applied
#      to WPI_mean after optional coverage filtering (--min_cov).
#    • Export: BigWig has sparse 1-bp entries at B_mid coordinates with value Z_WPI.
#    • No detrending: WPI encodes a local, bounded effect; robust Z standardizes distributions.
#
#  DEPENDENCIES
#    numpy, pandas, pyBigWig, (optional) statsmodels for robust MAD; matplotlib for QC histograms
#
#  FORMULAE
#    AB = |B-A|, BC = |C-B|, AC = |C-A|, denom = AB+BC
#    F  = max(0, min(1, 1 - AC/denom)) if denom>0 else NaN
#    s  = sign( (A+C)/2 - B )  # + upstream, 0 exactly at midpoint, − downstream
#    WPI = s * F
# =========================================================================================
from __future__ import annotations
import argparse, gzip, json, logging, hashlib, sys, math
from datetime import datetime
from pathlib import Path
from typing import Dict, List, Tuple

import numpy as np
import pandas as pd

# Optional deps
try:
    import pyBigWig
except Exception as e:
    pyBigWig = None

try:
    import importlib.util
    _sm = importlib.util.find_spec("statsmodels")
    if _sm is not None:
        from statsmodels import robust
        def mad(a): return robust.mad(a, c=1.4826)
    else:
        raise ImportError
except Exception:
    def mad(a, c: float = 1.4826) -> float:
        a = np.asarray(a, dtype=float)
        med = np.nanmedian(a)
        return c * np.nanmedian(np.abs(a - med))

try:
    import matplotlib
    matplotlib.use("Agg")
    import matplotlib.pyplot as plt
except Exception:
    plt = None

pd.options.mode.copy_on_write = True

# ---------------- helpers ------------------------------------------------------
def sha256_of_file(p: Path) -> str:
    h = hashlib.sha256(); h.update(p.read_bytes()); return h.hexdigest()

def parse_region(arg: str) -> Tuple[str, int, int]:
    chrom, rest = arg.split(":")
    start, end  = rest.replace(",", "").split("-")
    return chrom, int(float(start)), int(float(end))

def ensure_dir(p: Path): p.mkdir(parents=True, exist_ok=True)

def read_fai(path: str) -> Dict[str, int]:
    d: Dict[str, int] = {}
    with open(path) as fh:
        for line in fh:
            chrom, length, *_ = line.split()
            d[chrom] = int(length)
    return d

def robust_z_by_chrom(df: pd.DataFrame, col: str, chrom_col: str = "Chrom") -> pd.Series:
    def _z(s: pd.Series) -> pd.Series:
        m = s.median()
        mdev = mad(s.values)
        return (s - m) / mdev if mdev else np.nan
    return df.groupby(chrom_col)[col].transform(_z)

def write_bw(out_path: Path, chrom_sizes: Dict[str, int], df: pd.DataFrame, value_col: str):
    if pyBigWig is None:
        raise RuntimeError("pyBigWig is not available; cannot write BigWig.")
    bw = pyBigWig.open(str(out_path), "w")
    header = list(chrom_sizes.items())
    bw.addHeader(header)
    # Write sparse 1-bp entries per chromosome in header order
    for chrom, chrom_len in header:
        sub = df.loc[df["Chrom"] == chrom, ["Pos", value_col]].drop_duplicates("Pos")
        if sub.empty:
            bw.addEntries([chrom], [0], [1], [0.0])
            continue
        s = sub["Pos"].astype(np.int64).values
        e = s + 1
        v = sub[value_col].astype(float).values
        mask = (s >= 0) & (e <= chrom_len) & np.isfinite(v)
        s, e, v = s[mask], e[mask], v[mask]
        if len(v) == 0:
            bw.addEntries([chrom], [0], [1], [0.0])
            continue
        try:
            bw.addEntries([chrom] * len(v), s.tolist(), e.tolist(), v.tolist())
        except RuntimeError as err:
            bw.close()
            raise
    bw.close()

# ---------------- main ---------------------------------------------------------
def main():
    ap = argparse.ArgumentParser("Compute WPI and export Z_WPI BigWig (no binning).",
                                 formatter_class=argparse.ArgumentDefaultsHelpFormatter)
    ap.add_argument("--walks", required=True, help="Input walks .tsv or .tsv.gz with A_chrom,A_mid,B_mid,C_mid")
    ap.add_argument("--outdir", required=True, help="Output root directory (timestamped subdir will be created)")
    ap.add_argument("--fai", required=True, help="Genome FASTA index (.fai)")
    ap.add_argument("--chunk_rows", type=int, default=8_000_000, help="Rows per stream chunk")
    ap.add_argument("--min_cov", type=int, default=5, help="Minimum per-site coverage (n) before Z/bw")
    ap.add_argument("--exclude_random", action="store_true", help="Exclude *_random, chrUn, EBV, KI/GL scaffolds")
    ap.add_argument("--chrom", default=None, help="Process a single chromosome (e.g., chr1)")
    ap.add_argument("--region", default=None, help="Region filter chr:start-end (overrides --chrom)")
    args = ap.parse_args()

    ts = datetime.now().strftime("%Y%m%d_%H%M%S")
    outdir = Path(args.outdir).expanduser().resolve() / f"wpi_{ts}"
    ensure_dir(outdir)
    logging.basicConfig(filename=outdir / "run.log",
                        level=logging.INFO,
                        format="%(asctime)s | %(levelname)s | %(message)s")
    logging.info("Parameters: %s", vars(args))

    # region/chrom filters
    region_chr = None
    region_start = region_end = None
    if args.region:
        region_chr, region_start, region_end = parse_region(args.region)
    elif args.chrom:
        region_chr = args.chrom

    # temp partials dir
    partial_dir = outdir / "partials"
    ensure_dir(partial_dir)
    partial_paths: List[Path] = []

    usecols = ["A_chrom", "A_mid", "B_mid", "C_mid"]
    path = Path(args.walks).expanduser()
    open_fn = gzip.open if path.suffix.endswith(".gz") else open

    # stream + per-chunk reduction
    with open_fn(path, "rt") as fh:
        reader = pd.read_csv(fh, sep="\t", comment="#", header=0,
                             usecols=usecols, chunksize=args.chunk_rows, low_memory=False)
        for cno, chunk in enumerate(reader, 1):
            if region_chr:
                chunk = chunk[chunk["A_chrom"] == region_chr]
            if region_start is not None:
                m = ((chunk["A_mid"] >= region_start) & (chunk["A_mid"] < region_end)) \
                    | ((chunk["B_mid"] >= region_start) & (chunk["B_mid"] < region_end)) \
                    | ((chunk["C_mid"] >= region_start) & (chunk["C_mid"] < region_end))
                chunk = chunk[m]
            if chunk.empty:
                logging.info("chunk %d: empty after filters", cno)
                continue

            # numeric coercion
            for col in ("A_mid", "B_mid", "C_mid"):
                chunk[col] = pd.to_numeric(chunk[col], errors="coerce")
            chunk.dropna(subset=["A_mid", "B_mid", "C_mid"], inplace=True)
            if chunk.empty:
                logging.info("chunk %d: empty after numeric dropna", cno)
                continue
            chunk[["A_mid", "B_mid", "C_mid"]] = chunk[["A_mid", "B_mid", "C_mid"]].astype(np.int64)

            A = chunk["A_mid"].to_numpy(np.int64)
            B = chunk["B_mid"].to_numpy(np.int64)
            C = chunk["C_mid"].to_numpy(np.int64)

            AB = np.abs(B - A).astype(np.float64)
            BC = np.abs(C - B).astype(np.float64)
            AC = np.abs(C - A).astype(np.float64)
            denom = AB + BC

            # F = 1 - AC/denom  (guard denom>0); clamp [0,1]
            F = np.where(denom > 0, 1.0 - (AC / denom), np.nan)
            F = np.clip(F, 0.0, 1.0, out=F)
            s = np.sign(((A + C) / 2.0) - B)
            WPI = s * F
            # drop NaNs if any
            mask = np.isfinite(WPI)
            if not mask.any():
                logging.info("chunk %d: all WPI non-finite", cno)
                continue

            dfp = pd.DataFrame({
                "Chrom": chunk["A_chrom"].to_numpy()[mask],
                "Pos":   B[mask].astype(np.int64),
                "sum_WPI": WPI[mask].astype(np.float64),
                "n": np.ones(mask.sum(), dtype=np.int32)
            })
            # per-chunk aggregation to limit temp file size
            dfp = dfp.groupby(["Chrom", "Pos"], as_index=False).agg(sum_WPI=("sum_WPI", "sum"),
                                                                    n=("n", "sum"))
            ppath = partial_dir / f"partial_{cno:06d}.tsv.gz"
            dfp.to_csv(ppath, sep="\t", index=False, compression="gzip")
            partial_paths.append(ppath)
            logging.info("chunk %d: wrote partial %s (rows=%d)", cno, ppath.name, len(dfp))

    if not partial_paths:
        print("[FATAL] No data after filtering; aborting.", file=sys.stderr)
        sys.exit(2)

    # merge partials
    logging.info("Merging %d partials", len(partial_paths))
    agg = []
    for p in partial_paths:
        agg.append(pd.read_csv(p, sep="\t"))
    merged = (pd.concat(agg, ignore_index=True)
                .groupby(["Chrom", "Pos"], as_index=False)
                .agg(sum_WPI=("sum_WPI", "sum"), n=("n", "sum")))
    # compute mean
    merged["WPI_mean"] = merged["sum_WPI"] / merged["n"].replace(0, np.nan)

    # filters: chromosome naming + random scaffolds
    merged = merged[merged["Chrom"].astype(str).str.startswith("chr")]
    if args.exclude_random:
        merged = merged[~merged["Chrom"].str.contains("_random|_KI|_GL|chrUn|EBV")]

    # coverage filter
    before_cov = len(merged)
    if args.min_cov > 0:
        merged = merged[merged["n"] >= args.min_cov]
    logging.info("Coverage filter min_cov=%d kept %d/%d rows", args.min_cov, len(merged), before_cov)

    if merged.empty:
        print("[FATAL] No rows after coverage filter; aborting.", file=sys.stderr)
        sys.exit(3)

    # robust Z per chromosome
    merged["Z_WPI"] = robust_z_by_chrom(merged, "WPI_mean", "Chrom")

    # write site table
    sites_path = outdir / f"wpi_sites_{ts}.tsv.gz"
    merged[["Chrom", "Pos", "WPI_mean", "n", "Z_WPI"]].to_csv(sites_path, sep="\t",
                                                               index=False, compression="gzip")

    # BigWig export
    chrom_sizes = read_fai(args.fai)
    bw_path = outdir / "Z_WPI.bw"
    try:
        write_bw(bw_path, chrom_sizes, merged[["Chrom", "Pos", "Z_WPI"]], "Z_WPI")
    except Exception as e:
        print(f"[FATAL] BigWig export failed: {e}", file=sys.stderr)
        sys.exit(4)

    # QC histograms and summary
    try:
        summary_path = outdir / f"wpi_summary_{ts}.txt"
        with open(summary_path, "w") as fh:
            fh.write(f"rows_after_merge_and_filters: {len(merged)}\n")
            fh.write(f"min_cov: {args.min_cov}\n")
            fh.write("WPI_mean: min={:.3f} q25={:.3f} med={:.3f} q75={:.3f} max={:.3f}\n".format(
                np.nanmin(merged["WPI_mean"]), np.nanpercentile(merged["WPI_mean"], 25),
                np.nanmedian(merged["WPI_mean"]), np.nanpercentile(merged["WPI_mean"], 75),
                np.nanmax(merged["WPI_mean"])))
            fh.write("Z_WPI:    min={:.3f} q25={:.3f} med={:.3f} q75={:.3f} max={:.3f}\n".format(
                np.nanmin(merged["Z_WPI"]), np.nanpercentile(merged["Z_WPI"], 25),
                np.nanmedian(merged["Z_WPI"]), np.nanpercentile(merged["Z_WPI"], 75),
                np.nanmax(merged["Z_WPI"])))
        if plt is not None:
            # WPI_mean histogram [-1,1]
            fig1 = plt.figure(figsize=(6,4), dpi=120)
            plt.hist(merged["WPI_mean"].values[np.isfinite(merged["WPI_mean"].values)],
                     bins=101, range=(-1,1))
            plt.xlabel("WPI_mean"); plt.ylabel("count"); plt.title("WPI_mean distribution")
            fig1.tight_layout(); fig1.savefig(outdir / f"wpi_hist_{ts}.png"); plt.close(fig1)
            # Z_WPI histogram
            fig2 = plt.figure(figsize=(6,4), dpi=120)
            zvals = merged["Z_WPI"].values[np.isfinite(merged["Z_WPI"].values)]
            plt.hist(zvals, bins=101)
            plt.xlabel("Z_WPI"); plt.ylabel("count"); plt.title("Z_WPI distribution")
            fig2.tight_layout(); fig2.savefig(outdir / f"z_wpi_hist_{ts}.png"); plt.close(fig2)
    except Exception as e:
        logging.warning("QC summary/plots failed: %s", e)

    # metadata
    meta = {
        "timestamp": ts,
        "walks_file": str(path),
        "outdir": str(outdir),
        "fai": str(Path(args.fai).resolve()),
        "chunk_rows": args.chunk_rows,
        "min_cov": args.min_cov,
        "exclude_random": bool(args.exclude_random),
        "filters": args.region or args.chrom or "genome",
        "index_name": "WPI (Walk Polarity Index)",
        "formulas": {
            "AB": "|B-A|", "BC": "|C-B|", "AC": "|C-A|",
            "F": "1 - AC/(AB+BC)  (clamped to [0,1]; denom>0)",
            "sign": "sign((A+C)/2 - B)  (+ upstream, 0 midpoint, − downstream)",
            "WPI": "sign * F  ∈ [-1,1]"
        },
        "zscore": "per-chrom robust Z: (x - median) / MAD  (MAD scaled by 1.4826; NaN if MAD=0)",
        "script_sha256": sha256_of_file(Path(__file__))
    }
    with open(outdir / f"metadata_{ts}.json", "w") as fh:
        json.dump(meta, fh, indent=2)

    print(f"[DONE] Sites table   → {sites_path}")
    print(f"[DONE] BigWig        → {bw_path}")
    print(f"[DONE] QC summary    → {outdir / f'wpi_summary_{ts}.txt'}")
    if plt is not None:
        print(f"[DONE] QC plots     → {outdir / f'wpi_hist_{ts}.png'}, {outdir / f'z_wpi_hist_{ts}.png'}")

if __name__ == "__main__":
    main()
```
