## Supplementary material for "CAD-C: An engineered nuclease enables repair-free *in situ* proximity ligation and nucleosome-resolution chromosome walks in human cells": Custom code: BEI_flowchart_guide.txt

BEI WORKFLOW — Step‑by‑step runbook (2025‑09‑25)

0) PREREQUISITES (once per environment)
    • Python 3.8+ with: numpy, pandas, pyBigWig  (statsmodels optional)
        pip install --user numpy pandas pyBigWig statsmodels
    • Confirm you have a “walks” table with these columns:
        A_chrom, A_mid, B_mid, C_mid
      (pre‑oriented so A lies upstream of C; B may be anywhere).
    • FASTA index (.fai) for your genome (you already use hg38):
        FAI=/rugpfs/fs0/risc_lab/store/risc_data/downloaded/hg38/genome/Sequence/WholeGenomeFasta/genome.fa.fai

    • Make sure the three updated scripts are saved and executable (chmod +x):
        CADwalk_calculate_BEI_v2_20250925.py
        calculate_detrended_cadwalk_ratio_metrics_v4_20250925.py
        BEI_to_bigwig_v5_20250925.py

    • Decide where to store results. I recommend mirroring your ABI layout:
        ANALYSIS_ROOT=</absolute/path/to>/.../merged_all/BEI_analyses_2025
      (Create this if it doesn’t exist. The generator will make a timestamped subdir inside.)

1) GENERATE BEI BINS (walks → 1 kb bins with r_A, r_B, r_C, BEI, n_tot)
   You can run either A) direct Python or B) via SLURM (if you wrap it). The generator will print the exact “BEI table → …/bei_<ts>/bei_bins_<ts>.tsv.gz” path you’ll use downstream.

   REQUIRED inputs to decide:
     WALKS=</path/to/your/pre‑oriented_walks.tsv.gz>   # has A_chrom, A_mid, B_mid, C_mid
     OUTDIR=${ANALYSIS_ROOT}                            # parent folder; script adds bei_<ts>/

   A) Direct Python:
        SCRIPTS=/rugpfs/fs0/risc_lab/scratch/jsoroczyns/CAD/CAD-C/scripts/2025_09_25
        WALKS=__SET_THIS__: e.g.
          /rugpfs/fs0/risc_lab/scratch/jsoroczyns/CAD/CADwalk/.../oriented_walks_mids.tsv.gz
        OUTDIR=__SET_THIS__: e.g.
          /rugpfs/fs0/risc_lab/scratch/jsoroczyns/CAD/CADwalk/.../merged_all/BEI_analyses_2025

        python ${SCRIPTS}/CADwalk_calculate_BEI_v2_20250925.py \
            --walks  "${WALKS}" \
            --outdir "${OUTDIR}" \
            --bin_size 1000

        NOTE: Copy the “BEI table → …/bei_<ts>/bei_bins_<ts>.tsv.gz” path the script prints.

   B) (Optional) If you prefer, you can make an SBATCH wrapper for this step; but your existing workflow ran this as a plain Python step, so A) is simplest.

2) DETREND/Z APPEND (add Z_ACd, Z_ABd, Z_BEI to a TSV for QC)
   This step produces an augmented TSV with your two detrended ratio Z‑scores and the **robust Z of BEI (no detrend)**. You can run it directly or with the provided SBATCH wrapper.

   REQUIRED inputs to decide:
     BEI_BINS=</the/path/printed/in/step1/bei_<ts>/bei_bins_<ts>.tsv.gz>
     MIN_COV=20 (what you used before)
     OUTDIR_PARENT can be the same parent as in step 1 (a new detrended_<ts>/ will be created)

   A) Direct Python:
        SCRIPTS=/rugpfs/fs0/risc_lab/scratch/jsoroczyns/CAD/CAD-C/scripts/2025_09_25
        BEI_BINS=__SET_THIS__  # from step 1 output
        TS=$(date +%Y%m%d_%H%M%S)
        OUTDIR="$(dirname "${BEI_BINS}")/detrended_${TS}"
        mkdir -p "${OUTDIR}"
        OUT_PREFIX="${OUTDIR}/bei_bins_detrended_${TS}"

        python ${SCRIPTS}/calculate_detrended_cadwalk_ratio_metrics_v4_20250925.py \
            --bei "${BEI_BINS}" \
            --out "${OUT_PREFIX}" \
            --min_cov 20

        Outputs created:
          ${OUT_PREFIX}_with_detrend.tsv.gz       # contains Z_ACd, Z_ABd, Z_BEI
          ${OUT_PREFIX}_detrend_coeffs.json

   B) SBATCH wrapper (provided as init_detrended_v4_20250925_1.sh):
        • EDIT these lines in the wrapper:
            BEI_FILE="__REPLACE_WITH_NEW_BEI_BINS_TSV_GZ__"  → set to step‑1 BEI_BINS path
            SCRIPT_CMD=".../calculate_detrended_cadwalk_ratio_metrics_v4_20250925.py" → confirm path
            MIN_COV=20 → keep or change if needed
        • Run:
            sbatch init_detrended_v4_20250925_1.sh
        • The wrapper writes a timestamped detrended_<ts>/ folder alongside BEI_BINS and logs metadata_<ts>.txt.

   NOTE: This step is NOT required to export BigWigs, but it’s useful for QC tables and keeping per‑chrom detrend coefficients recorded.

3) EXPORT BIGWIGS (Z_ACd.bw, Z_ABd.bw, Z_BEI.bw)
   This reproduces your original “three‑track” behavior, now with BEI replacing ABI. BEI is standardized per chromosome (robust Z), NO detrend.

   REQUIRED inputs to decide:
     BEI_BINS=</the/path/printed/in/step1/bei_<ts>/bei_bins_<ts>.tsv.gz>
     FAI=/rugpfs/fs0/risc_lab/store/risc_data/downloaded/hg38/genome/Sequence/WholeGenomeFasta/genome.fa.fai
     MINCOV=50 (what you used before)
     OUT_ROOT=</where/to/collect/bigwigs>  (a new bigwig_<ts>/ will be created inside)

   A) Direct Python:
        SCRIPTS=/rugpfs/fs0/risc_lab/scratch/jsoroczyns/CAD/CAD-C/scripts/2025_09_25
        BEI_BINS=__SET_THIS__        # same as step 2
        OUT_ROOT=__SET_THIS__: e.g.
          /rugpfs/fs0/risc_lab/scratch/jsoroczyns/CAD/CADwalk/.../merged_all/BEI_analyses_2025

        TS=$(date +"%Y%m%d_%H%M%S")
        OUT_DIR=${OUT_ROOT}/bigwig_${TS}
        mkdir -p "${OUT_DIR}"
        PREFIX=${OUT_DIR}/bei_${TS}

        python ${SCRIPTS}/BEI_to_bigwig_v5_20250925.py \
            --bei "${BEI_BINS}" \
            --fai "${FAI}" \
            --out "${PREFIX}" \
            --min_cov ${MINCOV} \
            --exclude_random

        Outputs created:
          ${PREFIX}_Z_ACd.bw
          ${PREFIX}_Z_ABd.bw
          ${PREFIX}_Z_BEI.bw

   B) SBATCH wrapper (provided as submit_genomewide_BEI_to_bigwig_v5_20250925.sh):
        • EDIT these lines in the wrapper:
            FAI=.../genome.fa.fai            → confirm path
            BEI="__REPLACE_WITH_NEW_BEI_BINS_TSV_GZ__"  → set to step‑1 BEI_BINS path
            OUT_ROOT=__SET_THIS__            → choose the parent dir for outputs
            SCRIPT=.../BEI_to_bigwig_v5_20250925.py     → confirm path
            MINCOV=50                        → keep or change if needed
            EXCLUDE_RANDOM_FLAG=--exclude_random  → leave as is (matches your previous behavior)
        • Run:
            sbatch submit_genomewide_BEI_to_bigwig_v5_20250925.sh
        • Wrapper writes a timestamped bigwig_<ts>/ under OUT_ROOT with all three tracks.

4) WHAT YOU MUST DECIDE / FILL IN
    • WALKS: path to the oriented walks table with A_chrom, A_mid, B_mid, C_mid.
      (If you don’t already have mids, compute midpoints upstream once:
         A_mid=(A_start+A_end)//2 etc., then write the 4 required columns.)
    • ANALYSIS_ROOT / OUTDIR / OUT_ROOT: pick consistent root(s). A typical choice:
         /rugpfs/fs0/risc_lab/scratch/jsoroczyns/CAD/CADwalk/CADwalk_E1_GM12878_2x250/ \
           results_20250207_0034_nonexpanded/ABC_walk_processing_20250209/ \
           20250214_Bfilt_flattened_parse2_UU_direct_ABC_complete_consequent12/ \
           filtzerolenB_flip_sort_dedup_split_20250214/merged_all/BEI_analyses_2025
      The generator will add bei_<timestamp>/ inside this.
    • SCRIPTS: where you placed the three updated Python scripts.
    • FAI: confirm genome.fa.fai path.
    • MIN_COV / MINCOV: keep your existing thresholds (20 for modeling table; 50 for BigWigs).

5) QUICK SANITY CHECKS (takes seconds)
    • Inspect the BEI bins file header:
        zcat /path/to/bei_<ts>/bei_bins_<ts>.tsv.gz | head
      Expected columns: Chrom Start End r_A r_B r_C BEI n_tot
    • Confirm Z columns in the detrended TSV:
        zcat /path/to/detrended_<ts>/bei_bins_detrended_<ts>_with_detrend.tsv.gz | \
            head -n 1
      Expected to include: Z_ACd, Z_ABd, Z_BEI
    • Confirm BigWigs exist:
        ls -lh /path/to/bigwig_<ts>/bei_<ts>_Z_*.bw

6) EXACTLY WHAT CHANGED VS. YOUR OLD ABI PIPELINE
    • ABI renamed and sign‑flipped to BEI (BEI = 2*r_B − 1).
    • We still do NO detrend on the index; we DO compute per‑chrom robust Z of BEI.
    • You still get THREE tracks: Z_ACd.bw, Z_ABd.bw, Z_BEI.bw.
    • All other behavior is unchanged.

7) IF YOU HIT A BLOCKER
    • “Input missing columns” → verify your WALKS has A_chrom, A_mid, B_mid, C_mid (exact names).
    • “pyBigWig error” → ensure pyBigWig is installed and your .fai matches the chromosome naming used in BEI bins (chr*).
    • “No bins exported” → check min coverage thresholds (try lowering --min_cov temporarily to diagnose).

That’s it. If you paste your actual WALKS path and preferred ANALYSIS_ROOT, I’ll write out the precise ready‑to‑copy commands with your concrete directories substituted.
