## Supplementary material for "CAD-C: An engineered nuclease enables repair-free *in situ* proximity ligation and nucleosome-resolution chromosome walks in human cells": Custom code: BEI_to_bigwig_v5_20250925.py

```
#!/usr/bin/env python3
################################################################################
#  BEI_to_bigwig_v5_20250925.py
#  ---------------------------------------------------------------------------
#  Author   : Jan Soroczynski
#  Version  : 5.0  ·  2025-09-25
#
#  PURPOSE
#  =======
#  Create three 1 kb BigWig tracks from a BEI‑bin table:
#      • Z_ACd  = linear‑detrended, per‑chrom robust‑Z  [log₂(A/C)]
#      • Z_ABd  = linear‑detrended, per‑chrom robust‑Z  [log₂(A/B)]
#      • Z_BEI  = per‑chrom robust‑Z                    [BEI = 2*r_B−1]  (no detrend)
################################################################################
import argparse, sys, importlib.util
from pathlib import Path
from typing import Dict
import numpy as np, pandas as pd, pyBigWig

if importlib.util.find_spec("statsmodels") is not None:
    from statsmodels import robust
    mad = lambda a: robust.mad(a, c=1.4826)
else:
    def mad(a, c: float = 1.4826) -> float:
        med = np.nanmedian(a); return c * np.nanmedian(np.abs(a - med))

def robust_z_per_chrom(df: pd.DataFrame, col: str) -> pd.Series:
    def _z(s):
        m = s.median(); mdev = mad(s.values)
        return (s - m) / mdev if mdev else np.nan
    return df.groupby("Chrom")[col].transform(_z)

def detrend_linear(series: pd.Series, positions: pd.Series) -> pd.Series:
    x = positions.values.astype(float); y = series.values.astype(float)
    if len(x) < 2 or np.all(np.isnan(y)): return series * 0.0
    x0 = x - x.mean(); coeffs = np.polyfit(x0, y, 1); trend = np.polyval(coeffs, x0)
    return series - trend

def detect_bin(df: pd.DataFrame) -> int:
    if "End" in df.columns and len(df) > 0:
        bw = int(df["End"].iloc[0] - df["Start"].iloc[0])
        if bw > 0: 
            print(f"[INFO] bin size from End column: {bw}"); return bw
    diffs = []
    for _, sub in df.groupby("Chrom"):
        d = np.diff(np.sort(sub["Start"].unique())); diffs.extend(d[d > 0])
    bw = int(pd.Series(diffs).mode().iloc[0]) if diffs else 1
    print(f"[INFO] bin size from ΔStart: {bw}")
    return bw

def read_fai(path: str) -> Dict[str, int]:
    d = {}
    with open(path) as fh:
        for line in fh:
            chrom, length, *_ = line.split(); d[chrom] = int(length)
    return d

def write_bw(out_path: str, chrom_sizes: Dict[str, int],
             df: pd.DataFrame, col: str, bin_size: int):
    bw = pyBigWig.open(out_path, "w")
    header = list(chrom_sizes.items()); bw.addHeader(header)
    for chrom, chrom_len in header:
        sub = df[df["Chrom"] == chrom][["Start", col]]
        if sub.empty: bw.addEntries([chrom], [0], [1], [0.0]); continue
        sub = sub.drop_duplicates("Start").sort_values("Start")
        s = sub["Start"].astype(np.int64).values; e = s + bin_size
        v = sub[col].astype(float).values
        mask = (s >= 0) & (e <= chrom_len) & np.isfinite(v)
        s, e, v = s[mask], e[mask], v[mask]
        if len(v) == 0: bw.addEntries([chrom], [0], [1], [0.0]); continue
        try:
            bw.addEntries([chrom]*len(v), s.tolist(), e.tolist(), v.tolist())
        except RuntimeError as err:
            print(f"[FATAL] pyBigWig error on {chrom}: {err}", file=sys.stderr)
            bw.close(); sys.exit(1)
    bw.close()

def main(argv=None):
    ap = argparse.ArgumentParser()
    ap.add_argument("--bei", required=True,
                    help="BEI bins TSV(.gz) with Chrom/Start/End, r_A/B/C; BEI optional (will be computed if absent).")
    ap.add_argument("--fai", required=True, help="FASTA index (.fai)")
    ap.add_argument("--out", required=True, help="Output prefix for BigWigs")
    ap.add_argument("--min_cov", type=int, default=0, help="Minimum n_tot per bin for export")
    ap.add_argument("--exclude_random", action="store_true", help="Exclude *_random, chrUn, EBV, KI/GL scaffolds")
    args = ap.parse_args(argv)

    df = pd.read_csv(args.bei, sep="\t")
    df = df[df["Chrom"].astype(str).str.startswith("chr")]
    if args.exclude_random:
        mask_rand = df["Chrom"].str.contains("_random|_KI|_GL|chrUn|EBV"); df = df[~mask_rand]
    if args.min_cov and "n_tot" in df.columns:
        df = df[df["n_tot"] >= args.min_cov]
    elif args.min_cov:
        print("[WARN] n_tot absent – --min_cov ignored", file=sys.stderr)

    for col in ("r_A","r_B","r_C"):
        if col not in df.columns:
            print(f"[FATAL] Missing required column: {col}", file=sys.stderr); sys.exit(1)
    if "BEI" not in df.columns:
        df["BEI"] = 2.0 * df["r_B"].astype(float) - 1.0

    bin_size = detect_bin(df)

    df["log2_AC_raw"] = np.log2(df["r_A"].where(df["r_C"] > 0) / df["r_C"])
    df["log2_AB_raw"] = np.log2(df["r_A"].where(df["r_B"] > 0) / df["r_B"])
    df["log2_AC_detrend"] = df.groupby("Chrom", group_keys=False)\
                              .apply(lambda sub: detrend_linear(sub["log2_AC_raw"], sub["Start"]))
    df["log2_AB_detrend"] = df.groupby("Chrom", group_keys=False)\
                              .apply(lambda sub: detrend_linear(sub["log2_AB_raw"], sub["Start"]))
    df["Z_ACd"] = robust_z_per_chrom(df, "log2_AC_detrend")
    df["Z_ABd"] = robust_z_per_chrom(df, "log2_AB_detrend")
    df["Z_BEI"] = robust_z_per_chrom(df, "BEI")

    chrom_sizes = read_fai(args.fai)
    prefix = Path(args.out).as_posix()
    write_bw(f"{prefix}_Z_ACd.bw",  chrom_sizes, df, "Z_ACd", bin_size)
    write_bw(f"{prefix}_Z_ABd.bw",  chrom_sizes, df, "Z_ABd", bin_size)
    write_bw(f"{prefix}_Z_BEI.bw",  chrom_sizes, df, "Z_BEI", bin_size)
    print("[SUCCESS] BigWigs:", f"{prefix}_Z_ACd.bw  {prefix}_Z_ABd.bw  {prefix}_Z_BEI.bw")

if __name__ == "__main__":
    main()
```
