## Supplementary material for "CAD-C: An engineered nuclease enables repair-free *in situ* proximity ligation and nucleosome-resolution chromosome walks in human cells": Custom code: CADwalk_calculate_BEI_v2_20250925.py

```
#!/usr/bin/env python3
# =========================================================================================
#  CADwalk_calculate_BEI_v2_20250925.py
#  ----------------------------------------------------------------------------------------
#  Compute BEI (B‑Enrichment Index) from three‑fragment walks (A,B,C midpoints binned)
#
#  Version : 2.0   •   Date : 2025-09-25
#  Author  : Jan Soroczynski, Risca Lab, The Rockefeller University
#
#  README
#  ------
#  PURPOSE
#    Convert a large CAD‑walk TSV(.gz) into binned counts of A, B, C fragment midpoints,
#    normalise to compositions, and output the **B‑Enrichment Index (BEI)**:
#
#        BEI = 2 * r_B - 1
#
#    with r_X = (n_X + 0.5) / (n_tot + 1) for X ∈ {A,B} and r_C = 1 − r_A − r_B.
#    BEI is **independent of the A/C split** and is **not detrended**. It is the
#    sign‑reversed form of legacy ABI (ABI_legacy = 1 − 2 r_B), so higher BEI means
#    greater B‑fragment enrichment per genomic bin.
#
#  INPUT
#    TSV(.gz) with header (comment lines starting “#” allowed). Required fields:
#      A_chrom, A_mid, B_mid, C_mid. Walks pre‑oriented so A upstream of C.
#
#  OUTPUT (to outdir/bei_<timestamp>/)
#    bei_bins_<ts>.tsv.gz   : Chrom Start End r_A r_B r_C BEI n_tot
#    raw_counts_<ts>.tsv.gz : Chrom Start End n_A n_B n_C
#    metadata_<ts>.json     : parameters, checksum, and BEI semantics
# =========================================================================================
from __future__ import annotations
import argparse, gzip, json, logging, hashlib
from datetime import datetime
from pathlib import Path
from typing import Dict, Tuple
import numpy as np, pandas as pd
pd.options.mode.copy_on_write = True

def sha256_of_file(p: Path) -> str:
    h = hashlib.sha256(); h.update(p.read_bytes()); return h.hexdigest()

def parse_region(arg: str) -> Tuple[str, int, int]:
    chrom, rest = arg.split(":"); start, end = rest.replace(",", "").split("-")
    return chrom, int(float(start)), int(float(end))

def ensure_dir(p: Path): p.mkdir(parents=True, exist_ok=True)

def main():
    ap = argparse.ArgumentParser("BEI (B-positive) calculator",
        formatter_class=argparse.ArgumentDefaultsHelpFormatter)
    ap.add_argument("--walks", required=True)
    ap.add_argument("--outdir", required=True)
    ap.add_argument("--bin_size", type=int, default=10_000)
    ap.add_argument("--chrom", default=None)
    ap.add_argument("--region", default=None)
    ap.add_argument("--chunk_rows", type=int, default=2_000_000)
    args = ap.parse_args()

    ts = datetime.now().strftime("%Y%m%d_%H%M%S")
    outdir = Path(args.outdir).expanduser().resolve() / f"bei_{ts}"
    ensure_dir(outdir)
    logging.basicConfig(filename=outdir/"run.log", level=logging.INFO,
                        format="%(asctime)s | %(levelname)s | %(message)s")
    logging.info("Parameters: %s", vars(args))

    region_chr = region_start = region_end = None
    if args.region:
        region_chr, region_start, region_end = parse_region(args.region)
    elif args.chrom:
        region_chr = args.chrom

    counts_A: Dict[str, np.ndarray] = {}
    counts_B: Dict[str, np.ndarray] = {}
    counts_C: Dict[str, np.ndarray] = {}
    bin_size = args.bin_size

    def add_count(dct, chrom: str, bin_idx: np.ndarray):
        max_idx = int(bin_idx.max())
        arr = dct.setdefault(chrom, np.zeros(max_idx + 1, dtype=np.int64))
        if max_idx >= len(arr):
            arr.resize(max_idx + 1, refcheck=False)
        np.add.at(arr, bin_idx, 1)

    usecols = ["A_chrom","A_mid","B_mid","C_mid"]
    walks_path = Path(args.walks).expanduser()
    open_fn = gzip.open if walks_path.suffix.endswith(".gz") else open
    with open_fn(walks_path, "rt") as fh:
        reader = pd.read_csv(fh, sep="\t", comment="#", header=0, usecols=usecols,
                             chunksize=args.chunk_rows, low_memory=False)
        for cno, chunk in enumerate(reader, 1):
            if region_chr:
                chunk = chunk[chunk["A_chrom"] == region_chr]
            if region_start is not None:
                m = ((chunk["A_mid"] >= region_start) & (chunk["A_mid"] < region_end)) \
                    | ((chunk["B_mid"] >= region_start) & (chunk["B_mid"] < region_end)) \
                    | ((chunk["C_mid"] >= region_start) & (chunk["C_mid"] < region_end))
                chunk = chunk[m]
            if chunk.empty: 
                logging.info("chunk %-4d | rows 0 (post-filter)", cno); 
                continue

            for col in ("A_mid","B_mid","C_mid"):
                chunk[col] = pd.to_numeric(chunk[col], errors="coerce")
            chunk.dropna(subset=["A_mid","B_mid","C_mid"], inplace=True)
            if chunk.empty:
                logging.info("chunk %-4d | rows 0 (post-numeric)", cno); 
                continue
            chunk[["A_mid","B_mid","C_mid"]] = chunk[["A_mid","B_mid","C_mid"]].astype(np.int64)

            chroms = chunk["A_chrom"].to_numpy()
            a_bins = (chunk["A_mid"].to_numpy() // bin_size).astype(np.int64)
            b_bins = (chunk["B_mid"].to_numpy() // bin_size).astype(np.int64)
            c_bins = (chunk["C_mid"].to_numpy() // bin_size).astype(np.int64)

            for chrom in np.unique(chroms):
                m = chroms == chrom
                add_count(counts_A, chrom, a_bins[m])
                add_count(counts_B, chrom, b_bins[m])
                add_count(counts_C, chrom, c_bins[m])

            logging.info("chunk %-4d | rows %-7d", cno, len(chunk))

    rows = []
    for chrom in sorted(counts_A.keys()):
        n_bins = max(len(counts_A.get(chrom, [])),
                     len(counts_B.get(chrom, [])),
                     len(counts_C.get(chrom, [])))
        for d in (counts_A, counts_B, counts_C):
            if chrom not in d:
                d[chrom] = np.zeros(n_bins, dtype=np.int64)
            elif len(d[chrom]) < n_bins:
                d[chrom].resize(n_bins, refcheck=False)

        a, b, c = counts_A[chrom], counts_B[chrom], counts_C[chrom]
        n_tot = a + b + c
        sel = n_tot > 0
        if not np.any(sel): 
            continue
        idx = np.nonzero(sel)[0]
        starts = idx * bin_size
        ends   = starts + bin_size

        r_A = (a[sel] + 0.5) / (n_tot[sel] + 1.0)
        r_B = (b[sel] + 0.5) / (n_tot[sel] + 1.0)
        r_C = 1.0 - r_A - r_B
        bei = 2.0 * r_B - 1.0

        rows.append(pd.DataFrame({
            "Chrom": chrom, "Start": starts, "End": ends,
            "r_A": r_A, "r_B": r_B, "r_C": r_C, "BEI": bei, "n_tot": n_tot[sel]
        }))

    result_df = pd.concat(rows, ignore_index=True) if rows else pd.DataFrame(
        columns=["Chrom","Start","End","r_A","r_B","r_C","BEI","n_tot"])

    bei_path  = outdir / f"bei_bins_{ts}.tsv.gz"
    raw_path  = outdir / f"raw_counts_{ts}.tsv.gz"
    result_df.to_csv(bei_path, sep="\t", compression="gzip", index=False,
                     columns=["Chrom","Start","End","r_A","r_B","r_C","BEI","n_tot"])

    if rows:
        nA_list, nB_list, nC_list, cs_list, st_list = [], [], [], [], []
        for chrom in sorted(counts_A.keys()):
            a, b, c = counts_A[chrom], counts_B[chrom], counts_C[chrom]
            sel = (a + b + c) > 0
            if sel.any():
                idx = np.nonzero(sel)[0]; st = idx * bin_size
                nA_list.append(a[sel]); nB_list.append(b[sel]); nC_list.append(c[sel])
                cs_list.append(np.array([chrom]*len(idx), dtype=object))
                st_list.append(st)
        raw_df = pd.DataFrame({
            "Chrom": np.concatenate(cs_list) if cs_list else np.array([], dtype=object),
            "Start": np.concatenate(st_list) if st_list else np.array([], dtype=np.int64),
            "End":   (np.concatenate(st_list) + bin_size) if st_list else np.array([], dtype=np.int64),
            "n_A":   np.concatenate(nA_list) if nA_list else np.array([], dtype=np.int64),
            "n_B":   np.concatenate(nB_list) if nB_list else np.array([], dtype=np.int64),
            "n_C":   np.concatenate(nC_list) if nC_list else np.array([], dtype=np.int64),
        })
        raw_df.to_csv(raw_path, sep="\t", compression="gzip", index=False)
    else:
        pd.DataFrame(columns=["Chrom","Start","End","n_A","n_B","n_C"]).to_csv(
            raw_path, sep="\t", compression="gzip", index=False)

    meta = {
        "timestamp": ts, "bin_size": args.bin_size,
        "region": args.region or args.chrom or "genome",
        "script_sha256": sha256_of_file(Path(__file__)),
        "walks_file": str(walks_path),
        "index_name": "BEI (B-Enrichment Index)",
        "index_formula": "2*r_B - 1",
        "replaces_ABI_legacy": True,
        "legacy_relation": "BEI = -ABI_legacy"
    }
    with open(outdir/f"metadata_{ts}.json", "w") as fh:
        json.dump(meta, fh, indent=2)

    print(f"BEI table   → {bei_path}")
    print(f"Raw counts  → {raw_path}")

if __name__ == "__main__":
    main()
```
