## Supplementary material for "CAD-C: An engineered nuclease enables repair-free *in situ* proximity ligation and nucleosome-resolution chromosome walks in human cells": Custom code: calculate_detrended_cadwalk_ratio_metrics_v4_20250925.py

```
#!/usr/bin/env python3
################################################################################
#  calculate_detrended_cadwalk_ratio_metrics_v4_20250925.py
#  ---------------------------------------------------------------------------
#  Author   : Jan Soroczynski, Risca Lab, The Rockefeller University
#  Version  : 4.0  ·  2025-09-25
#
#  WHAT THIS SCRIPT DOES
#  =====================
#  Given a 1 kb BEI-bin table (plain or gzipped TSV) it appends:
#
#    ─── Raw ratios ──────────────────────────────────────────────────────────
#        • log2_AC_raw   = log₂( r_A / r_C )
#        • log2_AB_raw   = log₂( r_A / r_B )
#
#    ─── Chromosome-wise linear fit coefficients ────────────────────────────
#        For each chrom:  y = slope * (Start − mean) + intercept
#        • AC_slope , AC_intercept
#        • AB_slope , AB_intercept
#
#    ─── Residuals (detrended values) ───────────────────────────────────────
#        • log2_AC_detrend , log2_AB_detrend
#
#    ─── Robust‑Z (median 0, MAD 1, per chromosome) ─────────────────────────
#        • Z_ACd  – robust‑Z of detrended log₂(A/C)
#        • Z_ABd  – robust‑Z of detrended log₂(A/B)
#        • Z_BEI  – robust‑Z of **raw BEI** (no detrend)
#
#  IMPORTANT NOTES
#    • BEI = 2*r_B − 1 is independent of A/C and is not detrended; we only
#      standardize it per chromosome via robust Z.
#
#  OUTPUTS
#    <out_prefix>_with_detrend.tsv.gz   : augmented, gzip‑compressed table
#    <out_prefix>_detrend_coeffs.json   : per‑chrom slopes & intercepts
################################################################################
import argparse, json, sys, importlib.util
from typing import Dict
import numpy as np, pandas as pd

if importlib.util.find_spec("statsmodels") is not None:
    from statsmodels import robust
    def mad(arr): return robust.mad(arr, c=1.4826)
else:
    def mad(arr, c: float = 1.4826) -> float:
        med = np.nanmedian(arr)
        return c * np.nanmedian(np.abs(arr - med))

def robust_z(series: pd.Series) -> pd.Series:
    med = series.median()
    mdev = mad(series.values)
    return (series - med) / mdev if mdev else np.nan

def detrend(sub: pd.DataFrame, col: str):
    x = sub["Start"].astype(float).values
    y = sub[col].astype(float).values
    if len(x) < 2 or np.all(np.isnan(y)):
        return pd.Series(np.nan, index=sub.index), 0.0, 0.0
    x0 = x - x.mean()
    slope, intercept = np.polyfit(x0, y, 1)
    resid = y - (slope * x0 + intercept)
    return pd.Series(resid, index=sub.index), float(slope), float(intercept)

def main(argv=None):
    ap = argparse.ArgumentParser(
        description="Append detrended ratio Z-scores and Z_BEI (no detrend on BEI).")
    ap.add_argument("--bei", required=True,
                    help="BEI bin TSV(.gz) with Chrom/Start/End, r_A/r_B/r_C, and BEI (BEI will be computed if absent).")
    ap.add_argument("--out", required=True, help="output prefix (no extension)")
    ap.add_argument("--min_cov", type=int, default=0, help="minimum n_tot filter before modeling")
    args = ap.parse_args(argv)

    df = pd.read_csv(args.bei, sep="\t")
    if "Chrom" not in df.columns or "Start" not in df.columns:
        print("[FATAL] Input must have Chrom and Start columns.", file=sys.stderr); sys.exit(1)
    df = df[df["Chrom"].astype(str).str.startswith("chr")]

    if args.min_cov and "n_tot" in df.columns:
        before = len(df); df = df[df["n_tot"] >= args.min_cov]
        print(f"[INFO] coverage filter: kept {len(df)}/{before} bins (n_tot ≥ {args.min_cov})")
    elif args.min_cov:
        print("[WARN] n_tot column absent – --min_cov ignored", file=sys.stderr)

    for col in ("r_A","r_B","r_C"):
        if col not in df.columns:
            print(f"[FATAL] Missing required column: {col}", file=sys.stderr); sys.exit(1)
    if "BEI" not in df.columns:
        df["BEI"] = 2.0 * df["r_B"].astype(float) - 1.0

    df["log2_AC_raw"] = np.log2(df["r_A"].where(df["r_C"] > 0) / df["r_C"])
    df["log2_AB_raw"] = np.log2(df["r_A"].where(df["r_B"] > 0) / df["r_B"])

    coeffs: Dict[str, Dict[str, float]] = {}
    df["log2_AC_detrend"] = np.nan
    df["log2_AB_detrend"] = np.nan
    for chrom, sub in df.groupby("Chrom"):
        res_ac, s_ac, i_ac = detrend(sub, "log2_AC_raw")
        res_ab, s_ab, i_ab = detrend(sub, "log2_AB_raw")
        idx = sub.index
        df.loc[idx, "log2_AC_detrend"] = res_ac
        df.loc[idx, "log2_AB_detrend"] = res_ab
        coeffs[chrom] = dict(AC_slope=s_ac, AC_intercept=i_ac,
                             AB_slope=s_ab, AB_intercept=i_ab)

    df["Z_ACd"] = df.groupby("Chrom")["log2_AC_detrend"].transform(robust_z)
    df["Z_ABd"] = df.groupby("Chrom")["log2_AB_detrend"].transform(robust_z)
    df["Z_BEI"] = df.groupby("Chrom")["BEI"].transform(robust_z)  # no detrend

    out_tsv = f"{args.out}_with_detrend.tsv.gz"
    df.to_csv(out_tsv, sep="\t", index=False, compression="gzip")
    with open(f"{args.out}_detrend_coeffs.json", "w") as fh:
        json.dump(coeffs, fh, indent=2)

    print(f"[DONE] augmented table → {out_tsv}")
    print(f"[DONE] coeffs JSON    → {args.out}_detrend_coeffs.json")

if __name__ == "__main__":
    main()
```
