## Supplementary material for "CAD-C: An engineered nuclease enables repair-free *in situ* proximity ligation and nucleosome-resolution chromosome walks in human cells": Custom code: GM_deepmerge_highlights_3_darker55_600dpi.sh

```
#!/bin/bash
#SBATCH --job-name=plot_gm_9regions
#SBATCH --output=plot_gm_9regions_%j.log
#SBATCH --error=plot_gm_9regions_%j.err
#SBATCH --cpus-per-task=4

set -e

# ==============================================================================
# USER INPUTS
# ==============================================================================

# 1. Script Path (Using v3 with nearest-neighbor interpolation)
SCRIPT_PATH="/rugpfs/fs0/risc_lab/scratch/jsoroczyns/CAD/CAD-C/scripts/2025_11_26/replot_raw_balanced_N_matrices_waxes_nointerpol_v3_20251128.py"

# 2. Output Directory
BASE_OUT_DIR="/rugpfs/fs0/risc_lab/scratch/jsoroczyns/CAD/CAD-C/scripts/2025_11_26"
TIMESTAMP=$(date +"%Y%m%d_%H%M%S")
OUT_DIR="${BASE_OUT_DIR}/GM12878_Deep_9Regions_Final_${TIMESTAMP}"

# 3. Input File (GM12878 Deep Merge)
MCOOL="/rugpfs/fs0/risc_lab/scratch/jsoroczyns/CAD/CAD-C/CADC_GM12878_deepmerge_20250205/cooler_out_20250205_023558/GM12878_deepmerge_20250205_011600_dedup_500_bal_500N_igdiag1.mcool"

# 4. Label
LABEL="--label GM12878_Deepmerge"

# 5. Environment
export OMP_NUM_THREADS="${SLURM_CPUS_PER_TASK:-1}"

# ==============================================================================
# PLOTTING COMMANDS
# ==============================================================================

mkdir -p "$OUT_DIR"
echo "Starting job..."
echo "Outputting to: $OUT_DIR"
echo "Using DPI: 600 (High resolution for pixel-perfect rendering)"

# 1. chr1:202Mb (2.5kb) | Bins: ~1688 | Vmax: 0.0135
echo "Plotting 1..."
python "$SCRIPT_PATH" "$MCOOL" $LABEL \
    --region "chr1:202,374,167-206,595,716" \
    --res 2500 \
    --bal-vmax 0.0135 \
    --no-raw \
    --dpi 600 \
    --out "$OUT_DIR/01_chr1_202Mb_2.5kb.pdf"

# 2. chr1:24Mb (25kb) | Bins: ~843 | Vmax: 0.0315
echo "Plotting 2..."
python "$SCRIPT_PATH" "$MCOOL" $LABEL \
    --region "chr1:24,395,953-45,475,504" \
    --res 25000 \
    --bal-vmax 0.0315 \
    --no-raw \
    --dpi 600 \
    --out "$OUT_DIR/02_chr1_24Mb_25kb.pdf"

# 3. chr1:33Mb (5kb) | Bins: ~1346 | Vmax: 0.0225
echo "Plotting 3..."
python "$SCRIPT_PATH" "$MCOOL" $LABEL \
    --region "chr1:33,361,669-40,097,109" \
    --res 5000 \
    --bal-vmax 0.0225 \
    --no-raw \
    --dpi 600 \
    --out "$OUT_DIR/03_chr1_33Mb_5kb.pdf"

# 4. chr1:27Mb (2.5kb) | Bins: ~1652 | Vmax: 0.01125
echo "Plotting 4..."
python "$SCRIPT_PATH" "$MCOOL" $LABEL \
    --region "chr1:27,576,343-31,705,285" \
    --res 2500 \
    --bal-vmax 0.01125 \
    --no-raw \
    --dpi 600 \
    --out "$OUT_DIR/04_chr1_27Mb_2.5kb.pdf"

# 5. chr8:100Mb (2.5kb) | Bins: ~1500 | Vmax: 0.0081
echo "Plotting 5..."
python "$SCRIPT_PATH" "$MCOOL" $LABEL \
    --region "chr8:100,000,000-103,750,000" \
    --res 2500 \
    --bal-vmax 0.0081 \
    --no-raw \
    --dpi 600 \
    --out "$OUT_DIR/05_chr8_100Mb_2.5kb.pdf"

# 6. chr1:205Mb (500bp) | Bins: ~990 | Vmax: 0.0036
echo "Plotting 6..."
python "$SCRIPT_PATH" "$MCOOL" $LABEL \
    --region "chr1:205,315,697-205,810,791" \
    --res 500 \
    --bal-vmax 0.0036 \
    --no-raw \
    --dpi 600 \
    --out "$OUT_DIR/06_chr1_205Mb_500bp.pdf"

# 7. chr8:122Mb (2.5kb) | Bins: ~1768 | Vmax: 0.00675
echo "Plotting 7..."
python "$SCRIPT_PATH" "$MCOOL" $LABEL \
    --region "chr8:122,296,834-126,723,363" \
    --res 2500 \
    --bal-vmax 0.00675 \
    --no-raw \
    --dpi 600 \
    --out "$OUT_DIR/07_chr8_122Mb_2.5kb.pdf"

# 8. chr12:90Mb (2.5kb) | Bins: ~1500 | Vmax: 0.01125
echo "Plotting 8..."
python "$SCRIPT_PATH" "$MCOOL" $LABEL \
    --region "chr12:90,750,000-94,500,000" \
    --res 2500 \
    --bal-vmax 0.01125 \
    --no-raw \
    --dpi 600 \
    --out "$OUT_DIR/08_chr12_90Mb_2.5kb.pdf"

# 9. chr11:127Mb (1kb) | Bins: ~1340 | Vmax: 0.009
echo "Plotting 9..."
python "$SCRIPT_PATH" "$MCOOL" $LABEL \
    --region "chr11:127,832,229-129,177,988" \
    --res 1000 \
    --bal-vmax 0.009 \
    --no-raw \
    --dpi 600 \
    --out "$OUT_DIR/09_chr11_127Mb_1kb.pdf"

echo "Done."
```
