## Supplementary material for "CAD-C: An engineered nuclease enables repair-free *in situ* proximity ligation and nucleosome-resolution chromosome walks in human cells": Custom code: replot_raw_balanced_N_matrices_waxes_nointerpol_v3_20251128.py

```
#!/usr/bin/env python
"""
plot_triples v3: compare raw & balanced Hi‑C / Micro‑C contact matrices from multiple coolers.

Changes vs v2:
- Force nearest-neighbor (no interpolation) for all heatmaps → perfect square pixels.
- Keep FontType=42 + sans-serif defaults.
- Keep adaptive genomic ticks.
- Move colorbars to dedicated axes to the right of each row (no overlap with matrices).
"""

from __future__ import annotations

import os
import math
from dataclasses import dataclass
from datetime import datetime
from typing import Sequence, Optional, Tuple, List

import click
import numpy as np
import matplotlib as mpl
import matplotlib.pyplot as plt
from matplotlib.colors import LogNorm, Normalize

# Font / output configuration:
# - Force TrueType fonts in PDF/PS output (Type 42).
# - Prefer sans-serif fonts (Arial / DejaVu Sans / Helvetica).
mpl.rcParams["pdf.fonttype"] = 42
mpl.rcParams["ps.fonttype"] = 42
mpl.rcParams["text.usetex"] = False
mpl.rcParams.setdefault("font.family", "sans-serif")
mpl.rcParams.setdefault("font.sans-serif", ["Arial", "DejaVu Sans", "Helvetica"])

import cooler
from cooler import fileops as cfileops
import cooltools.lib.plotting  # registers 'fall' and other colormaps


# ---------------------------------------------------------------------------
# COOL_PATH resolution & region parsing
# ---------------------------------------------------------------------------


def _resolve_cooler_uri(path: str, res: Optional[int]) -> str:
    """
    Normalize a user-provided COOL_PATH into a proper Cooler URI.
    """
    # Already a cooler URI (e.g. file.mcool::/resolutions/250000)
    if "::" in path:
        return path

    # Single-resolution cooler?
    if cfileops.is_cooler(path):
        return path

    # Multi-resolution cooler?
    if cfileops.is_multires_file(path):
        coolers = cfileops.list_coolers(path)
        if not coolers:
            raise click.ClickException(
                f"File {path!r} is a multi-resolution cooler but contains no cooler groups."
            )

        if res is not None:
            # Look for a group whose trailing component matches the requested res
            target_suffix = f"/{res}"
            matches = [g for g in coolers if g.endswith(target_suffix)]
            if not matches:
                raise click.ClickException(
                    f"Resolution {res} not found in multi-res cooler {path!r}. "
                    f"Available groups: {coolers}"
                )
            grouppath = matches[0]
        else:
            # Auto: choose highest resolution = smallest bin size.
            parsed: List[Tuple[int, str]] = []
            for g in coolers:
                tail = g.rstrip("/").split("/")[-1]
                try:
                    binsize = int(tail)
                except ValueError:
                    continue
                parsed.append((binsize, g))

            if parsed:
                parsed.sort(key=lambda x: x[0])
                _, grouppath = parsed[0]
            else:
                # Fallback: just take the first cooler group.
                grouppath = coolers[0]

        return f"{path}::{grouppath.lstrip('/')}"  # ensure no duplicate '/'

    # Not recognized as cooler or multi-res
    raise click.ClickException(
        f"Path {path!r} is not a valid COOL_PATH or multi-resolution cooler file. "
        "Pass a .cool file, a cooler URI like 'file.mcool::/resolutions/250000', "
        "or a .mcool (optionally with --res to specify the resolution)."
    )


def _normalize_region(region: str, clr: cooler.Cooler) -> Tuple[str, str, int, int]:
    """
    Parse a genomic region string and normalize it to 'chrom:start-end' form.
    """
    region = region.strip()
    if ":" not in region:
        chrom = region.strip()
        start = 0
        try:
            chrom_len = int(clr.chromsizes[chrom])
        except KeyError as e:
            raise click.BadParameter(
                f"Chromosome {chrom!r} not found in cooler {clr.uri}"
            ) from e
        end = chrom_len
    else:
        chrom, rest = region.split(":", 1)
        chrom = chrom.strip()
        try:
            start_str, end_str = rest.split("-", 1)
        except ValueError as e:
            raise click.BadParameter(
                f"Region {region!r} must be of the form 'chrom:start-end'"
            ) from e

        def _parse_coord(s: str) -> int:
            s = s.replace(",", "").strip()
            if not s:
                raise click.BadParameter(
                    f"Empty coordinate in region spec {region!r}"
                )
            try:
                return int(s)
            except ValueError as e:
                raise click.BadParameter(
                    f"Could not parse coordinate {s!r} in region {region!r}"
                ) from e

        start = _parse_coord(start_str)
        end = _parse_coord(end_str)

        if start < 0 or end <= start:
            raise click.BadParameter(
                f"Region {region!r} has invalid coordinates: start={start}, end={end}"
            )

        # Ensure chromosome exists and that end is not beyond chrom length
        try:
            chrom_len = int(clr.chromsizes[chrom])
        except KeyError as e:
            raise click.BadParameter(
                f"Chromosome {chrom!r} not found in cooler {clr.uri}"
            ) from e
        if end > chrom_len:
            raise click.BadParameter(
                f"Region {region!r} extends beyond chromosome length "
                f"(len={chrom_len}); got end={end}."
            )

    region_str = f"{chrom}:{start}-{end}"
    return region_str, chrom, start, end


# ---------------------------------------------------------------------------
# Balancing & autoscaling
# ---------------------------------------------------------------------------

_MAX_VLIM_SAMPLES = 200_000


def _compute_balanced_from_weights(
    mat_raw: np.ndarray,
    weights: np.ndarray,
    *,
    uri: str,
    region_str: str,
    clr_weight_name: str,
) -> np.ndarray:
    """
    Apply bin-wise balancing weights to a raw contact matrix.
    """
    if mat_raw.shape[0] != mat_raw.shape[1]:
        raise ValueError("mat_raw must be square")
    if mat_raw.shape[0] != weights.shape[0]:
        raise ValueError(
            "Length of weights does not match matrix size: "
            f"{weights.shape[0]} vs {mat_raw.shape[0]}"
        )

    w = np.asarray(weights, dtype=float)
    valid = np.isfinite(w) & (w != 0)

    if not np.any(valid):
        raise click.ClickException(
            f"All weights in column {clr_weight_name!r} are invalid (NaN/Inf/0) "
            f"for region {region_str!r} in cooler {uri!r}."
        )

    # Replace invalid weights with 0 for multiplication
    w_clean = w.copy()
    w_clean[~valid] = 0.0

    mat_bal = mat_raw * w_clean[:, None] * w_clean[None, :]

    # Mask rows/cols corresponding to invalid bins back to NaN
    if np.any(~valid):
        mask = ~valid
        mat_bal = np.asarray(mat_bal, dtype=float)
        mat_bal[mask, :] = np.nan
        mat_bal[:, mask] = np.nan

    return mat_bal


def _compute_joint_vlim_log(
    matrices: Sequence[np.ndarray],
    vmin_opt: Optional[float],
    vmax_opt: Optional[float],
    *,
    q_low: float,
    q_high: float,
    floor: Optional[float] = None,
) -> Tuple[float, float]:
    """
    Compute vmin/vmax for log scaling (LogNorm) jointly across matrices.
    """
    all_vals = []
    rng = np.random.default_rng(0)

    for m in matrices:
        if m is None:
            continue
        arr = np.asarray(m, dtype=float)
        mask = np.isfinite(arr) & (arr > 0)
        if not np.any(mask):
            continue
        vals = arr[mask].ravel()
        if vals.size > _MAX_VLIM_SAMPLES:
            idx = rng.choice(vals.size, size=_MAX_VLIM_SAMPLES, replace=False)
            vals = vals[idx]
        all_vals.append(vals)

    if not all_vals:
        raise click.ClickException(
            "No positive finite values found to determine color scale "
            "for log scaling."
        )

    data = np.concatenate(all_vals)

    if vmin_opt is not None:
        vmin = float(vmin_opt)
    else:
        vmin = float(np.percentile(data, q_low))
        if floor is not None:
            vmin = max(vmin, floor)

    if vmin <= 0:
        raise click.BadParameter(
            f"For log scaling, vmin must be > 0. Got vmin={vmin}."
        )

    if vmax_opt is not None:
        vmax = float(vmax_opt)
    else:
        vmax = float(np.percentile(data, q_high))

    if vmax <= vmin:
        raise click.BadParameter(
            f"For log scaling, vmax must be greater than vmin. Got vmin={vmin}, vmax={vmax}."
        )

    return vmin, vmax


def _compute_joint_vlim_linear(
    matrices: Sequence[np.ndarray],
    vmin_opt: Optional[float],
    vmax_opt: Optional[float],
    *,
    q_low: float,
    q_high: float,
) -> Tuple[float, float]:
    """
    Compute vmin/vmax for linear scaling jointly across matrices.
    """
    all_vals = []
    rng = np.random.default_rng(0)

    for m in matrices:
        if m is None:
            continue
        arr = np.asarray(m, dtype=float)
        mask = np.isfinite(arr)
        if not np.any(mask):
            continue
        vals = arr[mask].ravel()
        if vals.size > _MAX_VLIM_SAMPLES:
            idx = rng.choice(vals.size, size=_MAX_VLIM_SAMPLES, replace=False)
            vals = vals[idx]
        all_vals.append(vals)

    if not all_vals:
        raise click.ClickException(
            "No finite values found to determine color scale for linear scaling."
        )

    data = np.concatenate(all_vals)

    if vmin_opt is not None:
        vmin = float(vmin_opt)
    else:
        vmin = float(np.percentile(data, q_low))

    if vmax_opt is not None:
        vmax = float(vmax_opt)
    else:
        vmax = float(np.percentile(data, q_high))

    if vmax <= vmin:
        raise click.BadParameter(
            f"For linear scaling, vmax must be greater than vmin. "
            f"Got vmin={vmin}, vmax={vmax}."
        )

    return vmin, vmax


# ---------------------------------------------------------------------------
# Genomic tick computation (adaptive Mb / kb / bp axes)
# ---------------------------------------------------------------------------


def _compute_genomic_ticks(
    start_bp: int,
    end_bp: int,
    n_bins: int,
    max_ticks: int = 6,
) -> Tuple[np.ndarray, List[str], str]:
    """
    Compute tick positions (in matrix bin coordinates) and labels (strings)
    for a genomic axis spanning [start_bp, end_bp).
    """
    length = end_bp - start_bp
    if length <= 0 or n_bins <= 0:
        return np.array([]), [], "bp"

    bin_size = length / float(n_bins)
    if bin_size <= 0:
        return np.array([]), [], "bp"

    # Target ~max_ticks labels, but never smaller than one bin
    raw_step = max(length / max_ticks, bin_size)

    # Use a 1–2–5×10^k pattern for "nice" steps
    exp = math.floor(math.log10(raw_step))
    base = 10 ** exp
    step_bp = None
    for m in (1, 2, 5, 10):
        step = m * base
        if step >= raw_step:
            step_bp = step
            break
    if step_bp is None:
        step_bp = 10 * base

    # Decide units from step size
    if step_bp >= 1_000_000:
        unit = "Mb"
        factor = 1_000_000.0
        fmt = "{:.1f}"
    elif step_bp >= 1_000:
        unit = "kb"
        factor = 1_000.0
        fmt = "{:.0f}"
    else:
        unit = "bp"
        factor = 1.0
        fmt = "{:.0f}"

    # First tick: first multiple of step_bp >= start_bp
    first_tick = math.ceil(start_bp / step_bp) * step_bp
    if first_tick < start_bp:
        first_tick += step_bp
    if first_tick >= end_bp:
        return np.array([]), [], unit

    coords_bp = np.arange(first_tick, end_bp + 1, step_bp, dtype=float)

    # Map genomic coords → bin index (0..n_bins-1); align to left bin edge
    tick_positions = (coords_bp - start_bp) / bin_size

    # Keep ticks inside the matrix
    mask = (tick_positions >= 0) & (tick_positions <= n_bins - 1)
    tick_positions = tick_positions[mask]
    coords_bp = coords_bp[mask]

    tick_labels = [fmt.format(c / factor) for c in coords_bp]

    return tick_positions, tick_labels, unit


# ---------------------------------------------------------------------------
# Dataset loading
# ---------------------------------------------------------------------------


def _load_dataset_matrices(
    clr: cooler.Cooler,
    uri: str,
    region_str: str,
    chrom: str,
    clr_weight_name: str,
    want_raw: bool,
    want_bal: bool,
) -> Tuple[Optional[np.ndarray], Optional[np.ndarray]]:
    """
    Fetch raw and/or balanced matrices for a single dataset.
    """
    if chrom not in clr.chromsizes:
        raise click.ClickException(
            f"Chromosome {chrom!r} not found in cooler {uri!r}."
        )

    mat_raw: Optional[np.ndarray] = None
    mat_bal: Optional[np.ndarray] = None

    if want_raw or want_bal:
        try:
            mat_raw = clr.matrix(balance=False).fetch(region_str)
        except Exception as e:
            raise click.ClickException(
                f"Failed to fetch raw matrix for region {region_str!r} from {uri!r}: {e}"
            ) from e

    if want_bal:
        try:
            bins_region = clr.bins().fetch(region_str)
        except Exception as e:
            raise click.ClickException(
                f"Failed to fetch bins for region {region_str!r} from {uri!r}: {e}"
            ) from e

        if clr_weight_name not in bins_region.columns:
            raise click.ClickException(
                f"Cooler {uri!r} has no bin weight column {clr_weight_name!r} "
                f"in clr.bins(). Available columns: {list(bins_region.columns)}"
            )

        weights = bins_region[clr_weight_name].to_numpy()
        mat_bal = _compute_balanced_from_weights(
            mat_raw,
            weights,
            uri=uri,
            region_str=region_str,
            clr_weight_name=clr_weight_name,
        )

    return mat_raw, mat_bal


# ---------------------------------------------------------------------------
# Core plotting logic (pure Python, no Click parsing)
# ---------------------------------------------------------------------------


@dataclass
class _DatasetMatrices:
    label: str
    uri: str
    mat_raw: Optional[np.ndarray]
    mat_bal: Optional[np.ndarray]


def plot_triples_core(
    clrs: Sequence[cooler.Cooler],
    uris: Sequence[str],
    region_str: str,
    chrom: str,
    labels: Sequence[str],
    clr_weight_name: str,
    raw_vmin_opt: Optional[float],
    raw_vmax_opt: Optional[float],
    bal_vmin_opt: Optional[float],
    bal_vmax_opt: Optional[float],
    raw_scale: str,
    bal_scale: str,
    cmap: str,
    fig_width: float,
    fig_height: float,
    dpi: int,
    plot_raw: bool,
    plot_bal: bool,
    q_low: float,
    q_high: float,
    verbose: bool,
    out_path: Optional[str] = None,
) -> str:
    """
    Core plotting routine for plot-triples.
    """
    if not clrs or not uris:
        raise click.ClickException("No coolers / URIs provided to plot_triples_core.")
    if len(clrs) != len(uris):
        raise click.ClickException("Number of coolers and URIs must match.")
    if labels and len(labels) != len(uris):
        raise click.ClickException("Number of labels must match number of URIs.")
    if not plot_raw and not plot_bal:
        raise click.BadParameter(
            "At least one of plot_raw or plot_bal must be True."
        )

    n_datasets = len(uris)
    labels_used = list(labels) if labels else [os.path.basename(u.split("::")[0]) for u in uris]

    # Load matrices
    datasets: List[_DatasetMatrices] = []
    for clr, uri, label in zip(clrs, uris, labels_used):
        if verbose:
            click.echo(f"[plot-triples] Loading {label!r} from {uri!r}", err=True)
        mat_raw, mat_bal = _load_dataset_matrices(
            clr=clr,
            uri=uri,
            region_str=region_str,
            chrom=chrom,
            clr_weight_name=clr_weight_name,
            want_raw=plot_raw or plot_bal,
            want_bal=plot_bal,
        )
        datasets.append(
            _DatasetMatrices(
                label=label,
                uri=uri,
                mat_raw=mat_raw,
                mat_bal=mat_bal,
            )
        )

    # Consistency checks on shapes per row
    if plot_raw:
        shapes_raw = [d.mat_raw.shape for d in datasets if d.mat_raw is not None]
        if len(set(shapes_raw)) > 1:
            raise click.ClickException(
                f"Inconsistent raw matrix shapes across datasets for region {region_str!r}: "
                f"{shapes_raw}"
            )
    if plot_bal:
        shapes_bal = [d.mat_bal.shape for d in datasets if d.mat_bal is not None]
        if len(set(shapes_bal)) > 1:
            raise click.ClickException(
                f"Inconsistent balanced matrix shapes across datasets for region {region_str!r}: "
                f"{shapes_bal}"
            )

    # Parse numeric start/end from region_str: chrom:start-end
    try:
        _chrom_part, _coords_part = region_str.split(":", 1)
        _start_str, _end_str = _coords_part.split("-", 1)
        start_bp = int(_start_str)
        end_bp = int(_end_str)
    except Exception as e:
        raise click.ClickException(
            f"Could not parse region_str {region_str!r} into 'chrom:start-end'."
        ) from e

    # Determine reference matrix size (number of bins) for genomic ticks
    if plot_raw:
        ref_mat = next((d.mat_raw for d in datasets if d.mat_raw is not None), None)
    else:
        ref_mat = next((d.mat_bal for d in datasets if d.mat_bal is not None), None)

    if ref_mat is None:
        raise click.ClickException(
            f"No matrices loaded for region {region_str!r}; nothing to plot."
        )

    n_bins = ref_mat.shape[0]

    # Compute genomic tick positions / labels once per figure
    x_tick_pos, x_tick_labels, x_unit = _compute_genomic_ticks(
        start_bp=start_bp,
        end_bp=end_bp,
        n_bins=n_bins,
        max_ticks=6,
    )
    # Reuse the same ticks for y-axis
    y_tick_pos, y_tick_labels, _y_unit = x_tick_pos, x_tick_labels, x_unit

    # Color limits and norms per row
    raw_norm: Optional[Normalize] = None
    bal_norm: Optional[Normalize] = None
    raw_vmin_eff = raw_vmax_eff = None
    bal_vmin_eff = bal_vmax_eff = None

    if plot_raw:
        raw_mats = [d.mat_raw for d in datasets if d.mat_raw is not None]
        if raw_scale == "log":
            raw_vmin_eff, raw_vmax_eff = _compute_joint_vlim_log(
                raw_mats,
                vmin_opt=raw_vmin_opt,
                vmax_opt=raw_vmax_opt,
                q_low=q_low,
                q_high=q_high,
                floor=1.0,
            )
            raw_norm = LogNorm(vmin=raw_vmin_eff, vmax=raw_vmax_eff)
        else:
            raw_vmin_eff, raw_vmax_eff = _compute_joint_vlim_linear(
                raw_mats,
                vmin_opt=raw_vmin_opt,
                vmax_opt=raw_vmax_opt,
                q_low=q_low,
                q_high=q_high,
            )
            raw_norm = Normalize(vmin=raw_vmin_eff, vmax=raw_vmax_eff)

        if verbose:
            click.echo(
                f"[plot-triples] Raw scale={raw_scale}, vmin={raw_vmin_eff}, vmax={raw_vmax_eff}",
                err=True,
            )

    if plot_bal:
        bal_mats = [d.mat_bal for d in datasets if d.mat_bal is not None]
        if bal_scale == "log":
            bal_vmin_eff, bal_vmax_eff = _compute_joint_vlim_log(
                bal_mats,
                vmin_opt=bal_vmin_opt,
                vmax_opt=bal_vmax_opt,
                q_low=q_low,
                q_high=q_high,
                floor=None,
            )
            bal_norm = LogNorm(vmin=bal_vmin_eff, vmax=bal_vmax_eff)
        else:
            bal_vmin_eff, bal_vmax_eff = _compute_joint_vlim_linear(
                bal_mats,
                vmin_opt=bal_vmin_opt,
                vmax_opt=bal_vmax_opt,
                q_low=q_low,
                q_high=q_high,
            )
            bal_norm = Normalize(vmin=bal_vmin_eff, vmax=bal_vmax_eff)

        if verbose:
            click.echo(
                f"[plot-triples] Balanced scale={bal_scale}, vmin={bal_vmin_eff}, vmax={bal_vmax_eff}",
                err=True,
            )

    # Row configuration
    row_names: List[str] = []
    if plot_raw:
        row_names.append("raw")
    if plot_bal:
        row_names.append("balanced")

    nrows = len(row_names)
    row_index = {name: i for i, name in enumerate(row_names)}
    top_row_idx = 0
    bottom_row_idx = nrows - 1
    raw_row_idx = row_index.get("raw")
    bal_row_idx = row_index.get("balanced")

    # Create figure & axes
    fig, axs = plt.subplots(
        nrows=nrows,
        ncols=n_datasets,
        figsize=(fig_width * n_datasets, fig_height * nrows),
        gridspec_kw={"height_ratios": [1] * nrows, "wspace": 0.3},
    )

    # Normalize axes to 2D array
    if nrows == 1 and n_datasets == 1:
        axs = np.array([[axs]])
    elif nrows == 1:
        axs = axs.reshape(1, n_datasets)
    elif n_datasets == 1:
        axs = axs.reshape(nrows, 1)

    # Leave room on the right for colorbars
    plt.subplots_adjust(left=0.05, right=0.9, top=0.90, bottom=0.18, hspace=0.3)

    # Top row titles: dataset labels
    for col, ds in enumerate(datasets):
        axs[top_row_idx, col].set_title(ds.label)

    # Plot
    example_im_raw = None
    example_im_bal = None

    for col, ds in enumerate(datasets):
        # Raw row
        if plot_raw and raw_row_idx is not None:
            ax_raw = axs[raw_row_idx, col]
            if raw_scale == "log":
                im_raw = ax_raw.matshow(
                    ds.mat_raw,
                    norm=raw_norm,
                    cmap=cmap,
                    aspect="equal",
                    interpolation="nearest",  # NO smoothing
                )
            else:
                im_raw = ax_raw.matshow(
                    ds.mat_raw,
                    vmin=raw_vmin_eff,
                    vmax=raw_vmax_eff,
                    cmap=cmap,
                    aspect="equal",
                    interpolation="nearest",
                )
            if example_im_raw is None:
                example_im_raw = im_raw

            # Y-axis ticks: only on first column, as genomic coords
            if col == 0:
                ax_raw.set_ylabel("Raw counts")
                if y_tick_pos.size > 0:
                    ax_raw.set_yticks(y_tick_pos)
                    ax_raw.set_yticklabels(y_tick_labels)
            else:
                ax_raw.set_ylabel("")
                ax_raw.set_yticklabels([])

            # X-axis: visible only if raw row is the bottom row
            is_bottom_raw = (raw_row_idx == bottom_row_idx)
            ax_raw.xaxis.set_visible(is_bottom_raw)
            if is_bottom_raw and x_tick_pos.size > 0:
                ax_raw.set_xticks(x_tick_pos)
                ax_raw.set_xticklabels(x_tick_labels, rotation=45, ha="right")
            else:
                ax_raw.set_xticklabels([])

        # Balanced row
        if plot_bal and bal_row_idx is not None:
            ax_bal = axs[bal_row_idx, col]
            if bal_scale == "log":
                im_bal = ax_bal.matshow(
                    ds.mat_bal,
                    norm=bal_norm,
                    cmap=cmap,
                    aspect="equal",
                    interpolation="nearest",
                )
            else:
                im_bal = ax_bal.matshow(
                    ds.mat_bal,
                    vmin=bal_vmin_eff,
                    vmax=bal_vmax_eff,
                    cmap=cmap,
                    aspect="equal",
                    interpolation="nearest",
                )
            if example_im_bal is None:
                example_im_bal = im_bal

            # Y-axis ticks: only on first column
            if col == 0:
                ax_bal.set_ylabel("Balanced frequencies")
                if y_tick_pos.size > 0:
                    ax_bal.set_yticks(y_tick_pos)
                    ax_bal.set_yticklabels(y_tick_labels)
            else:
                ax_bal.set_ylabel("")
                ax_bal.set_yticklabels([])

            # X-axis: visible only if balanced row is the bottom row
            is_bottom_bal = (bal_row_idx == bottom_row_idx)
            ax_bal.xaxis.set_visible(is_bottom_bal)
            if is_bottom_bal and x_tick_pos.size > 0:
                ax_bal.set_xticks(x_tick_pos)
                ax_bal.set_xticklabels(x_tick_labels, rotation=45, ha="right")
            else:
                ax_bal.set_xticklabels([])

    # Shared colorbars: put each in its own axes to the right of the last column
    if plot_raw and example_im_raw is not None and raw_row_idx is not None:
        last_ax_raw = axs[raw_row_idx, -1]
        pos = last_ax_raw.get_position()
        cax_raw = fig.add_axes([pos.x1 + 0.01, pos.y0, 0.02, pos.height])
        cbar_raw = fig.colorbar(example_im_raw, cax=cax_raw)
        cbar_raw.set_label("Raw counts", rotation=270, labelpad=10)

    if plot_bal and example_im_bal is not None and bal_row_idx is not None:
        last_ax_bal = axs[bal_row_idx, -1]
        pos = last_ax_bal.get_position()
        cax_bal = fig.add_axes([pos.x1 + 0.01, pos.y0, 0.02, pos.height])
        cbar_bal = fig.colorbar(example_im_bal, cax=cax_bal)
        cbar_bal.set_label("Balanced frequencies", rotation=270, labelpad=10)

    # Global region label at bottom (include units)
    fig.text(0.5, 0.02, f"{region_str} ({x_unit})", ha="center")

    # Output path
    if out_path is None:
        timestamp = datetime.now().strftime("%Y%m%d_%H%M%S")
        out_path = os.path.join(os.getcwd(), f"triples_{chrom}_{timestamp}.pdf")

    fig.savefig(out_path, bbox_inches="tight", dpi=dpi)
    plt.close(fig)

    if verbose:
        click.echo(f"[plot-triples] Saved plot to {out_path}", err=True)

    return out_path


# ---------------------------------------------------------------------------
# Click CLI wrapper
# ---------------------------------------------------------------------------


@click.command(
    name="plot-triples",
    short_help="Plot raw and/or balanced matrices for multiple coolers as a 1×N or 2×N grid.",
)
@click.argument(
    "cool_paths",
    nargs=-1,
    metavar="COOL_PATH ...",
)
@click.option(
    "-r",
    "--region",
    required=True,
    metavar="REGION",
    help=(
        "Genomic region to plot, e.g. 'chr19:0-58617616' "
        "or 'chr19:30,000,000-35,000,000'."
    ),
)
@click.option(
    "--res",
    type=int,
    default=None,
    metavar="BP",
    help=(
        "Resolution (bin size, in bp) to use when COOL_PATH points to a .mcool "
        "file without an explicit '::/resolutions/...' group. "
        "If omitted, the highest resolution (smallest bin size) in the .mcool "
        "file is chosen automatically. Ignored if COOL_PATH already includes '::'."
    ),
)
@click.option(
    "-l",
    "--label",
    "labels",
    multiple=True,
    help=(
        "Label for each COOL_PATH (repeat option for each dataset). "
        "If omitted, labels are derived from COOL_PATH basenames."
    ),
)
@click.option(
    "--clr-weight-name",
    "--clr_weight_name",
    "--weight-name",
    "--weight",
    "clr_weight_name",
    default="weight",
    show_default=True,
    metavar="NAME",
    help=(
        "Name of balancing weight column in the cooler bins table "
        "used for the balanced row. Aliases: --weight-name, --weight."
    ),
)
@click.option(
    "--raw-vmin",
    type=float,
    default=None,
    metavar="V",
    help=(
        "Lower bound for color scale of raw matrices. "
        "If omitted, chosen automatically from all datasets jointly."
    ),
)
@click.option(
    "--raw-vmax",
    type=float,
    default=None,
    metavar="V",
    help=(
        "Upper bound for color scale of raw matrices. "
        "If omitted, chosen automatically from all datasets jointly."
    ),
)
@click.option(
    "--bal-vmin",
    type=float,
    default=None,
    metavar="V",
    help=(
        "Lower bound for color scale of balanced matrices. "
        "If omitted, chosen automatically from all datasets jointly."
    ),
)
@click.option(
    "--bal-vmax",
    type=float,
    default=None,
    metavar="V",
    help=(
        "Upper bound for color scale of balanced matrices. "
        "If omitted, chosen automatically from all datasets jointly."
    ),
)
@click.option(
    "--raw-scale",
    type=click.Choice(["log", "linear"], case_sensitive=False),
    default="log",
    show_default=True,
    help="Scaling for raw matrices: 'log' = LogNorm, 'linear' = linear color scale.",
)
@click.option(
    "--balanced-scale",
    "bal_scale",
    type=click.Choice(["log", "linear"], case_sensitive=False),
    default="log",
    show_default=True,
    help="Scaling for balanced matrices: 'log' = LogNorm, 'linear' = linear color scale.",
)
@click.option(
    "--q-low",
    "q_low",
    type=float,
    default=1.0,
    show_default=True,
    metavar="PCT",
    help="Lower percentile for automatic vmin (when not provided).",
)
@click.option(
    "--q-high",
    "q_high",
    type=float,
    default=99.5,
    show_default=True,
    metavar="PCT",
    help="Upper percentile for automatic vmax (when not provided).",
)
@click.option(
    "--cmap",
    type=str,
    default="fall",
    show_default=True,
    help="Matplotlib colormap name to use for all panels.",
)
@click.option(
    "--fig-width",
    type=float,
    default=6.0,
    show_default=True,
    help="Width (in inches) of each column in the grid.",
)
@click.option(
    "--fig-height",
    type=float,
    default=6.0,
    show_default=True,
    help="Height (in inches) of each row in the grid.",
)
@click.option(
    "--dpi",
    type=int,
    default=150,
    show_default=True,
    help="DPI (resolution) for the output figure.",
)
@click.option(
    "--no-raw",
    is_flag=True,
    default=False,
    help="Do not plot the raw (unbalanced) row.",
)
@click.option(
    "--no-balanced",
    is_flag=True,
    default=False,
    help="Do not plot the balanced (weighted) row.",
)
@click.option(
    "-v",
    "--verbose",
    is_flag=True,
    default=False,
    help="Print resolved URIs, chosen scales, and other info to stderr.",
)
@click.option(
    "-o",
    "--out",
    "out_path",
    type=str,
    default=None,
    help=(
        "Output figure path (PDF/PNG/etc). "
        "If omitted, a timestamped PDF is written in the current directory."
    ),
)
def plot_triples(
    cool_paths: Sequence[str],
    region: str,
    res: Optional[int],
    labels: Sequence[str],
    clr_weight_name: str,
    raw_vmin: Optional[float],
    raw_vmax: Optional[float],
    bal_vmin: Optional[float],
    bal_vmax: Optional[float],
    raw_scale: str,
    bal_scale: str,
    q_low: float,
    q_high: float,
    cmap: str,
    fig_width: float,
    fig_height: float,
    dpi: int,
    no_raw: bool,
    no_balanced: bool,
    verbose: bool,
    out_path: Optional[str],
) -> None:
    """
    Click entry point for the 'plot-triples' command.
    """
    if not cool_paths:
        raise click.UsageError("Provide at least one COOL_PATH.")

    plot_raw = not no_raw
    plot_bal = not no_balanced
    if not plot_raw and not plot_bal:
        raise click.BadParameter(
            "Cannot disable both raw and balanced rows. "
            "Remove either --no-raw or --no-balanced."
        )

    if not (0.0 <= q_low < q_high <= 100.0):
        raise click.BadParameter(
            f"Invalid quantile range q_low={q_low}, q_high={q_high}; "
            "must satisfy 0 <= q_low < q_high <= 100."
        )

    n_datasets = len(cool_paths)
    if labels and len(labels) != n_datasets:
        raise click.BadParameter(
            f"Got {len(labels)} labels but {n_datasets} COOL_PATH arguments. "
            "Provide either no labels or exactly one per COOL_PATH."
        )

    raw_scale = raw_scale.lower()
    bal_scale = bal_scale.lower()

    # Resolve COOL_PATHs into URIs and open coolers
    uris: List[str] = []
    clrs: List[cooler.Cooler] = []
    for p in cool_paths:
        uri = _resolve_cooler_uri(p, res)
        uris.append(uri)
        try:
            clr = cooler.Cooler(uri)
        except Exception as e:
            raise click.ClickException(f"Could not open COOL_PATH {uri!r}: {e}") from e
        clrs.append(clr)
        if verbose:
            click.echo(f"[plot-triples] COOL_PATH {p!r} -> {uri!r}", err=True)

    # Normalize region vs first cooler
    region_str, chrom, start, end = _normalize_region(region, clrs[0])
    if verbose:
        click.echo(
            f"[plot-triples] region={region_str}, chrom={chrom}, "
            f"start={start}, end={end}",
            err=True,
        )

    # Derive labels if not provided
    if labels:
        labels_used = list(labels)
    else:
        labels_used = [os.path.basename(u.split("::")[0]) for u in uris]

    # Call core routine
    final_out = plot_triples_core(
        clrs=clrs,
        uris=uris,
        region_str=region_str,
        chrom=chrom,
        labels=labels_used,
        clr_weight_name=clr_weight_name,
        raw_vmin_opt=raw_vmin,
        raw_vmax_opt=raw_vmax,
        bal_vmin_opt=bal_vmin,
        bal_vmax_opt=bal_vmax,
        raw_scale=raw_scale,
        bal_scale=bal_scale,
        cmap=cmap,
        fig_width=fig_width,
        fig_height=fig_height,
        dpi=dpi,
        plot_raw=plot_raw,
        plot_bal=plot_bal,
        q_low=q_low,
        q_high=q_high,
        verbose=verbose,
        out_path=out_path,
    )

    click.echo(f"Saved plot to {final_out}")


if __name__ == "__main__":
    plot_triples()
```
