## Supplementary material for "CAD-C: An engineered nuclease enables repair-free *in situ* proximity ligation and nucleosome-resolution chromosome walks in human cells": Custom code: RPE1_darrow_Micro_CAD_v3_600dpi.sh

```
#!/bin/bash
#SBATCH --job-name=plot_rpe1_final
#SBATCH --output=plot_rpe1_final_%j.log
#SBATCH --error=plot_rpe1_final_%j.err
#SBATCH --cpus-per-task=4

set -e  # Exit if any plot fails

# ==============================================================================
# USER INPUTS
# ==============================================================================

# 1. Path to the v3 python script (Nearest Neighbor + Smart Axes)
SCRIPT_PATH="/rugpfs/fs0/risc_lab/scratch/jsoroczyns/CAD/CAD-C/scripts/2025_11_26/replot_raw_balanced_N_matrices_waxes_nointerpol_v3_20251128.py"

# 2. Base Output Directory (Updated to 11_26)
BASE_OUT_DIR="/rugpfs/fs0/risc_lab/scratch/jsoroczyns/CAD/CAD-C/scripts/2025_11_26"

# 3. Your .mcool files (Left, Middle, Right)
MCOOL_1="/rugpfs/fs0/risc_lab/scratch/jsoroczyns/CAD/CAD-C/Public_Datasets/Darrow_2016/20241125_merged_pairs/20241127_dedup_downsample_7222_igdiag2_coolers/Darrow2016_20241125_dedup_merged_downsample_7222_500_bal_500N_igdiag2.mcool"
MCOOL_2="/rugpfs/fs0/risc_lab/scratch/jsoroczyns/CAD/CAD-C/AO_MicroC_WT12and34_merge/20241127_dedup_downsample_5842_igdiag1_coolers/AO_MicroC_WT12and34_20241126_dedup_merge_downsample_5842_500_bal_500N_igdiag1.mcool"
MCOOL_3="/rugpfs/fs0/risc_lab/scratch/jsoroczyns/CAD/CAD-C/CADC_E1_P2andP3_merge/20241125_post_merge_rededup/20241127_rededup_igdiag1_coolers/CADC_E1_P2andP3_20241126_dedup_merged_rededup_500_bal_500N_igdiag1.mcool"

# 4. Labels
LABELS="--label Hi-C_Darrow_2016 --label Micro-C --label CAD-C"

# 5. Environment
export OMP_NUM_THREADS="${SLURM_CPUS_PER_TASK:-1}"

# ==============================================================================
# SETUP TIMESTAMPED DIRECTORY
# ==============================================================================

TIMESTAMP=$(date +"%Y%m%d_%H%M%S")
FINAL_OUT_DIR="${BASE_OUT_DIR}/RPE1_Triples_Final_600DPI_${TIMESTAMP}"
mkdir -p "$FINAL_OUT_DIR"

echo "Starting job..."
echo "Script: $SCRIPT_PATH"
echo "Saving outputs to: $FINAL_OUT_DIR"
echo "DPI: 600"

# ==============================================================================
# PLOTTING COMMANDS
# ==============================================================================

# 1. chr8 Zoom (1kb) | vmin 0.0005 | vmax 0.35
echo "Plotting 1..."
python "$SCRIPT_PATH" "$MCOOL_1" "$MCOOL_2" "$MCOOL_3" $LABELS \
    --region "chr8:102,594,246-102,889,522" \
    --res 1000 \
    --bal-vmin 0.0005 --bal-vmax 0.35 \
    --no-raw \
    --dpi 600 \
    --out "${FINAL_OUT_DIR}/01_chr8_zoom_1kb.pdf"

# 2. chr8 Wide (50kb) | vmin 0.00004 | vmax 2.0
echo "Plotting 2..."
python "$SCRIPT_PATH" "$MCOOL_1" "$MCOOL_2" "$MCOOL_3" $LABELS \
    --region "chr8:21,293,200-36,154,099" \
    --res 50000 \
    --bal-vmin 0.00004 --bal-vmax 2.0 \
    --no-raw \
    --dpi 600 \
    --out "${FINAL_OUT_DIR}/02_chr8_wide_50kb.pdf"

# 3. chr9 (2.5kb) | vmin 0.0002 | vmax 0.8
echo "Plotting 3..."
python "$SCRIPT_PATH" "$MCOOL_1" "$MCOOL_2" "$MCOOL_3" $LABELS \
    --region "chr9:129,042,198-129,780,074" \
    --res 2500 \
    --bal-vmin 0.0002 --bal-vmax 0.8 \
    --no-raw \
    --dpi 600 \
    --out "${FINAL_OUT_DIR}/03_chr9_2.5kb.pdf"

# 4. chr10 Top (5kb) | vmin 0.0002 | vmax 2.0
echo "Plotting 4..."
python "$SCRIPT_PATH" "$MCOOL_1" "$MCOOL_2" "$MCOOL_3" $LABELS \
    --region "chr10:3,110,091-5,087,278" \
    --res 5000 \
    --bal-vmin 0.0002 --bal-vmax 2.0 \
    --no-raw \
    --dpi 600 \
    --out "${FINAL_OUT_DIR}/04_chr10_top_5kb.pdf"

# 5. chr10 Bottom (2.5kb) | vmin 0.0003 | vmax 1.5
echo "Plotting 5..."
python "$SCRIPT_PATH" "$MCOOL_1" "$MCOOL_2" "$MCOOL_3" $LABELS \
    --region "chr10:88,496,048-89,665,189" \
    --res 2500 \
    --bal-vmin 0.0003 --bal-vmax 1.5 \
    --no-raw \
    --dpi 600 \
    --out "${FINAL_OUT_DIR}/05_chr10_bot_2.5kb.pdf"

echo "Done."
```
