## Supplementary material for "CAD-C: An engineered nuclease enables repair-free *in situ* proximity ligation and nucleosome-resolution chromosome walks in human cells": Custom code: v3_600dpi_OSCAR_GMdeepmerge.sh

```
#!/bin/bash
#SBATCH --job-name=plot_gm_oscar
#SBATCH --output=plot_gm_oscar_%j.log
#SBATCH --error=plot_gm_oscar_%j.err
#SBATCH --cpus-per-task=4

set -e

# ==============================================================================
# USER INPUTS
# ==============================================================================

# 1. Script Path (v3: No Interpolation + Smart Axes)
SCRIPT_PATH="/rugpfs/fs0/risc_lab/scratch/jsoroczyns/CAD/CAD-C/scripts/2025_11_26/replot_raw_balanced_N_matrices_waxes_nointerpol_v3_20251128.py"

# 2. Output Directory
BASE_OUT_DIR="/rugpfs/fs0/risc_lab/scratch/jsoroczyns/CAD/CAD-C/scripts/2025_11_26"
TIMESTAMP=$(date +"%Y%m%d_%H%M%S")
OUT_DIR="${BASE_OUT_DIR}/GM12878_OSCAR_Zoom_${TIMESTAMP}"

# 3. Input File (GM12878 Deep Merge)
MCOOL="/rugpfs/fs0/risc_lab/scratch/jsoroczyns/CAD/CAD-C/CADC_GM12878_deepmerge_20250205/cooler_out_20250205_023558/GM12878_deepmerge_20250205_011600_dedup_500_bal_500N_igdiag1.mcool"

# 4. Region Definition (hg38)
# OSCAR Gene (~35kb window)
REGION="chr19:54,093,528-54,128,248"

# 5. Environment
export OMP_NUM_THREADS="${SLURM_CPUS_PER_TASK:-1}"

# ==============================================================================
# PLOTTING COMMANDS
# ==============================================================================

mkdir -p "$OUT_DIR"
echo "Starting job..."
echo "Output directory: $OUT_DIR"
echo "DPI: 600"

# --- Plot: 500bp Resolution ---
# ~70 pixels wide (Very zoomed in)
echo "Plotting OSCAR region..."
python "$SCRIPT_PATH" "$MCOOL" \
    --label "CAD-C GM12878" \
    --region "$REGION" \
    --res 500 \
    --bal-vmax 0.03 \
    --no-raw \
    --dpi 600 \
    --out "$OUT_DIR/GM12878_OSCAR_500bp_vmax0.03.pdf"

echo "Done."
```
