## Supplementary material for "CAD-C: An engineered nuclease enables repair-free *in situ* proximity ligation and nucleosome-resolution chromosome walks in human cells": Custom code: v3_600dpi_PPIF_GMdeepmerge.sh

```
#!/bin/bash
#SBATCH --job-name=plot_gm_ppif
#SBATCH --output=plot_gm_ppif_%j.log
#SBATCH --error=plot_gm_ppif_%j.err
#SBATCH --cpus-per-task=4

set -e

# ==============================================================================
# USER INPUTS
# ==============================================================================

# 1. Script Path (v3: No Interpolation + Smart Axes)
SCRIPT_PATH="/rugpfs/fs0/risc_lab/scratch/jsoroczyns/CAD/CAD-C/scripts/2025_11_26/replot_raw_balanced_N_matrices_waxes_nointerpol_v3_20251128.py"

# 2. Output Directory (Updated to 11_26)
BASE_OUT_DIR="/rugpfs/fs0/risc_lab/scratch/jsoroczyns/CAD/CAD-C/scripts/2025_11_26"
TIMESTAMP=$(date +"%Y%m%d_%H%M%S")
OUT_DIR="${BASE_OUT_DIR}/GM12878_PPIF_Final_600DPI_${TIMESTAMP}"

# 3. Input File (GM12878 Deep Merge)
MCOOL="/rugpfs/fs0/risc_lab/scratch/jsoroczyns/CAD/CAD-C/CADC_GM12878_deepmerge_20250205/cooler_out_20250205_023558/GM12878_deepmerge_20250205_011600_dedup_500_bal_500N_igdiag1.mcool"

# 4. Region Definition (hg38)
# Width: ~128 kb
REGION="chr10:79,230,052-79,358,098"

# 5. Environment
export OMP_NUM_THREADS="${SLURM_CPUS_PER_TASK:-1}"

# ==============================================================================
# PLOTTING COMMANDS
# ==============================================================================

mkdir -p "$OUT_DIR"
echo "Starting job..."
echo "Output directory: $OUT_DIR"
echo "DPI: 600"

# --- Plot 1: 1kb Resolution ---
# Vmax lowered: 0.02 -> 0.009
echo "Plotting 1kb resolution..."
python "$SCRIPT_PATH" "$MCOOL" \
    --label "CAD-C GM12878" \
    --region "$REGION" \
    --res 1000 \
    --bal-vmax 0.009 \
    --dpi 600 \
    --out "$OUT_DIR/GM12878_PPIF_1kb_vmax0.009.pdf"

# --- Plot 2: 500bp Resolution ---
# Vmax lowered: 0.02 -> 0.009
echo "Plotting 500bp resolution..."
python "$SCRIPT_PATH" "$MCOOL" \
    --label "CAD-C GM12878" \
    --region "$REGION" \
    --res 500 \
    --bal-vmax 0.009 \
    --dpi 600 \
    --out "$OUT_DIR/GM12878_PPIF_500bp_vmax0.009.pdf"

echo "Done."
```
