## Supplementary material for "CAD-C: An engineered nuclease enables repair-free *in situ* proximity ligation and nucleosome-resolution chromosome walks in human cells": Custom code: bwamem2_pacbio_cadwalk_20250408.sh

```
#!/bin/bash

#SBATCH --job-name=parallel_bwamem2_pacbio
#SBATCH --output=bwamem2_parallel_%A_%a.out
#SBATCH --ntasks=1
#SBATCH --cpus-per-task=8
#SBATCH --partition=risc
#SBATCH --array=0-1

# Load environment (uncomment and modify if needed)
# source /path/to/conda.sh
# conda activate bwa-mem2 || { echo "Failed to activate bwa-mem2 environment"; exit 1; }

# Use the provided absolute paths
INPUT_DIR="/rugpfs/fs0/risc_lab/store/jsoroczyns/CAD/CADwalk/CADWALK_TR_20240404/demux/"
OUTPUT_DIR="/rugpfs/fs0/risc_lab/scratch/jsoroczyns/CAD/CADwalk/CADwalk_E1_GM12878/bwamem2_out_20250408/"
REF_GENOME="/rugpfs/fs0/risc_lab/scratch/jsoroczyns/Genome/hg38/bwa_mem_2/hg38_genome.fa"

# Define sample names corresponding to the barcodes in the demux directory
SAMPLES=("bc1001--bc1001" "bc1002--bc1002")

# Get sample name based on SLURM_ARRAY_TASK_ID
SAMPLE_NAME=${SAMPLES[$SLURM_ARRAY_TASK_ID]}

# Construct the FASTQ file path assuming the naming pattern:
# m64330e_240403_201627.<sample_name>.hifi_reads.fastq.gz
FASTQ_FILE="${INPUT_DIR}m64330e_240403_201627.${SAMPLE_NAME}.hifi_reads.fastq.gz"
OUTPUT_BAM="${OUTPUT_DIR}${SAMPLE_NAME}_bwamem2.bam"

# Debugging output
echo "DEBUG: SLURM_ARRAY_TASK_ID=${SLURM_ARRAY_TASK_ID}"
echo "DEBUG: SAMPLE_NAME=${SAMPLE_NAME}"
echo "DEBUG: FASTQ_FILE=${FASTQ_FILE}"
echo "DEBUG: OUTPUT_BAM=${OUTPUT_BAM}"
echo "DEBUG: REF_GENOME=${REF_GENOME}"

# Check if the FASTQ file exists
if [ ! -f "$FASTQ_FILE" ]; then
  echo "ERROR: FASTQ file not found: $FASTQ_FILE"
  exit 1
fi

# Create the output directory if it doesn't exist
if [ ! -d "$OUTPUT_DIR" ]; then
  echo "Output directory not found, creating: $OUTPUT_DIR"
  mkdir -p "$OUTPUT_DIR" || { echo "ERROR: Cannot create output directory: $OUTPUT_DIR"; exit 1; }
fi

# Perform alignment for PacBio HiFi (single-end)
echo "Starting alignment for ${SAMPLE_NAME}..."
bwa-mem2 mem -SP5M -t ${SLURM_CPUS_PER_TASK} "$REF_GENOME" "$FASTQ_FILE" | \
samtools view -@ ${SLURM_CPUS_PER_TASK} -b -o "$OUTPUT_BAM" - || { echo "Alignment failed for ${SAMPLE_NAME}"; exit 1; }

# Sort and index BAM file
SORTED_BAM="${OUTPUT_BAM%.bam}_sorted.bam"
echo "Sorting BAM file for ${SAMPLE_NAME}..."
samtools sort -@ ${SLURM_CPUS_PER_TASK} -o "$SORTED_BAM" "$OUTPUT_BAM" || { echo "Sorting failed for ${SAMPLE_NAME}"; exit 1; }

echo "Indexing sorted BAM file for ${SAMPLE_NAME}..."
samtools index "$SORTED_BAM" || { echo "Indexing failed for ${SAMPLE_NAME}"; exit 1; }

echo "Alignment, sorting, and indexing complete for ${SAMPLE_NAME}. Final BAM: ${SORTED_BAM}"
```
