## Supplementary material for "CAD-C: An engineered nuclease enables repair-free *in situ* proximity ligation and nucleosome-resolution chromosome walks in human cells": Custom code: cadwalk_pack_v9_20251020.py

```
#!/usr/bin/env python3
# -*- coding: utf-8 -*-
"""
CADwalk UU-run Packer (v9)
==========================

One model. One invariant. Self-certifying NPZ.

Purpose
-------
Stream pairtools *parse2* `.pairs.gz`, segment **maximal uninterrupted UU runs** per read,
and produce a compact **NPZ ("cadpack")** with runs, steps, fragments, and metadata for
downstream CADwalk analyses. v9 uses a **single orientation model**:

  **M‑terminal midpoint orientation (mode=3)**

  • Compute the **terminal fragment midpoints** from pos5/pos3 on:
      - FIRST step, side1  → A_mid (fragment A)
      - LAST  step, side2  → E_mid (fragment E)

  • The orientation rule is **applicable** iff both terminal midpoints exist and A/E are
    on the **same chromosome**. When applicable, **flip iff E_mid < A_mid**, so that the
    oriented run has E_mid ≥ A_mid. Otherwise, keep read order.

  • Flip = **reverse step order only** (and the induced fragment ordering).
    We **never** swap per-step `pos1/pos2`, mutate coordinates, or negate signs.
    The oriented anchors become:
      run_start_* = FIRST.pos1, run_end_* = LAST.pos2 (after orientation).

New in v9 (additive, NPZ-only QC & plotting)
--------------------------------------------
Run-level (per run):
  • run_term_left_mid_bp, run_term_right_mid_bp — the exact terminal **midpoints** (pos5/pos3) used for the rule.
  • run_mterm_applicable — 1 iff the rule applied (both mids present & same chromosome), else 0.
  • run_term_left_end_type, run_term_right_end_type — terminal end-typing for plotting:
      1 = anchor (AJ1/EJ2) is the fragment's LEFT end on the reference,
      2 = anchor is RIGHT end, 0 = unknown.
    (In v9 we set these deterministically from the oriented **step polarity**; see below.)
  • run_term_left_junc_bp, run_term_right_junc_bp — explicit terminal junction anchors (AJ1, EJ2).
  • run_orient_decision_src — always 3 (M‑terminal).

Fragment-level (per fragment):
  • frag_ref_chr_id, frag_ref_lo, frag_ref_hi — **canonical reference bounds** for every fragment:
      - Interiors: exact from adjacent junction endpoints (cis only).
      - Terminals: exact from pos5/pos3 when available; otherwise reconstructed from (anchor, length, end_type).
      - Unknown or trans → chr_id = −1, lo = hi = −1.
  • frag_len_bp, frag_len_src — unified per-fragment length and provenance:
      1 = from pos5/pos3 endpoints,
      2 = from junction endpoints (interior),
      3 = from terminal anchor ± run-level/frag length,
      4 = from step-level `algn_ref_spanX` fallback,
      0 = unknown.

Why this matters
----------------
The NPZ now **self-certifies** the orientation invariant (when applicable) via the stored
terminal midpoints, and provides **deterministic** fragment rectangles for plotting/QC
without guessing from polarity or reaching back to `.pairs.gz`.

Segmentation (unchanged)
------------------------
A **UU run** is a maximal block of consecutive rows with:
  • same `readID`,
  • `pair_type == 'UU'`,
  • strictly consecutive `walk_pair_index` (+1).
Missing or non-numeric `walk_pair_index` is a hard boundary.

Cis-only tortuosity & Dmax (unchanged in spirit)
------------------------------------------------
We compute step-level J/M distances/signs, cis-block tortuosity (length-weighted),
and within-run spans Dmax_J/M. Dmax_M uses **fragment midpoints** computed from
the canonical fragment bounds (terminals + interiors) when available.

CLI
---
python cadwalk_pack_v9.py \
  --pairs bc1001.pairs.gz bc1002.pairs.gz \
  --labels bc1001 bc1002 \
  --outnpz OUT/cadpack_bc1001_bc1002_v9.npz \
  [--no-verify-order] [--no-assume-sorted] [--limit-runs M] [--max-rows N] [--parquet]

Defaults:
  • --assume-sorted (True): rows are grouped by read and walk indices are monotone within the read.
  • --verify-order (True): enforce nondecreasing (readID, walk_index); use --no-verify-order if global monotonicity is not guaranteed by your pipeline.
  • No reorientation mode other than M‑terminal exists in v9.

Output (NPZ keys; all 1-D unless noted)
---------------------------------------
Global:
  chrom_names : S64[]
  tr_labels   : S64[]
  tr_sources  : S256[]
  meta_json   : uint8[] (UTF‑8 JSON, options & counts)

Runs (R):
  run_tr_id : int16                 run_read_uid : uint64
  run_L     : int32                 run_reversed : uint8
  run_orient_mode : int8 (=3)       run_orient_decision_src : int8 (=3)
  run_mterm_applicable : uint8      # 1 if A/E mids present & same chrom
  run_term_left_mid_bp,  run_term_right_mid_bp  : float64  # NaN if !applicable
  run_term_left_junc_bp, run_term_right_junc_bp : int64
  run_term_left_end_type, run_term_right_end_type : int8   # 1=left,2=right,0=unknown
  run_start_chr_id,pos : int32,int64  run_end_chr_id,pos : int32,int64
  run_n_cis_steps, run_n_trans_steps : int32,int32
  run_cis_longest_block_steps : int32
  run_sum_abs_dJ, run_max_dJ : int64,int32
  run_sum_abs_dM, run_max_dM : int64,int32
  run_Brun_J, run_Brun_M : float64,float64
  run_alt_rate_J, run_alt_rate_M : float64,float64
  run_Dmax_J, run_Dmax_M : float64,float64
  run_first_walk_idx, run_last_walk_idx : int32,int32
  run_first_step_row, run_n_steps : int64,int32
  run_first_frag_row, run_n_frags : int64,int32

Steps (S):
  step_run_idx : int32      step_order : int32      step_walk_idx : int32
  chrom1_id,pos1, chrom2_id,pos2 : int32,int64,int32,int64
  step_is_cis : uint8
  step_sgnJ, step_dJ : int8,int64
  step_sgnM, step_dM : int8,float64
  step_refspan1, step_refspan2 : int32,int32  # -1 if missing

Fragments (F = R+S):
  frag_run_idx : int32      frag_index : int32       # 0..L
  left_chr_id,pos  right_chr_id,pos  : int32,int64,int32,int64
  frag_is_cis : uint8       frag_spanJ : int64
  frag_len_bp : int64       frag_len_src : uint8     # 0/1/2/3/4 as above
  frag_ref_chr_id : int32   frag_ref_lo, frag_ref_hi : int64,int64

Invariants emitted
------------------
• If run_mterm_applicable==1:
    run_term_right_mid_bp >= run_term_left_mid_bp
    run_start_chr_id == run_end_chr_id
• Anchors match steps:
    run_term_left_junc_bp  == pos1[run_first_step_row]
    run_term_right_junc_bp == pos2[run_first_step_row + run_n_steps - 1]
• run_n_frags == run_L + 1
• Interior cis fragments with both endpoints:
    frag_len_bp == abs(right_pos - left_pos)
    frag_ref_lo/hi match min/max of endpoints

License: MIT
Author : Risca Lab + GPT‑5 Pro assistant
Version: 9.0.0
"""

from __future__ import annotations
import os, sys, gzip, argparse, json, hashlib, time
from typing import Dict, List, Tuple, Optional
import numpy as np
from datetime import datetime

VERSION = "9.0.0"

# -------------------- CLI --------------------
ap = argparse.ArgumentParser(
    description="Pack uninterrupted UU runs (v9: M-terminal midpoint orientation only) from parse2 .pairs.gz into a self-certifying NPZ.",
    formatter_class=argparse.ArgumentDefaultsHelpFormatter
)
ap.add_argument("--pairs", nargs="+", required=True, help=".pairs.gz file(s), one or more.")
ap.add_argument("--labels", nargs="*", default=None, help="Labels for each --pairs (default: basenames).")
ap.add_argument("--outnpz", required=True, help="Output NPZ path.")
ap.add_argument("--max-rows", type=int, default=0, help="Process at most N non-header rows per input (0=all).")
ap.add_argument("--limit-runs", type=int, default=0, help="Emit at most M UU runs per input (0=all).")
ap.add_argument("--verify-order", dest="verify_order", action="store_true", default=True, help="Verify (readID, walk_index) nondecreasing.")
ap.add_argument("--no-verify-order", dest="verify_order", action="store_false")
ap.add_argument("--assume-sorted", dest="assume_sorted", action="store_true", default=True, help="Assume rows are grouped by read and walk indices are monotone within read.")
ap.add_argument("--no-assume-sorted", dest="assume_sorted", action="store_false")
ap.add_argument("--parquet", action="store_true", help="Also write Parquet tables (requires pyarrow).")
args = ap.parse_args()

TS = datetime.now().strftime("%Y%m%d_%H%M%S")

# -------------------- Utilities --------------------
def safe_int(x) -> Optional[int]:
    try:
        if x is None: return None
        if x == "" or x == "!": return None
        return int(x)
    except Exception:
        return None

def read_uid64(read_id: str) -> int:
    h = hashlib.blake2b(read_id.encode("utf-8"), digest_size=8)
    return int.from_bytes(h.digest(), "big", signed=False)

def midpoint_from_pos53(p5: Optional[int], p3: Optional[int]) -> Optional[float]:
    if p5 is None or p3 is None: return None
    a,b = (p5,p3) if p5 <= p3 else (p3,p5)
    return 0.5*(a+b)

def parse_header(pairs_gz: str) -> Tuple[Dict[str,int], List[str]]:
    cols: Dict[str,int] = {}
    chroms: List[str] = []
    with gzip.open(pairs_gz, "rt", errors="replace") as fh:
        for ln in fh:
            if ln.startswith("#columns:"):
                fields = ln.strip().split()[1:]
                cols = {c:i for i,c in enumerate(fields)}
            elif ln.startswith("#chromsize:"):
                parts = ln.strip().split()
                if len(parts)>=2: chroms.append(parts[1])
            elif ln and not ln.startswith("#"):
                break
    need = ["readID","chrom1","pos1","chrom2","pos2","strand1","strand2","pair_type","walk_pair_index"]
    missing = [c for c in need if c not in cols]
    if missing:
        sys.exit(f"[ERROR] Missing required columns in header {pairs_gz}: {missing}")
    return cols, chroms

def make_ranker(chrom_order: List[str]):
    if chrom_order:
        order = {c:i for i,c in enumerate(chrom_order)}
        def rank_fn(c: str): return (order.get(c, 10**9), c)
    else:
        def rank_fn(c: str): return (0, c)
    return rank_fn

# -------------------- Global tables --------------------
chrom_to_id: Dict[str,int] = {}
chrom_names: List[bytes] = []
def ch_id(c: str) -> int:
    if c in chrom_to_id: return chrom_to_id[c]
    i = len(chrom_to_id); chrom_to_id[c] = i; chrom_names.append(c.encode("utf-8")); return i

# Runs
run_tr_id=[]; run_read_uid=[]; run_L=[]
run_reversed=[]; run_orient_mode=[]; run_orient_decision_src=[]
run_mterm_applicable=[]; run_term_left_mid_bp=[]; run_term_right_mid_bp=[]
run_term_left_junc_bp=[]; run_term_right_junc_bp=[]
run_term_left_end_type=[]; run_term_right_end_type=[]
run_start_chr_id=[]; run_start_pos=[]; run_end_chr_id=[]; run_end_pos=[]
run_n_cis_steps=[]; run_n_trans_steps=[]; run_cis_longest_block_steps=[]
run_sum_abs_dJ=[]; run_max_dJ=[]; run_sum_abs_dM=[]; run_max_dM=[]
run_Brun_J=[]; run_Brun_M=[]; run_alt_rate_J=[]; run_alt_rate_M=[]
run_Dmax_J=[]; run_Dmax_M=[]
run_first_walk_idx=[]; run_last_walk_idx=[]
run_first_step_row=[]; run_n_steps=[]; run_first_frag_row=[]; run_n_frags=[]

# Steps
step_run_idx=[]; step_order=[]; step_walk_idx=[]
chrom1_id=[]; pos1=[]; chrom2_id=[]; pos2=[]
step_is_cis=[]; step_sgnJ=[]; step_dJ=[]
step_sgnM=[]; step_dM=[]; step_refspan1=[]; step_refspan2=[]

# Fragments
frag_run_idx=[]; frag_index=[]
left_chr_id=[]; left_pos=[]; right_chr_id=[]; right_pos=[]
frag_is_cis=[]; frag_spanJ=[]
frag_len_bp=[]; frag_len_src=[]
frag_ref_chr_id=[]; frag_ref_lo=[]; frag_ref_hi=[]

# TR meta
TR_SOURCES = [os.path.abspath(p) for p in args.pairs]
TR_LABELS  = args.labels if (args.labels and len(args.labels)==len(TR_SOURCES)) else [os.path.basename(p).split(".")[0] for p in TR_SOURCES]

# -------------------- Tortuosity & stats --------------------
def compute_tortuosity_and_stats_JM(step_list: List[Tuple[int,int,int,Optional[float]]]):
    """
    step_list: (chrom_id, is_cis, dJ, sJ, dM) in oriented order.
    Returns:
      (B_J, altJ, sumJ, maxJ, longestJ), (B_M, altM, sumM, maxM)
    """
    blocksJ=[]; blocksM=[]
    curC=None; curJ=[]; curM=[]
    for c, cisf, dJ, sJ, dM in step_list:
        if not cisf:
            if curC is not None:
                blocksJ.append(curJ); blocksM.append(curM)
                curC=None; curJ=[]; curM=[]
            continue
        if curC is None or c!=curC:
            if curC is not None:
                blocksJ.append(curJ); blocksM.append(curM)
            curC=c; curJ=[(dJ,sJ)]; curM=[(dM, sJ if (dM==dM) else 0)]
        else:
            curJ.append((dJ,sJ)); curM.append((dM, sJ if (dM==dM) else 0))
    if curC is not None:
        blocksJ.append(curJ); blocksM.append(curM)

    # J
    B_num=0.0; B_den=0.0; alt_cnt=0; alt_den=0; sumJ=0; maxJ=0; longestJ=0
    for b in blocksJ:
        if not b: continue
        s_signed = sum(s*d for d,s in b)
        s_total  = sum(d for d,_ in b)
        B_num += abs(s_signed); B_den += s_total
        alt_cnt += sum(1 for i in range(len(b)-1) if b[i][1]!=b[i+1][1])
        alt_den += max(0, len(b)-1)
        sumJ += s_total; maxJ = max(maxJ, max(d for d,_ in b)); longestJ = max(longestJ, len(b))
    B_J = (1 - B_num/B_den) if B_den>0 else float("nan")
    altJ = (alt_cnt/alt_den) if alt_den>0 else float("nan")

    # M
    Bm_num=0.0; Bm_den=0.0; sumM=0.0; maxM=0.0; altM_cnt=0; altM_den=0
    for b in blocksM:
        dvals=[d for d,_ in b if d==d]; svals=[s for d,s in b if d==d]
        if not dvals: continue
        Bm_num += abs(sum(s*d for s,d in zip(svals,dvals)))
        Bm_den += sum(dvals); sumM += sum(dvals); maxM = max(maxM, max(dvals))
        altM_cnt += sum(1 for i in range(len(svals)-1) if svals[i]!=svals[i+1]); altM_den += max(0, len(svals)-1)
    B_M = (1 - Bm_num/Bm_den) if Bm_den>0 else float("nan")
    altM = (altM_cnt/altM_den) if altM_den>0 else float("nan")
    return (B_J, altJ, int(sumJ), int(maxJ), int(longestJ)), (B_M, altM, int(np.rint(sumM)), int(np.rint(maxM)))

# -------------------- Orientation (v9: M-terminal only) --------------------
def orient_run_mterm_only(rows: List[List[str]], COL: Dict[str,int]) -> Tuple[List[List[str]], int, int, Optional[float], Optional[float]]:
    """
    Compute A_mid (FIRST.side1 pos5/pos3), E_mid (LAST.side2 pos5/pos3).
    If both exist and FIRST.chrom1 == LAST.chrom2, reverse iff E_mid < A_mid.
    Returns (rows_oriented, reversed_flag, applicable, A_mid_oriented, E_mid_oriented).
    """
    first, last = rows[0], rows[-1]
    A_mid = midpoint_from_pos53(safe_int(first[COL.get("pos51")]) if "pos51" in COL else None,
                                safe_int(first[COL.get("pos31")]) if "pos31" in COL else None)
    E_mid = midpoint_from_pos53(safe_int(last[COL.get("pos52")]) if "pos52" in COL else None,
                                safe_int(last[COL.get("pos32")]) if "pos32" in COL else None)
    A_chr = first[COL["chrom1"]]; E_chr = last[COL["chrom2"]]
    applicable = int((A_mid is not None) and (E_mid is not None) and (A_chr == E_chr))
    if applicable and (E_mid < A_mid):
        rows = list(reversed(rows))
        # recompute mids after orientation for storage
        first, last = rows[0], rows[-1]
        A_mid = midpoint_from_pos53(safe_int(first[COL.get("pos51")]) if "pos51" in COL else None,
                                    safe_int(first[COL.get("pos31")]) if "pos31" in COL else None)
        E_mid = midpoint_from_pos53(safe_int(last[COL.get("pos52")]) if "pos52" in COL else None,
                                    safe_int(last[COL.get("pos32")]) if "pos32" in COL else None)
        return rows, 1, applicable, A_mid, E_mid
    else:
        return rows, 0, applicable, A_mid if applicable else None, E_mid if applicable else None

# -------------------- Per-run processing --------------------
def process_run(tr_id_i: int, read_id: str, rows: List[List[str]], COL: Dict[str,int]):
    if not rows: return
    # Orientation (v9 single model)
    rows_or, reversed_flag, applicable, A_mid, E_mid = orient_run_mterm_only(rows, COL)
    L = len(rows_or)
    klass = 4 if L>=4 else L

    first = rows_or[0]; last = rows_or[-1]
    s_ch = ch_id(first[COL["chrom1"]]); s_pos = safe_int(first[COL["pos1"]]) or -1
    e_ch = ch_id(last[COL["chrom2"]]);  e_pos = safe_int(last[COL["pos2"]])  or -1

    # Optional indices
    pos51_i = COL.get("pos51"); pos31_i = COL.get("pos31")
    pos52_i = COL.get("pos52"); pos32_i = COL.get("pos32")
    rs1_i   = COL.get("algn_ref_span1"); rs2_i = COL.get("algn_ref_span2")

    # Steps
    first_step_row_idx = len(step_run_idx)
    first_walk = safe_int(first[COL["walk_pair_index"]]) or -1
    last_walk  = safe_int(last[COL["walk_pair_index"]])  or -1

    n_cis=0; n_trans=0
    step_tuples=[]
    endpoints_J: Dict[int, List[int]] = {}
    # for Dmax_M from canonical fragment mids: we will compute after fragment phase

    for i, r in enumerate(rows_or):
        c1s, c2s = r[COL["chrom1"]], r[COL["chrom2"]]
        c1 = ch_id(c1s); c2 = ch_id(c2s)
        p1_raw = safe_int(r[COL["pos1"]]); p2_raw = safe_int(r[COL["pos2"]])
        cis = (c1s==c2s) and (p1_raw is not None) and (p2_raw is not None)

        # junction metrics
        if cis:
            dJ = abs(p2_raw - p1_raw); sJ = 1 if p2_raw>p1_raw else (-1 if p2_raw<p1_raw else 0)
            n_cis += 1; endpoints_J.setdefault(c1, []).extend([p1_raw, p2_raw])
        else:
            dJ = 0; sJ = 0; n_trans += 1

        # midpoint metrics (cis only; require both sides)
        m1 = midpoint_from_pos53(
            safe_int(r[pos51_i]) if pos51_i is not None and pos51_i < len(r) else None,
            safe_int(r[pos31_i]) if pos31_i is not None and pos31_i < len(r) else None
        )
        m2 = midpoint_from_pos53(
            safe_int(r[pos52_i]) if pos52_i is not None and pos52_i < len(r) else None,
            safe_int(r[pos32_i]) if pos32_i is not None and pos32_i < len(r) else None
        )
        if cis and (m1 is not None) and (m2 is not None):
            dM = abs(m2 - m1); sM = 1 if m2>m1 else (-1 if m2<m1 else 0)
        else:
            dM = float("nan"); sM = 0

        step_run_idx.append(len(run_L)); step_order.append(i)
        step_walk_idx.append(safe_int(r[COL["walk_pair_index"]]) or -1)
        chrom1_id.append(c1); chrom2_id.append(c2)
        pos1.append(p1_raw if p1_raw is not None else -1)
        pos2.append(p2_raw if p2_raw is not None else -1)
        step_is_cis.append(1 if cis else 0)
        step_sgnJ.append(sJ); step_dJ.append(int(dJ))
        step_sgnM.append(sM if dM==dM else 0); step_dM.append(float(dM))
        # step-level ref spans
        rs1 = safe_int(r[rs1_i]) if rs1_i is not None and rs1_i < len(r) else None
        rs2 = safe_int(r[rs2_i]) if rs2_i is not None and rs2_i < len(r) else None
        step_refspan1.append(rs1 if rs1 is not None else -1)
        step_refspan2.append(rs2 if rs2 is not None else -1)

        step_tuples.append((c1, 1 if cis else 0, int(dJ), sJ, float(dM)))

    n_steps = L

    # Tortuosity & Dmax_J from endpoints
    (B_J, altJ, sumJ, maxJ, longestJ), (B_M, altM, sumM, maxM) = compute_tortuosity_and_stats_JM(step_tuples)
    Dmax_J_val = 0.0
    for _, arr in endpoints_J.items():
        if len(arr)>=2:
            span = max(arr) - min(arr)
            if span > Dmax_J_val: Dmax_J_val = float(span)

    # Fragments (canonical bounds + lengths)
    first_frag_row_idx = len(frag_run_idx)

    # Terminal A (frag_index=0) — anchor AJ1 and end-type from first step polarity
    AJ1 = safe_int(first[COL["pos1"]]) or -1
    sgn_first = step_sgnJ[first_step_row_idx] if n_steps>0 else 0
    # A end-type: if sgn>0, unligated LEFT ⇒ anchor is RIGHT end (2); if sgn<0, anchor is LEFT end (1)
    A_et = 2 if sgn_first>0 else (1 if sgn_first<0 else 0)

    # Candidate A length
    A_lo_ps=None; A_hi_ps=None; A_len=None; A_len_src=0
    if (pos51_i is not None) and (pos31_i is not None):
        p5 = safe_int(first[pos51_i]); p3 = safe_int(first[pos31_i])
        if (p5 is not None) and (p3 is not None):
            A_lo_ps, A_hi_ps = (min(p5,p3), max(p5,p3))
            A_len = A_hi_ps - A_lo_ps; A_len_src=1
    if (A_len is None) and (rs1_i is not None) and (rs1_i < len(first)):
        Ltmp = safe_int(first[rs1_i]); 
        if Ltmp and Ltmp>0: A_len = Ltmp; A_len_src = 4

    # Emit A fragment
    frag_run_idx.append(len(run_L)); frag_index.append(0)
    left_chr_id.append(ch_id(first[COL["chrom1"]]))
    left_pos.append(AJ1); right_chr_id.append(-1); right_pos.append(-1)
    frag_is_cis.append(0); frag_spanJ.append(0)
    # frag_len_bp/src
    if A_len is not None and A_len>0: frag_len_bp.append(int(A_len)); frag_len_src.append(A_len_src)
    else: frag_len_bp.append(0); frag_len_src.append(0)
    # canonical bounds for A
    if (A_lo_ps is not None) and (A_hi_ps is not None):
        frag_ref_chr_id.append(ch_id(first[COL["chrom1"]]))
        frag_ref_lo.append(int(A_lo_ps)); frag_ref_hi.append(int(A_hi_ps))
    elif (A_len is not None) and (A_len>0) and (A_et in (1,2)) and AJ1>=0:
        if A_et == 1: lo,hi = AJ1, AJ1 + A_len
        else:         lo,hi = AJ1 - A_len, AJ1
        if lo>hi: lo,hi = hi,lo
        frag_ref_chr_id.append(ch_id(first[COL["chrom1"]]))
        frag_ref_lo.append(int(lo)); frag_ref_hi.append(int(hi))
    else:
        frag_ref_chr_id.append(-1); frag_ref_lo.append(-1); frag_ref_hi.append(-1)

    # Interiors 1..L-1
    for k in range(1, L):
        left = rows_or[k-1]; right = rows_or[k]
        la_c = ch_id(left[COL["chrom2"]]);  la_p = safe_int(left[COL["pos2"]])
        rb_c = ch_id(right[COL["chrom1"]]); rb_p = safe_int(right[COL["pos1"]])

        frag_run_idx.append(len(run_L)); frag_index.append(k)
        left_chr_id.append(la_c); left_pos.append(la_p if la_p is not None else -1)
        right_chr_id.append(rb_c); right_pos.append(rb_p if rb_p is not None else -1)

        if (la_p is not None) and (rb_p is not None) and (la_c == rb_c):
            frag_is_cis.append(1)
            span = abs(rb_p - la_p); frag_spanJ.append(int(span))
            # canonical bounds from endpoints
            lo,hi = (la_p, rb_p) if la_p <= rb_p else (rb_p, la_p)
            frag_ref_chr_id.append(int(la_c)); frag_ref_lo.append(int(lo)); frag_ref_hi.append(int(hi))
            # length from endpoints
            frag_len_bp.append(int(hi-lo)); frag_len_src.append(2)
        else:
            frag_is_cis.append(0); frag_spanJ.append(0)
            frag_ref_chr_id.append(-1); frag_ref_lo.append(-1); frag_ref_hi.append(-1)
            frag_len_bp.append(0); frag_len_src.append(0)

    # Terminal E (frag_index=L)
    EJ2 = safe_int(last[COL["pos2"]]) or -1
    sgn_last = step_sgnJ[first_step_row_idx + (L-1)] if n_steps>0 else 0
    # E end-type: if sgn>0, unligated RIGHT ⇒ anchor is LEFT end (1); if sgn<0, anchor is RIGHT end (2)
    E_et = 1 if sgn_last>0 else (2 if sgn_last<0 else 0)

    E_lo_ps=None; E_hi_ps=None; E_len=None; E_len_src=0
    if (pos52_i is not None) and (pos32_i is not None):
        p5 = safe_int(last[pos52_i]); p3 = safe_int(last[pos32_i])
        if (p5 is not None) and (p3 is not None):
            E_lo_ps, E_hi_ps = (min(p5,p3), max(p5,p3))
            E_len = E_hi_ps - E_lo_ps; E_len_src=1
    if (E_len is None) and (rs2_i is not None) and (rs2_i < len(last)):
        Ltmp = safe_int(last[rs2_i])
        if Ltmp and Ltmp>0: E_len = Ltmp; E_len_src = 4

    frag_run_idx.append(len(run_L)); frag_index.append(L)
    left_chr_id.append(ch_id(last[COL["chrom2"]]))
    left_pos.append(EJ2); right_chr_id.append(-1); right_pos.append(-1)
    frag_is_cis.append(0); frag_spanJ.append(0)
    if E_len is not None and E_len>0: frag_len_bp.append(int(E_len)); frag_len_src.append(E_len_src)
    else: frag_len_bp.append(0); frag_len_src.append(0)
    if (E_lo_ps is not None) and (E_hi_ps is not None):
        frag_ref_chr_id.append(ch_id(last[COL["chrom2"]]))
        frag_ref_lo.append(int(E_lo_ps)); frag_ref_hi.append(int(E_hi_ps))
    elif (E_len is not None) and (E_len>0) and (E_et in (1,2)) and EJ2>=0:
        if E_et == 1: lo,hi = EJ2, EJ2 + E_len
        else:         lo,hi = EJ2 - E_len, EJ2
        if lo>hi: lo,hi = hi,lo
        frag_ref_chr_id.append(ch_id(last[COL["chrom2"]]))
        frag_ref_lo.append(int(lo)); frag_ref_hi.append(int(hi))
    else:
        frag_ref_chr_id.append(-1); frag_ref_lo.append(-1); frag_ref_hi.append(-1)

    n_frags = L+1

    # Dmax_M from canonical fragment mids (cis only)
    mids_by_chr: Dict[int, List[float]] = {}
    for idx in range(first_frag_row_idx, first_frag_row_idx+n_frags):
        c = frag_ref_chr_id[idx]
        lo = frag_ref_lo[idx]; hi = frag_ref_hi[idx]
        if c >= 0 and lo >= 0 and hi >= 0 and hi > lo:
            mids_by_chr.setdefault(c, []).append(0.5*(lo+hi))
    Dmax_M_val = float("nan")
    if mids_by_chr:
        Dmax_M_val = 0.0
        for c, arr in mids_by_chr.items():
            if len(arr)>=2:
                span = max(arr) - min(arr)
                if span > Dmax_M_val: Dmax_M_val = float(span)

    # Emit run arrays
    run_tr_id.append(tr_id_i); run_read_uid.append(read_uid64(read_id)); run_L.append(L)
    run_reversed.append(reversed_flag); run_orient_mode.append(3); run_orient_decision_src.append(3)
    run_mterm_applicable.append(np.uint8(applicable))
    run_term_left_mid_bp.append(float(A_mid) if (applicable and A_mid is not None) else float("nan"))
    run_term_right_mid_bp.append(float(E_mid) if (applicable and E_mid is not None) else float("nan"))
    run_term_left_junc_bp.append(int(AJ1)); run_term_right_junc_bp.append(int(EJ2))
    run_term_left_end_type.append(np.int8(A_et)); run_term_right_end_type.append(np.int8(E_et))
    run_start_chr_id.append(s_ch); run_start_pos.append(int(s_pos))
    run_end_chr_id.append(e_ch);   run_end_pos.append(int(e_pos))
    run_n_cis_steps.append(n_cis); run_n_trans_steps.append(n_trans)
    run_cis_longest_block_steps.append(int(longestJ))
    run_sum_abs_dJ.append(int(sumJ)); run_max_dJ.append(int(maxJ))
    run_sum_abs_dM.append(int(sumM)); run_max_dM.append(int(maxM))
    run_Brun_J.append(float(B_J)); run_Brun_M.append(float(B_M))
    run_alt_rate_J.append(float(altJ)); run_alt_rate_M.append(float(altM))
    run_Dmax_J.append(float(Dmax_J_val)); run_Dmax_M.append(float(Dmax_M_val) if Dmax_M_val==Dmax_M_val else float("nan"))
    run_first_walk_idx.append(first_walk); run_last_walk_idx.append(last_walk)
    run_first_step_row.append(first_step_row_idx); run_n_steps.append(n_steps)
    run_first_frag_row.append(first_frag_row_idx); run_n_frags.append(n_frags)

# -------------------- Read streaming per input --------------------
def process_input(tr_id_i: int, pairs_path: str):
    COL, chrom_order = parse_header(pairs_path)
    rank_fn = make_ranker(chrom_order)  # reserved if needed
    rows_read=0; runs_emitted=0
    cur_read=None; buf=[]
    verify=args.verify_order; last_key=None

    class _PerInputLimit(Exception): pass

    def emit_runset(run_rows: List[List[str]], rid: str):
        nonlocal runs_emitted
        if run_rows:
            process_run(tr_id_i, rid, run_rows, COL)
            runs_emitted += 1
            if args.limit_runs and runs_emitted >= args.limit_runs:
                raise _PerInputLimit()

    def flush_read():
        if not buf: return
        rb = buf if args.assume_sorted else sorted(buf, key=lambda r: (safe_int(r[COL["walk_pair_index"]]) or -10**12))
        prev=None; run=[]
        rid = rb[0][COL["readID"]]
        for r in rb:
            if r[COL["pair_type"]] == "UU":
                wi = safe_int(r[COL["walk_pair_index"]])
                if wi is None:
                    emit_runset(run, rid); run=[]; prev=None
                    emit_runset([r], rid)
                elif prev is None:
                    emit_runset(run, rid); run=[r]; prev=wi
                elif run and wi == prev + 1:
                    run.append(r); prev=wi
                else:
                    emit_runset(run, rid); run=[r]; prev=wi
            else:
                emit_runset(run, rid); run=[]; prev=None
        emit_runset(run, rid)

    try:
        with gzip.open(pairs_path, "rt", errors="replace") as fh:
            for ln in fh:
                if ln.startswith("#"): continue
                if not ln.strip(): continue
                parts = ln.rstrip("\n").split("\t")
                if len(parts) <=  max(COL.values()): continue
                rid = parts[COL["readID"]]
                wi  = safe_int(parts[COL["walk_pair_index"]])
                if verify:
                    key=(rid, wi if wi is not None else -10**12)
                    if last_key and key < last_key:
                        sys.exit(f"[ERROR] Input not ordered at {pairs_path}: saw {key} < {last_key}. Use --no-verify-order or --no-assume-sorted.")
                    last_key=key
                if cur_read is None:
                    cur_read=rid; buf=[parts]
                elif rid != cur_read:
                    flush_read(); cur_read=rid; buf=[parts]
                else:
                    buf.append(parts)
                rows_read += 1
                if args.max_rows and rows_read >= args.max_rows:
                    break
            if buf: flush_read()
    except _PerInputLimit:
        pass
    return runs_emitted

# -------------------- Drive all inputs --------------------
t0=time.time()
total_runs=0
for ti, p in enumerate(TR_SOURCES):
    total_runs += process_input(ti, p)
elapsed = time.time()-t0

# -------------------- Save NPZ --------------------
def arr(x, dtype): return np.asarray(x, dtype=dtype)

np.savez_compressed(
    args.outnpz,
    # chrom table & TR
    chrom_names = arr(chrom_names, "S64"),
    tr_labels   = arr([s.encode("utf-8") for s in TR_LABELS], "S64"),
    tr_sources  = arr([s.encode("utf-8") for s in TR_SOURCES], "S256"),

    # runs
    run_tr_id = arr(run_tr_id, np.int16),
    run_read_uid = arr(run_read_uid, np.uint64),
    run_L = arr(run_L, np.int32),
    run_reversed = arr(run_reversed, np.uint8),
    run_orient_mode = arr(run_orient_mode, np.int8),
    run_orient_decision_src = arr(run_orient_decision_src, np.int8),
    run_mterm_applicable = arr(run_mterm_applicable, np.uint8),
    run_term_left_mid_bp  = arr(run_term_left_mid_bp,  np.float64),
    run_term_right_mid_bp = arr(run_term_right_mid_bp, np.float64),
    run_term_left_junc_bp  = arr(run_term_left_junc_bp,  np.int64),
    run_term_right_junc_bp = arr(run_term_right_junc_bp, np.int64),
    run_term_left_end_type  = arr(run_term_left_end_type,  np.int8),
    run_term_right_end_type = arr(run_term_right_end_type, np.int8),
    run_start_chr_id = arr(run_start_chr_id, np.int32),
    run_start_pos    = arr(run_start_pos,   np.int64),
    run_end_chr_id   = arr(run_end_chr_id,  np.int32),
    run_end_pos      = arr(run_end_pos,     np.int64),
    run_n_cis_steps  = arr(run_n_cis_steps, np.int32),
    run_n_trans_steps= arr(run_n_trans_steps, np.int32),
    run_cis_longest_block_steps = arr(run_cis_longest_block_steps, np.int32),
    run_sum_abs_dJ = arr(run_sum_abs_dJ, np.int64),
    run_max_dJ     = arr(run_max_dJ,   np.int32),
    run_sum_abs_dM = arr(run_sum_abs_dM, np.int64),
    run_max_dM     = arr(run_max_dM,     np.int32),
    run_Brun_J     = arr(run_Brun_J,   np.float64),
    run_Brun_M     = arr(run_Brun_M,   np.float64),
    run_alt_rate_J = arr(run_alt_rate_J, np.float64),
    run_alt_rate_M = arr(run_alt_rate_M, np.float64),
    run_Dmax_J     = arr(run_Dmax_J,   np.float64),
    run_Dmax_M     = arr(run_Dmax_M,   np.float64),
    run_first_walk_idx = arr(run_first_walk_idx, np.int32),
    run_last_walk_idx  = arr(run_last_walk_idx,  np.int32),
    run_first_step_row = arr(run_first_step_row, np.int64),
    run_n_steps        = arr(run_n_steps,       np.int32),
    run_first_frag_row = arr(run_first_frag_row, np.int64),
    run_n_frags        = arr(run_n_frags,        np.int32),

    # steps
    step_run_idx = arr(step_run_idx, np.int32),
    step_order   = arr(step_order,   np.int32),
    step_walk_idx= arr(step_walk_idx,np.int32),
    chrom1_id    = arr(chrom1_id,    np.int32),
    pos1         = arr(pos1,         np.int64),
    chrom2_id    = arr(chrom2_id,    np.int32),
    pos2         = arr(pos2,         np.int64),
    step_is_cis  = arr(step_is_cis,  np.uint8),
    step_sgnJ    = arr(step_sgnJ,    np.int8),
    step_dJ      = arr(step_dJ,      np.int64),
    step_sgnM    = arr(step_sgnM,    np.int8),
    step_dM      = arr(step_dM,      np.float64),
    step_refspan1= arr(step_refspan1,np.int32),
    step_refspan2= arr(step_refspan2,np.int32),

    # fragments
    frag_run_idx = arr(frag_run_idx, np.int32),
    frag_index   = arr(frag_index,   np.int32),
    left_chr_id  = arr(left_chr_id,  np.int32),
    left_pos     = arr(left_pos,     np.int64),
    right_chr_id = arr(right_chr_id, np.int32),
    right_pos    = arr(right_pos,    np.int64),
    frag_is_cis  = arr(frag_is_cis,  np.uint8),
    frag_spanJ   = arr(frag_spanJ,   np.int64),
    frag_len_bp  = arr(frag_len_bp,  np.int64),
    frag_len_src = arr(frag_len_src, np.uint8),
    frag_ref_chr_id = arr(frag_ref_chr_id, np.int32),
    frag_ref_lo     = arr(frag_ref_lo,     np.int64),
    frag_ref_hi     = arr(frag_ref_hi,     np.int64),

    # metadata
    meta_json = np.frombuffer(json.dumps({
        "version": VERSION,
        "timestamp": TS,
        "elapsed_sec": round(elapsed,2),
        "n_tr": len(TR_LABELS),
        "inputs": [{"label": l, "path": s} for l,s in zip(TR_LABELS, TR_SOURCES)],
        "options": {
            "verify_order": args.verify_order,
            "assume_sorted": args.assume_sorted,
            "max_rows": args.max_rows,
            "limit_runs_per_input": args.limit_runs
        },
        "orientation_model": "mterminal_midpoints",
        "fields_added_v9": [
            "run_term_left_mid_bp","run_term_right_mid_bp","run_mterm_applicable",
            "run_term_left_end_type","run_term_right_end_type",
            "run_term_left_junc_bp","run_term_right_junc_bp",
            "run_orient_decision_src",
            "frag_ref_chr_id","frag_ref_lo","frag_ref_hi",
            "frag_len_bp","frag_len_src"
        ]
    }, separators=(',',':')).encode('utf-8'), dtype=np.uint8)
)

print(f"[OK] v9 cadpack written: {args.outnpz} | runs={len(run_L):,} | steps={len(step_run_idx):,} | frags={len(frag_run_idx):,}")

# -------------------- Optional Parquet --------------------
if args.parquet:
    try:
        import pyarrow as pa, pyarrow.parquet as pq
        base = os.path.splitext(args.outnpz)[0]
        runs_tab = pa.table({
            "tr_id": arr(run_tr_id, np.int16),
            "read_uid": arr(run_read_uid, np.uint64),
            "L": arr(run_L, np.int32),
            "reversed": arr(run_reversed, np.uint8),
            "orient_mode": arr(run_orient_mode, np.int8),
            "orient_src": arr(run_orient_decision_src, np.int8),
            "mterm_applicable": arr(run_mterm_applicable, np.uint8),
            "A_mid": arr(run_term_left_mid_bp, np.float64),
            "E_mid": arr(run_term_right_mid_bp, np.float64),
            "AJ1": arr(run_term_left_junc_bp, np.int64),
            "EJ2": arr(run_term_right_junc_bp, np.int64),
            "A_end_type": arr(run_term_left_end_type, np.int8),
            "E_end_type": arr(run_term_right_end_type, np.int8),
            "start_chr_id": arr(run_start_chr_id, np.int32),
            "start_pos": arr(run_start_pos, np.int64),
            "end_chr_id": arr(run_end_chr_id, np.int32),
            "end_pos": arr(run_end_pos, np.int64),
            "cis_steps": arr(run_n_cis_steps, np.int32),
            "trans_steps": arr(run_n_trans_steps, np.int32),
            "longest_cis_block": arr(run_cis_longest_block_steps, np.int32),
            "sum_abs_dJ": arr(run_sum_abs_dJ, np.int64),
            "max_dJ": arr(run_max_dJ, np.int32),
            "sum_abs_dM": arr(run_sum_abs_dM, np.int64),
            "max_dM": arr(run_max_dM, np.int32),
            "Brun_J": arr(run_Brun_J, np.float64),
            "Brun_M": arr(run_Brun_M, np.float64),
            "alt_rate_J": arr(run_alt_rate_J, np.float64),
            "alt_rate_M": arr(run_alt_rate_M, np.float64),
            "Dmax_J": arr(run_Dmax_J, np.float64),
            "Dmax_M": arr(run_Dmax_M, np.float64),
            "first_walk_idx": arr(run_first_walk_idx, np.int32),
            "last_walk_idx": arr(run_last_walk_idx, np.int32),
            "first_step_row": arr(run_first_step_row, np.int64),
            "n_steps": arr(run_n_steps, np.int32),
            "first_frag_row": arr(run_first_frag_row, np.int64),
            "n_frags": arr(run_n_frags, np.int32),
        })
        steps_tab = pa.table({
            "run_idx": arr(step_run_idx, np.int32),
            "order": arr(step_order, np.int32),
            "walk_idx": arr(step_walk_idx, np.int32),
            "chrom1_id": arr(chrom1_id, np.int32),
            "pos1": arr(pos1, np.int64),
            "chrom2_id": arr(chrom2_id, np.int32),
            "pos2": arr(pos2, np.int64),
            "is_cis": arr(step_is_cis, np.uint8),
            "sgnJ": arr(step_sgnJ, np.int8),
            "dJ": arr(step_dJ, np.int64),
            "sgnM": arr(step_sgnM, np.int8),
            "dM": arr(step_dM, np.float64),
            "refspan1": arr(step_refspan1, np.int32),
            "refspan2": arr(step_refspan2, np.int32),
        })
        frags_tab = pa.table({
            "run_idx": arr(frag_run_idx, np.int32),
            "frag_index": arr(frag_index, np.int32),
            "left_chr_id": arr(left_chr_id, np.int32),
            "left_pos": arr(left_pos, np.int64),
            "right_chr_id": arr(right_chr_id, np.int32),
            "right_pos": arr(right_pos, np.int64),
            "is_cis": arr(frag_is_cis, np.uint8),
            "spanJ": arr(frag_spanJ, np.int64),
            "len_bp": arr(frag_len_bp, np.int64),
            "len_src": arr(frag_len_src, np.uint8),
            "ref_chr_id": arr(frag_ref_chr_id, np.int32),
            "ref_lo": arr(frag_ref_lo, np.int64),
            "ref_hi": arr(frag_ref_hi, np.int64),
        })
        pq.write_table(runs_tab,  base + ".runs.parquet",  compression="zstd")
        pq.write_table(steps_tab, base + ".steps.parquet", compression="zstd")
        pq.write_table(frags_tab, base + ".frags.parquet", compression="zstd")
        print(f"[OK] Parquet written under: {base}.[runs|steps|frags].parquet")
    except Exception as e:
        print("[WARN] --parquet requested but failed:", e)
```
