## Supplementary material for "CAD-C: An engineered nuclease enables repair-free *in situ* proximity ligation and nucleosome-resolution chromosome walks in human cells": Custom code: o1p_wEBVchromsizes_pairtools_pipe_noexpand_from_parse2_20250408.sh

```
#!/usr/bin/env bash
#
# pairtools_pipeline_from_pairs.sh
#
# USAGE:
#   ./pairtools_pipeline_from_pairs.sh <existing_pairs_dir> <output_dir>
#
# DESCRIPTION:
#   Given a directory of .pairs.gz files, run pairtools flip -> sort -> dedup -> select direct-ligation.
#   Each step is submitted as a Slurm job with dependencies in a chain.
#
# DEPENDENCIES:
#   - Slurm environment with 'sbatch' commands accessible.
#   - Pairtools installed (e.g., conda environment "open2c").
#   - A valid chrom.sizes file (HG38 or custom) for flipping, etc.
#
# AUTHOR:
#   Example author, 2025-04-08

###############################################################################
# 1) Make script robust to errors
###############################################################################
set -euo pipefail

###############################################################################
# 2) Capture inputs
###############################################################################
existing_pairs_dir="${1:-}"
output_dir="${2:-}"

if [ -z "$existing_pairs_dir" ] || [ -z "$output_dir" ]; then
  echo "Usage: $0 <existing_pairs_dir> <output_dir>"
  exit 1
fi

###############################################################################
# 3) Check for .pairs.gz input files
###############################################################################
shopt -s nullglob
pairs_files=("$existing_pairs_dir"/*.pairs.gz)
shopt -u nullglob

if [ ${#pairs_files[@]} -eq 0 ]; then
  echo "ERROR: No .pairs.gz files found in: $existing_pairs_dir"
  exit 1
fi

echo "==> Found the following .pairs.gz files in $existing_pairs_dir:"
for f in "${pairs_files[@]}"; do
  echo "     - $(basename "$f")"
done

###############################################################################
# 4) Setup output directories
###############################################################################
timestamp=$(date +%Y%m%d_%H%M)
log_dir="${output_dir}/logs_${timestamp}"
results_dir="${output_dir}/results_${timestamp}"

mkdir -p "$log_dir" "$results_dir"

echo "==> Logs will go to:    $log_dir"
echo "==> Results will go to: $results_dir"

###############################################################################
# 5) Path to your chrom.sizes file (adjust as needed)
###############################################################################
CHROM_SIZES="/rugpfs/fs0/risc_lab/scratch/jsoroczyns/CAD/CAD-C/scripts/2025_04_08/hg38_wEBV.chrom.sizes"
if [ ! -f "$CHROM_SIZES" ]; then
  echo "ERROR: Chrom.sizes file not found at '$CHROM_SIZES'!"
  exit 1
fi

###############################################################################
# 6) Function to process each .pairs.gz
###############################################################################
process_replicate() {
  local input_pairs_file="$1"
  local rdir="$2"
  local ldir="$3"

  local replicate
  replicate="$(basename "$input_pairs_file" .pairs.gz)"

  # Define output files
  local flipped_file="${rdir}/${replicate}_flipped_${timestamp}.pairs.gz"
  local sorted_file="${rdir}/${replicate}_sorted_${timestamp}.pairs.gz"
  local dedup_file="${rdir}/${replicate}_dedup_${timestamp}.pairs.gz"
  local dedup_stats_file="${rdir}/${replicate}_dedup_stats_${timestamp}.tsv"
  local direct_file="${rdir}/${replicate}_direct_only_${timestamp}.pairs.gz"

  echo "-------------------------------------------------------------------"
  echo "SUBMITTING JOBS FOR REPLICATE: $replicate"
  echo "Input:  $input_pairs_file"
  echo "-------------------------------------------------------------------"

  # ---------------------------------------------------------------------------
  # 1) Flip
  #    Use sbatch --parsable to capture job ID.
  # ---------------------------------------------------------------------------
  flip_job_id=$(
    sbatch --parsable <<EOF
#!/bin/bash
#SBATCH --job-name=flip_${replicate}
#SBATCH --output=${ldir}/flip_${replicate}_%j.out
#SBATCH --error=${ldir}/flip_${replicate}_%j.err
#SBATCH --ntasks=1
#SBATCH --cpus-per-task=8
#SBATCH --partition=risc

# Make conda available
# source /ru-auth/local/home/risc_soft2/miniconda3/etc/profile.d/conda.sh
# conda activate open2c

echo "==== [\$(date)] Starting pairtools flip for ${replicate}"
pairtools flip \
  --chroms-path "${CHROM_SIZES}" \
  --output "${flipped_file}" \
  --nproc-in 8 \
  --nproc-out 8 \
  "${input_pairs_file}"

echo "==== [\$(date)] Finished pairtools flip for ${replicate}"
EOF
  )
  echo "  -> flip job ID: $flip_job_id"

  # ---------------------------------------------------------------------------
  # 2) Sort (depends on flip)
  # ---------------------------------------------------------------------------
  sort_job_id=$(
    sbatch --parsable --dependency=afterok:${flip_job_id} <<EOF
#!/bin/bash
#SBATCH --job-name=sort_${replicate}
#SBATCH --output=${ldir}/sort_${replicate}_%j.out
#SBATCH --error=${ldir}/sort_${replicate}_%j.err
#SBATCH --ntasks=1
#SBATCH --cpus-per-task=8
#SBATCH --partition=risc

source /ru-auth/local/home/risc_soft2/miniconda3/etc/profile.d/conda.sh
conda activate open2c

echo "==== [\$(date)] Starting pairtools sort for ${replicate}"
pairtools sort \
  --output "${sorted_file}" \
  --nproc-in 8 \
  --nproc-out 8 \
  "${flipped_file}"

echo "==== [\$(date)] Finished pairtools sort for ${replicate}"
EOF
  )
  echo "  -> sort job ID: $sort_job_id"

  # ---------------------------------------------------------------------------
  # 3) Dedup (depends on sort)
  # ---------------------------------------------------------------------------
  dedup_job_id=$(
    sbatch --parsable --dependency=afterok:${sort_job_id} <<EOF
#!/bin/bash
#SBATCH --job-name=dedup_${replicate}
#SBATCH --output=${ldir}/dedup_${replicate}_%j.out
#SBATCH --error=${ldir}/dedup_${replicate}_%j.err
#SBATCH --ntasks=1
#SBATCH --cpus-per-task=8
#SBATCH --partition=risc

source /ru-auth/local/home/risc_soft2/miniconda3/etc/profile.d/conda.sh
conda activate open2c

echo "==== [\$(date)] Starting pairtools dedup for ${replicate}"
pairtools dedup \
  --output "${dedup_file}" \
  --output-stats "${dedup_stats_file}" \
  --nproc-in 8 \
  --nproc-out 8 \
  "${sorted_file}"

echo "==== [\$(date)] Finished pairtools dedup for ${replicate}"
EOF
  )
  echo "  -> dedup job ID: $dedup_job_id"

  # ---------------------------------------------------------------------------
  # 4) Select direct ligation pairs (depends on dedup)
  # ---------------------------------------------------------------------------
  select_job_id=$(
    sbatch --parsable --dependency=afterok:${dedup_job_id} <<EOF
#!/bin/bash
#SBATCH --job-name=selectDirect_${replicate}
#SBATCH --output=${ldir}/select_direct_${replicate}_%j.out
#SBATCH --error=${ldir}/select_direct_${replicate}_%j.err
#SBATCH --ntasks=1
#SBATCH --cpus-per-task=8
#SBATCH --partition=risc

source /ru-auth/local/home/risc_soft2/miniconda3/etc/profile.d/conda.sh
conda activate open2c

echo "==== [\$(date)] Starting pairtools select (direct) for ${replicate}"
pairtools select \
  --output "${direct_file}" \
  --nproc-in 8 \
  --nproc-out 8 \
  '(walk_pair_type == "R1" or walk_pair_type == "R2" or walk_pair_type == "R1&2")' \
  "${dedup_file}"

echo "==== [\$(date)] Finished pairtools select (direct) for ${replicate}"
EOF
  )
  echo "  -> select job ID: $select_job_id"

  echo "==> Submit chain for replicate: $replicate completed."
}

###############################################################################
# 7) Submit jobs for each .pairs.gz
###############################################################################
for pf in "${pairs_files[@]}"; do
  process_replicate "$pf" "$results_dir" "$log_dir" &
done

wait

###############################################################################
# 8) Done
###############################################################################
echo
echo "ALL job submission steps are complete."
echo "Check 'squeue' or Slurm logs in '$log_dir' to see progress."
```
