## Supplementary material for "CAD-C: An engineered nuclease enables repair-free *in situ* proximity ligation and nucleosome-resolution chromosome walks in human cells": Custom code: pairtools_pipe_noexpand_from_parse2_20250408.sh

```
#!/bin/bash

###############################################################################
# Modified Pipeline Script to Start from Existing .pairs.gz Files
#
# Usage:
#   ./modified_pairs_pipeline.sh <existing_pairs_dir> <output_dir>
#
# Description:
#   For each .pairs.gz file in <existing_pairs_dir>, perform:
#      - flip,
#      - sort,
#      - dedup,
#      - select direct-ligation pairs only.
#
# Produces .pairs.gz outputs and stats in <output_dir>.
#
# Notes / HPC environment:
#   - Uses risc_soft2/open2c environment.
#   - CPUs per job reduced to 8.
#
# Author: Jan Soroczynski (Risca Lab, The Rockefeller University)
# Date: 2025-04-08
###############################################################################

# --- [Command-line arguments] ---
existing_pairs_dir="$1"
output_dir="$2"

if [ -z "$existing_pairs_dir" ] || [ -z "$output_dir" ]; then
    echo "Usage: $0 <existing_pairs_dir> <output_dir>"
    exit 1
fi

# --- [Check for .pairs.gz files] ---
pairs_files=("$existing_pairs_dir"/*.pairs.gz)
if [ ! -e "${pairs_files[0]}" ]; then
    echo "No .pairs.gz files found in input directory: $existing_pairs_dir"
    exit 1
fi

echo "Processing these .pairs.gz files:"
for pairs_file in "${pairs_files[@]}"; do
    echo "  - $(basename "$pairs_file")"
done

timestamp=$(date +%Y%m%d_%H%M)
log_dir="${output_dir}/logs_${timestamp}"
results_dir="${output_dir}/results_${timestamp}"

mkdir -p "$log_dir" "$results_dir"

# -----------------------------------------------------------------------------
# Function: process_replicate from existing .pairs.gz file
# -----------------------------------------------------------------------------
process_replicate() {
    local existing_pairs_file="$1"
    local results_dir="$2"
    local log_dir="$3"

    local replicate
    replicate="$(basename "$existing_pairs_file" .pairs.gz)"

    local flipped_file="${results_dir}/${replicate}_flipped_${timestamp}.pairs.gz"
    local sorted_file="${results_dir}/${replicate}_sorted_${timestamp}.pairs.gz"
    local dedup_file="${results_dir}/${replicate}_dedup_${timestamp}.pairs.gz"
    local dedup_stats_file="${results_dir}/${replicate}_dedup_stats_${timestamp}.tsv"
    local direct_file="${results_dir}/${replicate}_direct_only_${timestamp}.pairs.gz"

    # flip
    flip_job_id=$(
        sbatch <<EOF | awk '{print \$4}'
#!/bin/bash
#SBATCH --job-name=flip_${replicate}
#SBATCH --output=${log_dir}/flip_${replicate}_%j.out
#SBATCH --error=${log_dir}/flip_${replicate}_%j.err
#SBATCH --ntasks=1
#SBATCH --cpus-per-task=8

use_risc_soft2
conda activate open2c

pairtools flip \
    --chroms-path "/rugpfs/fs0/risc_lab/scratch/jsoroczyns/CAD/CAD-C/CADC_E2_P1_Novogene/20240320_pt_flip_out/hg38.chrom.sizes" \
    --output "${flipped_file}" \
    --nproc-in 8 \
    --nproc-out 8 \
    "${existing_pairs_file}"
EOF
    )

    # sort
    sort_job_id=$(
        sbatch --dependency=afterok:"$flip_job_id" <<EOF | awk '{print \$4}'
#!/bin/bash
#SBATCH --job-name=sort_${replicate}
#SBATCH --output=${log_dir}/sort_${replicate}_%j.out
#SBATCH --error=${log_dir}/sort_${replicate}_%j.err
#SBATCH --ntasks=1
#SBATCH --cpus-per-task=8

use_risc_soft2
conda activate open2c

pairtools sort \
    --output "${sorted_file}" \
    --nproc-in 8 \
    --nproc-out 8 \
    "${flipped_file}"
EOF
    )

    # dedup
    dedup_job_id=$(
        sbatch --dependency=afterok:"$sort_job_id" <<EOF | awk '{print \$4}'
#!/bin/bash
#SBATCH --job-name=dedup_${replicate}
#SBATCH --output=${log_dir}/dedup_${replicate}_%j.out
#SBATCH --error=${log_dir}/dedup_${replicate}_%j.err
#SBATCH --ntasks=1
#SBATCH --cpus-per-task=8

use_risc_soft2
conda activate open2c

pairtools dedup \
    --output "${dedup_file}" \
    --output-stats "${dedup_stats_file}" \
    --nproc-in 8 \
    --nproc-out 8 \
    "${sorted_file}"
EOF
    )

    # select direct-ligation junctions
    sbatch --dependency=afterok:"$dedup_job_id" <<EOF
#!/bin/bash
#SBATCH --job-name=select_direct_${replicate}
#SBATCH --output=${log_dir}/select_direct_${replicate}_%j.out
#SBATCH --error=${log_dir}/select_direct_${replicate}_%j.err
#SBATCH --ntasks=1
#SBATCH --cpus-per-task=8

use_risc_soft2
conda activate open2c

pairtools select \
    --output "${direct_file}" \
    --nproc-in 8 \
    --nproc-out 8 \
    '(walk_pair_type == "R1" or walk_pair_type == "R2" or walk_pair_type == "R1&2")' \
    "${dedup_file}"
EOF
}

# -----------------------------------------------------------------------------
# Main Loop: process each existing .pairs.gz file
# -----------------------------------------------------------------------------
for pairs_file in "${pairs_files[@]}"; do
    process_replicate "$pairs_file" "$results_dir" "$log_dir" &
done

wait
echo "All jobs submitted. Check Slurm job status and logs for progress."
```
