## Supplementary material for "CAD-C: An engineered nuclease enables repair-free *in situ* proximity ligation and nucleosome-resolution chromosome walks in human cells": Custom code: pt_stats_20250408.sh

```
#!/usr/bin/env bash

###############################################################################
# run_pairtools_stats_direct.sh
#
# PURPOSE:
#   For each *_direct_only_*.pairs.gz in a specified directory,
#   run pairtools stats and produce a corresponding *_stats.tsv file.
#
# USAGE:
#   sbatch -p risc run_pairtools_stats_direct.sh /path/to/results_dir
#
# NOTES:
#   - Adjust #SBATCH directives (partition, CPUs, memory) as needed.
#   - Adjust conda environment loading if your HPC is different.
#   - Change CHROM_SIZES to your actual path if you want coverage stats by chromosome.
###############################################################################

#SBATCH --job-name=pt_stats_direct
#SBATCH --output=pt_stats_direct_%j.out
#SBATCH --error=pt_stats_direct_%j.err
#SBATCH --ntasks=1
#SBATCH --cpus-per-task=4
#SBATCH --partition=risc

# Exit on error, treat unset variables as errors, and fail on pipelines if any command fails
set -euo pipefail

# 1) Directory containing the direct-only .pairs.gz files
RESULTS_DIR="${1:-}"

if [ -z "$RESULTS_DIR" ]; then
  echo "Usage: sbatch run_pairtools_stats_direct.sh /path/to/results_dir"
  exit 1
fi

if [ ! -d "$RESULTS_DIR" ]; then
  echo "ERROR: Directory does not exist: $RESULTS_DIR"
  exit 1
fi

# 2) Chrom sizes file (OPTIONAL if you just want basic stats)
# CHROM_SIZES="/rugpfs/fs0/risc_lab/scratch/jsoroczyns/CAD/CAD-C/scripts/2025_04_08/hg38_wEBV.chrom.sizes"

# 3) Load your conda environment with pairtools
source /ru-auth/local/home/risc_soft2/miniconda3/etc/profile.d/conda.sh
conda activate open2c

echo "Running pairtools stats on direct-only files in: $RESULTS_DIR"
# echo "Using chrom sizes: $CHROM_SIZES"
echo "=========================================================="
cd "$RESULTS_DIR"

# 4) Loop over all *_direct_only_*.pairs.gz files
shopt -s nullglob
direct_only_files=(*_direct_only_*.pairs.gz)
shopt -u nullglob

if [ ${#direct_only_files[@]} -eq 0 ]; then
  echo "No *_direct_only_*.pairs.gz files found in $RESULTS_DIR!"
  exit 0
fi

for direct_file in "${direct_only_files[@]}"; do
  # Remove .pairs.gz suffix
  prefix="${direct_file%.pairs.gz}"
  out_stats="${prefix}_stats.tsv"

  echo "==> Computing stats for: $direct_file"
  echo "    Output to: $out_stats"
  
  # pairtools stats command
  pairtools stats \
  --nproc-in 4 \
  --nproc-out 4 \
  --with-chromsizes \
  -o "$out_stats" \
  "$direct_file"

  echo "Finished stats for: $direct_file"
  echo "---------------------------------------------------"
done

echo "All 'pairtools stats' commands complete!"
```
