## Supplementary material for "CAD-C: An engineered nuclease enables repair-free *in situ* proximity ligation and nucleosome-resolution chromosome walks in human cells": Custom code: sbatch_pairtools_pipe_noexpand_from_parse2_20250408.sh

```
#!/bin/bash
#SBATCH --job-name=pairtools_pipe_noexpand
#SBATCH --output=pairtools_pipe_noexpand_%j.out
#SBATCH --error=pairtools_pipe_noexpand_%j.err
#SBATCH --ntasks=1
#SBATCH --cpus-per-task=1

bash /rugpfs/fs0/risc_lab/scratch/jsoroczyns/CAD/CAD-C/scripts/2025_04_08/pairtools_pipe_noexpand_from_parse2_20250408.sh \
    /rugpfs/fs0/risc_lab/scratch/jsoroczyns/CAD/CADwalk/CADwalk_E1_GM12878/bwamem2_out_20250408/local_parse2_out \
    /rugpfs/fs0/risc_lab/scratch/jsoroczyns/CAD/CADwalk/CADwalk_E1_GM12878/bwamem2_out_20250408/pipeline_out_20250408
```
