## Supplementary material for "CAD-C: An engineered nuclease enables repair-free *in situ* proximity ligation and nucleosome-resolution chromosome walks in human cells": Custom code: submit_cadwalk_pack_v9_1_20251020.sh

```
#!/bin/bash
#SBATCH --job-name=cadpack_v9
#SBATCH --output=%x_%j.out
#SBATCH --error=%x_%j.err
#SBATCH --cpus-per-task=4

# v9 SBATCH — builds a single v9 NPZ with M-terminal orientation + canonical fragment bounds.
# Adjust paths below if your directory layout differs.

set -euo pipefail

# -------------------------
# (Optional) Python / conda
# -------------------------
# If you need a specific env, uncomment & edit:
# source /path/to/conda.sh
# conda activate open2c_v3
PYTHON="${PYTHON:-python}"

# -------------------------
# Inputs & outputs
# -------------------------
INDIR="/rugpfs/fs0/risc_lab/scratch/jsoroczyns/CAD/CADwalk/CADwalk_E1_GM12878/bwamem2_out_20250408/local_parse2_out"

P1="${INDIR}/bc1001_parse2_20250408.pairs.gz"
P2="${INDIR}/bc1002_parse2_20250408.pairs.gz"

# v9 packer script path (edit if you placed it elsewhere)
SCRIPT="/rugpfs/fs0/risc_lab/scratch/jsoroczyns/CAD/CAD-C/scripts/2025_10_20/cadwalk_pack_v9_20251020.py"

# Output NPZ
OUTDIR="${INDIR}/analyses_20251020"
mkdir -p "${OUTDIR}"
OUTNPZ="${OUTDIR}/cadpack_bc1001_bc1002_v9_${SLURM_JOB_ID}.npz"

# -------------------------
# Sanity checks
# -------------------------
if [[ ! -s "${P1}" ]]; then echo "[ERR] Missing ${P1}"; exit 2; fi
if [[ ! -s "${P2}" ]]; then echo "[ERR] Missing ${P2}"; exit 2; fi
if [[ ! -s "${SCRIPT}" ]]; then echo "[ERR] Missing packer script ${SCRIPT}"; exit 2; fi

# -------------------------
# Run v9 packer
# -------------------------
# Notes:
#  • --no-verify-order because global (readID,walk_idx) monotonicity isn’t guaranteed by many pipelines.
#  • --assume-sorted keeps speed if rows are grouped per read and walk_pair_index is monotone *within* a read.
#    If you see per-read disorder, switch to --no-assume-sorted (slower, but robust).
${PYTHON} "${SCRIPT}" \
  --pairs "${P1}" "${P2}" \
  --labels bc1001 bc1002 \
  --outnpz "${OUTNPZ}" \
  --no-verify-order \
  --assume-sorted \
  --limit-runs 0 \
  --max-rows 0

# If you want Parquet alongside the NPZ, add:  --parquet

echo "[OK] $(date) v9 NPZ → ${OUTNPZ}"
```
