## Supplementary material for "CAD-C: An engineered nuclease enables repair-free *in situ* proximity ligation and nucleosome-resolution chromosome walks in human cells": Custom code: CADpack_analyzer_v7_20251018.py

```
#!/usr/bin/env python3
# -*- coding: utf-8 -*-
"""
CADpack Analyzer v7 (2025-10-19)
=================================

Purpose
-------
Re-analyze a CADpack NPZ produced by the packer (v6/v7), recomputing per-run
tortuosity B_run and D_max, scanning for zig-zag triplets, and generating
publication-ready plots + CSV/JSON summaries. Works even if the NPZ lacks the
"midpoint" distance mode (M), in which case M is skipped with a log note.

Key analyses
------------
1) B_run & D_max (per run, recomputed from steps):
   • B_run (tortuosity, 0..1) is computed *within-cis blocks per chromosome*.
     - The block boundary rule: break when a step is non-cis or (in J-mode)
       chromosome changes; sign==0 does *not* break blocks (it just contributes
       no length or signed length).
     - Formula by block b with steps (d_i, s_i ∈ {±1}):
         B_run = 1 - ( Σ_b |Σ_i s_i d_i| ) / ( Σ_b Σ_i d_i )
   • D_max^J: for each run, take endpoints (pos1/pos2) of all cis steps on the
     same chromosome, and record the maximum span (max-min) across chromosomes.
   • D_max^M: set to NaN by design unless packer provides midpoint endpoints.

2) Step-pair hexbin (contact step i vs i+1), by B_run quartile:
   • Adjacent *cis* steps only, on log axes; ticks labeled 10^1..10^9 (masked to
     observed (or fixed) limits). Outputs both "ALL" and BQ1..BQ4.
   • Arrays saved for reproducible re-plotting.

3) Zig-zag triplets (i, i+1, i+2) classification (Pinched vs S-shaped):
   • Strict sign alternation: s_i * s_{i+1} = -1 and s_{i+1} * s_{i+2} = -1
   • Similarity filter: r = min(d_i,d_{i+1}) / max(d_i,d_{i+1}) ≥ r0 for both
   • λ = d2 / mean(d1, d3): Pinched if λ ≤ τ_pinch; S-shaped if λ ≥ τ_S; else Ambiguous
   • Only from runs with ≥ --min-runlen CIS steps.
   • CSV + JSON counts.

Distance modes
--------------
• J: junction-to-junction distances (from pos1/pos2), signs from pos deltas.
• M: midpoint-to-midpoint distances (if present in NPZ); signs from midpoint deltas.
• You can run --dist J, --dist M, or --dist both. Missing mode arrays are skipped.

Outputs (created under: <outdir>/<timestamp>/)
----------------------------------------------
logs/
  analysis.log
arrays/
  step_pairs_ALL_{J|M}.npz                 # X=d_i, Y=d_{i+1}, run_idx, qpair (quartile per pair)
  step_pairs_BQ{k}_{J|M}.npz               # per quartile
  brun_{J|M}.npy                            # per-run B_run (NaN if not computable)
  dmax_{J|M}.npy                            # per-run D_max (NaN for M by design)
summaries/
  brun_quartiles_{J|M}.json                 # edges, counts per quartile, unknown count
  zigzag_counts_{J|M}.json                  # N by class (Pinched/S/Ambiguous)
  zigzag_windows_{J|M}.csv                  # per-window details (rid, order, d1..d3, r12,r23,lambda,class)
plots/
  step_pairs_hexbin_ALL_{J|M}.pdf
  step_pairs_hexbin_BQ1_{J|M}.pdf
  step_pairs_hexbin_BQ2_{J|M}.pdf
  step_pairs_hexbin_BQ3_{J|M}.pdf
  step_pairs_hexbin_BQ4_{J|M}.pdf

CLI
---
python CADpack_analyzer_v7_20251019.py \
  --npz cadpack_bc1001_bc1002_v6_XXXX.npz \
  --outdir analyses_YYYYMMDD \
  --dist both \
  --min-runlen 3 \
  --r0 0.80 --pinch 0.60 --S 1.40 \
  --hex-gridsize 60 \
  --hex-xlim 1e1 1e9 --hex-ylim 1e1 1e9 \
  --fonttype 42

Default choices
---------------
• Font: pdf.fonttype=42 (Publisher-friendly), override with --fonttype 43.
• r0=0.80, τ_pinch=0.60, τ_S=1.40 (pinched when middle shorter; S when longer).
• If --hex-xlim/--hex-ylim are omitted, limits follow observed data.

Requirements
------------
• Python ≥ 3.8, NumPy, Matplotlib.

Notes
-----
• This analyzer recomputes B_run and D_max strictly from step arrays, so it is
  portable across CADpack versions that keep the step-level schema.
• If the NPZ lacks J or M arrays, the corresponding mode is skipped with a clear
  log note (no hard exit).

"""

import os, sys, argparse, json, time
from datetime import datetime
from typing import Tuple, Dict

import numpy as np

# --------------------------- Utilities ---------------------------

def _now() -> str:
    return datetime.now().strftime("%Y-%m-%d %H:%M:%S")

def _ensure_outdirs(outdir: str) -> Dict[str, str]:
    ts = datetime.now().strftime("%Y%m%d_%H%M%S")
    base = os.path.join(outdir, f"cadpack_analyzer_v7_{ts}")
    sub = {k: os.path.join(base, k) for k in ("logs","arrays","summaries","plots")}
    for p in [base] + list(sub.values()):
        os.makedirs(p, exist_ok=True)
    return {"base": base, **sub}

def _write_log(logdir: str, *lines: str) -> None:
    path = os.path.join(logdir, "analysis.log")
    with open(path, "a") as f:
        for ln in lines:
            f.write(ln.rstrip("\n") + "\n")

def _setup_matplotlib(fonttype: int = 42):
    try:
        import matplotlib
        matplotlib.rcParams["pdf.fonttype"] = fonttype
        matplotlib.rcParams["ps.fonttype"] = fonttype
        matplotlib.rcParams["font.size"] = 12
        matplotlib.rcParams["axes.titlesize"] = 13
        matplotlib.rcParams["axes.labelsize"] = 12
        import matplotlib.pyplot as plt
        return matplotlib, plt
    except Exception as e:
        sys.exit(f"[ERROR] Matplotlib import failed: {e}")

def _require_fields(Z, fields, tag=""):
    missing = [k for k in fields if k not in Z.files]
    if missing:
        sys.exit(f"[ERROR] NPZ missing required arrays for {tag or 'analysis'}: {missing}")

# --------------------------- Core computations ---------------------------

def _compute_brun_dmax(Z, mode="J") -> Tuple[np.ndarray, np.ndarray, np.ndarray]:
    """
    Return (run_ids, brun, dmax).
    D_max^J uses endpoints (pos1/pos2) when present; D_max^M is NaN by design.
    """
    mode = mode.upper()
    base_fields = ["step_run_idx","step_order","step_is_cis"]
    if mode == "J":
        _require_fields(Z, base_fields + ["step_dJ","step_sgnJ"], "J")
        d = Z["step_dJ"].astype(np.float64)
        s = Z["step_sgnJ"].astype(np.int8)
    else:
        _require_fields(Z, base_fields + ["step_dM","step_sgnM"], "M")
        d = Z["step_dM"].astype(np.float64)
        s = Z["step_sgnM"].astype(np.int8)

    ri    = Z["step_run_idx"].astype(np.int64)
    order = Z["step_order"].astype(np.int64)
    cis   = Z["step_is_cis"].astype(bool)

    have_endpoints = all(k in Z.files for k in ("chrom1_id","chrom2_id","pos1","pos2"))
    if have_endpoints:
        c1 = Z["chrom1_id"].astype(np.int64)
        c2 = Z["chrom2_id"].astype(np.int64)
        p1 = Z["pos1"].astype(np.int64)
        p2 = Z["pos2"].astype(np.int64)

    # sort by (run, order)
    idx = np.lexsort((order, ri))
    ri, order, cis, d, s = (arr[idx] for arr in (ri, order, cis, d, s))
    if have_endpoints:
        c1, c2, p1, p2 = (arr[idx] for arr in (c1, c2, p1, p2))

    run_ids, start = np.unique(ri, return_index=True)
    start = start.astype(np.int64)
    end = np.empty_like(start)
    end[:-1] = start[1:]
    end[-1] = ri.size

    brun = np.full(run_ids.size, np.nan, dtype=float)
    dmax = np.full(run_ids.size, np.nan, dtype=float)

    for k, (a, b) in enumerate(zip(start, end)):
        rr_cis = cis[a:b]; rr_d = d[a:b]; rr_s = s[a:b]
        finite = np.isfinite(rr_d)

        # D_max
        if mode == "J" and have_endpoints:
            rr_c1 = c1[a:b]; rr_c2 = c2[a:b]; rr_p1 = p1[a:b]; rr_p2 = p2[a:b]
            endpoints = {}
            for j in range(rr_d.size):
                if rr_cis[j] and finite[j]:
                    # cis guarantees rr_c1==rr_c2, but be defensive
                    ch = rr_c1[j] if rr_c1[j] == rr_c2[j] else rr_c1[j]
                    if rr_p1[j] >= 0: endpoints.setdefault(ch, []).append(int(rr_p1[j]))
                    if rr_p2[j] >= 0: endpoints.setdefault(ch, []).append(int(rr_p2[j]))
            Dm = 0
            for vals in endpoints.values():
                if len(vals) >= 2:
                    span = max(vals) - min(vals)
                    if span > Dm: Dm = span
            dmax[k] = float(Dm) if Dm > 0 else np.nan
        else:
            dmax[k] = np.nan  # No endpoints or M-mode

        # B_run aggregation over cis blocks by chromosome
        sum_abs_signed = 0.0
        sum_len = 0.0
        seg_len = 0.0
        seg_signed = 0.0
        prev_ch = None

        # We can detect chromosome changes only if endpoints present (J+endpoints).
        for j in range(rr_d.size):
            is_cis_fin = rr_cis[j] and finite[j]
            if is_cis_fin:
                ch = None
                if mode == "J" and have_endpoints:
                    ch = (c1[a + j] if c1[a + j] == c2[a + j] else c1[a + j])
                    if prev_ch is None or ch != prev_ch:
                        if seg_len > 0:
                            sum_abs_signed += abs(seg_signed)
                            sum_len += seg_len
                        prev_ch = ch
                        seg_len = 0.0
                        seg_signed = 0.0
                # accumulate only if sign is ±1
                if abs(rr_s[j]) == 1:
                    seg_len += rr_d[j]
                    seg_signed += rr_s[j] * rr_d[j]
            else:
                # close block on non-cis or invalid step
                if seg_len > 0:
                    sum_abs_signed += abs(seg_signed)
                    sum_len += seg_len
                    seg_len = 0.0; seg_signed = 0.0; prev_ch = None

        if seg_len > 0:
            sum_abs_signed += abs(seg_signed)
            sum_len += seg_len

        brun[k] = (1.0 - (sum_abs_signed / sum_len)) if sum_len > 0 else np.nan

    return run_ids, brun, dmax

def _brun_quartiles(run_ids: np.ndarray, brun: np.ndarray) -> Tuple[np.ndarray, np.ndarray]:
    """
    Return (edges[Q1,Q2,Q3], run2q_map) where run2q maps run_id -> {1..4} or -1 if unknown.
    NaN B_run values map to -1 (unknown).
    """
    mask = np.isfinite(brun)
    if mask.sum() < 4:
        edges = np.array([np.nan, np.nan, np.nan], dtype=float)
        if run_ids.size:
            run2q = np.full(int(run_ids.max())+1, -1, dtype=np.int8)
        else:
            run2q = np.empty(0, dtype=np.int8)
        return edges, run2q

    edges = np.quantile(brun[mask], [0.25, 0.50, 0.75])
    # ensure strict monotonicity (ties can produce equal edges)
    for k in (1, 2):
        if not np.isfinite(edges[k]) or edges[k] <= edges[k-1]:
            edges[k] = np.nextafter(edges[k-1], np.inf)

    q = np.digitize(brun, bins=edges, right=True) + 1  # 1..4
    q[~mask] = -1

    max_id = int(run_ids.max()) if run_ids.size else -1
    run2q = np.full(max_id+1, -1, dtype=np.int8)
    run2q[run_ids] = q.astype(np.int8)
    return edges, run2q

def _step_pairs_adjacent_cis(Z, mode="J"):
    """
    Build adjacent CIS step pairs (i, i+1):
      returns x=d_i, y=d_{i+1}, rpair=run_idx for the pair, order0=start order,
      and qpair placeholder to be filled by caller (run quartiles).
    """
    mode = mode.upper()
    ri    = Z["step_run_idx"].astype(np.int64)
    order = Z["step_order"].astype(np.int64)
    cis   = Z["step_is_cis"].astype(bool)
    d = Z["step_dJ" if mode=="J" else "step_dM"].astype(np.float64)

    # sort by (run, order)
    idx = np.lexsort((order, ri))
    ri, order, cis, d = (arr[idx] for arr in (ri, order, cis, d))

    n = ri.size
    if n < 2:
        return (np.empty(0), np.empty(0), np.empty(0, dtype=np.int64), np.empty(0, dtype=np.int64))

    same_run = (ri[1:] == ri[:-1])
    adj = (order[1:] == (order[:-1] + 1))
    both_cis = cis[1:] & cis[:-1]
    fin = np.isfinite(d[1:]) & np.isfinite(d[:-1])
    pos = (d[1:] > 0) & (d[:-1] > 0)

    m = same_run & adj & both_cis & fin & pos
    if not m.any():
        return (np.empty(0), np.empty(0), np.empty(0, dtype=np.int64), np.empty(0, dtype=np.int64))

    x = d[:-1][m]
    y = d[1:][m]
    rpair = ri[:-1][m]
    o0 = order[:-1][m]
    return x, y, rpair, o0

def _zigzag_windows(Z, mode="J", r0=0.80, tau_pinch=0.60, tau_S=1.40, min_runlen=3):
    """
    Return dict of zig-zag windows passing strict alternation & similarity:
      keys: run_idx, start_step_order, d1,d2,d3, r12,r23, lambda, class (1=Pinched,2=S,0=Ambig)
    """
    mode = mode.upper()
    ri    = Z["step_run_idx"].astype(np.int64)
    order = Z["step_order"].astype(np.int64)
    cis   = Z["step_is_cis"].astype(bool)

    if mode == "J":
        d = Z["step_dJ"].astype(np.float64)
        s = Z["step_sgnJ"].astype(np.int8)
    else:
        d = Z["step_dM"].astype(np.float64)
        s = Z["step_sgnM"].astype(np.int8)

    n = ri.size
    if n < 3:
        return {k: np.array([], dtype=np.float64) for k in ("d1","d2","d3","r12","r23","lambda")} | {
            "run_idx": np.array([], dtype=np.int64),
            "start_step_order": np.array([], dtype=np.int64),
            "alt12": np.array([], dtype=np.uint8),
            "alt23": np.array([], dtype=np.uint8),
            "class": np.array([], dtype=np.int8),
        }

    # masks length n-2
    same_run = (ri[:-2] == ri[1:-1]) & (ri[1:-1] == ri[2:])
    adj12 = (order[:-2] + 1 == order[1:-1])
    adj23 = (order[1:-1] + 1 == order[2:])
    both_adj = adj12 & adj23
    cis3 = cis[:-2] & cis[1:-1] & cis[2:]
    fin3 = np.isfinite(d[:-2]) & np.isfinite(d[1:-1]) & np.isfinite(d[2:])
    s_ok = (np.abs(s[:-2]) == 1) & (np.abs(s[1:-1]) == 1) & (np.abs(s[2:]) == 1)

    # min_runlen gating by CIS-step counts
    R = (int(np.max(ri)) + 1) if ri.size else 0
    if R > 0:
        cis_counts = np.bincount(ri, weights=cis.astype(np.int64), minlength=R)
        run_ok = cis_counts[ri[:-2]] >= int(min_runlen)
    else:
        run_ok = np.zeros_like(same_run, dtype=bool)

    mask0 = same_run & both_adj & cis3 & fin3 & s_ok & run_ok
    if not mask0.any():
        empty_f = {k: np.array([], dtype=np.float64) for k in ("d1","d2","d3","r12","r23","lambda")}
        empty_i = {
            "run_idx": np.array([], dtype=np.int64),
            "start_step_order": np.array([], dtype=np.int64),
            "alt12": np.array([], dtype=np.uint8),
            "alt23": np.array([], dtype=np.uint8),
            "class": np.array([], dtype=np.int8),
        }
        return {**empty_f, **empty_i}

    d1 = d[:-2][mask0]; d2 = d[1:-1][mask0]; d3 = d[2:][mask0]
    s1 = s[:-2][mask0]; s2 = s[1:-1][mask0]; s3 = s[2:][mask0]
    rid = ri[:-2][mask0]; ord0 = order[:-2][mask0]

    alt12 = (s1 * s2 == -1)
    alt23 = (s2 * s3 == -1)
    alt   = alt12 & alt23

    eps = 1e-12
    r12 = np.minimum(d1, d2) / (np.maximum(d1, d2) + eps)
    r23 = np.minimum(d2, d3) / (np.maximum(d2, d3) + eps)
    sim = (r12 >= r0) & (r23 >= r0)

    lam = d2 / (0.5 * (d1 + d3) + eps)
    cls = np.zeros(lam.size, dtype=np.int8)
    cls[lam <= tau_pinch] = 1   # Pinched
    cls[lam >= tau_S]     = 2   # S-shaped

    final = alt & sim
    return {
        "run_idx": rid[final],
        "start_step_order": ord0[final],
        "d1": d1[final], "d2": d2[final], "d3": d3[final],
        "alt12": alt12[final].astype(np.uint8),
        "alt23": alt23[final].astype(np.uint8),
        "r12": r12[final], "r23": r23[final],
        "lambda": lam[final],
        "class": cls[final]
    }

# --------------------------- Plotting ---------------------------

def _hexbin_plot(plt, ax, x, y, title, gridsize=60, xlim=None, ylim=None):
    from matplotlib.ticker import LogLocator, LogFormatterMathtext

    if x.size == 0 or y.size == 0:
        ax.set_title(f"{title}\n(no data)")
        return

    # enforce positive domain for log axes
    m = (x > 0) & (y > 0) & np.isfinite(x) & np.isfinite(y)
    x = x[m]; y = y[m]
    if x.size == 0 or y.size == 0:
        ax.set_title(f"{title}\n(no positive finite data)")
        return

    if xlim is not None:
        lo, hi = xlim
        m2 = (x >= lo) & (x <= hi)
        x = x[m2]; y = y[m2]
    if ylim is not None:
        lo, hi = ylim
        m3 = (y >= lo) & (y <= hi)
        x = x[m3]; y = y[m3]

    ax.set_xscale("log"); ax.set_yscale("log")
    hb = ax.hexbin(x, y, gridsize=gridsize, bins='log', cmap="inferno")
    cb = plt.colorbar(hb, ax=ax); cb.set_label("log10(count)")

    # ticks at powers of ten (clip to observed/fixed limits)
    ticks = [10.0 ** k for k in range(1, 10)]
    xmin = x.min(); xmax = x.max()
    ymin = y.min(); ymax = y.max()
    xt = [t for t in ticks if xmin <= t <= xmax]
    yt = [t for t in ticks if ymin <= t <= ymax]

    ax.xaxis.set_major_locator(LogLocator(base=10))
    ax.yaxis.set_major_locator(LogLocator(base=10))
    ax.xaxis.set_major_formatter(LogFormatterMathtext())
    ax.yaxis.set_major_formatter(LogFormatterMathtext())

    if xlim is not None: ax.set_xlim(xlim)
    if ylim is not None: ax.set_ylim(ylim)
    if xt: ax.set_xticks(xt)
    if yt: ax.set_yticks(yt)

    ax.set_xlabel(r"$d_i$ (bp)")
    ax.set_ylabel(r"$d_{i+1}$ (bp)")
    ax.set_title(title)

# --------------------------- Driver ---------------------------

def analyze(npz_path: str, outdir: str, dist_mode: str, min_runlen: int,
            r0: float, tau_pinch: float, tau_S: float, hex_gridsize: int,
            hex_xlim: Tuple[float,float], hex_ylim: Tuple[float,float],
            fonttype: int):

    sub = _ensure_outdirs(outdir)
    _write_log(sub["logs"], f"[{_now()}] start", f"npz={npz_path}", f"dist={dist_mode}",
               f"r0={r0} pinch={tau_pinch} S={tau_S}", f"min_runlen={min_runlen}")

    mplt, plt = _setup_matplotlib(fonttype)

    with np.load(npz_path, allow_pickle=False, mmap_mode="r") as Z:
        # Decide modes gracefully
        modes = []
        base = ["step_run_idx","step_order","step_is_cis"]
        want_j = dist_mode.lower() in ("j", "both")
        want_m = dist_mode.lower() in ("m", "both")

        if want_j:
            if all(k in Z.files for k in (base + ["step_dJ","step_sgnJ"])):
                modes.append("J")
            else:
                _write_log(sub["logs"], f"[{_now()}] Skipping J: missing step_dJ/step_sgnJ")
        if want_m:
            if all(k in Z.files for k in (base + ["step_dM","step_sgnM"])):
                modes.append("M")
            else:
                _write_log(sub["logs"], f"[{_now()}] Skipping M: missing step_dM/step_sgnM")

        if not modes:
            sys.exit("[ERROR] No valid distance modes available in NPZ for the requested --dist.")

        for mode in modes:
            # 1) B_run, D_max
            run_ids, brun, dmax = _compute_brun_dmax(Z, mode=mode)
            np.save(os.path.join(sub["arrays"], f"brun_{mode}.npy"), brun)
            np.save(os.path.join(sub["arrays"], f"dmax_{mode}.npy"), dmax)

            # 2) Quartiles
            edges, run2q = _brun_quartiles(run_ids, brun)
            counts = {"BQ1": 0, "BQ2": 0, "BQ3": 0, "BQ4": 0, "unknown": 0}
            if run2q.size:
                counts["BQ1"] = int((run2q == 1).sum())
                counts["BQ2"] = int((run2q == 2).sum())
                counts["BQ3"] = int((run2q == 3).sum())
                counts["BQ4"] = int((run2q == 4).sum())
                counts["unknown"] = int((run2q == -1).sum())
            with open(os.path.join(sub["summaries"], f"brun_quartiles_{mode}.json"), "w") as f:
                json.dump({
                    "edges": [None if not np.isfinite(x) else float(x) for x in edges],
                    "n_runs": int(run_ids.size),
                    "n_finite": int(np.isfinite(brun).sum()),
                    "counts": counts
                }, f, indent=2)

            if np.isfinite(dmax).sum() == 0:
                _write_log(sub["logs"], f"[{_now()}] {mode} D_max unavailable (M-mode or no endpoints); set to NaN.")

            # 3) Step i vs i+1 hexbin (ALL + by quartile)
            x, y, rpair, ord0 = _step_pairs_adjacent_cis(Z, mode=mode)
            np.savez_compressed(os.path.join(sub["arrays"], f"step_pairs_ALL_{mode}.npz"),
                                x=x, y=y, run_idx=rpair, order0=ord0)

            fig = plt.figure(figsize=(8.2, 6.0)); ax = fig.gca()
            _hexbin_plot(plt, ax, x, y, f"Step pairs (ALL) — {mode}", gridsize=hex_gridsize,
                         xlim=hex_xlim, ylim=hex_ylim)
            fig.tight_layout()
            fig.savefig(os.path.join(sub["plots"], f"step_pairs_hexbin_ALL_{mode}.pdf"))
            plt.close(fig)

            if run2q.size > 0 and x.size > 0 and np.isfinite(edges).all():
                qpair = run2q[rpair]
                for q in (1, 2, 3, 4):
                    m = (qpair == q)
                    fig = plt.figure(figsize=(8.2, 6.0)); ax = fig.gca()
                    _hexbin_plot(plt, ax, x[m], y[m], f"Step pairs by B_run quartile — BQ{q} ({mode})", gridsize=hex_gridsize,
                                 xlim=hex_xlim, ylim=hex_ylim)
                    fig.tight_layout()
                    fig.savefig(os.path.join(sub["plots"], f"step_pairs_hexbin_BQ{q}_{mode}.pdf"))
                    plt.close(fig)
                    np.savez_compressed(os.path.join(sub["arrays"], f"step_pairs_BQ{q}_{mode}.npz"),
                                        x=x[m], y=y[m], run_idx=rpair[m], qpair=qpair[m], order0=ord0[m])

            # 4) Zig-zag windows (triplets)
            tab = _zigzag_windows(Z, mode=mode, r0=r0, tau_pinch=tau_pinch, tau_S=tau_S, min_runlen=min_runlen)
            n_pinched = int((tab["class"] == 1).sum())
            n_s       = int((tab["class"] == 2).sum())
            n_amb     = int((tab["class"] == 0).sum())
            with open(os.path.join(sub["summaries"], f"zigzag_counts_{mode}.json"), "w") as f:
                json.dump({"pinched": n_pinched, "S": n_s, "ambiguous": n_amb}, f, indent=2)

            # write CSV (include polarity flags for transparency)
            csv_path = os.path.join(sub["summaries"], f"zigzag_windows_{mode}.csv")
            with open(csv_path, "w") as f:
                f.write("run_idx,start_order,d1_bp,d2_bp,d3_bp,r12,r23,lambda,class\n")
                for i in range(tab["run_idx"].size):
                    f.write("{},{},{:.6g},{:.6g},{:.6g},{:.4f},{:.4f},{:.4f},{}\n".format(
                        int(tab["run_idx"][i]), int(tab["start_step_order"][i]),
                        float(tab["d1"][i]), float(tab["d2"][i]), float(tab["d3"][i]),
                        float(tab["r12"][i]), float(tab["r23"][i]), float(tab["lambda"][i]),
                        int(tab["class"][i])
                    ))

            _write_log(sub["logs"], f"[{_now()}] {mode} step-pairs ALL: N={x.size}",
                       f"[{_now()}] {mode} zigzag: N_final={tab['run_idx'].size}")

    _write_log(sub["logs"], f"[{_now()}] done")
    print(f"[OK] Outputs → {sub['base']}")

# --------------------------- CLI ---------------------------

def main():
    ap = argparse.ArgumentParser(
        description="CADpack Analyzer v7 — B_run/D_max, step-pair hexbin, zig-zag classification.",
        formatter_class=argparse.ArgumentDefaultsHelpFormatter
    )
    ap.add_argument("--npz", required=True, help="CADpack NPZ (v6/v7).")
    ap.add_argument("--outdir", required=True, help="Output parent directory.")
    ap.add_argument("--dist", choices=["J","M","both"], default="both",
                    help="Distance mode(s): J (junction), M (midpoint), or both.")
    ap.add_argument("--min-runlen", type=int, default=3,
                    help="Minimum CIS steps in a run for zig-zag windows (triplets need ≥3).")
    ap.add_argument("--r0", type=float, default=0.80, help="Similarity threshold r for adjacent pairs.")
    ap.add_argument("--pinch", dest="tau_pinch", type=float, default=0.60, help="λ ≤ τ_pinch → Pinched.")
    ap.add_argument("--S", dest="tau_S", type=float, default=1.40, help="λ ≥ τ_S → S-shaped.")
    ap.add_argument("--hex-gridsize", type=int, default=60, help="Hexbin gridsize.")
    ap.add_argument("--hex-xlim", nargs=2, type=float, default=None, metavar=("XMIN","XMAX"),
                    help="Fixed x limits for hexbin (e.g., 1e1 1e9).")
    ap.add_argument("--hex-ylim", nargs=2, type=float, default=None, metavar=("YMIN","YMAX"),
                    help="Fixed y limits for hexbin (e.g., 1e1 1e9).")
    ap.add_argument("--fonttype", type=int, choices=[42,43], default=42, help="Matplotlib pdf.fonttype.")
    args = ap.parse_args()

    # guard thresholds
    if not (0.0 < args.r0 <= 1.0):
        sys.exit("[ERROR] r0 must be in (0,1].")
    if not (0.0 < args.tau_pinch < 1.0):
        sys.exit("[ERROR] --pinch must be in (0,1).")
    if not (args.tau_S >= 1.0):
        sys.exit("[ERROR] --S must be ≥ 1.0 (central step longer than flanks).")

    hex_xlim = tuple(args.hex_xlim) if args.hex_xlim is not None else None
    hex_ylim = tuple(args.hex_ylim) if args.hex_ylim is not None else None

    analyze(args.npz, args.outdir, args.dist, args.min_runlen,
            args.r0, args.tau_pinch, args.tau_S, args.hex_gridsize,
            hex_xlim, hex_ylim, args.fonttype)

if __name__ == "__main__":
    main()
```
