## Supplementary material for "CAD-C: An engineered nuclease enables repair-free *in situ* proximity ligation and nucleosome-resolution chromosome walks in human cells": Custom code: Dmax_empirical_vs_signedstep_resampling_null_Lstratified_v2_20251021.py

```
#!/usr/bin/env python3
# -*- coding: utf-8 -*-
"""
Dmax (M or J) — empirical vs. signed-step resampling null for CADpack v9
=======================================================================

Purpose
-------
Show that empirical Dmax distributions (by run length L) are more *discretized*
and have a lower median than chance, using a null that resamples **signed step
vectors** from the **same L stratum** and rebuilds synthetic runs.

Geometries (both supported; default = mids)
-------------------------------------------
• mids (DmaxM): fragments → midpoints from canonical bounds:
    mid_i = 0.5 * (frag_ref_lo_i + frag_ref_hi_i)  for i=0..L
    empirical steps  : Δmid_i = mid_{i+1} − mid_i             (L steps)
    empirical Dmax_M : max(mid) − min(mid)
    null steps       : sample L signed Δmid from the L-stratum pool

• junctions (DmaxJ): reconstruct junction positions from v9 step deltas:
    signed_dJ_i = step_sgnJ_i * step_dJ_i                      (i=0..L−1)
    jpos path    : [0, cumsum(signed_dJ)]                      (L+1 points)
    empirical Dmax_J : max(jpos) − min(jpos)
    null steps       : sample L signed_dJ from the L-stratum pool

CADpack v9 fields used (exact names)
------------------------------------
• Runs:      run_L, run_first_frag_row, run_n_frags, run_first_step_row, run_n_steps
• Steps:     step_is_cis, chrom1_id, chrom2_id, step_dJ, step_sgnJ
• Fragments: frag_ref_chr_id, frag_ref_lo, frag_ref_hi, frag_len_bp
• Nice-to-have: run_read_uid (optional), chrom_names (optional)

Run-level QC (per L stratum)
----------------------------
1) **Cis + single chromosome**: all steps cis and same chrom id.
2) **No fragment overlaps** on reference: sort by frag_ref_lo; fail if next.lo < prev.hi.
3) **Extreme fragment lengths** removed: compute [q_lo, q_hi] from **positive**
   `frag_len_bp` across cis runs in the L stratum; exclude any run containing a
   fragment outside this range. Runs with any invalid canonical bounds (−1 or hi<=lo)
   are excluded (needed to compute midpoints and overlaps).

Plotting style (Illustrator‑friendly)
-------------------------------------
• Histograms in log10(D) with density=True (probability per *dex*), then drawn as
  stairs vs 10**edges on a true log‑x axis, preserving area=1 in log‑space.
• Decade major ticks + minor ticks 2..9; outward bottom ticks only; no top/right spines.
• All PDFs/EPS embed TrueType: pdf.fonttype=42, ps.fonttype=42.

Outputs (all under --outprefix*)
--------------------------------
• *_empirical.csv : per-run rows with labels & QC flags
• *_null.csv      : per-replicate null Dmax
• *_hist.csv      : bin_lo_bp, bin_hi_bp, emp_density_perdex, null_density_perdex
• *_summary.json  : parameters, medians, KS(log10), entropies, quantiles
• *_summary.csv   : one-line, grep-friendly
• *_hist.pdf      : semilog-x overlay (per‑dex density)

CLI
---
python dmax_emp_vs_null_v9.py \
  --npz /path/to/cadpack_v9.npz \
  --L 3 \
  --geometry mids \
  --null-reps 100000 \
  --frag-qlo 0.01 --frag-qhi 0.99 \
  --log-min 2 --log-max 9 --bins 60 \
  --seed 13 \
  --outprefix out/v9_L3_mids

Switch to junctions:
  --geometry junctions

Optional overrides (only if your NPZ deviates from v9 names):
  --frag-chrom-key frag_ref_chr_id --frag-start-key frag_ref_lo --frag-end-key frag_ref_hi
  --run-first-frag-key run_first_frag_row --run-n-frags-key run_n_frags --run-len-key run_L
  --run-first-step-key run_first_step_row --run-n-steps-key run_n_steps
  --step-dj-key step_dJ --step-sgnj-key step_sgnJ
  --step-is-cis-key step_is_cis --chrom1-key chrom1_id --chrom2-key chrom2_id
  --run-read-uid-key run_read_uid

Reproducibility: summary.json captures seed, thresholds, L, geometry, and the
exact key mapping used.

License: MIT
"""

from __future__ import annotations
import argparse, json, math, os, sys
from typing import Dict, List, Optional, Tuple

import numpy as np
import pandas as pd

import matplotlib
matplotlib.use("Agg")
matplotlib.rcParams.update({
    "pdf.fonttype": 42,   # Illustrator-friendly
    "ps.fonttype": 42,
    "axes.unicode_minus": False,
})
import matplotlib.pyplot as plt
from matplotlib.ticker import LogLocator, LogFormatterMathtext


# ---------- helpers ----------
def _pick(keys: List[str], candidates: List[str]) -> Optional[str]:
    for c in candidates:
        c = c.lower()
        for k in keys:
            if c in k.lower(): return k
    return None

def _resolve_keys_v9(npz) -> Tuple[Dict[str, str], List[str]]:
    keys = list(npz.files); km: Dict[str,str] = {}
    # canonical v9 names first
    km['frag_chrom'] = 'frag_ref_chr_id' if 'frag_ref_chr_id' in keys else _pick(keys, ['frag_ref_chr','frag_chr_id','frag_chrom'])
    km['frag_start'] = 'frag_ref_lo'     if 'frag_ref_lo'     in keys else _pick(keys, ['frag_lo','frag_start'])
    km['frag_end']   = 'frag_ref_hi'     if 'frag_ref_hi'     in keys else _pick(keys, ['frag_hi','frag_end'])
    km['frag_len']   = 'frag_len_bp'     if 'frag_len_bp'     in keys else _pick(keys, ['frag_len'])
    km['run_len']        = 'run_L'               if 'run_L'               in keys else _pick(keys, ['run_len'])
    km['run_first_frag'] = 'run_first_frag_row'  if 'run_first_frag_row'  in keys else _pick(keys, ['run_first_frag'])
    km['run_n_frags']    = 'run_n_frags'         if 'run_n_frags'         in keys else _pick(keys, ['run_nfrag'])
    km['run_first_step'] = 'run_first_step_row'  if 'run_first_step_row'  in keys else _pick(keys, ['run_first_step'])
    km['run_n_steps']    = 'run_n_steps'         if 'run_n_steps'         in keys else _pick(keys, ['run_nstep'])
    km['step_is_cis'] = 'step_is_cis' if 'step_is_cis' in keys else None
    km['chrom1_id']   = 'chrom1_id'   if 'chrom1_id'   in keys else None
    km['chrom2_id']   = 'chrom2_id'   if 'chrom2_id'   in keys else None
    km['step_dJ']     = 'step_dJ'     if 'step_dJ'     in keys else _pick(keys, ['dJ'])
    km['step_sgnJ']   = 'step_sgnJ'   if 'step_sgnJ'   in keys else _pick(keys, ['sgnJ'])
    # optional niceties
    km['run_read_uid']= 'run_read_uid' if 'run_read_uid' in keys else None
    km['chrom_names'] = 'chrom_names'  if 'chrom_names'  in keys else None
    return km, keys

def _cis_singlechrom_mask(step_is_cis, chrom1_id, chrom2_id, r_first_s, r_n_steps):
    keep = np.zeros(r_n_steps.size, dtype=bool)
    for i in range(r_n_steps.size):
        s0 = int(r_first_s[i]); n = int(r_n_steps[i])
        if n <= 0: continue
        S = slice(s0, s0 + n)
        if not np.all(step_is_cis[S]): continue
        c0 = chrom1_id[s0]
        if np.all(chrom1_id[S]==c0) and np.all(chrom2_id[S]==c0):
            keep[i] = True
    return keep

def _any_overlap(starts, ends):
    o = np.argsort(starts); s = starts[o]; e = ends[o]
    return np.any(s[1:] < e[:-1])  # strict overlap; touching OK

def _path_range_from_steps(steps):
    path = np.concatenate([[0.0], np.cumsum(steps)])
    return float(np.max(path) - np.min(path))

def _ks_D(x_emp, x_null):
    xe = np.sort(x_emp); xn = np.sort(x_null)
    xs = np.sort(np.unique(np.concatenate([xe, xn])))
    Fe = np.searchsorted(xe, xs, side="right")/len(xe)
    Fn = np.searchsorted(xn, xs, side="right")/len(xn)
    return float(np.max(np.abs(Fe - Fn)))

def _entropy_from_counts(counts, eps=1e-12):
    p = counts/(counts.sum()+eps); p = p[p>0]
    return float(-(p*np.log(p)).sum())


# ---------- main ----------
def main():
    ap = argparse.ArgumentParser(formatter_class=argparse.ArgumentDefaultsHelpFormatter)
    ap.add_argument("--npz", required=True)
    ap.add_argument("--L", type=int, required=True)
    ap.add_argument("--geometry", choices=["mids","junctions"], default="mids")
    ap.add_argument("--null-reps", type=int, default=100000)
    ap.add_argument("--frag-qlo", type=float, default=0.01)
    ap.add_argument("--frag-qhi", type=float, default=0.99)
    ap.add_argument("--log-min", type=float, default=2.0)
    ap.add_argument("--log-max", type=float, default=9.0)
    ap.add_argument("--bins", type=int, default=60)
    ap.add_argument("--seed", type=int, default=13)
    ap.add_argument("--outprefix", required=True)
    # optional overrides
    ap.add_argument("--frag-chrom-key"); ap.add_argument("--frag-start-key"); ap.add_argument("--frag-end-key"); ap.add_argument("--frag-len-key")
    ap.add_argument("--run-first-frag-key"); ap.add_argument("--run-n-frags-key"); ap.add_argument("--run-len-key")
    ap.add_argument("--run-first-step-key"); ap.add_argument("--run-n-steps-key")
    ap.add_argument("--step-dj-key"); ap.add_argument("--step-sgnj-key"); ap.add_argument("--step-is-cis-key")
    ap.add_argument("--chrom1-key"); ap.add_argument("--chrom2-key"); ap.add_argument("--run-read-uid-key"); ap.add_argument("--chrom-names-key")
    args = ap.parse_args()

    rng = np.random.default_rng(args.seed)
    Z = np.load(args.npz, allow_pickle=False)
    km, all_keys = _resolve_keys_v9(Z)

    # apply overrides if provided
    overrides = {
        'frag_chrom': args.frag_chrom_key, 'frag_start': args.frag_start_key, 'frag_end': args.frag_end_key,
        'frag_len': args.frag_len_key, 'run_first_frag': args.run_first_frag_key, 'run_n_frags': args.run_n_frags_key,
        'run_len': args.run_len_key, 'run_first_step': args.run_first_step_key, 'run_n_steps': args.run_n_steps_key,
        'step_dJ': args.step_dj_key, 'step_sgnJ': args.step_sgnj_key, 'step_is_cis': args.step_is_cis_key,
        'chrom1_id': args.chrom1_key, 'chrom2_id': args.chrom2_key, 'run_read_uid': args.run_read_uid_key,
        'chrom_names': args.chrom_names_key
    }
    for k,v in overrides.items():
        if v: km[k]=v

    # required
    required = ['frag_chrom','frag_start','frag_end','frag_len','run_len',
                'run_first_frag','run_n_frags','run_first_step','run_n_steps',
                'step_is_cis','chrom1_id','chrom2_id','step_dJ','step_sgnJ']
    missing = [k for k in required if km.get(k) not in all_keys]
    if missing:
        print("[ERROR] Missing expected v9 keys (or provide overrides):", missing, file=sys.stderr)
        print("Detected map:", km, file=sys.stderr)
        print("NPZ keys:", *all_keys, sep="\n  ", file=sys.stderr)
        sys.exit(2)

    # load arrays
    frag_chr   = Z[km['frag_chrom']]
    frag_lo    = Z[km['frag_start']].astype(np.int64)
    frag_hi    = Z[km['frag_end']].astype(np.int64)
    frag_lenbp = Z[km['frag_len']].astype(np.int64)

    run_L      = Z[km['run_len']].astype(np.int64)
    r_first_f  = Z[km['run_first_frag']].astype(np.int64)
    r_n_frags  = Z[km['run_n_frags']].astype(np.int64)
    r_first_s  = Z[km['run_first_step']].astype(np.int64)
    r_n_steps  = Z[km['run_n_steps']].astype(np.int64)

    step_is_cis= Z[km['step_is_cis']].astype(bool)
    chrom1_id  = Z[km['chrom1_id']].astype(np.int32)
    chrom2_id  = Z[km['chrom2_id']].astype(np.int32)
    step_dJmag = Z[km['step_dJ']].astype(np.float64)   # magnitude
    step_sgnJ  = Z[km['step_sgnJ']].astype(np.int8)    # -1/0/+1
    step_dJsig = step_sgnJ * np.abs(step_dJmag)        # signed dJ

    run_read_uid = Z[km['run_read_uid']] if (km.get('run_read_uid') in all_keys) else None
    chrom_names  = Z[km['chrom_names']]  if (km.get('chrom_names')  in all_keys) else None

    # select L stratum + cis+single chrom gating
    idx_runs_allL = np.where(run_L == args.L)[0]
    keep_cis = _cis_singlechrom_mask(step_is_cis, chrom1_id, chrom2_id, r_first_s, r_n_steps)
    idx_runs = idx_runs_allL[keep_cis[idx_runs_allL]]
    if idx_runs.size == 0:
        print(f"[ERROR] No cis-confined single-chrom runs with L={args.L}.", file=sys.stderr)
        sys.exit(3)

    # fragment-length quantiles within L (positive lengths only)
    lens_pos = []
    for r in idx_runs:
        fs = int(r_first_f[r]); fe = int(fs + r_n_frags[r])
        flen = frag_lenbp[fs:fe]
        lens_pos.append(flen[flen > 0])
    lens_pos = np.concatenate(lens_pos) if len(lens_pos) else np.array([], dtype=np.int64)
    if lens_pos.size >= 20:
        qlo = float(np.quantile(lens_pos, args.frag_qlo))
        qhi = float(np.quantile(lens_pos, args.frag_qhi))
        use_len_filter = True
    else:
        # fallback: compute from hi-lo where valid
        hi_lo_lengths = []
        for r in idx_runs:
            fs = int(r_first_f[r]); fe = int(fs + r_n_frags[r])
            lo = frag_lo[fs:fe]; hi = frag_hi[fs:fe]
            valid = (lo>=0) & (hi>=0) & (hi>lo)
            hi_lo_lengths.append((hi - lo)[valid])
        hi_lo_lengths = np.concatenate(hi_lo_lengths) if hi_lo_lengths else np.array([], dtype=np.int64)
        if hi_lo_lengths.size >= 20:
            qlo = float(np.quantile(hi_lo_lengths, args.frag_qlo))
            qhi = float(np.quantile(hi_lo_lengths, args.frag_qhi))
            use_len_filter = True
        else:
            qlo = qhi = float("nan")
            use_len_filter = False

    # iterate runs: QC + compute empirical Dmax + collect signed steps
    rows_emp = []
    emp_vals = []
    pool_steps = []

    geometry = args.geometry  # 'mids' or 'junctions'
    nsteps_for_null = args.L  # both geometries sample L signed steps

    for r in idx_runs:
        fs = int(r_first_f[r]); fe = int(fs + r_n_frags[r])
        ss = int(r_first_s[r]); se = int(ss + r_n_steps[r])

        # require exact L and L+1
        if (se - ss) != args.L or (fe - fs) != (args.L + 1):
            continue

        # fragment bounds present and on same chrom
        cids = frag_chr[fs:fe]
        los  = frag_lo[fs:fe]
        his  = frag_hi[fs:fe]
        valid_bounds = (cids >= 0) & (los >= 0) & (his >= 0) & (his > los)

        # length filter (if enabled)
        if use_len_filter:
            lengths = frag_lenbp[fs:fe]
            if np.any((lengths <= 0) | (lengths < qlo) | (lengths > qhi)):
                continue

        # all frags must be valid to compute midpoints & overlap
        if not np.all(valid_bounds):
            continue

        # same chrom (should hold after cis mask)
        if not np.all(cids == cids[0]):
            continue

        # no overlaps
        if _any_overlap(los, his):
            continue

        # compute empirical & pool steps
        if geometry == "mids":
            mids = (los + his) / 2.0  # L+1 points
            steps = np.diff(mids)     # L signed steps
            D = float(np.max(mids) - np.min(mids))
        else:
            signed_dJ = step_dJsig[ss:se]   # length L
            # path has L+1 points starting at 0
            jpos = np.concatenate([[0.0], np.cumsum(signed_dJ)])
            steps = signed_dJ               # L signed steps
            D = float(np.max(jpos) - np.min(jpos))

        emp_vals.append(D)
        pool_steps.append(steps)

        chrom_label = None
        try:
            chr_id = int(cids[0])
            if chrom_names is not None and 0 <= chr_id < len(chrom_names):
                chrom_label = chrom_names[chr_id].decode("utf-8", errors="ignore")
        except Exception:
            pass

        rows_emp.append({
            "run_idx": int(r),
            "run_L": int(run_L[r]),
            "chrom_id": int(cids[0]),
            "chrom": chrom_label if chrom_label is not None else "",
            "geometry": geometry,
            "n_steps": int(se - ss),
            "n_frags": int(fe - fs),
            "Dmax": D,
            "qc_cis_singlechrom": True,
            "qc_no_overlap": True,
            "qc_fraglen_ok": bool(use_len_filter),
            "frag_len_qlo_bp": qlo if use_len_filter else "",
            "frag_len_qhi_bp": qhi if use_len_filter else "",
            "run_read_uid": (str(run_read_uid[r]) if (run_read_uid is not None) else "")
        })

    if len(emp_vals) == 0:
        print("[ERROR] All runs filtered out by QC.", file=sys.stderr)
        sys.exit(4)

    emp_vals = np.asarray(emp_vals, dtype=np.float64)
    pool = np.concatenate(pool_steps).astype(np.float64)

    # null resampling
    draws = rng.choice(pool, size=(args.null_reps, nsteps_for_null), replace=True)
    null_vals = np.apply_along_axis(_path_range_from_steps, 1, draws)

    # ---------- save tabular outputs ----------
    pd.DataFrame(rows_emp).to_csv(f"{args.outprefix}_empirical.csv", index=False)
    pd.DataFrame({"replicate": np.arange(1, args.null_reps+1, dtype=np.int64),
                  "geometry": geometry, "Dmax": null_vals}).to_csv(f"{args.outprefix}_null.csv", index=False)

    # histogram table (per-dex densities)
    edges_log = np.linspace(args.log_min, args.log_max, args.bins+1)
    edges_bp  = 10.0 ** edges_log
    dens_emp, _  = np.histogram(np.log10(emp_vals),  bins=edges_log, density=True)
    dens_null, _ = np.histogram(np.log10(null_vals), bins=edges_log, density=True)
    pd.DataFrame({
        "bin_lo_bp": edges_bp[:-1],
        "bin_hi_bp": edges_bp[1:],
        "emp_density_perdex": dens_emp,
        "null_density_perdex": dens_null
    }).to_csv(f"{args.outprefix}_hist.csv", index=False)

    # stats
    H_emp  = _entropy_from_counts(np.histogram(np.log10(emp_vals),  bins=edges_log)[0])
    H_null = _entropy_from_counts(np.histogram(np.log10(null_vals), bins=edges_log)[0])
    ksD    = _ks_D(np.log10(emp_vals), np.log10(null_vals))

    summary = {
        "npz_path": os.path.abspath(args.npz),
        "geometry": geometry,
        "L": int(args.L),
        "null_replicates": int(args.null_reps),
        "n_runs_empirical": int(emp_vals.size),
        "n_steps_in_pool": int(pool.size),
        "frag_len_filter_enabled": bool(use_len_filter),
        "frag_len_quantiles_bp": {"qlo": (float(qlo) if use_len_filter else None),
                                  "qhi": (float(qhi) if use_len_filter else None)},
        "log_bin_params": {"log_min": float(args.log_min), "log_max": float(args.log_max), "bins": int(args.bins)},
        "median_empirical_bp": float(np.median(emp_vals)),
        "p25_empirical_bp": float(np.percentile(emp_vals, 25)),
        "p75_empirical_bp": float(np.percentile(emp_vals, 75)),
        "median_null_bp": float(np.median(null_vals)),
        "p25_null_bp": float(np.percentile(null_vals, 25)),
        "p75_null_bp": float(np.percentile(null_vals, 75)),
        "median_ratio_null_over_emp": float(np.median(null_vals)/np.median(emp_vals)),
        "KS_D_log10": float(ksD),
        "entropy_empirical_nats": float(H_emp),
        "entropy_null_nats": float(H_null),
        "entropy_diff_emp_minus_null": float(H_emp - H_null),
        "rng_seed": int(args.seed),
        "resolved_keys": km,
    }
    with open(f"{args.outprefix}_summary.json","w") as fh:
        json.dump(summary, fh, indent=2)

    pd.DataFrame([{
        "npz": os.path.basename(args.npz),
        "geometry": geometry,
        "L": int(args.L),
        "n_emp": int(emp_vals.size),
        "n_null": int(args.null_reps),
        "med_emp_bp": summary["median_empirical_bp"],
        "med_null_bp": summary["median_null_bp"],
        "ratio_null_over_emp": summary["median_ratio_null_over_emp"],
        "KS_D_log10": ksD,
        "H_emp": H_emp,
        "H_null": H_null,
        "H_diff": H_emp - H_null
    }]).to_csv(f"{args.outprefix}_summary.csv", index=False)

    # ---------- plot: true semilog-x, per-dex density ----------
    fig = plt.figure(figsize=(6.8, 4.6), constrained_layout=True)
    ax = fig.add_subplot(1,1,1)
    ax.set_xscale("log", base=10.0)

    ax.step(edges_bp[:-1], dens_emp,  where="post", linewidth=1.8,
            label=f"empirical L={args.L} ({geometry})")
    ax.step(edges_bp[:-1], dens_null, where="post", linewidth=1.2, linestyle="dashed",
            label="null (resampled signed steps)")

    ax.set_xlim(10**args.log_min, 10**args.log_max)
    ax.xaxis.set_major_locator(LogLocator(base=10.0, numticks=8))
    ax.xaxis.set_major_formatter(LogFormatterMathtext(base=10.0))
    ax.xaxis.set_minor_locator(LogLocator(base=10.0, subs=np.arange(2,10)*0.1, numticks=80))
    ax.tick_params(axis="x", which="both", direction="out", top=False, bottom=True, labeltop=False)
    ax.tick_params(axis="y", which="both", direction="out")
    for spine in ("top","right"): ax.spines[spine].set_visible(False)

    ax.set_ylabel(r"density per log$_{10}$ bp (area = 1 in log-space)")
    xlab = r"Max within-run separation, D$_{max}$ (bp) " + ("[midpoints]" if geometry=="mids" else "[junctions]")
    ax.set_xlabel(xlab)

    ymax = float(max(np.nanmax(dens_emp) if dens_emp.size else 0.0,
                     np.nanmax(dens_null) if dens_null.size else 0.0))
    ax.set_ylim(0, ymax*1.08 if ymax>0 else 1)
    ax.legend(frameon=False, loc="upper right")

    fig.savefig(f"{args.outprefix}_hist.pdf", dpi=300)
    plt.close(fig)

if __name__ == "__main__":
    main()
```
