## Supplementary material for "CAD-C: An engineered nuclease enables repair-free *in situ* proximity ligation and nucleosome-resolution chromosome walks in human cells": Custom code: rgs_pattern_census_v9_cli_20251021_151529.py

```
#!/usr/bin/env python3
# -*- coding: utf-8 -*-
"""
Restricted-Growth Pattern Census from CADpack v9 — Baseline vs Strict Non-Overlap
=================================================================================

Author: Jan Soroczynski, Risca Lab, The Rockefeller University
Date: 2025_10_21

Purpose
-------
From a CADpack **v9** NPZ, compute, for **all run lengths L > 1** (i.e., at least 3 fragments),
the frequency of **chromosome-visit patterns** encoded as **restricted-growth strings** (RGS)
over the **fragment chromosomes** within each run (length L → L+1 fragments). For each pattern,
report **Baseline** counts (no overlap gating) and **Strict non-overlap** counts (within-chromosome
fragment intervals must be strictly separated; touching counts as overlap). Also report the
**retained fraction** (strict/baseline).

Key definitions
---------------
- Run length **L**: number of steps in a run (fragments per run = L+1).
- Fragment sequence per run: the ordered list of fragments’ reference chromosomes.
- **RGS pattern**: map the first chromosome encountered to '0', the next *new* chromosome to '1', etc.
  Example for L=4 (5 fragments):
    00000  (all 5 on the same chromosome),
    01000  (two-chrom alternating start then same), … up to 01234 (all 5 on distinct chromosomes).
  This avoids ambiguity of letter labels (A/B/…) and generalizes to >2 chromosomes gracefully.
- **Strict non-overlap** filter: within a run, for *each* chromosome, sort that chromosome’s
  fragment intervals by start and require `next_start > prev_end` (disallow touching and overlap).
  Long fragments are **not** filtered by length unless they cause overlap; this matches the user spec.

Inputs (required NPZ arrays)
----------------------------
- run_n_steps, run_first_frag_row, run_n_frags
- frag_ref_chr_id, frag_ref_lo, frag_ref_hi
- (Optional) chrom_names is read if present for diagnostics only

Outputs
-------
- A **timestamped output directory** inside --outdir with:
  1) **CSV** per-L pattern table (baseline_count, strict_nonoverlap_count, retained_fraction).
  2) **CSV** all-L concatenated table.
  3) **PDF** plots per-L:
      • counts side-by-side (Baseline vs Strict)
      • retained fraction bars
  4) **metadata.json** with script/version/env/CLI/config and aggregate counts.
  5) **rgs_census.log** structured log.
- PDFs are exported with **fonttype=42** (Illustrator-compatible), matplotlib-only.

CLI arguments
-------------
--npz PATH               Path to cadpack v9 NPZ.
--outdir DIR            Output root directory (script creates a timestamped subdir).
--lmin INT              Minimum L to include (default: 2).
--lmax INT              Maximum L to include (default: auto = max in NPZ).
--include-theoretical   If set, include **all theoretical RGS** of length (L+1) in the tables,
                        not just those observed in data (default: off).
--max-theoretical-L INT Safety cap: only enumerate all-theoretical RGS when L <= this cap (default: 6).
--config PATH           Optional YAML/JSON config with any of the above keys; CLI overrides config.
--allow-touching        Allow touching intervals to pass the non-overlap filter (default: touching fails).

Examples
--------
# Typical run (all L>1 found in the NPZ, only observed patterns)
python rgs_pattern_census_v9_cli.py \  --npz /path/to/cadpack_bc1001_bc1002_v9_14422039.npz \  --outdir ./out

# L restricted to 4..4 (replicates your L=4 analysis), include all theoretical RGS for length 5
python rgs_pattern_census_v9_cli.py \  --npz /path/to/cadpack_bc1001_bc1002_v9_14422039.npz \  --outdir ./out --lmin 4 --lmax 4 --include-theoretical --max-theoretical-L 8

Required nonstandard packages
-----------------------------
- numpy, pandas, matplotlib, pyyaml

Reproducibility & logging
-------------------------
- Writes a metadata.json with the precise CLI, timestamp, environment, L range, and counts.
- Uses deterministic RGS mapping per run by first-seen order.
- No randomization.

"""
from __future__ import annotations
import os, sys, json, argparse, datetime, platform, getpass, math, re, logging
from typing import Dict, List, Tuple
import numpy as np
import pandas as pd
import matplotlib
import matplotlib.pyplot as plt

SCRIPT_VERSION = "1.0.0"

def load_config(path: str) -> dict:
    if path is None: return {}
    # Try YAML then JSON
    try:
        import yaml
        with open(path, "r") as f:
            return yaml.safe_load(f) or {}
    except Exception:
        try:
            with open(path, "r") as f:
                return json.load(f)
        except Exception as e:
            raise RuntimeError(f"Could not parse config file {path}: {e}")

def ensure_dir(path: str):
    os.makedirs(path, exist_ok=True)

def now_stamp():
    return datetime.datetime.now().strftime("%Y%m%d_%H%M%S")

def strict_nonoverlap_for_run(lo: np.ndarray, hi: np.ndarray, chr_ids: np.ndarray, disallow_touching=True) -> bool:
    for c in np.unique(chr_ids):
        idx = np.where(chr_ids == c)[0]
        if idx.size <= 1:
            continue
        intervals = sorted([(int(lo[i]), int(hi[i])) for i in idx], key=lambda x: x[0])
        prev_lo, prev_hi = intervals[0]
        for (a,b) in intervals[1:]:
            if disallow_touching:
                if a <= prev_hi:
                    return False
            else:
                if a < prev_hi:
                    return False
            prev_lo, prev_hi = a, b
    return True

def rgs_label(chr_seq: np.ndarray) -> str:
    mapping: Dict[int,int] = {}
    next_code = 0
    out: List[str] = []
    for c in chr_seq.tolist():
        c = int(c)
        if c not in mapping:
            mapping[c] = next_code
            next_code += 1
        out.append(str(mapping[c]))
    return ''.join(out)

def enumerate_rgs(n_frag: int) -> List[str]:
    res: List[str] = []
    def rec(prefix: List[int], maxval: int):
        if len(prefix) == n_frag:
            res.append(''.join(map(str, prefix)))
            return
        for v in range(0, maxval + 2):
            if len(prefix) == 0 and v != 0:
                continue
            rec(prefix + [v], max(maxval, v))
    rec([0], 0)
    return res

def setup_logger(log_path: str) -> logging.Logger:
    logger = logging.getLogger("rgs_census")
    logger.setLevel(logging.INFO)
    fmt = logging.Formatter("%(asctime)s | %(levelname)s | %(message)s")
    fh = logging.FileHandler(log_path, mode="w")
    fh.setFormatter(fmt)
    logger.addHandler(fh)
    sh = logging.StreamHandler(sys.stdout)
    sh.setFormatter(fmt)
    logger.addHandler(sh)
    return logger

def analyze(npz_path: str, outdir: str, lmin: int, lmax: int|None,
            include_theoretical: bool, max_theoretical_L: int, disallow_touching: bool=True) -> dict:
    matplotlib.rcParams['pdf.fonttype'] = 42
    matplotlib.rcParams['ps.fonttype'] = 42

    Z = np.load(npz_path, allow_pickle=True)
    required = ["run_n_steps","run_first_frag_row","run_n_frags","frag_ref_chr_id","frag_ref_lo","frag_ref_hi"]
    missing = [k for k in required if k not in Z.keys()]
    if missing:
        raise RuntimeError(f"Required arrays missing in NPZ: {missing}")

    run_n_steps = Z["run_n_steps"]
    run_first_frag_row = Z["run_first_frag_row"]
    run_n_frags = Z["run_n_frags"]
    frag_chr = Z["frag_ref_chr_id"]
    frag_lo  = Z["frag_ref_lo"]
    frag_hi  = Z["frag_ref_hi"]

    L_max_in_data = int(np.max(run_n_steps)) if run_n_steps.size else 0
    if lmax is None:
        lmax = L_max_in_data
    lmin = max(2, int(lmin))
    lmax = max(lmin, int(lmax))

    stamp = now_stamp()
    out_root = os.path.join(outdir, f"rgs_census_{stamp}")
    os.makedirs(out_root, exist_ok=True)
    log_path = os.path.join(out_root, "rgs_census.log")
    logger = setup_logger(log_path)
    logger.info(f"Starting RGS census v{SCRIPT_VERSION}")
    logger.info(f"NPZ: {npz_path}")
    logger.info(f"L range: {lmin}..{lmax} (max in data = {L_max_in_data})")
    logger.info(f"include_theoretical={include_theoretical}, max_theoretical_L={max_theoretical_L}, disallow_touching={disallow_touching}")

    tables = []
    agg = []

    for L in range(lmin, lmax+1):
        n_frag = L + 1
        idxs = np.where(run_n_steps == L)[0]

        counts_base: Dict[str,int] = {}
        counts_strict: Dict[str,int] = {}
        total_base = 0
        total_strict = 0

        for i in idxs.tolist():
            f0 = int(run_first_frag_row[i])
            nF = int(run_n_frags[i])
            if nF != n_frag:
                continue
            chr_ids = frag_chr[f0:f0+n_frag]
            lo      = frag_lo[f0:f0+n_frag]
            hi      = frag_hi[f0:f0+n_frag]

            pat = rgs_label(chr_ids)
            counts_base[pat] = counts_base.get(pat,0) + 1
            total_base += 1

            if strict_nonoverlap_for_run(lo, hi, chr_ids, disallow_touching=disallow_touching):
                counts_strict[pat] = counts_strict.get(pat,0) + 1
                total_strict += 1

        if include_theoretical and L <= max_theoretical_L:
            patterns = enumerate_rgs(n_frag)
        else:
            patterns = sorted(counts_base.keys())

        rows = []
        for pat in patterns:
            nb = counts_base.get(pat, 0)
            ns = counts_strict.get(pat, 0)
            frac = (ns/nb) if nb>0 else float("nan")
            rows.append({
                "L": L,
                "pattern": pat,
                "n_unique_chroms": (1 + max(map(int, pat))) if len(pat) else 0,
                "baseline_count": int(nb),
                "strict_nonoverlap_count": int(ns),
                "retained_fraction": (round(frac, 3) if nb>0 else "NA")
            })

        df_L = pd.DataFrame(rows).sort_values(["n_unique_chroms","pattern"]).reset_index(drop=True)
        tot_row = pd.DataFrame([{
            "L": L, "pattern": "TOTAL", "n_unique_chroms": "",
            "baseline_count": int(total_base),
            "strict_nonoverlap_count": int(total_strict),
            "retained_fraction": (round(total_strict/total_base, 3) if total_base>0 else "NA")
        }])
        df_L_out = pd.concat([df_L, tot_row], ignore_index=True)

        csv_L = os.path.join(out_root, f"rgs_L{L}_patterns.csv")
        df_L_out.to_csv(csv_L, index=False)
        logger.info(f"L={L}: runs={len(idxs)} | patterns={len(patterns)} | CSV={os.path.basename(csv_L)}")

        df_plot = df_L.copy()
        if not df_plot.empty:
            X = np.arange(df_plot.shape[0])
            width = 0.4
            fig_w = max(10, 0.35 * df_plot.shape[0] * 2)
            plt.figure(figsize=(fig_w, 6))
            plt.bar(X - width/2, df_plot["baseline_count"].values, width=width)
            plt.bar(X + width/2, df_plot["strict_nonoverlap_count"].values, width=width)
            plt.xticks(X, df_plot["pattern"].tolist(), rotation=90)
            plt.ylabel("Count (runs)")
            plt.title(f"L={L} (n_frag={n_frag}) — RGS pattern counts (Baseline vs Strict)")
            from matplotlib.patches import Patch
            patches = [Patch(label="Baseline"), Patch(label="Strict non-overlap")]
            plt.legend(handles=patches, frameon=False)
            plt.tight_layout()
            pdf1 = os.path.join(out_root, f"rgs_L{L}_counts.pdf")
            png1 = os.path.join(out_root, f"rgs_L{L}_counts.png")
            plt.savefig(pdf1, bbox_inches='tight')
            plt.savefig(png1, dpi=200, bbox_inches='tight')
            plt.close()

            plt.figure(figsize=(fig_w, 4.5))
            vals = [ (float('nan') if v=='NA' else float(v)) for v in df_plot['retained_fraction'].tolist() ]
            plt.bar(X, vals)
            plt.xticks(X, df_plot["pattern"].tolist(), rotation=90)
            plt.ylim(0,1)
            plt.ylabel("Retained fraction")
            plt.title(f"L={L} (n_frag={n_frag}) — Fraction retained after strict non-overlap")
            plt.tight_layout()
            pdf2 = os.path.join(out_root, f"rgs_L{L}_retention.pdf")
            png2 = os.path.join(out_root, f"rgs_L{L}_retention.png")
            plt.savefig(pdf2, bbox_inches='tight')
            plt.savefig(png2, dpi=200, bbox_inches='tight')
            plt.close()

        tables.append(df_L_out)
        agg.append({
            "L": L, "n_runs": int(len(idxs)),
            "n_patterns_observed": int(len([k for k,v in counts_base.items() if v>0])),
            "total_baseline": int(total_base),
            "total_strict": int(total_strict),
            "retained_fraction_overall": (round(total_strict/total_base, 3) if total_base>0 else "NA")
        })

    all_table = pd.concat(tables, ignore_index=True) if tables else pd.DataFrame()
    csv_all = os.path.join(out_root, "rgs_all_L_patterns.csv")
    all_table.to_csv(csv_all, index=False)

    meta = {
        "script_version": SCRIPT_VERSION,
        "timestamp": now_stamp(),
        "python": platform.python_version(),
        "platform": platform.platform(),
        "npz_path": npz_path,
        "lmin": lmin, "lmax": lmax,
        "include_theoretical": include_theoretical,
        "max_theoretical_L": max_theoretical_L,
        "disallow_touching": disallow_touching,
        "outputs": {
            "out_root": out_root,
            "csv_all": csv_all,
            "per_L_csv": [os.path.join(out_root, f) for f in os.listdir(out_root) if re.match(r"rgs_L\d+_patterns\.csv$", f)],
            "plots_pdf": [os.path.join(out_root, f) for f in os.listdir(out_root) if f.endswith(".pdf")]
        },
        "aggregate": agg
    }
    with open(os.path.join(out_root, "metadata.json"), "w") as w:
        json.dump(meta, w, indent=2)

    return {"out_root": out_root, "csv_all": csv_all, "metadata": meta}

def parse_args(argv):
    import argparse
    p = argparse.ArgumentParser(description="RGS pattern census from CADpack v9 — baseline vs strict non-overlap (L>1).")
    p.add_argument("--npz", required=True, help="Path to cadpack v9 NPZ.")
    p.add_argument("--outdir", required=True, help="Output root directory (timestamped subdir will be created).")
    p.add_argument("--lmin", type=int, default=2, help="Minimum L to include (default: 2).")
    p.add_argument("--lmax", type=int, default=None, help="Maximum L to include (default: max in NPZ).")
    p.add_argument("--include-theoretical", action="store_true",
                   help="Include all theoretical RGS patterns for length (L+1) if L <= --max-theoretical-L.")
    p.add_argument("--max-theoretical-L", type=int, default=6,
                   help="Safety cap for theoretical enumeration (default: 6).")
    p.add_argument("--config", default=None, help="Optional YAML/JSON config; CLI overrides config.")
    p.add_argument("--allow-touching", action="store_true",
                   help="If set, allow touching intervals (next_start == prev_end) to pass the non-overlap filter.")
    return p.parse_args(argv)

def main(argv):
    args = parse_args(argv)
    # Merge config then override with CLI
    cfg = {}
    if args.config:
        try:
            import yaml, json as _json
            with open(args.config, "r") as f:
                try:
                    cfg = yaml.safe_load(f) or {}
                except Exception:
                    f.seek(0)
                    cfg = _json.load(f)
        except Exception:
            cfg = {}

    npz_path = args.npz or cfg.get("npz")
    outdir   = args.outdir or cfg.get("outdir", ".")
    lmin     = int(args.lmin if args.lmin is not None else cfg.get("lmin", 2))
    lmax     = int(args.lmax) if args.lmax is not None else cfg.get("lmax") if cfg.get("lmax") is not None else None
    include_theoretical = bool(args.include_theoretical or cfg.get("include_theoretical", False))
    max_theoretical_L   = int(args.max_theoretical_L if args.max_theoretical_L is not None else cfg.get("max_theoretical_L", 6))
    disallow_touching   = not bool(args.allow_touching or cfg.get("allow_touching", False))

    if not os.path.exists(npz_path):
        raise FileNotFoundError(f"NPZ not found: {npz_path}")
    if not os.path.isdir(outdir):
        os.makedirs(outdir, exist_ok=True)

    res = analyze(npz_path=npz_path, outdir=outdir, lmin=lmin, lmax=lmax,
                  include_theoretical=include_theoretical, max_theoretical_L=max_theoretical_L,
                  disallow_touching=disallow_touching)
    print(json.dumps(res["metadata"], indent=2))
    print(f"All tables CSV: {res['csv_all']}")
    print(f"Output directory: {res['out_root']}")
    return 0

if __name__ == "__main__":
    import sys, platform
    raise SystemExit(main(sys.argv[1:]))
```
