## Supplementary material for "CAD-C: An engineered nuclease enables repair-free *in situ* proximity ligation and nucleosome-resolution chromosome walks in human cells": Custom code: streaming_PBCADwalk_analyses_v7_20251017.py

```
#!/usr/bin/env python3
# -*- coding: utf-8 -*-
"""
CADWALK-CLI v7 — PacBio CADwalk streaming analytics (pairtools parse2 .pairs.gz)

Author : Jan Soroczynski (Risca Lab, Rockefeller University)
Version: 7.0.0
License: MIT

================================================================================
README (Executive Overview)
================================================================================
Purpose
-------
This tool streams one or more PacBio CADwalk `.pairs.gz` files (pairtools parse2,
"pinned" schema), performs *read-only* analytics, and writes reproducible artifacts:
NumPy arrays (.npy/.npz), CSV/JSON summaries, and vector PDFs (Matplotlib fonttype=42).

What’s fixed vs v6
------------------
• **Non-adjacent cis reservoir**: Correct Algorithm-R implementation.
  - Fill by appending until capacity (k) is reached.
  - For each subsequent value t (>k), draw `i ~ U{0..t}`; if `i<k`, replace `res[i]`.
  - Prevents out-of-bounds writes when the first batch is small.

What’s improved vs v6
---------------------
• **Step-pair hexbin by B_run quartiles (merged)**:
  - While parsing runs we now store each adjacent cis pair `(d_i, d_{i+1})` **with its run’s B_run**.
  - Merged analysis plots square hexbins for **ALL** and for **each B_run quartile**.
• Otherwise, v7 preserves v6 outputs and directory structure.

Kept from v6 (requested features)
---------------------------------
• **Tortuosity by uninterrupted CIS run length** (longest same-chrom block): strata {1,2,3,≥4}.
  Hexbin: D_max vs B_run for each TR and for merged.
• **Fragment length** (U fragments, dedup per read):
  - Histogram 0–250 bp (bin=3) PDF; store **bin=1** and **bin=3** arrays up to `--frag-ecdf-max`.
  - ECDFs with common x-limits; **stratified** by CIS-block length strata and ALL.
• **B_run deciles × D_max deciles (10×10)**:
  - For each cell: **query-overlap** (on-read U-interval reuse) vs global.
  - **Mann–Whitney U (two-sided)** p, **Cliff’s δ**, **BH–FDR** q; store decile edges.
• **Adjacent vs non-adjacent cis separations**:
  - Small-scale histograms (0–`--small-hist-max`, bin=`--small-hist-bin`) and **ECDFs to 10^9 bp (log)**.
  - Non-adjacent cis separations are reservoir-sampled (size=`--nonadj-reservoir`).
• **UU-run composition by read-length deciles** (contiguous UU runs; cis or trans allowed):
  - Stacked bars per TR and merged.
• All plots **PDF fonttype=42**; arrays/CSVs/JSON are replot-ready.

Pinned input schema (0-based indices)
-------------------------------------
 0  readID           1  chrom1          2  pos1           3  chrom2
 4  pos2             5  strand1         6  strand2        7  pair_type
 8  walk_pair_index  9  walk_pair_type 10  mapq1          11 mapq2
12  pos5_1          13  pos5_2          14 pos3_1        15 pos3_2
16 cigar1           17 cigar2           18 read_len1      19 read_len2
20 matched_bp1      21 matched_bp2      22 algn_ref_span1 23 algn_ref_span2
24 algn_read_span1  25 algn_read_span2  26 dist_to_5_1    27 dist_to_5_2
28 dist_to_3_1      29 dist_to_3_2      30 seq1           31 seq2
32 mismatches1      33 mismatches2

Key definitions
---------------
• Fragment genomic midpoint: mid = 0.5*(min(pos5,pos3) + max(pos5,pos3)).
• On-read U interval: [dist_to_5, dist_to_5 + algn_read_span - 1].
• UU-run: consecutive rows with same readID and pair_type=="UU".
• CIS block: consecutive steps in a UU-run where both fragments are on the same chromosome.
  Longest CIS block length (steps) defines the 4 strata: {1,2,3,≥4}.
• Run-level tortuosity:
    For block with steps (d_j, s_j), s_j ∈ {±1}, d_j>0
        B_seg = 1 - |Σ s_j d_j| / Σ d_j
    B_run = 1 - [ Σ_blocks |Σ s_j d_j| ] / [ Σ_blocks Σ d_j ]
• D_max: max within-run cis separation via (max(mid) - min(mid)) per chromosome, then max over chromosomes.
• Query-overlap per run (on-read): let union_bp be the union length across all U intervals
  in the run; overlap_bp is bp covered by depth ≥2. Primary metric q1 = overlap_bp / union_bp.

Outputs
-------
OUTDIR/cadwalk_v7_<TS>/
  ├─ <label>/            # per TR
  │   ├─ arrays/         # .npy/.npz replot-ready
  │   ├─ summaries/      # CSV/JSON tables
  │   ├─ metadata/       # run_meta.json, columns_detected.json
  │   └─ plots/          # PDF fonttype=42
  └─ merged/             # pooled across TRs
      ├─ arrays/
      ├─ summaries/
      ├─ metadata/
      └─ plots/

Major arrays/tables
-------------------
• Tortuosity per run:
    arrays: brun.npy, dmax_bp.npy, log10_dmax.npy, run_len.npy,
            cis_longest_block.npy, cis_steps.npy, trans_steps.npy,
            alt_rate.npy, q_overlap_frac.npy, q_overcov_idx.npy, read_len.npy
    table : tortuosity_per_run.csv
• Step pairs (adjacent cis):
    per TR : arrays/step_pairs_xy.npz (x,y)
             arrays/step_pairs_xy_brun.npz (x,y,brun)   # NEW in v7
    merged : arrays/step_pairs_xy_brun.npz pooled
• Fragment lengths:
    arrays: fraglen_bin1_ALL.npy, fraglen_bin1_strata.npy
            fraglen_bin3_ALL.npy, fraglen_bin3_strata.npy
    table : fraglen_meta.json
• Adj/non-adj cis separations:
    arrays: adj_dists.npy, nonadj_reservoir.npy
            adj_ecdf_x.npy, adj_ecdf_y.npy
            nonadj_ecdf_x.npy, nonadj_ecdf_y.npy
    meta  : contact_spectra_meta.json
• UU-run composition:
    arrays: uu_run_counts.npy, uu_run_pct.npy
    table : uu_run_composition.csv
• Brun×Dmax (10×10) query-overlap grid (merged):
    arrays: (stats via CSV), decile_edges.json
    table : grid_q1_tests.csv (N, med, q25, q75, Mann–Whitney p, Cliff’s δ, BH–FDR q)

================================================================================
USAGE
================================================================================
Two TRs + merged:
    python streaming_PBCADwalk_analyses_v7_20251018.py \
        -o OUTDIR \
        --pairs bc1001.pairs.gz bc1002.pairs.gz \
        --label bc1001 bc1002 \
        --small-hist-bin 5 --small-hist-max 2000 \
        --nonadj-reservoir 2000000 \
        --frag-ecdf-max 1000

All plots are PDF (fonttype=42). Arrays/CSVs/JSON are replot-ready.
================================================================================
"""
from __future__ import annotations
import os, sys, gzip, json, csv, math, hashlib, platform, random, argparse, time
from dataclasses import dataclass
from typing import List, Tuple, Dict, Optional
from collections import defaultdict, namedtuple

import numpy as np

# ---------------- Matplotlib (headless) ----------------
def _mpl():
    import matplotlib
    matplotlib.use("Agg")
    import matplotlib.pyplot as plt
    matplotlib.rcParams['pdf.fonttype'] = 42
    matplotlib.rcParams['ps.fonttype']  = 42
    matplotlib.rcParams['font.size']    = 12
    matplotlib.rcParams['axes.titlesize']= 13
    matplotlib.rcParams['axes.labelsize']= 12
    return matplotlib, plt

# ---------------- Pinned schema ----------------
FieldMap = namedtuple("FieldMap", [
    "read_id","chrom1","pos1","chrom2","pos2","strand1","strand2","pair_type",
    "walk_pair_index","walk_pair_type","mapq1","mapq2",
    "pos51","pos52","pos31","pos32",
    "cigar1","cigar2",
    "read_len1","read_len2","matched_bp1","matched_bp2",
    "algn_ref_span1","algn_ref_span2","algn_read_span1","algn_read_span2",
    "dist_to_51","dist_to_52","dist_to_31","dist_to_32",
    "seq1","seq2","mismatches1","mismatches2"
])
PIN = FieldMap(
    read_id=0,  chrom1=1,  pos1=2,  chrom2=3,  pos2=4,  strand1=5,  strand2=6,
    pair_type=7, walk_pair_index=8, walk_pair_type=9, mapq1=10, mapq2=11,
    pos51=12, pos52=13, pos31=14, pos32=15, cigar1=16, cigar2=17,
    read_len1=18, read_len2=19, matched_bp1=20, matched_bp2=21,
    algn_ref_span1=22, algn_ref_span2=23, algn_read_span1=24, algn_read_span2=25,
    dist_to_51=26, dist_to_52=27, dist_to_31=28, dist_to_32=29,
    seq1=30, seq2=31, mismatches1=32, mismatches2=33
)

# ---------------- Utilities ----------------
def _now_ts_compact() -> str:
    import datetime
    return datetime.datetime.now().strftime("%Y%m%d_%H%M%S")

def _ensure_dirs(*paths):
    for p in paths:
        os.makedirs(p, exist_ok=True)

def _sha256(path: str) -> Optional[str]:
    try:
        h=hashlib.sha256()
        with open(path, "rb") as f:
            for chunk in iter(lambda: f.read(1<<20), b""):
                h.update(chunk)
        return h.hexdigest()
    except Exception:
        return None

def _safe_int(x: str) -> Optional[int]:
    try: return int(x)
    except: return None

def _midpoint(p5: Optional[int], p3: Optional[int]) -> Optional[float]:
    if p5 is None or p3 is None: return None
    a=min(p5,p3); b=max(p5,p3)
    return 0.5*(a+b)

def _rankdata(a: np.ndarray) -> np.ndarray:
    n=a.size
    order=np.argsort(a, kind="mergesort")
    ranks=np.empty(n, dtype=float)
    ranks[order]=np.arange(1, n+1, dtype=float)
    srt=a[order]; i=0
    while i<n:
        j=i+1
        while j<n and srt[j]==srt[i]: j+=1
        if j-i>1: ranks[order[i:j]]=(i+1+j)/2.0
        i=j
    return ranks

def _pearsonr(x: np.ndarray, y: np.ndarray) -> float:
    x=np.asarray(x,float); y=np.asarray(y,float)
    xm=x-x.mean(); ym=y-y.mean()
    den=np.sqrt((xm*xm).sum())*np.sqrt((ym*ym).sum())
    return float((xm*ym).sum()/den) if den>0 else float('nan')

def _spearmanr(x: np.ndarray, y: np.ndarray) -> float:
    return _pearsonr(_rankdata(x), _rankdata(y))

def _mannwhitney_u_p(x: np.ndarray, y: np.ndarray) -> float:
    if x.size==0 or y.size==0: return float('nan')
    R=np.concatenate([x,y])
    ranks=_rankdata(R)
    rx=ranks[:x.size].sum()
    U = rx - x.size*(x.size+1)/2.0
    mu= x.size*y.size/2.0
    sigma=math.sqrt(x.size*y.size*(x.size+y.size+1)/12.0)
    if sigma==0: return float('nan')
    z= (U - mu)/sigma
    # two-sided p using error function
    p = 2.0*0.5*(1.0 - math.erf(abs(z)/math.sqrt(2.0)))
    return float(min(max(p, 0.0), 1.0))

def _cliffs_delta(x: np.ndarray, y: np.ndarray) -> float:
    if x.size==0 or y.size==0: return float('nan')
    gt=0; lt=0
    for a in x:
        gt += (a>y).sum()
        lt += (a<y).sum()
    return float((gt-lt)/(x.size*y.size))

def _bh_fdr(pvals: List[float]) -> List[float]:
    m=len(pvals)
    pv = np.array([p if np.isfinite(p) else 1.0 for p in pvals], float)
    idx=np.argsort(pv)
    q=np.empty(m, float)
    prev=1.0
    for rank,i in enumerate(idx, start=1):
        val = pv[i]*m/rank
        prev=min(prev, val)
        q[i]=prev
    return [float(min(max(qi,0.0),1.0)) for qi in q.tolist()]

def _interval_union_and_overlap(intervals: List[Tuple[int,int]]) -> Tuple[int,int,int]:
    if not intervals: return 0,0,0
    ev=[]
    for s,e in intervals:
        if s>e: s,e=e,s
        ev.append((s, +1)); ev.append((e+1, -1))
    ev.sort()
    depth=0; last=None
    union_bp=0; overlap_bp=0; maxd=0; overcov=0
    for x,delta in ev:
        if last is not None and x>last and depth>0:
            L=x-last
            union_bp += L
            if depth>=2: overlap_bp += L
            overcov += max(depth-1,0)*L
        depth += delta
        maxd=max(maxd, depth)
        last=x
    return int(union_bp), int(overlap_bp), int(maxd)

# ---------------- Data structures ----------------
@dataclass
class RunStats:
    read_id: str
    run_ord: int
    run_len_steps: int
    n_frags: int
    n_cis_steps: int
    n_trans_steps: int
    cis_longest_block: int
    n_cis_blocks: int
    n_chroms: int
    alt_rate: float
    brun: float
    dmax_bp: float
    log10_dmax: float
    q_overlap_frac: float
    q_overcov_idx: float
    read_len: int

# ---------------- Streaming processor ----------------
class TRProcessor:
    def __init__(self, pairs_path: str, label: str, outdir: str,
                 small_hist_bin: int=5, small_hist_max: int=2000,
                 nonadj_reservoir: int=2_000_000, frag_ecdf_max: int=1000,
                 seed: int=17):
        self.pairs_path=pairs_path
        self.label=label
        self.outdir=outdir
        self.rng = np.random.default_rng(seed)

        # Adjacent/non-adj arrays
        self.adj_dists=[]

        # Reservoir (Algorithm R) — FIXED
        self.nadj_reservoir_target = int(nonadj_reservoir)
        self.nadj_res_list: List[float] = []   # dynamic; grows to capacity; then replace
        self.nadj_seen = 0

        self.small_hist_bin=small_hist_bin
        self.small_hist_max=small_hist_max

        # Step pairs and run-level Brun for each pair (for quartiles later)
        self.step_pairs_x=[]; self.step_pairs_y=[]; self.step_pairs_b=[]

        # Per-run stats
        self.run_stats: List[RunStats]=[]

        # Fragment lengths (dedup per read)
        self.frag_ecdf_max = frag_ecdf_max
        self.frag_counts_all_bin1 = np.zeros(frag_ecdf_max+1, np.int64)
        self.frag_counts_strata_bin1 = np.zeros((4, frag_ecdf_max+1), np.int64)
        self.frag_counts_all_bin3 = np.zeros((frag_ecdf_max//3)+1, np.int64)
        self.frag_counts_strata_bin3 = np.zeros((4, (frag_ecdf_max//3)+1), np.int64)

        # UU-run composition
        self.uur_readlens=[]; self.uur_runlens=[]

    @staticmethod
    def _cisblock_category(x: int) -> int:
        if x<=1: return 1
        if x==2: return 2
        if x==3: return 3
        return 4

    def _reservoir_add_many(self, vals: np.ndarray):
        """Algorithm R: append until capacity; then replace with prob k/t."""
        if vals.size==0: return
        for v in vals:
            self.nadj_seen += 1
            if len(self.nadj_res_list) < self.nadj_reservoir_target:
                self.nadj_res_list.append(float(v))
            else:
                # t = self.nadj_seen, k = target
                i = int(self.rng.integers(0, self.nadj_seen))
                if i < self.nadj_reservoir_target:
                    self.nadj_res_list[i] = float(v)

    def _process_run(self, read_id: str, run_ord: int, rows: List[List[str]],
                     per_read_frag_map: Dict[Tuple[str,int,int], int],
                     per_read_frag_cat: Dict[Tuple[str,int,int], int],
                     per_read_intervals: List[Tuple[int,int]]):
        if not rows: return

        # Build fragment chain
        p0=rows[0]
        m1=_midpoint(_safe_int(p0[PIN.pos51]), _safe_int(p0[PIN.pos31]))
        if m1 is None: return
        frag_chr=[p0[PIN.chrom1]]; frag_mid=[m1]
        read_len=0
        for p in rows:
            m2=_midpoint(_safe_int(p[PIN.pos52]), _safe_int(p[PIN.pos32]))
            if m2 is None: return
            frag_chr.append(p[PIN.chrom2]); frag_mid.append(m2)
            rl=max(_safe_int(p[PIN.read_len1]) or 0, _safe_int(p[PIN.read_len2]) or 0)
            if rl>read_len: read_len=rl

        n_frags=len(frag_mid); steps=n_frags-1
        if steps<=0: return

        # CIS steps, blocks, step_ds/step_ch
        cis_steps=0; trans_steps=0
        blocks=[]; cur_block_ch=None; cur_block=[]
        adj_cis_d=[]; step_ds=[]; step_ch=[]
        # Collect step pairs locally; attach B_run later
        local_pairs=[]

        for i in range(n_frags-1):
            c1,c2=frag_chr[i],frag_chr[i+1]
            d=abs(frag_mid[i+1]-frag_mid[i])
            if c1==c2 and d>0:
                cis_steps+=1
                sgn= 1 if frag_mid[i+1]>=frag_mid[i] else -1
                adj_cis_d.append(d)
                step_ds.append(d); step_ch.append(c1)
                if cur_block_ch is None or cur_block_ch!=c1:
                    if cur_block: blocks.append((cur_block_ch, cur_block))
                    cur_block_ch=c1; cur_block=[]
                cur_block.append((d, sgn))
            else:
                trans_steps+=1
                if cur_block: blocks.append((cur_block_ch, cur_block)); cur_block=[]; cur_block_ch=None
                step_ds.append(None); step_ch.append(None)
        if cur_block: blocks.append((cur_block_ch, cur_block))

        # Step pairs (within same-chrom contiguous ds)
        for i in range(len(step_ds)-1):
            if (step_ds[i] is not None) and (step_ds[i+1] is not None) and (step_ch[i]==step_ch[i+1]):
                local_pairs.append((step_ds[i], step_ds[i+1]))

        longest_cis = max((len(Lst) for _, Lst in blocks), default=0)

        # Alternation
        alt_cnt=0; alt_den=0
        for _, Lst in blocks:
            for j in range(len(Lst)-1):
                alt_den+=1
                if Lst[j][1]!=Lst[j+1][1]: alt_cnt+=1
        alt_rate = (alt_den and (alt_cnt/alt_den)) or float('nan')

        # B_run
        sum_abs_signed=0.0; sum_len=0.0
        for _, Lst in blocks:
            seg_sum = sum(d for d,_ in Lst)
            seg_signed = sum((sgn*d) for d,sgn in Lst)
            if seg_sum>0:
                sum_abs_signed += abs(seg_signed)
                sum_len += seg_sum
        brun = (sum_len and (1.0 - (sum_abs_signed/sum_len))) or float('nan')

        # Dmax
        per_ch=defaultdict(list)
        for c,x in zip(frag_chr, frag_mid): per_ch[c].append(x)
        dmax=0.0
        for c, xs in per_ch.items():
            if len(xs)>=2:
                span=max(xs)-min(xs)
                if span>dmax: dmax=span
        log10_dmax = float(np.log10(dmax)) if dmax>0 else float('nan')

        # Store adjacent cis
        if adj_cis_d: self.adj_dists.extend(adj_cis_d)

        # Non-adj cis reservoir (per chromosome, exclude adjacent)
        NONADJ_SAMPLE_PER_CH_CAP = 50_000
        for c, xs in per_ch.items():
            n=len(xs)
            if n>=3:
                if n*(n-1)//2 - (n-1) <= NONADJ_SAMPLE_PER_CH_CAP:
                    vals=[]
                    for i in range(n-1):
                        for j in range(i+2, n):
                            vals.append(abs(xs[j]-xs[i]))
                    if vals: self._reservoir_add_many(np.array(vals, float))
                else:
                    to_take = NONADJ_SAMPLE_PER_CH_CAP
                    for _ in range(to_take):
                        i = int(self.rng.integers(0, n-2))
                        j = int(self.rng.integers(i+2, n))
                        self._reservoir_add_many(np.array([abs(xs[j]-xs[i])], float))

        # Query-overlap (on-read)
        intervals=[]
        for p in rows:
            s1=_safe_int(p[PIN.dist_to_51]); L1=_safe_int(p[PIN.algn_read_span1])
            if s1 is not None and L1 and L1>0: intervals.append((s1, s1+L1-1))
            s2=_safe_int(p[PIN.dist_to_52]); L2=_safe_int(p[PIN.algn_read_span2])
            if s2 is not None and L2 and L2>0: intervals.append((s2, s2+L2-1))
        union_bp, overlap_bp, _maxd = _interval_union_and_overlap(intervals)
        q1 = (union_bp and (overlap_bp/union_bp)) or float('nan')
        # Overcoverage index (reuse) approximated in sweep above:
        # We compute it inline in _interval_union_and_overlap for union/overlap only; for q2 we zero for now.
        # For consistency with v6’s reporting, keep q2 as NaN (or extend sweep to record overcov explicitly).
        # Here we recompute with overcov:
        ev=[]
        for s,e in intervals:
            if s>e: s,e=e,s
            ev.append((s, +1)); ev.append((e+1, -1))
        ev.sort()
        depth=0; last=None; overcov=0
        for x,delta in ev:
            if last is not None and x>last and depth>0:
                L=x-last; overcov += max(depth-1,0)*L
            depth += delta; last=x
        q2 = (union_bp and (overcov/union_bp)) or float('nan')

        # UU-run composition inputs
        self.uur_readlens.append(read_len)
        self.uur_runlens.append(len(rows))

        # Register run
        self.run_stats.append(RunStats(
            read_id=read_id, run_ord=run_ord, run_len_steps=len(rows),
            n_frags=n_frags, n_cis_steps=cis_steps, n_trans_steps=trans_steps,
            cis_longest_block=longest_cis, n_cis_blocks=len(blocks),
            n_chroms=len(per_ch), alt_rate=alt_rate, brun=brun,
            dmax_bp=dmax, log10_dmax=log10_dmax, q_overlap_frac=q1,
            q_overcov_idx=q2, read_len=read_len
        ))

        # Attach run’s brun to each local step pair
        if local_pairs and np.isfinite(brun):
            for (dx, dy) in local_pairs:
                self.step_pairs_x.append(dx)
                self.step_pairs_y.append(dy)
                self.step_pairs_b.append(brun)

        # Fragment length strata assignment (dedup per read)
        cat=self._cisblock_category(longest_cis)
        for p in rows:
            # side 1 U?
            if p[PIN.pair_type][0]=="U":
                a=_safe_int(p[PIN.pos51]); b=_safe_int(p[PIN.pos31])
                if a is not None and b is not None:
                    s=min(a,b); e=max(a,b); L=e-s+1
                    key=(p[PIN.chrom1], s, e)
                    if key not in per_read_frag_map: per_read_frag_map[key]=L
                    per_read_frag_cat[key]= max(per_read_frag_cat.get(key,1), cat)
            # side 2 U?
            if p[PIN.pair_type][1]=="U":
                a=_safe_int(p[PIN.pos52]); b=_safe_int(p[PIN.pos32])
                if a is not None and b is not None:
                    s=min(a,b); e=max(a,b); L=e-s+1
                    key=(p[PIN.chrom2], s, e)
                    if key not in per_read_frag_map: per_read_frag_map[key]=L
                    per_read_frag_cat[key]= max(per_read_frag_cat.get(key,1), cat)

        per_read_intervals.extend(intervals)

    def _flush_read_fragments(self, frag_map, frag_cat):
        if not frag_map: return
        for key, L in frag_map.items():
            Lc = max(0, min(L, self.frag_ecdf_max))
            # ALL
            self.frag_counts_all_bin1[Lc] += 1
            self.frag_counts_all_bin3[Lc//3] += 1
            # strata
            cat = frag_cat.get(key, None)
            if cat is not None:
                idx = self._cisblock_category(cat)-1
                self.frag_counts_strata_bin1[idx, Lc] += 1
                self.frag_counts_strata_bin3[idx, Lc//3] += 1

    def run(self):
        _ensure_dirs(self.outdir, os.path.join(self.outdir,"arrays"),
                     os.path.join(self.outdir,"summaries"),
                     os.path.join(self.outdir,"metadata"),
                     os.path.join(self.outdir,"plots"))

        # Metadata
        run_meta={
            "label": self.label,
            "pairs_path": self.pairs_path,
            "sha256": _sha256(self.pairs_path),
            "platform": {
                "python": sys.version.split()[0],
                "platform": platform.platform(),
                "machine": platform.machine(),
                "cpu_count": os.cpu_count()
            },
            "params": {
                "small_hist_bin": self.small_hist_bin,
                "small_hist_max": self.small_hist_max,
                "nonadj_reservoir": self.nadj_reservoir_target,
                "frag_ecdf_max": self.frag_ecdf_max
            }
        }
        with open(os.path.join(self.outdir,"metadata","run_meta.json"),"w") as f:
            json.dump(run_meta, f, indent=2)
        with open(os.path.join(self.outdir,"metadata","columns_detected.json"),"w") as f:
            json.dump(PIN._asdict(), f, indent=2)

        # Stream file
        t0=time.time()
        with gzip.open(self.pairs_path, "rt", encoding="utf-8", errors="replace") as fin:
            cur_read=None; run_rows=[]; run_ord=0; read_len=0
            frag_map={}; frag_cat={}; read_intervals=[]
            for line in fin:
                if not line or line[0]=="#": continue
                p=line.rstrip("\n").split("\t")
                rid=p[PIN.read_id]
                if rid!=cur_read:
                    if run_rows:
                        self._process_run(cur_read, run_ord, run_rows, frag_map, frag_cat, read_intervals)
                        run_rows=[]
                    self._flush_read_fragments(frag_map, frag_cat)
                    frag_map={}; frag_cat={}; read_intervals=[]
                    cur_read=rid; run_ord=0; read_len=0

                rl=max(_safe_int(p[PIN.read_len1]) or 0, _safe_int(p[PIN.read_len2]) or 0)
                if rl>read_len: read_len=rl

                if p[PIN.pair_type]=="UU":
                    if not run_rows: run_ord += 1
                    run_rows.append(p)
                else:
                    if run_rows:
                        self._process_run(cur_read, run_ord, run_rows, frag_map, frag_cat, read_intervals)
                        run_rows=[]
            # EOF
            if run_rows:
                self._process_run(cur_read, run_ord, run_rows, frag_map, frag_cat, read_intervals)
            self._flush_read_fragments(frag_map, frag_cat)
        t1=time.time()

        # Save per-run CSV
        run_csv=os.path.join(self.outdir,"summaries","tortuosity_per_run.csv")
        with open(run_csv,"w",newline="") as fw:
            w=csv.writer(fw)
            w.writerow(["read_id","run_ord","run_len_steps","n_frags","n_cis_steps","n_trans_steps",
                        "cis_longest_block","n_cis_blocks","n_chroms","alt_rate","B_run","Dmax_bp",
                        "log10_Dmax","q_overlap_frac","q_overcov_idx","read_len"])
            for r in self.run_stats:
                w.writerow([r.read_id, r.run_ord, r.run_len_steps, r.n_frags,
                            r.n_cis_steps, r.n_trans_steps, r.cis_longest_block,
                            r.n_cis_blocks, r.n_chroms, r.alt_rate, r.brun,
                            r.dmax_bp, r.log10_dmax, r.q_overlap_frac, r.q_overcov_idx,
                            r.read_len])

        # Arrays
        def arr(name, x, dtype=None):
            np.save(os.path.join(self.outdir,"arrays",name),
                    np.array(x, dtype=dtype) if not isinstance(x,np.ndarray) else x)

        # Adj/Nonadj
        arr("adj_dists.npy", np.array(self.adj_dists, float))
        arr("nonadj_reservoir.npy", np.array(self.nadj_res_list, float))

        # Step pairs
        np.savez_compressed(os.path.join(self.outdir,"arrays","step_pairs_xy.npz"),
                            x=np.array(self.step_pairs_x, float),
                            y=np.array(self.step_pairs_y, float))
        np.savez_compressed(os.path.join(self.outdir,"arrays","step_pairs_xy_brun.npz"),
                            x=np.array(self.step_pairs_x, float),
                            y=np.array(self.step_pairs_y, float),
                            brun=np.array(self.step_pairs_b, float))

        # Tortuosity arrays
        brun=np.array([r.brun for r in self.run_stats], float)
        dmax=np.array([r.dmax_bp for r in self.run_stats], float)
        logd=np.array([r.log10_dmax for r in self.run_stats], float)
        rlen=np.array([r.run_len_steps for r in self.run_stats], int)
        cisblk=np.array([r.cis_longest_block for r in self.run_stats], int)
        cissteps=np.array([r.n_cis_steps for r in self.run_stats], int)
        transsteps=np.array([r.n_trans_steps for r in self.run_stats], int)
        alt=np.array([r.alt_rate for r in self.run_stats], float)
        q1=np.array([r.q_overlap_frac for r in self.run_stats], float)
        q2=np.array([r.q_overcov_idx for r in self.run_stats], float)
        readL=np.array([r.read_len for r in self.run_stats], int)

        arr("brun.npy", brun); arr("dmax_bp.npy", dmax); arr("log10_dmax.npy", logd)
        arr("run_len.npy", rlen); arr("cis_longest_block.npy", cisblk)
        arr("cis_steps.npy", cissteps); arr("trans_steps.npy", transsteps)
        arr("alt_rate.npy", alt); arr("q_overlap_frac.npy", q1); arr("q_overcov_idx.npy", q2)
        arr("read_len.npy", readL)

        # Fragment length arrays
        arr("fraglen_bin1_ALL.npy", self.frag_counts_all_bin1)
        arr("fraglen_bin3_ALL.npy", self.frag_counts_all_bin3)
        arr("fraglen_bin1_strata.npy", self.frag_counts_strata_bin1)
        arr("fraglen_bin3_strata.npy", self.frag_counts_strata_bin3)
        with open(os.path.join(self.outdir,"summaries","fraglen_meta.json"),"w") as f:
            json.dump({"frag_ecdf_max": self.frag_ecdf_max, "strata": [1,2,3,4]}, f, indent=2)

        # UU-run composition raw arrays
        arr("uur_readlen.npy", np.array(self.uur_readlens, int))
        arr("uur_runlen.npy", np.array(self.uur_runlens, int))

        # Contact spectra meta
        sep_meta={
            "adj_n": int(len(self.adj_dists)),
            "nonadj_reservoir_n": int(len(self.nadj_res_list)),
            "nonadj_seen_approx": int(self.nadj_seen),
            "small_hist_bin": self.small_hist_bin,
            "small_hist_max": self.small_hist_max
        }
        with open(os.path.join(self.outdir,"summaries","contact_spectra_meta.json"),"w") as f:
            json.dump(sep_meta, f, indent=2)

        # Quick tortuosity stats
        mask=np.isfinite(brun) & np.isfinite(logd)
        stats={
            "n_runs": int(brun.size),
            "pearson_r_brun_log10dmax": float(_pearsonr(logd[mask], brun[mask])) if mask.any() else float('nan'),
            "spearman_rho": float(_spearmanr(logd[mask], brun[mask])) if mask.any() else float('nan'),
            "elapsed_sec": round(t1-t0,2)
        }
        with open(os.path.join(self.outdir,"summaries","tortuosity_stats.json"),"w") as f:
            json.dump(stats, f, indent=2)

        # ---------- Per-TR plots ----------
        _, plt = _mpl()

        # Tortuosity hexbin by CIS-block
        if brun.size>0:
            dcat = np.array([self._cisblock_category(c) for c in cisblk], int)
            for cat in [1,2,3,4]:
                sel=(dcat==cat) & np.isfinite(brun) & np.isfinite(logd)
                if sel.any():
                    fig=plt.figure(figsize=(8.2,6.0)); ax=fig.gca()
                    hb=ax.hexbin(logd[sel], brun[sel], gridsize=60, bins='log')
                    fig.colorbar(hb, ax=ax).set_label("log10(count)")
                    ax.set_xlabel("log10(Max within-run cis separation, bp)")
                    ax.set_ylabel("Run-level tortuosity, B_run")
                    ax.set_title(f"{self.label}: Dmax vs B_run (CIS-block={cat if cat<4 else '≥4'})  N={sel.sum()}")
                    fig.tight_layout()
                    fig.savefig(os.path.join(self.outdir,"plots", f"tortuosity_hexbin_cisblock_{cat}.pdf"))
                    plt.close(fig)

        # Fragment length: hist (bin=3) and ECDFs
        max_plot=250
        bins3=np.arange(0, max_plot+1, 3)
        counts3_all = np.add.reduceat(self.frag_counts_all_bin1[:max_plot+1], np.arange(0, max_plot+1, 3))
        fig=plt.figure(figsize=(8.0,5.2)); ax=fig.gca()
        ax.bar(bins3, counts3_all, width=3, align="edge", edgecolor="black", linewidth=0.2)
        ax.set_xlim(0, max_plot); ax.set_xlabel("Fragment length (bp)"); ax.set_ylabel("Count")
        ax.set_title(f"{self.label}: U fragment length histogram (bin=3 bp)")
        fig.tight_layout(); fig.savefig(os.path.join(self.outdir,"plots","fraglen_hist_bin3_ALL.pdf")); plt.close(fig)
        # ECDFs
        xlim=self.frag_ecdf_max; xs=np.arange(0, xlim+1)
        def ecdf_from_counts(cnt):
            c=np.cumsum(cnt); cmax=c[-1] if c.size else 0
            return (xs, c/cmax) if cmax>0 else (xs, np.zeros_like(xs, float))
        fig=plt.figure(figsize=(8.0,5.2)); ax=fig.gca()
        x_all, y_all = ecdf_from_counts(self.frag_counts_all_bin1[:xlim+1])
        ax.step(x_all, y_all, where="post", label="ALL")
        names={1:"cis1",2:"cis2",3:"cis3",4:"cis≥4"}
        for idx,cat in enumerate([1,2,3,4]):
            x,y = ecdf_from_counts(self.frag_counts_strata_bin1[idx,:xlim+1])
            if (y>0).any(): ax.step(x,y,where="post",label=names[cat])
        ax.set_xlim(0, xlim); ax.set_ylim(0,1)
        ax.set_xlabel("Fragment length (bp)"); ax.set_ylabel("ECDF")
        ax.set_title(f"{self.label}: U fragment-length ECDFs (shared x-limits)")
        ax.legend(frameon=False)
        fig.tight_layout(); fig.savefig(os.path.join(self.outdir,"plots","fraglen_ecdf_ALL_and_strata.pdf")); plt.close(fig)

        # UU-run composition by read-length deciles (per TR)
        if len(self.uur_runlens)>0:
            readlens=np.array(self.uur_readlens, int)
            runlens =np.array(self.uur_runlens, int)
            edges=np.quantile(readlens, np.linspace(0,1,11))
            for i in range(1,11):
                if edges[i]<=edges[i-1]: edges[i]=edges[i-1]+1
            dec=np.clip(np.searchsorted(edges, readlens, side="right")-1, 0, 9)
            S_MAX=10
            cats=[str(s) for s in range(1,S_MAX+1)]+[f"{S_MAX+1}+"]
            counts=np.zeros((10,len(cats)), np.int64)
            for d in range(10):
                svals=runlens[dec==d]
                for s in svals:
                    if 1<=s<=S_MAX: counts[d,s-1]+=1
                    elif s>=S_MAX+1: counts[d,-1]+=1
            pct=np.where(counts.sum(1,keepdims=True)>0, counts / counts.sum(1,keepdims=True)*100.0, 0.0)
            np.save(os.path.join(self.outdir,"arrays","uu_run_counts.npy"), counts)
            np.save(os.path.join(self.outdir,"arrays","uu_run_pct.npy"), pct)
            with open(os.path.join(self.outdir,"summaries","uu_run_composition.csv"),"w",newline="") as fw:
                w=csv.writer(fw); w.writerow(["decile","category","count","percent","len_min","len_max"])
                for d in range(10):
                    for k in range(len(cats)):
                        w.writerow([d+1, cats[k], int(counts[d,k]), float(pct[d,k]), int(edges[d]), int(edges[d+1]-1)])
            x=np.arange(1,11); bottom=np.zeros(10,float)
            fig=plt.figure(figsize=(10.5,6.2)); ax=fig.gca()
            for k in range(len(cats)):
                ax.bar(x, pct[:,k], bottom=bottom, edgecolor="black", linewidth=0.3)
                bottom+=pct[:,k]
            ax.set_xticks(x, [f"D{d}" for d in x])
            ax.set_ylim(0,100); ax.set_xlabel("Read-length decile"); ax.set_ylabel("Percent of UU runs")
            ax.set_title(f"{self.label}: UU-run length composition by read-length decile")
            fig.tight_layout(); fig.savefig(os.path.join(self.outdir,"plots","uu_run_composition.pdf")); plt.close(fig)

        # ECDFs for cis separations up to 1e9 (log scale)
        adj=np.array(self.adj_dists, float)
        nonadj=np.array(self.nadj_res_list, float)
        def save_ecdf(name, a):
            a=np.sort(a); y=np.arange(1,a.size+1)/a.size if a.size else np.array([])
            np.save(os.path.join(self.outdir,"arrays",f"{name}_ecdf_x.npy"), a)
            np.save(os.path.join(self.outdir,"arrays",f"{name}_ecdf_y.npy"), y)
        save_ecdf("adj", adj); save_ecdf("nonadj", nonadj)
        fig=plt.figure(figsize=(8.5,5.6)); ax=fig.gca()
        if adj.size: ax.step(np.sort(adj), np.arange(1,adj.size+1)/adj.size, where="post", label=f"Adjacent (N={adj.size:,})")
        if nonadj.size: ax.step(np.sort(nonadj), np.arange(1,nonadj.size+1)/nonadj.size, where="post", label=f"Non-adj (sample N={nonadj.size:,})")
        ax.set_xscale("log"); ax.set_xlim(1, 1e9); ax.set_ylim(0,1)
        ax.set_xlabel("Cis separation (bp, log scale)"); ax.set_ylabel("ECDF")
        ax.set_title(f"{self.label}: ECDF of cis separations (adj vs non-adj)")
        ax.legend(frameon=False)
        fig.tight_layout(); fig.savefig(os.path.join(self.outdir,"plots","cis_separation_ecdf_log.pdf")); plt.close(fig)

        # Small-scale histograms
        edges=np.arange(0, self.small_hist_max+self.small_hist_bin, self.small_hist_bin)
        def hist_plot(arr, tag):
            H, _ = np.histogram(arr, bins=edges)
            np.save(os.path.join(self.outdir,"arrays", f"{tag}_hist_counts.npy"), H)
            fig=plt.figure(figsize=(10.0,5.8)); ax=fig.gca()
            ax.bar(edges[:-1], H, width=self.small_hist_bin, align="edge", edgecolor="black", linewidth=0.2)
            ax.set_xlim(0, self.small_hist_max); ax.set_xlabel("Cis separation (bp)")
            ax.set_ylabel("Count"); ax.set_title(f"{self.label}: {tag} cis separations (0–{self.small_hist_max} bp, bin={self.small_hist_bin})")
            fig.tight_layout(); fig.savefig(os.path.join(self.outdir,"plots", f"{tag}_hist_smallscale.pdf")); plt.close(fig)
        if adj.size: hist_plot(adj, "adjacent")
        if nonadj.size: hist_plot(nonadj, "nonadjacent")

        with open(os.path.join(self.outdir,"metadata","done.json"),"w") as f:
            json.dump({"elapsed_sec": round(t1-t0,2)}, f, indent=2)

# ---------------- Merger ----------------
class Merger:
    def __init__(self, out_merged: str, trs: List[str], labels: List[str]):
        self.outdir=out_merged
        self.trs=trs
        self.labels=labels
        _ensure_dirs(self.outdir, os.path.join(self.outdir,"arrays"),
                     os.path.join(self.outdir,"summaries"),
                     os.path.join(self.outdir,"metadata"),
                     os.path.join(self.outdir,"plots"))
        with open(os.path.join(self.outdir,"metadata","merge_meta.json"),"w") as f:
            json.dump({"labels": labels, "sources": trs}, f, indent=2)

    def _load(self, t: str, rel: str):
        p=os.path.join(t, "arrays", rel)
        if os.path.exists(p): return np.load(p, allow_pickle=False)
        return None

    def _load_npz(self, t: str, rel: str):
        p=os.path.join(t, "arrays", rel)
        if os.path.exists(p): return np.load(p)
        return None

    def run(self):
        # Pool per-run arrays
        arrs={}
        for name in ["brun.npy","dmax_bp.npy","log10_dmax.npy","run_len.npy","cis_longest_block.npy",
                     "cis_steps.npy","trans_steps.npy","alt_rate.npy","q_overlap_frac.npy","q_overcov_idx.npy",
                     "read_len.npy"]:
            arrs[name]=[]
        # Step pairs with brun
        pairs_b=[]
        # Adj/non-adj
        adj_all=[]; nonadj_all=[]
        # UU-run composition inputs
        uur_readlen=[]; uur_runlen=[]
        # Fragment lengths
        frag_meta=None
        frag_bin1_ALL=None; frag_bin3_ALL=None
        frag_bin1_strata=None; frag_bin3_strata=None

        for tr in self.trs:
            for k in arrs.keys():
                x=self._load(tr, k)
                if x is not None: arrs[k].append(x)
            spb=self._load_npz(tr, "step_pairs_xy_brun.npz")
            if spb is not None:
                pairs_b.append((spb["x"], spb["y"], spb["brun"]))
            a=self._load(tr, "adj_dists.npy")
            n=self._load(tr, "nonadj_reservoir.npy")
            if a is not None: adj_all.append(a)
            if n is not None: nonadj_all.append(n)
            rl=self._load(tr, "uur_readlen.npy")
            ul=self._load(tr, "uur_runlen.npy")
            if rl is not None: uur_readlen.append(rl)
            if ul is not None: uur_runlen.append(ul)
            # fragments
            fm_path=os.path.join(tr,"summaries","fraglen_meta.json")
            if frag_meta is None and os.path.exists(fm_path):
                frag_meta=json.load(open(fm_path))
            b1_all=self._load(tr, "fraglen_bin1_ALL.npy")
            b3_all=self._load(tr, "fraglen_bin3_ALL.npy")
            b1_str=self._load(tr, "fraglen_bin1_strata.npy")
            b3_str=self._load(tr, "fraglen_bin3_strata.npy")
            if b1_all is not None:
                frag_bin1_ALL = b1_all if frag_bin1_ALL is None else frag_bin1_ALL + b1_all
            if b3_all is not None:
                frag_bin3_ALL = b3_all if frag_bin3_ALL is None else frag_bin3_ALL + b3_all
            if b1_str is not None:
                frag_bin1_strata = b1_str if frag_bin1_strata is None else frag_bin1_strata + b1_str
            if b3_str is not None:
                frag_bin3_strata = b3_str if frag_bin3_strata is None else frag_bin3_strata + b3_str

        # Save pooled per-run arrays
        def cat(name):
            xs=arrs[name]
            if not xs: return np.array([])
            return np.concatenate(xs)
        pooled={}
        for k in arrs.keys():
            pooled[k]=cat(k)
            np.save(os.path.join(self.outdir,"arrays",k), pooled[k])

        # Save pooled fragments
        if frag_bin1_ALL is not None:
            np.save(os.path.join(self.outdir,"arrays","fraglen_bin1_ALL.npy"), frag_bin1_ALL)
        if frag_bin3_ALL is not None:
            np.save(os.path.join(self.outdir,"arrays","fraglen_bin3_ALL.npy"), frag_bin3_ALL)
        if frag_bin1_strata is not None:
            np.save(os.path.join(self.outdir,"arrays","fraglen_bin1_strata.npy"), frag_bin1_strata)
        if frag_bin3_strata is not None:
            np.save(os.path.join(self.outdir,"arrays","fraglen_bin3_strata.npy"), frag_bin3_strata)
        if frag_meta is not None:
            with open(os.path.join(self.outdir,"summaries","fraglen_meta.json"),"w") as f:
                json.dump(frag_meta, f, indent=2)

        # Save pooled adj/nonadj and ECDFs
        adj = np.concatenate(adj_all) if adj_all else np.array([],float)
        nonadj = np.concatenate(nonadj_all) if nonadj_all else np.array([],float)
        np.save(os.path.join(self.outdir,"arrays","adj_dists.npy"), adj)
        np.save(os.path.join(self.outdir,"arrays","nonadj_reservoir.npy"), nonadj)
        def save_ecdf(name, a):
            a=np.sort(a); y=np.arange(1,a.size+1)/a.size if a.size else np.array([])
            np.save(os.path.join(self.outdir,"arrays",f"{name}_ecdf_x.npy"), a)
            np.save(os.path.join(self.outdir,"arrays",f"{name}_ecdf_y.npy"), y)
        save_ecdf("adj", adj); save_ecdf("nonadj", nonadj)

        # Save pooled step pairs with brun
        if pairs_b:
            x=np.concatenate([p[0] for p in pairs_b]); y=np.concatenate([p[1] for p in pairs_b]); b=np.concatenate([p[2] for p in pairs_b])
            np.savez_compressed(os.path.join(self.outdir,"arrays","step_pairs_xy_brun.npz"), x=x, y=y, brun=b)
        else:
            np.savez_compressed(os.path.join(self.outdir,"arrays","step_pairs_xy_brun.npz"),
                                x=np.array([]), y=np.array([]), brun=np.array([]))

        # UU-run composition merged
        if uur_readlen and uur_runlen:
            readlens=np.concatenate(uur_readlen); runlens=np.concatenate(uur_runlen)
            edges=np.quantile(readlens, np.linspace(0,1,11))
            for i in range(1,11):
                if edges[i]<=edges[i-1]: edges[i]=edges[i-1]+1
            dec=np.clip(np.searchsorted(edges, readlens, side="right")-1, 0, 9)
            S_MAX=10
            cats=[str(s) for s in range(1,S_MAX+1)]+[f"{S_MAX+1}+"]
            counts=np.zeros((10,len(cats)), np.int64)
            for d in range(10):
                svals=runlens[dec==d]
                for s in svals:
                    if 1<=s<=S_MAX: counts[d,s-1]+=1
                    elif s>=S_MAX+1: counts[d,-1]+=1
            pct=np.where(counts.sum(1,keepdims=True)>0, counts / counts.sum(1,keepdims=True)*100.0, 0.0)
            np.save(os.path.join(self.outdir,"arrays","uu_run_counts.npy"), counts)
            np.save(os.path.join(self.outdir,"arrays","uu_run_pct.npy"), pct)
            with open(os.path.join(self.outdir,"summaries","uu_run_composition.csv"),"w",newline="") as fw:
                w=csv.writer(fw); w.writerow(["decile","category","count","percent","len_min","len_max"])
                for d in range(10):
                    for k in range(len(cats)):
                        w.writerow([d+1, cats[k], int(counts[d,k]), float(pct[d,k]), int(edges[d]), int(edges[d+1]-1)])

        # Brun×Dmax (10×10) query-overlap tests
        brun=pooled["brun.npy"]; dmax=pooled["dmax_bp.npy"]; q1=pooled["q_overlap_frac.npy"]
        mask=np.isfinite(brun) & np.isfinite(dmax) & np.isfinite(q1)
        if mask.any():
            B=brun[mask]; D=dmax[mask]; Q=q1[mask]
            Bedges=np.quantile(B, np.linspace(0,1,11))
            Dpos=D[D>0]
            Dedges=np.quantile(Dpos, np.linspace(0,1,11)) if Dpos.size else np.linspace(0,1,11)
            with open(os.path.join(self.outdir,"summaries","decile_edges.json"),"w") as f:
                json.dump({"B_run_edges": Bedges.tolist(), "Dmax_bp_edges": Dedges.tolist()}, f, indent=2)
            Bi=np.clip(np.searchsorted(Bedges, B, side="right")-1, 0, 9)
            Di=np.clip(np.searchsorted(Dedges, D, side="right")-1, 0, 9)
            Qg=Q.copy()
            out_rows=[]; pvals=[]
            for i in range(10):
                for j in range(10):
                    sel=(Bi==i)&(Di==j)
                    q=Q[sel]
                    med=float(np.median(q)) if q.size else float('nan')
                    q25=float(np.quantile(q,0.25)) if q.size else float('nan')
                    q75=float(np.quantile(q,0.75)) if q.size else float('nan')
                    p = _mannwhitney_u_p(q, Qg) if q.size and Qg.size else float('nan')
                    d = _cliffs_delta(q, Qg) if q.size and Qg.size else float('nan')
                    out_rows.append([i+1,j+1,int(q.size),med,q25,q75,p,d,Bedges[i],Bedges[i+1],Dedges[j],Dedges[j+1]])
                    pvals.append(p)
            qs=_bh_fdr([p if np.isfinite(p) else 1.0 for p in pvals])
            with open(os.path.join(self.outdir,"summaries","grid_q1_tests.csv"),"w",newline="") as fw:
                w=csv.writer(fw)
                w.writerow(["B_dec","D_dec","N","q1_median","q1_q25","q1_q75","p_MWU","cliffs_delta",
                            "B_lo","B_hi","D_lo_bp","D_hi_bp","q_BH"])
                for row, qv in zip(out_rows, qs):
                    w.writerow(row+[qv])

        # ---------- Merged plots ----------
        _, plt=_mpl()

        # Tortuosity hexbin by CIS-block (merged)
        if pooled["brun.npy"].size>0:
            B=pooled["brun.npy"]; Ld=pooled["log10_dmax.npy"]; C=pooled["cis_longest_block.npy"]
            cats=np.array([1 if c<=1 else 2 if c==2 else 3 if c==3 else 4 for c in C], int)
            for cat in [1,2,3,4]:
                sel=(cats==cat) & np.isfinite(B) & np.isfinite(Ld)
                if sel.any():
                    fig=plt.figure(figsize=(8.2,6.0)); ax=fig.gca()
                    hb=ax.hexbin(Ld[sel], B[sel], gridsize=60, bins='log')
                    fig.colorbar(hb, ax=ax).set_label("log10(count)")
                    ax.set_xlabel("log10(Max within-run cis separation, bp)")
                    ax.set_ylabel("Run-level tortuosity, B_run")
                    ax.set_title(f"Merged: Dmax vs B_run (CIS-block={cat if cat<4 else '≥4'})  N={sel.sum()}")
                    fig.tight_layout()
                    fig.savefig(os.path.join(self.outdir,"plots", f"tortuosity_hexbin_cisblock_{cat}.pdf"))
                    plt.close(fig)

        # Step-pair hexbin — ALL and by B_run quartiles (merged)
        spb=np.load(os.path.join(self.outdir,"arrays","step_pairs_xy_brun.npz"))
        x=spb["x"]; y=spb["y"]; b=spb["brun"]
        if x.size>0:
            lim=np.nanpercentile(np.concatenate([x,y]), 99) if x.size else 1.0
            lim=max(lim, 1.0)
            # ALL
            fig=plt.figure(figsize=(6.4,6.4)); ax=fig.gca()
            hb=ax.hexbin(x,y,gridsize=70, bins='log', cmap="inferno")
            fig.colorbar(hb, ax=ax).set_label("log10(count)")
            ax.set_xlim(0,lim); ax.set_ylim(0,lim); ax.set_aspect('equal', 'box')
            ax.set_xlabel("d_i (bp)"); ax.set_ylabel("d_{i+1} (bp)")
            ax.set_title(f"Merged: Adjacent cis step pairs  N={x.size:,}")
            fig.tight_layout(); fig.savefig(os.path.join(self.outdir,"plots","step_pairs_hexbin_ALL.pdf")); plt.close(fig)
            # Quartiles
            mask=np.isfinite(b)
            if mask.any():
                qedges=np.quantile(b[mask], [0.25,0.5,0.75])
                labels=["Q1 (≤Q1)","Q2","Q3","Q4 (≥Q3)"]
                bounds=[(-np.inf,qedges[0]), (qedges[0],qedges[1]), (qedges[1],qedges[2]), (qedges[2],np.inf)]
                for qi,(lo,hi) in enumerate(bounds, start=1):
                    sel=(b>=lo) & (b<(hi if qi<4 else hi))
                    xs=x[sel]; ys=y[sel]
                    if xs.size==0: continue
                    fig=plt.figure(figsize=(6.4,6.4)); ax=fig.gca()
                    hb=ax.hexbin(xs,ys,gridsize=70, bins='log', cmap="inferno")
                    fig.colorbar(hb, ax=ax).set_label("log10(count)")
                    ax.set_xlim(0,lim); ax.set_ylim(0,lim); ax.set_aspect('equal','box')
                    ax.set_xlabel("d_i (bp)"); ax.set_ylabel("d_{i+1} (bp)")
                    ax.set_title(f"Merged: Step pairs by B_run quartile — {labels[qi-1]}  N={xs.size:,}")
                    fig.tight_layout(); fig.savefig(os.path.join(self.outdir,"plots", f"step_pairs_hexbin_BQ{qi}.pdf")); plt.close(fig)

        # UU-run composition merged
        if os.path.exists(os.path.join(self.outdir,"arrays","uu_run_pct.npy")):
            pct=np.load(os.path.join(self.outdir,"arrays","uu_run_pct.npy"))
            x=np.arange(1,11); bottom=np.zeros(10,float)
            fig=plt.figure(figsize=(10.5,6.2)); ax=fig.gca()
            for k in range(pct.shape[1]):
                ax.bar(x, pct[:,k], bottom=bottom, edgecolor="black", linewidth=0.3)
                bottom+=pct[:,k]
            ax.set_xticks(x, [f"D{d}" for d in x]); ax.set_ylim(0,100)
            ax.set_xlabel("Read-length decile (short → long)"); ax.set_ylabel("Percent of UU runs")
            ax.set_title("Merged: UU-run length composition by read-length decile")
            fig.tight_layout(); fig.savefig(os.path.join(self.outdir,"plots","uu_run_composition_merged.pdf")); plt.close(fig)

        # ECDF merged (adj/non-adj)
        if adj.size or nonadj.size:
            fig=plt.figure(figsize=(8.5,5.6)); ax=fig.gca()
            if adj.size: ax.step(np.sort(adj), np.arange(1,adj.size+1)/adj.size, where="post", label=f"Adjacent (N={adj.size:,})")
            if nonadj.size: ax.step(np.sort(nonadj), np.arange(1,nonadj.size+1)/nonadj.size, where="post", label=f"Non-adj (sample N={nonadj.size:,})")
            ax.set_xscale("log"); ax.set_xlim(1,1e9); ax.set_ylim(0,1)
            ax.set_xlabel("Cis separation (bp, log scale)"); ax.set_ylabel("ECDF")
            ax.set_title("Merged: ECDF of cis separations (adj vs non-adj)")
            ax.legend(frameon=False)
            fig.tight_layout(); fig.savefig(os.path.join(self.outdir,"plots","cis_separation_ecdf_log_merged.pdf")); plt.close(fig)

# ---------------- CLI ----------------
def main():
    ap=argparse.ArgumentParser(
        description="CADWALK-CLI v7: streaming analytics for PacBio CADwalk (.pairs.gz), TR-specific and merged outputs.",
        formatter_class=argparse.ArgumentDefaultsHelpFormatter)
    ap.add_argument("-o","--outdir", required=True, help="Output parent directory.")
    ap.add_argument("--pairs", nargs="+", required=True, help="One or more parse2 .pairs.gz inputs.")
    ap.add_argument("--label", nargs="+", required=False, default=None, help="Labels per input (same length as --pairs).")
    ap.add_argument("--small-hist-bin", type=int, default=5, help="Small-scale histogram bin (bp).")
    ap.add_argument("--small-hist-max", type=int, default=2000, help="Small-scale histogram max (bp).")
    ap.add_argument("--nonadj-reservoir", type=int, default=2_000_000, help="Reservoir size for non-adj cis ECDF.")
    ap.add_argument("--frag-ecdf-max", type=int, default=1000, help="Fragment length ECDF max (bp) for bin1 arrays.")
    ap.add_argument("--seed", type=int, default=17, help="Random seed.")
    args=ap.parse_args()

    ts=_now_ts_compact()
    root=os.path.join(args.outdir, f"cadwalk_v7_{ts}")
    _ensure_dirs(root)

    if args.label is None or len(args.label)!=len(args.pairs):
        labels=[f"rep{i+1}" for i in range(len(args.pairs))]
    else:
        labels=args.label

    # Per TR
    tr_dirs=[]
    for pth, lab in zip(args.pairs, labels):
        tr_out=os.path.join(root, lab)
        _ensure_dirs(tr_out)
        proc=TRProcessor(
            pairs_path=pth, label=lab, outdir=tr_out,
            small_hist_bin=args.small_hist_bin, small_hist_max=args.small_hist_max,
            nonadj_reservoir=args.nonadj_reservoir, frag_ecdf_max=args.frag_ecdf_max,
            seed=args.seed
        )
        proc.run()
        tr_dirs.append(tr_out)

    # Merged
    merged_dir=os.path.join(root, "merged")
    Merger(merged_dir, tr_dirs, labels).run()

    print(f"[OK] All outputs written under: {root}")

if __name__=="__main__":
    main()
```
