## Supplementary material for "CAD-C: An engineered nuclease enables repair-free *in situ* proximity ligation and nucleosome-resolution chromosome walks in human cells": Custom code: v9cadpack_transjumps_and_uniqchros_by_runLen_v4_20251021.py

```
#!/usr/bin/env python3
# -*- coding: utf-8 -*-
"""
v9cadpack_trans_and_uniqchr_by_runLen_palette_v3_20251021.py
------------------------------------------------------------
Changes vs v2:
- X label -> 'uninterrupted UU walk steps (=run)' (main + inset)
- Y label -> 'Count'
- Plot is capped at L <= 13 (run steps)
- Inset x-axis uses integer ticks only
- Also emits log10(count) versions (semilog base-10 ticks) with '_log10' suffix

From a v9 CADpack NPZ, produce two stacked-bar figures by run length L
(= uninterrupted UU walk steps). Stacks reflect:
  1) # of trans‑chromosomal jumps per run
  2) # of unique chromosomes explored per run

Both figures include an inset with per-length *normalized* fractions.
Palette control via --palette. Output: two PDFs (+ two log10 PDFs). Optional CSV.
"""

import argparse
import os
import numpy as np
import matplotlib
matplotlib.use("Agg")
# Editable text in PDF
matplotlib.rcParams["pdf.fonttype"] = 42
matplotlib.rcParams["ps.fonttype"]  = 42

import matplotlib.pyplot as plt
from matplotlib.patches import Patch
from mpl_toolkits.axes_grid1.inset_locator import inset_axes
from matplotlib.ticker import FuncFormatter, MaxNLocator, LogLocator, NullFormatter


def _fmt_int(x, _pos=None):
    try:
        return f"{int(round(x)):,}"
    except Exception:
        return str(x)


def _get_discrete_colors(K: int, palette: str):
    """
    Return a list of K RGBA colors from the requested palette.
    Supports seaborn palettes if installed; otherwise uses matplotlib colormaps.
    """
    if K <= 0:
        return []
    try:
        import seaborn as sns  # optional
        try:
            return list(sns.color_palette(palette, n_colors=K))
        except Exception:
            pass
    except Exception:
        pass

    try:
        cmap = plt.get_cmap(palette, K)
    except Exception:
        cmap = plt.get_cmap("GnBu", K)
    return [cmap(i) for i in range(K)]


def _tally_metrics(npz_path: str):
    """
    Return dicts keyed by run length L (steps):
       counts_trans[L][T], counts_uniq[L][U],
       L_vals, T_max, U_max, n_runs.
    """
    Z   = np.load(npz_path, allow_pickle=False, mmap_mode=None)
    run_L = Z["run_L"].astype(np.int64)
    rfs   = Z["run_first_step_row"].astype(np.int64)
    rns   = Z["run_n_steps"].astype(np.int64)
    c1    = Z["chrom1_id"].astype(np.int32)
    c2    = Z["chrom2_id"].astype(np.int32)

    L_list, T_list, U_list = [], [], []

    for r in range(run_L.size):
        L = int(run_L[r])           # uninterrupted UU steps (= run length)
        n = int(rns[r])
        if L <= 0 or n != L:
            continue
        s0 = int(rfs[r])
        sl = slice(s0, s0 + n)

        c1r = c1[sl]
        c2r = c2[sl]

        # Metric 1: # of trans steps
        T = int(np.sum(c1r != c2r))

        # Metric 2: # of unique chromosomes explored (ignore negatives)
        uniq = np.unique(np.concatenate([c1r[c1r >= 0], c2r[c2r >= 0]]))
        U = int(uniq.size)

        L_list.append(L); T_list.append(T); U_list.append(U)

    if not L_list:
        raise SystemExit("[ERROR] No valid runs found in NPZ.")

    L_arr = np.asarray(L_list, dtype=int)
    T_arr = np.asarray(T_list, dtype=int)
    U_arr = np.asarray(U_list, dtype=int)

    L_vals = np.unique(L_arr)
    T_max  = int(T_arr.max()) if T_arr.size else 0
    U_max  = int(U_arr.max()) if U_arr.size else 1

    counts_trans = {L: np.zeros(T_max + 1, dtype=np.int64) for L in L_vals}
    counts_uniq  = {L: np.zeros(U_max + 1, dtype=np.int64) for L in L_vals}

    for L, T, U in zip(L_arr, T_arr, U_arr):
        if 0 <= T <= T_max: counts_trans[L][T] += 1
        if 0 <= U <= U_max: counts_uniq[L][U]  += 1

    n_runs = int(L_arr.size)
    return counts_trans, counts_uniq, L_vals, T_max, U_max, n_runs


def _semilog10_axis(ax):
    """
    Configure Y as semilog (base 10) but label the major ticks as log10(count)
    i.e., 0,1,2,... where tick positions are 10^k.
    """
    ax.set_yscale("log", base=10, nonpositive="clip")
    ax.set_ylim(bottom=1)  # avoid log(0)
    ax.yaxis.set_major_locator(LogLocator(base=10))
    ax.yaxis.set_minor_locator(LogLocator(base=10, subs=np.arange(2, 10) * 0.1))
    # Format major ticks as exponents k = log10(value)
    def _fmt_log10(v, _pos=None):
        try:
            if v <= 0:
                return ""
            return f"{int(np.round(np.log10(v)))}"
        except Exception:
            return ""
    ax.yaxis.set_major_formatter(FuncFormatter(_fmt_log10))
    ax.yaxis.set_minor_formatter(NullFormatter())
    ax.set_ylabel("log10(Count)", fontsize=14)


def _plot_stacked(counts, L_vals, K_max, title, legend_title, out_pdf,
                  palette="GnBu", max_run_len=13, make_log10=False):
    """
    Generic stacked bar plot by run length L with normalized inset.
    counts[L][k] for k=0..K_max. Only L in [min(L_vals), min(max(L_vals), max_run_len)]
    are shown.
    """
    L_vals = np.sort(L_vals)
    L_min  = int(max(1, L_vals.min()))
    L_max  = int(min(L_vals.max(), max_run_len))
    L_full = np.arange(L_min, L_max + 1, dtype=int)

    # Build matrix M[x, k] with x indexed by L_full
    M = np.zeros((L_full.size, K_max + 1), dtype=np.int64)
    for i, L in enumerate(L_full):
        if L in counts:
            row = counts[L]
            M[i, :row.size] = row

    colors = _get_discrete_colors(K_max + 1, palette)

    fig = plt.figure(figsize=(8.6, 8.8), facecolor="white")
    ax  = fig.add_subplot(111)

    # Absolute stacked bars
    bottoms = np.zeros(L_full.size, dtype=float)
    for k in range(K_max + 1):
        vals = M[:, k].astype(float)
        ax.bar(L_full, vals, width=0.85, bottom=bottoms, color=colors[k],
               edgecolor="none", align="center")
        bottoms += vals

    # Inset: normalized fractions (per-length columns sum to 1)
    ax_in = inset_axes(ax, width="54%", height="42%", loc="upper center",
                       bbox_to_anchor=(0.13, 0.10, 0.74, 0.74),
                       bbox_transform=ax.transAxes, borderpad=0.6)
    col_sums = M.sum(axis=1).astype(float)
    col_sums[col_sums == 0] = 1.0
    M_norm = (M.T / col_sums).T
    bottoms = np.zeros(L_full.size, dtype=float)
    for k in range(K_max + 1):
        ax_in.bar(L_full, M_norm[:, k], width=0.85, bottom=bottoms,
                  color=colors[k], edgecolor="none", align="center")
        bottoms += M_norm[:, k]
    ax_in.set_ylim(0, 1.0)
    ax_in.set_yticks([0, 0.2, 0.4, 0.6, 0.8, 1.0])
    ax_in.set_ylabel("Frequency Normalized Fraction of Walks", fontsize=9)

    # ---- labels and tick control ----
    xlab = "uninterrupted UU walk steps (=run)"
    ax.set_xlabel(xlab, fontsize=14)
    ax_in.set_xlabel(xlab, fontsize=9)

    # integer ticks only
    ax.set_xticks(L_full)
    ax.xaxis.set_major_locator(MaxNLocator(integer=True))
    ax_in.set_xticks(L_full)
    ax_in.xaxis.set_major_locator(MaxNLocator(integer=True))

    # Y axis scaling/format
    if make_log10:
        _semilog10_axis(ax)
        ax.grid(axis="y", which="both", linestyle=":", alpha=0.35)
    else:
        ax.set_ylabel("Count", fontsize=14)
        ax.yaxis.set_major_formatter(FuncFormatter(_fmt_int))
        ax.grid(axis="y", linestyle=":", alpha=0.35)

    for s in ax.spines.values():
        s.set_linewidth(1.2)

    # Legend
    patches = [Patch(facecolor=colors[k], edgecolor="none", label=str(k))
               for k in range(K_max + 1)]
    leg = ax.legend(handles=patches, title=legend_title,
                    loc="center left", bbox_to_anchor=(1.02, 0.5),
                    frameon=False, borderaxespad=0.0)
    plt.setp(leg.get_title(), fontsize=11)

    ax.set_title(title, fontsize=18, weight="bold", pad=12)

    fig.tight_layout()
    fig.savefig(out_pdf, dpi=300)
    plt.close(fig)
    print(f"[OK] Wrote: {out_pdf}")


def _write_csvs(counts_trans, counts_uniq, L_vals, out_prefix: str):
    import csv, os
    os.makedirs(os.path.dirname(out_prefix), exist_ok=True)
    trans_csv = out_prefix + "_trans_counts.csv"
    uniq_csv  = out_prefix + "_uniqchr_counts.csv"

    with open(trans_csv, "w", newline="") as f:
        w = csv.writer(f); w.writerow(["run_length_L", "trans_jumps", "count"])
        for L in sorted(L_vals):
            row = counts_trans.get(L, None)
            if row is None: continue
            for T, cnt in enumerate(row):
                w.writerow([L, T, int(cnt)])

    with open(uniq_csv, "w", newline="") as f:
        w = csv.writer(f); w.writerow(["run_length_L", "unique_chromosomes", "count"])
        for L in sorted(counts_uniq.keys()):
            row = counts_uniq[L]
            for U, cnt in enumerate(row):
                w.writerow([L, U, int(cnt)])

    print(f"[OK] CSVs: {trans_csv} , {uniq_csv}")


def _with_log_suffix(path: str) -> str:
    root, ext = os.path.splitext(path)
    if not ext:
        ext = ".pdf"
    return f"{root}_log10{ext}"


def main():
    ap = argparse.ArgumentParser(
        description="v9 CADpack: trans-jumps & unique-chromosomes by run length (palette-aware).",
        formatter_class=argparse.ArgumentDefaultsHelpFormatter
    )
    ap.add_argument("--npz", required=True, help="Path to v9 CADpack NPZ")
    ap.add_argument("--out-trans", required=True, help="Output PDF for trans-jumps figure")
    ap.add_argument("--out-uniq",  required=True, help="Output PDF for unique-chromosomes figure")
    ap.add_argument("--csv-prefix", default=None, help="If set, write CSVs with this prefix (no extension)")
    ap.add_argument("--palette", default="GnBu",
                    help="Color palette name. Examples: 'mako_r', 'mako', 'GnBu', 'Blues', 'viridis'. "
                         "If using 'mako'/'mako_r', seaborn must be installed.")
    args = ap.parse_args()

    counts_trans, counts_uniq, L_vals, T_max, U_max, n_runs = _tally_metrics(args.npz)

    # Linear scale (Count)
    _plot_stacked(
        counts_trans, L_vals, T_max,
        title="trans‑Chromosomal Jumps per Run Length",
        legend_title="Number of\ntrans-chr\nJumps",
        out_pdf=args.out_trans,
        palette=args.palette,
        max_run_len=13,
        make_log10=False
    )

    _plot_stacked(
        counts_uniq, L_vals, U_max,
        title="Unique Chromosomes Explored per Run Length",
        legend_title="Number of\nUnique\nChromosomes",
        out_pdf=args.out_uniq,
        palette=args.palette,
        max_run_len=13,
        make_log10=False
    )

    # Log10(count) views with semilog ticks
    _plot_stacked(
        counts_trans, L_vals, T_max,
        title="trans‑Chromosomal Jumps per Run Length (log10 view)",
        legend_title="Number of\ntrans-chr\nJumps",
        out_pdf=_with_log_suffix(args.out_trans),
        palette=args.palette,
        max_run_len=13,
        make_log10=True
    )

    _plot_stacked(
        counts_uniq, L_vals, U_max,
        title="Unique Chromosomes per Run Length (log10 view)",
        legend_title="Number of\nUnique\nChromosomes",
        out_pdf=_with_log_suffix(args.out_uniq),
        palette=args.palette,
        max_run_len=13,
        make_log10=True
    )

    if args.csv_prefix:
        _write_csvs(counts_trans, counts_uniq, L_vals, args.csv_prefix)


if __name__ == "__main__":
    main()
```
