## Supplementary material for "CAD-C: An engineered nuclease enables repair-free *in situ* proximity ligation and nucleosome-resolution chromosome walks in human cells": Custom code: vis_indiv_cadpack_run_viewer_v4_20251020.py

```
#!/usr/bin/env python3
# -*- coding: utf-8 -*-
"""
README — plot_walk_v9_individual.py (v4)
=======================================

Purpose
-------
Replot a single CADwalk run from a **v9** CADpack NPZ and generate a figure with
**alternating rows**: fragment A band, AB contact, fragment B band, …, ending with
fragment E. This makes AB/BC/CD/… semantics explicit and includes terminal
fragments, addressing earlier aesthetic/interpretability concerns.

Key Guarantees
--------------
- **Orientation for display:** default `--display-repair auto` orients cis runs
  left→right (A upstream, E downstream). When a flip is applied, **both steps and
  fragments are reversed together** so that **S0 ≡ AB**, **S1 ≡ BC**, … always holds.
- **Fragment bands (visual):** by default bands are **solid black**; color/opacity/
  edge can be customized via `--frag-*` flags (see below).
- **Font policy:** all vector exports embed TrueType (**Type 42**) fonts
  (`pdf.fonttype=42`, `ps.fonttype=42`, `text.usetex=False`) for searchable text.
- **Layouts:** `--layout alt-rows` (default) for A/AB/B/…/E rows; `step-bands` reproduces
  the earlier “step‑bounded shaded” style if needed.
- **Fragment reference overlap filter:** optional check for **any** reuse/overlap of
  fragment reference intervals (same chromosome), including **non‑adjacent** fragments;
  configurable strictness and action.

Inputs (from NPZ v9)
--------------------
Per-run scalars (L; run_* indices/counts; AJ1/EJ2 terminal junctions; Amid/Emid midpoints),
step arrays (`pos1`, `pos2`, `step_sgnJ`, `chrom1_id`, `chrom2_id`),
fragment arrays (`frag_ref_chr_id`, `frag_ref_lo`, `frag_ref_hi`, `frag_len_bp`,
`frag_len_src`, `frag_index`), and `chrom_names`.

Outputs
-------
- PNG:  <outdir>/walk_v9_run{RUN}.png   (dpi=170)
- PDF:  <outdir>/walk_v9_run{RUN}.pdf   (**Type 42 fonts enforced**)

Usage
-----
python plot_walk_v9_individual.py \
  --npz /path/to/cadpack_v9.npz \
  --run 1234 \
  --outdir ./plots \
  [--layout alt-rows|step-bands] \
  [--require-cis] \
  [--display-repair auto|none|aj|mid] \
  [--ref-overlap-filter off|any|hybrid \
     --ref-overlap-bp-min 1 \
     --ref-overlap-pct-min 5 \
     --on-overlap error|skip|annotate] \
  [--frag-fill black --frag-edge black --frag-alpha 1.0 --frag-edgewidth 0.0]

Fragment Overlap Filter (semantics)
-----------------------------------
- `--ref-overlap-filter any`: exclude if **any** pair of fragments (k≠l) on the same chromosome
  overlap by ≥1 bp.
- `--ref-overlap-filter hybrid`: exclude if overlap length `ov` satisfies
  `ov ≥ max(ref_overlap_bp_min, ref_overlap_pct_min × larger_fragment_length)`.
  `ref_overlap_pct_min` accepts **percent** (e.g., 1,5,10) or **fraction** (0.01,0.05,0.10).
- Pairs considered only when both fragments have valid canonical bounds (`lo < hi`).
- Action: `--on-overlap error|skip|annotate` (default `error`).

Notes
-----
- Top x-axis shows distance (bp) from AJ1; bottom x-axis shows absolute genomic coordinates.
- Set `--frag-fill 0.15 --frag-edge 0.15` for a very dark gray look (still solid).
- `--require-cis` enforces single-chrom bounds for a clean absolute axis.

Changelog
---------
- v4 (2025‑10‑21): Solid dark fragment bands by default with CLI controls; preserved
  alternating‑rows layout; consistent flip of steps **and** fragments; reference overlap
  filter; strict **Type‑42** font export for PDF/EPS.

"""

import os
import argparse
import numpy as np
import matplotlib

# --------------------------- FONT POLICY ---------------------------------------
# PROJECT POLICY: All PDFs/EPS must use TrueType (Type 42) fonts.
matplotlib.use("Agg")
matplotlib.rcParams['pdf.fonttype'] = 42
matplotlib.rcParams['ps.fonttype']  = 42
matplotlib.rcParams['text.usetex']  = False
# ------------------------------------------------------------------------------

import matplotlib.pyplot as plt
from matplotlib.patches import Rectangle
from matplotlib.ticker import FuncFormatter


# ---------- helpers ----------
def _fmt_bp(x, _pos=None):
    try:
        return f"{int(round(x)):,}"
    except Exception:
        return str(x)

def _frag_label(k: int) -> str:
    # A, B, C ... Z, then F{k} beyond 26
    return chr(65 + k) if 0 <= k < 26 else f"F{k}"

def _step_label(k: int) -> str:
    return f"{_frag_label(k)}{_frag_label(k+1)}"

def _as_fraction(p: float) -> float:
    """Accept 5==5% and 0.05==5%; returns a fraction in [0,1]."""
    if p <= 1.0:
        return max(0.0, float(p))
    return float(p) / 100.0

def _find_ref_overlaps(refc, reflo, refhi, flen, mode: str, bp_min: int, pct_min: float):
    """
    Return list of (i, j, chrom_id, ov_len) pairs that violate the overlap policy.
    Only considers pairs with valid bounds on the *same* chromosome.
    mode: 'any' or 'hybrid'
    """
    n = len(refc)
    valid = (refc >= 0) & (reflo >= 0) & (refhi >= 0) & (refhi > reflo)
    pct = _as_fraction(pct_min)

    viol = []
    for i in range(n):
        if not valid[i]:
            continue
        for j in range(i + 1, n):
            if not valid[j]:
                continue
            if refc[i] != refc[j]:
                continue
            ov = min(refhi[i], refhi[j]) - max(reflo[i], reflo[j])
            if ov <= 0:
                continue
            if mode == "any":
                threshold = 1.0  # ≥1 bp
            else:  # hybrid
                max_len = float(max(flen[i], flen[j])) if (flen[i] > 0 and flen[j] > 0) else 0.0
                threshold = max(float(bp_min), _as_fraction(pct_min) * max_len)
            if ov >= threshold:
                viol.append((i, j, int(refc[i]), int(ov)))
    return viol

def _flip_arrays(pos1, pos2, sgnJ, AJ1, EJ2, Amid, Emid,
                 refc, reflo, refhi, flen, lensrc):
    """Reverse step & fragment order for display; maintain S0≡AB mapping."""
    pos1, pos2 = pos2[::-1], pos1[::-1]
    sgnJ = (-sgnJ[::-1]).astype(np.int8)
    AJ1, EJ2 = EJ2, AJ1
    Amid, Emid = Emid, Amid
    refc   = refc[::-1]
    reflo  = reflo[::-1]
    refhi  = refhi[::-1]
    flen   = flen[::-1]
    lensrc = lensrc[::-1]
    return pos1, pos2, sgnJ, AJ1, EJ2, Amid, Emid, refc, reflo, refhi, flen, lensrc


def load_args():
    ap = argparse.ArgumentParser(
        description="Replot one CADwalk run from a v9 NPZ with canonical fragment bounds.",
        formatter_class=argparse.ArgumentDefaultsHelpFormatter
    )
    ap.add_argument("--npz", required=True, help="Path to v9 NPZ")
    ap.add_argument("--run", type=int, required=True, help="Run index (0-based) to plot")
    ap.add_argument("--outdir", default=".", help="Output directory for PNG/PDF")

    # Layout & visuals
    ap.add_argument("--layout", choices=["alt-rows", "step-bands"], default="alt-rows",
                    help="Row layout: alternate fragment and step rows (default) or old step-bounded bands")
    ap.add_argument("--shade-margin", type=float, default=0.06,
                    help="Vertical margin fraction inside each fragment row")
    # Fragment band styling (v4 defaults = solid black)
    ap.add_argument("--frag-fill", default="black",
                    help="Matplotlib color for fragment fill (e.g. 'black', '0.15', '#111')")
    ap.add_argument("--frag-edge", default="black",
                    help="Edge color for fragments (e.g. 'black', 'none')")
    ap.add_argument("--frag-alpha", type=float, default=1.0,
                    help="Fill alpha for fragments")
    ap.add_argument("--frag-edgewidth", type=float, default=0.0,
                    help="Fragment edge line width")

    # Cohesion/orientation
    ap.add_argument("--require-cis", action="store_true",
                    help="Require all fragment bounds to be on a single chromosome")
    ap.add_argument("--display-repair", choices=["auto", "none", "aj", "mid"], default="auto",
                    help="Display-only reorientation rule")

    # Fragment reference overlap (reuse) filter
    ap.add_argument("--ref-overlap-filter", choices=["off", "any", "hybrid"], default="off",
                    help="off: ignore; any: exclude if any pair of fragments overlaps by ≥1 bp; "
                         "hybrid: exclude if overlap ≥ max(--ref-overlap-bp-min, --ref-overlap-pct-min × larger fragment)")
    ap.add_argument("--ref-overlap-bp-min", type=int, default=1,
                    help="Minimum bp overlap used in 'hybrid' mode (default 1 bp)")
    ap.add_argument("--ref-overlap-pct-min", type=float, default=0.0,
                    help="Minimum percent used in 'hybrid' mode; accepts 1 (1%%) or 0.01 (1%%)")
    ap.add_argument("--on-overlap", choices=["error", "skip", "annotate"], default="error",
                    help="What to do when an overlap violation is detected")

    # Raster export DPI (PNG)
    ap.add_argument("--png-dpi", type=int, default=170, help="DPI for PNG export")

    return ap.parse_args()


def main():
    args = load_args()
    os.makedirs(args.outdir, exist_ok=True)

    Z = np.load(args.npz, allow_pickle=False, mmap_mode=None)

    # ---- Pull run-level scalars ----
    r = int(args.run)
    run_L  = int(Z["run_L"][r])
    rfs    = int(Z["run_first_step_row"][r])
    rns    = int(Z["run_n_steps"][r])
    rff    = int(Z["run_first_frag_row"][r])
    rnf    = int(Z["run_n_frags"][r])

    AJ1 = int(Z["run_term_left_junc_bp"][r])
    EJ2 = int(Z["run_term_right_junc_bp"][r])
    Amid = float(Z["run_term_left_mid_bp"][r])   # may be NaN
    Emid = float(Z["run_term_right_mid_bp"][r])  # may be NaN

    # ---- Step slice ----
    s_sl = slice(rfs, rfs + rns)
    pos1 = Z["pos1"][s_sl].astype(np.int64).copy()
    pos2 = Z["pos2"][s_sl].astype(np.int64).copy()
    sgnJ = Z["step_sgnJ"][s_sl].astype(np.int8).copy()
    c1id = Z["chrom1_id"][s_sl].astype(np.int32)

    # ---- Fragment slice ----
    f_sl = slice(rff, rff + rnf)
    fidx = Z["frag_index"][f_sl].astype(np.int64)
    refc = Z["frag_ref_chr_id"][f_sl].astype(np.int32).copy()
    reflo= Z["frag_ref_lo"][f_sl].astype(np.int64).copy()
    refhi= Z["frag_ref_hi"][f_sl].astype(np.int64).copy()
    flen = Z["frag_len_bp"][f_sl].astype(np.int64).copy()
    lensrc = Z["frag_len_src"][f_sl].astype(np.uint8).copy()

    chrom_names = [c.decode("utf-8") if isinstance(c, (bytes, np.bytes_)) else str(c)
                   for c in Z["chrom_names"]]

    # ---- Sanity/robustness around L ----
    L_from_steps = int(pos1.shape[0])
    L_from_frags = int(refc.shape[0] - 1) if refc.shape[0] > 0 else 0
    L = min(run_L, L_from_steps, L_from_frags)
    if (rns != run_L) or (rnf != run_L + 1) or (L != run_L):
        print(f"[WARN] Run {r}: counts differ (run_L={run_L}, steps={rns}, frags={rnf}); using L={L}.")

    # Trim to L/L+1 to avoid any out-of-range issues
    pos1, pos2, sgnJ = pos1[:L], pos2[:L], sgnJ[:L]
    refc, reflo, refhi, flen, lensrc = refc[:L+1], reflo[:L+1], refhi[:L+1], flen[:L+1], lensrc[:L+1]

    # ---- Choose a chromosome for absolute axis (prefer majority of fragments with good bounds) ----
    good_bounds = (refc >= 0) & (reflo >= 0) & (refhi >= 0) & (refhi > reflo)
    frag_chr_vals = refc[good_bounds]
    if frag_chr_vals.size > 0:
        uniq, counts = np.unique(frag_chr_vals, return_counts=True)
        run_chr_id = int(uniq[np.argmax(counts)])
    else:
        run_chr_id = int(c1id[0])

    if args.require_cis:
        incoherent = np.any((refc >= 0) & (refc != run_chr_id))
        if incoherent:
            raise SystemExit(f"[ERROR] Run {r}: fragment bounds span multiple chromosomes; re-run without --require-cis.")

    run_chr_name = chrom_names[run_chr_id] if (0 <= run_chr_id < len(chrom_names)) else f"chr{run_chr_id}"

    # ---- Optional display-only reorientation (consistent: steps *and* fragments) ----
    def _should_flip(rule: str) -> bool:
        if rule == "none":
            return False
        if rule == "mid":
            return (np.isfinite(Amid) and np.isfinite(Emid) and (Emid < Amid))
        if rule == "aj":
            return (EJ2 < AJ1)
        # auto: prefer midpoints if defined, else AJ1/EJ2
        if np.isfinite(Amid) and np.isfinite(Emid):
            return (Emid < Amid)
        return (EJ2 < AJ1)

    if _should_flip(args.display_repair):
        pos1, pos2, sgnJ, AJ1, EJ2, Amid, Emid, refc, reflo, refhi, flen, lensrc = _flip_arrays(
            pos1, pos2, sgnJ, AJ1, EJ2, Amid, Emid, refc, reflo, refhi, flen, lensrc
        )

    # ---- Fragment reference overlap filtering (pairwise across all fragments) ----
    overlap_pairs = []
    if args.ref_overlap_filter != "off":
        overlap_pairs = _find_ref_overlaps(
            refc=refc, reflo=reflo, refhi=refhi, flen=flen,
            mode=args.ref_overlap_filter,
            bp_min=args.ref_overlap_bp_min,
            pct_min=args.ref_overlap_pct_min
        )
        if overlap_pairs:
            min_ov = min(p[3] for p in overlap_pairs)
            msg = (f"Run {r}: REF-OVERLAP violation — {len(overlap_pairs)} pair(s) overlap; "
                   f"min overlap = {min_ov:,} bp; mode={args.ref_overlap_filter}, "
                   f"bp_min={args.ref_overlap_bp_min}, pct_min={args.ref_overlap_pct_min}")
            if args.on_overlap == "skip":
                print("[SKIP]", msg)
                return
            elif args.on_overlap == "error":
                raise SystemExit("[ERROR] " + msg)
            else:
                print("[WARN]", msg, "— annotating figure.")

    # ---- Build rows ----
    gap = 2.0
    if args.layout == "alt-rows":
        n_rows = 2 * L + 1
        y_all  = np.array([(n_rows - 1 - i) * gap for i in range(n_rows)], dtype=float)
        y_frag = np.array([y_all[2*k]     for k in range(L + 1)], dtype=float)  # A...E
        y_step = np.array([y_all[2*k + 1] for k in range(L)], dtype=float)      # AB..DE
        def _frag_band(k: int):
            h = gap * (1.0 - 2.0 * args.shade_margin) * 0.60
            return (y_frag[k] - 0.5*h, y_frag[k] + 0.5*h)
        y_ticks = []
        y_labels = []
        for k in range(L + 1):
            y_ticks.append(y_frag[k]); y_labels.append(_frag_label(k))
            if k < L:
                y_ticks.append(y_step[k]); y_labels.append(_step_label(k))
    else:
        y_step = np.array([(L - j) * gap for j in range(L)], dtype=float)
        def _frag_band_between(k: int):
            if 1 <= k <= L - 1:
                y0, y1 = y_step[k-1], y_step[k]
                y_lo, y_hi = (min(y0, y1), max(y0, y1))
                pad = args.shade_margin * (y_hi - y_lo)
                return (y_lo + pad, y_hi - pad)
            elif k == 0:
                h = gap * (1.0 - 2.0 * args.shade_margin) * 0.6
                return (y_step[0] - 0.5*h, y_step[0] + 0.5*h)
            else:
                h = gap * (1.0 - 2.0 * args.shade_margin) * 0.6
                return (y_step[-1] - 0.5*h, y_step[-1] + 0.5*h)
        y_ticks = list(y_step)
        y_labels = [f"S{j}" for j in range(L)]

    # ---- Determine x limits from junctions + fragment bounds ----
    x_candidates = []
    x_candidates.extend(pos1.tolist()); x_candidates.extend(pos2.tolist())
    good_frags = (refc == run_chr_id) & (reflo >= 0) & (refhi >= 0) & (refhi > reflo)
    x_candidates.extend(reflo[good_frags].tolist())
    x_candidates.extend(refhi[good_frags].tolist())
    if len(x_candidates) == 0:
        raise SystemExit(f"[ERROR] Run {r}: no valid coordinates found to define x-limits.")
    xmin = float(np.min(x_candidates))
    xmax = float(np.max(x_candidates))
    pad = max(50.0, 0.05 * (xmax - xmin) if xmax > xmin else 50.0)

    # ---- Prepare figure ----
    n_display_rows = (2 * L + 1) if (args.layout == "alt-rows") else L
    W = 10.6
    H = max(4.0, 1.6 + 0.45 * n_display_rows)
    fig = plt.figure(figsize=(W, H), facecolor="white")
    ax = fig.add_subplot(111)

    # ---- Draw fragment rectangles (now solid dark by default) ----
    if args.layout == "alt-rows":
        for k in range(L + 1):
            if (refc[k] == run_chr_id) and (reflo[k] >= 0) and (refhi[k] >= 0) and (refhi[k] > reflo[k]):
                lo, hi = float(reflo[k]), float(refhi[k])
            else:
                continue
            y_lo, y_hi = _frag_band(k)
            ax.add_patch(Rectangle((lo, y_lo), hi - lo, y_hi - y_lo,
                                   facecolor=args.frag_fill, edgecolor=args.frag_edge,
                                   linewidth=args.frag_edgewidth, alpha=args.frag_alpha, zorder=0))
    else:
        for k in range(L + 1):
            if (refc[k] == run_chr_id) and (reflo[k] >= 0) and (refhi[k] >= 0) and (refhi[k] > reflo[k]):
                lo, hi = float(reflo[k]), float(refhi[k])
            else:
                continue
            y_lo, y_hi = _frag_band_between(k)
            ax.add_patch(Rectangle((lo, y_lo), hi - lo, y_hi - y_lo,
                                   facecolor=args.frag_fill, edgecolor=args.frag_edge,
                                   linewidth=args.frag_edgewidth, alpha=args.frag_alpha, zorder=0))

    # ---- Draw step arrows & endpoints ----
    for j in range(L):
        y = y_step[j] if args.layout == "alt-rows" else y_ticks[j]
        x1 = float(pos1[j]); x2 = float(pos2[j])
        color = 'red' if x2 > x1 else ('blue' if x2 < x1 else 'gray')
        ax.annotate("", xy=(x2, y), xytext=(x1, y),
                    arrowprops=dict(arrowstyle='-|>', lw=2.8, color=color), zorder=3)
        ax.plot([x1], [y], marker='o', ms=4, color=color, zorder=4)
        ax.plot([x2], [y], marker='o', ms=4, color=color, zorder=4)

    # ---- Axes, labels, and secondary (top) x-axis ----
    ax.set_xlim(xmin - pad, xmax + pad)
    ax.set_ylim((min(y_ticks) - gap, max(y_ticks) + gap) if len(y_ticks) > 0 else (0, gap))
    ax.set_yticks(y_ticks)
    ax.set_yticklabels(y_labels)
    ax.grid(axis='x', linestyle='--', alpha=0.30)

    # Bottom x-axis: absolute coordinate
    ax.set_xlabel(f"{run_chr_name} coordinate (bp)")
    ax.xaxis.set_major_formatter(FuncFormatter(_fmt_bp))

    # Top x-axis: Δ from AJ1
    def _to_rel(x): return x - AJ1
    def _to_abs(xrel): return xrel + AJ1
    ax_top = ax.secondary_xaxis("top", functions=(_to_rel, _to_abs))
    ax_top.xaxis.set_major_formatter(FuncFormatter(_fmt_bp))
    ax_top.set_xlabel("Distance from walk start (AJ1) [bp]")

    # ---- Title with quick stats (and overlap annotation if requested) ----
    patt = "".join("+" if s > 0 else ("-" if s < 0 else ".") for s in sgnJ[:L])
    deltaJ = EJ2 - AJ1
    deltaM = (Emid - Amid) if (np.isfinite(Amid) and np.isfinite(Emid)) else np.nan
    subtitle = f"ΔJ={deltaJ:,} bp"
    if np.isfinite(deltaM):
        subtitle += f"  |  ΔM={deltaM:,.0f} bp"
    title_left = f"Run {r} (L={L}) — pattern {patt}"
    if overlap_pairs and args.on_overlap == "annotate":
        min_ov = min(p[3] for p in overlap_pairs)
        title_left += f"  |  REF-OVERLAP ⚠︎ {len(overlap_pairs)} pair(s), min {min_ov:,} bp"
    ax.set_title(f"{title_left}  |  {subtitle}", fontsize=11.5)

    fig.tight_layout()
    base = os.path.join(args.outdir, f"walk_v9_run{r}")
    fig.savefig(base + ".png", dpi=args.png_dpi)
    fig.savefig(base + ".pdf")
    plt.close(fig)
    print(f"[OK] Saved: {base}.png/.pdf  |  layout={args.layout}, repair={args.display_repair}, "
          f"overlap={args.ref_overlap_filter}, action={args.on_overlap}, frag_fill={args.frag_fill}")


if __name__ == "__main__":
    main()
```
