## Supplementary material for "CAD-C: An engineered nuclease enables repair-free *in situ* proximity ligation and nucleosome-resolution chromosome walks in human cells": Custom code: mcool_Ps_curve_v1_20250411.py

```
#!/usr/bin/env python3
# P(s) Curve Calculation Script
#
# Description:
#   This script computes the P(s) curve (intrachromosomal contact probability as a function
#   of genomic separation) for one or multiple Hi-C contact maps provided in .mcool format.
#   By default, it works on hg38 but can adapt to other builds or skip centromere fetching
#   entirely. It uses the `cooler` and `cooltools` libraries to calculate expected contact
#   frequencies (P(s)) and then plots the decay of contact probability with distance.
#
#   For resolution handling:
#       1) If the user provides `--binsize XYZ`, the script tries to open that resolution
#          in each .mcool file. It will error out if any file lacks it.
#       2) If `--binsize` is not provided, the script looks for a common resolution among
#          all the input .mcool files:
#             - If 500 bp is found in all files, it uses 500 bp.
#             - Otherwise, it picks the smallest resolution (highest-res in base pairs)
#               that all files share.
#       3) If there is no common resolution, the script fails.
#
#   You can also specify `--threads <N>` to pass a multi-processing argument (`nproc=N`)
#   to `cooltools.expected_cis`, potentially speeding up certain steps if multiple cores
#   are available.
#
#   The user can now specify `--x-min`/`--x-max` to restrict the plotted data range (in bp).
#   By default, x_min=1e3 and x_max=None, so the plot starts at 10^3 bp with no forced upper limit.
#   The gridlines are drawn behind the data lines.
#
#   NEW: The slope panel’s y-axis defaults to an upper limit (y_max) of 0. By default, the lower
#   limit is auto-scaled. The user can override via `--slope-y-min` and `--slope-y-max`.
#
# Usage:
#   python mcool_Ps_curve_v1_20250411.py --mcool dataset1.mcool dataset2.mcool [...] \
#                             [--binsize 5000] \
#                             [--expected-tsv dataset1_expected.tsv dataset2_expected.tsv ...] \
#                             [--aliases alias1 alias2 ...] \
#                             [--colors color1 color2 ...] \
#                             [--order idx1 idx2 ...] \
#                             [--ps-ratio RATIO] \
#                             [--genome-build BUILD_OR_NONE] \
#                             [--threads N] \
#                             [--x-min VAL] [--x-max VAL] \
#                             [--slope-y-min VAL] [--slope-y-max VAL] \
#                             [--output-dir OUTPUT_DIR]
#
# Options:
#   --mcool            Paths to one or more .mcool file(s). Each can have multiple
#                      resolutions. If multiple are provided, the script processes them in
#                      order or as specified by --order.
#
#   --binsize          Optional integer specifying the resolution (binsize, in bp) to open
#                      within each .mcool file. If not provided, the script checks for a
#                      common resolution across all files:
#                        (a) If 500 is in all, uses 500 bp.
#                        (b) Else uses the smallest common resolution in all files.
#
#   --expected-tsv     Optional path(s) to precomputed expected P(s) TSV file(s), one per
#                      .mcool file. If omitted, an expected file is computed for each .mcool
#                      on the chosen binsize.
#
#   --aliases          Optional custom labels for each dataset to use in plot legends and
#                      output file names. Must match the number of .mcool files.
#
#   --colors           Optional list of colors for plot lines. If not provided, default
#                      matplotlib colors are used.
#
#   --order            Optional custom order (0-based indices) for plotting the datasets.
#                      By default, datasets are plotted in the order given under --mcool.
#
#   --ps-ratio         Optional float controlling the height:width ratio of the main P(s) plot
#                      (upper panel). Default=1.0 (square).
#
#   --genome-build     Genome assembly for centromere fetching, default is 'hg38'. If set to
#                      'none', the script will skip chromosome-arm calculations and compute
#                      on entire chromosomes.
#
#   --threads          Integer specifying the number of parallel processes for
#                      cooltools.expected_cis (nproc=...). Default=1 (no parallel partitioning).
#
#   --x-min            Lower bound (in bp) for the plotted separation range. Default=1e3
#                      (i.e., 10^3 bp). Any data below this is excluded from the figure.
#
#   --x-max            Upper bound (in bp) for the plotted separation range. Default=None
#                      (no forced upper limit). Any data above x-max is excluded from the figure.
#
#   --slope-y-min      Lower y-limit for the slope panel. Default=None (auto-scale).
#   --slope-y-max      Upper y-limit for the slope panel. Default=0. So by default, the slope
#                      plot will only show slope ≤ 0. If your data has slope>0 that you want
#                      to see, you can override this.
#
#   --output-dir       Directory where a timestamped subdirectory will be created to store
#                      all output files. Default is the current directory.
#
# Outputs:
#   - A PDF plot file named "Ps_<timestamp>.pdf", with:
#       * Upper panel: P(s) curves (log–log).
#       * Lower panel: slope of log(P(s)) (semi-log).
#   - For each dataset, a TSV file "<basename>_Ps.tsv" containing:
#       distance_bp        Contact separation in base pairs.
#       contact_frequency  The P(s) contact frequency at that distance.
#       slope              The local slope of log(P(s)) at that distance.
#   - If the script computed expected (rather than using --expected-tsv), it also saves
#     "<basename>_expected.tsv" with columns 'distance_bp' and 'contact_frequency'.
#
# Logging:
#   Informative, timestamped log messages are printed to the terminal (or to Slurm stdout/stderr).
#
# Requirements:
#   Python 3 with:
#     * numpy, pandas, matplotlib, argparse, datetime
#     * cooler, cooltools>=0.5.2, bioframe
#     * packaging (for version parsing)
#
# Author: Jan Soroczynski, Risca Lab, The Rockefeller University. 
# Date: 2025_04_11.

import os
import sys
import argparse
import logging
from datetime import datetime
import numpy as np
import pandas as pd
import matplotlib
import matplotlib.pyplot as plt
import cooler
import cooltools
import bioframe
from packaging import version

def main():
    # ------------------------------------------------
    # Configure logging with timestamp
    # ------------------------------------------------
    logging.basicConfig(level=logging.INFO,
                        format="[%(asctime)s] %(levelname)s: %(message)s",
                        datefmt="%Y-%m-%d %H:%M:%S")

    # ------------------------------------------------
    # Check cooltools version requirement
    # ------------------------------------------------
    if version.parse(cooltools.__version__) < version.parse("0.5.2"):
        logging.error("cooltools >=0.5.2 is required (found %s). Please update.",
                      cooltools.__version__)
        sys.exit(1)

    # ------------------------------------------------
    # Parse arguments
    # ------------------------------------------------
    parser = argparse.ArgumentParser(
        description="Compute and plot P(s) contact decay curves from .mcool Hi-C data."
    )
    parser.add_argument("--mcool", required=True, nargs="+",
                        help="Path(s) to one or more .mcool files with Hi-C data.")
    parser.add_argument("--binsize", type=int, default=None,
                        help="Resolution (bp) to open in each .mcool. If not provided, checks common resolutions.")
    parser.add_argument("--expected-tsv", nargs="+",
                        help="Optional path(s) to precomputed expected P(s) TSV file(s), one per .mcool.")
    parser.add_argument("--aliases", nargs="+",
                        help="Optional custom labels for each dataset.")
    parser.add_argument("--colors", nargs="+",
                        help="Optional list of colors for lines.")
    parser.add_argument("--order", nargs="+", type=int,
                        help="Optional custom plotting order (0-based).")
    parser.add_argument("--ps-ratio", type=float, default=1.0,
                        help="Height:width ratio of main P(s) plot. Default=1.0 (square).")
    parser.add_argument("--genome-build", default="hg38",
                        help="Genome for centromere fetching, or 'none' to skip.")
    parser.add_argument("--threads", type=int, default=1,
                        help="Number of parallel processes for cooltools.expected_cis. Default=1.")
    parser.add_argument("--x-min", type=float, default=1e3,
                        help="Minimum separation in bp to plot. Default=1e3.")
    parser.add_argument("--x-max", type=float, default=None,
                        help="Maximum separation in bp to plot (optional). Default=None.")
    parser.add_argument("--slope-y-min", type=float, default=None,
                        help="Lower y-limit for slope panel. Default=None (auto).")
    parser.add_argument("--slope-y-max", type=float, default=0,
                        help="Upper y-limit for slope panel. Default=0 (so slope ≤0 is visible).")
    parser.add_argument("--output-dir", default=".",
                        help="Directory where a timestamped subdirectory will store outputs.")
    args = parser.parse_args()

    # ------------------------------------------------
    # Validate inputs
    # ------------------------------------------------
    num_files = len(args.mcool)
    for path in args.mcool:
        if not os.path.isfile(path):
            logging.error("Input file not found: %s", path)
            sys.exit(1)

    if args.expected_tsv:
        if len(args.expected_tsv) != num_files:
            logging.error("Number of expected TSV files doesn't match .mcool count.")
            sys.exit(1)
        for exp in args.expected_tsv:
            if not os.path.isfile(exp):
                logging.error("Expected TSV file not found: %s", exp)
                sys.exit(1)

    if args.aliases and len(args.aliases) != num_files:
        logging.error("Number of aliases doesn't match number of .mcool files.")
        sys.exit(1)

    if args.threads < 1:
        logging.error("--threads must be >= 1.")
        sys.exit(1)

    # ------------------------------------------------
    # Order handling
    # ------------------------------------------------
    order = list(range(num_files))
    if args.order:
        if len(args.order) != num_files:
            logging.error("--order must have exactly %d indices.", num_files)
            sys.exit(1)
        if sorted(args.order) != list(range(num_files)):
            logging.error("--order must be a permutation of [0..%d].", num_files - 1)
            sys.exit(1)
        order = args.order

    # ------------------------------------------------
    # Output dir
    # ------------------------------------------------
    timestamp = datetime.now().strftime("%Y%m%d_%H%M%S")
    out_dir = os.path.join(args.output_dir, timestamp)
    try:
        os.makedirs(out_dir, exist_ok=False)
    except Exception as e:
        logging.error("Could not create output directory %s: %s", out_dir, str(e))
        sys.exit(1)
    logging.info("Created output directory: %s", out_dir)

    # ------------------------------------------------
    # Plotting style
    # ------------------------------------------------
    try:
        plt.style.use("seaborn-v0_8-poster")
    except Exception:
        logging.warning("Could not apply 'seaborn-v0_8-poster' style. Using default.")

    matplotlib.rcParams['font.family'] = 'Helvetica'
    matplotlib.rcParams['pdf.fonttype'] = 42  # text in PDF is editable

    # ------------------------------------------------
    # Genome build
    # ------------------------------------------------
    if args.genome_build.lower() != "none":
        try:
            logging.info("Fetching centromeres for genome: %s", args.genome_build)
            cens = bioframe.fetch_centromeres(args.genome_build)
        except Exception as e:
            logging.error("Failed to fetch centromeres for %s: %s", args.genome_build, str(e))
            sys.exit(1)
    else:
        logging.info("Skipping centromere fetching (genome-build='none').")
        cens = None

    # ------------------------------------------------
    # Determine binsize if needed
    # ------------------------------------------------
    desired_binsize = args.binsize
    if desired_binsize is not None:
        # Check each file
        for path in args.mcool:
            found = _get_mcool_resolutions(path)
            if desired_binsize not in found:
                logging.error("File '%s' lacks binsize=%d. Available: %s", path, desired_binsize, sorted(found))
                sys.exit(1)
        logging.info("Using user binsize=%d for all .mcool files.", desired_binsize)
    else:
        # Auto-detect
        logging.info("No --binsize provided. Checking common resolutions in all .mcool files...")
        sets_list = []
        for path in args.mcool:
            sets_list.append(_get_mcool_resolutions(path))
        common = set.intersection(*sets_list)
        if not common:
            logging.error("No common resolutions among all .mcool inputs.")
            sys.exit(1)
        if 500 in common:
            desired_binsize = 500
            logging.info("All have 500 bp. Using binsize=500.")
        else:
            fallback = min(common)
            desired_binsize = fallback
            logging.info("Using smallest common resolution=%d (500 not found).", desired_binsize)

    # ------------------------------------------------
    # Main data loop
    # ------------------------------------------------
    results_list = []
    expected_paths = args.expected_tsv if args.expected_tsv else [None]*num_files

    for i, mcool_path in enumerate(args.mcool):
        default_label = os.path.basename(mcool_path)
        base_name = default_label
        if base_name.endswith(".mcool") or base_name.endswith(".cool"):
            base_name = os.path.splitext(base_name)[0]
        alias_label = args.aliases[i] if args.aliases else default_label

        # If user provided expected
        if expected_paths[i] is not None:
            exp_path = expected_paths[i]
            logging.info("Using precomputed expected for %s from %s", default_label, exp_path)
            df = _read_expected_tsv(exp_path, desired_binsize)
            distances = df["distances"]
            contact_freq = df["contact_freq"]
            slope = df["slope"]

        else:
            logging.info("Computing expected for %s at %d bp", default_label, desired_binsize)
            distances, contact_freq, slope = _compute_expected(
                mcool_path, desired_binsize, cens, alias_label, out_dir, args.threads
            )

        results_list.append({
            "label": alias_label,
            "base_name": base_name,
            "distances": distances,
            "contact_freq": contact_freq,
            "slope": slope
        })

    # Reorder
    if order != list(range(num_files)):
        results_list = [results_list[i] for i in order]

    if args.colors:
        cycle = plt.cycler(color=args.colors)
        matplotlib.rcParams['axes.prop_cycle'] = cycle

    # ------------------------------------------------
    # Make figure
    # ------------------------------------------------
    plot_width = 6.5
    top_height_inches = plot_width * args.ps_ratio
    bottom_height_inches = plot_width / 3.0
    total_height = top_height_inches + bottom_height_inches

    fig, axs = plt.subplots(nrows=2, figsize=(plot_width, total_height),
                            sharex=True,
                            gridspec_kw={'height_ratios': [top_height_inches, bottom_height_inches]})
    plt.subplots_adjust(hspace=0.15)

    # Grid behind data
    axs[0].set_axisbelow(True)
    axs[1].set_axisbelow(True)

    # Plot
    for idx, rdict in enumerate(results_list):
        dist = rdict["distances"]
        freq = rdict["contact_freq"]
        slope = rdict["slope"]
        label = rdict["label"]

        # Subset by x_min/x_max
        mask = (dist >= args.x_min)
        if args.x_max is not None:
            mask &= (dist <= args.x_max)
        dist_plot = dist[mask]
        freq_plot = freq[mask]
        slope_plot = slope[mask]

        axs[0].loglog(dist_plot, freq_plot, '-', markersize=4, label=label, zorder=2+idx)
        axs[1].semilogx(dist_plot, slope_plot, '-', label=label, zorder=2+idx)

    axs[0].grid(linewidth=0.5, zorder=1)
    axs[1].grid(linewidth=0.5, zorder=1)

    axs[0].set_ylabel("IC contact frequency")
    axs[1].set_xlabel("Separation (bp)")
    axs[1].set_ylabel("Slope")

    axs[0].legend()
    axs[1].legend()

    # slope panel y-limits
    if (args.slope_y_min is not None) or (args.slope_y_max is not None):
        axs[1].set_ylim(args.slope_y_min, args.slope_y_max)
    else:
        axs[1].set_ylim(None, 0)

    # Save fig
    pdf_path = os.path.join(out_dir, f"Ps_{timestamp}.pdf")
    try:
        plt.savefig(pdf_path, format='pdf', bbox_inches='tight')
        logging.info("Saved P(s) plot to %s", pdf_path)
    except Exception as e:
        logging.error("Could not save PDF: %s", str(e))
    plt.close(fig)

    # Save numeric outputs
    for rd in results_list:
        label = rd["label"]
        df_out = pd.DataFrame({
            "distance_bp": rd["distances"],
            "contact_frequency": rd["contact_freq"],
            "slope": rd["slope"]
        })
        tsv_out = os.path.join(out_dir, f"{rd['base_name']}_Ps.tsv")
        try:
            df_out.to_csv(tsv_out, sep='\t', index=False)
            logging.info("Wrote data for %s to %s", label, tsv_out)
        except Exception as e:
            logging.error("Failed to write TSV for %s: %s", label, str(e))

    logging.info("Analysis complete.")


# ------------------------------------------------
# HELPER FUNCTIONS
# ------------------------------------------------

def _compute_expected(mcool_path, binsize, cens, label, out_dir, threads=1):
    """
    Open the .mcool at 'binsize', compute expected_cis with nproc=threads,
    return (distances, contact_freq, slope).
    Also saves a <basename>_expected.tsv to 'out_dir'.
    """
    import os

    base_name = os.path.basename(mcool_path)
    if base_name.endswith(".mcool") or base_name.endswith(".cool"):
        base_name = os.path.splitext(base_name)[0]

    try:
        clr = cooler.Cooler(f"{mcool_path}::resolutions/{binsize}")
    except Exception as e:
        logging.error("Failed to open cooler at %d for %s: %s", binsize, mcool_path, str(e))
        sys.exit(1)

    actual = clr.binsize
    if actual != binsize:
        logging.error("Cooler indicates binsize=%d, mismatch with requested %d for %s.", actual, binsize, label)
        sys.exit(1)

    # Build arms or entire chroms
    try:
        chrom_df = clr.chroms()[:]
        if 'name' in chrom_df.columns:
            chrom_df.rename(columns={'name': 'chrom'}, inplace=True)
        if 'length' not in chrom_df.columns:
            logging.error("No 'length' col in chroms table for %s.", mcool_path)
            sys.exit(1)
        chrom_sizes = chrom_df[['chrom', 'length']]
    except Exception as e:
        logging.error("Error reading chromosomes from %s: %s", mcool_path, str(e))
        sys.exit(1)

    if cens is not None:
        try:
            arms = bioframe.make_chromarms(chrom_sizes, cens)
        except Exception as e:
            logging.error("Error building chromosome arms: %s", str(e))
            sys.exit(1)
    else:
        arms = chrom_sizes.copy()
        arms['start'] = 0
        arms.rename(columns={'length': 'end'}, inplace=True)

    # Compute expected with multi-threading
    try:
        cvd = cooltools.expected_cis(
            clr=clr,
            view_df=arms,
            smooth=True,
            aggregate_smoothed=True,
            nproc=threads  # <--- CRITICAL FIX: pass nproc
        )
    except Exception as e:
        logging.error("Error in cooltools.expected_cis for %s: %s", label, str(e))
        sys.exit(1)

    if 'dist' not in cvd.columns:
        logging.error("'dist' col missing in expected data for %s.", label)
        sys.exit(1)

    cvd['distance_bp'] = cvd['dist'] * binsize
    if 'balanced.avg.smoothed.agg' in cvd.columns:
        col_sel = 'balanced.avg.smoothed.agg'
    else:
        col_sel = 'balanced.avg'

    df_uniq = cvd.drop_duplicates(subset=['dist'])[['distance_bp', col_sel]].copy()
    freq = df_uniq[col_sel].values
    dists = df_uniq['distance_bp'].values
    good_idx = ~np.isnan(freq) & (dists > 0)
    dists = dists[good_idx]
    freq = freq[good_idx]

    if len(dists) < 2:
        logging.error("Not enough valid points in computed expected for %s.", label)
        sys.exit(1)

    slope = np.gradient(np.log(freq), np.log(dists))

    # Save expected
    exp_out = os.path.join(out_dir, f"{base_name}_expected.tsv")
    try:
        pd.DataFrame({
            'distance_bp': dists,
            'contact_frequency': freq
        }).to_csv(exp_out, sep='\t', index=False)
        logging.info("Saved computed expected for %s -> %s", label, exp_out)
    except Exception as e:
        logging.warning("Could not save expected for %s: %s", label, str(e))

    return dists, freq, slope


def _read_expected_tsv(filepath, binsize):
    """
    Read a precomputed expected file from 'filepath', parse out distances & freq,
    compute slope. If the 'dist' column is in bin units, we multiply by binsize.
    Returns a dict with keys 'distances','contact_freq','slope'.
    """
    import pandas as pd
    import numpy as np

    try:
        df = pd.read_csv(filepath, sep='\t' if filepath.endswith(('.tsv','.txt')) else ',', comment='#')
    except Exception as e:
        logging.error("Failed to read expected TSV %s: %s", filepath, str(e))
        sys.exit(1)

    # distance column
    if 's_bp' in df.columns:
        dist = df['s_bp'].values
    elif 'distance' in df.columns:
        dist = df['distance'].values
    elif 'distance_bp' in df.columns:
        dist = df['distance_bp'].values
    elif 'dist' in df.columns:
        dist = df['dist'].values
        # Maybe in bins
        if dist.max() < 1e5:
            dist = dist * binsize
    else:
        logging.error("No recognized distance column in %s. "
                      "Expected one of s_bp, distance, distance_bp, dist.", filepath)
        sys.exit(1)

    # contact freq col
    freq_col = None
    for c in ['balanced.avg.smoothed.agg', 'balanced.avg', 'contact_frequency']:
        if c in df.columns:
            freq_col = c
            break
    if freq_col is None:
        logging.error("No recognized contact freq column in %s. "
                      "Expected one of balanced.avg.smoothed.agg, balanced.avg, contact_frequency.", filepath)
        sys.exit(1)

    freq = df[freq_col].values
    dist = dist.astype(float)
    freq = freq.astype(float)
    valid = ~np.isnan(freq) & (dist > 0)
    dist = dist[valid]
    freq = freq[valid]
    if len(dist) < 2:
        logging.error("Insufficient points in %s after cleanup.", filepath)
        sys.exit(1)

    slope = np.gradient(np.log(freq), np.log(dist))

    return {
        "distances": dist,
        "contact_freq": freq,
        "slope": slope
    }


def _get_mcool_resolutions(mcool_path):
    """
    Return a set of integer resolutions (binsizes) available in the given .mcool file.

    We use cooler.fileops.list_coolers to get subpaths like '/resolutions/500', etc.
    Strip leading '/', parse out integer from 'resolutions/XXX'.
    """
    from cooler.fileops import list_coolers
    subs = list_coolers(mcool_path)
    res_set = set()
    for sp in subs:
        sp_stripped = sp.lstrip('/')
        if sp_stripped.startswith("resolutions/"):
            try:
                val = int(sp_stripped.split('/')[1])
                res_set.add(val)
            except ValueError:
                pass
    return res_set


if __name__ == "__main__":
    main()
```
