## Supplementary material for "CAD-C: An engineered nuclease enables repair-free *in situ* proximity ligation and nucleosome-resolution chromosome walks in human cells": Custom code: pairs_pipe_20240822_v2.sh

```
#!/bin/bash

# Activate the conda environment
source /ru-auth/local/home/risc_soft/miniconda3/bin/activate open2c

# Set the chroms path as specified
chroms_path="/rugpfs/fs0/risc_lab/scratch/jsoroczyns/CAD/CAD-C/CADC_E2_P1_Novogene/20240320_pt_flip_out/hg38.chrom.sizes"

# Get the directory where BAM files are located, output directory, and partition
input_dir="$1"
output_dir="$2"
partition="$3"

# Ensure that both input, output directories, and partition are provided
if [ -z "$input_dir" ] || [ -z "$output_dir" ] || [ -z "$partition" ]; then
    echo "Usage: $0 <input_directory> <output_directory> <partition>"
    exit 1
fi

# List BAM files in the input directory
bam_files=($(ls "$input_dir"/*.bam))
if [ ${#bam_files[@]} -eq 0 ]; then
    echo "No BAM files found in the specified input directory: $input_dir"
    exit 1
fi

# Print the files that will be processed
echo "Processing the following BAM files:"
for bam_file in "${bam_files[@]}"; do
    echo "$(basename "$bam_file")"
done

# Create the output and log directories
timestamp=$(date +%Y%m%d)
log_dir="${output_dir}/logs_${timestamp}"
output_dir="${output_dir}/results_${timestamp}"

mkdir -p "$log_dir" "$output_dir"

process_replicate() {
    local bam_file=$1
    local output_dir=$2
    local log_dir=$3
    local partition=$4
    local timestamp=$(date +%Y%m%d)

    replicate=$(basename "$bam_file" .bam)

    pairs_file="${output_dir}/${replicate}_parse2_${timestamp}.pairs.gz"
    stats_file="${output_dir}/${replicate}_parse2_stats_${timestamp}.tsv"
    sorted_file="${output_dir}/${replicate}_sorted_${timestamp}.pairs.gz"
    dedup_file="${output_dir}/${replicate}_dedup_${timestamp}.pairs.gz"
    flipped_file="${output_dir}/${replicate}_flipped_${timestamp}.pairs.gz"
    selected_direct_file="${output_dir}/${replicate}_selected_direct_${timestamp}.pairs.gz"
    selected_UU_file="${output_dir}/${replicate}_selected_UU_${timestamp}.pairs.gz"
    dedup_final_file="${output_dir}/${replicate}_dedup_final_${timestamp}.pairs.gz"
    flipped_sorted_direct_file="${output_dir}/${replicate}_flipped_sorted_direct_${timestamp}.pairs.gz"
    dedup_stats_file="${output_dir}/${replicate}_dedup_stats_${timestamp}.tsv"
    dedup_final_stats_file="${output_dir}/${replicate}_dedup_final_stats_${timestamp}.tsv"

    # Step 1: Parse BAM file to Pairs using pairtools parse2
    parse_job_id=$(sbatch --partition=$partition <<EOF
#!/bin/bash
#SBATCH --job-name=parse2_${replicate}
#SBATCH --output=${log_dir}/parse2_${replicate}%j.out
#SBATCH --error=${log_dir}/parse2_${replicate}%j.err
#SBATCH --ntasks=1
#SBATCH --cpus-per-task=16

source /ru-auth/local/home/risc_soft/miniconda3/bin/activate open2c

pairtools parse2 \
    --chroms-path ${chroms_path} \
    --output ${pairs_file} \
    --max-inter-align-gap 30 \
    --min-mapq 30 \
    --drop-sam \
    --report-position junction \
    --report-orientation junction \
    --add-pair-index \
    --no-expand \
    --nproc-in 16 \
    --nproc-out 16 \
    --no-flip \
    --add-columns mapq \
    --add-columns cigar \
    --add-columns matched_bp \
    --add-columns algn_ref_span \
    --add-columns dist_to_5 \
    --add-columns dist_to_3 \
    --output-stats ${stats_file} \
    ${bam_file}
EOF
)
    parse_job_id=$(echo $parse_job_id | cut -d ' ' -f 4)

    # Step 2: Flip pairs (After Parse2 completes)
    flip_job_id=$(sbatch --partition=$partition --dependency=afterok:$parse_job_id <<EOF
#!/bin/bash
#SBATCH --job-name=pt_flip_${replicate}
#SBATCH --output=${log_dir}/flip_${replicate}%j.out
#SBATCH --error=${log_dir}/flip_${replicate}%j.err
#SBATCH --ntasks=1
#SBATCH --cpus-per-task=16

source /ru-auth/local/home/risc_soft/miniconda3/bin/activate open2c

pairtools flip --output ${flipped_file} --chroms-path ${chroms_path} --nproc-in 16 --nproc-out 16 ${pairs_file}
EOF
)
    flip_job_id=$(echo $flip_job_id | cut -d ' ' -f 4)

    # Step 3: Sort pairs
    sort_job_id=$(sbatch --partition=$partition --dependency=afterok:$flip_job_id <<EOF
#!/bin/bash
#SBATCH --job-name=pt_sort_${replicate}
#SBATCH --output=${log_dir}/sort_${replicate}%j.out
#SBATCH --error=${log_dir}/sort_${replicate}%j.err
#SBATCH --ntasks=1
#SBATCH --cpus-per-task=16

source /ru-auth/local/home/risc_soft/miniconda3/bin/activate open2c

pairtools sort --output ${sorted_file} --nproc-in 16 --nproc-out 16 ${flipped_file}
EOF
)
    sort_job_id=$(echo $sort_job_id | cut -d ' ' -f 4)

    # Step 4: Deduplicate pairs (first round)
    dedup_job_id=$(sbatch --partition=$partition --dependency=afterok:$sort_job_id <<EOF
#!/bin/bash
#SBATCH --job-name=pt_dedup_${replicate}
#SBATCH --output=${log_dir}/dedup_${replicate}%j.out
#SBATCH --error=${log_dir}/dedup_${replicate}%j.err
#SBATCH --ntasks=1
#SBATCH --cpus-per-task=16

source /ru-auth/local/home/risc_soft/miniconda3/bin/activate open2c

pairtools dedup --output ${dedup_file} --output-stats ${dedup_stats_file} --nproc-in 16 --nproc-out 16 ${sorted_file}
EOF
)
    dedup_job_id=$(echo $dedup_job_id | cut -d ' ' -f 4)

    # Step 5: Select direct pairs
    select_direct_job_id=$(sbatch --partition=$partition --dependency=afterok:$dedup_job_id <<EOF
#!/bin/bash
#SBATCH --job-name=pt_select_direct_${replicate}
#SBATCH --output=${log_dir}/select_direct_${replicate}%j.out
#SBATCH --error=${log_dir}/select_direct_${replicate}%j.err
#SBATCH --ntasks=1
#SBATCH --cpus-per-task=16

source /ru-auth/local/home/risc_soft/miniconda3/bin/activate open2c

pairtools select --output ${selected_direct_file} --nproc-in 16 --nproc-out 16 '(walk_pair_type == "R1" or walk_pair_type == "R2" or walk_pair_type == "R1&2")' ${dedup_file}
EOF
)
    select_direct_job_id=$(echo $select_direct_job_id | cut -d ' ' -f 4)

    # Step 6: Save the flipped and sorted direct pairs
    save_flipped_sorted_direct_job_id=$(sbatch --partition=$partition --dependency=afterok:$select_direct_job_id <<EOF
#!/bin/bash
#SBATCH --job-name=pt_save_flipped_sorted_direct_${replicate}
#SBATCH --output=${log_dir}/save_flipped_sorted_direct_${replicate}%j.out
#SBATCH --error=${log_dir}/save_flipped_sorted_direct_${replicate}%j.err
#SBATCH --ntasks=1
#SBATCH --cpus-per-task=16

cp ${selected_direct_file} ${flipped_sorted_direct_file}
EOF
)
    save_flipped_sorted_direct_job_id=$(echo $save_flipped_sorted_direct_job_id | cut -d ' ' -f 4)

    # Step 7: Select UU pairs
    select_UU_job_id=$(sbatch --partition=$partition --dependency=afterok:$save_flipped_sorted_direct_job_id <<EOF
#!/bin/bash
#SBATCH --job-name=pt_select_UU_${replicate}
#SBATCH --output=${log_dir}/select_UU_${replicate}%j.out
#SBATCH --error=${log_dir}/select_UU_${replicate}%j.err
#SBATCH --ntasks=1
#SBATCH --cpus-per-task=16

source /ru-auth/local/home/risc_soft/miniconda3/bin/activate open2c

pairtools select --output ${selected_UU_file} --nproc-in 16 --nproc-out 16 'pair_type == "UU"' ${selected_direct_file}
EOF
)
    select_UU_job_id=$(echo $select_UU_job_id | cut -d ' ' -f 4)

    # Step 8: Deduplicate UU pairs
    sbatch --partition=$partition --dependency=afterok:$select_UU_job_id <<EOF
#!/bin/bash
#SBATCH --job-name=pt_dedup_final_${replicate}
#SBATCH --output=${log_dir}/dedup_final_${replicate}%j.out
#SBATCH --error=${log_dir}/dedup_final_${replicate}%j.err
#SBATCH --ntasks=1
#SBATCH --cpus-per-task=16

source /ru-auth/local/home/risc_soft/miniconda3/bin/activate open2c

pairtools dedup --output ${dedup_final_file} --output-stats ${dedup_final_stats_file} --nproc-in 16 --nproc-out 16 ${selected_UU_file}
EOF
}

# Process each BAM file found in the input directory
for bam_file in "${bam_files[@]}"; do
    process_replicate $bam_file $output_dir $log_dir $partition &
done

wait
```
