## Supplementary material for "CAD-C: An engineered nuclease enables repair-free *in situ* proximity ligation and nucleosome-resolution chromosome walks in human cells": Custom code: pairs_to_500_igdiag1_bal_cool_500N_mcool_v2.sh

```
#!/bin/bash
#SBATCH --cpus-per-task=16
#SBATCH --job-name=cooler_pipeline
#SBATCH --output=%x_%j.out

# Check for input directory
if [ $# -ne 1 ]; then
  echo "Usage: $0 <input_directory>"
  exit 1
fi

input_dir=$1
output_dir="${input_dir}/cool_output"  # Create output directory within the input directory
timestamp=$(date +"%Y%m%d_%H%M%S")
report_file="${output_dir}/cooler_report_${timestamp}.txt"

# Create output directory if it does not exist
mkdir -p $output_dir

# Initialize report
echo "Cooler Pipeline Report - $timestamp" > $report_file
echo "Input Directory: $input_dir" >> $report_file
echo "Output Directory: $output_dir" >> $report_file
echo "----------------------------------" >> $report_file

# Iterate over .pairs.gz files in the input directory
for pairs_file in "$input_dir"/*.pairs.gz; do
  base_name=$(basename "$pairs_file" .pairs.gz)
  
  # Step 1: Submit job to create 500 bp .cool file
  cool_file="${output_dir}/${base_name}_500.cool"
  cool_job_id=$(sbatch --parsable -p vgl --cpus-per-task=16 <<EOF
#!/bin/bash
#SBATCH --job-name=cool_${base_name}
#SBATCH --output=cool_${base_name}_%j.out
#SBATCH --cpus-per-task=16

# Load required modules or activate conda environment
source /ru-auth/local/home/risc_soft/miniconda3/etc/profile.d/conda.sh
conda activate open2c

# Generate 500 bp .cool file using specified hg38 chromosome sizes path
cooler cload pairs -c1 2 -p1 3 -c2 4 -p2 5 --assembly hg38 /lustre/fs4/risc_lab/store/risc_data/downloaded/hg38/genome/chrom.sizes:500 \
  "$pairs_file" "$cool_file"
EOF
)

  # Step 2: Submit job to balance .cool file after Step 1 is complete
  balanced_file="${output_dir}/${base_name}_500_bal.cool"
  balance_job_id=$(sbatch --parsable --dependency=afterok:$cool_job_id -p vgl --cpus-per-task=16 <<EOF
#!/bin/bash
#SBATCH --job-name=balance_${base_name}
#SBATCH --output=balance_${base_name}_%j.out
#SBATCH --cpus-per-task=16

# Load required modules or activate conda environment
source /ru-auth/local/home/risc_soft/miniconda3/etc/profile.d/conda.sh
conda activate open2c

# Balance the .cool file (ICE balancing)
cp "$cool_file" "$balanced_file"
cooler balance -p 16 --ignore-diags 1 "$balanced_file"
EOF
)

  # Step 3: Submit job to zoomify after Step 2 (balancing) is complete
  mcool_file="${output_dir}/${base_name}_500_bal_500N_igdiag1.mcool"
  sbatch --dependency=afterok:$balance_job_id -p vgl --cpus-per-task=16 <<EOF
#!/bin/bash
#SBATCH --job-name=zoomify_${base_name}
#SBATCH --output=zoomify_${base_name}_%j.out
#SBATCH --cpus-per-task=16

# Load required modules or activate conda environment
source /ru-auth/local/home/risc_soft/miniconda3/etc/profile.d/conda.sh
conda activate open2c

# Run cooler zoomify with balance and 500N resolutions
cooler zoomify \
    -p 16 \
    -r 500N \
    --balance \
    --balance-args '--nproc 16 --ignore-diags 1' \
    -o "$mcool_file" \
    "$balanced_file"
EOF

  # Log the outputs in the report
  echo "Processed: $pairs_file" >> $report_file
  echo "  500 bp .cool file: $cool_file" >> $report_file
  echo "  Balanced .cool file: $balanced_file" >> $report_file
  echo "  Multi-resolution .mcool file: $mcool_file" >> $report_file
  echo "----------------------------------" >> $report_file

done

# Finalize report
echo "Pipeline finished at: $(date)" >> $report_file
```
