## Supplementary material for "CAD-C: An engineered nuclease enables repair-free *in situ* proximity ligation and nucleosome-resolution chromosome walks in human cells": Custom code: append_bed_annotated_walks_with_fraglen_AB_BC_AC_midpointsep_v2_20250312.py

```
#!/usr/bin/env python3

import sys
import argparse
import gzip
from datetime import datetime

def process_chunk(lines, metrics):
    """
    Parse each line, compute new columns (length, midpoints, distances),
    update metrics, and return lines with appended columns.
    """
    out_lines = []
    for line in lines:
        line = line.rstrip("\n")  # strip only newline at end, keep leading spaces if any

        # If it's a comment line, pass it through unchanged
        if line.startswith("#"):
            out_lines.append(line + "\n")
            continue

        # If empty line, skip or keep if you like. 
        # We'll skip here (common in TSV).
        if not line.strip():
            continue

        fields = line.split("\t")

        # If the first row is the header row (i.e., starts with "readID"), we rewrite with new columns
        if fields[0] == "readID":
            # We'll produce a new header with additional columns at the end:
            new_header = fields + [
                "A_len", "B_len", "C_len",
                "A_mid", "B_mid", "C_mid",
                "AB_dist", "BC_dist", "AC_dist"
            ]
            out_lines.append("\t".join(new_header) + "\n")
            continue
        
        # Parse numeric columns (assuming you always have correct indices)
        A_start = int(fields[2])
        A_end   = int(fields[3])
        B_start = int(fields[6])
        B_end   = int(fields[7])
        C_start = int(fields[10])
        C_end   = int(fields[11])

        # Compute lengths (inclusive)
        A_len = A_end - A_start + 1
        B_len = B_end - B_start + 1
        C_len = C_end - C_start + 1

        # Compute midpoints
        A_mid = (A_start + A_end) // 2
        B_mid = (B_start + B_end) // 2
        C_mid = (C_start + C_end) // 2

        # Chroms
        A_chrom = fields[1]
        B_chrom = fields[5]
        C_chrom = fields[9]

        # Distances if on same chrom, otherwise empty string
        if A_chrom == B_chrom:
            AB_dist = abs(A_mid - B_mid)
            metrics["cis_AB"] += 1
        else:
            AB_dist = ""  # or use "NA" or "-1"
            metrics["trans_AB"] += 1

        if B_chrom == C_chrom:
            BC_dist = abs(B_mid - C_mid)
            metrics["cis_BC"] += 1
        else:
            BC_dist = ""
            metrics["trans_BC"] += 1

        if A_chrom == C_chrom:
            AC_dist = abs(A_mid - C_mid)
            metrics["cis_AC"] += 1
        else:
            AC_dist = ""
            metrics["trans_AC"] += 1
        
        metrics["total_rows"] += 1

        # Create the augmented line (we append 9 new columns)
        new_fields = fields + [
            str(A_len), str(B_len), str(C_len),
            str(A_mid), str(B_mid), str(C_mid),
            str(AB_dist), str(BC_dist), str(AC_dist),
        ]
        out_lines.append("\t".join(new_fields) + "\n")
    
    return out_lines

def main():
    parser = argparse.ArgumentParser(
        description="Plain Python script to add fragment length / midpoint / cis-distance columns to a 3-fragment TSV, retaining # comment lines."
    )
    parser.add_argument("--output", required=True,
                        help="Output .tsv.gz filename (gzipped).")
    parser.add_argument("--chunksize", type=int, default=1_000_000,
                        help="Number of lines to read at a time.")
    args = parser.parse_args()

    # Create a timestamp for the metrics file
    timestamp_str = datetime.now().strftime('%Y%m%d_%H%M%S')
    metrics_filename = f"metrics_{timestamp_str}.txt"

    # We'll store counters in a dictionary
    metrics = {
        "total_rows": 0,
        "cis_AB": 0,
        "trans_AB": 0,
        "cis_BC": 0,
        "trans_BC": 0,
        "cis_AC": 0,
        "trans_AC": 0,
    }

    # Open the output in gzip mode
    with gzip.open(args.output, "wt") as gz_out:
        # We'll read lines from stdin in chunks
        chunk_lines = []
        for line in sys.stdin:
            chunk_lines.append(line)
            if len(chunk_lines) >= args.chunksize:
                # Process this chunk
                out_chunk = process_chunk(chunk_lines, metrics)
                # Write them out
                gz_out.writelines(out_chunk)
                # Clear chunk
                chunk_lines = []
        
        # Process any leftover lines
        if chunk_lines:
            out_chunk = process_chunk(chunk_lines, metrics)
            gz_out.writelines(out_chunk)

    # Write metrics
    total_rows = metrics["total_rows"]
    cis_AB_count = metrics["cis_AB"]
    trans_AB_count = metrics["trans_AB"]
    cis_BC_count = metrics["cis_BC"]
    trans_BC_count = metrics["trans_BC"]
    cis_AC_count = metrics["cis_AC"]
    trans_AC_count = metrics["trans_AC"]

    with open(metrics_filename, "w") as m_out:
        m_out.write(f"Timestamp: {timestamp_str}\n")
        m_out.write(f"Total rows processed: {total_rows}\n")
        m_out.write(f"AB cis contacts: {cis_AB_count}\n")
        m_out.write(f"AB trans contacts: {trans_AB_count}\n")
        m_out.write(f"BC cis contacts: {cis_BC_count}\n")
        m_out.write(f"BC trans contacts: {trans_BC_count}\n")
        m_out.write(f"AC cis contacts: {cis_AC_count}\n")
        m_out.write(f"AC trans contacts: {trans_AC_count}\n")

    print(f"Done! Wrote augmented (gzipped) data to {args.output}")
    print(f"Wrote metrics to {metrics_filename}")

if __name__ == "__main__":
    main()
```
