## Supplementary material for "CAD-C: An engineered nuclease enables repair-free *in situ* proximity ligation and nucleosome-resolution chromosome walks in human cells": Custom code: density_norm_chromHMM_colorkey_CLI_v4_augmented_yaxscaleopts_20250313_log10yax10e-5.py

```
#!/usr/bin/env python3

import os
import argparse
import pandas as pd
import numpy as np
import matplotlib.pyplot as plt
import matplotlib
from datetime import datetime

matplotlib.rcParams['pdf.fonttype'] = 42
matplotlib.rcParams['ps.fonttype'] = 42

# Color dictionary for recognized states:
short_label_colors = {
    "ActiveEnhancer": (240/255, 180/255, 90/255),
    "DNase": (250/255, 230/255, 100/255),
    "Het": (90/255, 120/255, 180/255),
    "PromBiv": (180/255, 120/255, 150/255),
    "PromP": (180/255, 120/255, 150/255),
    "PromoterActive": (230/255, 60/255, 50/255),
    "ReprPC": (130/255, 130/255, 130/255),
    "TxStrong": (30/255, 140/255, 50/255),
    "TxWeak": (190/255, 200/255, 80/255),
    "WeakEnhancer": (250/255, 230/255, 100/255),
    "ZNF_Rpts": (90/255, 120/255, 180/255),
    "NA": (0.5, 0.5, 0.5)
}

def parse_chromhmm_label(raw_label: str) -> str:
    """
    Convert a full path like '/path/ChromHMM_ActiveEnhancer_merged.bed'
    => 'ActiveEnhancer'
    """
    if pd.isna(raw_label):
        return "NA"
    fname = os.path.basename(str(raw_label))
    fname = fname.replace(".bed", "")
    fname = fname.replace("_merged", "")
    if fname.startswith("ChromHMM_"):
        fname = fname[len("ChromHMM_"):]
    if not fname:
        return "NA"
    return fname

def boxcar_smooth_by_rows(df, window_size):
    """
    For each clean_label:
      1) Sort by distance
      2) Perform a row-based rolling mean on 'normalized_count' with the given window_size
         (center=True).
      3) Re-combine the results
    Then, in case of duplicates on (clean_label, distance), we average them.
    """
    # We'll do it group by group
    out_list = []
    for lbl, sub in df.groupby("clean_label", group_keys=False):
        # Sort by distance
        sub = sub.sort_values("distance")
        # row-based rolling
        rolled = (sub["normalized_count"]
                  .rolling(window=window_size, center=True, min_periods=1)
                  .mean())
        sub = sub.copy()
        sub["normalized_count"] = rolled
        out_list.append(sub)

    df_rolled = pd.concat(out_list, ignore_index=True)
    # Now ensure unique (clean_label, distance) by averaging if duplicates remain
    df_rolled = (
        df_rolled.groupby(["clean_label", "distance"], as_index=False)["normalized_count"]
        .mean()
    )
    return df_rolled

def generate_plots(df, pair_type, output_prefix, timestamp, log_type=None, bin_size=None):
    """
    Plots for pair_type (e.g. 'AB'), normalizing counts within each clean_label.
    """
    # 1) Subset by pair
    df_pair = df[df["pair"] == pair_type].dropna(subset=["clean_label"])
    if df_pair.empty:
        print(f"[INFO] No data for pair='{pair_type}'. Skipping.")
        return None, None

    # 2) Aggregate raw counts => sum by (clean_label, distance)
    df_grouped = df_pair.groupby(["clean_label", "distance"], as_index=False)["count"].sum()

    # 3) Normalize
    total_sums = df_grouped.groupby("clean_label")["count"].transform("sum")
    df_grouped["normalized_count"] = df_grouped["count"] / total_sums

    # 4) Smoothing if specified
    if bin_size is not None:
        df_grouped = boxcar_smooth_by_rows(df_grouped, bin_size)

    # 5) Pivot
    #   index='distance', columns='clean_label', values='normalized_count'
    #   => one row per distance, one column per label
    df_pivot = df_grouped.pivot(index="distance", columns="clean_label", values="normalized_count")
    if df_pivot.empty:
        print(f"[WARN] pivot empty for pair='{pair_type}'. No plot.")
        return None, None

    y_max = df_pivot.max().max()

    # If log scale => shift
    if log_type is not None:
        df_pivot += 1e-5
        y_max = df_pivot.max().max()

    # -- SUMMARY PLOT --
    plt.figure(figsize=(12, 6))
    lines_plotted = False

    for lbl in df_pivot.columns:
        series = df_pivot[lbl]
        if np.isnan(series).all():
            continue
        color = short_label_colors.get(lbl, (0.5, 0.5, 0.5))
        plt.plot(df_pivot.index, series, label=lbl, color=color, alpha=0.8)
        lines_plotted = True

    plt.xlabel("Distance (bp)")
    plt.ylabel("Relative Density (Normalized)")
    plt.title(f"{pair_type}: Relative Density by Bed State")

    if log_type == "log10":
        plt.yscale("log", base=10)
    elif log_type == "log2":
        plt.yscale("log", base=2)
    elif log_type == "ln":
        plt.yscale("log", base=np.e)

    if lines_plotted:
        plt.legend(title="Bed State", bbox_to_anchor=(1.05, 1), loc="upper left")
    plt.grid(True)

    summary_pdf = f"{output_prefix}_{pair_type}_summary_{log_type if log_type else 'linear'}_{bin_size if bin_size else 'no_smooth'}_{timestamp}.pdf"
    plt.savefig(summary_pdf, bbox_inches="tight")
    plt.close()

    # -- INDIVIDUAL PLOTS --
    num_labels = len(df_pivot.columns)
    cols = 3
    rows = (num_labels + cols - 1) // cols
    fig, axes = plt.subplots(rows, cols, figsize=(15, 5*rows))
    axes = axes.flatten()

    idx = 0
    for lbl in df_pivot.columns:
        ax = axes[idx]
        idx += 1

        series = df_pivot[lbl]
        if np.isnan(series).all():
            ax.set_title(f"{lbl} (no data)")
            ax.set_axis_off()
            continue

        color = short_label_colors.get(lbl, (0.5, 0.5, 0.5))
        ax.plot(df_pivot.index, series, color=color)
        ax.set_title(lbl)
        ax.set_xlabel("Distance (bp)")
        ax.set_ylabel("Relative Density")

        if log_type == "log10":
            ax.set_yscale("log", base=10)
        elif log_type == "log2":
            ax.set_yscale("log", base=2)
        elif log_type == "ln":
            ax.set_yscale("log", base=np.e)

        if log_type:
            ax.set_ylim(1e-5, y_max)
        else:
            ax.set_ylim(0, y_max)
        ax.grid(True)

    # Hide unused subplots
    for j in range(idx, len(axes)):
        fig.delaxes(axes[j])

    grid_pdf = f"{output_prefix}_{pair_type}_individual_{log_type if log_type else 'linear'}_{bin_size if bin_size else 'no_smooth'}_{timestamp}.pdf"
    plt.tight_layout()
    plt.savefig(grid_pdf, bbox_inches="tight")
    plt.close()

    return summary_pdf, grid_pdf

if __name__ == "__main__":
    parser = argparse.ArgumentParser(
        description="Histogram of genomic separations with optional rolling smoothing. "
                    "Handles duplicates by row-based rolling & averaging."
    )
    parser.add_argument("--csv", required=True,
                        help="CSV file with columns: pair, bed_label, distance, count")
    parser.add_argument("--log", choices=["log10", "log2", "ln"], default=None)
    parser.add_argument("--smooth", type=int, default=None,
                        help="Integer window-size for row-based rolling.")
    parser.add_argument("--x_min", type=float, default=None)
    parser.add_argument("--x_max", type=float, default=None)
    parser.add_argument("--output_prefix", default="output")
    args = parser.parse_args()

    # Load
    df = pd.read_csv(args.csv, compression="infer")

    # Distance filter
    if args.x_min is not None:
        df = df[df["distance"] >= args.x_min]
    if args.x_max is not None:
        df = df[df["distance"] <= args.x_max]
    if df.empty:
        raise ValueError("No data after filtering.")

    # Create a 'clean_label' from bed_label
    df["clean_label"] = df["bed_label"].apply(parse_chromhmm_label)

    # Unique timestamp
    timestamp = datetime.now().strftime("%Y%m%d_%H%M%S")

    # If your CSV has pairs like AB, BC, AC:
    pair_types = ["AB", "BC", "AC"]
    found_pairs = df["pair"].unique()
    for ptype in pair_types:
        if ptype in found_pairs:
            generate_plots(
                df=df,
                pair_type=ptype,
                output_prefix=args.output_prefix,
                timestamp=timestamp,
                log_type=args.log,
                bin_size=args.smooth
            )
        else:
            print(f"[INFO] pair='{ptype}' not found. Skipping.")

    print(f"Done! Check PDFs with timestamp={timestamp}")
```
