## Supplementary material for "CAD-C: An engineered nuclease enables repair-free *in situ* proximity ligation and nucleosome-resolution chromosome walks in human cells": Custom code: filt_zerolenB_flip_sort_dedup_split_cis_trans_v2_20250214.py

```
#!/usr/bin/env python3

import os
import sys
import gzip
import subprocess
import datetime
import argparse

##############################################################################
# 1) Check + Flip Walks
##############################################################################

def check_segments(A_start, A_end, B_start, B_end, C_start, C_end):
    """
    Return True if all segments have start <= end, else False.
    """
    if A_start > A_end:
        return False
    if B_start > B_end:
        return False
    if C_start > C_end:
        return False
    return True

def flip_walk_if_needed(fields):
    """
    fields = [
      readID,
      A_chrom, A_start, A_end, A_strand,
      B_chrom, B_start, B_end, B_strand,
      C_chrom, C_start, C_end, C_strand
    ]

    1) Filter out B fragments of length 0 (B_start == B_end).
    2) Check that each segment has start <= end (pre-flip).
    3) If A_start > C_start, swap A <-> C & flip A/B/C strands.
    4) Check again that each segment has start <= end (post-flip).

    Returns (new_fields, flipped_bool) or (None, False) if we skip.
    """

    readID   = fields[0]
    A_chrom  = fields[1]
    A_start  = int(fields[2])
    A_end    = int(fields[3])
    A_strand = fields[4]

    B_chrom  = fields[5]
    B_start  = int(fields[6])
    B_end    = int(fields[7])
    B_strand = fields[8]

    C_chrom  = fields[9]
    C_start  = int(fields[10])
    C_end    = int(fields[11])
    C_strand = fields[12]

    # 1) Filter out B fragments with length 0
    if B_start == B_end:
        return None, False  # skip

    # 2) Pre‐Flip Sanity Check
    if not check_segments(A_start, A_end, B_start, B_end, C_start, C_end):
        return None, False

    # 3) Flip if A_start > C_start
    flipped = False
    if A_start > C_start:
        flipped = True
        # Swap entire A <-> C
        (A_chrom, C_chrom)   = (C_chrom, A_chrom)
        (A_start, C_start)   = (C_start, A_start)
        (A_end,   C_end)     = (C_end,   A_end)
        (A_strand, C_strand) = (C_strand, A_strand)

        # Flip all strands
        A_strand = flip_strand(A_strand)
        B_strand = flip_strand(B_strand)
        C_strand = flip_strand(C_strand)

    # 4) Post‐Flip Sanity Check
    if not check_segments(A_start, A_end, B_start, B_end, C_start, C_end):
        return None, flipped

    new_fields = [
        readID,
        A_chrom, str(A_start), str(A_end), A_strand,
        B_chrom, str(B_start), str(B_end), B_strand,
        C_chrom, str(C_start), str(C_end), C_strand
    ]
    return new_fields, flipped

def flip_strand(s):
    if s == '+':
        return '-'
    elif s == '-':
        return '+'
    return s  # If something else is present, just return it

##############################################################################
# 2) Deduplicate
##############################################################################

def deduplicate_tsv(input_tsv_gz, output_dedup_tsv_gz):
    """
    Reads a .tsv.gz (sorted by A_chrom, A_start),
    writes a deduplicated version to output_dedup_tsv_gz.

    Returns (n_in, n_out, duplicates).
    """

    if input_tsv_gz.endswith(".gz"):
        inf = gzip.open(input_tsv_gz, "rt")
    else:
        inf = open(input_tsv_gz, "r")

    if output_dedup_tsv_gz.endswith(".gz"):
        outf = gzip.open(output_dedup_tsv_gz, "wt")
    else:
        outf = open(output_dedup_tsv_gz, "w")

    header = next(inf).rstrip("\n")
    outf.write(header + "\n")

    last_key = None
    n_in = 0
    n_out = 0

    for line in inf:
        line_str = line.rstrip("\n")
        fields = line_str.split("\t")
        # dedup key = everything except readID
        key = tuple(fields[1:])

        n_in += 1
        if key != last_key:
            outf.write(line_str + "\n")
            n_out += 1
            last_key = key

    inf.close()
    outf.close()

    duplicates = n_in - n_out
    return (n_in, n_out, duplicates)

##############################################################################
# 3) Split cis/trans
##############################################################################

def split_cis_trans(input_dedup_tsv_gz, output_cis_tsv_gz, output_trans_tsv_gz):
    """
    Split the deduplicated .tsv.gz into cis (A/B/C same chrom) and trans (otherwise).
    Returns (n_in, n_cis, n_trans).
    """

    if input_dedup_tsv_gz.endswith(".gz"):
        inf = gzip.open(input_dedup_tsv_gz, "rt")
    else:
        inf = open(input_dedup_tsv_gz, "r")

    if output_cis_tsv_gz.endswith(".gz"):
        out_cis = gzip.open(output_cis_tsv_gz, "wt")
    else:
        out_cis = open(output_cis_tsv_gz, "w")

    if output_trans_tsv_gz.endswith(".gz"):
        out_trans = gzip.open(output_trans_tsv_gz, "wt")
    else:
        out_trans = open(output_trans_tsv_gz, "w")

    header = next(inf).rstrip("\n")
    out_cis.write(header + "\n")
    out_trans.write(header + "\n")

    n_in = 0
    n_cis = 0
    n_trans = 0

    for line in inf:
        line_str = line.rstrip("\n")
        fields = line_str.split("\t")
        # [readID, A_chrom, A_start, A_end, A_strand,
        #  B_chrom, B_start, B_end, B_strand,
        #  C_chrom, C_start, C_end, C_strand]
        A_chrom = fields[1]
        B_chrom = fields[5]
        C_chrom = fields[9]

        n_in += 1
        if A_chrom == B_chrom == C_chrom:
            out_cis.write(line_str + "\n")
            n_cis += 1
        else:
            out_trans.write(line_str + "\n")
            n_trans += 1

    inf.close()
    out_cis.close()
    out_trans.close()
    return (n_in, n_cis, n_trans)

##############################################################################
# 4) Main Pipeline
##############################################################################

def parse_args():
    parser = argparse.ArgumentParser(
        description="Flip (if needed) each ABC walk so A_start < C_start, exclude B-length=0, verify segments, then sort, deduplicate, and split into cis/trans."
    )
    parser.add_argument("--input", required=True,
                        help="Input flattened ABC .tsv.gz file.")
    parser.add_argument("--output-dir", required=True,
                        help="Directory to write all outputs.")
    parser.add_argument("--prefix", required=True,
                        help="File prefix for output files (e.g. 'MyFlattenedABC').")
    parser.add_argument("--threads", type=int, default=1,
                        help="Number of threads for GNU sort (default=1).")
    parser.add_argument("--skip-on-error", action="store_true",
                        help="Skip (omit) any row that fails sanity checks, rather than aborting.")
    return parser.parse_args()

def main():
    args = parse_args()
    os.makedirs(args.output_dir, exist_ok=True)

    flipped_tsv = os.path.join(args.output_dir, f"{args.prefix}.flipped.tsv.gz")
    sorted_tsv  = os.path.join(args.output_dir, f"{args.prefix}.sorted.tsv.gz")
    dedup_tsv   = os.path.join(args.output_dir, f"{args.prefix}.sorted.dedup.tsv.gz")
    cis_tsv     = os.path.join(args.output_dir, f"{args.prefix}.sorted.dedup.cis.tsv.gz")
    trans_tsv   = os.path.join(args.output_dir, f"{args.prefix}.sorted.dedup.trans.tsv.gz")

    timestamp_str = datetime.datetime.now().strftime("%Y%m%d_%H%M%S")
    metrics_file = os.path.join(args.output_dir, f"{args.prefix}.metrics_{timestamp_str}.txt")

    ########################################################################
    # A) Flip Step (and filter out B-length=0)
    ########################################################################

    n_in_flips = 0
    n_flipped = 0
    n_sanity_errors = 0
    n_bzero = 0  # count B fragment length=0

    # Read input, skip the header, flip if needed, write to flipped_tsv
    if args.input.endswith(".gz"):
        inf = gzip.open(args.input, "rt")
    else:
        inf = open(args.input, "r")

    if flipped_tsv.endswith(".gz"):
        outf = gzip.open(flipped_tsv, "wt")
    else:
        outf = open(flipped_tsv, "w")

    # Write the same header line
    header_line = next(inf).rstrip("\n")
    outf.write(header_line + "\n")

    for line in inf:
        line_str = line.rstrip("\n")
        fields = line_str.split("\t")
        if len(fields) < 13:
            # skip malformed
            n_sanity_errors += 1
            continue

        n_in_flips += 1
        new_fields, did_flip = flip_walk_if_needed(fields)

        if new_fields is None:
            # Could be B=0 or segment sanity fail
            # We'll see which:
            B_start = int(fields[6])
            B_end   = int(fields[7])
            if B_start == B_end:
                n_bzero += 1
            else:
                n_sanity_errors += 1

            if not args.skip_on_error:
                # Hard fail
                print(f"ERROR: Found B=0 or invalid segment. Offending line:\n{line_str}", file=sys.stderr)
                sys.exit(1)
            else:
                # skip the line
                continue

        if did_flip:
            n_flipped += 1

        outf.write("\t".join(new_fields) + "\n")

    inf.close()
    outf.close()

    ########################################################################
    # B) Sort by (A_chrom, A_start) with version sort on A_chrom
    ########################################################################
    tmp_sorted = os.path.join(args.output_dir, f"{args.prefix}.sorted.NO_HEADER.tmp")

    # We'll skip the first line of flipped_tsv, then do:
    #    -k2,2V => version sort on column 2 (A_chrom) 
    #    -k3,3n => numeric sort on column 3 (A_start)
    sort_cmd = [
        "bash", "-c",
        (
            "set -o pipefail; "
            f"(tail -n +2 <(zcat -f '{flipped_tsv}')) "
            "| sort -t $'\\t' -k2,2V -k3,3n "
            f"--parallel={args.threads} "
            f"> '{tmp_sorted}'"
        )
    ]

    ret = subprocess.run(" ".join(sort_cmd), shell=True, executable="/bin/bash")
    if ret.returncode != 0:
        print("ERROR: Sorting step failed.", file=sys.stderr)
        sys.exit(1)

    # Re-add header, compress
    with open(tmp_sorted, "r") as ftmp:
        sorted_lines = ftmp.read()
    os.remove(tmp_sorted)

    with gzip.open(sorted_tsv, "wt") as outf2:
        outf2.write(header_line + "\n")
        outf2.write(sorted_lines)

    ########################################################################
    # C) Deduplicate
    ########################################################################
    n_in, n_out, duplicates = deduplicate_tsv(sorted_tsv, dedup_tsv)

    ########################################################################
    # D) Split cis/trans
    ########################################################################
    n_split_in, n_cis, n_trans = split_cis_trans(dedup_tsv, cis_tsv, trans_tsv)

    ########################################################################
    # E) Metrics
    ########################################################################
    now_str = datetime.datetime.now().strftime("%Y-%m-%d %H:%M:%S")
    with open(metrics_file, "w") as mf:
        mf.write(f"Timestamp: {now_str}\n")
        mf.write(f"Input file: {os.path.abspath(args.input)}\n\n")

        mf.write(f"Flipped file:   {os.path.abspath(flipped_tsv)}\n")
        mf.write(f"  Lines read (excluding header): {n_in_flips}\n")
        mf.write(f"  Lines flipped: {n_flipped} "
                 f"({(n_flipped / n_in_flips * 100) if n_in_flips else 0:.2f}% of input)\n")
        mf.write(f"  B=0 fragments skipped: {n_bzero} "
                 f"({(n_bzero / n_in_flips * 100) if n_in_flips else 0:.2f}% of input)\n")
        mf.write(f"  Other sanity errors: {n_sanity_errors} "
                 f"({(n_sanity_errors / n_in_flips * 100) if n_in_flips else 0:.2f}% of input)\n\n")

        mf.write(f"Sorted file:    {os.path.abspath(sorted_tsv)}\n")
        mf.write(f"Deduplicated:   {os.path.abspath(dedup_tsv)}\n")
        mf.write(f"CIS file:       {os.path.abspath(cis_tsv)}\n")
        mf.write(f"TRANS file:     {os.path.abspath(trans_tsv)}\n\n")

        mf.write(f"Lines read from sorted file (excluding header): {n_in}\n")
        mf.write(f"Lines after dedup (excluding header): {n_out}\n")
        mf.write(f"Duplicates removed: {duplicates} "
                 f"({(duplicates / n_in * 100) if n_in else 0:.2f}% of input)\n\n")

        mf.write(f"Lines read for splitting (should match dedup output): {n_split_in}\n")
        mf.write(f"Cis lines: {n_cis}\n")
        mf.write(f"Trans lines: {n_trans}\n")
        if n_split_in > 0:
            mf.write(f"Percent cis: {n_cis / n_split_in * 100:.2f}%\n")
            mf.write(f"Percent trans: {n_trans / n_split_in * 100:.2f}%\n")

    print("===================================================")
    print("Pipeline Complete!")
    print(f"  Flipped file:   {flipped_tsv} (read {n_in_flips}, flipped {n_flipped}, B=0 {n_bzero}, other errors {n_sanity_errors})")
    print(f"  Sorted file:    {sorted_tsv}")
    print(f"  Deduplicated:   {dedup_tsv} (duplicates removed: {duplicates})")
    print(f"  CIS: {cis_tsv} ({n_cis} lines), TRANS: {trans_tsv} ({n_trans} lines)")
    print(f"  Metrics:        {metrics_file}")
    print("===================================================")

if __name__ == "__main__":
    main()
```
