## Supplementary material for "CAD-C: An engineered nuclease enables repair-free *in situ* proximity ligation and nucleosome-resolution chromosome walks in human cells": Custom code: merge_flat_flipped_sorted_20250214.py

```
#!/usr/bin/env python3

import sys
import os
import gzip
import argparse
import datetime
import heapq

def parse_args():
    parser = argparse.ArgumentParser(
        description="Merge multiple flipped+sorted+deduped ABC .tsv(.gz) files into one, preserving natural/“version” order on A_chrom and numeric order on A_start."
    )
    parser.add_argument("--inputs", nargs="+", required=True,
                        help="Two or more sorted .tsv(.gz) files (same header).")
    parser.add_argument("--output", required=True,
                        help="Output merged .tsv.gz file.")
    parser.add_argument("--metrics", required=True,
                        help="Path to a .txt file containing merge metrics.")
    return parser.parse_args()

###############################################################################
# Version Sort Key for A_chrom
###############################################################################
# We want a function that turns "chr1" -> (1, False), "chrX" -> (100001, False)
# so numeric < X < Y < M < etc. Then we keep A_start numeric after that.
# If your actual files might contain "chrUn_random" or scaffolds, you'll want
# to adapt this logic. This is a minimal example.

def version_sort_chrom(chrom):
    """
    Convert A_chrom string into a tuple that Python can sort naturally:
    e.g. 'chr1' -> (1, 0)
         'chr2' -> (2, 0)
         'chr10'-> (10, 0)
         'chrX' -> (100001, 0)
         'chrY' -> (100002, 0)
         'chrM' -> (100003, 0)
    If we can't parse numeric, we assign some big code for leftover.
    """
    # remove "chr" prefix if present
    if chrom.startswith("chr"):
        chrom_str = chrom[3:]
    else:
        chrom_str = chrom

    # handle special cases
    if chrom_str == "X":
        return (100001, 0)
    elif chrom_str == "Y":
        return (100002, 0)
    elif chrom_str in ("M", "MT"):  # mtDNA
        return (100003, 0)

    # else try numeric
    # if not numeric, push it “after” main autosomes
    try:
        val = int(chrom_str)
        return (val, 0)
    except ValueError:
        # fallback for random/other scaffolds
        # e.g. "14_KI270723v1_random"
        # We'll store them at big code 200000, then lexicographically after that
        return (200000, chrom_str)

###############################################################################
# Parsing each line -> Merge key
###############################################################################
def parse_line_to_key(line):
    """
    Given a flattened ABC line:
      readID, A_chrom, A_start, A_end, A_strand,
              B_chrom, B_start, B_end, B_strand,
              C_chrom, C_start, C_end, C_strand
    we want a key = (version_sort_chrom(A_chrom), A_start).
    Returns ((chrom_key, Astart), line).
    """
    fields = line.rstrip("\n").split("\t")
    if len(fields) < 13:
        # skip malformed lines
        return None

    A_chrom = fields[1]
    try:
        A_start = int(fields[2])
    except ValueError:
        # skip malformed
        return None

    chrom_key = version_sort_chrom(A_chrom)
    return ((chrom_key, A_start), line)

###############################################################################
# Generator to read lines from each input file (skipping the header)
###############################################################################
def gen_lines_sorted(file_path, counts_dict):
    """
    Opens file_path (.tsv or .tsv.gz),
    reads+skips the header,
    then yields ((chrom_key, A_start), line_str).
    increments counts_dict[file_path] for each valid line.
    """
    # open
    if file_path.endswith(".gz"):
        fobj = gzip.open(file_path, "rt")
    else:
        fobj = open(file_path, "r")

    # skip header
    header_line = next(fobj, None)

    for raw_line in fobj:
        parsed = parse_line_to_key(raw_line)
        if parsed is None:
            # skip malformed
            continue
        (sort_key, line_str) = parsed
        counts_dict[file_path] += 1
        yield (sort_key, line_str)

    fobj.close()

###############################################################################
# Main merging function
###############################################################################
def main():
    args = parse_args()

    input_files = args.inputs
    if len(input_files) < 2:
        print("ERROR: Need at least 2 input files to merge.", file=sys.stderr)
        sys.exit(1)

    # We'll store line counts in a dict
    line_counts = {f: 0 for f in input_files}

    # We also want to store a single header. We'll just copy the header from
    # the first file to the output. We assume all files share the same header.
    # If you prefer a more robust approach, you'd read all headers + compare them.
    first_file = input_files[0]
    if first_file.endswith(".gz"):
        with gzip.open(first_file, "rt") as test_in:
            header_line = next(test_in, "").rstrip("\n")
    else:
        with open(first_file, "r") as test_in:
            header_line = next(test_in, "").rstrip("\n")

    # Prepare iterators, skipping the header in each
    iterators = []
    for f in input_files:
        iterators.append(gen_lines_sorted(f, line_counts))

    # Merge them with heapq.merge, using the (chrom_key, A_start) as the sorting key
    # All input files are *already* sorted in the same manner, so this is effectively
    # a standard multi-way merge. If there's any mismatch in sort order, this
    # will preserve the order specified by the key function.
    merged_iter = heapq.merge(*iterators, key=lambda x: x[0])

    # Open output
    # We'll write a .gz file either way
    out_f = gzip.open(args.output, "wt") if args.output.endswith(".gz") \
            else open(args.output, "w")

    # Write the single header
    out_f.write(header_line + "\n")

    # Now stream
    merged_count = 0
    for (sort_key, line_str) in merged_iter:
        out_f.write(line_str)
        merged_count += 1

    out_f.close()

    # Build metrics
    now_str = datetime.datetime.now().strftime("%Y-%m-%d %H:%M:%S")
    with open(args.metrics, "w") as mf:
        mf.write(f"Timestamp: {now_str}\n\n")

        total_in = 0
        for f in input_files:
            c = line_counts[f]
            total_in += c
            mf.write(f"Input file: {os.path.abspath(f)} => {c} lines\n")
        mf.write("\n")

        mf.write(f"Total lines merged in output (excluding header): {merged_count}\n\n")
        mf.write(f"Output file: {os.path.abspath(args.output)}\n")
        mf.write(f"Metrics file: {os.path.abspath(args.metrics)}\n")

    print("=======================================")
    print("Merge complete!")
    for f in input_files:
        print(f"  {f}: {line_counts[f]} lines")
    print(f"Combined => {args.output} ({merged_count} lines)")
    print(f"Metrics => {args.metrics}")
    print("=======================================")

if __name__ == "__main__":
    main()
```
