## Supplementary material for "CAD-C: An engineered nuclease enables repair-free *in situ* proximity ligation and nucleosome-resolution chromosome walks in human cells": Custom code: parse_AB_BC_AC_seps_by_leftbedstate_v2_20250312.py

```
#!/usr/bin/env python3

import sys
import argparse
import gzip
import re
from collections import defaultdict
from datetime import datetime

def main():
    parser = argparse.ArgumentParser(
        description="Collect single-bp resolution midpoint separations (AB, BC, AC) from precomputed columns "
                    "22,23,24, keyed by the bed annotation of the 'origin' fragment "
                    "(A for AB/AC, B for BC). Skips trans if the column is empty/NA."
    )
    parser.add_argument("--sep-out", required=True,
                        help="Output CSV or CSV.gz with columns (pair,bed_label,distance,count).")
    parser.add_argument("--bed-map-out", default="bed_map_info.txt",
                        help="Where to write the bed_index => label lines from # bed_index=... comments.")
    parser.add_argument("--retain-comments", action="store_true",
                        help="If set, print # lines and header row to stdout.")
    parser.add_argument("--chunksize", type=int, default=1_000_000,
                        help="Number of lines to read at a time (avoids large memory usage).")
    parser.add_argument("--gz-output", action="store_true",
                        help="If set, write sep-out in gzip format on-the-fly (sep-out must end with .gz).")
    args = parser.parse_args()

    # bed_map: e.g. "bed_0" => "/path/to/file"
    bed_map = {}

    # histograms[(pair, bed_label)][distance] = count
    # pair ∈ {"AB", "BC", "AC"}
    histograms = defaultdict(lambda: defaultdict(int))

    bed_index_pattern = re.compile(r"^#\s*bed_index\s*=\s*(\d+)\s*=>\s*(.*)")
    chunk_buffer = []

    def process_chunk(lines):
        for line in lines:
            line_stripped = line.rstrip("\n")

            # 1) bed_index=... => ...
            m = bed_index_pattern.match(line_stripped)
            if m:
                idx_num = m.group(1)
                path_label = m.group(2).strip()
                bed_map[f"bed_{idx_num}"] = path_label
                if args.retain_comments:
                    print(line_stripped)
                continue

            # 2) Other comment lines
            if line_stripped.startswith("#"):
                if args.retain_comments:
                    print(line_stripped)
                continue

            # 3) Possibly the header
            if line_stripped.startswith("readID"):
                if args.retain_comments:
                    print(line_stripped)
                continue

            # 4) Data line
            fields = line_stripped.split("\t")
            # We expect columns up to 24 => index 24 (0-based)
            if len(fields) < 25:
                continue

            A_bed = fields[13]
            B_bed = fields[14]
            # For AC, also A_bed (since A is the 'origin')
            # If you wanted to use C_bed, you'd adjust logic, but typically we follow your earlier approach.

            # Convert bed_x => label if known
            def get_bed_label(x):
                if x == "NA":
                    return "NA"
                return bed_map.get(x, x)

            A_label = get_bed_label(A_bed)
            B_label = get_bed_label(B_bed)

            AB_str = fields[22].strip()
            BC_str = fields[23].strip()
            AC_str = fields[24].strip()

            # If AB_str is not empty => cis => store it under ("AB", A_label)
            if AB_str not in ("", "NA"):
                try:
                    dist_AB = int(AB_str)
                    histograms[("AB", A_label)][dist_AB] += 1
                except ValueError:
                    pass

            # If BC_str is not empty => cis => store it under ("BC", B_label)
            if BC_str not in ("", "NA"):
                try:
                    dist_BC = int(BC_str)
                    histograms[("BC", B_label)][dist_BC] += 1
                except ValueError:
                    pass

            # If AC_str is not empty => cis => store it under ("AC", A_label)
            if AC_str not in ("", "NA"):
                try:
                    dist_AC = int(AC_str)
                    histograms[("AC", A_label)][dist_AC] += 1
                except ValueError:
                    pass

    # chunk reading
    for line in sys.stdin:
        chunk_buffer.append(line)
        if len(chunk_buffer) >= args.chunksize:
            process_chunk(chunk_buffer)
            chunk_buffer.clear()

    # leftover
    if chunk_buffer:
        process_chunk(chunk_buffer)
        chunk_buffer.clear()

    # Write bed_map
    with open(args.bed_map_out, "w") as bm:
        for k in sorted(bed_map.keys()):
            bm.write(f"{k}\t{bed_map[k]}\n")

    # Write the final histogram
    # If user gave --gz-output, we do gzip.open; else normal open
    if args.gz_output:
        import gzip
        with gzip.open(args.sep_out, "wt") as out_f:
            out_f.write("pair,bed_label,distance,count\n")
            for (pair_name, bed_lbl), dist_dict in histograms.items():
                for dist_val, ct in dist_dict.items():
                    out_f.write(f"{pair_name},{bed_lbl},{dist_val},{ct}\n")
    else:
        with open(args.sep_out, "w") as out_f:
            out_f.write("pair,bed_label,distance,count\n")
            for (pair_name, bed_lbl), dist_dict in histograms.items():
                for dist_val, ct in dist_dict.items():
                    out_f.write(f"{pair_name},{bed_lbl},{dist_val},{ct}\n")

    print(f"Done! Wrote separation histogram to {args.sep_out}")
    print(f"Bed map => {args.bed_map_out}")

if __name__ == "__main__":
    main()
```
