## Supplementary material for "CAD-C: An engineered nuclease enables repair-free *in situ* proximity ligation and nucleosome-resolution chromosome walks in human cells": Custom code: parse_frag_lens_by_bed_v2_20250312.py

```
#!/usr/bin/env python3

import sys
import argparse
import gzip
import re
from collections import defaultdict
from datetime import datetime

def main():
    parser = argparse.ArgumentParser(
        description="Collect single-bp resolution fragment-length histograms for A_len/B_len/C_len columns, "
                    "keyed by the bed annotation of each fragment (A_bed, B_bed, C_bed). "
                    "Assumes columns 16,17,18 are precomputed A_len,B_len,C_len."
    )
    parser.add_argument("--hist-out", required=True,
                        help="Path to CSV file with columns (position,bed_label,length,count).")
    parser.add_argument("--bed-map-out", default="bed_map_info.txt",
                        help="Where to write the bed_index => label lines (from # bed_index=...).")
    parser.add_argument("--retain-comments", action="store_true",
                        help="If set, prints # lines and the header row to stdout.")
    parser.add_argument("--chunksize", type=int, default=1_000_000,
                        help="Number of lines to read at a time.")
    args = parser.parse_args()

    # Dictionary e.g. bed_map["bed_0"] = "/path/to/ChromHMM_ActiveEnhancer_merged.bed"
    bed_map = {}

    # Histograms stored as: histograms[(pos, bed_label)][length] = count
    # e.g. histograms[("A", "bed_0")][200] = 15
    histograms = defaultdict(lambda: defaultdict(int))

    # Regex for lines like "# bed_index=0 => /some/path"
    bed_index_pattern = re.compile(r"^#\s*bed_index\s*=\s*(\d+)\s*=>\s*(.*)")

    chunk_buffer = []

    def process_chunk(lines):
        for line in lines:
            line_stripped = line.rstrip("\n")

            # 1) Check if it's a # bed_index=... => ...
            match = bed_index_pattern.match(line_stripped)
            if match:
                idx_num = match.group(1)
                path_label = match.group(2).strip()
                bed_map[f"bed_{idx_num}"] = path_label
                if args.retain_comments:
                    print(line_stripped)
                continue

            # 2) Other # lines
            if line_stripped.startswith("#"):
                if args.retain_comments:
                    print(line_stripped)
                continue

            # 3) Possibly the header
            if line_stripped.startswith("readID"):
                if args.retain_comments:
                    print(line_stripped)
                continue

            # 4) Actual data line
            fields = line_stripped.split("\t")
            # We expect at least 19 columns: up to col 18 (0-based) => A_len,B_len,C_len are 16,17,18
            if len(fields) < 19:
                continue

            A_bed = fields[13]
            B_bed = fields[14]
            C_bed = fields[15]

            # Convert e.g. "bed_0" => descriptive path if known
            def get_bed_label(x):
                if x == "NA":
                    return "NA"
                return bed_map.get(x, x)

            A_label = get_bed_label(A_bed)
            B_label = get_bed_label(B_bed)
            C_label = get_bed_label(C_bed)

            A_len_str = fields[16].strip()
            B_len_str = fields[17].strip()
            C_len_str = fields[18].strip()

            # If these columns might be "" or "NA" for missing data, skip those
            # Typically they should be integers if precomputed
            if A_len_str not in ("", "NA"):
                try:
                    A_len = int(A_len_str)
                    histograms[("A", A_label)][A_len] += 1
                except ValueError:
                    pass

            if B_len_str not in ("", "NA"):
                try:
                    B_len = int(B_len_str)
                    histograms[("B", B_label)][B_len] += 1
                except ValueError:
                    pass

            if C_len_str not in ("", "NA"):
                try:
                    C_len = int(C_len_str)
                    histograms[("C", C_label)][C_len] += 1
                except ValueError:
                    pass
        # end for

    # -- read lines in chunks
    for line in sys.stdin:
        chunk_buffer.append(line)
        if len(chunk_buffer) >= args.chunksize:
            process_chunk(chunk_buffer)
            chunk_buffer.clear()

    # leftover
    if chunk_buffer:
        process_chunk(chunk_buffer)
        chunk_buffer.clear()

    # Write the bed_map
    with open(args.bed_map_out, "w") as bm:
        for k in sorted(bed_map.keys()):
            bm.write(f"{k}\t{bed_map[k]}\n")

    # Write final histogram CSV: position,bed_label,length,count
    with open(args.hist_out, "w") as out_f:
        out_f.write("position,bed_label,length,count\n")
        for (pos, lbl), length_dict in histograms.items():
            for length_val, ct in length_dict.items():
                out_f.write(f"{pos},{lbl},{length_val},{ct}\n")

    print(f"Done! Wrote fragment length histograms to {args.hist_out}.")
    print(f"Bed map => {args.bed_map_out}.")

if __name__ == "__main__":
    main()
```
