## Supplementary material for "CAD-C: An engineered nuclease enables repair-free *in situ* proximity ligation and nucleosome-resolution chromosome walks in human cells": Custom code: split_flattened_ABC_by_order_classes_1_20250218.py

```
#!/usr/bin/env python3

import argparse
import gzip
import csv
import sys
from pathlib import Path

def process_chunk(rows,
                  out_abc,
                  out_acb,
                  out_bac,
                  total_in,
                  abc_count,
                  acb_count,
                  bac_count):
    """
    Classify each row in 'rows' and write it to the correct output file.
    Update counters and return them.
    """
    for row in rows:
        # row layout (13 columns):
        #  0: readID
        #  1: A_chrom    2: A_start   3: A_end     4: A_strand
        #  5: B_chrom    6: B_start   7: B_end     8: B_strand
        #  9: C_chrom   10: C_start  11: C_end    12: C_strand

        total_in += 1  # Count lines (excluding header)

        # Convert the relevant columns to integers
        A_start = int(row[2])
        B_start = int(row[6])
        C_start = int(row[10])

        # Classify
        if A_start < B_start < C_start:
            # ABC
            out_abc.write("\t".join(row) + "\n")
            abc_count += 1
        elif A_start < C_start < B_start:
            # ACB
            out_acb.write("\t".join(row) + "\n")
            acb_count += 1
        elif B_start < A_start < C_start:
            # BAC
            out_bac.write("\t".join(row) + "\n")
            bac_count += 1
        else:
            # If your data always ensures A_start < C_start
            # you may not expect other orders. We simply skip or you could raise an error.
            pass

    return total_in, abc_count, acb_count, bac_count


def split_3way_walks(input_tsv_gz, output_dir, out_prefix, chunk_size=100_000):
    """
    Read a large gzipped TSV of "A-B-C" chromosome walks and split lines
    into three order classes (ABC, ACB, BAC), writing each class to its
    own gzipped file.

    Parameters
    ----------
    input_tsv_gz : Path or str
        Path to the input .tsv.gz file.
    output_dir : Path or str
        Directory to which output files will be written.
    out_prefix : str
        Prefix for the three output files. For example, if out_prefix="ABC_split",
        the output will be:
            - ABC_split.ABC.tsv.gz
            - ABC_split.ACB.tsv.gz
            - ABC_split.BAC.tsv.gz
    chunk_size : int, optional
        Number of lines to read in each iteration (streaming). Default: 100,000.
    """

    output_dir = Path(output_dir)
    output_dir.mkdir(parents=True, exist_ok=True)

    # Construct output file paths
    path_abc = output_dir / f"{out_prefix}.ABC.tsv.gz"
    path_acb = output_dir / f"{out_prefix}.ACB.tsv.gz"
    path_bac = output_dir / f"{out_prefix}.BAC.tsv.gz"

    total_in = 0
    abc_count = 0
    acb_count = 0
    bac_count = 0

    # Open input and outputs
    with gzip.open(input_tsv_gz, mode="rt") as infile, \
         gzip.open(path_abc, mode="wt") as out_abc, \
         gzip.open(path_acb, mode="wt") as out_acb, \
         gzip.open(path_bac, mode="wt") as out_bac:

        reader = csv.reader(infile, delimiter="\t")

        # Read header
        header = next(reader)
        # Write header to each output
        out_abc.write("\t".join(header) + "\n")
        out_acb.write("\t".join(header) + "\n")
        out_bac.write("\t".join(header) + "\n")

        # Stream through in chunks
        chunk = []
        for row in reader:
            chunk.append(row)
            if len(chunk) >= chunk_size:
                total_in, abc_count, acb_count, bac_count = process_chunk(
                    chunk, out_abc, out_acb, out_bac,
                    total_in, abc_count, acb_count, bac_count
                )
                chunk = []  # Reset chunk

        # Process any leftovers
        if chunk:
            total_in, abc_count, acb_count, bac_count = process_chunk(
                chunk, out_abc, out_acb, out_bac,
                total_in, abc_count, acb_count, bac_count
            )

    # Print summary metrics
    print(f"Done processing {input_tsv_gz}")
    print(f"Total lines (excluding header): {total_in}")
    print(f"  -> ABC: {abc_count}")
    print(f"  -> ACB: {acb_count}")
    print(f"  -> BAC: {bac_count}")


def main():
    parser = argparse.ArgumentParser(
        description="Split a large ABC-type walks .tsv.gz file into three order classes (ABC, ACB, BAC)."
    )
    parser.add_argument(
        "-i", "--input", required=True,
        help="Path to the input .tsv.gz file."
    )
    parser.add_argument(
        "-o", "--outdir", required=True,
        help="Path to the output directory."
    )
    parser.add_argument(
        "-p", "--prefix", required=True,
        help="Prefix for output files, e.g. 'myABC'."
    )
    parser.add_argument(
        "-c", "--chunk_size", type=int, default=100_000,
        help="Number of lines to read at a time (default: 100000)."
    )

    args = parser.parse_args()

    # Call the main function
    split_3way_walks(
        input_tsv_gz=args.input,
        output_dir=args.outdir,
        out_prefix=args.prefix,
        chunk_size=args.chunk_size
    )

if __name__ == "__main__":
    main()
```
