## Supplementary material for "CAD-C: An engineered nuclease enables repair-free *in situ* proximity ligation and nucleosome-resolution chromosome walks in human cells": Custom code: truncate_sep_hist_table_1_20250312.py

```
#!/usr/bin/env python3

import sys
import argparse
import gzip

def main():
    parser = argparse.ArgumentParser(
        description="Truncate a separation CSV(.gz) to rows with distance <= 1500, preserving comments and header."
    )
    parser.add_argument("--input", required=True,
                        help="Input file: a CSV or CSV.gz with columns: pair,bed_label,distance,count.")
    parser.add_argument("--output", required=True,
                        help="Output file (e.g. short_1500.csv.gz). If it ends with .gz, it will be gzipped.")
    parser.add_argument("--maxdist", type=int, default=1500,
                        help="Maximum distance cutoff (default=1500). Rows with distance <= this are kept.")
    args = parser.parse_args()

    # Decide how to open input (possibly gzipped)
    if args.input.endswith(".gz"):
        in_f = gzip.open(args.input, "rt")
    else:
        in_f = open(args.input, "r")

    # Decide how to open output
    if args.output.endswith(".gz"):
        out_f = gzip.open(args.output, "wt")
    else:
        out_f = open(args.output, "w")

    with in_f, out_f:
        header_seen = False

        for line in in_f:
            line_stripped = line.rstrip("\n")

            # Preserve comment lines that begin with '#'
            if line_stripped.startswith("#"):
                out_f.write(line)
                continue

            # Possibly the header row: "pair,bed_label,distance,count"
            if not header_seen and line_stripped.startswith("pair,"):
                # Write the header row as-is
                out_f.write(line)
                header_seen = True
                continue

            # For data rows, parse the distance
            # Only if we've seen the header do we expect data
            if header_seen and line_stripped:
                # e.g. "AB,ChromHMM_ActiveEnhancer_merged.bed,123,456"
                cols = line_stripped.split(",")
                if len(cols) < 4:
                    # skip malformed lines
                    continue

                try:
                    distance_val = int(cols[2])  # third column is 'distance'
                except ValueError:
                    # skip if distance is not an integer
                    continue

                # Keep row if distance <= maxdist
                if distance_val <= args.maxdist:
                    out_f.write(line)

    print(f"Done! Wrote truncated file to {args.output}")

if __name__ == "__main__":
    main()
```
