## Supplementary material for "CAD-C: An engineered nuclease enables repair-free *in situ* proximity ligation and nucleosome-resolution chromosome walks in human cells": Custom code: filter_consequent_12_ABC_walks_20250211.sh

```
#!/bin/bash
#SBATCH --job-name=filter_groups
#SBATCH --output=filter_groups_%A_%a.out
#SBATCH --error=filter_groups_%A_%a.err
#SBATCH --ntasks=1
#SBATCH --cpus-per-task=4
#SBATCH --array=1-4  # We have 4 input files

# Load pigz if needed (depends on your HPC environment).
# module load pigz

# Use gawk (or default awk) if mawk is not available
AWK_CMD=gawk

# Directory containing your input files:
INPUT_DIR="/rugpfs/fs0/risc_lab/scratch/jsoroczyns/CAD/CADwalk/CADwalk_E1_GM12878_2x250/results_20250207_0034_nonexpanded/ABC_walk_processing_20250209/20250209_parse2_UU_direct"

# Directory to write outputs:
OUTPUT_DIR="/rugpfs/fs0/risc_lab/scratch/jsoroczyns/CAD/CADwalk/CADwalk_E1_GM12878_2x250/results_20250207_0034_nonexpanded/ABC_walk_processing_20250209/20250211_parse2_UU_direct_ABC_complete_consequent12"
mkdir -p "$OUTPUT_DIR"

# 1) Put your 4 input filenames in an array:
FILES=(
  "medium_TR1_bwamem2_parse2_UU_direct_20250207_0034.pairs.gz"
  "medium_TR2_bwamem2_parse2_UU_direct_20250207_0034.pairs.gz"
  "small_TR1_bwamem2_parse2_UU_direct_20250207_0034.pairs.gz"
  "small_TR2_bwamem2_parse2_UU_direct_20250207_0034.pairs.gz"
)

# 2) Select the file for the current SLURM_ARRAY_TASK_ID:
INPUT_FILE="${FILES[$SLURM_ARRAY_TASK_ID-1]}"
INPUT_PATH="$INPUT_DIR/$INPUT_FILE"

# 3) Construct the output filename:
#    e.g., medium_TR1_bwamem2_parse2_UU_direct_20250207_0034_20250211_ABC_complete.pairs.gz
BASENAME="${INPUT_FILE%.pairs.gz}"
OUTPUT_FILE="${BASENAME}_20250211_ABC_complete.pairs.gz"
OUTPUT_PATH="$OUTPUT_DIR/$OUTPUT_FILE"

echo "Processing file: $INPUT_PATH"
echo "Output file    : $OUTPUT_PATH"

# 4) Stream filtering:
#    - pigz -dc = parallel decompress
#    - gawk = filter logic
#    - pigz -c = parallel compress
pigz -dc "$INPUT_PATH" | \
"$AWK_CMD" '
BEGIN {
    prev = "";
    count = 0;
    invalid = 0;
}

(/^#/) {
    print;
    next;
}

{
    # When readID changes, process the old group
    if ($1 != prev) {
        if (prev != "") {
            if (count == 2 && invalid == 0 && first_index == "1" && second_index == "2") {
                print group1;
                print group2;
            }
        }
        # Start a new group
        prev = $1;
        count = 1;
        invalid = 0;
        group1 = $0;
        first_index = $9;
    } else {
        count++;
        if (count == 2) {
            group2 = $0;
            second_index = $9;
        } else {
            # More than 2 lines => invalid
            invalid = 1;
        }
    }
}

END {
    # Check the last group
    if (prev != "" && count == 2 && invalid == 0 && first_index == "1" && second_index == "2") {
        print group1;
        print group2;
    }
}' | pigz -c > "$OUTPUT_PATH"

echo "Done filtering $INPUT_PATH."
```
