## Supplementary material for "CAD-C: An engineered nuclease enables repair-free *in situ* proximity ligation and nucleosome-resolution chromosome walks in human cells": Custom code: parallel_tsv_flatten_ABC_Bfrag_filter_base_v2_20250214.py

```
#!/usr/bin/env python3

import os
import sys
import csv
import gzip
import glob
import datetime
import argparse
import concurrent.futures

def parse_int(s, default=0):
    try:
        return int(s)
    except ValueError:
        return default

def flatten_file(input_file, output_file, metrics_file):
    """
    Process a single .pairs(.gz) file, flatten ABC, apply filters on B fragment:
      1) strand2( line1 ) != strand1( line2 )
      2) chrom2( line1 ) == chrom1( line2 )
      3) (B_end - B_start) < 500
    Write the passing ABC records to output_tsv.gz and record filter metrics.
    """

    # Decide if we need "rt" for gzip input
    if input_file.endswith(".gz"):
        in_f = gzip.open(input_file, "rt")
    else:
        in_f = open(input_file, "r")

    # Decide if we need "wt" for gzip output
    if output_file.endswith(".gz"):
        out_f = gzip.open(output_file, "wt", newline="")
    else:
        out_f = open(output_file, "w", newline="")

    writer = csv.writer(out_f, delimiter="\t")

    # Header for the flattened output
    header = [
        "readID",
        "A_chrom", "A_start", "A_end", "A_strand",
        "B_chrom", "B_start", "B_end", "B_strand",
        "C_chrom", "C_start", "C_end", "C_strand"
    ]
    writer.writerow(header)

    # Counters
    total_B_fragments = 0
    n_fail_strand = 0
    n_fail_chrom = 0
    n_fail_length = 0

    current_read_id = None
    line1_fields = None
    n_passed = 0  # final number of ABC records written

    for raw_line in in_f:
        line_str = raw_line.strip()
        # Skip headers or empty lines
        if not line_str or line_str.startswith("#"):
            continue

        fields = line_str.split("\t")
        if len(fields) < 24:
            continue  # skip malformed lines

        read_id = fields[0]

        # If we don't have a stored line1 yet => store
        if current_read_id is None:
            current_read_id = read_id
            line1_fields = fields
            continue

        # If we found line2 for the same readID => flatten
        if read_id == current_read_id:
            line2_fields = fields

            # ----- A logic (from line1) -----
            A_chrom = line1_fields[1]
            A_pos   = parse_int(line1_fields[2])
            A_strand1 = line1_fields[5]
            A_span  = parse_int(line1_fields[22])

            if A_strand1 == '-':
                # A is plus
                A_strand = '+'
                A_start  = A_pos - A_span
                A_end    = A_pos
            else:
                # A_strand1 == '+'
                # A is minus
                A_strand = '-'
                A_start  = A_pos
                A_end    = A_pos + A_span

            if A_start > A_end:
                A_start, A_end = A_end, A_start

            # ----- B logic -----
            B_chrom = line1_fields[3]   # line1.chrom2
            B_start = parse_int(line1_fields[4])  # line1.pos2
            B_end   = parse_int(line2_fields[2])  # line2.pos1
            B_strand_parse2 = line1_fields[6]     # line1.strand2
            B_strand = B_strand_parse2

            # reorder start/end
            if B_start > B_end:
                B_start, B_end = B_end, B_start

            # We'll compare for filters:
            total_B_fragments += 1
            fail_strand = (line1_fields[6] == line2_fields[5])  # want them !=
            fail_chrom  = (line1_fields[3] != line2_fields[1])  # want them ==
            b_length    = B_end - B_start
            fail_length = (b_length >= 500)                     # want < 500

            # If any fail => skip
            if fail_strand or fail_chrom or fail_length:
                if fail_strand:
                    n_fail_strand += 1
                if fail_chrom:
                    n_fail_chrom += 1
                if fail_length:
                    n_fail_length += 1
                # skip writing
            else:
                # pass => proceed with C logic
                # ----- C logic (from line2) -----
                C_chrom = line2_fields[3]
                C_pos   = parse_int(line2_fields[4])
                C_strand2 = line2_fields[6]
                C_span = parse_int(line2_fields[23])

                if C_strand2 == '+':
                    C_strand = '+'
                    C_start  = C_pos
                    C_end    = C_pos + C_span
                else:
                    # C_strand2 == '-'
                    C_strand = '-'
                    C_start  = C_pos - C_span
                    C_end    = C_pos

                if C_start > C_end:
                    C_start, C_end = C_end, C_start

                # Write the flattened row
                writer.writerow([
                    current_read_id,
                    A_chrom, A_start, A_end, A_strand,
                    B_chrom, B_start, B_end, B_strand,
                    C_chrom, C_start, C_end, C_strand
                ])
                n_passed += 1

            # Reset for next read
            current_read_id = None
            line1_fields = None

        else:
            # New read => skip old single-liner
            current_read_id = read_id
            line1_fields = fields

    in_f.close()
    out_f.close()

    # Write metrics
    now_str = datetime.datetime.now().strftime('%Y-%m-%d %H:%M:%S')
    with open(metrics_file, "w") as mf:
        mf.write(f"Timestamp: {now_str}\n")
        mf.write(f"Input file: {os.path.abspath(input_file)}\n")
        mf.write(f"Output file: {os.path.abspath(output_file)}\n")
        mf.write(f"Total B fragments encountered: {total_B_fragments}\n")
        mf.write(f"Final ABC records passed: {n_passed}\n")
        if total_B_fragments > 0:
            pass_percent = (n_passed / total_B_fragments) * 100
        else:
            pass_percent = 0.0
        mf.write(f"  => Pass %: {pass_percent:.2f}%\n\n")

        # Separate fails
        mf.write(f"Filter 1 fails (strand same): {n_fail_strand} "
                 f"({(n_fail_strand/total_B_fragments*100) if total_B_fragments else 0:.2f}% of input)\n")
        mf.write(f"Filter 2 fails (chrom mismatch): {n_fail_chrom} "
                 f"({(n_fail_chrom/total_B_fragments*100) if total_B_fragments else 0:.2f}% of input)\n")
        mf.write(f"Filter 3 fails (length >= 500): {n_fail_length} "
                 f"({(n_fail_length/total_B_fragments*100) if total_B_fragments else 0:.2f}% of input)\n")

    print(f"[DONE] {input_file} => {output_file}")
    print(f"   Passed: {n_passed} / {total_B_fragments} B fragments")

##############################################################################
# The rest of the script: parse_args, parallel flatten, etc. remains the same
##############################################################################

def parse_args():
    parser = argparse.ArgumentParser(
        description="Flatten .pairs(.gz) files into ABC records with B-fragment filters, produce TSV + metrics."
    )
    parser.add_argument(
        "--input-dir", required=True,
        help="Directory containing input .pairs(.gz) files."
    )
    parser.add_argument(
        "--pattern", default="*.pairs.gz",
        help="Filename pattern to match in --input-dir (default: '*.pairs.gz')."
    )
    parser.add_argument(
        "--output-dir", required=True,
        help="Directory to write output .tsv(.gz) and metrics files."
    )
    parser.add_argument(
        "--threads", type=int, default=1,
        help="Number of parallel worker processes (default=1)."
    )
    return parser.parse_args()

def make_output_paths(infile, output_dir):
    """
    Given an input file, return the corresponding output file path
    and metrics file path in output_dir.
    """
    base = os.path.basename(infile)
    # Remove .pairs or .pairs.gz, then add .tsv.gz
    if base.endswith(".pairs.gz"):
        stem = base.replace(".pairs.gz", "")
        out_tsv = f"{stem}.flattened.tsv.gz"
    elif base.endswith(".pairs"):
        stem = base.replace(".pairs", "")
        out_tsv = f"{stem}.flattened.tsv"
    else:
        # fallback
        stem, _ = os.path.splitext(base)
        out_tsv = f"{stem}.flattened.tsv"

    out_file = os.path.join(output_dir, out_tsv)
    now_str = datetime.datetime.now().strftime('%Y%m%d_%H%M%S')
    metrics_file = os.path.join(output_dir, f"{stem}_metrics_{now_str}.txt")
    return out_file, metrics_file

def main():
    args = parse_args()
    os.makedirs(args.output_dir, exist_ok=True)
    input_pattern = os.path.join(args.input_dir, args.pattern)
    input_files = sorted(glob.glob(input_pattern))
    if not input_files:
        print(f"No files found matching {input_pattern}", file=sys.stderr)
        sys.exit(1)

    with concurrent.futures.ProcessPoolExecutor(max_workers=args.threads) as executor:
        futures = []
        for infile in input_files:
            out_file, metrics_file = make_output_paths(infile, args.output_dir)
            f = executor.submit(flatten_file, infile, out_file, metrics_file)
            futures.append(f)

        # Wait for all to complete, check for errors
        for f in concurrent.futures.as_completed(futures):
            exc = f.exception()
            if exc:
                print(f"ERROR in a worker: {exc}", file=sys.stderr)

if __name__ == "__main__":
    main()
```
