## Supplementary material for "CAD-C: An engineered nuclease enables repair-free *in situ* proximity ligation and nucleosome-resolution chromosome walks in human cells": Custom code: README_FLAT_UPLOAD_20260511.txt

This folder is a flat upload-oriented derivative of CADC_Code_Handoff_20260511_STAGING.

Only byte-verified staged copies were used as inputs; original Dropbox/source files were not read again or modified.
Python scripts are in py/ and shell scripts are in sh/.

If multiple workbook entries pointed to the same script filename with the same SHA256 hash, the script was copied once and duplicate references are recorded in FLAT_UPLOAD_MANIFEST_20260511.tsv.
If same-named scripts had different SHA256 hashes, the later copy was disambiguated with parent-folder context and an SHA256 prefix so no file was overwritten.
.R/.r and .ipynb entries, URL-only entries, and provenance-only records are listed in the manifest but intentionally not copied into this two-folder flat upload package.
