## Supplementary Figures for "CAD-C: An engineered nuclease enables repair-free *in situ* proximity ligation and nucleosome-resolution chromosome walks in human cells"

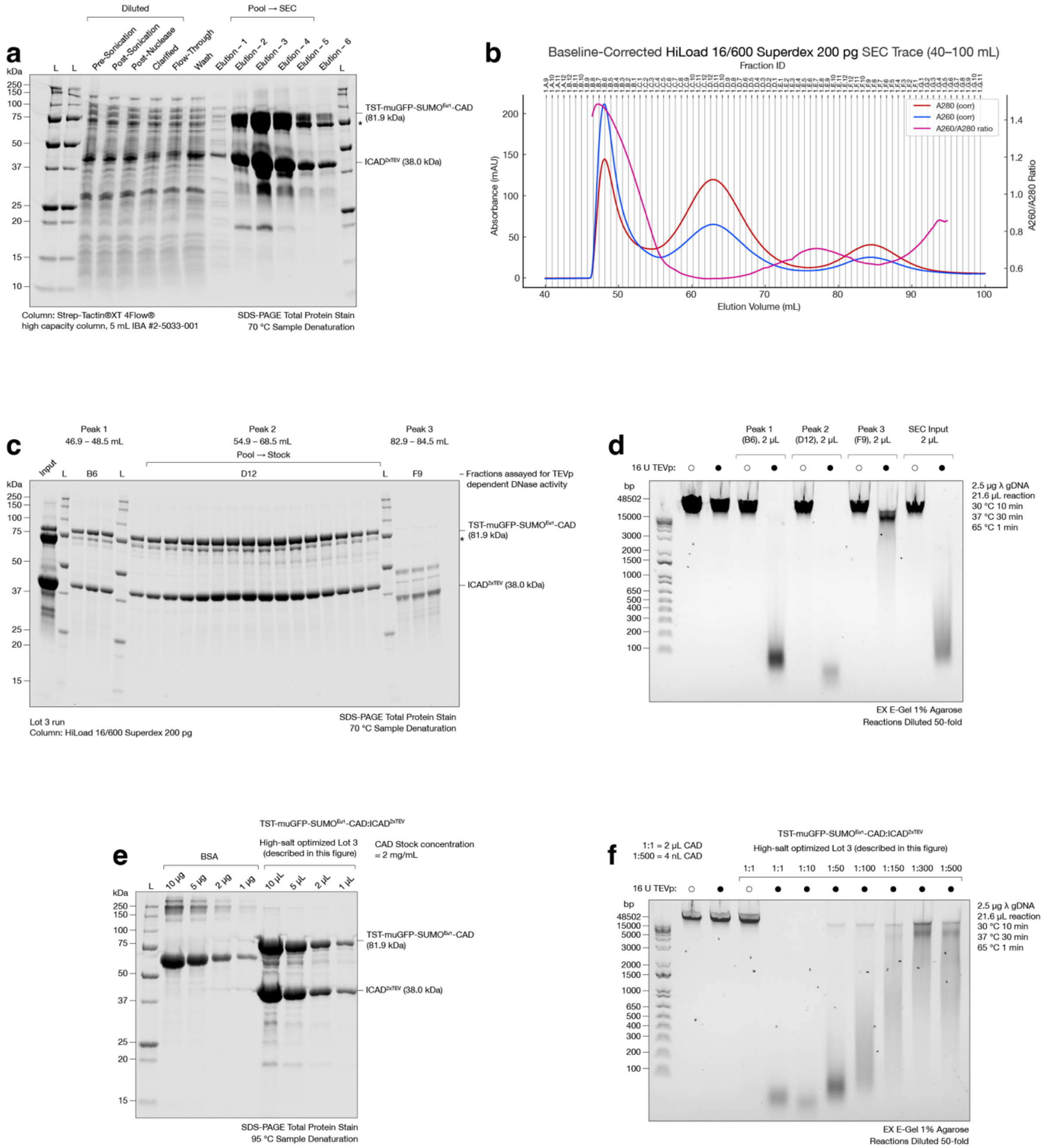


**Supplementary Figure 1. High-salt optimized CAD:ICAD$_{2xTEV}$ stock (Lot 3): purification checks, concentration estimate, and λ-DNA cleavage activity.**

(a) Coomassie-stained SDS–PAGE across the Strep–SEC workflow for Lot 3 (high-salt optimized; described in Methods), purified using high-salt lysis and capture on a 5 mL Strep-TactinXT 4Flow high-capacity column. For routine SDS–PAGE in panels (a,c), samples were denatured at lower temperature (70–75°C, rather than boiling) to minimize aberrant heat induced cleavage; under these conditions the TST-muGFP-$SUMO_{Eu1}$-CAD fusion reproducibly appears as a closely spaced band doublet (upper band plus a faster, anomalously migrating species), which we interpret as incomplete denaturation/altered SDS–PAGE mobility of the highly stable $muGFP_{70}$-containing fusion.

(b) Baseline-corrected size-exclusion chromatography trace (A280, A260, and A260/A280) of 5 mL Strep-column eluate injected onto a HiLoad 16/600 Superdex 200 pg column (see high-salt Lot 3 Methods).

(c) Coomassie-stained SDS–PAGE of size-exclusion chromatography fractions spanning Peak 1 (void), Peak 2, and Peak 3. Peak 2 contains the predominant intact TST-muGFP-$SUMO_{Eu1}$-CAD (81.9 kDa) and $ICAD_{2xTEV}$ (38.0 kDa) species used to prepare the working stock (Lot 3).

(d) TEVp-dependent λ-DNA digestion assay on representative SEC fractions (±TEVp) using a shortened single-tube reaction (2.5 µg λ gDNA in 21.6 µL modified 1×WDB-70 (0.1 mM DTT); 30°C 10 min, 37°C 30 min; 65°C 1 min), followed by 50-fold dilution and analysis on 1% agarose E-Gel.

(e) BSA calibration ("concentration") gel for Lot 3 (BSA 10, 5, 2, 1 µg; Lot 3 loaded at 10, 5, 2, 1 µL), estimating ~2 $mg·mL_{-1}$ total protein (~1 $mg·mL_{-1}$ with respect to tagged CAD). For this gel, samples were denatured at 95°C, which eliminates the faster anomalously migrating band and collapses the TST-muGFP-$SUMO_{Eu1}$-CAD doublet to a single band, consistent with full denaturation of the muGFP-containing fusion.

(f) Serial dilution titration of the Lot 3 Peak 2 stock in the shortened λ-DNA assay, 1×WDB-70 with standard 1 mM DTT (±16 U TEVp; 1:1–1:500; "1:1 = 2 µL stock per reaction"). CAD dilutions were made in 1×CSEC buffer supplemented with 0.1 $mg·ml_{-1}$ BSA, this is done to reduce non-specific adsorption of CAD onto reaction tube walls, i.e. when using reaction vessels other than Eppendorf Protein LoBind tubes. Complete digestion to predominantly <100 bp fragments persists to at least a 1:50 dilution in +TEVp reactions with minimal/undetectable digestion in matched –TEVp controls, corresponding to an estimated activity of ≥62.5 $U·µL^{-1}$ (≥2.5× higher than Lot 1 at ~25 $U·µL^{-1}$, under the standard unit definition).

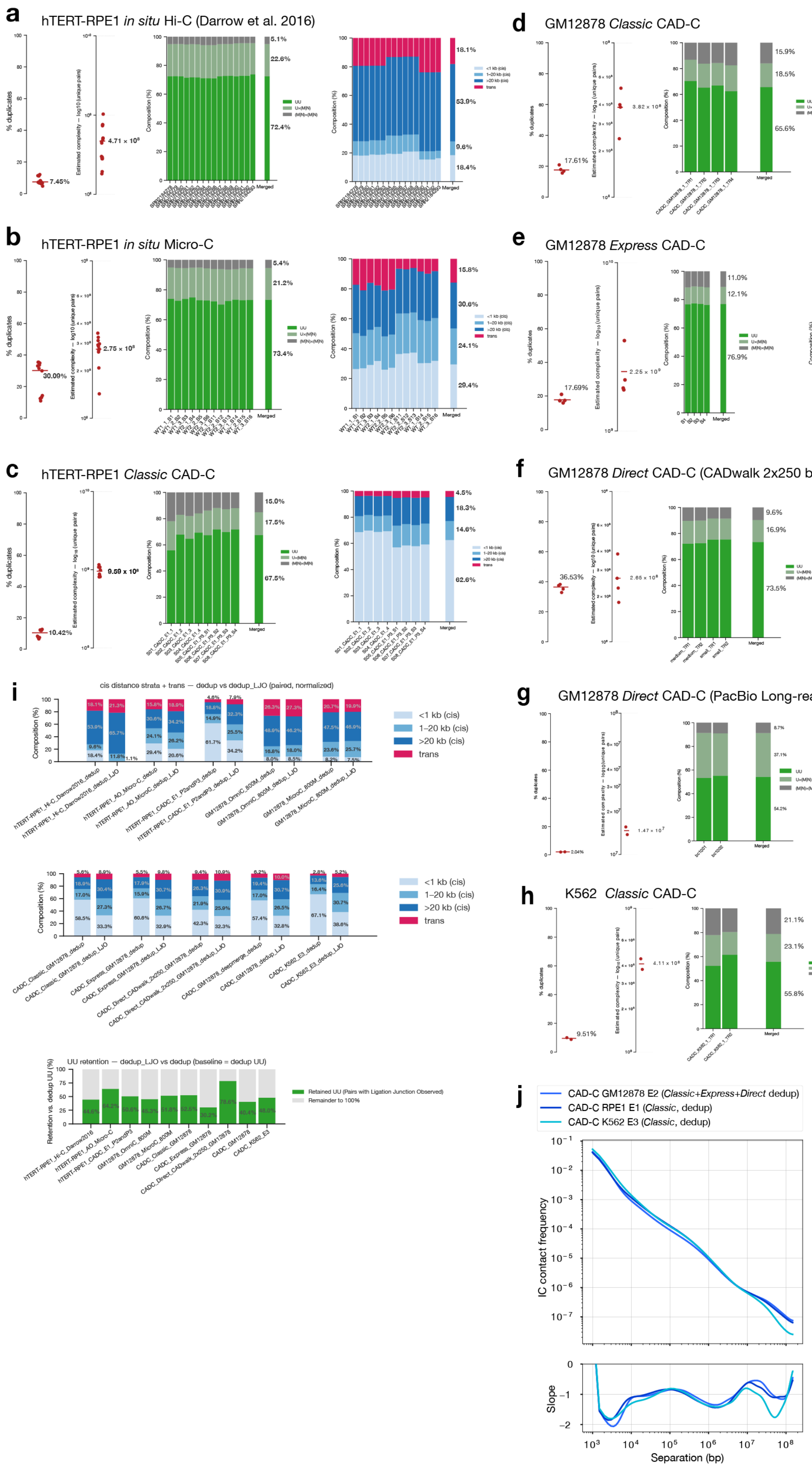
a
hTERT-RPE1 in situ Hi-C (Darrow et al. 2016)
b
hTERT-RPE1 in situ Micro-C
c
hTERT-RPE1 Classic CAD-C
d
GM12878 Classic CAD-C
e
GM12878 Express CAD-C
f
GM12878 Direct CAD-C (CADwalk 2x250 bp)
g
GM12878 Direct CAD-C (PacBio Long-read CADwalk )
h
K562 Classic CAD-C
i
cis distance strata + trans — dedup vs dedup_LJO (paired, normalized)
<1 kb (cis)
1–20 kb (cis)
>20 kb (cis)
trans
UU retention — dedup_LJO vs dedup (baseline = dedup UU)
Retained UU (Pairs with Ligation Junction Observed)
Remainder to 100%
j
CAD-C GM12878 E2 (Classic+Express+Direct dedup)
CAD-C RPE1 E1 (Classic, dedup)
CAD-C K562 E3 (Classic, dedup)
IC contact frequency
Slope
Separation (bp)

**Supplementary Figure 2. Contact pair summary statistics across Hi-C, Micro-C, CAD-C, and CADwalk datasets.**

(a-h) Pairtools summary statistics for each dataset (hTERT–RPE1 in situ Hi-C[51], hTERT–RPE1 in situ Micro-C, hTERT–RPE1 Classic CAD-C; GM12878 Classic CAD-C, Express CAD-C, and Direct CAD-C (CADwalk 2×250 bp); GM12878 Direct CAD-C (PacBio long-read CADwalk); and K562 Classic CAD-C). For each dataset, panels report duplicate rate, estimated library complexity, pair-type composition (collapsed to UU, U×(M/N), and (M/N)×(M/N)), and contact-distance composition (cis <1 kb, cis 1–20 kb, cis >20 kb, and trans).

(i) Cross-dataset comparison of contact-distance composition before and after ligation-junction–observed filtering (dedup vs dedup_LJO), and the corresponding UU retention after LJO filtering.

(j) Intrachromosomal contact-probability decay P(s) and the local slope for representative CAD-C datasets (GM12878, hTERT–RPE1, and K562; deduplicated).

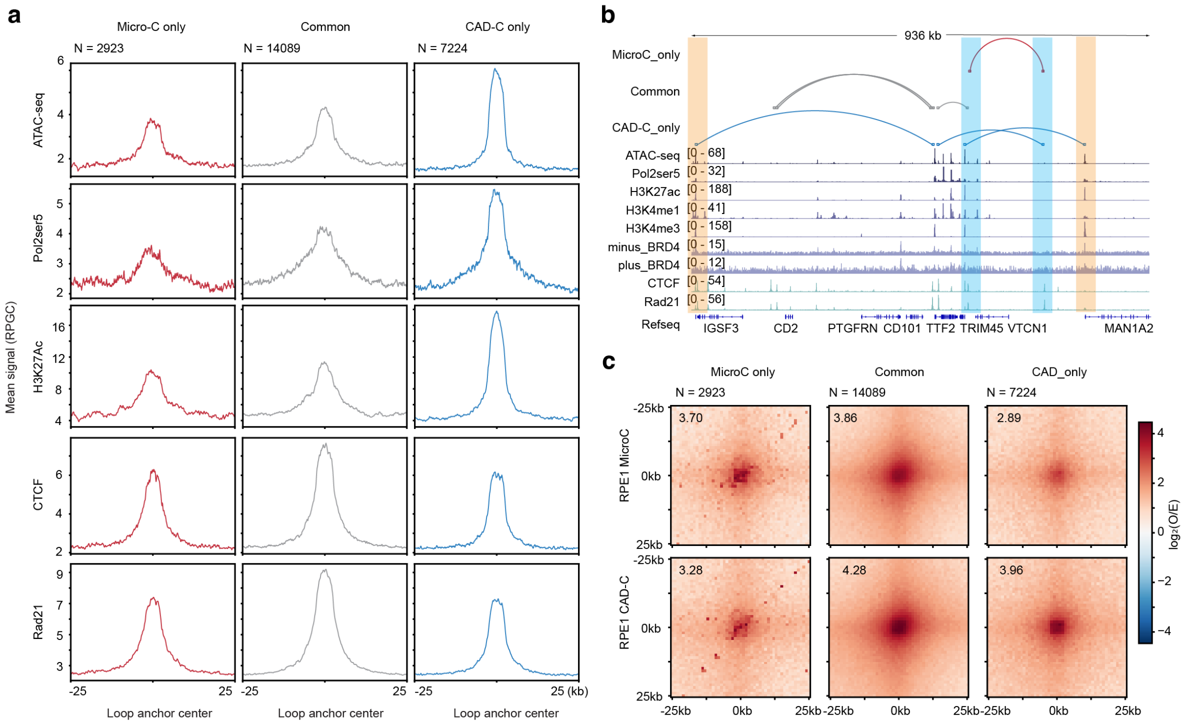


**Supplementary Figure 3. CAD-C preferentially captures the active transcription loops compared to in situ Micro-C in hTERT-RPE1 cells**

(a) Pileup line plots for chromatin accessibility (ATAC-seq), Pol2Ser5, H3K27ac and CTCF, Rad21 at the anchors called at the resolution of 5kb with equal depth of CAD-C and Micro-C data from hTERT-RPE1 cell line.

(b) Genomic tracks across the chr1:116.56Mb-117.50Mb region showing Micro-C and CAD-C loop calls together with ATAC-seq, Pol2Ser5, H3K27ac, H3K4me1, H3K4me3, BRD4, CTCF and Rad21 profiles.

(c) Pileup analysis of the three loop groups in panel a, centered on loops (±25 kb).

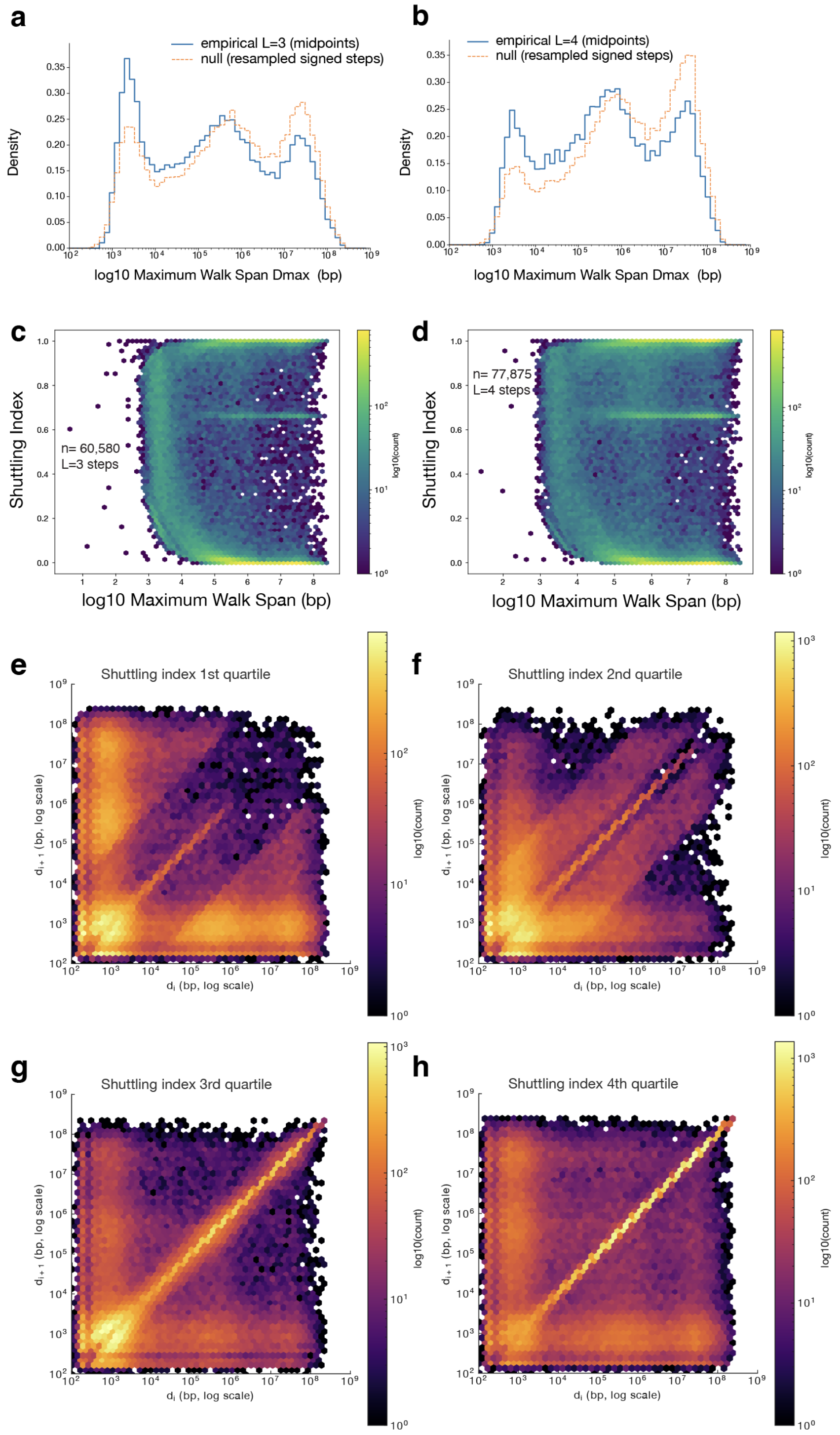


**Supplementary Figure 4. CADwalk span and shuttling behavior across long-read walk lengths.**

(a,b) Distributions of maximum walk span (Dmax) for long-read cis, single-chromosome CADwalks with L = 3 steps (a) or L = 4 steps (b), compared with a resampling null generated by drawing signed step lengths from the empirical distribution within each L class (see Methods).

(c,d) Hexbin density of shuttling index versus $\log_{10}(D_{max})$ for L = 3 (c) and L = 4 (d) CADwalks (color indicates $\log_{10}$ counts).
(e–h) Joint distributions of successive cis step lengths ($d_i$, $d_{i+1}$) pooled across long-read CADwalks and stratified by shuttling index quartile (Q1–Q4), Q1 = lowest shuttling index quartile, Q4 = highest. Axes are in bp on log–log scales; color indicates $\log_{10}$ counts.

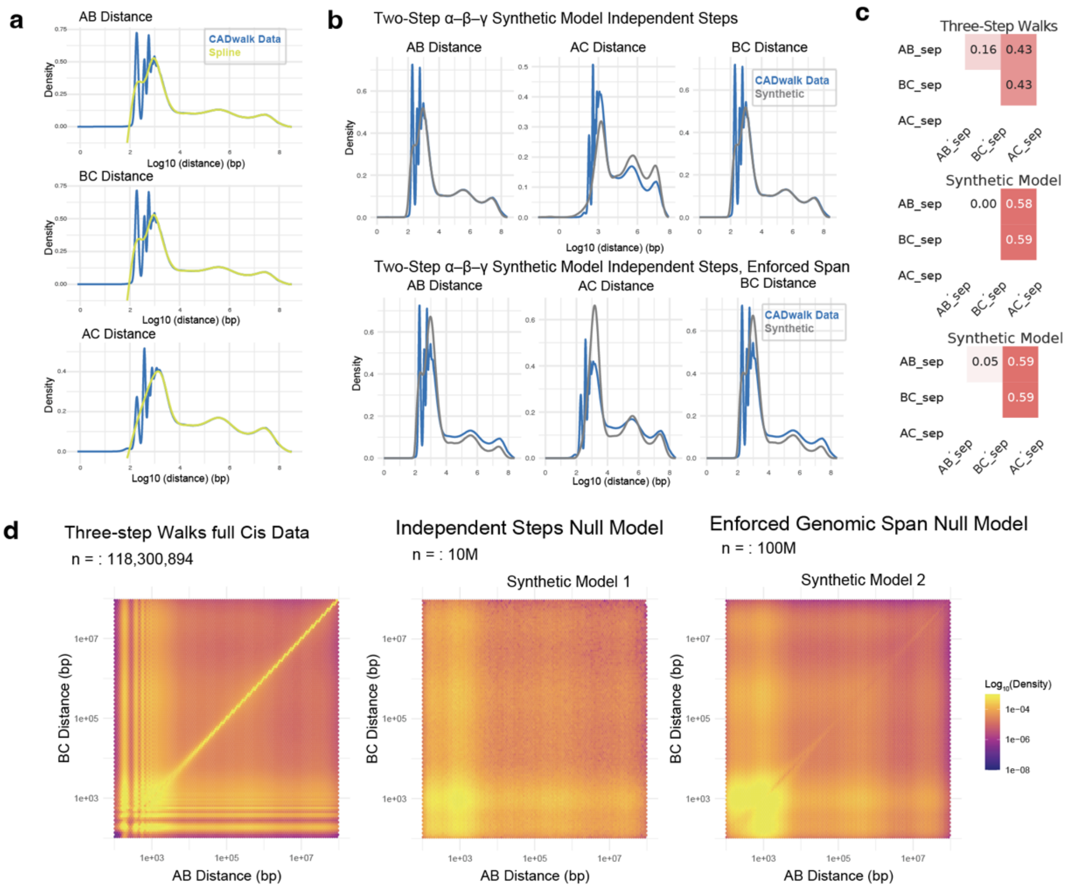


**Supplementary Figure 5. Synthetic random-sampling null model of short-read CADwalks.**
Development of Two-Step α-β-γ Synthetic Null Models 1 + 2 (see Methods)
(a) Density distributions of distances for AB, BC, AC contacts from observed step length distributions (blue) fitted spline (lime). Here and in panels below A,B,C, refer to the α,β,γ fragments in short-read CADwalks.
(b) Density distributions of AB, BC, AC of observed data (blue) and synthetic models (gray). Top: Model 1, independent sampling, Bottom: enforced α-γ distance distributions to match empirical data.
(c) Spearman correlation heatmaps of separation from the density distributions in B.
(d) Joint distribution hexagonal-binned density maps of αβ distance vs βγ distance from cis-chromosomal two-step α-β-γ walks, Model 1, and Model 2. Hexbin intensities are normalized to 1 within each panel. Hexbin fill values are shown with a $\log_{10}$ color transform.
