## Supplementary Methods for "CAD-C: An engineered nuclease enables repair-free *in situ* proximity ligation and nucleosome-resolution chromosome walks in human cells"

**CADwalk Code Documentation for:**

**Short, two-step A-β-γ CADwalk analysis**

To analyze short-range three-fragment chromatin walks, we focused on two consecutive ligation junctions derived from individual proximity-ligation concatemers, corresponding to A–B–C fragment configurations. Reads were restricted to those containing exactly two consecutive junctions, ensuring that the two interaction steps shared a single intermediate fragment. Candidate walks were required to traverse a single digestion fragment at the intermediate position, enforced by genomic contiguity and strand-orientation constraints, thereby isolating bona fide two-step walks rather than independent ligation events. Validated A–B–C walks were reconstructed into three genomic fragments with standardized coordinates and orientations. Records were deduplicated, normalized to a consistent genomic order, and separated into cis and trans configurations based on chromosomal identity. Cis walks were further stratified by the relative genomic ordering of the three fragments (ABC, ACB, or BAC), enabling positional analysis of multi-fragment contact structures.

Fragments from ABC walks were annotated by intersecting fragment start positions with genomic state annotations, including chromatin compartments and ChromHMM states. These annotations were used to stratify subsequent analyses by chromatin context. Fragment length distributions were computed separately for A, B, and C fragments within each genomic state, providing state-specific fragment size profiles. To characterize the spatial organization of two-step walks, we analyzed the distances between fragments and local contact patterns. Separation histograms were truncated to short genomic ranges to emphasize local contact structure and were visualized as state-stratified density profiles. For cis walks, fragment midpoint–midpoint contacts were aggregated within fixed windows centered on genomic features of interest, yielding local contact matrices for direct (A–B, B–C) and indirect (A–C) interactions. Strand-specific profiles were combined by symmetry to generate strand-agnostic representations.

We further examined motif-centered organization by aligning ABC walks to fixed windows around CTCF motifs. Fragment coverage, midpoint distributions, and fragment length–distance relationships were quantified relative to motif centers, producing aggregated coverage profiles and V-plots. All analyses pooled fragments across walk positions unless otherwise specified and were performed in a fully reproducible manner.

**Directionality scores, B-Enrichment Index (BEI) computation, detrending, and genome-wide track export**

**Input and orientation.**

We started from a tab-delimited CAD-walk file (.tsv/.tsv.gz) that includes a header line and may contain leading comment lines beginning with # (ignored by the parser). The BEI generator (CADwalk_calculate_BEI_v2_20250925.py) reads exactly the named columns A_chrom, A_mid, B_mid, and C_mid. Walks are assumed to be pre-oriented such that A lies upstream of C (as documented inside the script). No B_chrom/C_chrom columns are consulted; all counting is keyed by A_chrom.

**Binning and streaming.**

Next, Genomic space is discretized into fixed-width bins, and counts are accumulated per chromosome in three int64 arrays ($n_A$, $n_B$, $n_C$). For each read row, the midpoint in base pairs is converted to a bin index by integer division, bin = mid // bin_size, and incremented. Files are processed in streaming chunks to cap memory. Parameters actually used: BEI was run genome-wide (no --region, no --chrom filters) with 1 kb bins (--bin_size 1000) and the default streaming size of 2,000,000 rows per chunk (--chunk_rows 2000000).

**Compositional normalization and BEI.**

For each non-empty bin (where $n_{\text{tot}} = n_A + n_B + n_C > 0$), compositional proportions are computed with Laplace smoothing:

$$r_X = \frac{n_X + 0.5}{n_\text{tot} + 1},\ \ X \in \{A,B,C\},\qquad r_C = 1 - r_A - r_B.$$

The B-Enrichment Index (BEI) is then

$$\text{BEI} = 2,r_B - 1.$$

High BEI indicates interior (B) enrichment; low or negative BEI indicates anchor (A+C) dominance. (BEI is the sign-reversed legacy ABI.)

**Outputs from the BEI generator.**

The script writes a timestamped subfolder containing:

• bei_bins_<ts>.tsv.gz with columns Chrom, Start, End, r_A, r_B, r_C, BEI, n_tot (only bins with $n_\text{tot}>0$);

• raw_counts_<ts>.tsv.gz with Chrom, Start, End, n_A, n_B, n_C;

• metadata_<ts>.json with parameters and script checksum; a run log is also produced.

(Wrapper: submit_generate_BEI_bins_v2_20250925.sh invoked the generator with --bin_size 1000 and no region/chrom filters.)

**Detrending and robust Z-scoring of ratios.**

For downstream analysis, calculate_detrended_cadwalk_ratio_metrics_v4_20250925.py ingests the BEI table, filters to chromosomes whose names start with chr, and (optionally) applies a coverage threshold before any modeling. It computes raw log ratios

$$\log_2(A/C) = \log_2(r_A/r_C),\quad \log_2(A/B) = \log_2(r_A/r_B),$$

fits a per-chromosome linear trend (degree-1 polynomial in centered genomic position) for each ratio, and stores residuals as detrended values. Finally, it converts per-chromosome residuals to robust Z-scores using MAD×1.4826; Z_ACd and Z_ABd are derived from the detrended ratios, and Z_BEI is the robust Z-score of the raw BEI (no detrending). Parameters actually used: --min_cov 50 (bins with $n_\text{tot} < 50$ were excluded), invoked via the detrend/Z-score wrapper init_detrended_v4_20250925_1.sh. Outputs: <prefix>_with_detrend.tsv.gz plus <prefix>_detrend_coeffs.json (per-chromosome slopes/intercepts).

**Genome-wide BigWig export.**

BEI_to_bigwig_v5_20250925.py converts the BEI table into three 1 kb BigWig tracks:

• Z_ACd.bw — per-chromosome robust-Z of detrended $\log_2(A/C)$,

• Z_ABd.bw — per-chromosome robust-Z of detrended $\log_2(A/B)$,

• Z_BEI.bw — per-chromosome robust-Z of BEI (no detrend).

The script filters to chromosomes starting with chr, optionally excludes *random, chrUn, etc. (via*

*--exclude_random; not used here), detects the bin size from the table (End−Start or modal ΔStart), recomputes \log_2(A/C) and \log_2(A/B), applies the same per-chromosome linear detrend and MAD-based robust Z, and writes tracks using chromosome sizes from an input FASTA index (.fai). Parameters actually used: --min_cov 50 (only bins with n*\text{tot} \ge 50 exported) and no --exclude_random; FASTA index supplied via --fai. This run was driven by submit_genomewide_BEI_to_bigwig_v5_20250925.sh, which produced *_Z_ACd.bw, *_Z_ABd.bw, and *_Z_BEI.bw in a timestamped output directory.

**Track resolution note (1 kb binned vs 1 bp per-site tracks).**
The BEI-derived BigWig tracks described above (Z_ACd.bw, Z_ABd.bw, Z_BEI.bw) are exported at the BEI bin size (here 1 kb bins). In contrast, per-site walk metrics (e.g., WPI and AWB) were exported as sparse, single-base BigWigs with 1-bp intervals (“bin1” tracks). All meta-coverage pileups of WPI and AWB reported in this study used these 1-bp-resolution BigWigs (not 1 kb-binned summaries); accordingly, because Z_WPI.bw is stored as sparse 1-bp intervals [Pos, Pos+1), we used this single-base (“bin1”) Z_WPI track directly asi input for strand-aware pileups over CTCF motif instances and RAMPAGE TSSs.

**Implementation notes tied to the code.**
• Counting for A, B, and C uses A_chrom as the key; the code does not inspect B_chrom/C_chrom and therefore bins B and C midpoints under A_chrom (consistent with cis-only use cases).
• Only bins with n_\text{tot}>0 appear in the BEI table.
• Robust Z is computed per chromosome as (x - \text{median}) / \text{MAD}, with MAD scaled by 1.4826; if MAD is zero, results are set to NaN by the helper.
• BigWig export inserts a placeholder (length-1 zero entry) for chromosomes lacking valid rows to keep headers consistent.

**Walk Polarity Index (WPI) computation and genome-wide export**

**Input, orientation, and streaming.**
We quantified the polarity and strength of non-linear returning walks in three-fragment CAD data with WPI_compute_and_bigwig_v1_20250926.py. The program ingests a tab-delimited walks file (.tsv/.tsv.gz; leading “#” comments ignored) containing the named columns A_chrom, A_mid, B_mid, C_mid; walks are assumed to be pre-oriented such that A lies upstream of C. Processing is streamed in large chunks (we used 8,000,000 rows per chunk) to bound memory.

**Per-walk WPI definition.**
For each walk, absolute genomic distances are computed as AB=|B{-}A|, BC=|C{-}B| and AC=|C{-}A|. A scale-free fold fraction is defined as F = 1 - AC/(AB{+}BC), clamped to [0,1] and set to NaN when AB{+}BC=0; F=0 for linear A–B–C ordering and increases toward 1 as the walk approaches a non-linear returning walk (i.e., B lies outside the A–C interval and the two legs are of similar length). Polarity is the sign of the displacement of B from the A–C midpoint, s=\mathrm{sign}!\big((A{+}C)/2 - B\big) (+ when B is upstream of the midpoint, − when downstream). The per-walk Walk Polarity Index is \mathrm{WPI}=s\cdot F \in [-1,1].

**Per-site aggregation, filtering, and robust Z-scoring.**

Each walk's WPI is deposited at the B fragment midpoint coordinate (site defined as (Chrom=A_chrom, Pos=B_mid)). Within each chunk we collapse identical sites to partial sums \sum \mathrm{WPI} and counts n, write gzipped partial tables, and after streaming completes we merge all partials and re-aggregate across chunks to obtain per-site WPI_mean = ( \sum \mathrm{WPI} )/n and coverage n. Sites are restricted to chromosomes whose names start with "chr"; optional exclusion of *random/chrUn/KI/GL/EBV scaffolds is available but was not used here. A coverage filter is then applied; for the analyses reported here we set --min_cov 1, retaining all observed sites. To standardize across chromosomes without imposing a trend model, we compute a per-chromosome robust Z-score of the site means, Z*{\mathrm{WPI}} = \big(\mathrm{WPI_mean} - \mathrm{median}\big)/\mathrm{MAD}, where MAD is the median absolute deviation scaled by 1.4826; when MAD is zero, Z values are set to NaN.

**Outputs.**

Outputs are written to a timestamped subdirectory and include: (i) a per-site table wpi_sites_.tsv.gz with columns Chrom, Pos, WPI_mean, n, Z_WPI; (ii) a sparse single-base BigWig Z_WPI.bw with 1-bp intervals [,\mathrm{Pos}, \mathrm{Pos}{+}1) carrying per-chromosome robust-Z values (no detrending); (iii) QC summaries (histograms of WPI_mean and Z_WPI and a text report of quantiles and counts); and (iv) a metadata JSON that records parameters, formulas, the input path, and the script checksum. By construction, WPI captures both magnitude (fold fraction) and direction (upstream vs downstream of the A–C midpoint) of non-linear returning walks, and—because values are placed at B midpoints without binning—produces a genome-wide, per-site polarity track that is unique to three-fragment CAD-walks.

**BigWig pileups over genomic annotations (RAMPAGE TSSs and CTCF motifs)**

We computed strand-aware meta-coverage ("pileup") profiles of single-base BigWig tracks over genomic annotation intervals using smart_bigwig_over_bed_v10p6_20250927.py (SCRIPT_VERSION=v10p6_2025_09_26). Analyses used the "pileup" subcommand with BED inputs and midpoint mode to produce per-interval matrices, average profiles, and PDF metaplots with a timestamped output directory and a run metadata record.

**BED parsing, midpoint windows, and single-base extraction.**

BED files are read with pybedtools. Strand is inferred by auto-detecting a column containing only {+, −, .} in the first 50 non-comment lines, requiring at least one observed +/−. In midpoint mode (--mode midpoint), each feature is summarized by extracting a fixed-width window centered on the interval midpoint mid = floor((start+end)/2). For a half-window W (bp; --window), the extracted window is [mid−W, mid+W) and has length 2W. Signal is extracted at single-base spacing with pyBigWig.values (default --read-mode values). Windows that extend beyond chromosome boundaries are truncated and zero-padded to preserve constant length (2W positions). For minus-strand intervals, the extracted per-feature vector is reversed (x-axis flip) so that all profiles are aligned to a plus-strand frame.

**Zero-centered tracks and strand-dependent sign handling.**

The pileup engine supports auto detection of zero-centered signed tracks (--zero-centered-detect auto) and manual overrides (--force-zero-centered). In addition to the standard x-axis flip for minus-strand features, v10p6 supports optional sign inversion: --flip-sign multiplies a selected BigWig by −1 for all features, and --flip-sign-if-minus multiplies by −1 only for minus-strand features. Strand-dependent sign inversion was applied for directional, zero-centered Z-score tracks where the sign encodes directionality (e.g., Z_WPI), but was not applied to non-directional magnitude-only tracks (e.g., zAWB).

**Outputs.**
For each BED×BigWig run, outputs are written to a timestamped subdirectory and include: (i) coverage_matrix_*.tsv.gz (rows=features; columns=bp positions relative to the midpoint, from −W to W−1); (ii) coverage_average_profile_*.tsv (raw mean profile; no smoothing); (iii) metaplot_*.pdf (mean profile plot; Savitzky–Golay smoothing[75] is applied at render time with default settings unless disabled); and (iv) run_metadata_<ts>.txt recording the script version, the full command-line arguments, the effective configuration, and software/package versions.

**Parameters actually used (CTCF motif pileup of Z_WPI).**
Z_WPI.bw (single-base, zero-centered robust Z) was piled up over GM12878_top_5_percent_CTCF_motifs.bed using midpoint mode with W=2500 bp (±2.5 kb), auto zero-centered detection plus an explicit override, and strand-dependent sign inversion for minus-strand motifs:

```
python smart_bigwig_over_bed_v10p6_20250927.py pileup \
  -b GM12878_top_5_percent_CTCF_motifs.bed \
  -w Z_WPI.bw \
  -o <OUTDIR> \
  -p 8 \
  --mode midpoint \
  --window 2500 \
  --zero-centered-detect auto \
  --bw-labels ZSCORE \
  --force-zero-centered ZSCORE \
  --flip-sign-if-minus ZSCORE \
  --plot-format pdf \
  --fonttype 42 \
  --font-size 11 --title-size 13 --label-size 12 --tick-size 10 --legend-size 10
```

**Parameters used for strand-aware RAMPAGE TSS pileups (Z_WPI).**
The same v10p6 pileup framework was used to pile up Z_WPI over all RAMPAGE TSSs using the SBATCH wrapper v10p6_ZWPIbw_over_RAMPAGEall_2500_flipZscore_if_TSSstrandnegative_20250927.sh (±2.5 kb) with strand-dependent sign inversion for minus-strand TSSs.

**Parameters used for zAWB pileups.**

For “bin1” zAWB pileups over CTCF motifs, the same midpoint pileup framework was used, but strand-dependent sign inversion was not applied (i.e., minus-strand features were oriented by x-axis flipping only, without multiplying the signal by −1).

**α-β-γ distance calculations**

Genomic distance features were computed from a gzipped TSV file using a custom R script. The script reads the input TSV into R and, for each three-fragment walk (A, B, C), calculates pairwise cis separations as absolute differences between fragment start coordinates A_start, B_start, C_start: AB_distance = abs(B_start-A_start), BC_distance = abs(B_start-A_start), and AC_distance = abs (C_start-A_start). The resulting distance columns (AB_distance, BC_distance, AC_distance) were appended to the input table for downstream analyses.

**Compartment Annotation**

Genomic distance tables were read from a gzipped TSV file using data.table::fread in R. Each fragment in the three-fragment walk (A, B, C) was assigned an A/B compartment label based on its intersection annotation: entries matching compartment bed files (Methods- compartment bed file generation). Fragments that did not intersect with A or B compartment were treated as missing/invalid. For each walk, a compartment “walk type” label was constructed as A_comp → B_comp → C_comp. For walk-type–stratified analyses, records were retained only if AB_distance, BC_distance, and AC_distance were present and the walk-type label contained no missing compartment calls (i.e., no “NA”).

**Step 1 vs Step 2 Density Hexbin**

We plotted cis-chromosomal α–β–γ walk step-lengths AB_distance (step 1; x-axis) versus BC_distance (step 2; y-axis) using ggplot2::geom_hex(bins = 100). Distances are shown in base pairs (bp). For walk-type–stratified plots, rows were filtered to retain complete cases (non-missing AB_distance, BC_distance, and AC_distance) and to exclude walks with undefined compartment labels (walk_type containing “NA”), where walk_type was defined from A/B compartment assignments of fragments A, B, and C. To make hexbin intensities comparable across panels, each walk contributed equal weight w = 1/N so that the total weight sums to 1 within each plotted panel (all-walk plot: N = nrow(data); per-walk-type plots: N = nrow(walk_data)). Both axes were displayed on log10 scales (scale_x_log10 and scale_y_log10). Hexbin fill used scale_fill_viridis_c(option = “plasma”, trans = “log10”), applying a log10 color transform to the normalized per-bin weights.

**ChromHMM Annotation and Per-Fragment Enrichment**

Each fragment in the cis-chromosomal two-start α-β-γ data was annotated by GM12878 imputed ChromHMM state. See Methods section “Fragment-lengths, midpoints, and pairwise separations appended to ChromHMM-annotated ABC walks.” Diagonal walks are defined by similarity in magnitude ($|\log10(AB) - \log10(BC)| < 0.2$), and all remaining walks were classified as other. For each position (α, β, or γ), fragments were counted by ChromHMM state and converted to percent frequencies. The full cis-chromosomal ChromHMM walk set was used as a baseline reference to calculate per state, per position log2 enrichment of diagonal and other walks. Diagonal log2 enrichment (over full data) was divided by “other” log2 enrichment (over full data) to visualize differences in per-position ChromHMM state.

**Spearman Correlation of Stepwise Walks**

Genomic separations AB_distance and BC_distance on the same walk were tested for correlation by walk classes defined by genomic geometry in R. Walks with missing AB_distance or BC_distance were removed. Diagonal walks are defined by similarity in magnitude ($|\log10(AB) - \log10(BC)| < 0.2$). Off-diagonal walks were classified either as linear if fragment start coordinates were monotonic along the chromosome ($\alpha\_start < \beta\_start < \gamma\_start$), otherwise they were defined as nonlinear. This analysis thus masks the diagonal from off-diagonal walks. Spearman correlation was calculated using stats::cor() within 100 quantile bins of AB_distance (x-axis; 1–100) separately for each walk class using cut() with breakpoints from quantile(AB_distance, probs = seq(0,1,0.01)).For each bin, the number of walks and the corresponding AB distance range (min–max within bin) was recorded.

**Two-Step α-β-γ Synthetic Null Model - Model 1- Independent Assortment**

**See Supplementary Figure 14**

We generated a synthetic null model of three-fragment chromosome walks by fitting a spline to the absolute genomic distance in bp between Alpha and Beta (αβ distance), and to absolute genomic distance in bp between Beta and Gamma (βγ distance) (Plot A). From those splines, we used inverse transform sampling to pick synthetic αβ and βγ distance, which yield an implied αγ distance. This model assumes independence between αβ step and the βγ step. The model was generated using custom R code by (1) building spline distributions for the absolute genomic distance (step length) >100 base pairs between AB **(**αβ) and BC **(**βγ), splines are sampled on a regular grid of 500 log-distance values, normalized to the area under the curve, and cumulatively integrated, thus creating a normalized probability distribution function (PDF), and cumulative distribution function (CDF). (2) computing a kernel density estimate on log10-transformed distances, (3) fitting a cubic regression spline (GAM) with 20 knots using mgcv::gam. To generate synthetic data, it (4) constructs an anchor pool from all observed fragments annotated to A or B compartment in Two-Step α-β-γ data, (5) randomly selects an “A” fragment anchor, (6) samples AB distance using inverse CDF sampling and placing fragment B length from the anchor, (7) samples BC distance using inverse CDF sampling and places fragment C from B, (8) computes geometry-based AC distance (|C-A|), (9) confirms that AC distance falls within chromosome length, then accepting the row (a walk). Distance values are exponentiated from log space back to linear space to get the distance in base pairs, dist_bp. The model iterates until the synthetic data reaches 100M rows (walks). Chromosome length is hard-coded from hg38.

**Two-Step α-β-γ Synthetic Null Model - Model 2- Enforced αγ distance (displacement)**

**See Supplementary Figure 14**

This model is built in custom R code, as above, but with the following modifications: the model builds spline distributions for all distances in a two-step walk; AB, BC, and AC. It generates synthetic data as described above but adds an additional step (10), checking that the synthetic AC distance falls within the real data’s AC probability distribution.

### LONG-READ CADwalk ANALYSES

#### CADwalk UU-run packing and canonical fragment definition

(cadwalk_pack_v9_20251020.py)

The script cadwalk_pack_v9_20251020.py parsed one or more pairtools parse2 .pairs.gz files with a pinned schema, segmented each read into maximal uninterrupted UU runs, and wrote a self-contained CADpack v9 NPZ containing run-, step-, and fragment-level tensors for downstream analyses. The parser first read the #columns header, required the presence of readID, genomic coordinates, pair types, and walk_pair_index, and then streamedthefilebyread,enforcingmonotonic(readID, walk_pair_index)orderunless -no-verify-order or -no-assume-sorted were specified. Uninterrupted runs were defined as consecutive rows from the same read with pair_type=="UU" and strictly consecutive numeric walk_pair_index; non-numeric indices or other pair types terminated a run. For each run, the code applied a single “M-terminal midpoint” orientation model, computing midpoints from pos5/pos3 on the first side 1 and last side 2 steps, reversing the step order only when both midpoints existed on the same chromosome and the terminal midpoint on the right lay upstream of the left. It then emitted step-level cis/trans flags, signed and absolute junction ($dJ$) and midpoint ($dM$) separations, and step-level reference spans; run-level tortuosity, alternation rates, longitudinal spans $D$max,$J$ and $D$max,$M$, and cis/trans counts; and a fragment chain of length $L$ + 1 per run with canonical reference intervals and length provenance (terminal fragments from pos5/pos3 or alignment spans; interior fragments from junction endpoints). All arrays were written to a compressed NPZ together with a compact JSON metadata record, and, if requested with -parquet, mirrored into run, step, and fragment Parquet tables for downstream use.

#### Streaming PacBio CADwalk analytics and merged summaries

The script streaming_PBCADwalk_analyses_v7_20251017.py performed read-only, streaming analytics on pairtools parse2 PacBio CADwalk .pairs.gz files and produced re- producible per–technical replicate (TR) and merged outputs. Using a pinned column map, the TRProcessor class read each file linearly, grouped rows by readID, and segmented contigu- ous UU runs (consecutive walk_pair_index values) without modifying the underlying data. For each run, it reconstructed a fragment chain from pos5/pos3-derived genomic midpoints, computed per-run cis/trans step counts, longest cis-block length, a block-wise tortuosity $B$run (us- ing signed step directions), maximum cis span $D$max per chromosome, alternation rates, and an on-read query-overlap fraction and overcoverage index from the union of U-intervals defined by dist_to_5 and algn_read_span. The script reservoir-sampled non-adjacent cis separations using a fixed-size Algorithm R reservoir per TR, while retaining all adjacent cis separations, and accumulated fragment-length histograms and ECDFs from deduplicated on-read U fragments, stratified by longest cis-block length. It wrote per-run metrics to CSV and NumPy arrays, together with TRlevel PDFs (tortuosity–$D$max hexbins, fragment-length histograms and ECDFs, cis separa- tion ECDFs and small-scale histograms, and stacked UU-run composition by read-length decile) using Matplotlib with Type 42 fonts. A Merger class pooled arrays and metadata across TRs, recomputed merged UU-run composition and cis-separation ECDFs, and constructed a 10 × 10 grid of $B$run and $D$max deciles with Mann–Whitney and Cliff's $\delta$ statistics and BH–FDR-adjusted $q$-values written to CSV for downstream interpretation.

**Computation of trans-step rate and chromosome novelty versus CADwalk length**

Trans-step analyses were performed using the Python script v9cadpack_readDrop_refOverlap_transrate_novelty_plots_exact.py (executed as python3 v9cadpack_readDrop_refOverlap_transrate_novelty_plots_exact.py cadpack_bc1001_bc1002_v9_14422039.npz --out_prefix v9cadpack_readDrop_refOverlap --make_png) under Python 3.11.2 with NumPy 1.24.0, pandas 2.2.3 and Matplotlib 3.7.5. The v9 cadpack NPZ was loaded with NumPy, and stringent read-level filtering was applied using reference coordinates only: within each run, fragment reference intervals (frag_ref_chr_id, frag_ref_lo, frag_ref_hi) were sorted by (run, chromosome, start) and runs were flagged if any interval started before the running maximum end coordinate on the same chromosome (positive-length overlap); reads were then flagged as bad if any of their runs were flagged, and all runs from bad reads were removed. For each retained run of length L (run_L), the number of trans steps was taken from run_n_trans_steps and aggregated by L to compute the per-step trans rate as total trans steps divided by total steps. Novelty among trans steps was computed per run as (number of unique chromosomes visited − 1), where unique chromosomes were collected from the initial chrom1_id and all chrom2_id step endpoints, and was aggregated by L and normalized by total trans steps. Matplotlib was configured to embed TrueType fonts (pdf.fonttype=42) and was used to write a dual-axis line plot of trans rate and novelty versus L and a log10-scaled walk-length barchart; per-L summaries (CSV) and filter masks (NPY) were saved alongside the figures.

**Run-length stratified trans-jump and unique-chromosome counts**

The script v9cadpack_trans_and_uniqchr_by_runLen_palette_v3_20251021.py read a CADpack v9 NPZ and, for each uninterrupted UU run with consistent length run_n_steps == run_L, tallied the distribution of trans-chromosomal jumps and distinct chromosomes explored as a function of run length $L$. For every run, it sliced the corresponding step rows using run_first_step_row and run_n_steps, counted trans steps as those with chrom1_id != chrom2_id, and counted unique chromosomes as the number of distinct non-negative chromosome IDs encountered across all step endpoints. These per-run values were aggregated into two length-stratified count tables countstrans($L$,$T$) and countsuniq($L$,$U$), where $T$ is the number of trans steps and $U$ the number of unique chromosomes, with maximum $T$ and $U$ inferred from the data. The script then generated stacked-bar PDFs for both metrics on a linear count scale and a semilogarithmic log10(Count) scale, restricting the x-axis to $1 \le L \le 13$ and colouring stack components by $T$ or $U$ using a user-selectable palette. Insets in each figure plotted per-$L$ columns normalized to unit height to visualize the fractional composition of runs of a given length. Optionally, when -csv-prefix was supplied, the script wrote companion CSV files listing ($L$,$T$,count) and ($L$,$U$,count) triplets for re-plotting or further statistical analysis.

**Step-distance hexbins stratified by run length and Brun (Shuttling index)**

(Lx_hexbin_robustZ_and_counts_v3.py)

Note: “Brun” is referred to as “shuttling index” in the text

The script Lx_hexbin_robustZ_and_counts_v3.py generated log-space hexbin maps of cis step separations coloured by robust-z–scored run-level tortuosity for CADpack NPZ files. It first loaded step- and run-level arrays, including chrom1_id, chrom2_id, pos1, pos2, run_first_step_row, run_n_steps, and a run-level run_Brun_* field (prefer- ring run_Brun_M when present), and identified eligible runs whose step count lay between -runlen-min and -runlen-max and for which all steps were cis and confined to a single chromosome. For each such run of length $L$, it computed per-step cis distances |pos2 − pos1| in J-mode and, for every ordered pair of steps (0, $k$) with 1 ≤ $k$ < $L$, collected the base-10 logarithms of their distances as (log10 $d$0, log10 $dk$ ) together with the run’s Brun value. Across all runs and step pairs, the script transformed Brun into a robust z-score using a median and MAD-based scaling, then used Matplotlib hexbins on the fixed log–log extent 102–109 bp (with user-selectable grid size) to compute per-hex medians and means of robust-$z$(Brun), using a global symmetric colour scale (TwoSlopeNorm) centred at zero. It generated separate PDFs for median and mean robust-$z$ hexbins (RdBu_r palette) and for log10(count) hexbins (Inferno, LogNorm) for each ($L$, $k$) pair, enforcing square aspect ratios, power-of-ten tick labels, rasterised hex collections, and Type 42 fonts. A JSON sidecar recorded the NPZ path, run-length range, grid size, Brun key, global colour scale, and list of output files.

**Empirical versus resampled Dmax distributions by run length**

(Dmax_empirical_vs_signedstep_resampling_null_Lstratified_v1_20251021.py)

The Python script tested empirical Dmax distributions for CADpack v9 runs against a signed-step resampling null within fixed run-length strata. Given a v9 NPZ and a target run length $L$, it auto-detected fragment-level keys (canonical chromosome, start, end, run-wise fragment offsets and counts) and, whenrequestedvia-geometry junctions,junction-levelkeysforstepanchors;fieldnames couldalsobeoverriddenexplicitly.Foreachrunwithrun_n_frags == L+1,itrequiredthat all fragments be cis on a single chromosome, that canonical fragment lengths fall within per-stratum quantiles [ql, qh] (-frag-qlo, -frag-qhi), and that no fragment intervals overlapped on the reference; runs failing any criterion were excluded. For geometry mids, it computed midpoints from canonical fragment bounds, derived signed inter-fragment steps, and recorded the empirical Dmax as the range of the path; for geometry junctions, it performed the analogous calculation on junction positions. Across all kept runs, it pooled signed steps within the $L$ stratum and generated -null-reps synthetic paths by sampling steps with replacement and recomputing path ranges, thereby obtaining null Dmax samples of matching step count. The script wrote empirical and null Dmax values to separate CSV files, summarised counts, quantiles, entropy and Kolmogorov–Smirnov distances in a JSON file, and produced log10-scale overlaid histograms. The Slurm wrapper submit_Dmax_empirical_vs_resample_v1_20251021.sh configured the cluster environment, pointed -npz at the GM12878 v9 CADpack NPZ, set $L$ = 4, -null-reps = 150000, and other parameters, and invoked the Python script twice (midpoint and optional junction geometries) under srun.

**Per-run CADwalk visualization with consistent AB–E semantics**

The plotting script documented in plot_walk_v9_individual.py v4 took a CADpack v9 NPZ and a run index and generated publication-ready visualizations of individual UU runs with explicit A–B–C–D–E fragment semantics. Given -npz, -run, and -outdir, it loaded run-level scalars (including run_L, step and fragment row offsets, terminal junction positions AJ1/EJ2, and terminal midpoints), step-level arrays (pos1, pos2, step_sgnJ, chromosome IDs), and fragment-level canonical bounds (frag_ref_chr_id, frag_ref_lo, frag_ref_hi, frag_len_bp, frag_len_src). It reconciled the nominal run length with available steps and fragments, chose a dominant chromosome for the absolute x-axis from fragments with valid bounds, and, when requested via -require-cis, enforced single-chromosome consistency for allfragmentrectangles.Adisplay-onlyorientationrule(-display-repair auto|aj|mid) optionally flipped both step and fragment order, ensuring that the plotted first step corresponded to AB, the second to BC, and so on, while keeping genomic coordinates and step polarity self- consistent. The script offered two layouts, an alternating fragment/step-row arrangement and a legacy step-bands mode, and could optionally screen for any canonical fragment overlaps via a configurable -ref-overlap-filter with user-defined base-pair and fractional thresholds, either erroring, skipping, or annotating overlapping runs. It drew fragment rectangles in solid, user-configurable colours with Type 42 fonts, plotted coloured arrows for each step on a shared absolute coordinate axis with a secondary axis showing distance from AJ1, summarised pattern and span (including $\Delta J$ and, when available, $\Delta M$) in the title, and exported both PNG and PDF files for each run.
